# Bridging Stem Cell Models and Medicine: Integrated 3D Human Cerebral Organoids and Pediatric Serum Reveal Mechanisms and Biomarkers of Anesthetic-Induced Neurotoxicity

**DOI:** 10.1101/2025.04.23.650256

**Authors:** Tarun Pant, Congshan Jiang, Yasheng Yan, Bailey Schultz, Daniel Laws, Sarah Logan, Amina Bedrat, Matea Juric, Richard J Berens, Susan P Taylor, Amy M Henry, Zeljko J. Bosnjak, Xiaowen Bai

**Author notes:** Equal contribution (Tarun Pant, Congshan Jiang). Correspondence to: Xiaowen Bai, MD, PhD Department of Cell Biology, Neurobiology & Anatomy Medical College of Wisconsin 8701 Watertown Plank Road Milwaukee, WI 53226.

## Abstract

**Background:** General anesthetics have been shown to cause acute pathological changes in the developing brain, including neuronal cell death. These effects contribute to long-term cognitive and behavioral impairments observed in animal models. Epidemiological and prospective clinical studies have also reported associations between early-life exposure to anesthesia and neurodevelopmental deficits in children, raising significant concerns about pediatric anesthesia and highlighting the urgent need to understand the molecular mechanisms and biomarkers of anesthetic-induced developmental neurotoxicity (AIDN) using human-relevant models.

**Methods:** This study employed human induced pluripotent stem cell-derived cerebral organoids that were exposed to varying doses of anesthetic propofol for 1 to 6 hours, including single and repeated exposures. In parallel, serum samples were collected pre- and post-surgery from pediatric patients under 4 years old (n = 10 per group) who underwent either short (<1 hour) or prolonged anesthesia (>3 hours) at the Children’s Hospital of Wisconsin between September 2018 and October 2019. Pathological changes in organoids were assessed using chemical assays, electron microscopy, and western blotting. Brain injury-related proteins in patient serum were quantified via ELISA. Genome-wide expression profiling was conducted on 18,855 mRNAs and 27,427 lncRNAs in organoids and serum using microarray and bioinformatics.

**Results:** Higher doses, longer duration, and repeated exposures to propofol led to increased apoptosis in organoids. Six-hour exposure induced autophagy as evidenced by LC3-II elevation and led to dysregulation of 553 mRNAs and 792 lncRNAs in organoids, along with their co-expressed signaling networks, affecting pathways related to synaptic integrity, mitochondrial function, and inflammation. ELISA and serum analysis of pediatric patients (<4 years old) exposed to anesthesia (>3 hours) demonstrated elevated brain cell injury-associated proteins (e.g., increased NSE) and brain cell type-specific gene expression changes. Serum findings corroborated organoid data, identifying 21 mRNAs and 12 lncRNAs that were dysregulated in both models and associated with cell injury, neuronal development, inflammation, and learning deficits.

**Conclusions:** This study represents the first integrative transcriptomic analysis of AIDN using both 3D human cerebral organoids and pediatric patient serum—two complementary, human-relevant models. By identifying consistently dysregulated coding and non-coding RNAs across both platforms, we provide compelling evidence of shared molecular signatures linked to neuronal injury, inflammation, and neurodevelopmental disruption. These findings offer not only mechanistic insights but also lay the groundwork for the development of minimally invasive biomarkers and future therapeutic strategies. This dual-model approach bridges experimental discovery with clinical relevance, advancing the translational understanding of pediatric anesthetic neurotoxicity and supporting efforts to improve long-term neurological outcomes in vulnerable patient populations.

## Background

Each year in the USA, an estimated 6 million children—including 1.5 million infants undergoing surgery—are exposed to anesthesia.^1^ While necessary for surgical procedures, compelling evidence from studies on neonatal rodents and other vertebrate species, including non-human primates, indicates that prolonged (e.g., >3 hours) or repeated exposure to general anesthetics during brain development can lead to acute molecular and cellular pathological changes, followed by long-term cognitive deficits and behavioral problems.^2–4^ These cellular changes include neuroapoptosis, synaptic loss, reduced dendritic spine numbers and stability, inhibition of neurogenesis, inflammation, decreased mitochondrial density, increased mitochondrial fission, and excessive intracellular calcium concentration.^5–7^ Several retrospective and prospective clinical studies have been conducted to explore the adverse effects of general anesthetics on immature brains. Notably, the General Anesthesia vs. Spinal Anesthesia (GAS) study, the Pediatric Anesthesia and Neurodevelopment Assessment (PANDA) study, and the Mayo Anesthesia Safety in Kids (MASK) study provided evidence that a single, short exposure to general anesthetics (45– 84 minutes) in children under 3 years old does not result in measurable deficits in neurocognitive function or behavior at follow-ups between 5 and 15 years of age.^8^ However, the MASK study revealed that children with multiple exposures performed significantly worse in processing speed and motor ability tests.^9–11^ Additionally, parents in all three studies reported worse scores in behavior, socio-affective, and executive functions in anesthesia-exposed children.^10^

Despite these compelling epidemiological findings, direct investigation of anesthetic-induced neuropathology in human brain tissue remains unfeasible due to ethical and logistical constraints. This gap necessitates the use of alternative human-relevant models, such as human induced pluripotent stem cell (iPSC)-derived cerebral organoids, which closely mimic early human brain development and cellular responses to anesthesia. While extensive preclinical studies have demonstrated anesthetic-induced neurotoxicity in developing animal brains, the translatability of these findings to human observational studies remains inconsistent. Additionally, ethical and practical barriers prevent direct assessment of anesthetic-induced developmental pathological changes in developing human brains. The advent of iPSC technology, particularly the development of three-dimensional (3D) human cerebral organoids has revolutionized neurodevelopmental research. We and others have shown that these organoids closely recapitulate key aspects of early human brain development, including cellular composition, structure organization, neurogenesis, neuronal migration, and functional responses to both neurotransmitters, alcohol, and intravenous anesthetic propofol.^12–15^ Recent studies further demonstrate their efficacy in modeling various developmental neurological disorders, including autism, offering valuable insights into disease etiology, underlying mechanisms, and potential therapeutic strategies.^16, 17^ Given these advantages, iPSC-derived cerebral organoids represent a powerful, physiologically relevant platform for investigating anesthetic-induced developmental neurotoxicity (AIDN).

In this study, we investigated anesthetic-induced pathological changes in 2-month-old human iPSC-derived cerebral organoids and clinical serum samples from pediatric patients. By integrating high-throughput RNA profiling and bioinformatics, we analyzed the roles of coding RNAs (mRNAs) and long non-coding RNAs (lncRNAs) in mediating anesthesia-induced neurotoxicity. LncRNA research is rapidly advancing, with over 100,000 human lncRNA transcripts identified, approximately 40% of which are brain-specific and exhibit distinct spatiotemporal expression patterns.^18, 19^ These molecules play essential regulatory roles in neurodevelopment, synaptic plasticity, mitochondrial function, and inflammatory responses. Given their involvement in brain homeostasis, lncRNA dysregulation may contribute significantly to AIDN pathology. To enhance our translational relevance, we employed a dual-model strategy combining advanced human organoid systems with serum samples from pediatric patients exposed to anesthesia. By linking anesthesia-induced pathological changes to lncRNA and mRNA signaling, we aim to provide crucial insights into the molecular mechanisms underlying AIDN. These findings have the potential to inform the identification of novel neuroprotective targets and serum biomarkers, ultimately advancing pediatric anesthesia safety.

## Methods

### Human iPSCs derived cerebral organoid and characterization

All the experiments in the present study utilized a healthy donor-derived iPSC line (1013^12, 20^) and were approved by the Medical College of Wisconsin Institutional Review Board (Protocol number PRO00027064). The 3D cerebral organoids were generated from iPSCs and further characterized as depicted in (Fig. 1a-e) following the previously established protocols in our lab,^12, 15, 20^ described in detail in the Supplementary Materials and Data.

**Fig. 1.**
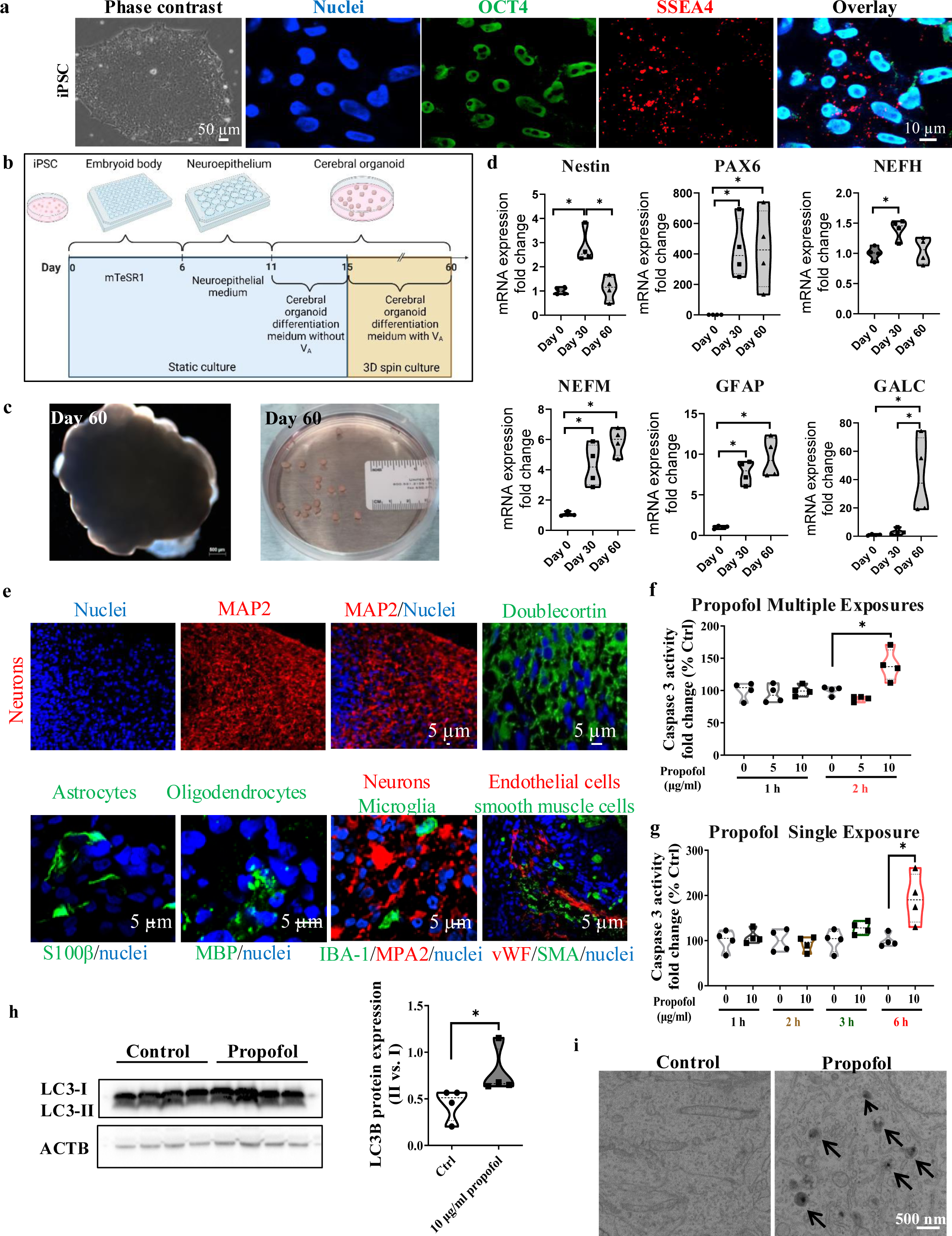
Characterization of cerebral organoids over the 60-day differentiation process from human induced pluripotent stem cells (iPSCs) and analysis of the neurotoxic effect of propofol on day 60 organoids. (a) iPSC characterization. Phase contrast and confocal images of the iPSCs stained with stem cell markers OCT4 (green) and SSEA4 (red) and nuclei stained with Hoechst 33342 (blue). Scale bar = 50 or 10 µm. (b) The scheme for generating cerebral organoids over the 60-day differentiation period (Created with Biorender.com). (c) Images of day 60 single iPSC-derived 3D sphere-like cerebral organoids with scale bar=500μm or multiple organoids cultured in a petri dish with a ruler as calibrator. (d) Reverse transcription-quantitative polymerase chain reaction (RT-qPCR) analysis of the expression of the following genes related to various types of brain cells: paired box 6 (PAX6), nestin, neurofilament heavy chain (NEFH), neurofilament medium chain (NEFM), glial fibrillary acidic protein (GFAP), and galactosylceramidase (GALC) over time during the 60-day differentiation process. n=4 (n=four independently differentiated tissue samples used in each group, with each n sample pooled from 2 organoids). *: P < 0.05. (e) Confocal images of the day 60 organoid tissue sections with Immunofluorescent staining, showing the presence of various types of brain cells within the organoids, using specific markers for neurons, astrocytes, oligodendrocytes, microglia, endothelial cells, smooth muscle cells. Nuclei were stained in blue. Scale bar: 5μm. (f) Caspase 3 activity, an indicator of apoptosis, was analyzed in cerebral organoids with multiple single (10 μg/ml) propofol exposure for 1 to 6 hours, and (g) 5 or 10 μg/ml propofol for 1 and 2 hours per day over three consecutive days by an enzyme assay kit. (h) Western blot assay displays the LC3 I/II ratio (16kD to 14kD) in organoids treated with 10 μg/ml propofol for 6 hours. Beta-actin (ACTB) served as an internal control. n=4, *: P < 0.05 vs. DMSO control. (i) Electron microscopy images display autophagosome formation following a 6-hour propofol treatment (10 μg/ml)—scale bar: 500 nm. See Supplementary Materials and Data for more details.

### Analysis of anesthetic induced apoptosis and ultrastructure changes on cerebral organoids

The cerebral organoids were treated with propofol (2,6-diisopropylphenol) (Sigma Aldrich) under different condition and further assessed for apoptotic and ultrastructural changes following our previously reported protocols^12, 20–22^ and described in detail in the Supplementary Materials and Data.

### Patient characteristics and demographics

The data included in this prospective study were derived from blood samples obtained from pediatric patients aged < 4 years of either sex. The patient profiles and details regarding the inclusion and exclusion criteria are discussed in detail in the Table S1, and Supplementary Materials and Data.

### Clinical study design and analysis of neurotoxicity in pediatric patients

The detailed description about the sample collection and processing is in the Supplementary Materials and Data. Protein levels of serum markers of brain cell injury or stress^23^, were quantified using ELISA kits (R&D Systems, Minneapolis, MN, USA) following the manufacturer’s instructions (Supplementary Materials and Data).

### Transcriptional and bioinformatic analysis of dysregulated lncRNA and mRNA

The microarray and bioinformatics analysis for the anesthetic-dysregulated lncRNA and mRNA transcripts in the cerebral organoids and serum was described in detail in the Supplementary Materials and Data. and in our previous publications.^15, 24–26^ To further identify the brain cell sources of the anesthesia/surgery -dysregulated mRNAs, we used the dataset of Zhang et al.^27, 28^ to sort brain cell type-specific (neuron, astrocyte, microglial, oligodendrocyte, and endothelial cell) mRNAs dysregulated by anesthesia/surgery in patient serum.

### Reverse transcription-quantitative PCR (RT-qPCR)

The RT-qPCR was used to analyze expression of neuronal markers and validate microarray results. The sequence of the primers is listed in the Table S2. Further details regarding the RT-qPCR assay are described in the Supplementary Materials and Data.

### Statistical analysis

All the statistical analyses that will be performed using GraphPad Prism 8 (GraphPad Software, Inc., La Jolla, CA, USA). The data in the present study were presented as mean ± standard error of mean (SEM). The sample sizes were determined based on previous experience and standard practices in the fields of neurotoxicity, stem cell, and organoid research, where sample sizes of n=3 to 4 per group have consistently yielded statistically significant results^12, 15, 29–31^. To further validate our approach, we performed a power analysis based on the pilot data of apoptosis assay. The analysis indicated that a sample size of at least 3 per group was sufficient to detect significant differences between the control and propofol groups at a significance level of 0.05 with a power of 0.8. In this study, data were obtained from n = 3 samples per group for serum studies and n = 4 samples per group for organoid studies. Each serum sample was pooled from 3 to 4 patients, resulting in a total of 10 patient serum samples per group. Each organoid sample was obtained from independent differentiations and pooled from 2 organoids, resulting in 8 organoids per group, unless otherwise specified. Student’s t-test or non-parametric Mann Whitney test was applied to identify significant difference between two groups. ANOVA was used for comparing the means of three or more groups. A p-value less than 0.05 was considered statistically significant.

## Results

### Characterization of iPSC-derived 3D cerebral organoids and anesthetic-induced neurotoxicity on cerebral organoids

Human iPSCs grew as colonies in the mTeSR1 stem cell culture medium and expressed pluripotent stem cell markers, OCT4 and SSEA4 (Fig. 1a, b). The 3D cerebral organoids derived from iPSCs exhibited a sphere-like shape (Fig. 1b, c). The expression level of neuronal markers was found to be remarkably high in day 60-cerebral organoids (Fig. 1d, e). RT-qPCR analysis showed that the gene expression of nestin, a neural stem cell marker, increased by 2.8-fold on day 30 following the initiation of iPSC differentiation, then decreased on day 60. The gene expression of PAX6, a transcription factor involved in neural development, increased dramatically on days 30 and 60 (430.5- and 431.6-fold vs. day 0 group, respectively). The expression of NEFH and NEFM, which comprise the cytoskeleton in mature neurons, showed slight increases on days 20 and 60. The expression of GFAP, an astrocyte marker, increased slightly by 3.3-fold on day 30 and significantly by 42.1-fold on day 60. The GALC, an oligodendrocyte marker, gradually increased by 7.8-and 9.5-fold on days 30 and 60, respectively (Fig. 1d). These results demonstrate that organoids develop over time in culture, showing the presence of various types of brain cells. Immunofluorescent staining and confocal imaging of cerebral organoids revealed widespread positive staining for MAP2, a neuronal marker, along with sporadic positive staining for S100B, MBP, IBA-1, VWF, and SMA, markers of astrocytes, oligodendrocytes, microglia, endothelial cells, and smooth muscle cells, respectively, further confirming the presence of various types of brain cells within the organoids. Additionally, neurons in organoids on day 60 expressed the immature neuronal marker doublecortin (Fig. 1e). Therefore, in this study, we used day 60-cerebral organoids to study whether the intravenous anesthetic propofol induced pathological changes and the underlying mechanisms.

The studies of anesthetic-dose (0, 5, or 10 µg/ml), exposure duration (1, 2, 3, or 6 hours), and exposure frequency (single or multiple) showed that either single 10 μg/ml propofol exposure for 6 hours or three exposures for 2 hours per day over three consecutive days increased the activity of caspase 3, an apoptotic marker, in the day 60-cerebral organoids (Fig. 1f, g). Western blotting results (Fig.1h) demonstrated that single propofol (10 μg/ml) exposure for 6 hours also increased LC3B I/II, an autophagy marker, in organoids. The propofol-increased autophagy was further confirmed by the electron microscopy images, displaying autophagosomes in the organoids compared to controls (Fig. 1i).

### Microarray and bioinformatic analysis of anesthetic-induced dysregulated mRNA in organoids

The integrity of RNA isolated from the day 60-organoids treated with either propofol (10 μg/ml for 6 hours) or control DMSO was confirmed to be suitable for further microarray assay, as indicated by clear and distinct 28S and 18S rRNA bands on agarose gels (Fig. 2a). The box and scatter plot results of the microarray detection signals demonstrated a similar distribution of normalized intensity between the propofol and control groups, suggesting that the gene expression signal intensity is reproducible, and the microarray results are suitable for further analysis (Fig. 2b, c). The hierarchical clustering showed that 553 mRNAs was dysregulated by propofol with 307 upregulated and 246 downregulated (P<0.05, fold change>±1.2) (Fig. 2d) documented in detail in (Table S3). Ingenuity pathway analysis (IPA)-based bioinformatic analysis of the dysregulated mRNA transcripts in propofol-treated cerebral organoids revealed associations with several critical signaling pathways. These pathways include: 1) Inflammation: Involving cytokines such as IL-2, IL-4, and IL-13. 2) Mitochondrial Function: Pathways linked to mitochondrial activities. 3) Stress Responses: Including autophagy and unfolded protein response. 3) Cellular Damage and Repair: Encompassing DNA damage, telomere maintenance, and apoptosis (Fig. 2e). 4) Cellular Function: Various pathways affecting normal cell operations. 5) Neurological Diseases and Disorders: Developmental and psychological disorders (Fig. 2f). These signaling pathways corroborate our observations of propofol-induced apoptosis and autophagy in the cerebral organoids (Fig. 1f-i) and align with previous findings from animal studies.^3, 32^ Notably, the propofol-increased expression (2.3-fold) of CRH, the gene encoding corticotropin-releasing hormone, was linked to four neural phenotypes and activities: neurite morphology, hippocampus CA3 neuron excitation, abnormal neurotransmitter release, and the initiation of apoptosis (Table S4).

**Fig. 2.**
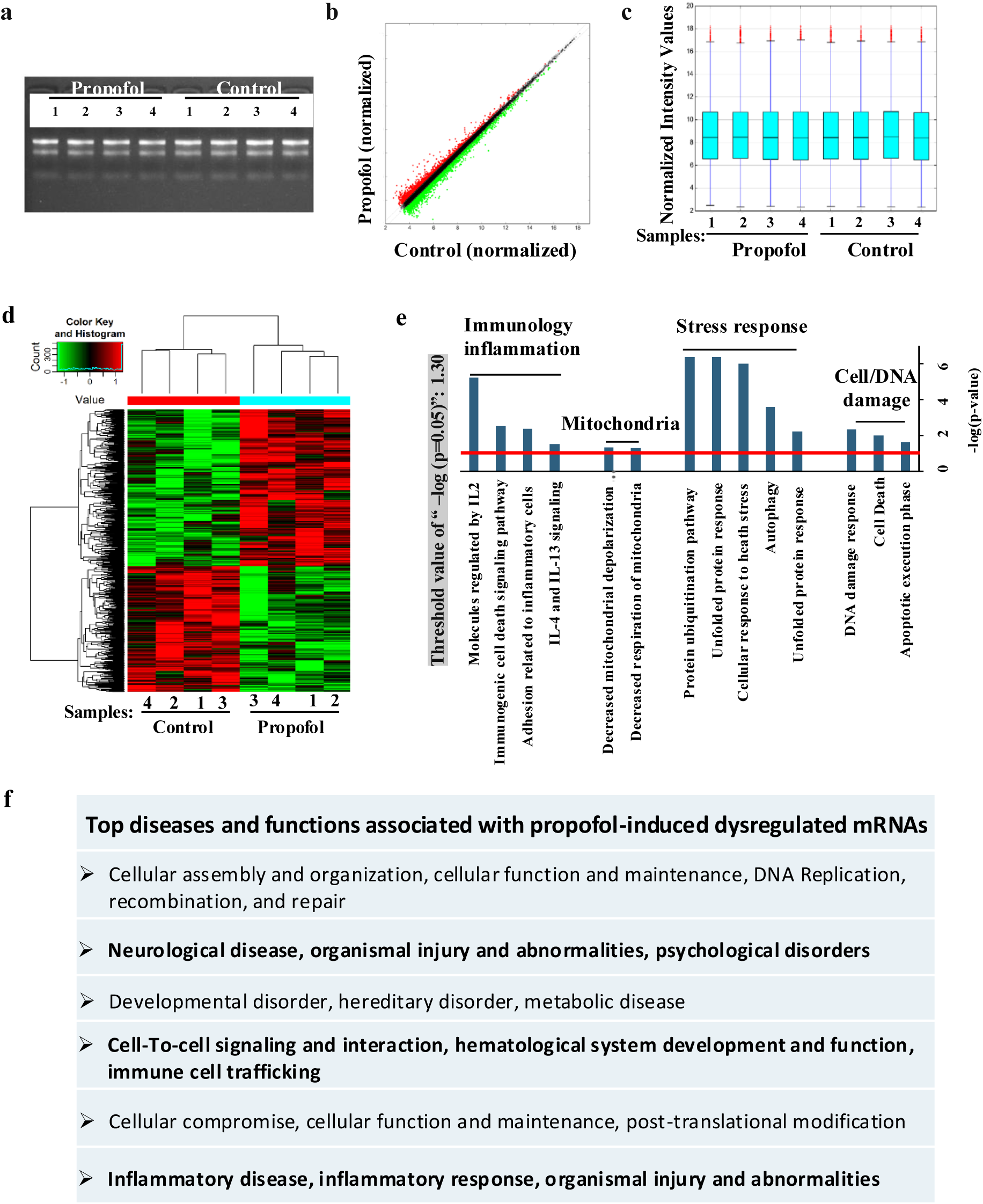
Transcriptomic analysis of propofol (10 μg/ml) exposure for six hours-induced dysregulations of inflammatory, mitochondrial, and neurological genes and pathways in day 60 cerebral organoids. (a) RNA integrity assessment using agarose gel electrophoresis of RNA extracted from control and propofol-treated organoids. (b) Box and scatter plots show normalized gene expression intensity values distribution for propofol-treated and control organoids. (c) Scatter plot comparing normalized gene expression values between propofol-treated and control organoids. (d) Heatmap representing hierarchical clustering of 553 propofol-dysregulated mRNAs (307 upregulated and 246 downregulated, p < 0.05, fold change > ±1.2) in propofol-treated organoids compared to control. (e) IPA reveals the propofol-dysregulated mRNA-associated canonical pathways with -log(p-value) above 1.30 (p < 0.05) displayed. (f) List of top diseases and functions linked to the dysregulated mRNAs. Note: The dysregulated mRNAs are listed in (Supplementary Table 3).

### Anesthetic-dysregulated synaptic, mitochondrial, and inflammatory gene expression in organoids

To further dissect the IPA-analyzed signaling above (Fig. 2e, f), we used various bioinformatic tools and databases to enrich the dysregulated synaptic, mitochondrial, and inflammatory genes from the total propofol-induced 553 abnormally expressed genes. The SynGO database analysis revealed that the propofol-induced dysregulated mRNAs include 44 synaptic genes (Fig. 3a), and these synaptic genes (CALB2, ATP2B4, NRP2, DNAJB1, NPY2R, SLC17A8, ERC2, and FILIP1) are specifically localized to the presynaptic and postsynaptic regions and are known to play crucial role in various synaptic activities, including intracellular calcium (Ca^2+^) homeostasis, presynaptic cytosolic Ca^2+^ levels, and postsynaptic organization (Fig. 3b, c, and Table S5). Further IPA-based analysis showed that these synapse-related genes are associated with: 1) behavior-related networks (e.g., anxiety-like behavior, fear, spatial, associative, and motor learning), 2) neurodevelopment and function (e.g., synaptic transmission, morphology, and development of neurons (Fig. 3d, e), and 3) calcium handling (Fig. 3f). The mitoXplorer database identified a total of 39 propofol-dysregulated mitochondrial genes, with 21 up-regulated and 18 down-regulated in organoids (Fig. 3g, and Table S6). Most of these genes are linked to mitochondrial protein translation (e.g., WARS2, CCDC58, MRPL12, AURKAIP1, PUS1, PTRH2, and EARS2), while other mitochondrial genes were implicated in essential mitochondrial functions, mainly involving oxidative phosphorylation (OXPHOS), Fe-S cluster biosynthesis, importing and sorting, and mitochondrial dynamics (Fig. 3h). Additionally, IPA analysis of these mitochondrial genes strongly implicates their role in neurological diseases and psychological disorders (Fig. 3i). Further, through the inflammation gene lists included in the NCBI database, we identified 23 propofol-induced dysregulated mRNA transcripts (Fig. 3j, and Table S7). These genes are related to distinct immune cell functions such as cell migration, activation, attachment, chemotaxis, and homing involved in processes related to neurological diseases including loss of neurons, neurogenic atrophy, and cerebrovascular dysfunction (Fig. 3k).

**Fig. 3.**
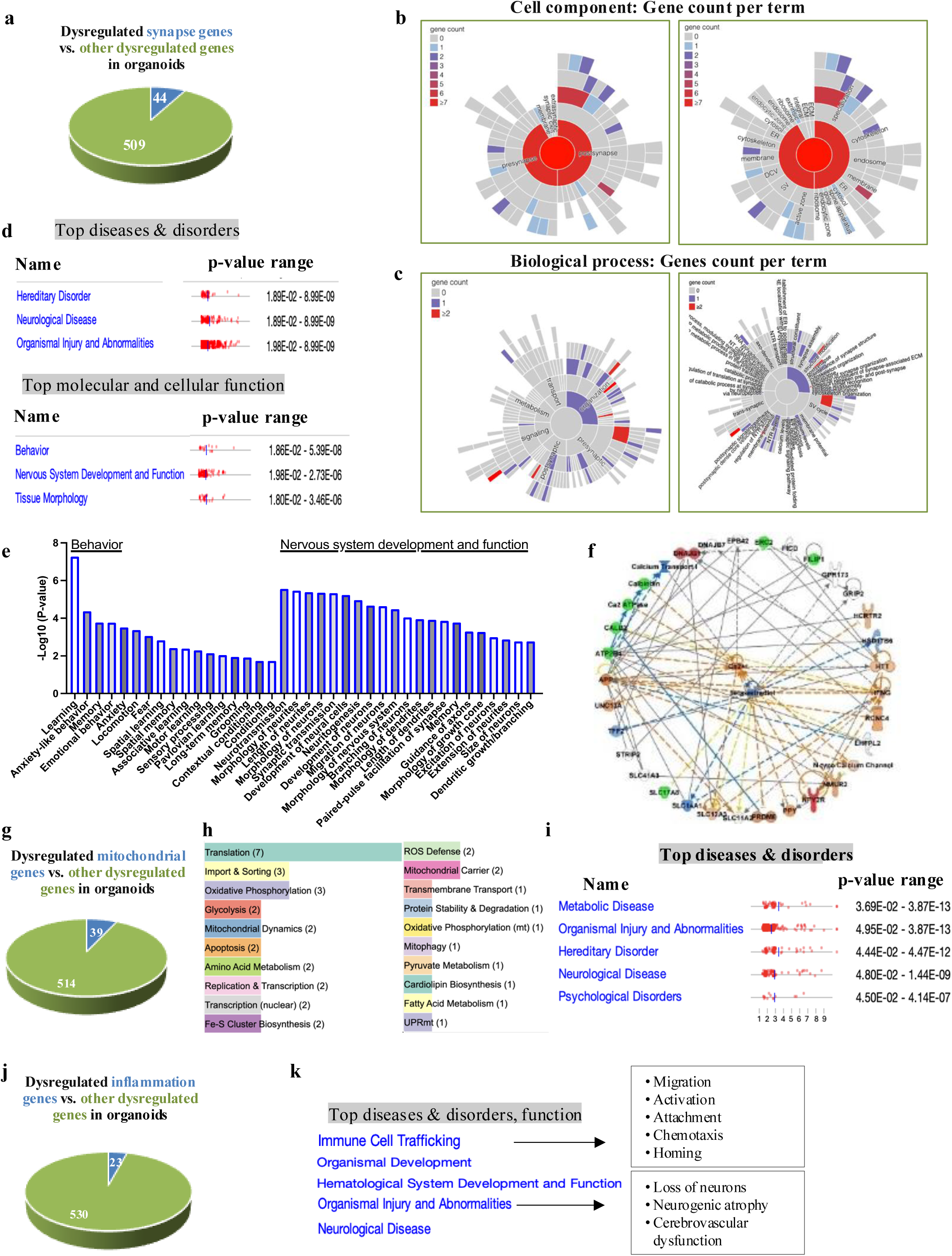
Bioinformatic analysis of propofol (10 μg/ml) exposure for six hours-induced dysregulations of synaptic, mitochondrial, and inflammatory genes (mRNAs) in day 60 cerebral organoids. (a) Bioinformatic analysis using the SynGO database identified 44 propofol-dysregulated synaptic mRNAs compared to other 509 dysregulated genes. (b) Sunburst plots represent the gene counts per term for cell components of synapse-related genes as identified in the SynGO database. (c) Sunburst plots represent the gene counts per term for biological processes of synapse-related genes as identified in the SynGO database. (d) IPA analysis predicted top diseases and disorders, and top molecular and cellular functions associated with the dysregulated synaptic mRNAs, highlighting their links to various types of cognition, behavior, and nervous system development. (e) IPA analysis highlights their links to various types of cognition, behavior, and nervous system development. (f) Network analysis using IPA predicts the mechanistic regulatory networks of propofol-induced dysregulated synaptic genes. (g) Bioinformatic analysis using the mitoXplorer database identified 39 propofol-dysregulated mitochondrial genes. (h) These genes are involved in 20 mitochondrial pathways, as calculated using the mitoXplorer database. (i) IPA analysis predicted the top diseases and disorders related to the dysregulated mitochondrial genes. (j) Bioinformatics analysis using NCBI identified 23 propofol-dysregulated inflammatory mRNAs among the total dysregulated genes (k). IPA analysis predicted top diseases, disorders, and functions related to the dysregulated inflammatory mRNAs. Note: The dysregulated synaptic, mitochondrial, and inflammatory genes are included in Supplementary Tables 5, 6, and 7. See Supplementary Materials and Data for more details.

### Anesthetic-dysregulated lncRNA profiles in organoids

Out of 27,427 lncRNAs examined, a total of 792 lncRNA were dysregulated (above ±1.2-fold difference, p < 0.05) by propofol, including 330 upregulated and 462 downregulated lncRNAs (Table S8). Hierarchical clustering suggests high dissimilarity of lncRNA profiles between propofol-treated and control organoids (Fig. 4a). The lncRNA differential expression analysis was performed between propofol-treated and control organoids (Fig. 4a, and Table S8), and by calculating the correlation coefficients between 792 dysregulated lncRNAs and abnormally expressed synaptic, mitochondrial, and inflammatory mRNA transcripts (Fig. 4b), we identified networks between abnormally expressed lncRNAs and dysregulated synaptic, mitochondrial, and inflammatory mRNAs involved in regulating crucial mechanisms associated with neurotoxicity in organoids (Fig. 4c-e, and Tables S9, S10 and S11).

**Fig. 4.**
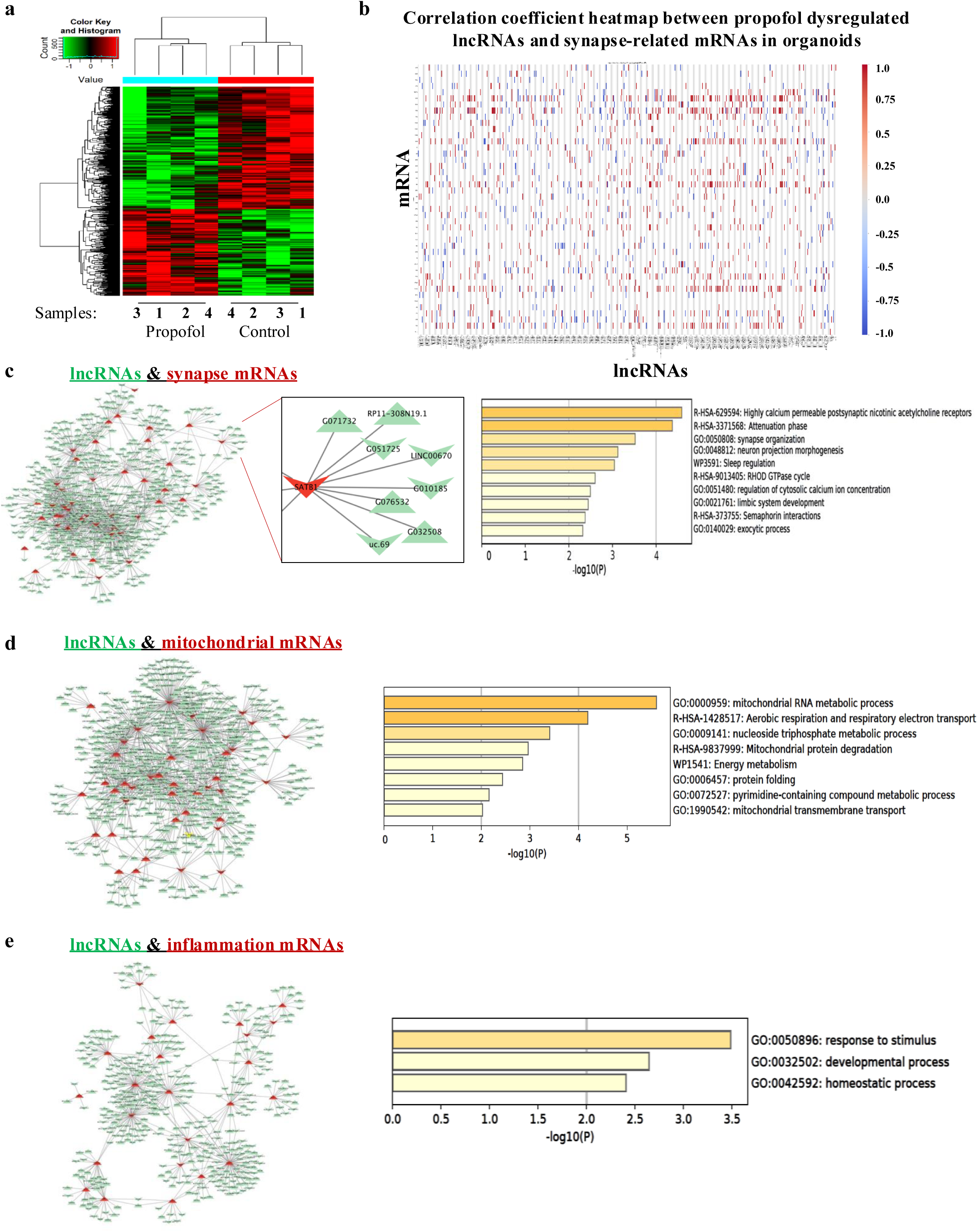
Propofol (10 μg/ml) 6 hrs. exposure dysregulated expression profile of lncRNAs in day 60 organoids, coexpression with dysregulated synaptic, mitochondrial, and inflammatory mRNAs in multiple pathways. (a) Heatmap representing the differential expression (│fold change│>1.2 and p-value <0.05) of propofol-dysregulated lncRNAs in organoids. (b) Correlation coefficient heatmap between propofol-dysregulated lncRNAs and synaptic mRNAs in organoids. The coexpression network and related pathways of dysregulated lncRNA with their co-expressing mRNA transcripts were analyzed using the Cytoscape 3.10.2 platform. (c) The network demonstrates the coexpression relationships between propofol-dysregulated lncRNAs and synaptic mRNAs in organoids, with a cutoff of coexpression efficiency >0.9 and P<0.05 from Fig. 4b. Metascape platform-based Enriched Ontology Clusters of the propofol-dysregulated synaptic mRNAs that were co-expressed with dysregulated lncRNAs are shown alongside the coexpression network. (d) Coexpression network between the propofol-deregulated lncRNAs and mitochondrial mRNA. (e) Inflammatory mRNAs are shown alongside the related Enriched Ontology Clusters of the propofol-dysregulated mitochondrial and inflammatory genes included in the networks. Note: The dysregulated lncRNAs and mRNAs in the networks are listed in Supplementary Tables 9, 10, and 11. See Supplementary Materials and Data for more details.

### Anesthesia-induced increase of brain injury-associated protein levels and dysregulated transcriptomic profiles in the serum of pediatric patients

The serum samples were collected pre- and post-surgery from pediatric patients under 4 years old (10 patient serum samples per group) who underwent anesthesia (<1 hour or >3 hours) with propofol, ketamine, isoflurane, or sevoflurane at the Children’s Hospital of Wisconsin between September 2018 and October 2019. The studies from the patient serums, as a complementary models of stem cell-derived human cerebral organoids (Fig. 5a), showed that less than one-hour anesthesia/surgery did not change the protein levels of BDNF, NSE, and S100B, serum markers of brain cell injury or stress.^23^ However, prolonged (>3hours) exposure to general anesthesia/surgery significantly altered the levels of these proteins in serum (Fig. 5b), suggesting the brain injury of the patients with longer surgery and anesthesia.

**Fig. 5.**
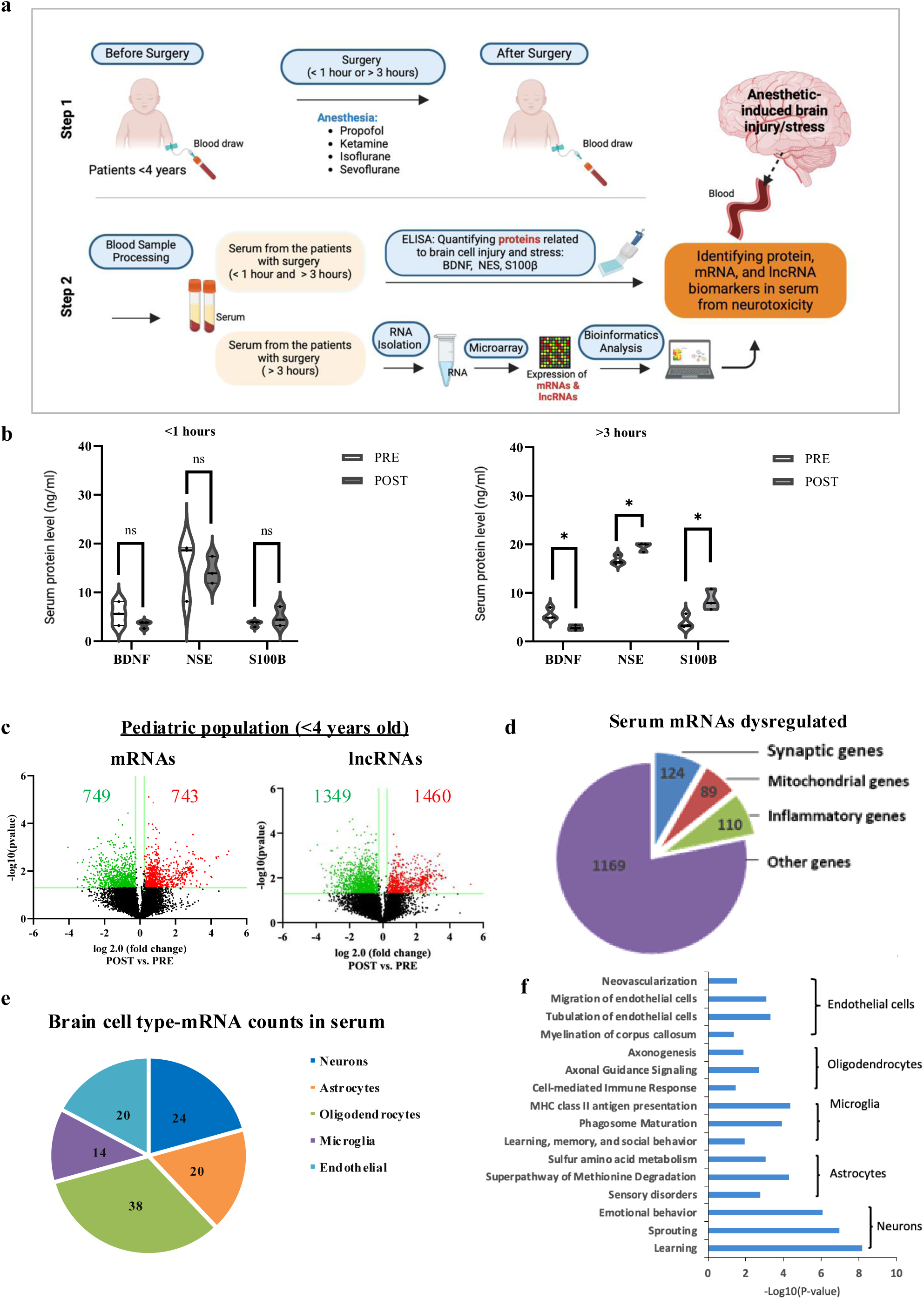
General anesthesia/surgery (>3 hours) in pediatric patients (< 4 years) resulted in abnormal levels of serum brain injury markers and dysregulated profiles of mRNAs and lncRNAs. (a) Schematic depicting the experimental design for clinical studies, including patient information on anesthesia/surgery duration (< 1 hour or > 3 hours), anesthetics used, age, and blood sample collection time points (pre- and post-surgery). (b) ELISA quantified protein levels of brain injury biomarkers in serum collected from pediatric patients undergoing non-cardiac and neuronal surgeries before (pre-surgery) and after (post-surgery) with anesthesia/surgery < 1 hour or > 3 hours: brain-derived neurotrophic factor (BDNF), neuron serum enolase (NSE), and S100 calcium-binding protein B (S100B). n=3, with each n representing serum samples from 3 to 4 patient-derived pooled samples, totaling 10 patients. *: P < 0.05. (c) Volcano plot showing dysregulated mRNA and lncRNA profiles in serum from pediatric patients with anesthesia/surgery duration longer than 3 hours compared with pre-surgery control samples. Green and red dots represent downregulated and upregulated mRNAs and lncRNAs (above 1.2-fold change and p < 0.05). (d) Pie chart displaying dysregulated synaptic, mitochondrial, and inflammatory mRNAs in patients’ serum (> 3-hour anesthesia). (e) Pie chart displaying the sorted 116 brain cell type-specific anesthesia/surgery (> 3 hours)-dysregulated mRNA counts from the total dysregulated 1492 mRNAs in patients’ serum based on Zhang’s data set (f) IPA analysis of neuron- and oligodendrocyte-associated anesthesia/surgery-dysregulated genes. ^27, 28^ Note: The dysregulated lncRNAs and mRNAs in the serum and the brain cell type-specific dysregulated mRNAs in serum are listed in Supplementary Tables 12, 13 and 18. See Supplementary Materials and Data for more details.

The microarray assay also detected the mRNA and lncRNA expression profiling for pediatric patients who underwent anesthesia (>3 hours). The box and scatter plot for serum samples showed that the mRNA and lncRNA normalized gene expression intensity values were consistent for all the samples, confirming data integrity and consistency (Fig.S1). Our microarray analysis further showed that compared with the pre-surgery control samples of pediatric patients, 1492 mRNA transcripts (Fig.5c, and Table S12) and 2809 lncRNAs (Fig.5c, and Table S13) were dysregulated (above 1.2-fold change and p< 0.05) in the post-surgery serum samples. Consistent with prior report bioinformatic analysis showed that anesthesia/surgery-dysregulated mRNA had enrichment of transcripts related to the pathways associated with initiation and progression of neurodegeneration (Table S14). Further, the SynGO, mitoXplorer, and NCBI database-based analysis identified that 1492 dysregulated mRNAs include 124 synaptic genes, 89 mitochondrial genes, and 110 inflammatory genes (Fig.5d, and S15, S16 and S17). The microarray assay was further validated by RT-qPCR analysis of randomly selected mRNAs (NDUFA11, PGAM5, DRD1, and APC2) and two lncRNAs in serum, showing the same expression trend (up or down) for these genes (Fig. S2 and S3) as we observed in the microarray results (Table S12 and 13).

### Anesthesia/surgery-dysregulated brain cell type-specific mRNAs in serum

Through the dataset of Zhang et al.^27, 28^ we sorted brain cell type-specific mRNAs from the 1,492 anesthesia/surgery-induced dysregulated mRNAs. We identified 116 brain cell-specific mRNAs in the serum from patients with >3-hour anesthesia/surgery: 38 genes associated with oligodendrocytes, 24 with neurons, 14 with microglia, 20 with astrocytes, and 20 with endothelial cells (Fig.5e, and Table S18). IPA analysis indicates that the neuron- and oligodendrocyte-associated anesthesia/surgery-dysregulated genes were mainly enriched for emotional behavior, learning, sensory disorders, and brain development (e.g., axon genesis, axon guidance, and sprouting). In contrast, microglia-, astrocyte-, and endothelial cell-associated genes were mainly enriched in immune responses, phagosome maturation, metabolism, neovascularization, myelination, and angiogenesis (Fig.5f).

### Overlapping anesthetic-dysregulated mRNAs and lncRNAs in anesthetic-exposed cerebral organoids and patient serum

To investigate the therapeutic and prognostic applications of cerebral organoids in the clinical setting, we characterized the overlapping anesthesia-dysregulated mRNAs and lncRNAs between cerebral organoids and patient serum samples. Specifically, we identified 21 overlapping mRNAs (Fig. 6a, b, and Tables S19, S20 and S21) between the organoid dataset (553 total) and the patient serum dataset (1492 total). Among these, two were related to synapse function (ACTN1 and PTPRN2) and one to mitochondrial function (CKMT1B) (Fig.6c, and Tables S20, S21). Additionally, there were 12 overlapping lncRNAs (Fig. 6d, e), between the organoid dataset (792 total) and the patient serum dataset (2809 total). All 21 overlapping mRNAs and 12 lncRNAs showed the same expression pattern (either upregulated or downregulated) (Fig. 6a-e, and Table S22). IPA analysis indicates that these 21 dysregulated overlapping mRNAs are associated with top canonical pathways, including response to stress, melatonin degradation (involved in the regulation of sleep and circadian rhythms), angiogenesis, energy metabolism, and inflammation (Fig. 6f). Metascape enrichment analysis also suggests the involvement of these dysregulated mRNAs in various other pathways, including the regulation of cell activities such as cell division, organelle fusion and organization, and vesicle-mediated transport (Fig. 6g). Notably, seven of the 12 overlapping lncRNAs and 11 of the 21 overlapping mRNAs were found to co-express (Pearson correlation > 0.9 or < −0.9, p < 0.05), forming crucial regulatory networks (Fig. 6h, i). The top network is related to cell morphology, neurological diseases, organismal injury, and abnormalities (Fig. 6j), and disease pathways and functions related to various neuronal activities and functions, tissue development, cell compromise, cell injury and death, inflammation, and the quantity of phagocytes (Fig. 6k).

**Fig. 6.**
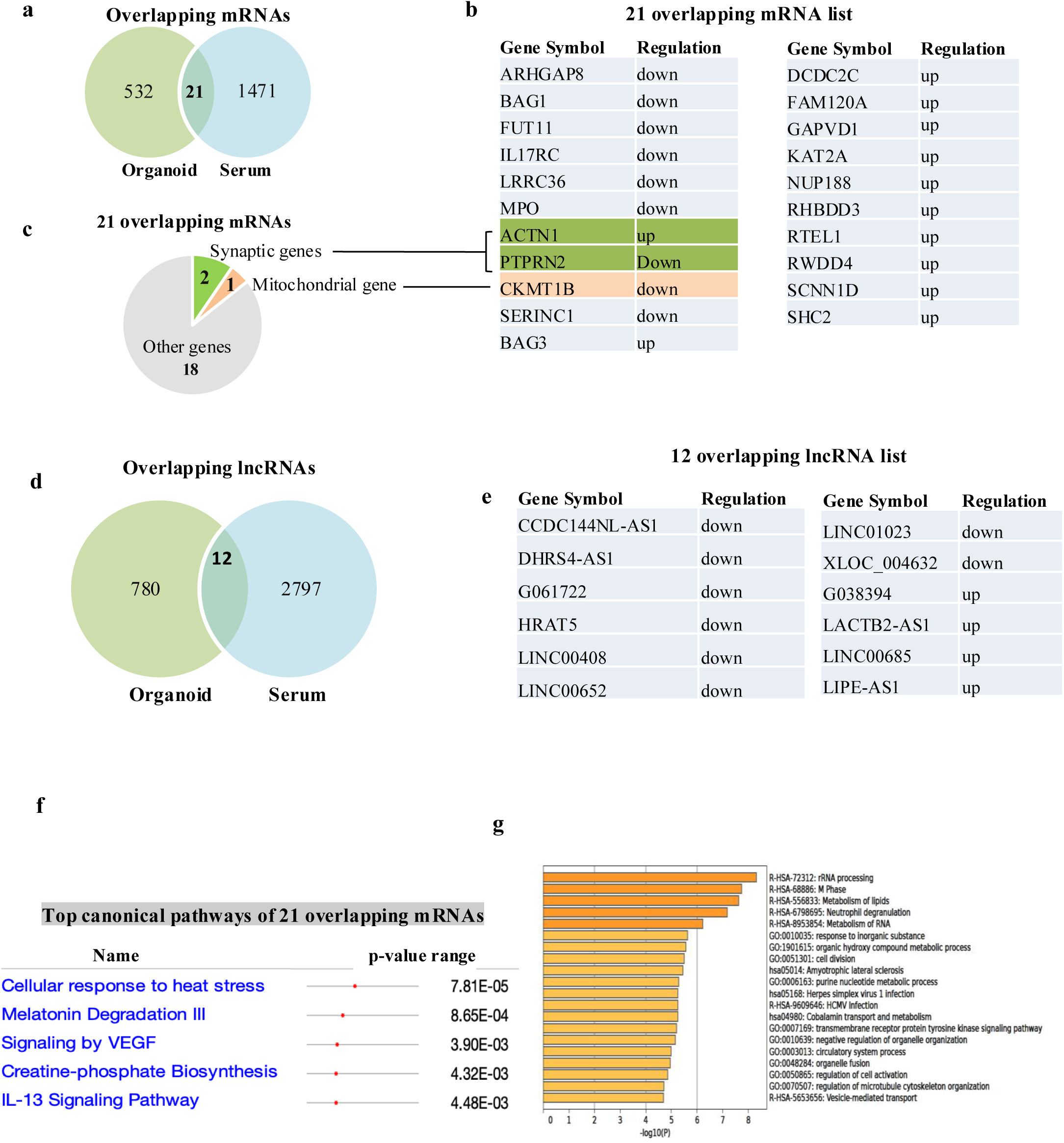
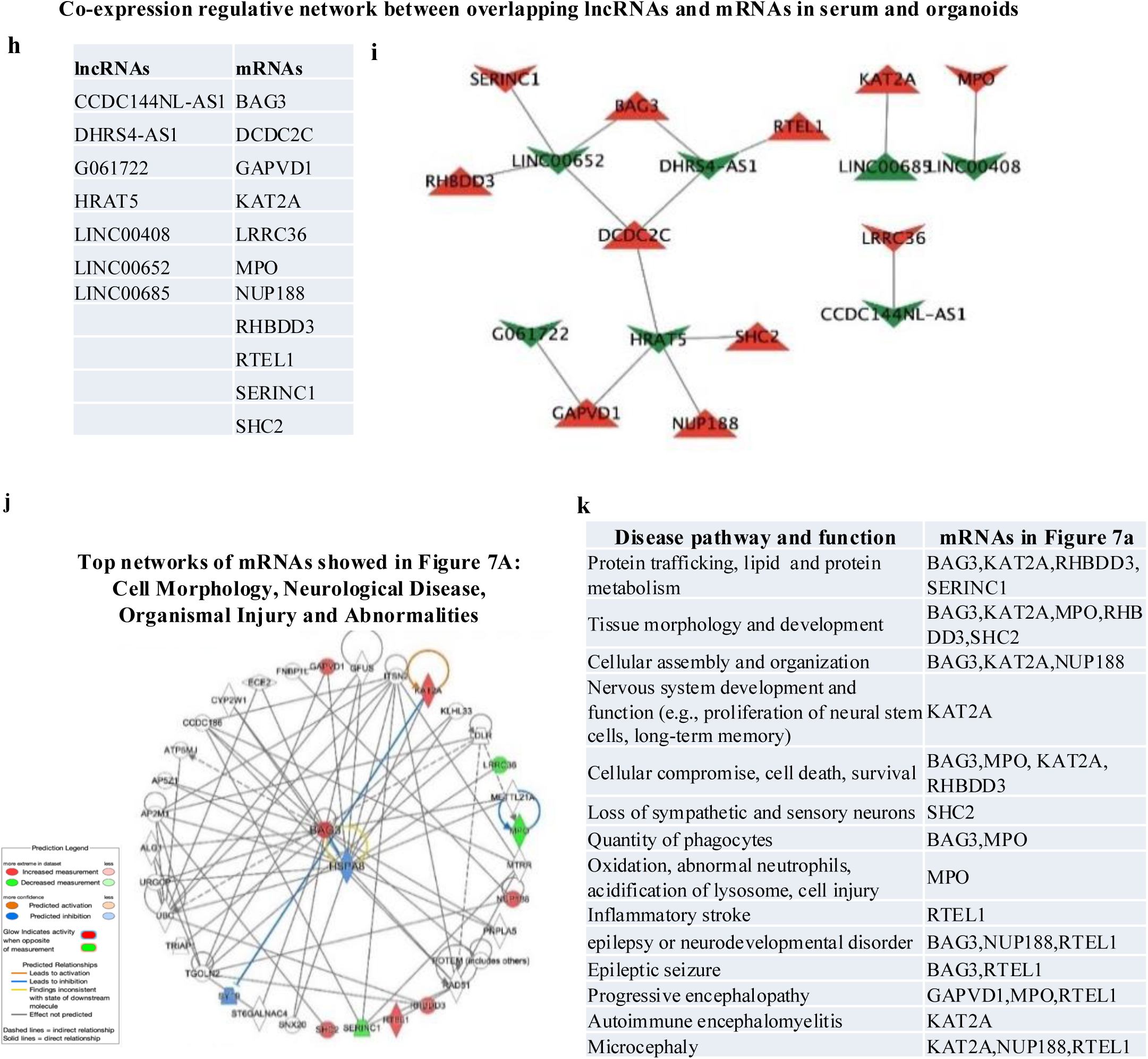
The Shared dysregulated mRNAs and lncRNAs identified in pediatric patient serum following general anesthesia (>3 hours) and in day-60 cerebral organoids treated with propofol (10 µg/ml for 6 hours), along with the associated signaling pathways. (a) Venn plot displaying the 21 differentially modulated mRNAs overlapping in both cerebral organoids and pediatric serum samples after anesthetic exposure. (b) List of the 21 overlapping mRNAs, including their gene symbols and the direction of regulation (up or down) in both organoids and serum. (c) Pie chart illustrating the 21 overlapping mRNAs, highlighting 2 synaptic genes (ACTN1 and PTPRN2) and 1 mitochondrial gene (CKMT1B) among the overlapping transcripts. (d) Venn plot shows 12 differentially modulated lncRNAs overlap between cerebral organoids and pediatric serum samples after anesthetic exposure. (e) Top canonical pathways and enriched biological processes (p<0.05) associated with the 21 overlapping mRNAs identified via IPA-based pathway analysis. (f) Top canonical pathways and enriched biological processes (p<0.05) associated with the 21overlapping mRNAs identified via IPA-based pathway analysis. (g) Metascape enrichment analysis of 21 overlapping mRNAs in various other pathways, including regulating cell activities such as cell division, organelle fusion and organization, and vesicle-mediated transport. See Supplementary Materials and Data for more details. (h) List of the 12 overlapping lncRNAs, including their gene symbols and the direction of regulation (up or down) in both organoids and serum. (i) Coexpression analysis between the overlapping anesthesia-dysregulated lncRNAs and mRNAs in serum and organoids using Cytoscape 3.10.2 platform identified 7 lncRNAs and 11 mRNAs coexpressed, forming a regulative network. A cutoff of coexpression efficiency > 0.9 and p < 0.05 was used. (j) IPA predicted the top network of mRNAs listed in Fig. 6h, suggesting the involvement of the lncRNAs in these pathways through regulation of the mRNAs in the coexpression network. (k) Table summarizing the top disease pathways and functions associated with the mRNAs included in Fig. 6h, i. See Supplementary Materials and Data for more details.

## Discussion

In this study, we leveraged two complementary, human relevant models systems (stem cell-derived 3D cerebral organoids) and pediatric patient serum samples—to perform the first integrated molecular analysis of how general anesthetics impact neurodevelopment and induce neurotoxic effects in the developing brain. Through a multifaceted approach—including imaging, protein assays, transcriptomic profiling, and bioinformatics—we present direct evidence that anesthetic exposure induces dose- and duration-dependent cellular injury and stress responses in human tissues. These findings provide key insights into the molecular mechanisms driving anesthesia-induced neurotoxicity and support the identification of serum biomarkers for early detection. Furthermore, this study establishes a foundation for developing targeted neuroprotective strategies to prevent anesthesia-related neurodevelopmental impairment.

### Direct evidence of anesthetic-induced pathology changes in human brain tissue

Mounting animal studies have provided strong evidence of general anesthetic-induced acute neurotoxic effects (e.g., apoptosis, reduced synaptogenesis, mitochondrial dysfunction, neurogenesis, and inflammation) in the developing brain followed by long-term learning and memory disability as well as behavioral abnormalities. However, there is inconsistent human epidemiology data related to the association between early general anesthesia and long-term brain dysfunctions, raising the concerns of the translational potential of animal findings into human patient settings. This study pioneers the extension of our prior animal studies on AIDN to a 3D human model using iPSC-derived cerebral organoids. The organoids can mimic developing brains in multiple aspects, including cell components with immature neurons, various types of glial cells (astrocytes, oligodendrocytes, and microglia), and vascular cells (Fig. 1a-e). They exhibit complex tissue structures and functional electrophysiological responses to propofol and neurotransmitters.^12^ Our data revealed that propofol, whether as a 6-hour single exposure or as triple exposures for a much shorter duration (2 hours/day) at the same concentration (10 µg/ml), increased apoptosis in organoids in a dose-, concentration-, and exposure frequency-dependent manner (Fig. 1f, g). In addition to apoptosis, we found that propofol resulted in an increase in autophagosomes and the autophagy marker LC3B I/II turnover in the organoids (Fig.1h, i). These findings are significant because the propofol concentration, exposure duration, and frequency used in our study closely mirror clinical practices in pediatric anesthesia. Our results align with animal studies demonstrating that both prolonged exposure and increased frequency of exposure, even at shorter durations, can lead to developmental neurotoxicity in rodents.^33, 34^ By replicating clinical exposure conditions, our study provides valuable insights into the potential risks of AIDN. Furthermore, we found that propofol increased autophagy in the cerebral organoids. This observation is consistent with findings from various animal models studying propofol-induced developmental neurotoxicity. Both in vivo animal studies and in vitro experiments with cultured rodent neurons have reported similar autophagy responses following propofol exposure.^8, 35^ Additionally, comparable autophagy alterations have been observed in developing rodent neurons or human stem cell-derived neurons treated with other anesthetics,^29, 36^ such as sevoflurane and ketamine, highlighting the significant role of autophagy in anesthetic-induced neurotoxicity.

### Mechanisms of anesthetic-induced neurotoxicity in developing brain organoids

Our findings in cerebral organoids revealed significant mechanisms of anesthetic-induced neurotoxicity, which remain largely unknown, especially in human brain tissue. Microarray studies and bioinformatic analysis revealed that propofol exposure resulted in an abnormal organoid transcriptome (307 upregulated and 246 downregulated genes), exhibiting enrichment for calcium signaling, mitochondrial dysfunction, inflammatory activity, autophagy, and apoptosis in the bioinformatics analysis (Fig. 2a-f, and Tables S3, S4, S5, S6, and S7). Calcium signaling and synaptic genes are critical for maintaining the structure and function of synapses, which are essential for neural communication, learning, and memory. For example, among 44 propofol-dysregulated synaptic genes (Fig. 3, a-c, and Table S5), ATP2B4, CALB2, and NRP2 are involved in calcium signaling, neuronal excitability, and synaptic function. This aligns with our recent findings of abnormal excitatory and inhibitory balances in the propofol-treated developing mouse brain tissue.^3^ Collectively, these dysregulated synaptic genes at either transcriptional or protein levels indicate compromised synaptic integrity by propofol in human cerebral organoid tissues, ultimately contributing to cognitive and behavioral deficits (Fig. 3d-f).

Mitochondrial genes are involved in energy production, oxidative stress regulation, intracellular calcium homeostasis, and apoptosis. Moreover, mitochondria are extremely dynamic and able to divide, fuse, and move along microtubule tracks to ensure their distribution to the neuronal periphery. Mitochondria participate in all activities of neurons, including birth, regulating the outgrowth of axons and dendrites, neurogenesis, synaptogenesis, synapse function, and cell death. Mitochondrial dysfunction and altered mitochondrial dynamics are observed in a wide range of conditions, from impaired neuronal development to various neurodegenerative diseases and neuroinflammation. In the current study, we found that propofol dysregulated 39 mitochondrial genes involved in various mitochondrial activities, including transition, mitochondrial dynamics, apoptosis, energy generation, and metabolisms, indicating the toxic effect of propofol on mitochondria in human brain cells (Fig. 3g, h, and Table S6). Notably, the brain displays a high energy demand. In humans, the brain consumes approximately 20% of our metabolic energy, despite comprising only 2% of our body weight. Specifically, synapses are sites of high energy demand, dependent on elevated levels of mitochondrial-derived adenosine triphosphate. Therefore, brain tissue is particularly vulnerable to propofol-induced mitochondrial dysfunction, resulting in brain cells undergoing stress, injury, synapse dysfunction, impaired neuronal development, and even death, such as apoptosis, as we and others observed in animal models. The increased expression of autophagy further supports the notion of mitochondrial stress and the activation of compensatory mechanisms to remove damaged mitochondria (Fig. 3g-i).

In addition to the propofol-deregulated synaptic and mitochondrial genes in organoids, bioinformatic analysis also revealed 23 genes related to inflammation (Fig. 3j, and Table S7). Inflammatory processes are often activated in response to cellular stress and injury, and chronic inflammation can exacerbate neuronal damage. Upregulation of pro-inflammatory cytokines and chemokines can lead to neuroinflammation, contributing to the pathology of various neurodegenerative conditions. The altered expression of these genes indicates that propofol exposure may initiate or exacerbate inflammatory pathways, as observed in animal models,^37^ further compromising brain health. The dysregulation of these key gene categories highlights the multifaceted nature of propofol-induced neurotoxicity. IPA-based analysis revealed several insights: 1) Propofol-dysregulated synaptic genes are related to behavior and nervous system development and function. 2) The top unique diseases and disorders related to dysregulated mitochondrial genes are metabolic diseases, psychological disorders, and hereditary disorders. 3) Dysregulated inflammatory genes are uniquely associated with immune cell trafficking. Interestingly, we found that dysregulated mitochondrial and inflammatory genes share the same two top disease categories: "organoid injury and abnormalities" and "neurological disorders” (Fig.3i, k). The observed changes in synaptic, mitochondrial, and inflammatory genes, along with their associated unique and common top diseases and disorders, indicate a coordinated response of human brain tissue to propofol-induced stress. This suggests complex molecular and cellular mechanisms of neurotoxicity, including pathological changes and cognitive and behavioral deficits.

### LncRNA mechanisms in anesthetic neurotoxicity

LncRNAs are a class of RNA molecules that do not encode proteins but play crucial roles in regulating gene expression and various cellular processes. LncRNAs have very low sequence conservation across different species compared to protein-coding genes, and over 48% of the discovered lncRNAs are specifically expressed in the brain. A growing number of studies indicate that lncRNAs are involved in brain development and the pathogenesis of various neurodegenerative diseases, including Alzheimer’s disease and schizophrenia.^38^ The roles of lncRNAs in AIDN in human brain tissue are unknown, making it important to study lncRNA mechanisms in AIDN using the model of human cerebral organoids. LncRNAs can regulate the transcription of nearby or distant protein-coding genes. Therefore, we used dysregulated lncRNAs co-expressed with dysregulated protein-coding genes to predict the lncRNA mechanisms in neurotoxicity. The findings from the current study showed, for the first time, that propofol exposure elicited changes in the organoid lncRNAs (downregulation of 462, and upregulation of 330; Fig. 4a, and Table S8). Co-expression analysis identified that propofol-dysregulated lncRNAs form regulatory networks with the dysregulated mRNAs, including synaptic, mitochondrial, and inflammatory genes (Fig. 4b-e). These data suggest the roles of lncRNAs in various pathways and highlight the complexity of their interactions with coding genes in neurotoxicity.

### Patient serum-derived supportive evidence related to neurotoxicity by anesthesia

In addition to the neurotoxic findings from cerebral organoids, our study provides supportive evidence from patient serum samples, further corroborating the neurotoxic effects of anesthesia. Our results revealed significant alterations in serum biomarkers associated with brain injury. Specifically, we observed elevated protein levels of S100B and NSE, and decreased levels of BDNF in the serum of pediatric patients aged less than 4 years who underwent anesthesia/surgery for longer than 3 hours, but not for less than one hour (Fig. 5a). S100B is a calcium-binding protein primarily found in glial cells and is considered a marker of astrocyte activation and brain damage. NSE is a glycolytic enzyme found in neurons, and elevated levels in the serum are indicative of neuronal injury. BDNF is crucial for neuronal survival, development, and plasticity. The decreased levels of BDNF following anesthesia suggest a reduction in neurotrophic support, which may contribute to neuronal vulnerability and impair recovery processes following injury.^39–41^ The changes in these biomarkers strongly indicate that general anesthesia induces significant neurotoxic effects and brain injury in pediatric patients. These biomarkers provide a measurable indication of the extent of brain injury, supporting the observations made in the cerebral organoids (Fig. 5b). The serum biomarkers corroborate our findings in organoids, suggesting that prolonged anesthesia exposure has detrimental effects on the developing human brain.

In addition to the altered serum markers of brain injury (Fig. 5b), the relatively low number of overlapping mRNAs (21) and lncRNAs (12) identified in cerebral organoids and patient serum, and their involvement in critical pathways (e.g., synaptic integrity, mitochondrial health, inflammation, neurodevelopment, and neurological diseases), underscore the relevance of stem cell-derived human organoids in modeling AIDN in human brain tissues. The biological differences between tissue types (brain organoids vs. serum) are likely the primary reason for the low overlap of dysregulated mRNAs and lncRNAs between the datasets. Organoids represent localized effects on specific brain cell types (e.g., neurons and glial cells), whereas serum contains systemic biomarkers influenced by broader physiological responses, including inflammation, stress, and immune activation. These systemic responses may not be fully captured by organoid models, which focus primarily on localized tissue-specific effects. Additionally, the blood-brain barrier in patients restricts molecular exchange between the brain and systemic circulation, creating inherent differences in gene expression profiles. Despite the low overlap, both organoid and serum analyses provide valuable and complementary insights. The organoid model allows us to explore direct effects of anesthetics on brain-specific pathways, such as synaptic integrity and mitochondrial function, while serum analysis reveals systemic biomarkers indicative of broader physiological responses, like inflammation and stress. Together, these models enhance our understanding of AIDN and its impact on neurodevelopment. By combining insights from tissue-specific and systemic analyses, our study provides a comprehensive framework for investigating AIDN, with potential implications for improving anesthetic safety in pediatric populations.

### Serum biomarkers and therapeutic targets for anesthesia-induced neurotoxicity

The lack of biomarkers that identify brain injury due to exposure to general anesthetics makes detecting anesthesia-induced neurodevelopmental outcomes challenging in clinical practice. The possibility of easily detecting changes in the altered mRNAs and lncRNA profile due to anesthetic exposure in the serum of pediatric patients, which can aid in predicting the severity of neurotoxicity, provides an opportunity to use them as biomarkers for AIDN in clinical practice. This concept is supported by recent studies showing that circulating proteins, mRNAs, and lncRNAs in the serum and plasma of patients correlate with brain injury from various diseases, including traumatic brain injury, inflammation, and Alzheimer’s disease.^39–43^ Our studies reveal the following potential serum biomarkers of AIDN: 1) Serum proteins BDNF, NSE, and S100B, well-known serum biomarkers of brain cell injury,^39–41^ were altered following prolonged anesthesia exposure. 2) 21 dysregulated mRNAs and 12 lncRNAs overlapping between anesthetic-induced organoids and serum from patients receiving anesthesia for longer than 3 hours (Fig. 6a-e). 3) The abnormally expressed 116 serum mRNAs that are brain cell type (neurons, astrocytes, oligodendrocytes, microglial, and endothelial cells)-specific genes (Fig. 5e, and Table S18). Both animal studies and human stem cell-based in vitro models have shown that certain volatile anesthetics (e.g., isoflurane, sevoflurane) and intravenous anesthetics like propofol can damage the blood-brain barrier by altering crucial barrier properties.^44, 45^ Emerging evidence also suggests that extracellular vesicles are an alternative transportation tool for delivering brain cell content, including proteins and RNAs, to the serum. Therefore, it is possible that these brain injury protein markers and overlapping RNAs in patient serum were from leaking BBB and extracellular vesicles.

The overlapping dysregulated mRNAs include two synaptic genes (ACTN1 and PTPRN2), one mitochondrial gene (CKMT1B, as creatine kinase mitochondrial 1B), and 18 other genes involved in various cellular processes (e.g., signal transduction, immune response, and cell survival). ACTN1 (Alpha-actinin 1) is involved in the organization of the cytoskeleton and is crucial for maintaining synaptic structure and function. Its upregulation might indicate synaptic remodeling or stress response. PTPRN2 (Protein tyrosine phosphatase receptor type N2) plays a role in synaptic vesicle recycling and neurotransmitter release. Its downregulation suggests impaired synaptic transmission, which could lead to cognitive deficits. CKMT1B is essential for energy metabolism in neurons. Its downregulation indicates mitochondrial dysfunction, which can lead to energy deficits and neuronal damage. Among the 12 dysregulated lncRNAs and 21 mRNAs overlapping in both organoids and patient serum, 7 lncRNAs were co-expressed with 11 protein-coding genes, forming regulatory networks in response to anesthesia stress and contributing to brain cell injury, death, development, and cognition (Fig. 6h-k). This further supports these genes as potential serum biomarkers of neurotoxicity. Together, these potential serum biomarkers emphasize the multifaceted nature of AIDN. This includes apoptosis, synapse disruption, unhealthy mitochondria, inflammation, and their associated dysregulated coding and lncRNA genes, as well as brain cell type-specific signaling. These insights not only deepen our understanding of underlying mechanisms of AIDN but also lay the groundwork for translational applications, including serum-based diagnostics and RNA-targeted therapeutic strategies. Such interventions could include anti-inflammation, mitochondrial stabilizers, and RNA-based therapeutics that modulate expression of anesthetic-dysregulated lncRNAs and mRNAs.

## Conclusion

Our study utilized two complementary models to investigate anesthesia-induced neurotoxicity: human stem cell-derived cerebral organoids and serum samples from pediatric patients. This dual approach allowed us to explore the neurotoxic effects of anesthetics under controlled conditions using the organoids, while also considering real-world confounding factors present in patient serum samples. Cerebral organoids provide a unique and controlled environment to study the direct impact of anesthetics on human brain tissue. They mimic various aspects of human brain development, including cell components, tissue structure, and functional responses. Our studies revealed significant mechanisms of anesthetic-induced neurotoxicity, such as cell apoptosis, synaptic dysfunction, mitochondrial stress, and inflammation. This model allowed us to pinpoint specific dysregulated coding genes and lncRNAs, and their related pathways, offering a clear understanding of the underlying molecular and cellular mechanisms. Serum samples from pediatric patients exposed to anesthesia provided critical insights into the systemic effects of anesthetics, reflecting the complex interplay of various physiological factors. We identified altered levels of well-known serum biomarkers of brain injury (BDNF, NSE, and S100B) and overlapping dysregulated mRNAs and lncRNAs between organoids and patient serum. These findings suggest potential serum biomarkers for anesthesia-induced neurotoxicity, supporting the clinical relevance of our in vitro human organoid model. Additionally, the serum analysis reinforced the organoid findings, revealing dysregulated genes associated with synaptic integrity, mitochondrial function, inflammation, and brain cell type-specific signaling. Our study highlights the potential of targeting specific pathways and genes (e.g., synaptic, mitochondrial, and inflammatory pathways) to mitigate the adverse effects of anesthesia. Therapeutic interventions could include anti-inflammatory treatments, protection of mitochondrial function, and modulation of synaptic activity through drugs or gene expression manipulation. The identification of specific serum biomarkers provides a valuable tool for diagnosing and prognosticating anesthesia-induced neurotoxicity.

In conclusion, this study provides the first comprehensive molecular and pathological characterization of AIDN using dual human relevant platforms. We demonstrated a clear dose- and exposure duration-dependent neurotoxic response in iPSC-derived human cerebral organoids and identified shared dysregulated of lncRNA-mRNA signatures in both organoids and serum. These findings provide a critical translational bridge between mechanistic research and clinical relevance, validating the utility of these models for biomarker discovery and therapeutic development. Future research should investigate the long-term consequences of anesthetic exposure on brain development, particularly its potential role in neurodevelopmental disorders such as autism spectrum disorder and attention-deficit/hyperactivity disorder. Such longitudinal efforts will be essential for defining long-term risks and developing targeted neuroprotective strategies to safeguard neurodevelopment in vulnerable pediatric populations.

## Acknowledgements

We would like to thank all the participants enrolled in the studies. The authors would like to thank Roland James (Department of Medicine, Medical College of Wisconsin, Milwaukee, WI 53226, USA) for providing administrative, technical and laboratory support.

## Author contributions

This study was initiated and designed by T.P; C.J; X.B. Experiments were performed by T.P; C.J; X.B; Y.Y; B.S; D.L; S.L. The data collection and interpretation were performed by T.P; C.J; X.B. T.P; X.B. Supervision. T.P; C.J; X.B. reviewed the literature and wrote the original manuscript. T.P; C.J; X.B; Z.J.B; B.S; D.L; S.L; A.B; M.J; R.J.B; S.P.T; A.M.H. Read discussed and revised the original manuscript.

## Availability of data and materials

All data needed to evaluate the conclusions in the paper are present in the paper and/or the Supplementary Materials. All microarray gene expression data files have been deposited at The National Center for Biotechnology Information Gene Expression Omnibus (accession number: GSE271591 and GSE271593). Drs Bai, Pant and Jiang have full access to all the data in the study and take responsibility for its integrity and accuracy of its analysis.

## Declarations

### Ethics approval and consent to participate

The study was approved by the Institutional Review Board of the Medical College of Wisconsin (Protocol number 1077311). The nature and purpose of the study were explained to the parents and participants, and informed consent/assent was obtained from those who agreed to participate.

### Consent for publication

All authors have reached consent for publication.

### Author Information

Tarun Pant and Congshan Jiang have contributed equally to this work.

