## Supplementary file for "Bridging Stem Cell Models and Medicine: Integrated 3D Human Cerebral Organoids and Pediatric Serum Reveal Mechanisms and Biomarkers of Anesthetic-Induced Neurotoxicity"

**Supplementary Data**

**Supplementary Figures and Figure legends**

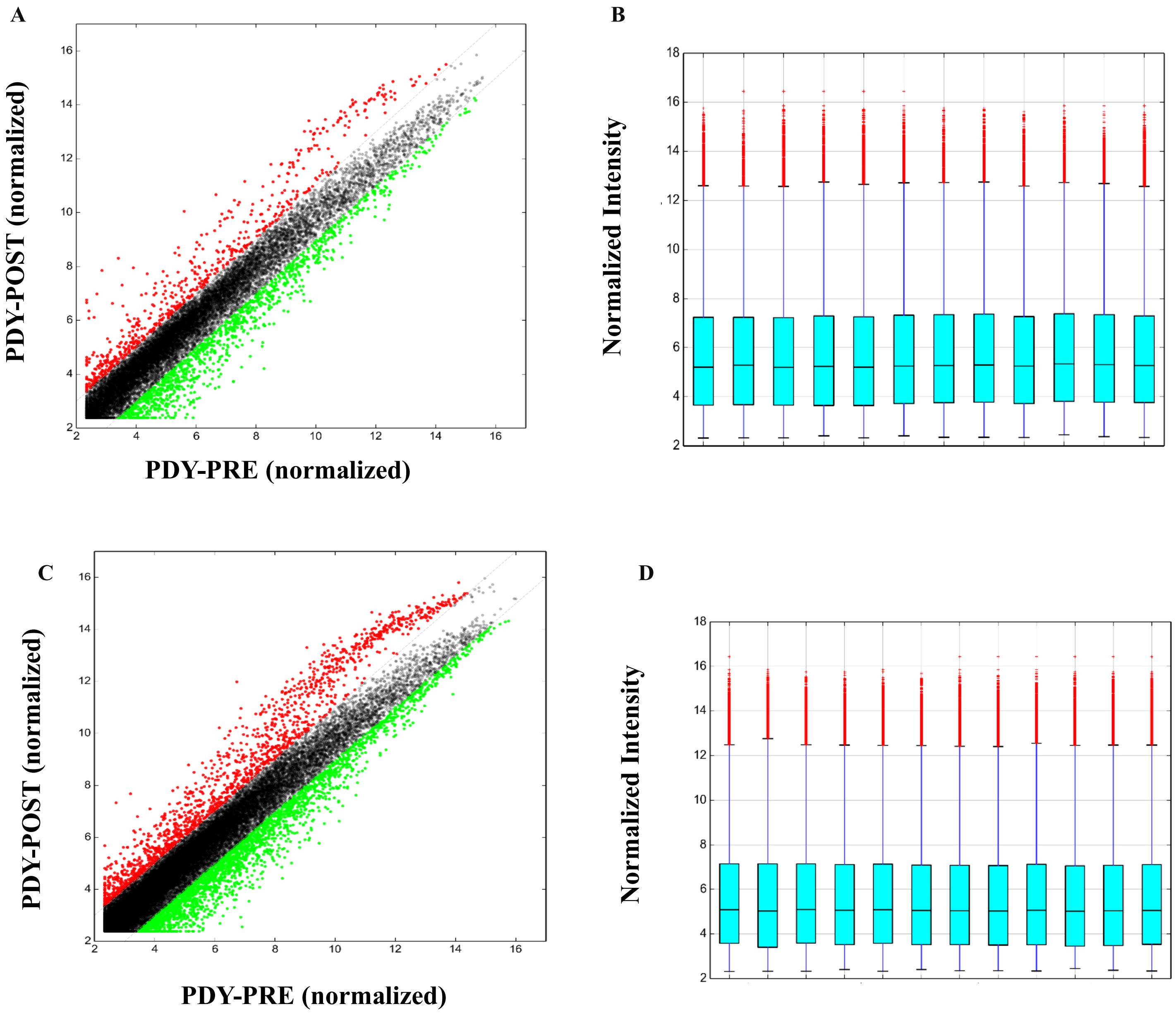


**Figure S1. Microarray and Bioinformatic analysis of serum mRNA and lncRNAs.**

Box and scatter plots for mRNA (A&B); and lncRNAs (C&D); showing the distribution of normalized gene expression intensity values for mRNA in serum of pediatric patient receiving general anesthesia (> 3 hours; n=3 from 10 pooled patient serum samples).

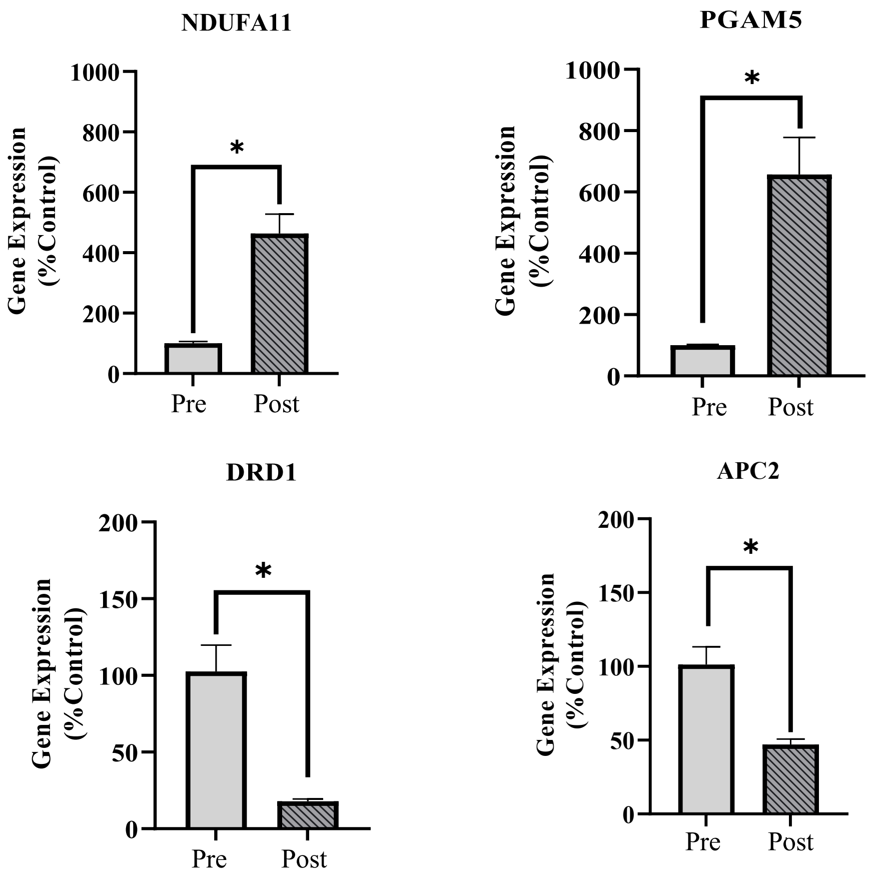


**Figure S2. RT-qPCR validation of microarray data of mRNAs from the serum of pediatric patients (> 3-hour anesthesia/surgery).** qRT-PCR validation of four mRNAs (upregulated: NDUFA11; PGAM5; and downregulated: DRD1; and APC2) from array data from serum of pediatric patients (n=3 from 10 pooled patient serum samples; *P < 0.05).

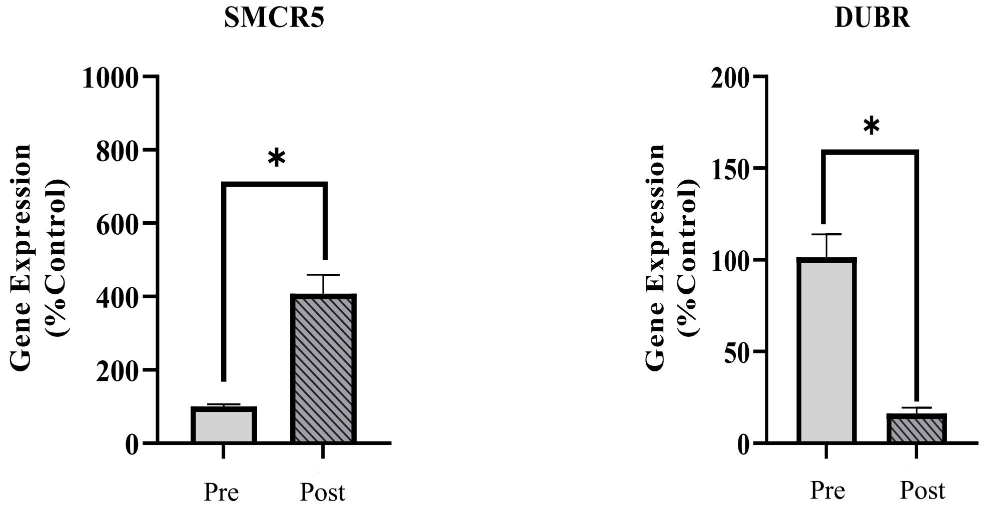


**Figure S3. RT-qPCR validation of microarray data of lncRNAs from the serum of pediatric patients (> 3-hour anesthesia/surgery).** RT-qPCR validation of two anesthesia/surgery-altered lncRNAs (upregulated: SMCR5; downregulated: DUBR) from array data in pediatric patients (n=3 from 10 pooled patient serum samples; *P < 0.05).

### Supplementary Methods

#### Human iPSCs derived cerebral organoid and characterization

In brief, singularized iPSCs were plated at a seeding density of 12,000 cells/well in an ultra-low attachment 96-well plate and cultured with mTeSR1 medium (STEMCELL Technologies, Vancouver, Canada) in the incubator (5% $CO_2$, 21% $O_2$) at 37 °C. The medium was changed every day for 5 days. On Day 6, embryonic bodies were transferred into each well of an ultra-low attachment 24-well plate and cultured with neural induction media for the formation of neuroepithelial tissues. On Day 11, each neuroepithelial tissue was embedded as a Matrigel droplet, and every 16 droplets were cultured with organoid differentiation media without vitamin A in 100-mm Petri dishes. On Day 15, the cerebral organoids were further cultured in the organoid differentiation media with vitamin A on an orbital shaker (at 80 rpm). The media were changed every 3 days. We utilized the MycoAlert® Mycoplasma Detection Kit (Lonza, Basel, Switzerland) to assess potential mycoplasma contamination in our iPSC and organoid cultures. The analysis confirmed that our iPSC cultures were free from mycoplasma contamination. Organoid experiments below were from days 30 and 60 cerebral organoids. Each sample (n = 4) per group was pooled from two organoids for protein and RNA assays. Each sample was generated from 4 independent differentiations of iPSCs cultured in separate dishes.

Briefly, iPSCs cultured on coverslips were fixed and stained with the primary antibodies: anti-stage-specific embryonic antigen 4 (SSEA4, Abcam, Cambridge, MA, USA; ab16287) and anti-octamer-binding transcription factor 4 (OCT4, Sigma Aldrich, St. Louis, MO, USA; AB3209). SSEA4 and OCT4 are pluripotent stem cell markers. Two-month-old cerebral organoids were fixed with 10% zinc formalin (Sigma Aldrich), embedded in paraffin, and sectioned into 4 µm thick slices. The sections were blocked in 10% donkey serum for 30 minutes at room temperature. The tissue sections were then incubated with the following primary antibodies at 4 °C overnight to characterize the presence of the various types of brain cells within organoids: 1) Anti-microtubule-associated protein-2 (MAP2: a neuron marker; Abcam, ab11267), anti-doublecortin (an immature and migrating neuron marker; Abcam ab18723), 2) anti-S100 calcium-binding protein B (S100B: an astrocyte marker; Abcam, ab868), 3) anti-allograft inflammatory factor 1/Ionized Calcium-Binding Adapter Molecule 1 (AIF-1/IBA1: a microglia marker; Novus Biologicals, CO, USA; NB100-1028), 4) anti-von Willebrand Factor (VWF: and endothelial cell marker) (DAKO, CA,

USA; A0082), 5) anti-smooth muscle actin (SMA: a smooth muscle cell marker) (Novus Biologicals, MAB343c), and 6) anti-myelin basic protein (MBP: oligodendrocyte marker; Santa Cruz, Dallas, TX, USA; sc-66064). After washing with phosphate-buffered saline (PBS) three times, the sections were further stained with Alexa Fluor 488-conjugated anti-mouse immunoglobulin G (IgG) or goat IgG and Alexa Fluor 594-conjugated donkey anti-rabbit (Thermo Fisher Scientific, Waltham, MA, USA) for 45 minutes at 37 °C. After three washes with PBS, the nuclei were stained with Hoechst 33342 (Thermo Fisher Scientific). Images of the stained sections were captured using laser scanning confocal microscopy Nikon Eclipse TE2000-U (Nikon, Tokyo, Japan).

**Analysis of anesthetic induced apoptosis and ultrastructure changes on cerebral organoids**

<u>Anesthetic exposure of 3D cerebral organoids:</u> Intravenous anesthetic propofol (2,6-diisopropylphenol) (Sigma Aldrich) is widely used in pediatric anesthesia. The blood concentration of propofol in children varies widely, typically ranging from about 1 to 10 μg/ml. Propofol is a highly lipophilic agent and, therefore, the concentration of propofol is relatively high in lipid-rich tissues such as the brain, with estimated concentrations ranging from 4 to 20 μg/ml.(1-3) In this study, day 60-organoids were exposed to propofol under different conditions. Organoids received either a single exposure to propofol (10 μg/ml) for various lengths of time, including 1, 2, 3, or 6 hours to represents short or prolonged anesthesia during lengthy surgical procedures in pediatric patients, or three exposures to propofol (5 or 10 μg/ml) for 1 or 2 hours per day over three consecutive days to simulates repeated short-duration anesthetic exposures in pediatric patients requiring multiple procedures within a short timeframe. Dimethyl sulfide (DMSO, Sigma Aldrich) (0.1%) served as a vehicle control, as DMSO is the solvent for propofol. Cell apoptosis in the organoids was then evaluated as described in the following section to examine the effects of propofol dose, exposure duration, and frequency on cell viability. Other organoid studies, including protein and gene expression analysis, were conducted based on the single 10 μg/ml propofol exposure condition.

<u>Electron Microscopy:</u> Cerebral organoids were fixed with 2% glutaraldehyde buffer at 4 °C and further prepared as sections with 60 nm thickness for electron microscopy analysis as previously described. The sections were stained with lead citrate and uranyl acetate, and imaged with the H600 Electron Microscope (Hitachi, Japen).

Caspase 3 activity assay: The whole-cell protein lysate of the organoids was used both for the Western blotting and Caspase 3 activity assay. Caspase 3 activity was detected using a caspase 3 colorimetric assay kit (Sigma Aldrich) in cerebral organoids exposed to propofol for various length of time, either once or three times. Manufacturer's instructions were followed. The following formula was used for the final data presentation:

$$\frac{\mu\text{mol (caspase 3 reaction product)} \times \text{dilution factor} \times 100\%}{\text{ml (sample lysate)} \times \text{minutes (reaction time)} \times \text{mg (protein)}}$$

Western blotting: Organoid protein lysates were prepared using RIPA buffer (Cell Signaling Technology, Danvers, MA, USA; #9806), freshly supplemented with phosphatase inhibitor cocktails (Roche Diagnostics, Indianapolis, IN, USA) and phenylmethylsulfonyl fluoride. Primary antibodies used included: anti-microtubule associated protein 1 light chain 3 beta (LC3B, Cell Signaling Technology, #3868), and anti-beta actin (Cell Signaling Technology, #4967S). Secondary antibodies included anti-rabbit and anti-mouse IgG conjugated with horseradish peroxidase (Cell Signaling Technology). The signals on the PVDF (polyvinylidene fluoride or polyvinylidene difluoride) membranes (Bio-Rad, Hercules, CA, USA) were developed using an ECL Detection kit (GE Healthcare Life Sciences, Marlborough, MA, USA) and further captured using a ChemiDoc imaging system (Bio-Rad). Beta actin was used as the endogenous control for protein normalization. The relative expression fold change of each protein was calculated by capturing and analyzing the optical densities of specific bands using the ChemiDoc imaging system (Bio-Rad) and normalizing with beta actin as the endogenous control protein.

**Patient characteristics and demographics**

In total, twenty pediatric patients were enrolled at the Children's Hospital of Wisconsin between September 2018 and October 2019. The study included 10 patients with a surgery duration of less than one hour and 10 patients with surgeries lasting over 3 hours. Patient inclusion criteria were: 1) Patients aged < 4 years and underwent non-neuronal surgeries. 2) Patients were treated with the general anesthetics (propofol, ketamine, isoflurane, or sevoflurane) for less than one hour or over 3 hours and had an existing intravenous (IV) line during the surgical procedure. 3) Patients were treatment-naive of any medications known to affect brain function. There were no drugs involved in this study beyond the anesthetics utilized for the subjects' standard of care surgical procedures. The rationale for selecting general anesthesia lasting more than 3 hours and patients aged under 4 years was based on previous findings indicating that prolonged anesthesia exposure in early life (>3 hours) poses a significant neurotoxicity risk before age five. Patients with anesthesia duration of less than one hour were used as controls for comparison. Exclusion Criteria were: 1) Patients who did not require an IV for their surgery. 2) Anesthetic duration prolonged beyond what was expected due to the blood draw, such as when the actual surgery duration extended beyond one hour but less than three hours. 3) Patients who underwent brain surgery. 4) Patients with a medical history that could affect serum biomarker levels. The study was approved by the Institutional Review Board of the Medical College of Wisconsin (Protocol number 1077311). The nature and purpose of the study were explained to the parents and participants, and informed consent/assent was obtained from those who agreed to participate.

**Clinical study design and analysis of neurotoxicity in pediatric patients**

Whole blood samples were collected from each patient at two time points (one ml per time point per patient) 1) Before IV fluids were administered pre-surgery. 2) Post-surgery, before the patient awoke, and the IV was removed. The tubes with collected blood were immediately placed on ice and allowed to clot undisturbed for 20-30 minutes. They were then centrifuged at 4°C at 300g for 15-20 minutes. Following centrifugation, the serum samples were aliquoted into small volumes in plastic screw-cap vials (100 µL per vial) and immediately stored at -80°C. All patient serum samples stored at -80 °C were thawed on the same day to avoid batch effects. The 10 serum samples from patients receiving either less than one hour or over 3 hours of anesthesia were divided into three replicate pools (3+3+4 samples) to assess protein levels of brain-derived neurotrophic factor (BDNF), neuron-specific enolase (NSE), and S100 calcium-binding protein B (S100B), as serum markers of brain cell injury or stress.

**Reverse transcription-quantitative PCR (RT-qPCR)**

RT-qPCR was used to quantify the expression of coding genes (mRNAs) related to neural stem cells, neurons, and astrocytes within the organoids and to confirm the expression tendency of the randomly selected mRNAs and lncRNAs from microarray assays. Briefly, cDNA was synthesized from total RNA using the Revert Aid™ First Strand cDNA Synthesis Kit (Thermo Fisher Scientific) according to the manufacturer's instructions. Each qPCR reaction included the mixed cDNA, PowerUp™ SYBR™ Green Master Mix (Applied Biosystems), primers, and water, and was conducted in the QuantStudio™ 6 Real-Time PCR machine (Applied Biosystems, Foster City, CA, USA). The sequence of the following primers were listed in (appendix pp 5-6): neurofilament heavy chain (NEFH), neurofilament medium chain (NEFM), galactosylceramidase (GALC), glial fibrillary acidic protein (GFAP), paired box 6 (PAX6), nestin (NES), NADH: ubiquinone oxidoreductase subunit A11 (NDUFA11), mitochondrial serine/threonine protein phosphatase (PGAM5), dopamine receptor D1 (DRD1), APC regulator of WNT signaling pathway 2 (APC2), Smith-Magenis syndrome chromosome region candidate 5 (SMCR5), and DPPA2 upstream binding RNA (DUBR). The relative gene expression fold change was calculated with the mean cycle threshold (Ct) of PCR triplicates for each gene using the $2^{-\Delta\Delta Ct}$ formula and further normalized against glyceraldehyde-3-phosphate dehydrogenase (GAPDH) and actin beta (ACTB) endogenous controls.

**Congruence of the microarray and bioinformatic analyses in organoid and serum studies**

To ensure consistency across datasets, we applied the following uniform methodologies and quality control measures for both organoid and serum studies: **1) Microarray Analysis**: Identical microarray assays were performed by Arraystar Inc. to evaluate anesthetic-induced dysregulation of lncRNAs and mRNAs across organoid and serum samples, ensuring comparable data acquisition. **2) Bioinformatic Analysis**: Normalization and filtering: Raw mRNA data were normalized using the quantile normalization method in GeneSpring GX v12.1, and low-intensity mRNAs were filtered to ensure data reliability. Data quality assessment: Quality control of microarray data included Scatter and Box Plot analyses for mRNAs and lncRNAs (Figures 2B-C; Supplementary Figures 3A-D), confirming data integrity and consistency. Differential expression screening: Differentially expressed mRNAs were identified using Volcano Plot filtering (fold change $\geq$ 1.2, p-value $\leq$ 0.05). Hierarchical clustering and pathway analysis: Hierarchical clustering of differentially expressed mRNAs and pathway analyses (using IPA, mitoXplorer, SynGO, Metascape, KEGG, and Cytoscape analysis) provided insights into key pathways affected by anesthesia exposure. This consistent workflow demonstrated congruence between the organoid and serum analyses, identifying anesthetic-induced overlapping dysregulated gene profiles and pathways.
