## Supplementary tables for "Bridging Stem Cell Models and Medicine: Integrated 3D Human Cerebral Organoids and Pediatric Serum Reveal Mechanisms and Biomarkers of Anesthetic-Induced Neurotoxicity"

Supplementary Table 1. Characteristic of patients in the clinical study

| | | | | | | | | | | | | | | | | | | | | | |
|---|---|---|---|---|---|---|---|---|---|---|---|---|---|---|---|---|---|---|---|---|---|
| | Subject identifiers | BM009 | BM016 | BM018 | BM003 | BM004 | BM047 | BM048 | BM053 | BM056 | BM58 | BM030 | BM035 | BM046 | BM050 | BM055 | BM59 | BM60 | BM65 | BM66 | BM67 |
| | Age (Years) | <4 | <4 | <4 | <4 | <4 | <4 | <4 | <4 | <4 | <4 | <4 | <4 | <4 | <4 | <4 | <4 | <4 | <4 | <4 | <4 |
| | Sex | Female | Male | Male | Male | Male | Female | Female | Male | Male | Male | Male | Female | Female | Male | Male | Male | Male | Male | Female | Female |
| | (hours) | <1 | <1 | <1 | <1 | <1 | <1 | <1 | <1 | <1 | <1 | >3 | >3 | >3 | >3 | >3 | >3 | >3 | >3 | >3 | >3 |
| Characteristics of patient | Surgery type | Non-cardiac | Non-cardiac | Non-cardiac | Non-cardiac | Non-cardiac | Non-cardiac | Non-cardiac | Non-cardiac | Non-cardiac | Non-cardiac | Non-cardiac | Non-cardiac | Non-cardiac | Non-cardiac | Non-cardiac | Non-cardiac | Non-cardiac | Non-cardiac | Non-cardiac | Non-cardiac |
| Intravenous anesthetic agents | PROPOFOL | X | X | X | X | X | X | X | X | X | X | X | X | X | X | X | X | X | X | X | X |
| Inhalational Anesthetic | SEVOFLURANE | X | X | X | X | | X | X | X | X | X | | X | | X | | X | X | X | X | X |
| Local Anesthetic | ROPIVACAINE | | | | | | | | | | | | | | | | X | | | | |
| | NITROUS OXIDE | X | X | X | X | X | | X | X | X | | X | | X | X | | X | | X | X | X |
| | OPIOID | X | X | | X | X | X | | X | X | | X | X | X | | X | X | X | X | X | X |
| Analgesics | DEXMEDETOMIDINE | X | X | | | | X | | | | | | X | X | | X | | X | | X | X |
| | DEXAMETHASONE | X | | | | | | | X | X | | X | X | | | | | | | | |
| | ONDANSETRON | X | | | | | X | | X | X | | | | | | | | | | | |
| | ACETAMINOPHEN | X | | | | | | | | | | | | X | | | X | | | | X |
| | EPINEPHRINE | | | | | | | | | | | | | | | | | | | | X |
| | ROCURONIUM | | | | | | | | | | | | | X | | | | | | | |
| | SUGAMMADEX | | | | | | | | | | | | | X | | | | | X | | |
| | KETOROLAC | X | | | | | | | | | | X | | | | | | | | | |
| | TYLENOL | | | | | | X | | X | X | | | X | | | | | | X | | |
| | CEFAZOLIN | | | | | | | | | | X | | | | | | | | | | |
| | PIPERACILLIN | | | | | | | | | | | | X | | | | | | | | |
| Other medications administered IV | TAZOBACTAM | | | | | | | | | | | | X | | | | | | | | |

**Supplementary Table 2. Primer information**

| Gene ID | Primer sequence (from 5' to 3') | Product length (bps) |
|---|---|---|
| *NEFH* | F: CAGCCAAGGTGAACACAGAC<br>R: GCTGCTGAATGGCTTCCT | 192 |
| *NEFM* | F: GACGGCGCTGAAGGAAATC<br>R: CTCTTCGCCCTGGTGCATAT | 70 |
| *GALC* | F: ACTCTCACCACTGGTCGCAAAG<br>R: GATCAGCAAAGTTTGGAGCTTCAC | 127 |
| *GFAP* | F: CTGGAGAGGAAGATTGAGTCGC<br>R: ACGTCAAGCTCCACATGGACCT | 122 |
| *PAX6* | F: CTGAGGAATCAGAGAAGACAGGC<br>R: ATGGAGCCAGATGTGAAGGAGG | 131 |
| *NES* | F: TGCGGGCTACTGAAAAGTTC<br>R: AGGCTGAGGGACATCTTGAG | 130 |
| *NDUFA11* | F: AGAATCCATCCCTGGAGACA<br>R: GTAATCCCAGTACTTTGGGAGG | 104 |
| *PGAM5* | F: CCGGAAGCTGTGCAGTATTA<br>R: GAGGAGGACAGTTACGAGATC | 106 |
| *DRD1* | F: GACTCCTGGAAATCTGACTGAC<br>R: GAAGATGAGGACTCTGAACACC | 103 |
| *APC2* | F: CAGACAAGGACGGCTCAAA<br>R: GTGTACTGCTTCTACGGCAA | 106 |
| *SMCR5* | F: GTCTCAGTGCTCCTCAAACA<br>R: GTAAGGAACTCCCAGGTCAAG | 111 |
| *DUBR* | F: CGCTAAAGATCCCAGAAAGGT<br>R: CTACACTACACTTGCTCCTCAC | 84 |
| *GAPDH* | F: GTCTCCTCTGACTTCAACAGCG<br>R: ACCACCCTGTTGCTGTAGCCAA | 131 |

Note: bps: base pairs; NEFH: neurofilament heavy chain; NEFM: neurofilament medium chain; GALC: galactosylceramidase; GFAP: glial fibrillary acidic protein; PAX6: paired box 6; NES: nestin; NDUFA11:NADH: ubiquinone oxidoreductase subunit A11; PGAM5: mitochondrial serine/threonine protein phosphatase; DRD1: Dopamine receptor D1;APC2:APC regulator of WNT signaling pathway 2; SMCR5:Smith-Magenis syndrome chromosome region candidate 5; DUBR: DPPA2 upstream binding RNA; GAPDH: glyceraldehyde-3-phosphate dehydrogenase

**Supplementary Table 3 Dysregulated coding genes (mRNAs) in propofol (6 h)-treated cerebral organoids**

| Gene ID | Fold change | P value | Regulation |
|---|---|---|---|
| HOXD11 | 6.001 | 0.030 | Up |
| CRYGC | 4.839 | 0.039 | Down |
| HIST1H1C | 3.645 | 0.001 | Down |
| CRYBA1 | 3.269 | 0.027 | Down |
| HIST1H2BG | 3.050 | 0.004 | Down |
| ANKRD34A | 3.023 | 0.041 | Down |
| CCDC61 | 2.937 | 0.009 | Down |
| CTSF | 2.917 | 0.030 | Up |
| NODAL | 2.655 | 0.021 | Up |
| CCDC141 | 2.613 | 0.017 | Down |
| HIST1H2AE | 2.526 | 0.003 | Down |
| CCDC144NL | 2.515 | 0.001 | Down |
| SLC39A12 | 2.513 | 0.039 | Down |
| POLR3B | 2.477 | 0.016 | Up |
| RP11-1070N10.3 | 2.411 | 0.028 | Down |
| MCIDAS | 2.391 | 0.032 | Down |
| HIST1H3H | 2.385 | 0.008 | Down |
| ELTD1 | 2.365 | 0.050 | Up |
| CRH | 2.344 | 0.014 | Up |
| HIST2H4B | 2.341 | 0.001 | Down |
| HIST1H2BD | 2.312 | 0.011 | Down |
| ACOT4 | 2.263 | 0.039 | Down |
| HIST1H2AI | 2.246 | 0.033 | Down |
| DKK1 | 2.226 | 0.048 | Up |
| HIST2H2AA4 | 2.119 | 0.016 | Down |
| HIST1H1D | 2.073 | 0.008 | Down |
| DNAH17 | 2.037 | 0.013 | Up |
| OR2M2 | 2.034 | 0.014 | Up |
| HIST2H2BE | 2.022 | 0.006 | Down |
| HIST1H2AM | 1.991 | 0.033 | Down |
| ICAM2 | 1.982 | 0.038 | Up |
| FCER2 | 1.956 | 0.036 | Down |
| HIST1H4D | 1.944 | 0.032 | Down |
| SPIN2A | 1.941 | 0.003 | Down |
| HIST2H2AA3 | 1.939 | 0.014 | Down |
| CCDC102B | 1.928 | 0.048 | Up |
| NPY2R | 1.924 | 0.031 | Up |
| HSPA1B | 1.920 | 0.015 | Up |
| ATP6V0D2 | 1.917 | 0.044 | Up |
| KCNJ13 | 1.903 | 0.018 | Down |
| SFRP5 | 1.895 | 0.030 | Down |
| PLVAP | 1.883 | 0.013 | Up |
| UNKL | 1.865 | 0.036 | Down |
| AQP1 | 1.863 | 0.029 | Down |
| HIST2H3C | 1.859 | 0.034 | Down |
| HENMT1 | 1.857 | 0.008 | Down |
| RTDR1 | 1.843 | 0.004 | Down |
| HIST1H2AK | 1.839 | 0.015 | Down |
| PRSS57 | 1.839 | 0.012 | Up |
| HIST1H2AH | 1.828 | 0.008 | Down |
| ZNF853 | 1.826 | 0.010 | Up |
| H1F0 | 1.824 | 0.010 | Down |
| FW339998 | 1.821 | 0.032 | Down |
| C20orf112 | 1.800 | 0.002 | Down |
| GPRC6A | 1.773 | 0.032 | Up |
| GPRIN2 | 1.758 | 0.031 | Down |
| SDIM1 | 1.739 | 0.008 | Up |
| H1FX | 1.736 | 0.041 | Down |
| CCDC151 | 1.725 | 0.035 | Down |
| CELF2 | 1.722 | 0.037 | Down |
| CDC20B | 1.718 | 0.050 | Up |

| | | | |
|---|---|---|---|
| CMKLR1 | 1.714 | 0.043 | Up |
| CABP5 | 1.702 | 0.021 | Up |
| GPSM1 | 1.702 | 0.035 | Up |
| AC110771.1 | 1.699 | 0.042 | Down |
| FLJ41484 | 1.687 | 0.036 | Down |
| TMCC2 | 1.682 | 0.038 | Down |
| BC033739 | 1.678 | 0.002 | Up |
| NLRP14 | 1.671 | 0.007 | Down |
| RP11-834C11.12 | 1.663 | 0.027 | Up |
| AL050303.1 | 1.661 | 0.039 | Down |
| TAS2R10 | 1.639 | 0.025 | Up |
| ATP2B4 | 1.639 | 0.026 | Down |
| ABTB1 | 1.636 | 0.035 | Down |
| ADCY4 | 1.636 | 0.043 | Up |
| RP11-201K10.3 | 1.630 | 0.027 | Down |
| RWDD2B | 1.630 | 0.017 | Down |
| EEPD1 | 1.625 | 0.027 | Down |
| LOH12CR2 | 1.608 | 0.049 | Down |
| PTCH1 | 1.601 | 0.006 | Down |
| MFSD4 | 1.597 | 0.006 | Down |
| IL18RAP | 1.592 | 0.019 | Up |
| NAT16 | 1.587 | 0.007 | Down |
| ERC2 | 1.586 | 0.014 | Down |
| IBA57 | 1.585 | 0.044 | Down |
| KLB | 1.584 | 0.026 | Up |
| FILIP1 | 1.581 | 0.047 | Down |
| GPSM2 | 1.579 | 0.049 | Down |
| MYOM2 | 1.573 | 0.023 | Down |
| HIST1H2BC | 1.557 | 0.039 | Down |
| DNAJB1 | 1.555 | 0.046 | Up |
| CALB2 | 1.549 | 0.031 | Down |
| NLRP10 | 1.549 | 0.048 | Up |
| ARIH2OS | 1.547 | 0.003 | Down |
| FAM13C | 1.545 | 0.040 | Down |
| HIST1H2BM | 1.544 | 0.048 | Down |
| LOC643355 | 1.543 | 0.004 | Down |
| FBXL22 | 1.534 | 0.026 | Down |
| C16orf3 | 1.530 | 0.037 | Up |
| CCDC110 | 1.529 | 0.015 | Down |
| F13A1 | 1.528 | 0.042 | Up |
| CHORDC1 | 1.518 | 0.009 | Up |
| CTC-241N9.1 | 1.516 | 0.047 | Down |
| PIGL | 1.516 | 0.005 | Up |
| CCDC28A | 1.515 | 0.027 | Down |
| KCNF1 | 1.515 | 0.040 | Down |
| SFTPC | 1.514 | 0.033 | Down |
| PNPLA1 | 1.511 | 0.034 | Up |
| SLC17A8 | 1.506 | 0.047 | Down |
| PABPC1L | 1.506 | 0.020 | Up |
| MTCP1 | 1.505 | 0.026 | Down |
| BAG3 | 1.502 | 0.012 | Up |
| RGS1 | 1.501 | 0.038 | Up |
| RNF182 | 1.498 | 0.018 | Down |
| DAB1 | 1.495 | 0.010 | Up |
| TXNRD3NB | 1.492 | 0.023 | Up |
| TMEM52B | 1.491 | 0.037 | Up |
| PAQR9 | 1.490 | 0.038 | Down |
| CALCB | 1.489 | 0.003 | Up |
| AX747379 | 1.486 | 0.010 | Down |
| AK304826 | 1.485 | 0.015 | Down |
| TSPY4 | 1.484 | 0.035 | Down |
| CHURC1 | 1.483 | 0.032 | Down |
| PBXIP1 | 1.483 | 0.050 | Down |
| TAS2R60 | 1.481 | 0.015 | Up |
| XBP1 | 1.480 | 0.044 | Up |

| | | | |
|---|---|---|---|
| TP53I3 | 1.480 | 0.019 | Down |
| DQ576756 | 1.479 | 0.023 | Down |
| SEC31B | 1.479 | 0.003 | Up |
| BMP2 | 1.478 | 0.025 | Up |
| HELB | 1.477 | 0.039 | Down |
| SPTB | 1.476 | 0.019 | Down |
| NTN5 | 1.475 | 0.011 | Up |
| PTPRN2 | 1.474 | 0.029 | Down |
| FAM46D | 1.472 | 0.030 | Down |
| MPO | 1.471 | 0.042 | Down |
| FUT11 | 1.469 | 0.030 | Down |
| SDF2L1 | 1.468 | 0.005 | Up |
| CCDC78 | 1.467 | 0.017 | Up |
| ACER2 | 1.466 | 0.043 | Down |
| TCP10L | 1.462 | 0.005 | Up |
| TTC30A | 1.456 | 0.003 | Down |
| PTH2 | 1.453 | 0.011 | Up |
| CACYBP | 1.452 | 0.013 | Up |
| AP001468.1 | 1.451 | 0.026 | Up |
| ZC3H18 | 1.450 | 0.016 | Up |
| ZC3H4 | 1.449 | 0.019 | Down |
| LOC81691 | 1.447 | 0.030 | Down |
| ATAD3C | 1.445 | 0.019 | Up |
| TNFSF18 | 1.444 | 0.010 | Down |
| FPGT | 1.441 | 0.028 | Down |
| DSC1 | 1.439 | 0.014 | Up |
| DNAJB3 | 1.438 | 0.008 | Up |
| KLHL24 | 1.437 | 0.016 | Down |
| CHRNA3 | 1.436 | 0.000 | Up |
| FAM195A | 1.433 | 0.003 | Up |
| UCP2 | 1.429 | 0.021 | Down |
| HSPA4L | 1.429 | 0.005 | Up |
| DNAJC19 | 1.428 | 0.041 | Down |
| SEPSECS | 1.427 | 0.009 | Down |
| UBE4A | 1.427 | 0.028 | Up |
| MANF | 1.424 | 0.004 | Up |
| FAM118A | 1.421 | 0.029 | Up |
| HBP1 | 1.419 | 0.011 | Down |
| GTF2IRD2 | 1.413 | 0.011 | Down |
| ACTN1 | 1.413 | 0.040 | Up |
| MICALL2 | 1.413 | 0.029 | Up |
| ALDOC | 1.410 | 0.046 | Down |
| HIST1H4H | 1.410 | 0.012 | Down |
| QSOX2 | 1.409 | 0.003 | Up |
| FAM120A | 1.409 | 0.035 | Up |
| CRELD2 | 1.406 | 0.019 | Up |
| RASL10A | 1.406 | 0.043 | Down |
| CHODL | 1.405 | 0.028 | Down |
| KIRREL | 1.404 | 0.021 | Up |
| PDE3B | 1.404 | 0.014 | Up |
| SCNN1D | 1.403 | 0.026 | Up |
| C1QTNF9B | 1.400 | 0.020 | Down |
| AC004899.1 | 1.400 | 0.025 | Down |
| CTIF | 1.399 | 0.005 | Up |
| S100A7A | 1.399 | 0.008 | Down |
| IPO7 | 1.398 | 0.003 | Up |
| RP11-1105G2.3 | 1.397 | 0.024 | Up |
| ANAPC15 | 1.396 | 0.034 | Down |
| RHOF | 1.395 | 0.006 | Up |
| STIP1 | 1.395 | 0.023 | Up |
| PDGFD | 1.395 | 0.045 | Down |
| IFRD2 | 1.394 | 0.041 | Up |
| IBA57-AS1 | 1.392 | 0.039 | Down |
| C21orf119 | 1.392 | 0.043 | Down |
| ST3GAL2 | 1.388 | 0.036 | Down |

| | | | |
|---|---|---|---|
| VSNL1 | 1.386 | 0.005 | Down |
| LINGO3 | 1.384 | 0.050 | Down |
| SEC31A | 1.383 | 0.039 | Up |
| ALDH8A1 | 1.382 | 0.012 | Up |
| AARD | 1.382 | 0.009 | Down |
| JMJD6 | 1.381 | 0.023 | Up |
| FAM218A | 1.380 | 0.030 | Down |
| BIVM-ERCC5 | 1.378 | 0.021 | Up |
| SIPA1L2 | 1.378 | 0.007 | Up |
| TCP10 | 1.376 | 0.002 | Up |
| SYT14 | 1.373 | 0.027 | Down |
| CMPK2 | 1.371 | 0.041 | Down |
| PPRC1 | 1.371 | 0.015 | Up |
| TSPAN7 | 1.371 | 0.041 | Down |
| AL592284.1 | 1.371 | 0.045 | Down |
| ELFN1 | 1.371 | 0.050 | Up |
| CASP10 | 1.370 | 0.021 | Up |
| NOP56 | 1.370 | 0.029 | Up |
| HSPE1 | 1.369 | 0.000 | Up |
| SRCAP | 1.368 | 0.024 | Up |
| C21orf2 | 1.368 | 0.001 | Up |
| MAP2 | 1.367 | 0.026 | Down |
| MTR | 1.366 | 0.001 | Up |
| TPST2 | 1.366 | 0.023 | Up |
| DHX40 | 1.365 | 0.010 | Down |
| ARHGAP8 | 1.364 | 0.038 | Down |
| LRRC36 | 1.364 | 0.039 | Down |
| SRPRB | 1.363 | 0.010 | Up |
| DUS1L | 1.362 | 0.032 | Up |
| LRRC17 | 1.360 | 0.043 | Down |
| PDCD4 | 1.360 | 0.029 | Down |
| FKBP4 | 1.360 | 0.038 | Up |
| MYO19 | 1.360 | 0.001 | Up |
| TMEM14A | 1.359 | 0.050 | Down |
| ABCC9 | 1.359 | 0.021 | Down |
| NRP2 | 1.359 | 0.002 | Up |
| CREBRF | 1.358 | 0.020 | Down |
| HSH2D | 1.357 | 0.021 | Up |
| C15orf52 | 1.355 | 0.037 | Down |
| GPR27 | 1.354 | 0.050 | Down |
| ADGRL2 | 1.354 | 0.004 | Up |
| LAS1L | 1.352 | 0.012 | Up |
| DCDC2C | 1.352 | 0.028 | Up |
| YPEL5 | 1.352 | 0.029 | Down |
| ENTPD7 | 1.351 | 0.021 | Up |
| METTL1 | 1.350 | 0.006 | Up |
| OPLAH | 1.350 | 0.049 | Up |
| MALAT1 | 1.350 | 0.013 | Up |
| PKM | 1.350 | 0.039 | Down |
| CABP1 | 1.349 | 0.032 | Down |
| IRF3 | 1.349 | 0.011 | Up |
| TMEM17 | 1.348 | 0.015 | Down |
| FKBP10 | 1.345 | 0.014 | Up |
| ATAD3A | 1.343 | 0.040 | Up |
| WWC2 | 1.343 | 0.018 | Up |
| FAM214A | 1.343 | 0.021 | Down |
| C6orf226 | 1.342 | 0.031 | Down |
| KY | 1.342 | 0.027 | Down |
| NRL | 1.341 | 0.022 | Down |
| LRLE1 | 1.341 | 0.048 | Down |
| SUDS3 | 1.340 | 0.029 | Up |
| SLC2A8 | 1.340 | 0.030 | Up |
| Em:AC005003.4 | 1.338 | 0.013 | Down |
| HSP90AA1 | 1.338 | 0.001 | Up |
| PUS7 | 1.337 | 0.039 | Up |

| | | | |
|---|---|---|---|
| AC103801.2 | 1.336 | 0.048 | Down |
| RHD | 1.336 | 0.004 | Up |
| PNN | 1.334 | 0.037 | Up |
| SMKR1 | 1.334 | 0.007 | Down |
| VCAM1 | 1.333 | 0.041 | Down |
| PRPF38B | 1.331 | 0.019 | Up |
| PDCD11 | 1.330 | 0.037 | Up |
| PNPLA8 | 1.330 | 0.031 | Down |
| AP1G2 | 1.329 | 0.047 | Up |
| NFAT5 | 1.328 | 0.008 | Up |
| CHRNB3 | 1.327 | 0.005 | Up |
| NME9 | 1.325 | 0.008 | Down |
| MCM3AP | 1.325 | 0.048 | Up |
| MYF6 | 1.325 | 0.021 | Down |
| C12orf76 | 1.324 | 0.034 | Down |
| TMTC2 | 1.324 | 0.023 | Down |
| PDK1 | 1.324 | 0.011 | Down |
| NOV | 1.324 | 0.043 | Down |
| IL17RC | 1.323 | 0.021 | Down |
| EPDR1 | 1.323 | 0.006 | Down |
| GDPD3 | 1.323 | 0.009 | Up |
| OGG1 | 1.321 | 0.005 | Down |
| TSEN54 | 1.320 | 0.012 | Up |
| WARS2 | 1.320 | 0.036 | Down |
| CCDC154 | 1.320 | 0.028 | Up |
| AC131935.1 | 1.318 | 0.043 | Up |
| ATAD3B | 1.318 | 0.018 | Up |
| COL7A1 | 1.318 | 0.021 | Up |
| LRRC55 | 1.318 | 0.045 | Down |
| BANP | 1.315 | 0.034 | Up |
| SEPT9 | 1.315 | 0.025 | Up |
| RANBP17 | 1.315 | 0.021 | Up |
| UTP14C | 1.314 | 0.009 | Up |
| DPYSL5 | 1.314 | 0.039 | Down |
| LUZP1 | 1.313 | 0.044 | Up |
| MCTP2 | 1.313 | 0.047 | Up |
| CHD3 | 1.312 | 0.001 | Down |
| CNN2 | 1.311 | 0.032 | Up |
| ADGRF3 | 1.311 | 0.011 | Up |
| DNAJB11 | 1.309 | 0.000 | Up |
| CCDC57 | 1.309 | 0.003 | Up |
| YARS | 1.308 | 0.004 | Up |
| POLR3E | 1.307 | 0.010 | Up |
| DDX31 | 1.307 | 0.011 | Up |
| ATMIN | 1.307 | 0.023 | Up |
| AGER | 1.306 | 0.011 | Up |
| AKNAD1 | 1.305 | 0.025 | Down |
| UTP18 | 1.303 | 0.009 | Up |
| GAS1 | 1.303 | 0.013 | Down |
| BNIP3L | 1.303 | 0.046 | Down |
| CCNL2 | 1.303 | 0.029 | Up |
| APITD1 | 1.301 | 0.007 | Down |
| MAP3K6 | 1.301 | 0.028 | Up |
| ZNF154 | 1.300 | 0.048 | Up |
| NRDE2 | 1.300 | 0.012 | Down |
| KDM5C | 1.299 | 0.015 | Up |
| ABCG1 | 1.298 | 0.016 | Down |
| MDN1 | 1.298 | 0.010 | Up |
| CCDC86 | 1.298 | 0.006 | Up |
| PTRH2 | 1.295 | 0.000 | Up |
| BAZ2A | 1.295 | 0.004 | Up |
| KAT2A | 1.295 | 0.025 | Up |
| NANS | 1.294 | 0.032 | Up |
| STOML1 | 1.294 | 0.013 | Down |
| PCGF5 | 1.292 | 0.028 | Down |

| | | | |
|---|---|---|---|
| KIAA1024L | 1.292 | 0.007 | Down |
| HSPA8 | 1.292 | 0.045 | Up |
| ATP6V0A2 | 1.292 | 0.028 | Up |
| HERC4 | 1.291 | 0.003 | Up |
| PUS1 | 1.291 | 0.003 | Up |
| GATAD2A | 1.291 | 0.047 | Up |
| PNP | 1.291 | 0.036 | Up |
| PNMA6C | 1.290 | 0.036 | Down |
| LATS2 | 1.290 | 0.015 | Down |
| C1GALT1C1L | 1.290 | 0.042 | Down |
| PPAN-P2RY11 | 1.290 | 0.041 | Up |
| DUSP28 | 1.290 | 0.047 | Down |
| SLC26A6 | 1.289 | 0.030 | Up |
| ATG10 | 1.288 | 0.031 | Down |
| AK096982 | 1.288 | 0.045 | Up |
| DUSP4 | 1.287 | 0.027 | Up |
| NOL6 | 1.286 | 0.003 | Up |
| ELF2 | 1.286 | 0.023 | Down |
| GABARAPL3 | 1.286 | 0.011 | Down |
| NOCT | 1.286 | 0.030 | Up |
| SERINC1 | 1.284 | 0.046 | Down |
| LRWD1 | 1.284 | 0.017 | Up |
| CTAGE8 | 1.284 | 0.010 | Up |
| KHDC3L | 1.284 | 0.021 | Up |
| CHMP1B | 1.283 | 0.025 | Down |
| IGF2R | 1.282 | 0.048 | Up |
| ANKRD32 | 1.282 | 0.005 | Up |
| ZCCHC2 | 1.281 | 0.008 | Up |
| DNAJC10 | 1.281 | 0.028 | Up |
| GABPB2 | 1.281 | 0.017 | Down |
| NELL1 | 1.281 | 0.011 | Down |
| CCDC17 | 1.280 | 0.039 | Up |
| RNASET2 | 1.279 | 0.020 | Up |
| GOLGA8T | 1.279 | 0.027 | Up |
| VPS13B | 1.279 | 0.007 | Up |
| RCVRN | 1.279 | 0.014 | Up |
| LYRM7 | 1.279 | 0.033 | Down |
| CCBL2 | 1.278 | 0.046 | Down |
| MTRNR2L4 | 1.278 | 0.011 | Down |
| NOP16 | 1.278 | 0.027 | Up |
| POMGNT1 | 1.278 | 0.048 | Up |
| LGALS16 | 1.278 | 0.045 | Up |
| RPS6KA5 | 1.277 | 0.046 | Down |
| NAT10 | 1.277 | 0.013 | Up |
| SMYD4 | 1.277 | 0.046 | Down |
| EPT1 | 1.277 | 0.045 | Up |
| SYTL3 | 1.276 | 0.034 | Down |
| DPYSL3 | 1.276 | 0.048 | Down |
| STK39 | 1.275 | 0.039 | Down |
| DKC1 | 1.275 | 0.022 | Up |
| MT-ND1 | 1.274 | 0.046 | Up |
| C14orf93 | 1.274 | 0.044 | Down |
| AC132192.1 | 1.274 | 0.044 | Down |
| MGEA5 | 1.273 | 0.031 | Down |
| EARS2 | 1.272 | 0.040 | Up |
| FKBP1B | 1.272 | 0.041 | Down |
| ZDHHC11B | 1.271 | 0.047 | Up |
| KRTAP4-12 | 1.271 | 0.032 | Up |
| RRP1 | 1.271 | 0.007 | Up |
| FOXO3 | 1.271 | 0.020 | Down |
| CORIN | 1.270 | 0.031 | Up |
| WWP2 | 1.270 | 0.021 | Down |
| CBX7 | 1.270 | 0.031 | Up |
| ADAM15 | 1.269 | 0.006 | Up |
| HSP90AB1 | 1.269 | 0.037 | Up |

| | | | |
|---|---|---|---|
| OLFM1 | 1.269 | 0.033 | Down |
| TMEM104 | 1.269 | 0.041 | Up |
| FAM156B | 1.269 | 0.013 | Up |
| TNFRSF25 | 1.268 | 0.034 | Up |
| MIER2 | 1.268 | 0.040 | Up |
| SPOP | 1.267 | 0.001 | Down |
| ARHGAP32 | 1.267 | 0.015 | Up |
| TAF15 | 1.267 | 0.034 | Up |
| NLRP2 | 1.266 | 0.000 | Up |
| CKMT1A | 1.265 | 0.001 | Down |
| CPSF6 | 1.264 | 0.046 | Up |
| CRELD1 | 1.263 | 0.006 | Up |
| CEP290 | 1.263 | 0.026 | Up |
| DUSP18 | 1.262 | 0.030 | Down |
| PGS1 | 1.262 | 0.007 | Up |
| LDAH | 1.260 | 0.030 | Up |
| RP11-231C14.4 | 1.259 | 0.017 | Up |
| AC114546.1 | 1.258 | 0.050 | Up |
| ZDHHC11 | 1.258 | 0.008 | Up |
| ABCF2 | 1.258 | 0.045 | Up |
| FAM27E2 | 1.257 | 0.012 | Up |
| OTULIN | 1.257 | 0.049 | Up |
| EXOSC6 | 1.257 | 0.006 | Up |
| CREBL2 | 1.256 | 0.017 | Down |
| ELMO3 | 1.256 | 0.046 | Up |
| MRPL12 | 1.256 | 0.037 | Up |
| RAB40AL | 1.255 | 0.032 | Up |
| CHKA | 1.255 | 0.006 | Up |
| SNRPA | 1.255 | 0.022 | Down |
| DSCAML1 | 1.255 | 0.024 | Down |
| NUP188 | 1.254 | 0.030 | Up |
| GPR89C | 1.254 | 0.029 | Up |
| SEC23IP | 1.253 | 0.040 | Up |
| PMPCA | 1.253 | 0.002 | Up |
| FAM107B | 1.253 | 0.042 | Down |
| FGF20 | 1.252 | 0.040 | Up |
| SATB1 | 1.251 | 0.019 | Down |
| NDUFAF7 | 1.251 | 0.043 | Up |
| TBC1D14 | 1.251 | 0.021 | Down |
| ING4 | 1.251 | 0.037 | Down |
| UNC13C | 1.250 | 0.046 | Up |
| WRB | 1.250 | 0.008 | Down |
| EXOC7 | 1.249 | 0.010 | Up |
| GABARAPL1 | 1.249 | 0.013 | Down |
| VEZF1 | 1.248 | 0.013 | Down |
| ZNF672 | 1.248 | 0.001 | Up |
| FAM217B | 1.248 | 0.037 | Down |
| PTPRK | 1.248 | 0.029 | Up |
| DUSP7 | 1.247 | 0.042 | Up |
| HSPA1L | 1.246 | 0.031 | Up |
| SGCB | 1.246 | 0.014 | Down |
| LYPD3 | 1.246 | 0.042 | Down |
| H2AFV | 1.245 | 0.020 | Down |
| SDAD1 | 1.245 | 0.019 | Down |
| ISOC2 | 1.245 | 0.007 | Up |
| TRAP1 | 1.244 | 0.027 | Up |
| SRRM1 | 1.244 | 0.000 | Up |
| TTC17 | 1.244 | 0.014 | Up |
| IGF2BP2 | 1.244 | 0.030 | Up |
| AHSA1 | 1.244 | 0.043 | Up |
| XCR1 | 1.244 | 0.027 | Up |
| SPRYD3 | 1.243 | 0.008 | Down |
| PGM3 | 1.243 | 0.000 | Up |
| BTBD9 | 1.242 | 0.013 | Down |
| LARP1B | 1.242 | 0.010 | Up |

| | | | |
|---|---|---|---|
| BRCA1 | 1.242 | 0.021 | Down |
| MBTPS2 | 1.241 | 0.049 | Down |
| CCDC87 | 1.241 | 0.021 | Down |
| KIAA1658 | 1.240 | 0.048 | Up |
| USP2 | 1.239 | 0.030 | Down |
| AC068533.7 | 1.239 | 0.002 | Up |
| M6PR | 1.238 | 0.018 | Up |
| GAPVD1 | 1.238 | 0.009 | Up |
| BAG1 | 1.238 | 0.025 | Down |
| SLC9B2 | 1.237 | 0.028 | Up |
| AURKAIP1 | 1.237 | 0.005 | Up |
| GNL3 | 1.237 | 0.010 | Up |
| CHRNA7 | 1.235 | 0.048 | Down |
| PHB | 1.235 | 0.034 | Up |
| C4orf48 | 1.234 | 0.014 | Down |
| SNUPN | 1.234 | 0.008 | Down |
| MEST | 1.233 | 0.022 | Down |
| FICD | 1.232 | 0.005 | Up |
| RHBDD3 | 1.232 | 0.012 | Up |
| UBE2Q2 | 1.231 | 0.048 | Down |
| INPPL1 | 1.231 | 0.010 | Up |
| RTEL1 | 1.231 | 0.001 | Up |
| PAIP2 | 1.231 | 0.002 | Down |
| KIAA1257 | 1.231 | 0.031 | Down |
| PRRC1 | 1.230 | 0.027 | Up |
| CKMT1B | 1.230 | 0.006 | Down |
| FBXO18 | 1.230 | 0.038 | Up |
| NOP2 | 1.230 | 0.037 | Up |
| PNO1 | 1.229 | 0.003 | Up |
| ID4 | 1.229 | 0.002 | Up |
| ROBO3 | 1.229 | 0.049 | Up |
| HM13 | 1.229 | 0.029 | Up |
| KIAA1731 | 1.229 | 0.043 | Up |
| SLC4A1AP | 1.229 | 0.031 | Up |
| CHAMP1 | 1.228 | 0.015 | Down |
| MANSC1 | 1.228 | 0.031 | Down |
| BC150535 | 1.227 | 0.018 | Down |
| ZBTB44 | 1.227 | 0.026 | Down |
| SHC2 | 1.226 | 0.029 | Up |
| PPP2R1A | 1.226 | 0.026 | Down |
| MSRA | 1.224 | 0.017 | Down |
| DTX2 | 1.223 | 0.044 | Up |
| BX648961 | 1.222 | 0.037 | Up |
| DNAJC25 | 1.222 | 0.038 | Up |
| FXYD6 | 1.222 | 0.047 | Down |
| HSD17B3 | 1.222 | 0.005 | Up |
| SLC25A37 | 1.221 | 0.033 | Up |
| ZSWIM8 | 1.220 | 0.039 | Up |
| RP11-176H8.1 | 1.220 | 0.042 | Down |
| TRPV6 | 1.220 | 0.039 | Up |
| MXI1 | 1.219 | 0.004 | Down |
| ZBED6 | 1.218 | 0.049 | Up |
| DNAJC16 | 1.217 | 0.032 | Up |
| HSPA4 | 1.217 | 0.038 | Up |
| NBR1 | 1.217 | 0.006 | Down |
| RABEPK | 1.216 | 0.006 | Up |
| CCDC85B | 1.216 | 0.026 | Up |
| KCNK7 | 1.216 | 0.047 | Up |
| PON2 | 1.215 | 0.019 | Up |
| RP11-343C2.11 | 1.215 | 0.041 | Up |
| SERP1 | 1.215 | 0.037 | Up |
| DDX51 | 1.214 | 0.020 | Up |
| NUDT15 | 1.214 | 0.017 | Down |
| SLC45A4 | 1.214 | 0.032 | Up |
| HLA-DOB | 1.213 | 0.035 | Up |

| | | | |
|---|---|---|---|
| LBH | 1.212 | 0.050 | Up |
| SRRD | 1.212 | 0.040 | Up |
| DNAJB4 | 1.211 | 0.007 | Up |
| FLNC | 1.211 | 0.011 | Up |
| AHSA2 | 1.211 | 0.039 | Up |
| SRFBP1 | 1.210 | 0.042 | Up |
| CCDC58 | 1.210 | 0.006 | Up |
| ZNF662 | 1.210 | 0.039 | Up |
| RWDD4 | 1.210 | 0.045 | Up |
| HIPK4 | 1.210 | 0.039 | Up |
| CDC37 | 1.209 | 0.037 | Up |
| INPP5K | 1.209 | 0.036 | Down |
| NDUFAF4 | 1.208 | 0.006 | Up |
| KCTD9 | 1.207 | 0.040 | Down |
| METTL3 | 1.207 | 0.030 | Up |
| PITPNM2 | 1.207 | 0.048 | Up |
| LCE1C | 1.206 | 0.024 | Up |
| SCYL1 | 1.205 | 0.049 | Up |
| TENM4 | 1.205 | 0.011 | Up |
| PRR18 | 1.204 | 0.040 | Down |
| TYSND1 | 1.204 | 0.001 | Up |
| NEK1 | 1.204 | 0.027 | Up |
| MON2 | 1.203 | 0.019 | Up |
| CCS | 1.203 | 0.042 | Up |
| B9D2 | 1.203 | 0.005 | Down |
| CRBN | 1.202 | 0.033 | Down |
| GATA4 | 1.202 | 0.039 | Up |
| EIF2B4 | 1.202 | 0.013 | Up |
| SLC25A19 | 1.202 | 0.041 | Up |
| ZNF700 | 1.201 | 0.028 | Up |
| RAD52 | 1.201 | 0.022 | Up |
| CTTNBP2NL | 1.201 | 0.045 | Down |
| SLC31A2 | 1.201 | 0.049 | Up |
| TUBG2 | 1.201 | 0.037 | Down |
| SOS1 | 1.200 | 0.003 | Up |
| AC020922.1 | 1.200 | 0.037 | Down |
| MICAL3 | 1.200 | 0.017 | Up |

**Supplementary Table 4. IPA analysis predicts association of the propofol-induced dysregulated mRNAs transcripts with neural phenotypes in propofol (6 hours)-treated cerebral organoids**

| Phenotype observed | Predicted genes involved (vs. control) |
|---|---|
| *Calcium uptake* | ↑CGA,↓NEB, ↑PTH,↓S100G,↓TTN |
| Neurite morphology | ↑BMP2,↑CRH,↓DPYSL3,↓FCER2, ↓MAP2,↑NRP2,↓ SLC39A12,↑TENM4 |
| *Autophagy* | ↑BAG3,↓BRCA1,↓GABARAPL1,↓TBC1D14,↓UCP2,↑XBP1 |
| *Endoplasmic reticulum stress* | ↑CMKLR1,↑GPSM1,↑ICAM2,↑NLRP10 |
| *Dendrite projection* | ↑CMKLR1,↑GPSM1,↑ICAM2,↑NLRP10 |
| *Hippocampus CA3 neuron excitation* | ↑CRH |
| *Abnormal neurotransmitter release* | ↑CABP5,↑CRH,↓ERC2,↑NPY2R |
| *CNS afterhyperpolarization* | ↓ATP2B4 |
| *Peri- or neonatal death* | ↓ACTA1,↑APOB,↓CASP8,↑CTSF,↑DKK1,↓DLX5,↑GDF3,↓ F5,↑HAND2, ↑HAPLN1,↓HIST1H1C,↓ HIST2H2AA3/HIST2H2AA4,↑HOXD11,↑HPR,↑IDO1,↓ |
| *Calcium absorption* | ↑PTH,↓S100G,↓TNNC1 |
| *Apoptosis initiation* | ↓CASP8,↑CRH,↑PTH |

The genes expressing more than ±1.2-fold difference between the propofol and vehicle control groups were analyzed by IPA software to predict neural phenotypes.

↑: up-regulated following propofol exposure vs. control group.

↓: downregulated following propofol exposure vs. control group.

**Supplementary Table 5 Dysregulated synaptic genes in propofol exposed organoids**

| Gene ID | P value | Fold change | Regulation |
|---|---|---|---|
| ATP2B4 | 0.026 | 1.639 | Down |
| PTCH1 | 0.006 | 1.601 | Down |
| ERC2 | 0.014 | 1.586 | Down |
| FILIP1 | 0.047 | 1.581 | Down |
| GPSM2 | 0.049 | 1.579 | Down |
| CALB2 | 0.031 | 1.549 | Down |
| SLC17A8 | 0.047 | 1.506 | Down |
| SPTB | 0.019 | 1.476 | Down |
| PTPRN2 | 0.029 | 1.474 | Down |
| SEPSECS | 0.009 | 1.427 | Down |
| ALDOC | 0.046 | 1.410 | Down |
| TSPAN7 | 0.041 | 1.371 | Down |
| MAP2 | 0.026 | 1.367 | Down |
| CABP1 | 0.032 | 1.349 | Down |
| DPYSL5 | 0.039 | 1.314 | Down |
| RPS6KA5 | 0.046 | 1.277 | Down |
| DPYSL3 | 0.048 | 1.276 | Down |
| OLFM1 | 0.033 | 1.269 | Down |
| BTBD9 | 0.013 | 1.242 | Down |
| CHRNA7 | 0.048 | 1.235 | Down |
| PPP2R1A | 0.026 | 1.226 | Down |
| FXYD6 | 0.047 | 1.222 | Down |
| SOS1 | 0.003 | 1.200 | Up |
| TENM4 | 0.011 | 1.205 | Up |
| PHB | 0.034 | 1.235 | Up |
| EXOC7 | 0.010 | 1.249 | Up |
| UNC13C | 0.046 | 1.250 | Up |
| CRELD1 | 0.006 | 1.263 | Up |
| ARHGAP32 | 0.015 | 1.267 | Up |
| HSPA8 | 0.045 | 1.292 | Up |
| MCTP2 | 0.047 | 1.313 | Up |
| CHRNB3 | 0.005 | 1.327 | Up |
| AP1G2 | 0.047 | 1.329 | Up |
| HSP90AA1 | 0.001 | 1.338 | Up |
| ADGRL2 | 0.004 | 1.354 | Up |
| NRP2 | 0.002 | 1.359 | Up |
| ELFN1 | 0.050 | 1.371 | Up |
| SIPA1L2 | 0.007 | 1.378 | Up |
| ACTN1 | 0.040 | 1.413 | Up |
| CHRNA3 | 0.000 | 1.436 | Up |
| DAB1 | 0.010 | 1.495 | Up |
| DNAJB1 | 0.046 | 1.555 | Up |
| NPY2R | 0.031 | 1.924 | Up |
| CRH | 0.014 | 2.344 | Up |

**Supplementary Table 6. Dysregulated mitochondrial genes in propofol exposed organoids**

| Gene ID | Gene name | Biological process | Fold change | Regulation |
|---|---|---|---|---|
| IBA57 | iron-sulfur cluster assembly factor IBA57 | Fe-S Cluster Biosynthesis | 1.585 | Down |
| UCP2 | uncoupling protein 2 | Mitochondrial Carrier | 1.429 | Down |
| DNAJC19 | DnaJ heat shock protein family (Hsp40) member C19 | Import & Sorting | 1.428 | Down |
| ALDOC | aldolase, fructose-bisphosphate C | Glycolysis | 1.410 | Down |
| CMPK2 | cytidine/uridine monophosphate kinase 2 | Replication & Transcription | 1.371 | Down |
| PKM | pyruvate kinase M1/2 | Glycolysis | 1.350 | Down |
| PNPLA8 | patatin like phospholipase domain containing 8 | Fatty Acid Metabolism | 1.330 | Down |
| PDK1 | pyruvate dehydrogenase kinase 1 | Pyruvate Metabolism | 1.324 | Down |
| OGG1 | 8-oxoguanine DNA glycosylase | ROS Defense | 1.321 | Down |
| WARS2 | tryptophanyl tRNA synthetase 2, mitochondrial | Translation | 1.320 | Down |
| BNIP3L | BCL2 interacting protein 3 | Apoptosis | 1.303 | Down |
| LYRM7 | LYR motif containing 7 | Oxidative Phosphorylation | 1.279 | Down |
| FOXO3 | forkhead box O3 | Transcription (nuclear) | 1.271 | Down |
| CKMT1A | creatine kinase, mitochondrial 1A | Amino Acid Metabolism | 1.265 | Down |
| SATB1 | SATB homeobox 1 | UPRmt | 1.251 | Down |
| CKMT1B | creatine kinase, mitochondrial 1B | Amino Acid Metabolism | 1.230 | Down |
| MSRA | methionine sulfoxide reductase A | ROS Defense | 1.224 | Down |
| NBR1 | NBR1 autophagy cargo receptor | Mitophagy | 1.217 | Down |
| TRAP1 | TNF receptor associated protein 1 | Apoptosis | 1.203 | Up |
| MT-ND1 | mitochondrially encoded NADH dehydrogenase 1 | Oxidative Phosphorylation (mt) | 1.206 | Up |
| CCDC58 | coiled-coil domain containing 58 | Translation | 1.211 | Up |
| PHB | prohibitin | Replication & Transcription | 1.221 | Up |
| MRPL12 | mitochondrial ribosomal protein L12 | Translation | 1.252 | Up |
| PMPCA | peptidase, mitochondrial processing alpha subunit | Import & Sorting | 1.260 | Up |
| PPRC1 | PPARG related coactivator 1 | Transcription (nuclear) | 1.264 | Up |
| NDUFAF7 | NADH:ubiquinone oxidoreductase complex assembly factor 7 | Oxidative Phosphorylation | 1.266 | Up |
| ATAD3A | ATPase family AAA domain containing 3A | Mitochondrial Dynamics | 1.278 | Up |
| HSPA1B | heat shock protein family A (Hsp70) member 1B | Protein Stability & Degradation | 1.307 | Up |
| HSPE1 | heat shock protein family E (Hsp10) member 1 | Import & Sorting | 1.320 | Up |
| AURKAIP1 | aurora kinase A interacting protein 1 | Translation | 1.325 | Up |
| PGS1 | phosphatidylglycerophosphate synthase 1 | Cardiolipin Biosynthesis | 1.336 | Up |
| SLC25A19 | solute carrier family 25 member 19 | Mitochondrial Carrier | 1.338 | Up |

| | | | | |
|---|---|---|---|---|
| SLC25A37 | solute carrier family 25 member 37 | Fe-S Cluster Biosynthesis | 1.350 | Up |
| PUS1 | pseudouridine synthase 1 | Translation | 1.360 | Up |
| PTRH2 | peptidyl-tRNA hydrolase 2 | Translation | 1.475 | Up |
| ATAD3B | ATPase family AAA domain containing 3B | Mitochondrial Dynamics | 1.495 | Up |
| EARS2 | glutamyl-tRNA synthetase 2, mitochondrial | Translation | 1.663 | Up |
| ABCF2 | ATP binding cassette subfamily F member 2 | Transmembrane Transport | 1.739 | Up |
| NDUFAF4 | NADH:ubiquinone oxidoreductase complex assembly factor 4 | Oxidative Phosphorylation | 2.034 | Up |

**Supplementary Table 7. Dysregulated inflammatory genes in propofol exposed organoids**

| Gene ID | P value | Fold change | Regulation |
|---|---|---|---|
| HSPA4 | 0.038 | 1.217 | Up |
| F13A1 | 0.042 | 1.528 | Up |
| RGS1 | 0.038 | 1.501 | Up |
| NLRP10 | 0.048 | 1.549 | Up |
| SLC26A6 | 0.030 | 1.289 | Up |
| XBP1 | 0.044 | 1.480 | Up |
| MALAT1 | 0.013 | 1.350 | Up |
| FKBP4 | 0.038 | 1.360 | Up |
| ATP6V0A2 | 0.028 | 1.292 | Up |
| CMKLR1 | 0.043 | 1.714 | Up |
| SCYL1 | 0.049 | 1.205 | Up |
| ADAM15 | 0.006 | 1.269 | Up |
| HSP90AB1 | 0.037 | 1.269 | Up |
| NFAT5 | 0.008 | 1.328 | Up |
| GATA4 | 0.039 | 1.202 | Up |
| NLRP2 | 0.000 | 1.266 | Up |
| PON2 | 0.019 | 1.215 | Up |
| IGF2R | 0.048 | 1.282 | Up |
| NELL1 | 0.011 | 1.281 | Down |
| VCAM1 | 0.041 | 1.333 | Down |
| UCP2 | 0.021 | 1.429 | Down |
| NBR1 | 0.006 | 1.217 | Down |
| C1QTNF9B | 0.020 | 1.400 | Down |

**Supplementary Table 8. Dysregulated lncRNAs in propofol exposed organoids**

| Gene ID | Fold change | P value | Regulation |
|---|---|---|---|
| G037251 | 1.476 | 0.001 | Up |
| G000844 | 1.381 | 0.017 | Up |
| RN7SL417P | 1.288 | 0.030 | Up |
| NADK2-AS1 | 1.287 | 0.033 | Up |
| G003271 | 1.323 | 0.044 | Up |
| XLOC_012303 | 1.372 | 0.021 | Up |
| HCG18 | 1.375 | 0.014 | Up |
| RP4-548D19.3 | 1.267 | 0.027 | Up |
| RP11-308N19.1 | 1.213 | 0.026 | Up |
| RP1-296L11.1 | 1.394 | 0.005 | Up |
| G042193 | 1.292 | 0.047 | Up |
| LOC100499484-C9ORF174 | 1.380 | 0.022 | Up |
| G056910 | 1.485 | 0.020 | Up |
| SLC7A11-AS1 | 1.416 | 0.026 | Up |
| RP11-21M24.3 | 1.359 | 0.007 | Up |
| LINC00473 | 1.888 | 0.030 | Up |
| RP11-674P19.2 | 1.481 | 0.045 | Up |
| uc.172 | 2.152 | 0.039 | Up |
| CTC-462L7.1 | 1.486 | 0.018 | Up |
| RP11-70P17.1 | 1.434 | 0.049 | Up |
| G076532 | 1.285 | 0.045 | Up |
| ITGA9-AS1 | 1.244 | 0.009 | Up |
| PPIEL | 1.285 | 0.049 | Up |
| LOC100505736 | 1.382 | 0.050 | Up |
| HNRNPU-AS1 | 1.427 | 0.037 | Up |
| G075352 | 1.226 | 0.021 | Up |
| G038103 | 1.298 | 0.007 | Up |
| G074385 | 1.306 | 0.027 | Up |
| G038095 | 1.463 | 0.028 | Up |
| CTA-217C2.2 | 1.253 | 0.034 | Up |
| G066551 | 1.666 | 0.020 | Up |
| XLOC_l2_009332 | 1.374 | 0.021 | Up |
| Alpha_250_dna | 1.375 | 0.009 | Up |
| LOC102723439 | 1.213 | 0.024 | Up |
| G038394 | 1.292 | 0.048 | Up |
| CTB-193M12.1 | 1.390 | 0.016 | Up |
| G089550 | 1.506 | 0.039 | Up |
| G053736 | 1.327 | 0.035 | Up |
| G058736 | 1.351 | 0.020 | Up |
| G060633 | 1.489 | 0.020 | Up |
| G029454 | 1.326 | 0.048 | Up |
| G020731 | 1.597 | 0.028 | Up |
| IGBP1-AS2 | 1.774 | 0.001 | Up |
| G011310 | 1.649 | 0.030 | Up |
| LOC171391 | 1.234 | 0.043 | Up |
| ASAP1-IT1 | 1.254 | 0.011 | Up |
| G032927 | 1.700 | 0.037 | Up |
| RP11-498C9.12 | 1.245 | 0.028 | Up |
| AK124233 | 1.224 | 0.034 | Up |
| G061035 | 1.450 | 0.039 | Up |
| XLOC_001223 | 1.410 | 0.012 | Up |
| RP11-3K16.2 | 1.229 | 0.001 | Up |
| G008916 | 1.228 | 0.003 | Up |
| G062538 | 1.295 | 0.048 | Up |
| RP11-187C18.2 | 1.907 | 0.003 | Up |
| RP11-142J21.2 | 1.296 | 0.023 | Up |
| G061982 | 1.442 | 0.048 | Up |
| G076011 | 1.316 | 0.050 | Up |
| BC033330 | 1.284 | 0.050 | Up |
| G053606 | 1.436 | 0.032 | Up |
| RP11-543A18.1 | 1.623 | 0.027 | Up |

| | | | |
|---|---|---|---|
| XLOC_006150 | 1.443 | 0.030 | Up |
| UNC5B-AS1 | 1.219 | 0.009 | Up |
| G006738 | 1.266 | 0.045 | Up |
| G000285 | 1.851 | 0.019 | Up |
| G087508 | 1.415 | 0.018 | Up |
| RP11-461L13.5 | 1.450 | 0.015 | Up |
| G068163 | 1.482 | 0.022 | Up |
| LACTB2-AS1 | 1.687 | 0.010 | Up |
| G049993 | 1.400 | 0.009 | Up |
| RP3-416H24.4 | 1.243 | 0.049 | Up |
| G073552 | 1.308 | 0.045 | Up |
| G058275 | 1.205 | 0.037 | Up |
| LOC101927305 | 1.320 | 0.035 | Up |
| G086585 | 1.222 | 0.041 | Up |
| RP11-445F12.1 | 1.368 | 0.047 | Up |
| RP4-761J14.9 | 1.251 | 0.035 | Up |
| RP4-816N1.7 | 1.449 | 0.031 | Up |
| AQP7P3 | 1.369 | 0.022 | Up |
| RP5-1065P14.2 | 1.472 | 0.024 | Up |
| GSE61474_XLOC_020343 | 1.265 | 0.016 | Up |
| G033744 | 1.703 | 0.002 | Up |
| G009303 | 1.235 | 0.048 | Up |
| G057625 | 1.480 | 0.047 | Up |
| G021277 | 1.439 | 0.020 | Up |
| CHKB-CPT1B | 1.212 | 0.039 | Up |
| G080487 | 1.282 | 0.031 | Up |
| AP000438.2 | 1.418 | 0.033 | Up |
| AC074363.1 | 1.592 | 0.002 | Up |
| G046678 | 1.450 | 0.002 | Up |
| G075558 | 1.258 | 0.021 | Up |
| G031832 | 1.228 | 0.044 | Up |
| PHF2P2 | 1.284 | 0.006 | Up |
| G081487 | 1.269 | 0.031 | Up |
| G055386 | 1.439 | 0.029 | Up |
| AC016682.1 | 1.220 | 0.013 | Up |
| GSE61474_XLOC_051316 | 1.870 | 0.048 | Up |
| RP11-466A19.1 | 1.222 | 0.020 | Up |
| AP001625.6 | 1.221 | 0.009 | Up |
| G063174 | 1.237 | 0.047 | Up |
| RP11-484O2.1 | 1.373 | 0.011 | Up |
| XLOC_008025 | 1.258 | 0.006 | Up |
| G050702 | 1.303 | 0.021 | Up |
| uc.281 | 1.522 | 0.014 | Up |
| PSG10P | 1.220 | 0.043 | Up |
| RP11-587H10.1 | 1.430 | 0.014 | Up |
| G082315 | 1.668 | 0.009 | Up |
| RP11-737O24.3 | 1.444 | 0.000 | Up |
| XLOC_003405 | 1.449 | 0.024 | Up |
| G015131 | 1.333 | 0.034 | Up |
| G009235 | 1.323 | 0.045 | Up |
| G063995 | 1.414 | 0.020 | Up |
| AADACL2-AS1 | 1.458 | 0.024 | Up |
| RP11-669E14.6 | 1.270 | 0.007 | Up |
| NEAT1 | 1.333 | 0.017 | Up |
| XLOC_012579 | 1.349 | 0.009 | Up |
| AL832615 | 1.526 | 0.029 | Up |
| G016487 | 1.330 | 0.043 | Up |
| XLOC_l2_011118 | 1.242 | 0.030 | Up |
| G012913 | 1.296 | 0.032 | Up |
| RN7SKP264 | 2.632 | 0.009 | Up |
| LINC00379 | 1.605 | 0.021 | Up |
| RP11-514F8.2 | 1.340 | 0.022 | Up |
| RP11-131K5.1 | 1.443 | 0.041 | Up |
| LIPE-AS1 | 1.364 | 0.025 | Up |
| PMCHL2 | 2.982 | 0.034 | Up |

| | | | |
|---|---|---|---|
| AX747064 | 1.231 | 0.041 | Up |
| RP11-219C20.3 | 1.252 | 0.007 | Up |
| G068416 | 1.361 | 0.048 | Up |
| G070831 | 1.312 | 0.037 | Up |
| ZFAT-AS1 | 1.277 | 0.045 | Up |
| G040015 | 1.442 | 0.006 | Up |
| G087395 | 1.330 | 0.043 | Up |
| G024041 | 1.305 | 0.005 | Up |
| G087265 | 1.282 | 0.015 | Up |
| G072062 | 1.340 | 0.008 | Up |
| Z69720.2 | 1.454 | 0.006 | Up |
| CTC-261N6.1 | 1.404 | 0.024 | Up |
| G081302 | 1.318 | 0.034 | Up |
| XLOC_l2_001890 | 1.234 | 0.036 | Up |
| BC034456 | 1.701 | 0.012 | Up |
| XLOC_006182 | 1.214 | 0.049 | Up |
| G067669 | 1.465 | 0.035 | Up |
| RP11-18D7.2 | 1.223 | 0.032 | Up |
| G071732 | 1.311 | 0.042 | Up |
| G071640 | 1.367 | 0.043 | Up |
| G082861 | 1.679 | 0.026 | Up |
| G056023 | 1.273 | 0.035 | Up |
| XLOC_l2_003787 | 1.296 | 0.030 | Up |
| AC107218.3 | 1.578 | 0.024 | Up |
| CTB-102L5.8 | 1.347 | 0.025 | Up |
| G047392 | 2.137 | 0.015 | Up |
| RP11-717D12.1 | 1.241 | 0.021 | Up |
| AK093193 | 1.275 | 0.033 | Up |
| LOC101927164 | 1.304 | 0.015 | Up |
| KLK8 | 1.723 | 0.040 | Up |
| G000948 | 1.440 | 0.048 | Up |
| G045594 | 1.513 | 0.031 | Up |
| G014972 | 1.214 | 0.020 | Up |
| CTD-2050E21.1 | 1.409 | 0.023 | Up |
| LOC727896 | 1.538 | 0.027 | Up |
| G036779 | 1.394 | 0.007 | Up |
| AC003682.16 | 1.214 | 0.020 | Up |
| LOC100506858 | 1.324 | 0.013 | Up |
| CRAT37 | 1.314 | 0.047 | Up |
| SNHG7 | 1.266 | 0.023 | Up |
| G046027 | 1.209 | 0.032 | Up |
| RP1-124C6.1 | 1.364 | 0.014 | Up |
| G022034 | 1.457 | 0.027 | Up |
| G062685 | 1.468 | 0.045 | Up |
| G080055 | 1.215 | 0.040 | Up |
| uc.49 | 1.567 | 0.005 | Up |
| G043880 | 1.263 | 0.021 | Up |
| RP11-424I19.2 | 1.266 | 0.014 | Up |
| G006360 | 1.650 | 0.047 | Up |
| G053959 | 1.566 | 0.007 | Up |
| XLOC_013958 | 2.256 | 0.011 | Up |
| RP11-166B2.5 | 1.305 | 0.023 | Up |
| G021365 | 1.519 | 0.018 | Up |
| XLOC_009306 | 1.556 | 0.028 | Up |
| G060178 | 1.253 | 0.045 | Up |
| G063524 | 1.313 | 0.022 | Up |
| XLOC_005405 | 1.693 | 0.003 | Up |
| GSE61474_XLOC_028076 | 1.231 | 0.042 | Up |
| G071944 | 1.232 | 0.029 | Up |
| G011388 | 1.430 | 0.048 | Up |
| XLOC_003837 | 1.474 | 0.012 | Up |
| BX648501 | 1.275 | 0.006 | Up |
| G060010 | 1.692 | 0.037 | Up |
| G073689 | 1.290 | 0.014 | Up |
| G072364 | 1.656 | 0.048 | Up |

| | | | |
|---|---|---|---|
| RP11-567E21.3 | 1.618 | 0.046 | Up |
| G036156 | 1.579 | 0.006 | Up |
| G032508 | 1.265 | 0.012 | Up |
| CYP2D7 | 1.261 | 0.015 | Up |
| CTD-3118D11.2 | 1.794 | 0.022 | Up |
| RP11-374P20.4 | 1.479 | 0.029 | Up |
| LOC401134 | 1.293 | 0.012 | Up |
| G063659 | 1.355 | 0.003 | Up |
| G068469 | 1.465 | 0.048 | Up |
| G020793 | 1.246 | 0.013 | Up |
| G054530 | 1.540 | 0.016 | Up |
| CTD-2023M8.1 | 1.439 | 0.011 | Up |
| G045736 | 1.846 | 0.016 | Up |
| EML2-AS1 | 1.231 | 0.030 | Up |
| LINC01149 | 2.116 | 0.024 | Up |
| G030124 | 1.454 | 0.002 | Up |
| G065302 | 1.580 | 0.032 | Up |
| XLOC_006124 | 1.259 | 0.018 | Up |
| G072890 | 1.418 | 0.018 | Up |
| RP11-94I2.4 | 1.602 | 0.034 | Up |
| NBPF8 | 1.298 | 0.035 | Up |
| LOC101927691 | 1.415 | 0.004 | Up |
| LINC00930 | 1.393 | 0.008 | Up |
| LOC101927285 | 1.376 | 0.035 | Up |
| AK025321 | 1.484 | 0.001 | Up |
| G002461 | 1.204 | 0.043 | Up |
| LOC101926908 | 1.283 | 0.029 | Up |
| G003431 | 1.865 | 0.021 | Up |
| G001795 | 1.518 | 0.003 | Up |
| G016657 | 1.397 | 0.011 | Up |
| AP000344.4 | 1.228 | 0.002 | Up |
| RMRPP3 | 1.204 | 0.018 | Up |
| XLOC_012740 | 1.388 | 0.003 | Up |
| G026517 | 1.339 | 0.026 | Up |
| RP11-96H17.3 | 1.692 | 0.037 | Up |
| ELOVL2-AS1 | 1.364 | 0.030 | Up |
| RP11-650L12.2 | 1.363 | 0.035 | Up |
| G024638 | 1.220 | 0.039 | Up |
| uc.474 | 1.398 | 0.032 | Up |
| GOLGA8B | 1.336 | 0.003 | Up |
| G035803 | 1.503 | 0.036 | Up |
| XLOC_006485 | 1.344 | 0.000 | Up |
| TMLHE-AS1 | 1.835 | 0.019 | Up |
| RP1-65P5.3 | 1.362 | 0.038 | Up |
| G060484 | 1.613 | 0.023 | Up |
| LOC100129917 | 1.269 | 0.016 | Up |
| RP11-45M22.5 | 1.243 | 0.041 | Up |
| RP11-30O15.1 | 1.288 | 0.023 | Up |
| RP11-768G7.2 | 1.461 | 0.028 | Up |
| G058409 | 2.070 | 0.013 | Up |
| G009329 | 1.626 | 0.011 | Up |
| G067215 | 1.343 | 0.048 | Up |
| LOC642846 | 1.373 | 0.043 | Up |
| RP11-10J18.3 | 1.272 | 0.046 | Up |
| G020470 | 1.268 | 0.048 | Up |
| XX-C283C717.1 | 1.265 | 0.034 | Up |
| LINC00685 | 1.308 | 0.031 | Up |
| G015181 | 1.395 | 0.007 | Up |
| G037029 | 1.575 | 0.021 | Up |
| G082713 | 1.703 | 0.015 | Up |
| G010088 | 1.290 | 0.040 | Up |
| XLOC_008270 | 1.240 | 0.030 | Up |
| G060726 | 1.321 | 0.016 | Up |
| G046492 | 1.260 | 0.013 | Up |
| G002410 | 1.412 | 0.045 | Up |

| | | | |
|---|---|---|---|
| AC026188.1 | 1.667 | 0.003 | Up |
| G052905 | 1.414 | 0.007 | Up |
| RP11-551L14.6 | 1.321 | 0.008 | Up |
| uc.50 | 1.273 | 0.012 | Up |
| LOC100131564 | 1.277 | 0.028 | Up |
| G080747 | 1.273 | 0.020 | Up |
| G013234 | 1.343 | 0.035 | Up |
| RP4-760C5.3 | 1.304 | 0.041 | Up |
| CTD-3193O13.1 | 1.289 | 0.030 | Up |
| G003928 | 1.638 | 0.019 | Up |
| G072257 | 1.289 | 0.037 | Up |
| AF086132 | 1.201 | 0.035 | Up |
| CTC-250I14.3 | 1.243 | 0.007 | Up |
| G023385 | 1.701 | 0.039 | Up |
| G051586 | 2.158 | 0.000 | Up |
| RP11-126O1.4 | 1.287 | 0.044 | Up |
| RP11-651L5.2 | 1.239 | 0.009 | Up |
| GSE61474_XLOC_042934 | 1.258 | 0.043 | Up |
| G067882 | 1.255 | 0.026 | Up |
| AK025564 | 1.265 | 0.021 | Up |
| XLOC_007237 | 1.295 | 0.036 | Up |
| XLOC_000048 | 1.222 | 0.024 | Up |
| LOC101926941 | 1.492 | 0.032 | Up |
| G053358 | 1.310 | 0.042 | Up |
| G068224 | 1.487 | 0.035 | Up |
| BCRP3 | 1.264 | 0.014 | Up |
| HM13-IT1 | 1.302 | 0.020 | Up |
| LOC105373609 | 1.653 | 0.012 | Up |
| G009199 | 2.721 | 0.042 | Up |
| G033096 | 1.454 | 0.006 | Up |
| G089283 | 1.433 | 0.009 | Up |
| XLOC_007868 | 1.394 | 0.026 | Up |
| AC025442.3 | 1.782 | 0.015 | Up |
| G013465 | 2.113 | 0.042 | Up |
| G071259 | 1.540 | 0.020 | Up |
| AC004054.1 | 1.449 | 0.048 | Up |
| RP5-1112D6.7 | 1.329 | 0.036 | Up |
| MINOS1P1 | 1.320 | 0.044 | Up |
| MIMT1 | 1.344 | 0.042 | Up |
| G079644 | 1.480 | 0.034 | Up |
| G068890 | 1.371 | 0.039 | Up |
| G010941 | 1.314 | 0.001 | Up |
| SPTY2D1-AS1 | 1.233 | 0.035 | Up |
| G031271 | 1.619 | 0.020 | Up |
| G029241 | 1.600 | 0.010 | Up |
| AK022848 | 1.410 | 0.031 | Up |
| LOC100506834 | 1.469 | 0.021 | Up |
| G051433 | 1.294 | 0.014 | Up |
| NBPF25P | 1.251 | 0.034 | Up |
| DLG3-AS1 | 1.382 | 0.000 | Up |
| EF565122 | 2.238 | 0.033 | Up |
| G088912 | 1.397 | 0.009 | Up |
| CTD-2015A6.1 | 1.227 | 0.020 | Up |
| G084746 | 1.407 | 0.009 | Up |
| G013179 | 1.470 | 0.039 | Up |
| GSE61474_XLOC_028262 | 1.320 | 0.013 | Up |
| LOC101929427 | 1.410 | 0.040 | Up |
| RN7SL176P | 1.341 | 0.039 | Up |
| G074298 | 1.412 | 0.013 | Up |
| LOC101927460 | 1.384 | 0.036 | Up |
| RP11-354H21.2 | 2.276 | 0.039 | Up |
| LINC01602 | 1.623 | 0.017 | Up |
| G010533 | 4.968 | 0.003 | Up |
| G010224 | 1.227 | 0.019 | Up |
| G030745 | 1.403 | 0.007 | Up |

| | | | |
|---|---|---|---|
| G022519 | 1.381 | 0.042 | Up |
| AK024936 | 1.550 | 0.022 | Up |
| XLOC_003048 | 1.274 | 0.010 | Up |
| XLOC_006353 | 1.334 | 0.047 | Up |
| G061888 | 1.345 | 0.022 | Up |
| XLOC_009429 | 1.299 | 0.033 | Up |
| RP11-461K13.1 | 1.778 | 0.050 | Up |
| G041587 | 1.409 | 0.029 | Up |
| G059595 | 1.385 | 0.005 | Up |
| IFT74-AS1 | 2.010 | 0.038 | Down |
| FAM74A4 | 1.606 | 0.018 | Down |
| RP11-644F5.16 | 1.253 | 0.048 | Down |
| RP11-602N24.3 | 1.338 | 0.020 | Down |
| AL110181 | 1.438 | 0.009 | Down |
| TIPARP-AS1 | 2.146 | 0.026 | Down |
| RP11-255E6.6 | 1.450 | 0.002 | Down |
| XLOC_010525 | 1.468 | 0.026 | Down |
| RP11-744I24.3 | 4.765 | 0.020 | Down |
| XLOC_006779 | 1.421 | 0.037 | Down |
| G039106 | 4.103 | 0.044 | Down |
| GSE61474_XLOC_054139 | 11.775 | 0.004 | Down |
| XLOC_003724 | 1.757 | 0.049 | Down |
| RP11-363E6.3 | 1.453 | 0.037 | Down |
| AK055712 | 1.469 | 0.017 | Down |
| RP11-230C9.2 | 1.384 | 0.032 | Down |
| RP11-49K24.8 | 1.233 | 0.045 | Down |
| LOC100130417 | 1.330 | 0.016 | Down |
| RPL23AP7 | 1.251 | 0.010 | Down |
| G090511 | 1.830 | 0.019 | Down |
| RP11-44N22.3 | 1.314 | 0.003 | Down |
| CTD-2302E22.4 | 1.632 | 0.009 | Down |
| OR2L1P | 1.669 | 0.036 | Down |
| PSMG3-AS1 | 1.447 | 0.045 | Down |
| GS1-433O24.1 | 1.762 | 0.005 | Down |
| GSE61474_XLOC_042356 | 1.449 | 0.042 | Down |
| RP11-196G18.23 | 1.880 | 0.008 | Down |
| LINC00652 | 1.441 | 0.021 | Down |
| RP11-525A16.4 | 1.674 | 0.017 | Down |
| G017718 | 1.257 | 0.020 | Down |
| RP11-689C9.1 | 1.340 | 0.035 | Down |
| RP11-303E16.7 | 1.908 | 0.028 | Down |
| XLOC_008005 | 1.709 | 0.037 | Down |
| G020528 | 2.793 | 0.029 | Down |
| CTD-2023N9.3 | 2.160 | 0.042 | Down |
| RP11-350N15.5 | 1.411 | 0.046 | Down |
| RP11-762I7.4 | 1.218 | 0.033 | Down |
| RP4-549L20.3 | 1.247 | 0.039 | Down |
| RP11-686O6.2 | 1.472 | 0.029 | Down |
| G061591 | 1.325 | 0.015 | Down |
| RP11-356J5.12 | 1.815 | 0.028 | Down |
| RP11-624D11.2 | 1.459 | 0.049 | Down |
| FAM230C | 1.567 | 0.002 | Down |
| USP27X-AS1 | 1.254 | 0.037 | Down |
| G018241 | 1.471 | 0.041 | Down |
| G070136 | 1.243 | 0.048 | Down |
| EPHA5-AS1 | 3.768 | 0.019 | Down |
| RP1-40E16.12 | 1.627 | 0.011 | Down |
| BRWD1-IT2 | 1.699 | 0.014 | Down |
| RP11-355B11.2 | 1.253 | 0.002 | Down |
| XLOC_l2_002067 | 1.568 | 0.028 | Down |
| RP11-10J21.4 | 1.340 | 0.012 | Down |
| XLOC_002675 | 1.441 | 0.006 | Down |
| G021652 | 1.422 | 0.007 | Down |
| GBAT2 | 1.641 | 0.010 | Down |
| RP11-490O6.2 | 1.917 | 0.018 | Down |

| | | | |
|---|---|---|---|
| G047503 | 1.248 | 0.022 | Down |
| AP000769.7 | 1.561 | 0.046 | Down |
| AL137474 | 2.358 | 0.049 | Down |
| ZFX-AS1 | 1.298 | 0.048 | Down |
| BC047090 | 1.357 | 0.038 | Down |
| RP11-1017G21.6 | 1.666 | 0.016 | Down |
| PLBD1-AS1 | 1.531 | 0.045 | Down |
| G002862 | 1.665 | 0.037 | Down |
| G033752 | 1.226 | 0.034 | Down |
| LOC101060498 | 1.369 | 0.045 | Down |
| G075198 | 1.270 | 0.034 | Down |
| DLG5-AS1 | 1.313 | 0.017 | Down |
| XLOC_011336 | 1.430 | 0.042 | Down |
| G049752 | 1.800 | 0.030 | Down |
| SH3GL1P2 | 1.519 | 0.026 | Down |
| AC138655.6 | 1.456 | 0.018 | Down |
| LOC729083 | 2.845 | 0.006 | Down |
| TM4SF1-AS1 | 1.245 | 0.024 | Down |
| TPTE2P3 | 1.794 | 0.022 | Down |
| CTD-2033A16.3 | 1.281 | 0.010 | Down |
| RP11-599B13.7 | 1.406 | 0.039 | Down |
| LINC00662 | 1.876 | 0.011 | Down |
| RP11-297J22.1 | 1.597 | 0.040 | Down |
| RP11-179B2.2 | 1.707 | 0.010 | Down |
| EP300-AS1 | 1.372 | 0.028 | Down |
| G064655 | 2.134 | 0.020 | Down |
| G017886 | 1.206 | 0.031 | Down |
| RP11-182N22.8 | 1.998 | 0.010 | Down |
| CTD-2588C8.8 | 1.920 | 0.021 | Down |
| G057990 | 1.730 | 0.019 | Down |
| G047225 | 2.136 | 0.036 | Down |
| RP1-309F20.3 | 1.332 | 0.040 | Down |
| RP11-643M14.1 | 1.662 | 0.046 | Down |
| RP11-344E13.4 | 1.213 | 0.042 | Down |
| SCAMP1-AS1 | 1.418 | 0.011 | Down |
| G017670 | 1.466 | 0.045 | Down |
| RP11-21M24.2 | 1.763 | 0.025 | Down |
| RP11-252A24.3 | 1.563 | 0.015 | Down |
| XLOC_013660 | 1.815 | 0.009 | Down |
| AC073321.4 | 1.284 | 0.004 | Down |
| CTC-336P14.1 | 1.581 | 0.047 | Down |
| XLOC_008875 | 1.341 | 0.049 | Down |
| G077330 | 4.686 | 0.013 | Down |
| RP11-46J23.1 | 1.218 | 0.007 | Down |
| RP11-715J22.6 | 1.796 | 0.003 | Down |
| RP11-277L2.3 | 1.339 | 0.037 | Down |
| RP11-191L17.1 | 1.398 | 0.009 | Down |
| G043416 | 1.384 | 0.044 | Down |
| G037472 | 1.467 | 0.048 | Down |
| CCDC144NL-AS1 | 1.913 | 0.023 | Down |
| WWC2-AS2 | 1.505 | 0.015 | Down |
| AC099754.1 | 2.464 | 0.010 | Down |
| RP11-165A20.3 | 1.738 | 0.038 | Down |
| G018162 | 1.367 | 0.013 | Down |
| G079009 | 1.274 | 0.015 | Down |
| RP11-396C23.4 | 1.676 | 0.005 | Down |
| G000481 | 2.507 | 0.035 | Down |
| RP11-285F16.1 | 1.704 | 0.006 | Down |
| G050681 | 1.621 | 0.012 | Down |
| G048723 | 1.346 | 0.039 | Down |
| CTC-232P5.3 | 1.410 | 0.002 | Down |
| G047031 | 1.605 | 0.044 | Down |
| RP5-991G20.4 | 1.267 | 0.025 | Down |
| RP5-1092A3.4 | 1.356 | 0.006 | Down |
| BC112270 | 1.299 | 0.017 | Down |

| | | | |
|---|---|---|---|
| G014329 | 1.583 | 0.025 | Down |
| RP11-351D16.3 | 1.550 | 0.020 | Down |
| MGC2889 | 1.677 | 0.011 | Down |
| AK123758 | 1.479 | 0.014 | Down |
| RP4-683L5.1 | 1.630 | 0.030 | Down |
| G086828 | 1.389 | 0.030 | Down |
| RP11-353K11.1 | 1.478 | 0.017 | Down |
| G082153 | 1.212 | 0.048 | Down |
| HRAT5 | 1.737 | 0.017 | Down |
| RP11-209D14.2 | 1.643 | 0.049 | Down |
| G050931 | 1.398 | 0.043 | Down |
| CTA-212D2.2 | 2.008 | 0.039 | Down |
| FAM138A | 1.309 | 0.043 | Down |
| AC002467.7 | 1.533 | 0.029 | Down |
| G015380 | 1.637 | 0.022 | Down |
| CTD-2005H7.1 | 1.267 | 0.008 | Down |
| TAF1A-AS1 | 1.224 | 0.030 | Down |
| G020605 | 1.249 | 0.032 | Down |
| uc.85 | 1.255 | 0.011 | Down |
| XLOC_001189 | 1.593 | 0.012 | Down |
| XLOC_003595 | 2.141 | 0.036 | Down |
| RP11-671J11.6 | 2.140 | 0.036 | Down |
| LINC00883 | 1.615 | 0.033 | Down |
| G066319 | 1.330 | 0.004 | Down |
| XLOC_006554 | 1.270 | 0.004 | Down |
| G051725 | 1.420 | 0.042 | Down |
| FAM66A | 1.270 | 0.045 | Down |
| G042247 | 1.603 | 0.010 | Down |
| RP11-475I24.3 | 1.221 | 0.040 | Down |
| AC006273.4 | 1.297 | 0.017 | Down |
| CTB-113I20.2 | 1.377 | 0.027 | Down |
| RP11-312B8.1 | 1.559 | 0.018 | Down |
| RP5-968P14.2 | 1.498 | 0.023 | Down |
| GAS5-AS1 | 1.406 | 0.045 | Down |
| AC144652.1 | 1.791 | 0.046 | Down |
| RP11-204E9.1 | 2.091 | 0.014 | Down |
| G085410 | 1.773 | 0.001 | Down |
| uc.446 | 1.818 | 0.046 | Down |
| DHRS4-AS1 | 1.450 | 0.005 | Down |
| RP3-410B11.1 | 1.414 | 0.023 | Down |
| G079888 | 1.220 | 0.042 | Down |
| RP11-326A19.3 | 1.244 | 0.025 | Down |
| G089005 | 2.621 | 0.005 | Down |
| G048385 | 1.340 | 0.012 | Down |
| G019879 | 1.661 | 0.033 | Down |
| OLMALINC | 1.548 | 0.003 | Down |
| G000619 | 2.743 | 0.038 | Down |
| AC093391.2 | 1.303 | 0.035 | Down |
| LOC101929741 | 1.520 | 0.014 | Down |
| G079654 | 1.849 | 0.005 | Down |
| TEX26-AS1 | 2.864 | 0.037 | Down |
| RP11-496H1.2 | 1.644 | 0.025 | Down |
| RP11-510M2.2 | 2.279 | 0.015 | Down |
| G010185 | 1.282 | 0.031 | Down |
| XLOC_006928 | 1.593 | 0.049 | Down |
| G009018 | 2.531 | 0.007 | Down |
| SUCLA2-AS1 | 2.732 | 0.020 | Down |
| G029844 | 3.768 | 0.030 | Down |
| RP11-2E11.10 | 1.900 | 0.032 | Down |
| FRMD6-AS1 | 1.978 | 0.033 | Down |
| RP11-1055B8.8 | 1.545 | 0.045 | Down |
| G052526 | 1.361 | 0.007 | Down |
| LINC00408 | 1.662 | 0.017 | Down |
| RP11-128P10.1 | 1.326 | 0.005 | Down |
| G011337 | 1.353 | 0.047 | Down |

| | | | |
|---|---|---|---|
| RP11-382D12.2 | 1.330 | 0.014 | Down |
| RP11-174G6.5 | 1.487 | 0.030 | Down |
| TCR-alpha | 1.230 | 0.047 | Down |
| AL355711 | 1.413 | 0.011 | Down |
| CTD-2256P15.2 | 1.672 | 0.011 | Down |
| TUB-AS1 | 1.519 | 0.010 | Down |
| G053260 | 1.329 | 0.034 | Down |
| BC070391 | 1.675 | 0.025 | Down |
| SIX3-AS1 | 2.169 | 0.040 | Down |
| G041948 | 1.295 | 0.037 | Down |
| G034470 | 1.503 | 0.022 | Down |
| MAPT-IT1 | 2.094 | 0.026 | Down |
| LINC01132 | 3.539 | 0.022 | Down |
| G041225 | 1.405 | 0.025 | Down |
| G073576 | 1.801 | 0.031 | Down |
| G045191 | 1.644 | 0.035 | Down |
| XLOC_013222 | 1.346 | 0.033 | Down |
| LINC-PINT | 1.622 | 0.012 | Down |
| RP11-299H21.1 | 2.041 | 0.016 | Down |
| ITFG1-AS1 | 1.441 | 0.010 | Down |
| G081251 | 2.388 | 0.007 | Down |
| G074033 | 3.239 | 0.006 | Down |
| CTD-2517M22.16 | 2.092 | 0.020 | Down |
| TPI1P3 | 1.342 | 0.044 | Down |
| RP11-424N24.2 | 1.488 | 0.004 | Down |
| CCNT2-AS1 | 1.370 | 0.003 | Down |
| XLOC_011052 | 1.446 | 0.003 | Down |
| G061579 | 1.661 | 0.015 | Down |
| RP11-502I4.3 | 1.564 | 0.001 | Down |
| G023041 | 1.458 | 0.008 | Down |
| RP1-202O8.3 | 1.741 | 0.016 | Down |
| G087757 | 2.008 | 0.042 | Down |
| RP13-131K19.2 | 1.385 | 0.008 | Down |
| G082866 | 1.797 | 0.037 | Down |
| XLOC_000308 | 1.327 | 0.027 | Down |
| G013286 | 1.401 | 0.040 | Down |
| CTD-2555K7.2 | 1.380 | 0.032 | Down |
| FOXN3-AS1 | 1.411 | 0.040 | Down |
| RP11-290L1.2 | 1.635 | 0.002 | Down |
| TNRC18P1 | 1.563 | 0.017 | Down |
| GSE61474_XLOC_038226 | 1.665 | 0.046 | Down |
| IDH1-AS1 | 1.389 | 0.002 | Down |
| G065067 | 1.544 | 0.045 | Down |
| INAFM1 | 2.793 | 0.032 | Down |
| G019632 | 1.518 | 0.024 | Down |
| NIPBL-AS1 | 1.263 | 0.041 | Down |
| RP11-145A3.2 | 1.289 | 0.035 | Down |
| KB-1592A4.14 | 1.388 | 0.036 | Down |
| LOC100506083 | 1.346 | 0.035 | Down |
| G038972 | 1.493 | 0.028 | Down |
| LOC100379224 | 1.325 | 0.029 | Down |
| LOC101928322 | 2.324 | 0.044 | Down |
| RP11-505K9.1 | 1.669 | 0.024 | Down |
| RP11-792A8.4 | 1.370 | 0.008 | Down |
| G063668 | 1.408 | 0.017 | Down |
| RP11-138E2.1 | 2.139 | 0.014 | Down |
| RP11-775D22.3 | 1.448 | 0.016 | Down |
| G003798 | 2.163 | 0.021 | Down |
| G053697 | 1.428 | 0.014 | Down |
| G006537 | 1.297 | 0.004 | Down |
| G066606 | 1.439 | 0.023 | Down |
| LOC730102 | 1.202 | 0.036 | Down |
| AK026244 | 1.368 | 0.032 | Down |
| G059716 | 1.364 | 0.016 | Down |
| RP11-2E11.5 | 1.452 | 0.010 | Down |

| | | | |
|---|---|---|---|
| LOC148709 | 1.652 | 0.023 | Down |
| G044581 | 1.474 | 0.009 | Down |
| RP11-567G11.1 | 1.928 | 0.027 | Down |
| G019668 | 1.257 | 0.049 | Down |
| RP5-884G6.2 | 2.083 | 0.019 | Down |
| PVRL3-AS1 | 1.516 | 0.002 | Down |
| AC114803.3 | 1.958 | 0.016 | Down |
| AC099850.1 | 1.406 | 0.040 | Down |
| G010782 | 1.344 | 0.009 | Down |
| BC042589 | 2.451 | 0.007 | Down |
| LINC00634 | 1.747 | 0.015 | Down |
| CTD-2383M3.1 | 2.290 | 0.037 | Down |
| AC092667.2 | 1.974 | 0.019 | Down |
| GYG2P1 | 1.772 | 0.023 | Down |
| RP11-376P6.3 | 1.491 | 0.048 | Down |
| FGF13-AS1 | 1.564 | 0.031 | Down |
| RP1-97J1.2 | 1.419 | 0.032 | Down |
| RP11-383C5.5 | 1.751 | 0.016 | Down |
| LOC644656 | 1.376 | 0.038 | Down |
| G001739 | 1.626 | 0.045 | Down |
| AC005220.3 | 2.248 | 0.005 | Down |
| RP11-22N19.2 | 1.674 | 0.023 | Down |
| RP11-51J9.5 | 1.327 | 0.042 | Down |
| LOC102724661 | 1.214 | 0.027 | Down |
| LINC01023 | 1.507 | 0.029 | Down |
| uc.95 | 2.574 | 0.029 | Down |
| LOC101928402 | 1.472 | 0.042 | Down |
| G090038 | 2.523 | 0.011 | Down |
| GFOD1 | 1.711 | 0.045 | Down |
| LOC101928417 | 1.401 | 0.048 | Down |
| GPR50-AS1 | 1.498 | 0.050 | Down |
| FAM138E | 1.580 | 0.029 | Down |
| G078120 | 1.358 | 0.012 | Down |
| RP4-616B8.4 | 1.487 | 0.036 | Down |
| RP4-555D20.3 | 2.806 | 0.004 | Down |
| RP11-507K2.3 | 1.530 | 0.042 | Down |
| DQ594967 | 1.325 | 0.046 | Down |
| TOB1-AS1 | 1.454 | 0.005 | Down |
| AP000255.6 | 1.744 | 0.010 | Down |
| AK000953 | 1.263 | 0.014 | Down |
| RP4-798A10.7 | 1.512 | 0.045 | Down |
| CTD-2377O17.1 | 2.237 | 0.015 | Down |
| LINC00630 | 1.395 | 0.047 | Down |
| G049105 | 2.007 | 0.013 | Down |
| RP11-254F7.2 | 1.393 | 0.007 | Down |
| G031140 | 1.314 | 0.044 | Down |
| uc.76 | 1.627 | 0.036 | Down |
| PRR36 | 1.574 | 0.032 | Down |
| LL22NC03-86G7.1 | 1.375 | 0.044 | Down |
| RP11-332H14.1 | 1.774 | 0.024 | Down |
| ULK4P1 | 1.289 | 0.021 | Down |
| G066437 | 1.332 | 0.001 | Down |
| G057325 | 1.207 | 0.029 | Down |
| ENO1-AS1 | 2.145 | 0.000 | Down |
| RP1-261G23.7 | 1.240 | 0.042 | Down |
| G069105 | 1.592 | 0.011 | Down |
| G023087 | 1.840 | 0.022 | Down |
| RP11-1275H24.1 | 1.311 | 0.040 | Down |
| RP1-223E5.4 | 1.546 | 0.012 | Down |
| LINC01152 | 1.228 | 0.047 | Down |
| AACSP1 | 1.571 | 0.030 | Down |
| G037470 | 1.396 | 0.039 | Down |
| RP11-680F8.1 | 1.219 | 0.031 | Down |
| DGCR5 | 1.841 | 0.001 | Down |
| RP11-413M3.4 | 1.262 | 0.028 | Down |

| | | | |
|---|---|---|---|
| FAM66D | 1.316 | 0.039 | Down |
| LINC00463 | 1.617 | 0.042 | Down |
| G010838 | 1.555 | 0.037 | Down |
| G027543 | 1.330 | 0.005 | Down |
| RP11-573G6.8 | 2.080 | 0.024 | Down |
| XLOC_007467 | 1.223 | 0.020 | Down |
| RP11-107N15.1 | 1.571 | 0.047 | Down |
| LL09NC01-139C3.1 | 2.099 | 0.002 | Down |
| G061722 | 1.202 | 0.002 | Down |
| G046707 | 3.146 | 0.033 | Down |
| G042996 | 1.248 | 0.046 | Down |
| LOC100288846 | 1.493 | 0.024 | Down |
| G028263 | 1.256 | 0.029 | Down |
| RP11-798M19.6 | 1.543 | 0.022 | Down |
| G068576 | 1.501 | 0.009 | Down |
| RP11-872D17.4 | 1.329 | 0.029 | Down |
| XLOC_004632 | 1.294 | 0.023 | Down |
| G036568 | 1.449 | 0.035 | Down |
| CXADRP3 | 1.217 | 0.003 | Down |
| UBAC2-AS1 | 1.390 | 0.007 | Down |
| RP11-53B2.6 | 2.831 | 0.007 | Down |
| CTD-2527I21.7 | 1.231 | 0.003 | Down |
| XLOC_005289 | 1.430 | 0.024 | Down |
| DDX39B-AS1 | 1.398 | 0.029 | Down |
| RP11-177G23.2 | 1.704 | 0.029 | Down |
| G000672 | 2.096 | 0.012 | Down |
| CTD-2382E5.2 | 1.549 | 0.000 | Down |
| EZR-AS1 | 1.328 | 0.002 | Down |
| RP1-86C11.7 | 3.053 | 0.004 | Down |
| GS1-600G8.3 | 1.957 | 0.029 | Down |
| G081642 | 1.398 | 0.005 | Down |
| G058460 | 1.369 | 0.043 | Down |
| CTD-2653B5.1 | 1.330 | 0.035 | Down |
| G047565 | 1.265 | 0.005 | Down |
| RP11-793H13.11 | 1.297 | 0.037 | Down |
| G064611 | 1.368 | 0.034 | Down |
| FLJ20021 | 1.636 | 0.008 | Down |
| uc.247 | 1.554 | 0.045 | Down |
| RP11-109D9.4 | 1.761 | 0.001 | Down |
| RP11-110G21.1 | 2.092 | 0.003 | Down |
| CTC-444N24.13 | 1.390 | 0.016 | Down |
| RP11-339B21.10 | 1.407 | 0.021 | Down |
| LINC00449 | 1.415 | 0.009 | Down |
| RP11-61L19.2 | 1.635 | 0.016 | Down |
| LOC101929066 | 1.301 | 0.032 | Down |
| G054532 | 2.237 | 0.018 | Down |
| RP11-85A1.3 | 1.794 | 0.050 | Down |
| G061596 | 2.589 | 0.010 | Down |
| AK125078 | 1.220 | 0.001 | Down |
| BX248253 | 1.634 | 0.001 | Down |
| uc.349 | 2.582 | 0.002 | Down |
| RP11-344P13.6 | 2.216 | 0.023 | Down |
| CTC-359M8.1 | 1.418 | 0.025 | Down |
| uc.69 | 1.346 | 0.016 | Down |
| LOC101927822 | 1.492 | 0.019 | Down |
| UBE2V1 | 1.290 | 0.018 | Down |
| G070405 | 1.811 | 0.002 | Down |
| RP11-565F19.2 | 1.418 | 0.020 | Down |
| G054079 | 1.539 | 0.001 | Down |
| ANKRD20A12P | 1.434 | 0.008 | Down |
| G003603 | 1.364 | 0.011 | Down |
| RP11-53I6.3 | 1.321 | 0.003 | Down |
| RP11-483I13.5 | 1.300 | 0.013 | Down |
| RP11-314A20.5 | 1.352 | 0.026 | Down |
| AK055974 | 2.113 | 0.011 | Down |

| | | | |
|---|---|---|---|
| G060549 | 1.318 | 0.028 | Down |
| RP11-474O21.5 | 2.237 | 0.020 | Down |
| G046528 | 1.519 | 0.028 | Down |
| CRYBB2P1 | 1.228 | 0.032 | Down |
| RP11-196G18.24 | 1.732 | 0.017 | Down |
| LOC101927811 | 1.364 | 0.006 | Down |
| CTD-2623N2.11 | 1.418 | 0.045 | Down |
| RP11-254F7.1 | 1.234 | 0.044 | Down |
| G064225 | 1.571 | 0.027 | Down |
| uc.412 | 1.525 | 0.021 | Down |
| CTC-350I8.1 | 1.410 | 0.045 | Down |
| G023340 | 1.552 | 0.024 | Down |
| G000727 | 1.390 | 0.027 | Down |
| H1FX-AS1 | 1.298 | 0.005 | Down |
| GSE61474_XLOC_052497 | 1.587 | 0.027 | Down |
| KB-318B8.7 | 1.601 | 0.023 | Down |
| G063514 | 1.217 | 0.026 | Down |
| RP6-109B7.4 | 1.324 | 0.030 | Down |
| G070622 | 1.300 | 0.030 | Down |
| RP11-30L3.2 | 1.810 | 0.038 | Down |
| G006089 | 1.670 | 0.039 | Down |
| G014106 | 1.704 | 0.001 | Down |
| G020860 | 1.317 | 0.022 | Down |
| RP1-232L24.3 | 1.972 | 0.043 | Down |
| RP11-362L22.1 | 1.463 | 0.040 | Down |
| RP11-467P9.1 | 1.721 | 0.009 | Down |
| XXbac-B476C20.9 | 1.701 | 0.013 | Down |
| G002695 | 1.575 | 0.007 | Down |
| G021588 | 2.147 | 0.002 | Down |
| G006578 | 1.688 | 0.038 | Down |
| RP11-371A19.2 | 1.816 | 0.039 | Down |
| RP11-95P2.3 | 2.466 | 0.013 | Down |
| RP11-118K6.3 | 1.462 | 0.006 | Down |
| G061238 | 1.404 | 0.015 | Down |
| RP11-895M11.3 | 1.809 | 0.019 | Down |
| LOC101928489 | 1.967 | 0.007 | Down |
| RP11-355F22.1 | 1.398 | 0.010 | Down |
| JARID2-AS1 | 1.269 | 0.046 | Down |
| G036604 | 1.357 | 0.041 | Down |
| G075319 | 1.249 | 0.024 | Down |
| XLOC_010219 | 1.541 | 0.028 | Down |
| GSE61474_XLOC_067608 | 1.663 | 0.022 | Down |
| RP11-330O11.3 | 2.547 | 0.039 | Down |
| LOC283140 | 2.176 | 0.020 | Down |
| DNAJC9-AS1 | 1.639 | 0.001 | Down |
| G065507 | 1.221 | 0.044 | Down |
| RP11-820I16.4 | 1.690 | 0.050 | Down |
| G021721 | 1.298 | 0.036 | Down |
| G026062 | 1.748 | 0.036 | Down |
| RP11-79P5.9 | 1.341 | 0.022 | Down |
| G057611 | 1.819 | 0.041 | Down |
| CTB-181F24.1 | 1.288 | 0.014 | Down |
| RP11-166B2.3 | 1.298 | 0.012 | Down |
| RP11-168O10.6 | 1.297 | 0.043 | Down |
| LOC100270746 | 1.305 | 0.006 | Down |
| G075703 | 2.144 | 0.002 | Down |
| LINC00858 | 2.557 | 0.016 | Down |
| BC047644 | 1.745 | 0.015 | Down |
| G022520 | 1.727 | 0.040 | Down |
| RN7SL832P | 1.291 | 0.030 | Down |
| RP11-820L6.1 | 1.864 | 0.000 | Down |
| LINC01604 | 1.306 | 0.025 | Down |
| LOC102723385 | 2.323 | 0.002 | Down |
| RP11-803B1.8 | 1.365 | 0.018 | Down |
| G005619 | 1.277 | 0.007 | Down |

| | | | |
|---|---|---|---|
| G063721 | 1.461 | 0.017 | Down |
| XLOC_004894 | 1.578 | 0.017 | Down |
| G015529 | 1.735 | 0.008 | Down |
| CTD-2517O10.6 | 1.583 | 0.029 | Down |
| G001497 | 1.393 | 0.013 | Down |
| GLIDR | 1.322 | 0.024 | Down |
| RP11-439C15.4 | 1.861 | 0.032 | Down |
| G007298 | 1.215 | 0.011 | Down |
| LOC101927358 | 1.259 | 0.034 | Down |
| BC040628 | 1.397 | 0.045 | Down |
| RP11-196G18.22 | 1.731 | 0.046 | Down |
| CTC-248O19.1 | 1.456 | 0.016 | Down |
| RP11-379F4.9 | 1.413 | 0.025 | Down |
| G014762 | 1.584 | 0.027 | Down |
| G085202 | 1.551 | 0.026 | Down |
| LINC00670 | 2.057 | 0.037 | Down |

**Supplementary Table 9. Dysregulated lncRNAs-synaptic mRNAs co-expression network in propofol exposed organoids**

| LncRNAs (source genes) | Synapse mRNAs (target gene) | Correlation coefficient | P value | Positive/Negative Correlation Coefficient | Regulation |
|---|---|---|---|---|---|
| RN7SL417P | EXOC7 | -0.954 | 0.000 | -1.000 | up |
| NADK2-AS1 | ACTN1 | -0.912 | 0.002 | -1.000 | up |
| XLOC_012303 | ATP2B4 | -0.927 | 0.001 | -1.000 | up |
| XLOC_012303 | SEPSECS | -0.914 | 0.001 | -1.000 | up |
| XLOC_012303 | CABP1 | -0.925 | 0.001 | -1.000 | up |
| RP11-308N19.1 | CHRNB3 | -0.912 | 0.002 | -1.000 | up |
| RP1-296L11.1 | CRELD1 | -0.966 | 0.000 | -1.000 | up |
| RP1-296L11.1 | SIPA1L2 | -0.937 | 0.001 | -1.000 | up |
| RP1-296L11.1 | FILIP1 | -0.905 | 0.002 | -1.000 | up |
| LOC100499484-C9ORF1 | CRELD1 | -0.900 | 0.002 | -1.000 | up |
| LOC100499484-C9ORF1 | HSP90AA1 | -0.922 | 0.001 | -1.000 | up |
| LOC100499484-C9ORF1 | DPYSL5 | -0.902 | 0.002 | -1.000 | up |
| LOC100499484-C9ORF1 | DPYSL3 | -0.964 | 0.000 | -1.000 | up |
| G056910 | EXOC7 | -0.915 | 0.001 | -1.000 | up |
| SLC7A11-AS1 | CRELD1 | -0.931 | 0.001 | -1.000 | up |
| SLC7A11-AS1 | RPS6KA5 | -0.903 | 0.002 | -1.000 | up |
| SLC7A11-AS1 | DPYSL3 | -0.987 | 0.000 | -1.000 | up |
| RP11-21M24.3 | EXOC7 | -0.945 | 0.000 | -1.000 | up |
| RP11-21M24.3 | CHRNB3 | -0.921 | 0.001 | -1.000 | up |
| RP11-674P19.2 | FXYD6 | -0.956 | 0.000 | -1.000 | up |
| uc.172 | HSP90AA1 | -0.912 | 0.002 | -1.000 | up |
| CTC-462L7.1 | OLFM1 | -0.933 | 0.001 | -1.000 | up |
| RP11-70P17.1 | CALB2 | -0.947 | 0.000 | -1.000 | up |
| RP11-70P17.1 | PPP2R1A | -0.907 | 0.002 | -1.000 | up |
| ITGA9-AS1 | CRELD1 | -0.939 | 0.001 | -1.000 | up |
| ITGA9-AS1 | ADGRL2 | -0.906 | 0.002 | -1.000 | up |
| PPIEL | SPTB | -0.903 | 0.002 | -1.000 | up |
| LOC100505736 | ATP2B4 | -0.918 | 0.001 | -1.000 | up |
| G075352 | SOS1 | -0.908 | 0.002 | -1.000 | up |
| G038103 | BTBD9 | -0.927 | 0.001 | -1.000 | up |
| CTA-217C2.2 | SIPA1L2 | -0.924 | 0.001 | -1.000 | up |
| XLOC_l2_009332 | SPTB | -0.917 | 0.001 | -1.000 | up |
| XLOC_l2_009332 | RPS6KA5 | -0.969 | 0.000 | -1.000 | up |
| Alpha_250_dna | DAB1 | -0.920 | 0.001 | -1.000 | up |
| CTB-193M12.1 | ACTN1 | -0.902 | 0.002 | -1.000 | up |
| G089550 | FXYD6 | -0.945 | 0.000 | -1.000 | up |
| G060633 | CRELD1 | -0.906 | 0.002 | -1.000 | up |
| G060633 | ATP2B4 | -0.925 | 0.001 | -1.000 | up |
| G060633 | PTCH1 | -0.909 | 0.002 | -1.000 | up |
| G060633 | ERC2 | -0.944 | 0.000 | -1.000 | up |
| G060633 | FILIP1 | -0.935 | 0.001 | -1.000 | up |
| G060633 | PPP2R1A | -0.905 | 0.002 | -1.000 | up |
| G029454 | ATP2B4 | -0.945 | 0.000 | -1.000 | up |
| G029454 | TSPAN7 | -0.901 | 0.002 | -1.000 | up |
| G020731 | TENM4 | -0.930 | 0.001 | -1.000 | up |
| IGBP1-AS2 | EXOC7 | -0.943 | 0.000 | -1.000 | up |
| IGBP1-AS2 | NRP2 | -0.943 | 0.000 | -1.000 | up |
| IGBP1-AS2 | CHRNA3 | -0.919 | 0.001 | -1.000 | up |
| G011310 | DAB1 | -0.928 | 0.001 | -1.000 | up |
| G011310 | ERC2 | -0.922 | 0.001 | -1.000 | up |
| G011310 | GPSM2 | -0.953 | 0.000 | -1.000 | up |
| G011310 | CALB2 | -0.943 | 0.000 | -1.000 | up |
| G011310 | PPP2R1A | -0.946 | 0.000 | -1.000 | up |
| ASAP1-IT1 | HSPA8 | -0.901 | 0.002 | -1.000 | up |
| ASAP1-IT1 | NRP2 | -0.914 | 0.001 | -1.000 | up |
| ASAP1-IT1 | DAB1 | -0.953 | 0.000 | -1.000 | up |
| ASAP1-IT1 | DNAJB1 | -0.914 | 0.001 | -1.000 | up |
| RP11-498C9.12 | BTBD9 | -0.907 | 0.002 | -1.000 | up |

| | | | | | |
|---|---|---|---|---|---|
| AK124233 | ACTN1 | -0.905 | 0.002 | -1.000 | up |
| AK124233 | OLFM1 | -0.924 | 0.001 | -1.000 | up |
| G061035 | PTCH1 | -0.906 | 0.002 | -1.000 | up |
| XLOC_001223 | CRELD1 | -0.964 | 0.000 | -1.000 | up |
| XLOC_001223 | ARHGAP32 | -0.907 | 0.002 | -1.000 | up |
| RP11-3K16.2 | ADGRL2 | -0.911 | 0.002 | -1.000 | up |
| G008916 | HSP90AA1 | -0.929 | 0.001 | -1.000 | up |
| G008916 | GPSM2 | -0.912 | 0.002 | -1.000 | up |
| G008916 | PPP2R1A | -0.941 | 0.000 | -1.000 | up |
| RP11-187C18.2 | CHRNB3 | -0.907 | 0.002 | -1.000 | up |
| RP11-187C18.2 | NRP2 | -0.927 | 0.001 | -1.000 | up |
| BC033330 | ELFN1 | -0.930 | 0.001 | -1.000 | up |
| G053606 | TSPAN7 | -0.907 | 0.002 | -1.000 | up |
| XLOC_006150 | GPSM2 | -0.921 | 0.001 | -1.000 | up |
| G006738 | TSPAN7 | -0.905 | 0.002 | -1.000 | up |
| G087508 | SPTB | -0.914 | 0.002 | -1.000 | up |
| G087508 | SEPSECS | -0.901 | 0.002 | -1.000 | up |
| G087508 | CABP1 | -0.932 | 0.001 | -1.000 | up |
| G068163 | ATP2B4 | -0.928 | 0.001 | -1.000 | up |
| G049993 | NRP2 | -0.930 | 0.001 | -1.000 | up |
| G073552 | TENM4 | -0.978 | 0.000 | -1.000 | up |
| G073552 | AP1G2 | -0.917 | 0.001 | -1.000 | up |
| G073552 | ACTN1 | -0.924 | 0.001 | -1.000 | up |
| G086585 | HSP90AA1 | -0.907 | 0.002 | -1.000 | up |
| RP4-761J14.9 | PTCH1 | -0.921 | 0.001 | -1.000 | up |
| RP4-816N1.7 | OLFM1 | -0.912 | 0.002 | -1.000 | up |
| RP4-816N1.7 | FXYD6 | -0.961 | 0.000 | -1.000 | up |
| AQP7P3 | OLFM1 | -0.946 | 0.000 | -1.000 | up |
| GSE61474_XLOC_0203 | PHB | -0.963 | 0.000 | -1.000 | up |
| G009303 | HSPA8 | -0.904 | 0.002 | -1.000 | up |
| G009303 | CRH | -0.916 | 0.001 | -1.000 | up |
| G057625 | ALDOC | -0.917 | 0.001 | -1.000 | up |
| G021277 | ATP2B4 | -0.921 | 0.001 | -1.000 | up |
| CHKB-CPT1B | DNAJB1 | -0.900 | 0.002 | -1.000 | up |
| G080487 | ELFN1 | -0.963 | 0.000 | -1.000 | up |
| G080487 | ATP2B4 | -0.915 | 0.001 | -1.000 | up |
| G080487 | CABP1 | -0.922 | 0.001 | -1.000 | up |
| AP000438.2 | ARHGAP32 | -0.904 | 0.002 | -1.000 | up |
| AP000438.2 | ERC2 | -0.944 | 0.000 | -1.000 | up |
| AP000438.2 | PTPRN2 | -0.921 | 0.001 | -1.000 | up |
| AP000438.2 | MAP2 | -0.972 | 0.000 | -1.000 | up |
| AP000438.2 | DPYSL5 | -0.933 | 0.001 | -1.000 | up |
| AP000438.2 | PPP2R1A | -0.957 | 0.000 | -1.000 | up |
| AC074363.1 | PHB | -0.904 | 0.002 | -1.000 | up |
| AC074363.1 | HSP90AA1 | -0.914 | 0.002 | -1.000 | up |
| AC074363.1 | CHRNA3 | -0.923 | 0.001 | -1.000 | up |
| AC074363.1 | OLFM1 | -0.900 | 0.002 | -1.000 | up |
| G046678 | TENM4 | -0.936 | 0.001 | -1.000 | up |
| G046678 | NRP2 | -0.986 | 0.000 | -1.000 | up |
| G031832 | HSPA8 | -0.901 | 0.002 | -1.000 | up |
| PHF2P2 | DAB1 | -0.905 | 0.002 | -1.000 | up |
| G081487 | DAB1 | -0.921 | 0.001 | -1.000 | up |
| G081487 | DNAJB1 | -0.982 | 0.000 | -1.000 | up |
| AC016682.1 | TENM4 | -0.967 | 0.000 | -1.000 | up |
| AC016682.1 | AP1G2 | -0.943 | 0.000 | -1.000 | up |
| AC016682.1 | NRP2 | -0.909 | 0.002 | -1.000 | up |
| RP11-466A19.1 | CRELD1 | -0.944 | 0.000 | -1.000 | up |
| RP11-466A19.1 | PTPRN2 | -0.931 | 0.001 | -1.000 | up |
| RP11-466A19.1 | DPYSL5 | -0.946 | 0.000 | -1.000 | up |
| RP11-466A19.1 | DPYSL3 | -0.967 | 0.000 | -1.000 | up |
| AP001625.6 | DAB1 | -0.907 | 0.002 | -1.000 | up |
| AP001625.6 | PTCH1 | -0.946 | 0.000 | -1.000 | up |
| AP001625.6 | BTBD9 | -0.923 | 0.001 | -1.000 | up |
| AP001625.6 | PPP2R1A | -0.924 | 0.001 | -1.000 | up |
| G063174 | TSPAN7 | -0.918 | 0.001 | -1.000 | up |

| | | | | | |
|---|---|---|---|---|---|
| RP11-484O2.1 | TENM4 | -0.921 | 0.001 | -1.000 | up |
| RP11-484O2.1 | AP1G2 | -0.926 | 0.001 | -1.000 | up |
| RP11-484O2.1 | NRP2 | -0.932 | 0.001 | -1.000 | up |
| XLOC_008025 | CRELD1 | -0.922 | 0.001 | -1.000 | up |
| XLOC_008025 | ARHGAP32 | -0.927 | 0.001 | -1.000 | up |
| XLOC_008025 | ATP2B4 | -0.966 | 0.000 | -1.000 | up |
| XLOC_008025 | PTCH1 | -0.940 | 0.001 | -1.000 | up |
| G050702 | TENM4 | -0.952 | 0.000 | -1.000 | up |
| G050702 | NRP2 | -0.942 | 0.000 | -1.000 | up |
| G050702 | SPTB | -0.933 | 0.001 | -1.000 | up |
| G050702 | SEPSECS | -0.903 | 0.002 | -1.000 | up |
| G082315 | OLFM1 | -0.935 | 0.001 | -1.000 | up |
| RP11-737O24.3 | HSP90AA1 | -0.960 | 0.000 | -1.000 | up |
| RP11-737O24.3 | CHRNA3 | -0.961 | 0.000 | -1.000 | up |
| XLOC_003405 | DAB1 | -0.930 | 0.001 | -1.000 | up |
| XLOC_003405 | DNAJB1 | -0.903 | 0.002 | -1.000 | up |
| G009235 | EXOC7 | -0.946 | 0.000 | -1.000 | up |
| G009235 | AP1G2 | -0.916 | 0.001 | -1.000 | up |
| G063995 | TENM4 | -0.959 | 0.000 | -1.000 | up |
| G063995 | NRP2 | -0.934 | 0.001 | -1.000 | up |
| G063995 | ACTN1 | -0.936 | 0.001 | -1.000 | up |
| AADACL2-AS1 | DAB1 | -0.932 | 0.001 | -1.000 | up |
| RP11-669E14.6 | HSPA8 | -0.908 | 0.002 | -1.000 | up |
| RP11-669E14.6 | DAB1 | -0.987 | 0.000 | -1.000 | up |
| RP11-669E14.6 | BTBD9 | -0.924 | 0.001 | -1.000 | up |
| NEAT1 | BTBD9 | -0.978 | 0.000 | -1.000 | up |
| NEAT1 | PPP2R1A | -0.905 | 0.002 | -1.000 | up |
| XLOC_012579 | MCTP2 | -0.965 | 0.000 | -1.000 | up |
| AL832615 | SOS1 | -0.909 | 0.002 | -1.000 | up |
| XLOC_l2_011118 | ATP2B4 | -0.913 | 0.002 | -1.000 | up |
| XLOC_l2_011118 | PTCH1 | -0.925 | 0.001 | -1.000 | up |
| RN7SKP264 | HSP90AA1 | -0.956 | 0.000 | -1.000 | up |
| RN7SKP264 | SIPA1L2 | -0.939 | 0.001 | -1.000 | up |
| LINC00379 | ATP2B4 | -0.910 | 0.002 | -1.000 | up |
| LINC00379 | TSPAN7 | -0.915 | 0.001 | -1.000 | up |
| LINC00379 | CABP1 | -0.909 | 0.002 | -1.000 | up |
| LIPE-AS1 | FXYD6 | -0.911 | 0.002 | -1.000 | up |
| PMCHL2 | PPP2R1A | -0.919 | 0.001 | -1.000 | up |
| AX747064 | OLFM1 | -0.977 | 0.000 | -1.000 | up |
| RP11-219C20.3 | PHB | -0.906 | 0.002 | -1.000 | up |
| RP11-219C20.3 | HSP90AA1 | -0.969 | 0.000 | -1.000 | up |
| RP11-219C20.3 | CHRNA3 | -0.910 | 0.002 | -1.000 | up |
| G070831 | TSPAN7 | -0.900 | 0.002 | -1.000 | up |
| G070831 | CABP1 | -0.954 | 0.000 | -1.000 | up |
| ZFAT-AS1 | TENM4 | -0.905 | 0.002 | -1.000 | up |
| ZFAT-AS1 | AP1G2 | -0.951 | 0.000 | -1.000 | up |
| ZFAT-AS1 | SPTB | -0.915 | 0.001 | -1.000 | up |
| G040015 | CHRNA3 | -0.916 | 0.001 | -1.000 | up |
| G087395 | DAB1 | -0.906 | 0.002 | -1.000 | up |
| G087395 | BTBD9 | -0.924 | 0.001 | -1.000 | up |
| G087265 | TENM4 | -0.944 | 0.000 | -1.000 | up |
| G087265 | EXOC7 | -0.922 | 0.001 | -1.000 | up |
| G087265 | AP1G2 | -0.903 | 0.002 | -1.000 | up |
| G087265 | NRP2 | -0.920 | 0.001 | -1.000 | up |
| G087265 | ACTN1 | -0.907 | 0.002 | -1.000 | up |
| G087265 | CHRNA3 | -0.905 | 0.002 | -1.000 | up |
| G072062 | CRELD1 | -0.970 | 0.000 | -1.000 | up |
| G072062 | ARHGAP32 | -0.905 | 0.002 | -1.000 | up |
| G072062 | ATP2B4 | -0.944 | 0.000 | -1.000 | up |
| Z69720.2 | SOS1 | -0.902 | 0.002 | -1.000 | up |
| Z69720.2 | HSP90AA1 | -0.910 | 0.002 | -1.000 | up |
| Z69720.2 | CHRNA3 | -0.941 | 0.000 | -1.000 | up |
| CTC-261N6.1 | DNAJB1 | -0.914 | 0.001 | -1.000 | up |
| CTC-261N6.1 | CRH | -0.907 | 0.002 | -1.000 | up |
| G081302 | DNAJB1 | -0.907 | 0.002 | -1.000 | up |

| | | | | | |
|---|---|---|---|---|---|
| G081302 | PTCH1 | -0.955 | 0.000 | -1.000 | up |
| BC034456 | HSP90AA1 | -0.911 | 0.002 | -1.000 | up |
| G067669 | MCTP2 | -0.928 | 0.001 | -1.000 | up |
| RP11-18D7.2 | SPTB | -0.904 | 0.002 | -1.000 | up |
| G071732 | CHRNA7 | -0.909 | 0.002 | -1.000 | up |
| G071640 | CALB2 | -0.903 | 0.002 | -1.000 | up |
| AC107218.3 | PHB | -0.902 | 0.002 | -1.000 | up |
| CTB-102L5.8 | CHRNB3 | -0.914 | 0.002 | -1.000 | up |
| CTB-102L5.8 | CRH | -0.931 | 0.001 | -1.000 | up |
| G047392 | NRP2 | -0.901 | 0.002 | -1.000 | up |
| RP11-717D12.1 | ATP2B4 | -0.967 | 0.000 | -1.000 | up |
| RP11-717D12.1 | CABP1 | -0.952 | 0.000 | -1.000 | up |
| AK093193 | TENM4 | -0.905 | 0.002 | -1.000 | up |
| AK093193 | SPTB | -0.949 | 0.000 | -1.000 | up |
| AK093193 | SEPSECS | -0.973 | 0.000 | -1.000 | up |
| AK093193 | TSPAN7 | -0.930 | 0.001 | -1.000 | up |
| AK093193 | CABP1 | -0.915 | 0.001 | -1.000 | up |
| LOC101927164 | TENM4 | -0.911 | 0.002 | -1.000 | up |
| LOC101927164 | NRP2 | -0.921 | 0.001 | -1.000 | up |
| LOC101927164 | SPTB | -0.947 | 0.000 | -1.000 | up |
| KLK8 | NPY2R | -0.918 | 0.001 | -1.000 | up |
| G000948 | FXYD6 | -0.954 | 0.000 | -1.000 | up |
| G045594 | ARHGAP32 | -0.962 | 0.000 | -1.000 | up |
| G045594 | MAP2 | -0.905 | 0.002 | -1.000 | up |
| CTD-2050E21.1 | SPTB | -0.940 | 0.001 | -1.000 | up |
| CTD-2050E21.1 | CABP1 | -0.910 | 0.002 | -1.000 | up |
| LOC727896 | OLFM1 | -0.964 | 0.000 | -1.000 | up |
| G036779 | SIPA1L2 | -0.964 | 0.000 | -1.000 | up |
| G036779 | ALDOC | -0.907 | 0.002 | -1.000 | up |
| AC003682.16 | SIPA1L2 | -0.942 | 0.000 | -1.000 | up |
| AC003682.16 | ALDOC | -0.964 | 0.000 | -1.000 | up |
| CRAT37 | NPY2R | -0.925 | 0.001 | -1.000 | up |
| SNHG7 | CABP1 | -0.924 | 0.001 | -1.000 | up |
| G046027 | SIPA1L2 | -0.900 | 0.002 | -1.000 | up |
| RP1-124C6.1 | NRP2 | -0.924 | 0.001 | -1.000 | up |
| RP1-124C6.1 | CRH | -0.926 | 0.001 | -1.000 | up |
| G022034 | ALDOC | -0.971 | 0.000 | -1.000 | up |
| uc.49 | ERC2 | -0.915 | 0.001 | -1.000 | up |
| uc.49 | CALB2 | -0.927 | 0.001 | -1.000 | up |
| uc.49 | MAP2 | -0.909 | 0.002 | -1.000 | up |
| uc.49 | BTBD9 | -0.913 | 0.002 | -1.000 | up |
| uc.49 | PPP2R1A | -0.926 | 0.001 | -1.000 | up |
| G043880 | CHRNB3 | -0.929 | 0.001 | -1.000 | up |
| RP11-424I19.2 | TENM4 | -0.915 | 0.001 | -1.000 | up |
| RP11-424I19.2 | SEPSECS | -0.923 | 0.001 | -1.000 | up |
| G006360 | ATP2B4 | -0.988 | 0.000 | -1.000 | up |
| G006360 | ERC2 | -0.900 | 0.002 | -1.000 | up |
| G006360 | SEPSECS | -0.917 | 0.001 | -1.000 | up |
| G006360 | CABP1 | -0.916 | 0.001 | -1.000 | up |
| G053959 | EXOC7 | -0.930 | 0.001 | -1.000 | up |
| XLOC_013958 | SPTB | -0.956 | 0.000 | -1.000 | up |
| XLOC_013958 | SEPSECS | -0.954 | 0.000 | -1.000 | up |
| XLOC_013958 | TSPAN7 | -0.945 | 0.000 | -1.000 | up |
| XLOC_013958 | CABP1 | -0.983 | 0.000 | -1.000 | up |
| RP11-166B2.5 | NRP2 | -0.906 | 0.002 | -1.000 | up |
| G021365 | CRH | -0.979 | 0.000 | -1.000 | up |
| XLOC_009306 | CRELD1 | -0.920 | 0.001 | -1.000 | up |
| XLOC_009306 | ARHGAP32 | -0.929 | 0.001 | -1.000 | up |
| XLOC_009306 | MAP2 | -0.904 | 0.002 | -1.000 | up |
| XLOC_009306 | DPYSL5 | -0.914 | 0.001 | -1.000 | up |
| G060178 | TENM4 | -0.906 | 0.002 | -1.000 | up |
| G063524 | CABP1 | -0.916 | 0.001 | -1.000 | up |
| GSE61474_XLOC_0280 | PTCH1 | -0.911 | 0.002 | -1.000 | up |
| G071944 | HSPA8 | -0.922 | 0.001 | -1.000 | up |
| G071944 | CRH | -0.967 | 0.000 | -1.000 | up |

| | | | | | |
|---|---|---|---|---|---|
| G011388 | ALDOC | -0.939 | 0.001 | -1.000 | up |
| XLOC_003837 | NRP2 | -0.940 | 0.001 | -1.000 | up |
| XLOC_003837 | DAB1 | -0.918 | 0.001 | -1.000 | up |
| BX648501 | TENM4 | -0.926 | 0.001 | -1.000 | up |
| BX648501 | EXOC7 | -0.948 | 0.000 | -1.000 | up |
| BX648501 | AP1G2 | -0.930 | 0.001 | -1.000 | up |
| BX648501 | NRP2 | -0.910 | 0.002 | -1.000 | up |
| BX648501 | SPTB | -0.931 | 0.001 | -1.000 | up |
| G060010 | TENM4 | -0.906 | 0.002 | -1.000 | up |
| G060010 | ACTN1 | -0.923 | 0.001 | -1.000 | up |
| G073689 | ERC2 | -0.944 | 0.000 | -1.000 | up |
| G073689 | GPSM2 | -0.913 | 0.002 | -1.000 | up |
| G073689 | CALB2 | -0.936 | 0.001 | -1.000 | up |
| G073689 | PPP2R1A | -0.963 | 0.000 | -1.000 | up |
| G072364 | ARHGAP32 | -0.967 | 0.000 | -1.000 | up |
| RP11-567E21.3 | NPY2R | -0.903 | 0.002 | -1.000 | up |
| G036156 | PHB | -0.934 | 0.001 | -1.000 | up |
| G036156 | HSP90AA1 | -0.908 | 0.002 | -1.000 | up |
| G036156 | CHRNA3 | -0.921 | 0.001 | -1.000 | up |
| G032508 | ELFN1 | -0.948 | 0.000 | -1.000 | up |
| CYP2D7 | TENM4 | -0.947 | 0.000 | -1.000 | up |
| CYP2D7 | ACTN1 | -0.966 | 0.000 | -1.000 | up |
| CTD-3118D11.2 | OLFM1 | -0.905 | 0.002 | -1.000 | up |
| RP11-374P20.4 | SPTB | -0.905 | 0.002 | -1.000 | up |
| RP11-374P20.4 | CABP1 | -0.947 | 0.000 | -1.000 | up |
| G063659 | PTCH1 | -0.934 | 0.001 | -1.000 | up |
| G068469 | OLFM1 | -0.972 | 0.000 | -1.000 | up |
| G020793 | ARHGAP32 | -0.994 | 0.000 | -1.000 | up |
| G020793 | MAP2 | -0.911 | 0.002 | -1.000 | up |
| G054530 | ERC2 | -0.916 | 0.001 | -1.000 | up |
| G054530 | PPP2R1A | -0.934 | 0.001 | -1.000 | up |
| CTD-2023M8.1 | ADGRL2 | -0.908 | 0.002 | -1.000 | up |
| CTD-2023M8.1 | DAB1 | -0.942 | 0.000 | -1.000 | up |
| G045736 | FILIP1 | -0.920 | 0.001 | -1.000 | up |
| G045736 | SPTB | -0.939 | 0.001 | -1.000 | up |
| G045736 | SEPSECS | -0.971 | 0.000 | -1.000 | up |
| G045736 | CABP1 | -0.917 | 0.001 | -1.000 | up |
| G045736 | RPS6KA5 | -0.935 | 0.001 | -1.000 | up |
| EML2-AS1 | CRELD1 | -0.913 | 0.002 | -1.000 | up |
| EML2-AS1 | ERC2 | -0.967 | 0.000 | -1.000 | up |
| EML2-AS1 | CALB2 | -0.910 | 0.002 | -1.000 | up |
| EML2-AS1 | PTPRN2 | -0.987 | 0.000 | -1.000 | up |
| EML2-AS1 | MAP2 | -0.962 | 0.000 | -1.000 | up |
| EML2-AS1 | DPYSL5 | -0.983 | 0.000 | -1.000 | up |
| LINC01149 | ARHGAP32 | -0.910 | 0.002 | -1.000 | up |
| LINC01149 | ATP2B4 | -0.929 | 0.001 | -1.000 | up |
| G030124 | CRELD1 | -0.916 | 0.001 | -1.000 | up |
| G030124 | HSP90AA1 | -0.906 | 0.002 | -1.000 | up |
| G030124 | ADGRL2 | -0.981 | 0.000 | -1.000 | up |
| G030124 | SIPA1L2 | -0.965 | 0.000 | -1.000 | up |
| G065302 | DNAJB1 | -0.912 | 0.002 | -1.000 | up |
| G065302 | BTBD9 | -0.920 | 0.001 | -1.000 | up |
| XLOC_006124 | RPS6KA5 | -0.909 | 0.002 | -1.000 | up |
| G072890 | PTCH1 | -0.930 | 0.001 | -1.000 | up |
| G072890 | BTBD9 | -0.946 | 0.000 | -1.000 | up |
| LOC101927691 | PTCH1 | -0.971 | 0.000 | -1.000 | up |
| LINC00930 | NRP2 | -0.913 | 0.002 | -1.000 | up |
| LINC00930 | CRH | -0.967 | 0.000 | -1.000 | up |
| AK025321 | CHRNA3 | -0.923 | 0.001 | -1.000 | up |
| G002461 | ALDOC | -0.938 | 0.001 | -1.000 | up |
| LOC101926908 | DAB1 | -0.976 | 0.000 | -1.000 | up |
| LOC101926908 | DNAJB1 | -0.920 | 0.001 | -1.000 | up |
| G003431 | CRELD1 | -0.940 | 0.001 | -1.000 | up |
| G003431 | SIPA1L2 | -0.911 | 0.002 | -1.000 | up |
| G003431 | GPSM2 | -0.908 | 0.002 | -1.000 | up |

| | | | | | |
|---|---|---|---|---|---|
| G003431 | DPYSL3 | -0.946 | 0.000 | -1.000 | up |
| G016657 | ATP2B4 | -0.947 | 0.000 | -1.000 | up |
| G016657 | SEPSECS | -0.922 | 0.001 | -1.000 | up |
| G016657 | TSPAN7 | -0.916 | 0.001 | -1.000 | up |
| G016657 | CABP1 | -0.979 | 0.000 | -1.000 | up |
| RMRPP3 | PTCH1 | -0.960 | 0.000 | -1.000 | up |
| RMRPP3 | BTBD9 | -0.932 | 0.001 | -1.000 | up |
| XLOC_012740 | CRELD1 | -0.911 | 0.002 | -1.000 | up |
| XLOC_012740 | DAB1 | -0.914 | 0.002 | -1.000 | up |
| XLOC_012740 | ERC2 | -0.942 | 0.000 | -1.000 | up |
| XLOC_012740 | CALB2 | -0.901 | 0.002 | -1.000 | up |
| XLOC_012740 | PTPRN2 | -0.911 | 0.002 | -1.000 | up |
| XLOC_012740 | MAP2 | -0.923 | 0.001 | -1.000 | up |
| XLOC_012740 | DPYSL5 | -0.908 | 0.002 | -1.000 | up |
| XLOC_012740 | PPP2R1A | -0.919 | 0.001 | -1.000 | up |
| RP11-96H17.3 | CRH | -0.921 | 0.001 | -1.000 | up |
| ELOVL2-AS1 | CRELD1 | -0.919 | 0.001 | -1.000 | up |
| uc.474 | HSPA8 | -0.928 | 0.001 | -1.000 | up |
| uc.474 | DAB1 | -0.927 | 0.001 | -1.000 | up |
| uc.474 | BTBD9 | -0.953 | 0.000 | -1.000 | up |
| GOLGA8B | BTBD9 | -0.942 | 0.000 | -1.000 | up |
| XLOC_006485 | SOS1 | -0.910 | 0.002 | -1.000 | up |
| XLOC_006485 | CHRNB3 | -0.923 | 0.001 | -1.000 | up |
| TMLHE-AS1 | DNAJB1 | -0.918 | 0.001 | -1.000 | up |
| RP11-30O15.1 | SPTB | -0.920 | 0.001 | -1.000 | up |
| RP11-30O15.1 | TSPAN7 | -0.927 | 0.001 | -1.000 | up |
| RP11-30O15.1 | CABP1 | -0.921 | 0.001 | -1.000 | up |
| RP11-768G7.2 | EXOC7 | -0.909 | 0.002 | -1.000 | up |
| RP11-768G7.2 | ACTN1 | -0.962 | 0.000 | -1.000 | up |
| G058409 | PTCH1 | -0.914 | 0.002 | -1.000 | up |
| G009329 | PTCH1 | -0.914 | 0.001 | -1.000 | up |
| G067215 | SPTB | -0.920 | 0.001 | -1.000 | up |
| G067215 | SEPSECS | -0.906 | 0.002 | -1.000 | up |
| G067215 | CABP1 | -0.948 | 0.000 | -1.000 | up |
| LOC642846 | ACTN1 | -0.911 | 0.002 | -1.000 | up |
| G020470 | ACTN1 | -0.901 | 0.002 | -1.000 | up |
| G015181 | ARHGAP32 | -0.957 | 0.000 | -1.000 | up |
| G015181 | ATP2B4 | -0.912 | 0.002 | -1.000 | up |
| G015181 | PTCH1 | -0.956 | 0.000 | -1.000 | up |
| G037029 | ATP2B4 | -0.952 | 0.000 | -1.000 | up |
| G037029 | SPTB | -0.900 | 0.002 | -1.000 | up |
| G037029 | SEPSECS | -0.954 | 0.000 | -1.000 | up |
| G037029 | TSPAN7 | -0.938 | 0.001 | -1.000 | up |
| G037029 | CABP1 | -0.981 | 0.000 | -1.000 | up |
| G082713 | HSP90AA1 | -0.901 | 0.002 | -1.000 | up |
| G010088 | SPTB | -0.909 | 0.002 | -1.000 | up |
| G010088 | TSPAN7 | -0.925 | 0.001 | -1.000 | up |
| G010088 | CABP1 | -0.931 | 0.001 | -1.000 | up |
| G010088 | CHRNA7 | -0.917 | 0.001 | -1.000 | up |
| XLOC_008270 | CRH | -0.935 | 0.001 | -1.000 | up |
| G046492 | ATP2B4 | -0.922 | 0.001 | -1.000 | up |
| G046492 | SEPSECS | -0.907 | 0.002 | -1.000 | up |
| G046492 | TSPAN7 | -0.916 | 0.001 | -1.000 | up |
| G046492 | CABP1 | -0.902 | 0.002 | -1.000 | up |
| AC026188.1 | CHRNA3 | -0.914 | 0.001 | -1.000 | up |
| G052905 | NRP2 | -0.941 | 0.000 | -1.000 | up |
| RP11-551L14.6 | ELFN1 | -0.949 | 0.000 | -1.000 | up |
| RP11-551L14.6 | CHRNA7 | -0.926 | 0.001 | -1.000 | up |
| uc.50 | TENM4 | -0.932 | 0.001 | -1.000 | up |
| uc.50 | SPTB | -0.902 | 0.002 | -1.000 | up |
| G080747 | BTBD9 | -0.945 | 0.000 | -1.000 | up |
| RP4-760C5.3 | TENM4 | -0.935 | 0.001 | -1.000 | up |
| RP4-760C5.3 | AP1G2 | -0.952 | 0.000 | -1.000 | up |
| CTD-3193O13.1 | CALB2 | -0.926 | 0.001 | -1.000 | up |
| CTD-3193O13.1 | TSPAN7 | -0.905 | 0.002 | -1.000 | up |

| | | | | | |
|---|---|---|---|---|---|
| G072257 | ATP2B4 | -0.912 | 0.002 | -1.000 | up |
| AF086132 | ALDOC | -0.944 | 0.000 | -1.000 | up |
| CTC-250I14.3 | HSP90AA1 | -0.914 | 0.001 | -1.000 | up |
| CTC-250I14.3 | ADGRL2 | -0.903 | 0.002 | -1.000 | up |
| CTC-250I14.3 | NPY2R | -0.911 | 0.002 | -1.000 | up |
| G023385 | SOS1 | -0.923 | 0.001 | -1.000 | up |
| G051586 | ARHGAP32 | -0.911 | 0.002 | -1.000 | up |
| G051586 | PTCH1 | -0.923 | 0.001 | -1.000 | up |
| G051586 | ERC2 | -0.919 | 0.001 | -1.000 | up |
| G051586 | MAP2 | -0.906 | 0.002 | -1.000 | up |
| RP11-126O1.4 | DNAJB1 | -0.942 | 0.000 | -1.000 | up |
| RP11-651L5.2 | SPTB | -0.904 | 0.002 | -1.000 | up |
| RP11-651L5.2 | CHRNA7 | -0.911 | 0.002 | -1.000 | up |
| XLOC_000048 | CABP1 | -0.928 | 0.001 | -1.000 | up |
| LOC101926941 | BTBD9 | -0.920 | 0.001 | -1.000 | up |
| LOC101926941 | PPP2R1A | -0.901 | 0.002 | -1.000 | up |
| G053358 | HSPA8 | -0.934 | 0.001 | -1.000 | up |
| G053358 | CRH | -0.913 | 0.002 | -1.000 | up |
| G068224 | CHRNA7 | -0.952 | 0.000 | -1.000 | up |
| BCRP3 | AP1G2 | -0.900 | 0.002 | -1.000 | up |
| BCRP3 | NRP2 | -0.949 | 0.000 | -1.000 | up |
| HM13-IT1 | TENM4 | -0.914 | 0.002 | -1.000 | up |
| HM13-IT1 | EXOC7 | -0.914 | 0.002 | -1.000 | up |
| HM13-IT1 | ACTN1 | -0.950 | 0.000 | -1.000 | up |
| G009199 | BTBD9 | -0.938 | 0.001 | -1.000 | up |
| G033096 | FXYD6 | -0.951 | 0.000 | -1.000 | up |
| G089283 | CALB2 | -0.928 | 0.001 | -1.000 | up |
| AC025442.3 | SIPA1L2 | -0.909 | 0.002 | -1.000 | up |
| AC025442.3 | GPSM2 | -0.934 | 0.001 | -1.000 | up |
| G013465 | GPSM2 | -0.982 | 0.000 | -1.000 | up |
| G013465 | DPYSL5 | -0.905 | 0.002 | -1.000 | up |
| G013465 | DPYSL3 | -0.943 | 0.000 | -1.000 | up |
| G071259 | SPTB | -0.915 | 0.001 | -1.000 | up |
| G071259 | CABP1 | -0.963 | 0.000 | -1.000 | up |
| AC004054.1 | SOS1 | -0.902 | 0.002 | -1.000 | up |
| RP5-1112D6.7 | ERC2 | -0.916 | 0.001 | -1.000 | up |
| RP5-1112D6.7 | CALB2 | -0.937 | 0.001 | -1.000 | up |
| MIMT1 | DAB1 | -0.918 | 0.001 | -1.000 | up |
| MIMT1 | CRH | -0.902 | 0.002 | -1.000 | up |
| G068890 | TENM4 | -0.928 | 0.001 | -1.000 | up |
| G068890 | EXOC7 | -0.916 | 0.001 | -1.000 | up |
| G068890 | AP1G2 | -0.966 | 0.000 | -1.000 | up |
| G068890 | SPTB | -0.904 | 0.002 | -1.000 | up |
| G010941 | CHRNA3 | -0.914 | 0.002 | -1.000 | up |
| SPTY2D1-AS1 | CHRNA7 | -0.960 | 0.000 | -1.000 | up |
| G031271 | CRH | -0.911 | 0.002 | -1.000 | up |
| G029241 | ADGRL2 | -0.925 | 0.001 | -1.000 | up |
| G029241 | DPYSL5 | -0.904 | 0.002 | -1.000 | up |
| G029241 | DPYSL3 | -0.914 | 0.002 | -1.000 | up |
| AK022848 | EXOC7 | -0.958 | 0.000 | -1.000 | up |
| AK022848 | AP1G2 | -0.924 | 0.001 | -1.000 | up |
| LOC100506834 | CRELD1 | -0.966 | 0.000 | -1.000 | up |
| LOC100506834 | ARHGAP32 | -0.925 | 0.001 | -1.000 | up |
| LOC100506834 | PTPRN2 | -0.928 | 0.001 | -1.000 | up |
| LOC100506834 | DPYSL5 | -0.939 | 0.001 | -1.000 | up |
| G051433 | TENM4 | -0.903 | 0.002 | -1.000 | up |
| G051433 | NRP2 | -0.922 | 0.001 | -1.000 | up |
| DLG3-AS1 | HSP90AA1 | -0.914 | 0.001 | -1.000 | up |
| DLG3-AS1 | CHRNA3 | -0.911 | 0.002 | -1.000 | up |
| DLG3-AS1 | BTBD9 | -0.906 | 0.002 | -1.000 | up |
| EF565122 | CRELD1 | -0.916 | 0.001 | -1.000 | up |
| EF565122 | PTPRN2 | -0.932 | 0.001 | -1.000 | up |
| EF565122 | DPYSL5 | -0.959 | 0.000 | -1.000 | up |
| EF565122 | DPYSL3 | -0.966 | 0.000 | -1.000 | up |
| G088912 | PHB | -0.903 | 0.002 | -1.000 | up |

| | | | | | |
|---|---|---|---|---|---|
| G088912 | HSP90AA1 | -0.927 | 0.001 | -1.000 | up |
| G088912 | ADGRL2 | -0.905 | 0.002 | -1.000 | up |
| G088912 | SIPA1L2 | -0.912 | 0.002 | -1.000 | up |
| CTD-2015A6.1 | HSP90AA1 | -0.916 | 0.001 | -1.000 | up |
| CTD-2015A6.1 | DPYSL3 | -0.936 | 0.001 | -1.000 | up |
| G084746 | SOS1 | -0.968 | 0.000 | -1.000 | up |
| G013179 | DPYSL3 | -0.926 | 0.001 | -1.000 | up |
| LOC101929427 | HSP90AA1 | -0.903 | 0.002 | -1.000 | up |
| G074298 | CRH | -0.943 | 0.000 | -1.000 | up |
| G010533 | PTCH1 | -0.989 | 0.000 | -1.000 | up |
| G010224 | ADGRL2 | -0.920 | 0.001 | -1.000 | up |
| G010224 | DAB1 | -0.910 | 0.002 | -1.000 | up |
| G030745 | SOS1 | -0.958 | 0.000 | -1.000 | up |
| G030745 | EXOC7 | -0.946 | 0.000 | -1.000 | up |
| G030745 | CHRNB3 | -0.950 | 0.000 | -1.000 | up |
| G030745 | CHRNA3 | -0.906 | 0.002 | -1.000 | up |
| AK024936 | OLFM1 | -0.946 | 0.000 | -1.000 | up |
| XLOC_003048 | PTCH1 | -0.928 | 0.001 | -1.000 | up |
| XLOC_003048 | BTBD9 | -0.959 | 0.000 | -1.000 | up |
| XLOC_006353 | NPY2R | -0.907 | 0.002 | -1.000 | up |
| G061888 | CRELD1 | -0.976 | 0.000 | -1.000 | up |
| G061888 | FILIP1 | -0.904 | 0.002 | -1.000 | up |
| XLOC_009429 | DNAJB1 | -0.923 | 0.001 | -1.000 | up |
| RP11-461K13.1 | ERC2 | -0.924 | 0.001 | -1.000 | up |
| RP11-461K13.1 | CALB2 | -0.906 | 0.002 | -1.000 | up |
| RP11-461K13.1 | DPYSL5 | -0.915 | 0.001 | -1.000 | up |
| RP11-461K13.1 | DPYSL3 | -0.921 | 0.001 | -1.000 | up |
| G041587 | RPS6KA5 | -0.949 | 0.000 | -1.000 | up |
| G041587 | DPYSL3 | -0.920 | 0.001 | -1.000 | up |
| G059595 | TENM4 | -0.910 | 0.002 | -1.000 | up |
| G059595 | NRP2 | -0.957 | 0.000 | -1.000 | up |
| G059595 | DNAJB1 | -0.914 | 0.001 | -1.000 | up |
| G059595 | SPTB | -0.908 | 0.002 | -1.000 | up |
| G059595 | SEPSECS | -0.917 | 0.001 | -1.000 | up |
| SOS1 | G075352 | -0.908 | 0.002 | -1.000 | up |
| SOS1 | AL832615 | -0.909 | 0.002 | -1.000 | up |
| SOS1 | Z69720.2 | -0.902 | 0.002 | -1.000 | up |
| SOS1 | XLOC_006485 | -0.910 | 0.002 | -1.000 | up |
| SOS1 | G023385 | -0.923 | 0.001 | -1.000 | up |
| SOS1 | AC004054.1 | -0.902 | 0.002 | -1.000 | up |
| SOS1 | G084746 | -0.968 | 0.000 | -1.000 | up |
| SOS1 | G030745 | -0.958 | 0.000 | -1.000 | up |
| SOS1 | RP11-277L2.3 | 0.904 | 0.002 | 1.000 | up |
| SOS1 | WWC2-AS2 | 0.911 | 0.002 | 1.000 | up |
| SOS1 | AC006273.4 | 0.928 | 0.001 | 1.000 | up |
| SOS1 | G046707 | 0.901 | 0.002 | 1.000 | up |
| SOS1 | CTD-2527I21.7 | 0.922 | 0.001 | 1.000 | up |
| SOS1 | CTD-2382E5.2 | 0.960 | 0.000 | 1.000 | up |
| SOS1 | G081642 | 0.925 | 0.001 | 1.000 | up |
| SOS1 | AK125078 | 0.919 | 0.001 | 1.000 | up |
| SOS1 | H1FX-AS1 | 0.908 | 0.002 | 1.000 | up |
| SOS1 | RP11-820L6.1 | 0.926 | 0.001 | 1.000 | up |
| TENM4 | G020731 | -0.930 | 0.001 | -1.000 | up |
| TENM4 | G073552 | -0.978 | 0.000 | -1.000 | up |
| TENM4 | G046678 | -0.936 | 0.001 | -1.000 | up |
| TENM4 | AC016682.1 | -0.967 | 0.000 | -1.000 | up |
| TENM4 | RP11-484O2.1 | -0.921 | 0.001 | -1.000 | up |
| TENM4 | G050702 | -0.952 | 0.000 | -1.000 | up |
| TENM4 | G063995 | -0.959 | 0.000 | -1.000 | up |
| TENM4 | ZFAT-AS1 | -0.905 | 0.002 | -1.000 | up |
| TENM4 | G087265 | -0.944 | 0.000 | -1.000 | up |
| TENM4 | AK093193 | -0.905 | 0.002 | -1.000 | up |
| TENM4 | LOC101927164 | -0.911 | 0.002 | -1.000 | up |
| TENM4 | RP11-424I19.2 | -0.915 | 0.001 | -1.000 | up |
| TENM4 | G060178 | -0.906 | 0.002 | -1.000 | up |

| | | | | | |
|---|---|---|---|---|---|
| TENM4 | BX648501 | -0.926 | 0.001 | -1.000 | up |
| TENM4 | G060010 | -0.906 | 0.002 | -1.000 | up |
| TENM4 | CYP2D7 | -0.947 | 0.000 | -1.000 | up |
| TENM4 | uc.50 | -0.932 | 0.001 | -1.000 | up |
| TENM4 | RP4-760C5.3 | -0.935 | 0.001 | -1.000 | up |
| TENM4 | HM13-IT1 | -0.914 | 0.002 | -1.000 | up |
| TENM4 | G068890 | -0.928 | 0.001 | -1.000 | up |
| TENM4 | G051433 | -0.903 | 0.002 | -1.000 | up |
| TENM4 | G059595 | -0.910 | 0.002 | -1.000 | up |
| TENM4 | FAM74A4 | 0.929 | 0.001 | 1.000 | up |
| TENM4 | RP11-744I24.3 | 0.925 | 0.001 | 1.000 | up |
| TENM4 | GSE61474_XLOC_054139 | 0.980 | 0.000 | 1.000 | up |
| TENM4 | G017718 | 0.942 | 0.000 | 1.000 | up |
| TENM4 | G021652 | 0.907 | 0.002 | 1.000 | up |
| TENM4 | G002862 | 0.930 | 0.001 | 1.000 | up |
| TENM4 | uc.446 | 0.916 | 0.001 | 1.000 | up |
| TENM4 | RP11-128P10.1 | 0.929 | 0.001 | 1.000 | up |
| TENM4 | G011337 | 0.909 | 0.002 | 1.000 | up |
| TENM4 | TUB-AS1 | 0.952 | 0.000 | 1.000 | up |
| TENM4 | XLOC_000308 | 0.905 | 0.002 | 1.000 | up |
| TENM4 | RP11-290L1.2 | 0.912 | 0.002 | 1.000 | up |
| TENM4 | IDH1-AS1 | 0.901 | 0.002 | 1.000 | up |
| TENM4 | KB-1592A4.14 | 0.930 | 0.001 | 1.000 | up |
| TENM4 | RP11-792A8.4 | 0.901 | 0.002 | 1.000 | up |
| TENM4 | AK000953 | 0.915 | 0.001 | 1.000 | up |
| TENM4 | G057325 | 0.905 | 0.002 | 1.000 | up |
| TENM4 | G063514 | 0.925 | 0.001 | 1.000 | up |
| TENM4 | RP11-95P2.3 | 0.925 | 0.001 | 1.000 | up |
| TENM4 | G005619 | 0.928 | 0.001 | 1.000 | up |
| PHB | GSE61474_XLOC_020343 | -0.963 | 0.000 | -1.000 | up |
| PHB | AC074363.1 | -0.904 | 0.002 | -1.000 | up |
| PHB | RP11-219C20.3 | -0.906 | 0.002 | -1.000 | up |
| PHB | AC107218.3 | -0.902 | 0.002 | -1.000 | up |
| PHB | G036156 | -0.934 | 0.001 | -1.000 | up |
| PHB | G088912 | -0.903 | 0.002 | -1.000 | up |
| PHB | G079009 | 0.914 | 0.002 | 1.000 | up |
| PHB | FAM66A | 0.921 | 0.001 | 1.000 | up |
| PHB | RP11-475I24.3 | 0.922 | 0.001 | 1.000 | up |
| PHB | G061579 | 0.914 | 0.001 | 1.000 | up |
| PHB | G031140 | 0.914 | 0.002 | 1.000 | up |
| PHB | G070622 | 0.911 | 0.002 | 1.000 | up |
| PHB | RP11-355F22.1 | 0.903 | 0.002 | 1.000 | up |
| EXOC7 | RN7SL417P | -0.954 | 0.000 | -1.000 | up |
| EXOC7 | G056910 | -0.915 | 0.001 | -1.000 | up |
| EXOC7 | RP11-21M24.3 | -0.945 | 0.000 | -1.000 | up |
| EXOC7 | IGBP1-AS2 | -0.943 | 0.000 | -1.000 | up |
| EXOC7 | G009235 | -0.946 | 0.000 | -1.000 | up |
| EXOC7 | G087265 | -0.922 | 0.001 | -1.000 | up |
| EXOC7 | G053959 | -0.930 | 0.001 | -1.000 | up |
| EXOC7 | BX648501 | -0.948 | 0.000 | -1.000 | up |
| EXOC7 | RP11-768G7.2 | -0.909 | 0.002 | -1.000 | up |
| EXOC7 | HM13-IT1 | -0.914 | 0.002 | -1.000 | up |
| EXOC7 | G068890 | -0.916 | 0.001 | -1.000 | up |
| EXOC7 | AK022848 | -0.958 | 0.000 | -1.000 | up |
| EXOC7 | G030745 | -0.946 | 0.000 | -1.000 | up |
| EXOC7 | FAM230C | 0.911 | 0.002 | 1.000 | up |
| EXOC7 | XLOC_002675 | 0.900 | 0.002 | 1.000 | up |
| EXOC7 | G002862 | 0.912 | 0.002 | 1.000 | up |
| EXOC7 | SCAMP1-AS1 | 0.949 | 0.000 | 1.000 | up |
| EXOC7 | DLG5-AS1 (2) | 0.906 | 0.002 | 1.000 | up |
| EXOC7 | BC112270 | 0.912 | 0.002 | 1.000 | up |
| EXOC7 | AC114803.3 | 0.915 | 0.001 | 1.000 | up |
| EXOC7 | RP11-872D17.4 | 0.900 | 0.002 | 1.000 | up |

| | | | | | |
|---|---|---|---|---|---|
| EXOC7 | XLOC_005289 | 0.930 | 0.001 | 1.000 | up |
| EXOC7 | G081642 | 0.951 | 0.000 | 1.000 | up |
| EXOC7 | CTC-350I8.1 | 0.920 | 0.001 | 1.000 | up |
| EXOC7 | XLOC_010219 | 0.907 | 0.002 | 1.000 | up |
| EXOC7 | RP11-168O10.6 | 0.922 | 0.001 | 1.000 | up |
| EXOC7 | XLOC_004894 | 0.937 | 0.001 | 1.000 | up |
| UNC13C | uc.247 | 0.942 | 0.000 | 1.000 | up |
| CRELD1 | RP1-296L11.1 | -0.966 | 0.000 | -1.000 | up |
| CRELD1 | LOC100499484-C9ORF174 | -0.900 | 0.002 | -1.000 | up |
| CRELD1 | SLC7A11-AS1 | -0.931 | 0.001 | -1.000 | up |
| CRELD1 | ITGA9-AS1 | -0.939 | 0.001 | -1.000 | up |
| CRELD1 | G060633 | -0.906 | 0.002 | -1.000 | up |
| CRELD1 | XLOC_001223 | -0.964 | 0.000 | -1.000 | up |
| CRELD1 | RP11-466A19.1 | -0.944 | 0.000 | -1.000 | up |
| CRELD1 | XLOC_008025 | -0.922 | 0.001 | -1.000 | up |
| CRELD1 | G072062 | -0.970 | 0.000 | -1.000 | up |
| CRELD1 | XLOC_009306 | -0.920 | 0.001 | -1.000 | up |
| CRELD1 | EML2-AS1 | -0.913 | 0.002 | -1.000 | up |
| CRELD1 | G030124 | -0.916 | 0.001 | -1.000 | up |
| CRELD1 | G003431 | -0.940 | 0.001 | -1.000 | up |
| CRELD1 | XLOC_012740 | -0.911 | 0.002 | -1.000 | up |
| CRELD1 | ELOVL2-AS1 | -0.919 | 0.001 | -1.000 | up |
| CRELD1 | LOC100506834 | -0.966 | 0.000 | -1.000 | up |
| CRELD1 | EF565122 | -0.916 | 0.001 | -1.000 | up |
| CRELD1 | G061888 | -0.976 | 0.000 | -1.000 | up |
| CRELD1 | AK055712 | 0.914 | 0.002 | 1.000 | up |
| CRELD1 | RP11-303E16.7 | 0.934 | 0.001 | 1.000 | up |
| CRELD1 | CTD-2023N9.3 | 0.905 | 0.002 | 1.000 | up |
| CRELD1 | RP11-686O6.2 | 0.929 | 0.001 | 1.000 | up |
| CRELD1 | RP11-356J5.12 | 0.916 | 0.001 | 1.000 | up |
| CRELD1 | RP1-40E16.12 | 0.925 | 0.001 | 1.000 | up |
| CRELD1 | GBAT2 | 0.918 | 0.001 | 1.000 | up |
| CRELD1 | RP11- | 0.914 | 0.002 | 1.000 | up |
| CRELD1 | RP11-297J22.1 | 0.923 | 0.001 | 1.000 | up |
| CRELD1 | RP11-179B2.2 | 0.900 | 0.002 | 1.000 | up |
| CRELD1 | G047225 | 0.912 | 0.002 | 1.000 | up |
| CRELD1 | RP11-191L17.1 | 0.936 | 0.001 | 1.000 | up |
| CRELD1 | G018162 | 0.912 | 0.002 | 1.000 | up |
| CRELD1 | RP11-285F16.1 | 0.902 | 0.002 | 1.000 | up |
| CRELD1 | RP5-1092A3.4 | 0.932 | 0.001 | 1.000 | up |
| CRELD1 | RP11-351D16.3 | 0.918 | 0.001 | 1.000 | up |
| CRELD1 | RP4-683L5.1 | 0.938 | 0.001 | 1.000 | up |
| CRELD1 | RP11-353K11.1 | 0.958 | 0.000 | 1.000 | up |
| CRELD1 | RP11-209D14.2 | 0.905 | 0.002 | 1.000 | up |
| CRELD1 | AC002467.7 | 0.932 | 0.001 | 1.000 | up |
| CRELD1 | G015380 | 0.930 | 0.001 | 1.000 | up |
| CRELD1 | XLOC_001189 | 0.913 | 0.002 | 1.000 | up |
| CRELD1 | XLOC_006554 | 0.927 | 0.001 | 1.000 | up |
| CRELD1 | AC144652.1 | 0.910 | 0.002 | 1.000 | up |
| CRELD1 | G085410 | 0.960 | 0.000 | 1.000 | up |
| CRELD1 | G089005 | 0.987 | 0.000 | 1.000 | up |
| CRELD1 | LINC00408 | 0.904 | 0.002 | 1.000 | up |
| CRELD1 | BC070391 | 0.934 | 0.001 | 1.000 | up |
| CRELD1 | G034470 | 0.956 | 0.000 | 1.000 | up |
| CRELD1 | G073576 | 0.944 | 0.000 | 1.000 | up |
| CRELD1 | XLOC_013222 | 0.908 | 0.002 | 1.000 | up |
| CRELD1 | XLOC_011052 | 0.911 | 0.002 | 1.000 | up |
| CRELD1 | TNRC18P1 | 0.939 | 0.001 | 1.000 | up |
| CRELD1 | RP11-138E2.1 | 0.945 | 0.000 | 1.000 | up |
| CRELD1 | G059716 | 0.940 | 0.001 | 1.000 | up |
| CRELD1 | G044581 | 0.968 | 0.000 | 1.000 | up |
| CRELD1 | PVRL3-AS1 | 0.924 | 0.001 | 1.000 | up |
| CRELD1 | BC042589 | 0.957 | 0.000 | 1.000 | up |
| CRELD1 | LINC00634 | 0.955 | 0.000 | 1.000 | up |

| | | | | | |
|---|---|---|---|---|---|
| CRELD1 | RP11-383C5.5 | 0.916 | 0.001 | 1.000 | up |
| CRELD1 | G001739 | 0.925 | 0.001 | 1.000 | up |
| CRELD1 | RP11-22N19.2 | 0.938 | 0.001 | 1.000 | up |
| CRELD1 | RP4-555D20.3 | 0.956 | 0.000 | 1.000 | up |
| CRELD1 | RP11-507K2.3 | 0.902 | 0.002 | 1.000 | up |
| CRELD1 | TOB1-AS1 (1) | 0.948 | 0.000 | 1.000 | up |
| CRELD1 | RP4-798A10.7 | 0.928 | 0.001 | 1.000 | up |
| CRELD1 | RP1-223E5.4 | 0.950 | 0.000 | 1.000 | up |
| CRELD1 | RP11-573G6.8 | 0.916 | 0.001 | 1.000 | up |
| CRELD1 | LL09NC01-139C3.1 | 0.953 | 0.000 | 1.000 | up |
| CRELD1 | RP1-86C11.7 | 0.954 | 0.000 | 1.000 | up |
| CRELD1 | FLJ20021 | 0.949 | 0.000 | 1.000 | up |
| CRELD1 | RP11-109D9.4 | 0.906 | 0.002 | 1.000 | up |
| CRELD1 | CTC- | 0.976 | 0.000 | 1.000 | up |
| CRELD1 | RP11-61L19.2 | 0.925 | 0.001 | 1.000 | up |
| CRELD1 | RP11-85A1.3 | 0.918 | 0.001 | 1.000 | up |
| CRELD1 | RP11-344P13.6 | 0.972 | 0.000 | 1.000 | up |
| CRELD1 | G070405 | 0.922 | 0.001 | 1.000 | up |
| CRELD1 | ANKRD20A12 | 0.919 | 0.001 | 1.000 | up |
| CRELD1 | G014106 | 0.907 | 0.002 | 1.000 | up |
| CRELD1 | G002695 | 0.944 | 0.000 | 1.000 | up |
| CRELD1 | G006578 | 0.940 | 0.001 | 1.000 | up |
| CRELD1 | LOC101928489 | 0.944 | 0.000 | 1.000 | up |
| CRELD1 | G075319 | 0.936 | 0.001 | 1.000 | up |
| CRELD1 | RP11-820I16.4 | 0.909 | 0.002 | 1.000 | up |
| CRELD1 | RP11-79P5.9 | 0.928 | 0.001 | 1.000 | up |
| CRELD1 | RP11-166B2.3 | 0.936 | 0.001 | 1.000 | up |
| CRELD1 | LOC100270746 | 0.918 | 0.001 | 1.000 | up |
| CRELD1 | LINC01604 | 0.903 | 0.002 | 1.000 | up |
| CRELD1 | RP11-439C15.4 | 0.957 | 0.000 | 1.000 | up |
| CRELD1 | RP11-379F4.9 | 0.917 | 0.001 | 1.000 | up |
| ARHGAP32 | XLOC_001223 | -0.907 | 0.002 | -1.000 | up |
| ARHGAP32 | AP000438.2 | -0.904 | 0.002 | -1.000 | up |
| ARHGAP32 | XLOC_008025 | -0.927 | 0.001 | -1.000 | up |
| ARHGAP32 | G072062 | -0.905 | 0.002 | -1.000 | up |
| ARHGAP32 | G045594 | -0.962 | 0.000 | -1.000 | up |
| ARHGAP32 | XLOC_009306 | -0.929 | 0.001 | -1.000 | up |
| ARHGAP32 | G072364 | -0.967 | 0.000 | -1.000 | up |
| ARHGAP32 | G020793 | -0.994 | 0.000 | -1.000 | up |
| ARHGAP32 | LINC01149 | -0.910 | 0.002 | -1.000 | up |
| ARHGAP32 | G015181 | -0.957 | 0.000 | -1.000 | up |
| ARHGAP32 | G051586 | -0.911 | 0.002 | -1.000 | up |
| ARHGAP32 | LOC100506834 | -0.925 | 0.001 | -1.000 | up |
| ARHGAP32 | CTD- | 0.938 | 0.001 | 1.000 | up |
| ARHGAP32 | RP11-350N15.5 | 0.903 | 0.002 | 1.000 | up |
| ARHGAP32 | G061591 | 0.914 | 0.002 | 1.000 | up |
| ARHGAP32 | ZFX-AS1 | 0.915 | 0.001 | 1.000 | up |
| ARHGAP32 | RP11-179B2.2 | 0.911 | 0.002 | 1.000 | up |
| ARHGAP32 | XLOC_008875 | 0.917 | 0.001 | 1.000 | up |
| ARHGAP32 | RP5-1092A3.4 | 0.902 | 0.002 | 1.000 | up |
| ARHGAP32 | RP11-353K11.1 | 0.931 | 0.001 | 1.000 | up |
| ARHGAP32 | TAF1A-AS1 | 0.950 | 0.000 | 1.000 | up |
| ARHGAP32 | XLOC_006554 | 0.961 | 0.000 | 1.000 | up |
| ARHGAP32 | G085410 | 0.918 | 0.001 | 1.000 | up |
| ARHGAP32 | G048385 | 0.940 | 0.001 | 1.000 | up |
| ARHGAP32 | RP11-496H1.2 | 0.908 | 0.002 | 1.000 | up |
| ARHGAP32 | TCR-alpha | 0.901 | 0.002 | 1.000 | up |
| ARHGAP32 | SIX3-AS1 | 0.914 | 0.001 | 1.000 | up |
| ARHGAP32 | TNRC18P1 | 0.940 | 0.001 | 1.000 | up |
| ARHGAP32 | uc.76 | 0.952 | 0.000 | 1.000 | up |
| ARHGAP32 | G010838 | 0.937 | 0.001 | 1.000 | up |
| ARHGAP32 | RP11-573G6.8 | 0.910 | 0.002 | 1.000 | up |
| ARHGAP32 | LL09NC01-139C3.1 | 0.904 | 0.002 | 1.000 | up |

| | | | | | |
|---|---|---|---|---|---|
| ARHGAP32 | LINC00449 | 0.933 | 0.001 | 1.000 | up |
| ARHGAP32 | LOC101929066 | 0.945 | 0.000 | 1.000 | up |
| ARHGAP32 | DHRS4-AS1 | 0.931 | 0.001 | 1.000 | up |
| ARHGAP32 | RP11-483I13.5 | 0.905 | 0.002 | 1.000 | up |
| ARHGAP32 | RP6-109B7.4 | 0.926 | 0.001 | 1.000 | up |
| ARHGAP32 | LINC01604 | 0.914 | 0.002 | 1.000 | up |
| ARHGAP32 | RP11-379F4.9 | 0.942 | 0.000 | 1.000 | up |
| HSPA8 | ASAP1-IT1 | -0.901 | 0.002 | -1.000 | up |
| HSPA8 | G009303 | -0.904 | 0.002 | -1.000 | up |
| HSPA8 | G031832 | -0.901 | 0.002 | -1.000 | up |
| HSPA8 | RP11-669E14.6 | -0.908 | 0.002 | -1.000 | up |
| HSPA8 | G071944 | -0.922 | 0.001 | -1.000 | up |
| HSPA8 | uc.474 | -0.928 | 0.001 | -1.000 | up |
| HSPA8 | G053358 | -0.934 | 0.001 | -1.000 | up |
| HSPA8 | SH3GL1P2 | 0.960 | 0.000 | 1.000 | up |
| HSPA8 | RP11-396C23.4 | 0.932 | 0.001 | 1.000 | up |
| HSPA8 | G047031 | 0.961 | 0.000 | 1.000 | up |
| HSPA8 | G082153 | 0.903 | 0.002 | 1.000 | up |
| HSPA8 | G041225 | 0.931 | 0.001 | 1.000 | up |
| HSPA8 | CCNT2-AS1 | 0.904 | 0.002 | 1.000 | up |
| HSPA8 | LOC101927822 | 0.906 | 0.002 | 1.000 | up |
| HSPA8 | uc.412 | 0.912 | 0.002 | 1.000 | up |
| HSPA8 | RP11-362L22.1 | 0.912 | 0.002 | 1.000 | up |
| HSPA8 | G061238 | 0.911 | 0.002 | 1.000 | up |
| HSPA8 | G057611 | 0.905 | 0.002 | 1.000 | up |
| MCTP2 | XLOC_012579 | -0.965 | 0.000 | -1.000 | up |
| MCTP2 | G067669 | -0.928 | 0.001 | -1.000 | up |
| MCTP2 | RP11-10J21.4 | 0.939 | 0.001 | 1.000 | up |
| MCTP2 | RP11-21M24.2 | 0.974 | 0.000 | 1.000 | up |
| MCTP2 | G086828 | 0.910 | 0.002 | 1.000 | up |
| MCTP2 | CTD-2517M22.16 | 0.914 | 0.002 | 1.000 | up |
| MCTP2 | G066606 | 0.910 | 0.002 | 1.000 | up |
| MCTP2 | ULK4P1 | 0.937 | 0.001 | 1.000 | up |
| MCTP2 | G036568 | 0.946 | 0.000 | 1.000 | up |
| MCTP2 | G064611 | 0.906 | 0.002 | 1.000 | up |
| MCTP2 | XLOC_010219 | 0.954 | 0.000 | 1.000 | up |
| MCTP2 | G022520 | 0.910 | 0.002 | 1.000 | up |
| CHRNB3 | RP11-308N19.1 | -0.912 | 0.002 | -1.000 | up |
| CHRNB3 | RP11-21M24.3 | -0.921 | 0.001 | -1.000 | up |
| CHRNB3 | RP11-187C18.2 | -0.907 | 0.002 | -1.000 | up |
| CHRNB3 | CTB-102L5.8 | -0.914 | 0.002 | -1.000 | up |
| CHRNB3 | G043880 | -0.929 | 0.001 | -1.000 | up |
| CHRNB3 | XLOC_006485 | -0.923 | 0.001 | -1.000 | up |
| CHRNB3 | G030745 | -0.950 | 0.000 | -1.000 | up |
| CHRNB3 | FAM230C | 0.929 | 0.001 | 1.000 | up |
| CHRNB3 | G077330 | 0.915 | 0.001 | 1.000 | up |
| CHRNB3 | AC006273.4 | 0.970 | 0.000 | 1.000 | up |
| CHRNB3 | G006537 | 0.956 | 0.000 | 1.000 | up |
| CHRNB3 | CTD-2527I21.7 | 0.946 | 0.000 | 1.000 | up |
| CHRNB3 | CTD-2382E5.2 | 0.938 | 0.001 | 1.000 | up |
| CHRNB3 | RP11-254F7.1 | 0.905 | 0.002 | 1.000 | up |
| CHRNB3 | H1FX-AS1 | 0.904 | 0.002 | 1.000 | up |
| AP1G2 | G073552 | -0.917 | 0.001 | -1.000 | up |
| AP1G2 | AC016682.1 | -0.943 | 0.000 | -1.000 | up |
| AP1G2 | RP11-484O2.1 | -0.926 | 0.001 | -1.000 | up |
| AP1G2 | G009235 | -0.916 | 0.001 | -1.000 | up |
| AP1G2 | ZFAT-AS1 | -0.951 | 0.000 | -1.000 | up |
| AP1G2 | G087265 | -0.903 | 0.002 | -1.000 | up |
| AP1G2 | BX648501 | -0.930 | 0.001 | -1.000 | up |
| AP1G2 | RP4-760C5.3 | -0.952 | 0.000 | -1.000 | up |
| AP1G2 | BCRP3 | -0.900 | 0.002 | -1.000 | up |
| AP1G2 | G068890 | -0.966 | 0.000 | -1.000 | up |
| AP1G2 | AK022848 | -0.924 | 0.001 | -1.000 | up |
| AP1G2 | FAM74A4 | 0.903 | 0.002 | 1.000 | up |

| | | | | | |
|---|---|---|---|---|---|
| AP1G2 | USP27X-AS1 | 0.914 | 0.001 | 1.000 | up |
| AP1G2 | G002862 | 0.969 | 0.000 | 1.000 | up |
| AP1G2 | RP11-128P10.1 | 0.924 | 0.001 | 1.000 | up |
| AP1G2 | GYG2P1 | 0.928 | 0.001 | 1.000 | up |
| AP1G2 | FGF13-AS1 | 0.914 | 0.001 | 1.000 | up |
| AP1G2 | PRR36 | 0.907 | 0.002 | 1.000 | up |
| AP1G2 | G046707 | 0.904 | 0.002 | 1.000 | up |
| AP1G2 | CRYBB2P1 | 0.917 | 0.001 | 1.000 | up |
| AP1G2 | G063514 | 0.927 | 0.001 | 1.000 | up |
| AP1G2 | RP11-168O10.6 | 0.926 | 0.001 | 1.000 | up |
| HSP90AA1 | LOC100499484-C9ORF174 | -0.922 | 0.001 | -1.000 | up |
| HSP90AA1 | uc.172 | -0.912 | 0.002 | -1.000 | up |
| HSP90AA1 | G008916 | -0.929 | 0.001 | -1.000 | up |
| HSP90AA1 | G086585 | -0.907 | 0.002 | -1.000 | up |
| HSP90AA1 | AC074363.1 | -0.914 | 0.002 | -1.000 | up |
| HSP90AA1 | RP11-737O24.3 | -0.960 | 0.000 | -1.000 | up |
| HSP90AA1 | RN7SKP264 | -0.956 | 0.000 | -1.000 | up |
| HSP90AA1 | RP11-219C20.3 | -0.969 | 0.000 | -1.000 | up |
| HSP90AA1 | Z69720.2 | -0.910 | 0.002 | -1.000 | up |
| HSP90AA1 | BC034456 | -0.911 | 0.002 | -1.000 | up |
| HSP90AA1 | G036156 | -0.908 | 0.002 | -1.000 | up |
| HSP90AA1 | G030124 | -0.906 | 0.002 | -1.000 | up |
| HSP90AA1 | G082713 | -0.901 | 0.002 | -1.000 | up |
| HSP90AA1 | CTC-250I14.3 | -0.914 | 0.001 | -1.000 | up |
| HSP90AA1 | DLG3-AS1 | -0.914 | 0.001 | -1.000 | up |
| HSP90AA1 | G088912 | -0.927 | 0.001 | -1.000 | up |
| HSP90AA1 | CTD-2015A6.1 | -0.916 | 0.001 | -1.000 | up |
| HSP90AA1 | LOC101929427 | -0.903 | 0.002 | -1.000 | up |
| HSP90AA1 | GBAT2 | 0.902 | 0.002 | 1.000 | up |
| HSP90AA1 | G066319 | 0.966 | 0.000 | 1.000 | up |
| HSP90AA1 | RP11-204E9.1 | 0.927 | 0.001 | 1.000 | up |
| HSP90AA1 | G085410 | 0.907 | 0.002 | 1.000 | up |
| HSP90AA1 | LINC00408 | 0.908 | 0.002 | 1.000 | up |
| HSP90AA1 | RP11-792A8.4 | 0.903 | 0.002 | 1.000 | up |
| HSP90AA1 | G063668 | 0.918 | 0.001 | 1.000 | up |
| HSP90AA1 | BC042589 | 0.900 | 0.002 | 1.000 | up |
| HSP90AA1 | G049105 | 0.905 | 0.002 | 1.000 | up |
| HSP90AA1 | ENO1-AS1 | 0.948 | 0.000 | 1.000 | up |
| HSP90AA1 | DGCR5 | 0.944 | 0.000 | 1.000 | up |
| HSP90AA1 | G027543 | 0.941 | 0.000 | 1.000 | up |
| HSP90AA1 | ANKRD20A12 | 0.917 | 0.001 | 1.000 | up |
| HSP90AA1 | LOC100270746 | 0.908 | 0.002 | 1.000 | up |
| ADGRL2 | ITGA9-AS1 | -0.906 | 0.002 | -1.000 | up |
| ADGRL2 | RP11-3K16.2 | -0.911 | 0.002 | -1.000 | up |
| ADGRL2 | CTD-2023M8.1 | -0.908 | 0.002 | -1.000 | up |
| ADGRL2 | G030124 | -0.981 | 0.000 | -1.000 | up |
| ADGRL2 | CTC-250I14.3 | -0.903 | 0.002 | -1.000 | up |
| ADGRL2 | G029241 | -0.925 | 0.001 | -1.000 | up |
| ADGRL2 | G088912 | -0.905 | 0.002 | -1.000 | up |
| ADGRL2 | G010224 | -0.920 | 0.001 | -1.000 | up |
| ADGRL2 | G014329 | 0.901 | 0.002 | 1.000 | up |
| ADGRL2 | G066319 | 0.922 | 0.001 | 1.000 | up |
| ADGRL2 | RP11-475I24.3 | 0.902 | 0.002 | 1.000 | up |
| ADGRL2 | G085410 | 0.914 | 0.001 | 1.000 | up |
| ADGRL2 | LINC00408 | 0.924 | 0.001 | 1.000 | up |
| ADGRL2 | G053697 | 0.922 | 0.001 | 1.000 | up |
| ADGRL2 | G010782 | 0.935 | 0.001 | 1.000 | up |
| ADGRL2 | RP11- | 0.902 | 0.002 | 1.000 | up |
| ADGRL2 | EZR-AS1 | 0.917 | 0.001 | 1.000 | up |
| ADGRL2 | ANKRD20A12 | 0.915 | 0.001 | 1.000 | up |
| ADGRL2 | G014106 | 0.937 | 0.001 | 1.000 | up |
| NRP2 | IGBP1-AS2 | -0.943 | 0.000 | -1.000 | up |
| NRP2 | ASAP1-IT1 | -0.914 | 0.001 | -1.000 | up |
| NRP2 | RP11-187C18.2 | -0.927 | 0.001 | -1.000 | up |

| | | | | | |
|---|---|---|---|---|---|
| NRP2 | G049993 | -0.930 | 0.001 | -1.000 | up |
| NRP2 | G046678 | -0.986 | 0.000 | -1.000 | up |
| NRP2 | AC016682.1 | -0.909 | 0.002 | -1.000 | up |
| NRP2 | RP11-484O2.1 | -0.932 | 0.001 | -1.000 | up |
| NRP2 | G050702 | -0.942 | 0.000 | -1.000 | up |
| NRP2 | G063995 | -0.934 | 0.001 | -1.000 | up |
| NRP2 | G087265 | -0.920 | 0.001 | -1.000 | up |
| NRP2 | G047392 | -0.901 | 0.002 | -1.000 | up |
| NRP2 | LOC101927164 | -0.921 | 0.001 | -1.000 | up |
| NRP2 | RP1-124C6.1 | -0.924 | 0.001 | -1.000 | up |
| NRP2 | RP11-166B2.5 | -0.906 | 0.002 | -1.000 | up |
| NRP2 | XLOC_003837 | -0.940 | 0.001 | -1.000 | up |
| NRP2 | BX648501 | -0.910 | 0.002 | -1.000 | up |
| NRP2 | LINC00930 | -0.913 | 0.002 | -1.000 | up |
| NRP2 | G052905 | -0.941 | 0.000 | -1.000 | up |
| NRP2 | BCRP3 | -0.949 | 0.000 | -1.000 | up |
| NRP2 | G051433 | -0.922 | 0.001 | -1.000 | up |
| NRP2 | G059595 | -0.957 | 0.000 | -1.000 | up |
| NRP2 | RP11-255E6.6 | 0.925 | 0.001 | 1.000 | up |
| NRP2 | GSE61474_XLOC_054139 | 0.925 | 0.001 | 1.000 | up |
| NRP2 | RP11-44N22.3 | 0.928 | 0.001 | 1.000 | up |
| NRP2 | FAM230C | 0.944 | 0.000 | 1.000 | up |
| NRP2 | G021652 | 0.909 | 0.002 | 1.000 | up |
| NRP2 | RP11-715J22.6 | 0.907 | 0.002 | 1.000 | up |
| NRP2 | AC093391.2 | 0.902 | 0.002 | 1.000 | up |
| NRP2 | RP11-128P10.1 | 0.947 | 0.000 | 1.000 | up |
| NRP2 | TUB-AS1 | 0.913 | 0.002 | 1.000 | up |
| NRP2 | IDH1-AS1 | 0.917 | 0.001 | 1.000 | up |
| NRP2 | RP11- | 0.918 | 0.001 | 1.000 | up |
| NRP2 | BX248253 | 0.951 | 0.000 | 1.000 | up |
| NRP2 | G063514 | 0.919 | 0.001 | 1.000 | up |
| NRP2 | RP11-118K6.3 | 0.909 | 0.002 | 1.000 | up |
| NRP2 | G005619 | 0.942 | 0.000 | 1.000 | up |
| NRP2 | G015529 | 0.904 | 0.002 | 1.000 | up |
| ELFN1 | BC033330 | -0.930 | 0.001 | -1.000 | up |
| ELFN1 | G080487 | -0.963 | 0.000 | -1.000 | up |
| ELFN1 | G032508 | -0.948 | 0.000 | -1.000 | up |
| ELFN1 | RP11-551L14.6 | -0.949 | 0.000 | -1.000 | up |
| ELFN1 | G047225 | 0.922 | 0.001 | 1.000 | up |
| ELFN1 | G015380 | 0.904 | 0.002 | 1.000 | up |
| ELFN1 | XLOC_001189 | 0.942 | 0.000 | 1.000 | up |
| ELFN1 | LOC101929741 | 0.928 | 0.001 | 1.000 | up |
| ELFN1 | G081251 | 0.930 | 0.001 | 1.000 | up |
| ELFN1 | LINC-PINT (2) | 0.902 | 0.002 | 1.000 | up |
| ELFN1 | AK026244 | 0.902 | 0.002 | 1.000 | up |
| ELFN1 | RP11-376P6.3 | 0.917 | 0.001 | 1.000 | up |
| ELFN1 | RP11-383C5.5 | 0.907 | 0.002 | 1.000 | up |
| ELFN1 | G001739 | 0.909 | 0.002 | 1.000 | up |
| ELFN1 | LINC01023 | 0.909 | 0.002 | 1.000 | up |
| ELFN1 | LOC101928417 | 0.914 | 0.002 | 1.000 | up |
| ELFN1 | RP11-107N15.1 | 0.947 | 0.000 | 1.000 | up |
| ELFN1 | RP11-61L19.2 | 0.909 | 0.002 | 1.000 | up |
| ELFN1 | G070405 | 0.914 | 0.002 | 1.000 | up |
| ELFN1 | G002695 | 0.917 | 0.001 | 1.000 | up |
| ELFN1 | G075703 | 0.905 | 0.002 | 1.000 | up |
| ELFN1 | CTD- | 0.902 | 0.002 | 1.000 | up |
| SIPA1L2 | RP1-296L11.1 | -0.937 | 0.001 | -1.000 | up |
| SIPA1L2 | CTA-217C2.2 | -0.924 | 0.001 | -1.000 | up |
| SIPA1L2 | RN7SKP264 | -0.939 | 0.001 | -1.000 | up |
| SIPA1L2 | G036779 | -0.964 | 0.000 | -1.000 | up |
| SIPA1L2 | AC003682.16 | -0.942 | 0.000 | -1.000 | up |
| SIPA1L2 | G046027 | -0.900 | 0.002 | -1.000 | up |
| SIPA1L2 | G030124 | -0.965 | 0.000 | -1.000 | up |
| SIPA1L2 | G003431 | -0.911 | 0.002 | -1.000 | up |

| | | | | | |
|---|---|---|---|---|---|
| SIPA1L2 | AC025442.3 | -0.909 | 0.002 | -1.000 | up |
| SIPA1L2 | G088912 | -0.912 | 0.002 | -1.000 | up |
| SIPA1L2 | EPHA5-AS1 | 0.909 | 0.002 | 1.000 | up |
| SIPA1L2 | G018162 | 0.903 | 0.002 | 1.000 | up |
| SIPA1L2 | G066319 | 0.923 | 0.001 | 1.000 | up |
| SIPA1L2 | LINC00408 | 0.944 | 0.000 | 1.000 | up |
| SIPA1L2 | RP11-680F8.1 | 0.929 | 0.001 | 1.000 | up |
| SIPA1L2 | RP11- | 0.934 | 0.001 | 1.000 | up |
| SIPA1L2 | AK125078 | 0.918 | 0.001 | 1.000 | up |
| SIPA1L2 | ALDOC | 0.906 | 0.002 | 1.000 | up |
| ACTN1 | NADK2-AS1 | -0.912 | 0.002 | -1.000 | up |
| ACTN1 | CTB-193M12.1 | -0.902 | 0.002 | -1.000 | up |
| ACTN1 | AK124233 | -0.905 | 0.002 | -1.000 | up |
| ACTN1 | G073552 | -0.924 | 0.001 | -1.000 | up |
| ACTN1 | G063995 | -0.936 | 0.001 | -1.000 | up |
| ACTN1 | G087265 | -0.907 | 0.002 | -1.000 | up |
| ACTN1 | G060010 | -0.923 | 0.001 | -1.000 | up |
| ACTN1 | CYP2D7 | -0.966 | 0.000 | -1.000 | up |
| ACTN1 | RP11-768G7.2 | -0.962 | 0.000 | -1.000 | up |
| ACTN1 | LOC642846 | -0.911 | 0.002 | -1.000 | up |
| ACTN1 | G020470 | -0.901 | 0.002 | -1.000 | up |
| ACTN1 | HM13-IT1 | -0.950 | 0.000 | -1.000 | up |
| ACTN1 | G090511 | 0.910 | 0.002 | 1.000 | up |
| ACTN1 | G017718 | 0.952 | 0.000 | 1.000 | up |
| ACTN1 | G021652 | 0.902 | 0.002 | 1.000 | up |
| ACTN1 | uc.446 | 0.982 | 0.000 | 1.000 | up |
| ACTN1 | TUB-AS1 | 0.955 | 0.000 | 1.000 | up |
| ACTN1 | XLOC_000308 | 0.971 | 0.000 | 1.000 | up |
| ACTN1 | G063668 | 0.922 | 0.001 | 1.000 | up |
| ACTN1 | AC114803.3 | 0.950 | 0.000 | 1.000 | up |
| ACTN1 | uc.95 | 0.909 | 0.002 | 1.000 | up |
| ACTN1 | G057325 | 0.959 | 0.000 | 1.000 | up |
| CHRNA3 | IGBP1-AS2 | -0.919 | 0.001 | -1.000 | up |
| CHRNA3 | AC074363.1 | -0.923 | 0.001 | -1.000 | up |
| CHRNA3 | RP11-737O24.3 | -0.961 | 0.000 | -1.000 | up |
| CHRNA3 | RP11-219C20.3 | -0.910 | 0.002 | -1.000 | up |
| CHRNA3 | G040015 | -0.916 | 0.001 | -1.000 | up |
| CHRNA3 | G087265 | -0.905 | 0.002 | -1.000 | up |
| CHRNA3 | Z69720.2 | -0.941 | 0.000 | -1.000 | up |
| CHRNA3 | G036156 | -0.921 | 0.001 | -1.000 | up |
| CHRNA3 | AK025321 | -0.923 | 0.001 | -1.000 | up |
| CHRNA3 | AC026188.1 | -0.914 | 0.001 | -1.000 | up |
| CHRNA3 | G010941 | -0.914 | 0.002 | -1.000 | up |
| CHRNA3 | DLG3-AS1 | -0.911 | 0.002 | -1.000 | up |
| CHRNA3 | G030745 | -0.906 | 0.002 | -1.000 | up |
| CHRNA3 | GSE61474_XLOC_054139 | 0.921 | 0.001 | 1.000 | up |
| CHRNA3 | RP11-44N22.3 | 0.904 | 0.002 | 1.000 | up |
| CHRNA3 | FAM230C | 0.907 | 0.002 | 1.000 | up |
| CHRNA3 | XLOC_002675 | 0.914 | 0.001 | 1.000 | up |
| CHRNA3 | RP11-502I4.3 | 0.934 | 0.001 | 1.000 | up |
| CHRNA3 | RP11-290L1.2 | 0.912 | 0.002 | 1.000 | up |
| CHRNA3 | ENO1-AS1 | 0.952 | 0.000 | 1.000 | up |
| CHRNA3 | G027543 | 0.902 | 0.002 | 1.000 | up |
| CHRNA3 | G061722 | 0.907 | 0.002 | 1.000 | up |
| CHRNA3 | CTD-2382E5.2 | 0.931 | 0.001 | 1.000 | up |
| CHRNA3 | G081642 | 0.924 | 0.001 | 1.000 | up |
| CHRNA3 | AK125078 | 0.902 | 0.002 | 1.000 | up |
| CHRNA3 | DNAJC9-AS1 | 0.947 | 0.000 | 1.000 | up |
| CHRNA3 | RP11-820L6.1 | 0.958 | 0.000 | 1.000 | up |
| CHRNA3 | XLOC_004894 | 0.907 | 0.002 | 1.000 | up |
| DAB1 | Alpha_250_dna | -0.920 | 0.001 | -1.000 | up |
| DAB1 | G011310 | -0.928 | 0.001 | -1.000 | up |
| DAB1 | ASAP1-IT1 | -0.953 | 0.000 | -1.000 | up |
| DAB1 | PHF2P2 | -0.905 | 0.002 | -1.000 | up |

| | | | | | |
|---|---|---|---|---|---|
| DAB1 | G081487 | -0.921 | 0.001 | -1.000 | up |
| DAB1 | AP001625.6 | -0.907 | 0.002 | -1.000 | up |
| DAB1 | XLOC_003405 | -0.930 | 0.001 | -1.000 | up |
| DAB1 | AADACL2- | -0.932 | 0.001 | -1.000 | up |
| DAB1 | RP11-669E14.6 | -0.987 | 0.000 | -1.000 | up |
| DAB1 | G087395 | -0.906 | 0.002 | -1.000 | up |
| DAB1 | XLOC_003837 | -0.918 | 0.001 | -1.000 | up |
| DAB1 | CTD-2023M8.1 | -0.942 | 0.000 | -1.000 | up |
| DAB1 | LOC101926908 | -0.976 | 0.000 | -1.000 | up |
| DAB1 | XLOC_012740 | -0.914 | 0.002 | -1.000 | up |
| DAB1 | uc.474 | -0.927 | 0.001 | -1.000 | up |
| DAB1 | MIMT1 | -0.918 | 0.001 | -1.000 | up |
| DAB1 | G010224 | -0.910 | 0.002 | -1.000 | up |
| DAB1 | TIPARP-AS1 | 0.928 | 0.001 | 1.000 | up |
| DAB1 | LINC00662 | 0.920 | 0.001 | 1.000 | up |
| DAB1 | G064655 | 0.912 | 0.002 | 1.000 | up |
| DAB1 | RP11-715J22.6 | 0.904 | 0.002 | 1.000 | up |
| DAB1 | RP11-396C23.4 | 0.929 | 0.001 | 1.000 | up |
| DAB1 | MGC2889 | 0.920 | 0.001 | 1.000 | up |
| DAB1 | CTD-2005H7.1 | 0.951 | 0.000 | 1.000 | up |
| DAB1 | G042247 | 0.927 | 0.001 | 1.000 | up |
| DAB1 | OLMALINC | 0.925 | 0.001 | 1.000 | up |
| DAB1 | CCNT2-AS1 | 0.949 | 0.000 | 1.000 | up |
| DAB1 | G023041 | 0.924 | 0.001 | 1.000 | up |
| DAB1 | LOC101928322 | 0.951 | 0.000 | 1.000 | up |
| DAB1 | G053697 | 0.925 | 0.001 | 1.000 | up |
| DAB1 | LOC730102 | 0.954 | 0.000 | 1.000 | up |
| DAB1 | G066437 | 0.947 | 0.000 | 1.000 | up |
| DAB1 | CXADRP3 | 0.914 | 0.001 | 1.000 | up |
| DAB1 | G054532 | 0.918 | 0.001 | 1.000 | up |
| DAB1 | LOC101927811 | 0.925 | 0.001 | 1.000 | up |
| DAB1 | uc.412 | 0.910 | 0.002 | 1.000 | up |
| DAB1 | G014106 | 0.929 | 0.001 | 1.000 | up |
| DAB1 | RP11-166B2.3 | 0.913 | 0.002 | 1.000 | up |
| DAB1 | BC047644 | 0.937 | 0.001 | 1.000 | up |
| DAB1 | LOC101927358 | 0.904 | 0.002 | 1.000 | up |
| DNAJB1 | ASAP1-IT1 | -0.914 | 0.001 | -1.000 | up |
| DNAJB1 | CHKB-CPT1B | -0.900 | 0.002 | -1.000 | up |
| DNAJB1 | G081487 | -0.982 | 0.000 | -1.000 | up |
| DNAJB1 | XLOC_003405 | -0.903 | 0.002 | -1.000 | up |
| DNAJB1 | CTC-261N6.1 | -0.914 | 0.001 | -1.000 | up |
| DNAJB1 | G081302 | -0.907 | 0.002 | -1.000 | up |
| DNAJB1 | G065302 | -0.912 | 0.002 | -1.000 | up |
| DNAJB1 | LOC101926908 | -0.920 | 0.001 | -1.000 | up |
| DNAJB1 | TMLHE-AS1 | -0.918 | 0.001 | -1.000 | up |
| DNAJB1 | RP11-126O1.4 | -0.942 | 0.000 | -1.000 | up |
| DNAJB1 | XLOC_009429 | -0.923 | 0.001 | -1.000 | up |
| DNAJB1 | G059595 | -0.914 | 0.001 | -1.000 | up |
| DNAJB1 | SH3GL1P2 | 0.951 | 0.000 | 1.000 | up |
| DNAJB1 | G064655 | 0.964 | 0.000 | 1.000 | up |
| DNAJB1 | RP5-991G20.4 | 0.952 | 0.000 | 1.000 | up |
| DNAJB1 | RP11-775D22.3 | 0.944 | 0.000 | 1.000 | up |
| DNAJB1 | GPR50-AS1 | 0.925 | 0.001 | 1.000 | up |
| DNAJB1 | G054532 | 0.949 | 0.000 | 1.000 | up |
| DNAJB1 | RP11-314A20.5 | 0.914 | 0.001 | 1.000 | up |
| DNAJB1 | LOC101927811 | 0.941 | 0.001 | 1.000 | up |
| DNAJB1 | uc.412 | 0.913 | 0.002 | 1.000 | up |
| DNAJB1 | JARID2-AS1 | 0.915 | 0.001 | 1.000 | up |
| DNAJB1 | LOC283140 | 0.959 | 0.000 | 1.000 | up |
| DNAJB1 | G057611 | 0.914 | 0.001 | 1.000 | up |
| NPY2R | KLK8 | -0.918 | 0.001 | -1.000 | up |
| NPY2R | CRAT37 | -0.925 | 0.001 | -1.000 | up |
| NPY2R | RP11-567E21.3 | -0.903 | 0.002 | -1.000 | up |
| NPY2R | CTC-250I14.3 | -0.911 | 0.002 | -1.000 | up |
| NPY2R | XLOC_006353 | -0.907 | 0.002 | -1.000 | up |

| | | | | | |
|---|---|---|---|---|---|
| NPY2R | XLOC_010525 | 0.920 | 0.001 | 1.000 | up |
| NPY2R | RP11-762I7.4 | 0.923 | 0.001 | 1.000 | up |
| NPY2R | BC047090 | 0.951 | 0.000 | 1.000 | up |
| NPY2R | ITFG1-AS1 | 0.939 | 0.001 | 1.000 | up |
| CRH | G009303 | -0.916 | 0.001 | -1.000 | up |
| CRH | CTC-261N6.1 | -0.907 | 0.002 | -1.000 | up |
| CRH | CTB-102L5.8 | -0.931 | 0.001 | -1.000 | up |
| CRH | RP1-124C6.1 | -0.926 | 0.001 | -1.000 | up |
| CRH | G021365 | -0.979 | 0.000 | -1.000 | up |
| CRH | G071944 | -0.967 | 0.000 | -1.000 | up |
| CRH | LINC00930 | -0.967 | 0.000 | -1.000 | up |
| CRH | RP11-96H17.3 | -0.921 | 0.001 | -1.000 | up |
| CRH | XLOC_008270 | -0.935 | 0.001 | -1.000 | up |
| CRH | G053358 | -0.913 | 0.002 | -1.000 | up |
| CRH | MIMT1 | -0.902 | 0.002 | -1.000 | up |
| CRH | G031271 | -0.911 | 0.002 | -1.000 | up |
| CRH | G074298 | -0.943 | 0.000 | -1.000 | up |
| CRH | FAM230C | 0.907 | 0.002 | 1.000 | up |
| CRH | G077330 | 0.900 | 0.002 | 1.000 | up |
| CRH | RP11-396C23.4 | 0.912 | 0.002 | 1.000 | up |
| CRH | uc.85 | 0.921 | 0.001 | 1.000 | up |
| CRH | G053260 | 0.919 | 0.001 | 1.000 | up |
| CRH | G041225 | 0.942 | 0.000 | 1.000 | up |
| CRH | CCNT2-AS1 | 0.914 | 0.001 | 1.000 | up |
| CRH | G006537 | 0.923 | 0.001 | 1.000 | up |
| CRH | FAM138E | 0.904 | 0.002 | 1.000 | up |
| CRH | LOC101927822 | 0.948 | 0.000 | 1.000 | up |
| CRH | uc.412 | 0.959 | 0.000 | 1.000 | up |
| CRH | G063721 | 0.904 | 0.002 | 1.000 | up |
| IFT74-AS1 | CALB2 | 0.916 | 0.001 | 1.000 | down |
| IFT74-AS1 | SEPSECS | 0.912 | 0.002 | 1.000 | down |
| IFT74-AS1 | TSPAN7 | 0.910 | 0.002 | 1.000 | down |
| FAM74A4 | TENM4 | 0.929 | 0.001 | 1.000 | down |
| FAM74A4 | AP1G2 | 0.903 | 0.002 | 1.000 | down |
| FAM74A4 | SPTB | 0.922 | 0.001 | 1.000 | down |
| RP11-602N24.3 | ERC2 | 0.903 | 0.002 | 1.000 | down |
| RP11-602N24.3 | CALB2 | 0.917 | 0.001 | 1.000 | down |
| RP11-602N24.3 | PTPRN2 | 0.931 | 0.001 | 1.000 | down |
| RP11-602N24.3 | MAP2 | 0.901 | 0.002 | 1.000 | down |
| RP11-602N24.3 | DPYSL5 | 0.919 | 0.001 | 1.000 | down |
| AL110181 | ERC2 | 0.942 | 0.000 | 1.000 | down |
| AL110181 | GPSM2 | 0.904 | 0.002 | 1.000 | down |
| AL110181 | CALB2 | 0.957 | 0.000 | 1.000 | down |
| AL110181 | PTPRN2 | 0.906 | 0.002 | 1.000 | down |
| AL110181 | DPYSL5 | 0.913 | 0.002 | 1.000 | down |
| AL110181 | DPYSL3 | 0.904 | 0.002 | 1.000 | down |
| TIPARP-AS1 | DAB1 | 0.928 | 0.001 | 1.000 | down |
| TIPARP-AS1 | ERC2 | 0.900 | 0.002 | 1.000 | down |
| TIPARP-AS1 | GPSM2 | 0.952 | 0.000 | 1.000 | down |
| TIPARP-AS1 | PPP2R1A | 0.963 | 0.000 | 1.000 | down |
| RP11-255E6.6 | NRP2 | 0.925 | 0.001 | 1.000 | down |
| RP11-255E6.6 | SPTB | 0.912 | 0.002 | 1.000 | down |
| RP11-255E6.6 | SEPSECS | 0.930 | 0.001 | 1.000 | down |
| XLOC_010525 | NPY2R | 0.920 | 0.001 | 1.000 | down |
| RP11-744I24.3 | TENM4 | 0.925 | 0.001 | 1.000 | down |
| RP11-744I24.3 | SPTB | 0.962 | 0.000 | 1.000 | down |
| RP11-744I24.3 | SEPSECS | 0.948 | 0.000 | 1.000 | down |
| RP11-744I24.3 | TSPAN7 | 0.963 | 0.000 | 1.000 | down |
| XLOC_006779 | SPTB | 0.912 | 0.002 | 1.000 | down |
| G039106 | MAP2 | 0.905 | 0.002 | 1.000 | down |
| G039106 | PPP2R1A | 0.909 | 0.002 | 1.000 | down |
| GSE61474_XLOC_0541[illegible] | TENM4 | 0.980 | 0.000 | 1.000 | down |
| GSE61474_XLOC_0541[illegible] | NRP2 | 0.925 | 0.001 | 1.000 | down |
| GSE61474_XLOC_0541[illegible] | CHRNA3 | 0.921 | 0.001 | 1.000 | down |
| GSE61474_XLOC_0541[illegible] | SPTB | 0.942 | 0.000 | 1.000 | down |

| | | | | | |
|---|---|---|---|---|---|
| GSE61474_XLOC_0541[illegible] | SEPSECS | 0.947 | 0.000 | 1.000 | down |
| RP11-363E6.3 | ERC2 | 0.946 | 0.000 | 1.000 | down |
| RP11-363E6.3 | CALB2 | 0.942 | 0.000 | 1.000 | down |
| RP11-363E6.3 | PTPRN2 | 0.917 | 0.001 | 1.000 | down |
| AK055712 | CRELD1 | 0.914 | 0.002 | 1.000 | down |
| RP11-49K24.8 | FXYD6 | 0.914 | 0.001 | 1.000 | down |
| LOC100130417 | SPTB | 0.947 | 0.000 | 1.000 | down |
| LOC100130417 | SEPSECS | 0.935 | 0.001 | 1.000 | down |
| LOC100130417 | TSPAN7 | 0.978 | 0.000 | 1.000 | down |
| LOC100130417 | CABP1 | 0.962 | 0.000 | 1.000 | down |
| G090511 | ACTN1 | 0.910 | 0.002 | 1.000 | down |
| G090511 | OLFM1 | 0.933 | 0.001 | 1.000 | down |
| RP11-44N22.3 | NRP2 | 0.928 | 0.001 | 1.000 | down |
| RP11-44N22.3 | CHRNA3 | 0.904 | 0.002 | 1.000 | down |
| CTD-2302E22.4 | ARHGAP32 | 0.938 | 0.001 | 1.000 | down |
| CTD-2302E22.4 | ATP2B4 | 0.964 | 0.000 | 1.000 | down |
| CTD-2302E22.4 | PTCH1 | 0.914 | 0.001 | 1.000 | down |
| CTD-2302E22.4 | ERC2 | 0.952 | 0.000 | 1.000 | down |
| CTD-2302E22.4 | PTPRN2 | 0.913 | 0.002 | 1.000 | down |
| CTD-2302E22.4 | MAP2 | 0.930 | 0.001 | 1.000 | down |
| OR2L1P | TSPAN7 | 0.929 | 0.001 | 1.000 | down |
| LINC00652 | ERC2 | 0.970 | 0.000 | 1.000 | down |
| LINC00652 | CALB2 | 0.957 | 0.000 | 1.000 | down |
| LINC00652 | MAP2 | 0.933 | 0.001 | 1.000 | down |
| LINC00652 | PPP2R1A | 0.945 | 0.000 | 1.000 | down |
| RP11-525A16.4 | ATP2B4 | 0.918 | 0.001 | 1.000 | down |
| RP11-525A16.4 | SEPSECS | 0.956 | 0.000 | 1.000 | down |
| RP11-525A16.4 | TSPAN7 | 0.911 | 0.002 | 1.000 | down |
| G017718 | TENM4 | 0.942 | 0.000 | 1.000 | down |
| G017718 | ACTN1 | 0.952 | 0.000 | 1.000 | down |
| RP11-303E16.7 | CRELD1 | 0.934 | 0.001 | 1.000 | down |
| RP11-303E16.7 | ERC2 | 0.913 | 0.002 | 1.000 | down |
| RP11-303E16.7 | FILIP1 | 0.908 | 0.002 | 1.000 | down |
| RP11-303E16.7 | DPYSL5 | 0.902 | 0.002 | 1.000 | down |
| RP11-303E16.7 | RPS6KA5 | 0.917 | 0.001 | 1.000 | down |
| RP11-303E16.7 | DPYSL3 | 0.935 | 0.001 | 1.000 | down |
| XLOC_008005 | SEPSECS | 0.917 | 0.001 | 1.000 | down |
| XLOC_008005 | RPS6KA5 | 0.950 | 0.000 | 1.000 | down |
| CTD-2023N9.3 | CRELD1 | 0.905 | 0.002 | 1.000 | down |
| CTD-2023N9.3 | GPSM2 | 0.912 | 0.002 | 1.000 | down |
| RP11-350N15.5 | ARHGAP32 | 0.903 | 0.002 | 1.000 | down |
| RP11-350N15.5 | ATP2B4 | 0.965 | 0.000 | 1.000 | down |
| RP11-350N15.5 | ERC2 | 0.932 | 0.001 | 1.000 | down |
| RP11-350N15.5 | PTPRN2 | 0.928 | 0.001 | 1.000 | down |
| RP11-350N15.5 | MAP2 | 0.911 | 0.002 | 1.000 | down |
| RP11-762I7.4 | NPY2R | 0.923 | 0.001 | 1.000 | down |
| RP4-549L20.3 | ATP2B4 | 0.915 | 0.001 | 1.000 | down |
| RP4-549L20.3 | ERC2 | 0.904 | 0.002 | 1.000 | down |
| RP4-549L20.3 | CABP1 | 0.927 | 0.001 | 1.000 | down |
| RP11-686O6.2 | CRELD1 | 0.929 | 0.001 | 1.000 | down |
| RP11-686O6.2 | ERC2 | 0.935 | 0.001 | 1.000 | down |
| RP11-686O6.2 | PTPRN2 | 0.957 | 0.000 | 1.000 | down |
| RP11-686O6.2 | DPYSL5 | 0.959 | 0.000 | 1.000 | down |
| RP11-686O6.2 | DPYSL3 | 0.937 | 0.001 | 1.000 | down |
| G061591 | ARHGAP32 | 0.914 | 0.002 | 1.000 | down |
| G061591 | ERC2 | 0.967 | 0.000 | 1.000 | down |
| G061591 | CALB2 | 0.927 | 0.001 | 1.000 | down |
| G061591 | PTPRN2 | 0.984 | 0.000 | 1.000 | down |
| G061591 | MAP2 | 0.977 | 0.000 | 1.000 | down |
| G061591 | DPYSL5 | 0.975 | 0.000 | 1.000 | down |
| G061591 | PPP2R1A | 0.924 | 0.001 | 1.000 | down |
| RP11-356J5.12 | CRELD1 | 0.916 | 0.001 | 1.000 | down |
| RP11-356J5.12 | GPSM2 | 0.908 | 0.002 | 1.000 | down |
| RP11-356J5.12 | DPYSL3 | 0.936 | 0.001 | 1.000 | down |
| FAM230C | EXOC7 | 0.911 | 0.002 | 1.000 | down |

| | | | | | |
|---|---|---|---|---|---|
| FAM230C | CHRNB3 | 0.929 | 0.001 | 1.000 | down |
| FAM230C | NRP2 | 0.944 | 0.000 | 1.000 | down |
| FAM230C | CHRNA3 | 0.907 | 0.002 | 1.000 | down |
| FAM230C | CRH | 0.907 | 0.002 | 1.000 | down |
| USP27X-AS1 | AP1G2 | 0.914 | 0.001 | 1.000 | down |
| USP27X-AS1 | SPTB | 0.960 | 0.000 | 1.000 | down |
| USP27X-AS1 | SEPSECS | 0.909 | 0.002 | 1.000 | down |
| USP27X-AS1 | TSPAN7 | 0.927 | 0.001 | 1.000 | down |
| USP27X-AS1 | CABP1 | 0.903 | 0.002 | 1.000 | down |
| G070136 | TSPAN7 | 0.939 | 0.001 | 1.000 | down |
| EPHA5-AS1 | SIPA1L2 | 0.909 | 0.002 | 1.000 | down |
| RP1-40E16.12 | CRELD1 | 0.925 | 0.001 | 1.000 | down |
| RP1-40E16.12 | ERC2 | 0.982 | 0.000 | 1.000 | down |
| RP1-40E16.12 | CALB2 | 0.949 | 0.000 | 1.000 | down |
| RP1-40E16.12 | PTPRN2 | 0.974 | 0.000 | 1.000 | down |
| RP1-40E16.12 | MAP2 | 0.960 | 0.000 | 1.000 | down |
| RP1-40E16.12 | DPYSL5 | 0.969 | 0.000 | 1.000 | down |
| RP1-40E16.12 | PPP2R1A | 0.920 | 0.001 | 1.000 | down |
| BRWD1-IT2 | ATP2B4 | 0.947 | 0.000 | 1.000 | down |
| BRWD1-IT2 | SEPSECS | 0.917 | 0.001 | 1.000 | down |
| BRWD1-IT2 | TSPAN7 | 0.913 | 0.002 | 1.000 | down |
| BRWD1-IT2 | CABP1 | 0.964 | 0.000 | 1.000 | down |
| RP11-355B11.2 | BTBD9 | 0.973 | 0.000 | 1.000 | down |
| RP11-10J21.4 | MCTP2 | 0.939 | 0.001 | 1.000 | down |
| XLOC_002675 | EXOC7 | 0.900 | 0.002 | 1.000 | down |
| XLOC_002675 | CHRNA3 | 0.914 | 0.001 | 1.000 | down |
| G021652 | TENM4 | 0.907 | 0.002 | 1.000 | down |
| G021652 | NRP2 | 0.909 | 0.002 | 1.000 | down |
| G021652 | ACTN1 | 0.902 | 0.002 | 1.000 | down |
| GBAT2 | CRELD1 | 0.918 | 0.001 | 1.000 | down |
| GBAT2 | HSP90AA1 | 0.902 | 0.002 | 1.000 | down |
| GBAT2 | ERC2 | 0.970 | 0.000 | 1.000 | down |
| GBAT2 | GPSM2 | 0.902 | 0.002 | 1.000 | down |
| GBAT2 | CALB2 | 0.943 | 0.000 | 1.000 | down |
| GBAT2 | PTPRN2 | 0.958 | 0.000 | 1.000 | down |
| GBAT2 | MAP2 | 0.958 | 0.000 | 1.000 | down |
| GBAT2 | DPYSL5 | 0.967 | 0.000 | 1.000 | down |
| GBAT2 | DPYSL3 | 0.900 | 0.002 | 1.000 | down |
| GBAT2 | PPP2R1A | 0.945 | 0.000 | 1.000 | down |
| RP11-490O6.2 | ATP2B4 | 0.957 | 0.000 | 1.000 | down |
| RP11-490O6.2 | SPTB | 0.908 | 0.002 | 1.000 | down |
| RP11-490O6.2 | SEPSECS | 0.953 | 0.000 | 1.000 | down |
| RP11-490O6.2 | TSPAN7 | 0.939 | 0.001 | 1.000 | down |
| RP11-490O6.2 | CABP1 | 0.986 | 0.000 | 1.000 | down |
| ZFX-AS1 | ARHGAP32 | 0.915 | 0.001 | 1.000 | down |
| ZFX-AS1 | ATP2B4 | 0.914 | 0.001 | 1.000 | down |
| ZFX-AS1 | ERC2 | 0.946 | 0.000 | 1.000 | down |
| ZFX-AS1 | PTPRN2 | 0.945 | 0.000 | 1.000 | down |
| ZFX-AS1 | MAP2 | 0.945 | 0.000 | 1.000 | down |
| BC047090 | NPY2R | 0.951 | 0.000 | 1.000 | down |
| RP11-1017G21.6 | CRELD1 | 0.914 | 0.002 | 1.000 | down |
| RP11-1017G21.6 | ATP2B4 | 0.967 | 0.000 | 1.000 | down |
| RP11-1017G21.6 | CABP1 | 0.938 | 0.001 | 1.000 | down |
| G002862 | TENM4 | 0.930 | 0.001 | 1.000 | down |
| G002862 | EXOC7 | 0.912 | 0.002 | 1.000 | down |
| G002862 | AP1G2 | 0.969 | 0.000 | 1.000 | down |
| G002862 | SPTB | 0.916 | 0.001 | 1.000 | down |
| PSMG3-AS1 (2) | SEPSECS | 0.942 | 0.000 | 1.000 | down |
| PSMG3-AS1 (2) | TSPAN7 | 0.913 | 0.002 | 1.000 | down |
| SH3GL1P2 | HSPA8 | 0.960 | 0.000 | 1.000 | down |
| SH3GL1P2 | DNAJB1 | 0.951 | 0.000 | 1.000 | down |
| SH3GL1P2 | BTBD9 | 0.902 | 0.002 | 1.000 | down |
| AC138655.6 | PTPRN2 | 0.920 | 0.001 | 1.000 | down |
| LOC729083 | PTCH1 | 0.916 | 0.001 | 1.000 | down |
| TM4SF1-AS1 | ERC2 | 0.942 | 0.000 | 1.000 | down |

| | | | | | |
|---|---|---|---|---|---|
| TM4SF1-AS1 | CALB2 | 0.906 | 0.002 | 1.000 | down |
| TM4SF1-AS1 | SEPSECS | 0.906 | 0.002 | 1.000 | down |
| CTD-2033A16.3 | ATP2B4 | 0.908 | 0.002 | 1.000 | down |
| CTD-2033A16.3 | TSPAN7 | 0.938 | 0.001 | 1.000 | down |
| CTD-2033A16.3 | CABP1 | 0.901 | 0.002 | 1.000 | down |
| LINC00662 | DAB1 | 0.920 | 0.001 | 1.000 | down |
| LINC00662 | ERC2 | 0.932 | 0.001 | 1.000 | down |
| LINC00662 | FILIP1 | 0.907 | 0.002 | 1.000 | down |
| LINC00662 | CALB2 | 0.904 | 0.002 | 1.000 | down |
| RP11-297J22.1 | CRELD1 | 0.923 | 0.001 | 1.000 | down |
| RP11-297J22.1 | ERC2 | 0.938 | 0.001 | 1.000 | down |
| RP11-297J22.1 | GPSM2 | 0.903 | 0.002 | 1.000 | down |
| RP11-297J22.1 | PTPRN2 | 0.946 | 0.000 | 1.000 | down |
| RP11-297J22.1 | MAP2 | 0.966 | 0.000 | 1.000 | down |
| RP11-297J22.1 | DPYSL5 | 0.977 | 0.000 | 1.000 | down |
| RP11-297J22.1 | PPP2R1A | 0.938 | 0.001 | 1.000 | down |
| RP11-179B2.2 | CRELD1 | 0.900 | 0.002 | 1.000 | down |
| RP11-179B2.2 | ARHGAP32 | 0.911 | 0.002 | 1.000 | down |
| RP11-179B2.2 | ERC2 | 0.960 | 0.000 | 1.000 | down |
| RP11-179B2.2 | CALB2 | 0.905 | 0.002 | 1.000 | down |
| RP11-179B2.2 | PTPRN2 | 0.962 | 0.000 | 1.000 | down |
| RP11-179B2.2 | MAP2 | 0.936 | 0.001 | 1.000 | down |
| RP11-179B2.2 | DPYSL5 | 0.927 | 0.001 | 1.000 | down |
| G064655 | DAB1 | 0.912 | 0.002 | 1.000 | down |
| G064655 | DNAJB1 | 0.964 | 0.000 | 1.000 | down |
| G017886 | CALB2 | 0.909 | 0.002 | 1.000 | down |
| CTD-2588C8.8 | ATP2B4 | 0.902 | 0.002 | 1.000 | down |
| CTD-2588C8.8 | SEPSECS | 0.902 | 0.002 | 1.000 | down |
| CTD-2588C8.8 | TSPAN7 | 0.901 | 0.002 | 1.000 | down |
| CTD-2588C8.8 | CABP1 | 0.912 | 0.002 | 1.000 | down |
| G047225 | CRELD1 | 0.912 | 0.002 | 1.000 | down |
| G047225 | ELFN1 | 0.922 | 0.001 | 1.000 | down |
| G047225 | ATP2B4 | 0.916 | 0.001 | 1.000 | down |
| RP1-309F20.3 | ATP2B4 | 0.919 | 0.001 | 1.000 | down |
| RP1-309F20.3 | SPTB | 0.902 | 0.002 | 1.000 | down |
| RP1-309F20.3 | SEPSECS | 0.935 | 0.001 | 1.000 | down |
| RP1-309F20.3 | TSPAN7 | 0.906 | 0.002 | 1.000 | down |
| RP1-309F20.3 | CABP1 | 0.980 | 0.000 | 1.000 | down |
| RP1-309F20.3 | RPS6KA5 | 0.915 | 0.001 | 1.000 | down |
| RP11-643M14.1 | ATP2B4 | 0.905 | 0.002 | 1.000 | down |
| RP11-643M14.1 | TSPAN7 | 0.918 | 0.001 | 1.000 | down |
| RP11-643M14.1 | CABP1 | 0.946 | 0.000 | 1.000 | down |
| RP11-643M14.1 | CHRNA7 | 0.914 | 0.001 | 1.000 | down |
| RP11-344E13.4 | PPP2R1A | 0.911 | 0.002 | 1.000 | down |
| SCAMP1-AS1 | EXOC7 | 0.949 | 0.000 | 1.000 | down |
| G017670 | ALDOC | 0.902 | 0.002 | 1.000 | down |
| G017670 | OLFM1 | 0.927 | 0.001 | 1.000 | down |
| RP11-21M24.2 | MCTP2 | 0.974 | 0.000 | 1.000 | down |
| RP11-252A24.3 | SEPSECS | 0.906 | 0.002 | 1.000 | down |
| AC073321.4 | OLFM1 | 0.936 | 0.001 | 1.000 | down |
| XLOC_008875 | ARHGAP32 | 0.917 | 0.001 | 1.000 | down |
| XLOC_008875 | ATP2B4 | 0.926 | 0.001 | 1.000 | down |
| G077330 | CHRNB3 | 0.915 | 0.001 | 1.000 | down |
| G077330 | CRH | 0.900 | 0.002 | 1.000 | down |
| RP11-46J23.1 | BTBD9 | 0.945 | 0.000 | 1.000 | down |
| RP11-715J22.6 | NRP2 | 0.907 | 0.002 | 1.000 | down |
| RP11-715J22.6 | DAB1 | 0.904 | 0.002 | 1.000 | down |
| RP11-277L2.3 | SOS1 | 0.904 | 0.002 | 1.000 | down |
| RP11-191L17.1 | CRELD1 | 0.936 | 0.001 | 1.000 | down |
| RP11-191L17.1 | ATP2B4 | 0.959 | 0.000 | 1.000 | down |
| RP11-191L17.1 | ERC2 | 0.965 | 0.000 | 1.000 | down |
| RP11-191L17.1 | PTPRN2 | 0.922 | 0.001 | 1.000 | down |
| RP11-191L17.1 | SEPSECS | 0.946 | 0.000 | 1.000 | down |
| RP11-191L17.1 | TSPAN7 | 0.906 | 0.002 | 1.000 | down |
| RP11-191L17.1 | MAP2 | 0.908 | 0.002 | 1.000 | down |

| | | | | | |
|---|---|---|---|---|---|
| RP11-191L17.1 | CABP1 | 0.929 | 0.001 | 1.000 | down |
| RP11-191L17.1 | DPYSL5 | 0.904 | 0.002 | 1.000 | down |
| DLG5-AS1 (2) | EXOC7 | 0.906 | 0.002 | 1.000 | down |
| WWC2-AS2 | SOS1 | 0.911 | 0.002 | 1.000 | down |
| AC099754.1 | ERC2 | 0.922 | 0.001 | 1.000 | down |
| AC099754.1 | CALB2 | 0.922 | 0.001 | 1.000 | down |
| AC099754.1 | PTPRN2 | 0.905 | 0.002 | 1.000 | down |
| RP11-165A20.3 | ATP2B4 | 0.938 | 0.001 | 1.000 | down |
| RP11-165A20.3 | SEPSECS | 0.906 | 0.002 | 1.000 | down |
| RP11-165A20.3 | TSPAN7 | 0.930 | 0.001 | 1.000 | down |
| RP11-165A20.3 | CABP1 | 0.976 | 0.000 | 1.000 | down |
| G018162 | CRELD1 | 0.912 | 0.002 | 1.000 | down |
| G018162 | SIPA1L2 | 0.903 | 0.002 | 1.000 | down |
| G018162 | FILIP1 | 0.965 | 0.000 | 1.000 | down |
| G079009 | PHB | 0.914 | 0.002 | 1.000 | down |
| RP11-396C23.4 | HSPA8 | 0.932 | 0.001 | 1.000 | down |
| RP11-396C23.4 | DAB1 | 0.929 | 0.001 | 1.000 | down |
| RP11-396C23.4 | CRH | 0.912 | 0.002 | 1.000 | down |
| RP11-396C23.4 | BTBD9 | 0.941 | 0.000 | 1.000 | down |
| G000481 | ATP2B4 | 0.917 | 0.001 | 1.000 | down |
| G000481 | TSPAN7 | 0.923 | 0.001 | 1.000 | down |
| G000481 | CABP1 | 0.972 | 0.000 | 1.000 | down |
| G000481 | CHRNA7 | 0.902 | 0.002 | 1.000 | down |
| RP11-285F16.1 | CRELD1 | 0.902 | 0.002 | 1.000 | down |
| RP11-285F16.1 | ATP2B4 | 0.964 | 0.000 | 1.000 | down |
| RP11-285F16.1 | PTCH1 | 0.908 | 0.002 | 1.000 | down |
| RP11-285F16.1 | ERC2 | 0.937 | 0.001 | 1.000 | down |
| RP11-285F16.1 | SEPSECS | 0.944 | 0.000 | 1.000 | down |
| RP11-285F16.1 | TSPAN7 | 0.927 | 0.001 | 1.000 | down |
| RP11-285F16.1 | CABP1 | 0.936 | 0.001 | 1.000 | down |
| G050681 | SPTB | 0.974 | 0.000 | 1.000 | down |
| G050681 | SEPSECS | 0.928 | 0.001 | 1.000 | down |
| G050681 | TSPAN7 | 0.912 | 0.002 | 1.000 | down |
| G050681 | CABP1 | 0.942 | 0.000 | 1.000 | down |
| CTC-232P5.3 | BTBD9 | 0.960 | 0.000 | 1.000 | down |
| G047031 | HSPA8 | 0.961 | 0.000 | 1.000 | down |
| RP5-991G20.4 | DNAJB1 | 0.952 | 0.000 | 1.000 | down |
| RP5-1092A3.4 | CRELD1 | 0.932 | 0.001 | 1.000 | down |
| RP5-1092A3.4 | ARHGAP32 | 0.902 | 0.002 | 1.000 | down |
| BC112270 | EXOC7 | 0.912 | 0.002 | 1.000 | down |
| G014329 | ADGRL2 | 0.901 | 0.002 | 1.000 | down |
| RP11-351D16.3 | CRELD1 | 0.918 | 0.001 | 1.000 | down |
| RP11-351D16.3 | ATP2B4 | 0.952 | 0.000 | 1.000 | down |
| RP11-351D16.3 | SEPSECS | 0.901 | 0.002 | 1.000 | down |
| RP11-351D16.3 | CABP1 | 0.965 | 0.000 | 1.000 | down |
| MGC2889 | DAB1 | 0.920 | 0.001 | 1.000 | down |
| MGC2889 | ERC2 | 0.927 | 0.001 | 1.000 | down |
| MGC2889 | CALB2 | 0.954 | 0.000 | 1.000 | down |
| MGC2889 | BTBD9 | 0.913 | 0.002 | 1.000 | down |
| MGC2889 | PPP2R1A | 0.933 | 0.001 | 1.000 | down |
| AK123758 | ATP2B4 | 0.935 | 0.001 | 1.000 | down |
| AK123758 | SEPSECS | 0.913 | 0.002 | 1.000 | down |
| AK123758 | CABP1 | 0.916 | 0.001 | 1.000 | down |
| RP4-683L5.1 | CRELD1 | 0.938 | 0.001 | 1.000 | down |
| RP4-683L5.1 | ATP2B4 | 0.936 | 0.001 | 1.000 | down |
| RP4-683L5.1 | ERC2 | 0.917 | 0.001 | 1.000 | down |
| RP4-683L5.1 | PTPRN2 | 0.907 | 0.002 | 1.000 | down |
| RP4-683L5.1 | CABP1 | 0.908 | 0.002 | 1.000 | down |
| RP4-683L5.1 | DPYSL5 | 0.907 | 0.002 | 1.000 | down |
| G086828 | MCTP2 | 0.910 | 0.002 | 1.000 | down |
| G086828 | FXYD6 | 0.907 | 0.002 | 1.000 | down |
| RP11-353K11.1 | CRELD1 | 0.958 | 0.000 | 1.000 | down |
| RP11-353K11.1 | ARHGAP32 | 0.931 | 0.001 | 1.000 | down |
| RP11-353K11.1 | ATP2B4 | 0.966 | 0.000 | 1.000 | down |
| G082153 | HSPA8 | 0.903 | 0.002 | 1.000 | down |

| | | | | | |
|---|---|---|---|---|---|
| HRAT5 | ATP2B4 | 0.922 | 0.001 | 1.000 | down |
| HRAT5 | TSPAN7 | 0.938 | 0.001 | 1.000 | down |
| HRAT5 | CABP1 | 0.957 | 0.000 | 1.000 | down |
| RP11-209D14.2 | CRELD1 | 0.905 | 0.002 | 1.000 | down |
| AC002467.7 | CRELD1 | 0.932 | 0.001 | 1.000 | down |
| AC002467.7 | FILIP1 | 0.960 | 0.000 | 1.000 | down |
| AC002467.7 | RPS6KA5 | 0.908 | 0.002 | 1.000 | down |
| G015380 | CRELD1 | 0.930 | 0.001 | 1.000 | down |
| G015380 | ELFN1 | 0.904 | 0.002 | 1.000 | down |
| CTD-2005H7.1 | DAB1 | 0.951 | 0.000 | 1.000 | down |
| TAF1A-AS1 | ARHGAP32 | 0.950 | 0.000 | 1.000 | down |
| uc.85 | CRH | 0.921 | 0.001 | 1.000 | down |
| uc.85 | BTBD9 | 0.903 | 0.002 | 1.000 | down |
| XLOC_001189 | CRELD1 | 0.913 | 0.002 | 1.000 | down |
| XLOC_001189 | ELFN1 | 0.942 | 0.000 | 1.000 | down |
| XLOC_003595 | OLFM1 | 0.969 | 0.000 | 1.000 | down |
| G066319 | HSP90AA1 | 0.966 | 0.000 | 1.000 | down |
| G066319 | ADGRL2 | 0.922 | 0.001 | 1.000 | down |
| G066319 | SIPA1L2 | 0.923 | 0.001 | 1.000 | down |
| XLOC_006554 | CRELD1 | 0.927 | 0.001 | 1.000 | down |
| XLOC_006554 | ARHGAP32 | 0.961 | 0.000 | 1.000 | down |
| FAM66A | PHB | 0.921 | 0.001 | 1.000 | down |
| G042247 | DAB1 | 0.927 | 0.001 | 1.000 | down |
| RP11-475I24.3 | PHB | 0.922 | 0.001 | 1.000 | down |
| RP11-475I24.3 | ADGRL2 | 0.902 | 0.002 | 1.000 | down |
| AC006273.4 | SOS1 | 0.928 | 0.001 | 1.000 | down |
| AC006273.4 | CHRNB3 | 0.970 | 0.000 | 1.000 | down |
| RP11-312B8.1 | ATP2B4 | 0.977 | 0.000 | 1.000 | down |
| RP11-312B8.1 | SEPSECS | 0.923 | 0.001 | 1.000 | down |
| RP11-312B8.1 | CABP1 | 0.920 | 0.001 | 1.000 | down |
| GAS5-AS1 | ATP2B4 | 0.929 | 0.001 | 1.000 | down |
| GAS5-AS1 | ERC2 | 0.930 | 0.001 | 1.000 | down |
| GAS5-AS1 | PTPRN2 | 0.919 | 0.001 | 1.000 | down |
| GAS5-AS1 | TSPAN7 | 0.911 | 0.002 | 1.000 | down |
| GAS5-AS1 | CABP1 | 0.918 | 0.001 | 1.000 | down |
| AC144652.1 | CRELD1 | 0.910 | 0.002 | 1.000 | down |
| AC144652.1 | GPSM2 | 0.968 | 0.000 | 1.000 | down |
| AC144652.1 | DPYSL5 | 0.932 | 0.001 | 1.000 | down |
| AC144652.1 | DPYSL3 | 0.949 | 0.000 | 1.000 | down |
| RP11-204E9.1 | HSP90AA1 | 0.927 | 0.001 | 1.000 | down |
| RP11-204E9.1 | ERC2 | 0.909 | 0.002 | 1.000 | down |
| RP11-204E9.1 | PPP2R1A | 0.913 | 0.002 | 1.000 | down |
| G085410 | CRELD1 | 0.960 | 0.000 | 1.000 | down |
| G085410 | ARHGAP32 | 0.918 | 0.001 | 1.000 | down |
| G085410 | HSP90AA1 | 0.907 | 0.002 | 1.000 | down |
| G085410 | ADGRL2 | 0.914 | 0.001 | 1.000 | down |
| G085410 | ERC2 | 0.942 | 0.000 | 1.000 | down |
| G085410 | PTPRN2 | 0.934 | 0.001 | 1.000 | down |
| G085410 | MAP2 | 0.924 | 0.001 | 1.000 | down |
| G085410 | DPYSL5 | 0.927 | 0.001 | 1.000 | down |
| uc.446 | TENM4 | 0.916 | 0.001 | 1.000 | down |
| uc.446 | ACTN1 | 0.982 | 0.000 | 1.000 | down |
| DHRS4-AS1 (1) | ERC2 | 0.926 | 0.001 | 1.000 | down |
| DHRS4-AS1 (1) | SEPSECS | 0.928 | 0.001 | 1.000 | down |
| RP3-410B11.1 | ATP2B4 | 0.921 | 0.001 | 1.000 | down |
| RP3-410B11.1 | SPTB | 0.904 | 0.002 | 1.000 | down |
| RP3-410B11.1 | SEPSECS | 0.917 | 0.001 | 1.000 | down |
| RP3-410B11.1 | TSPAN7 | 0.949 | 0.000 | 1.000 | down |
| RP3-410B11.1 | CABP1 | 0.984 | 0.000 | 1.000 | down |
| G089005 | CRELD1 | 0.987 | 0.000 | 1.000 | down |
| G089005 | ATP2B4 | 0.941 | 0.000 | 1.000 | down |
| G089005 | ERC2 | 0.948 | 0.000 | 1.000 | down |
| G089005 | PTPRN2 | 0.911 | 0.002 | 1.000 | down |
| G089005 | SEPSECS | 0.901 | 0.002 | 1.000 | down |
| G089005 | MAP2 | 0.910 | 0.002 | 1.000 | down |

| | | | | | |
|---|---|---|---|---|---|
| G089005 | DPYSL5 | 0.919 | 0.001 | 1.000 | down |
| G048385 | ARHGAP32 | 0.940 | 0.001 | 1.000 | down |
| OLMALINC | DAB1 | 0.925 | 0.001 | 1.000 | down |
| OLMALINC | ERC2 | 0.915 | 0.001 | 1.000 | down |
| OLMALINC | GPSM2 | 0.913 | 0.002 | 1.000 | down |
| OLMALINC | BTBD9 | 0.934 | 0.001 | 1.000 | down |
| OLMALINC | PPP2R1A | 0.961 | 0.000 | 1.000 | down |
| G000619 | CABP1 | 0.940 | 0.001 | 1.000 | down |
| G000619 | CHRNA7 | 0.903 | 0.002 | 1.000 | down |
| AC093391.2 | NRP2 | 0.902 | 0.002 | 1.000 | down |
| AC093391.2 | SPTB | 0.952 | 0.000 | 1.000 | down |
| AC093391.2 | RPS6KA5 | 0.923 | 0.001 | 1.000 | down |
| LOC101929741 | ELFN1 | 0.928 | 0.001 | 1.000 | down |
| LOC101929741 | ATP2B4 | 0.904 | 0.002 | 1.000 | down |
| LOC101929741 | CABP1 | 0.935 | 0.001 | 1.000 | down |
| G079654 | ATP2B4 | 0.956 | 0.000 | 1.000 | down |
| G079654 | PTCH1 | 0.915 | 0.001 | 1.000 | down |
| G079654 | SEPSECS | 0.940 | 0.001 | 1.000 | down |
| G079654 | TSPAN7 | 0.905 | 0.002 | 1.000 | down |
| G079654 | CABP1 | 0.943 | 0.000 | 1.000 | down |
| TEX26-AS1 | ATP2B4 | 0.957 | 0.000 | 1.000 | down |
| TEX26-AS1 | PTCH1 | 0.914 | 0.002 | 1.000 | down |
| RP11-496H1.2 | ARHGAP32 | 0.908 | 0.002 | 1.000 | down |
| RP11-496H1.2 | ATP2B4 | 0.916 | 0.001 | 1.000 | down |
| RP11-496H1.2 | ERC2 | 0.955 | 0.000 | 1.000 | down |
| RP11-496H1.2 | PTPRN2 | 0.944 | 0.000 | 1.000 | down |
| RP11-496H1.2 | MAP2 | 0.937 | 0.001 | 1.000 | down |
| RP11-510M2.2 | TSPAN7 | 0.912 | 0.002 | 1.000 | down |
| RP11-510M2.2 | CABP1 | 0.900 | 0.002 | 1.000 | down |
| XLOC_006928 | SPTB | 0.956 | 0.000 | 1.000 | down |
| XLOC_006928 | SEPSECS | 0.928 | 0.001 | 1.000 | down |
| XLOC_006928 | CABP1 | 0.905 | 0.002 | 1.000 | down |
| G009018 | PTCH1 | 0.997 | 0.000 | 1.000 | down |
| RP11-2E11.10 | CABP1 | 0.931 | 0.001 | 1.000 | down |
| FRMD6-AS1 | ATP2B4 | 0.935 | 0.001 | 1.000 | down |
| FRMD6-AS1 | TSPAN7 | 0.921 | 0.001 | 1.000 | down |
| FRMD6-AS1 | CABP1 | 0.964 | 0.000 | 1.000 | down |
| FRMD6-AS1 | CHRNA7 | 0.911 | 0.002 | 1.000 | down |
| RP11-1055B8.8 | PTCH1 | 0.916 | 0.001 | 1.000 | down |
| LINC00408 | CRELD1 | 0.904 | 0.002 | 1.000 | down |
| LINC00408 | HSP90AA1 | 0.908 | 0.002 | 1.000 | down |
| LINC00408 | ADGRL2 | 0.924 | 0.001 | 1.000 | down |
| LINC00408 | SIPA1L2 | 0.944 | 0.000 | 1.000 | down |
| LINC00408 | GPSM2 | 0.931 | 0.001 | 1.000 | down |
| LINC00408 | DPYSL3 | 0.932 | 0.001 | 1.000 | down |
| RP11-128P10.1 | TENM4 | 0.929 | 0.001 | 1.000 | down |
| RP11-128P10.1 | AP1G2 | 0.924 | 0.001 | 1.000 | down |
| RP11-128P10.1 | NRP2 | 0.947 | 0.000 | 1.000 | down |
| RP11-128P10.1 | SPTB | 0.957 | 0.000 | 1.000 | down |
| G011337 | TENM4 | 0.909 | 0.002 | 1.000 | down |
| RP11-382D12.2 | OLFM1 | 0.937 | 0.001 | 1.000 | down |
| RP11-174G6.5 | FILIP1 | 0.920 | 0.001 | 1.000 | down |
| RP11-174G6.5 | SPTB | 0.925 | 0.001 | 1.000 | down |
| RP11-174G6.5 | SEPSECS | 0.979 | 0.000 | 1.000 | down |
| RP11-174G6.5 | TSPAN7 | 0.907 | 0.002 | 1.000 | down |
| RP11-174G6.5 | CABP1 | 0.927 | 0.001 | 1.000 | down |
| RP11-174G6.5 | RPS6KA5 | 0.904 | 0.002 | 1.000 | down |
| TCR-alpha | ARHGAP32 | 0.901 | 0.002 | 1.000 | down |
| AL355711 | ATP2B4 | 0.948 | 0.000 | 1.000 | down |
| AL355711 | SPTB | 0.911 | 0.002 | 1.000 | down |
| AL355711 | SEPSECS | 0.941 | 0.001 | 1.000 | down |
| AL355711 | TSPAN7 | 0.912 | 0.002 | 1.000 | down |
| AL355711 | CABP1 | 0.983 | 0.000 | 1.000 | down |
| CTD-2256P15.2 | ATP2B4 | 0.967 | 0.000 | 1.000 | down |
| CTD-2256P15.2 | SEPSECS | 0.932 | 0.001 | 1.000 | down |

| | | | | | |
|---|---|---|---|---|---|
| CTD-2256P15.2 | TSPAN7 | 0.914 | 0.001 | 1.000 | down |
| CTD-2256P15.2 | CABP1 | 0.975 | 0.000 | 1.000 | down |
| TUB-AS1 | TENM4 | 0.952 | 0.000 | 1.000 | down |
| TUB-AS1 | NRP2 | 0.913 | 0.002 | 1.000 | down |
| TUB-AS1 | ACTN1 | 0.955 | 0.000 | 1.000 | down |
| G053260 | CRH | 0.919 | 0.001 | 1.000 | down |
| BC070391 | CRELD1 | 0.934 | 0.001 | 1.000 | down |
| BC070391 | ATP2B4 | 0.935 | 0.001 | 1.000 | down |
| BC070391 | CABP1 | 0.913 | 0.002 | 1.000 | down |
| SIX3-AS1 | ARHGAP32 | 0.914 | 0.001 | 1.000 | down |
| G041948 | ATP2B4 | 0.937 | 0.001 | 1.000 | down |
| G034470 | CRELD1 | 0.956 | 0.000 | 1.000 | down |
| G034470 | ATP2B4 | 0.939 | 0.001 | 1.000 | down |
| G034470 | FILIP1 | 0.918 | 0.001 | 1.000 | down |
| MAPT-IT1 | ERC2 | 0.917 | 0.001 | 1.000 | down |
| G041225 | HSPA8 | 0.931 | 0.001 | 1.000 | down |
| G041225 | CRH | 0.942 | 0.000 | 1.000 | down |
| G073576 | CRELD1 | 0.944 | 0.000 | 1.000 | down |
| G073576 | ATP2B4 | 0.943 | 0.000 | 1.000 | down |
| G073576 | ERC2 | 0.961 | 0.000 | 1.000 | down |
| G073576 | PTPRN2 | 0.948 | 0.000 | 1.000 | down |
| G073576 | MAP2 | 0.926 | 0.001 | 1.000 | down |
| G073576 | DPYSL5 | 0.951 | 0.000 | 1.000 | down |
| XLOC_013222 | CRELD1 | 0.908 | 0.002 | 1.000 | down |
| XLOC_013222 | ERC2 | 0.944 | 0.000 | 1.000 | down |
| XLOC_013222 | PTPRN2 | 0.910 | 0.002 | 1.000 | down |
| XLOC_013222 | DPYSL5 | 0.913 | 0.002 | 1.000 | down |
| LINC-PINT (1) | PTCH1 | 0.916 | 0.001 | 1.000 | down |
| RP11-299H21.1 | PTCH1 | 0.908 | 0.002 | 1.000 | down |
| ITFG1-AS1 | NPY2R | 0.939 | 0.001 | 1.000 | down |
| G081251 | ELFN1 | 0.930 | 0.001 | 1.000 | down |
| G081251 | CHRNA7 | 0.940 | 0.001 | 1.000 | down |
| CTD-2517M22.16 | MCTP2 | 0.914 | 0.002 | 1.000 | down |
| RP11-424N24.2 | ATP2B4 | 0.910 | 0.002 | 1.000 | down |
| RP11-424N24.2 | SEPSECS | 0.967 | 0.000 | 1.000 | down |
| CCNT2-AS1 | HSPA8 | 0.904 | 0.002 | 1.000 | down |
| CCNT2-AS1 | DAB1 | 0.949 | 0.000 | 1.000 | down |
| CCNT2-AS1 | CRH | 0.914 | 0.001 | 1.000 | down |
| CCNT2-AS1 | BTBD9 | 0.906 | 0.002 | 1.000 | down |
| XLOC_011052 | CRELD1 | 0.911 | 0.002 | 1.000 | down |
| XLOC_011052 | ATP2B4 | 0.947 | 0.000 | 1.000 | down |
| XLOC_011052 | PTCH1 | 0.910 | 0.002 | 1.000 | down |
| XLOC_011052 | ERC2 | 0.960 | 0.000 | 1.000 | down |
| XLOC_011052 | SEPSECS | 0.947 | 0.000 | 1.000 | down |
| XLOC_011052 | TSPAN7 | 0.903 | 0.002 | 1.000 | down |
| XLOC_011052 | MAP2 | 0.914 | 0.002 | 1.000 | down |
| G061579 | PHB | 0.914 | 0.001 | 1.000 | down |
| RP11-502I4.3 | CHRNA3 | 0.934 | 0.001 | 1.000 | down |
| RP11-502I4.3 | SEPSECS | 0.911 | 0.002 | 1.000 | down |
| G023041 | DAB1 | 0.924 | 0.001 | 1.000 | down |
| G023041 | GPSM2 | 0.908 | 0.002 | 1.000 | down |
| G023041 | BTBD9 | 0.901 | 0.002 | 1.000 | down |
| G023041 | PPP2R1A | 0.921 | 0.001 | 1.000 | down |
| RP1-202O8.3 | ATP2B4 | 0.952 | 0.000 | 1.000 | down |
| RP1-202O8.3 | SPTB | 0.912 | 0.002 | 1.000 | down |
| RP1-202O8.3 | SEPSECS | 0.952 | 0.000 | 1.000 | down |
| RP1-202O8.3 | TSPAN7 | 0.966 | 0.000 | 1.000 | down |
| RP1-202O8.3 | CABP1 | 0.976 | 0.000 | 1.000 | down |
| RP13-131K19.2 | SPTB | 0.921 | 0.001 | 1.000 | down |
| RP13-131K19.2 | SEPSECS | 0.918 | 0.001 | 1.000 | down |
| RP13-131K19.2 | TSPAN7 | 0.904 | 0.002 | 1.000 | down |
| RP13-131K19.2 | CABP1 | 0.901 | 0.002 | 1.000 | down |
| G082866 | RPS6KA5 | 0.931 | 0.001 | 1.000 | down |
| XLOC_000308 | TENM4 | 0.905 | 0.002 | 1.000 | down |
| XLOC_000308 | ACTN1 | 0.971 | 0.000 | 1.000 | down |

| | | | | | |
|---|---|---|---|---|---|
| FOXN3-AS1 | SPTB | 0.957 | 0.000 | 1.000 | down |
| FOXN3-AS1 | SEPSECS | 0.950 | 0.000 | 1.000 | down |
| FOXN3-AS1 | TSPAN7 | 0.982 | 0.000 | 1.000 | down |
| FOXN3-AS1 | CABP1 | 0.973 | 0.000 | 1.000 | down |
| RP11-290L1.2 | TENM4 | 0.912 | 0.002 | 1.000 | down |
| RP11-290L1.2 | CHRNA3 | 0.912 | 0.002 | 1.000 | down |
| RP11-290L1.2 | SEPSECS | 0.916 | 0.001 | 1.000 | down |
| TNRC18P1 | CRELD1 | 0.939 | 0.001 | 1.000 | down |
| TNRC18P1 | ARHGAP32 | 0.940 | 0.001 | 1.000 | down |
| TNRC18P1 | ERC2 | 0.927 | 0.001 | 1.000 | down |
| TNRC18P1 | PTPRN2 | 0.933 | 0.001 | 1.000 | down |
| TNRC18P1 | MAP2 | 0.947 | 0.000 | 1.000 | down |
| TNRC18P1 | DPYSL5 | 0.941 | 0.000 | 1.000 | down |
| GSE61474_XLOC_0382[illegible] | CABP1 | 0.958 | 0.000 | 1.000 | down |
| IDH1-AS1 | TENM4 | 0.901 | 0.002 | 1.000 | down |
| IDH1-AS1 | NRP2 | 0.917 | 0.001 | 1.000 | down |
| IDH1-AS1 | SPTB | 0.921 | 0.001 | 1.000 | down |
| IDH1-AS1 | SEPSECS | 0.903 | 0.002 | 1.000 | down |
| G065067 | ERC2 | 0.940 | 0.001 | 1.000 | down |
| G065067 | CALB2 | 0.934 | 0.001 | 1.000 | down |
| G065067 | PTPRN2 | 0.947 | 0.000 | 1.000 | down |
| G065067 | MAP2 | 0.924 | 0.001 | 1.000 | down |
| G019632 | ATP2B4 | 0.902 | 0.002 | 1.000 | down |
| G019632 | TSPAN7 | 0.918 | 0.001 | 1.000 | down |
| NIPBL-AS1 | CALB2 | 0.916 | 0.001 | 1.000 | down |
| NIPBL-AS1 | BTBD9 | 0.909 | 0.002 | 1.000 | down |
| NIPBL-AS1 | PPP2R1A | 0.934 | 0.001 | 1.000 | down |
| KB-1592A4.14 | TENM4 | 0.930 | 0.001 | 1.000 | down |
| G038972 | ATP2B4 | 0.948 | 0.000 | 1.000 | down |
| G038972 | SEPSECS | 0.917 | 0.001 | 1.000 | down |
| G038972 | TSPAN7 | 0.932 | 0.001 | 1.000 | down |
| G038972 | CABP1 | 0.969 | 0.000 | 1.000 | down |
| LINC-PINT (2) | ELFN1 | 0.902 | 0.002 | 1.000 | down |
| LINC-PINT (2) | ATP2B4 | 0.945 | 0.000 | 1.000 | down |
| LINC-PINT (2) | PTCH1 | 0.902 | 0.002 | 1.000 | down |
| LOC101928322 | DAB1 | 0.951 | 0.000 | 1.000 | down |
| RP11-505K9.1 | ATP2B4 | 0.957 | 0.000 | 1.000 | down |
| RP11-505K9.1 | SPTB | 0.905 | 0.002 | 1.000 | down |
| RP11-505K9.1 | SEPSECS | 0.956 | 0.000 | 1.000 | down |
| RP11-505K9.1 | TSPAN7 | 0.974 | 0.000 | 1.000 | down |
| RP11-505K9.1 | CABP1 | 0.971 | 0.000 | 1.000 | down |
| RP11-792A8.4 | TENM4 | 0.901 | 0.002 | 1.000 | down |
| RP11-792A8.4 | HSP90AA1 | 0.903 | 0.002 | 1.000 | down |
| RP11-792A8.4 | SEPSECS | 0.916 | 0.001 | 1.000 | down |
| G063668 | HSP90AA1 | 0.918 | 0.001 | 1.000 | down |
| G063668 | ACTN1 | 0.922 | 0.001 | 1.000 | down |
| RP11-138E2.1 | CRELD1 | 0.945 | 0.000 | 1.000 | down |
| RP11-138E2.1 | ATP2B4 | 0.930 | 0.001 | 1.000 | down |
| RP11-138E2.1 | ERC2 | 0.931 | 0.001 | 1.000 | down |
| RP11-138E2.1 | PTPRN2 | 0.903 | 0.002 | 1.000 | down |
| RP11-138E2.1 | CABP1 | 0.903 | 0.002 | 1.000 | down |
| RP11-138E2.1 | DPYSL5 | 0.901 | 0.002 | 1.000 | down |
| RP11-775D22.3 | DNAJB1 | 0.944 | 0.000 | 1.000 | down |
| RP11-775D22.3 | SEPSECS | 0.908 | 0.002 | 1.000 | down |
| G003798 | ATP2B4 | 0.964 | 0.000 | 1.000 | down |
| G003798 | CABP1 | 0.933 | 0.001 | 1.000 | down |
| G053697 | ADGRL2 | 0.922 | 0.001 | 1.000 | down |
| G053697 | DAB1 | 0.925 | 0.001 | 1.000 | down |
| G053697 | GPSM2 | 0.916 | 0.001 | 1.000 | down |
| G006537 | CHRNB3 | 0.956 | 0.000 | 1.000 | down |
| G006537 | CRH | 0.923 | 0.001 | 1.000 | down |
| G066606 | MCTP2 | 0.910 | 0.002 | 1.000 | down |
| G066606 | PTCH1 | 0.900 | 0.002 | 1.000 | down |
| LOC730102 | DAB1 | 0.954 | 0.000 | 1.000 | down |
| LOC730102 | FILIP1 | 0.907 | 0.002 | 1.000 | down |

| | | | | | |
|---|---|---|---|---|---|
| LOC730102 | GPSM2 | 0.956 | 0.000 | 1.000 | down |
| AK026244 | ELFN1 | 0.902 | 0.002 | 1.000 | down |
| AK026244 | CABP1 | 0.933 | 0.001 | 1.000 | down |
| AK026244 | CHRNA7 | 0.948 | 0.000 | 1.000 | down |
| G059716 | CRELD1 | 0.940 | 0.001 | 1.000 | down |
| G059716 | DPYSL3 | 0.927 | 0.001 | 1.000 | down |
| RP11-2E11.5 | SEPSECS | 0.905 | 0.002 | 1.000 | down |
| RP11-2E11.5 | TSPAN7 | 0.949 | 0.000 | 1.000 | down |
| RP11-2E11.5 | CABP1 | 0.933 | 0.001 | 1.000 | down |
| LOC148709 | ATP2B4 | 0.901 | 0.002 | 1.000 | down |
| LOC148709 | SPTB | 0.914 | 0.001 | 1.000 | down |
| LOC148709 | SEPSECS | 0.903 | 0.002 | 1.000 | down |
| LOC148709 | TSPAN7 | 0.912 | 0.002 | 1.000 | down |
| LOC148709 | CABP1 | 0.949 | 0.000 | 1.000 | down |
| LOC148709 | CHRNA7 | 0.918 | 0.001 | 1.000 | down |
| G044581 | CRELD1 | 0.968 | 0.000 | 1.000 | down |
| G044581 | ATP2B4 | 0.955 | 0.000 | 1.000 | down |
| G044581 | ERC2 | 0.908 | 0.002 | 1.000 | down |
| G044581 | FILIP1 | 0.912 | 0.002 | 1.000 | down |
| G044581 | SEPSECS | 0.927 | 0.001 | 1.000 | down |
| G044581 | CABP1 | 0.916 | 0.001 | 1.000 | down |
| G019668 | PTCH1 | 0.915 | 0.001 | 1.000 | down |
| RP5-884G6.2 | ATP2B4 | 0.934 | 0.001 | 1.000 | down |
| PVRL3-AS1 | CRELD1 | 0.924 | 0.001 | 1.000 | down |
| PVRL3-AS1 | CABP1 | 0.906 | 0.002 | 1.000 | down |
| AC114803.3 | EXOC7 | 0.915 | 0.001 | 1.000 | down |
| AC114803.3 | ACTN1 | 0.950 | 0.000 | 1.000 | down |
| AC099850.1 | ERC2 | 0.926 | 0.001 | 1.000 | down |
| AC099850.1 | PTPRN2 | 0.943 | 0.000 | 1.000 | down |
| AC099850.1 | MAP2 | 0.912 | 0.002 | 1.000 | down |
| G010782 | ADGRL2 | 0.935 | 0.001 | 1.000 | down |
| BC042589 | CRELD1 | 0.957 | 0.000 | 1.000 | down |
| BC042589 | HSP90AA1 | 0.900 | 0.002 | 1.000 | down |
| BC042589 | ERC2 | 0.944 | 0.000 | 1.000 | down |
| BC042589 | FILIP1 | 0.941 | 0.000 | 1.000 | down |
| BC042589 | SEPSECS | 0.917 | 0.001 | 1.000 | down |
| LINC00634 | CRELD1 | 0.955 | 0.000 | 1.000 | down |
| LINC00634 | ERC2 | 0.961 | 0.000 | 1.000 | down |
| LINC00634 | GPSM2 | 0.948 | 0.000 | 1.000 | down |
| LINC00634 | CALB2 | 0.905 | 0.002 | 1.000 | down |
| LINC00634 | PTPRN2 | 0.928 | 0.001 | 1.000 | down |
| LINC00634 | MAP2 | 0.930 | 0.001 | 1.000 | down |
| LINC00634 | DPYSL5 | 0.958 | 0.000 | 1.000 | down |
| LINC00634 | DPYSL3 | 0.926 | 0.001 | 1.000 | down |
| LINC00634 | PPP2R1A | 0.932 | 0.001 | 1.000 | down |
| CTD-2383M3.1 | ATP2B4 | 0.914 | 0.001 | 1.000 | down |
| CTD-2383M3.1 | CABP1 | 0.932 | 0.001 | 1.000 | down |
| AC092667.2 | ATP2B4 | 0.935 | 0.001 | 1.000 | down |
| AC092667.2 | ERC2 | 0.903 | 0.002 | 1.000 | down |
| AC092667.2 | PTPRN2 | 0.900 | 0.002 | 1.000 | down |
| AC092667.2 | SEPSECS | 0.916 | 0.001 | 1.000 | down |
| AC092667.2 | TSPAN7 | 0.938 | 0.001 | 1.000 | down |
| AC092667.2 | CABP1 | 0.961 | 0.000 | 1.000 | down |
| GYG2P1 | AP1G2 | 0.928 | 0.001 | 1.000 | down |
| GYG2P1 | SPTB | 0.938 | 0.001 | 1.000 | down |
| RP11-376P6.3 | ELFN1 | 0.917 | 0.001 | 1.000 | down |
| RP11-376P6.3 | CABP1 | 0.955 | 0.000 | 1.000 | down |
| RP11-376P6.3 | CHRNA7 | 0.917 | 0.001 | 1.000 | down |
| FGF13-AS1 | AP1G2 | 0.914 | 0.001 | 1.000 | down |
| FGF13-AS1 | SPTB | 0.940 | 0.001 | 1.000 | down |
| RP11-383C5.5 | CRELD1 | 0.916 | 0.001 | 1.000 | down |
| RP11-383C5.5 | ELFN1 | 0.907 | 0.002 | 1.000 | down |
| RP11-383C5.5 | ATP2B4 | 0.965 | 0.000 | 1.000 | down |
| RP11-383C5.5 | SEPSECS | 0.924 | 0.001 | 1.000 | down |
| RP11-383C5.5 | CABP1 | 0.967 | 0.000 | 1.000 | down |

| | | | | | |
|---|---|---|---|---|---|
| LOC644656 | ATP2B4 | 0.924 | 0.001 | 1.000 | down |
| G001739 | CRELD1 | 0.925 | 0.001 | 1.000 | down |
| G001739 | ELFN1 | 0.909 | 0.002 | 1.000 | down |
| G001739 | ATP2B4 | 0.960 | 0.000 | 1.000 | down |
| G001739 | CABP1 | 0.906 | 0.002 | 1.000 | down |
| AC005220.3 | PTCH1 | 0.916 | 0.001 | 1.000 | down |
| AC005220.3 | FILIP1 | 0.927 | 0.001 | 1.000 | down |
| AC005220.3 | SEPSECS | 0.927 | 0.001 | 1.000 | down |
| RP11-22N19.2 | CRELD1 | 0.938 | 0.001 | 1.000 | down |
| RP11-22N19.2 | ATP2B4 | 0.954 | 0.000 | 1.000 | down |
| RP11-22N19.2 | ERC2 | 0.951 | 0.000 | 1.000 | down |
| RP11-22N19.2 | FILIP1 | 0.906 | 0.002 | 1.000 | down |
| RP11-51J9.5 | ERC2 | 0.923 | 0.001 | 1.000 | down |
| RP11-51J9.5 | GPSM2 | 0.941 | 0.000 | 1.000 | down |
| RP11-51J9.5 | MAP2 | 0.900 | 0.002 | 1.000 | down |
| RP11-51J9.5 | PPP2R1A | 0.966 | 0.000 | 1.000 | down |
| LOC102724661 | OLFM1 | 0.930 | 0.001 | 1.000 | down |
| LOC102724661 | FXYD6 | 0.923 | 0.001 | 1.000 | down |
| LINC01023 | ELFN1 | 0.909 | 0.002 | 1.000 | down |
| uc.95 | ACTN1 | 0.909 | 0.002 | 1.000 | down |
| LOC101928402 | ATP2B4 | 0.912 | 0.002 | 1.000 | down |
| LOC101928402 | SEPSECS | 0.913 | 0.002 | 1.000 | down |
| LOC101928402 | TSPAN7 | 0.942 | 0.000 | 1.000 | down |
| LOC101928402 | CABP1 | 0.933 | 0.001 | 1.000 | down |
| GFOD1 | ATP2B4 | 0.938 | 0.001 | 1.000 | down |
| GFOD1 | TSPAN7 | 0.906 | 0.002 | 1.000 | down |
| GFOD1 | CABP1 | 0.944 | 0.000 | 1.000 | down |
| LOC101928417 | ELFN1 | 0.914 | 0.002 | 1.000 | down |
| LOC101928417 | ATP2B4 | 0.926 | 0.001 | 1.000 | down |
| LOC101928417 | CABP1 | 0.937 | 0.001 | 1.000 | down |
| GPR50-AS1 | DNAJB1 | 0.925 | 0.001 | 1.000 | down |
| FAM138E | CRH | 0.904 | 0.002 | 1.000 | down |
| FAM138E | SLC17A8 | 0.914 | 0.002 | 1.000 | down |
| G078120 | CALB2 | 0.905 | 0.002 | 1.000 | down |
| RP4-616B8.4 | ATP2B4 | 0.937 | 0.001 | 1.000 | down |
| RP4-616B8.4 | ERC2 | 0.921 | 0.001 | 1.000 | down |
| RP4-616B8.4 | TSPAN7 | 0.903 | 0.002 | 1.000 | down |
| RP4-555D20.3 | CRELD1 | 0.956 | 0.000 | 1.000 | down |
| RP4-555D20.3 | ATP2B4 | 0.934 | 0.001 | 1.000 | down |
| RP4-555D20.3 | ERC2 | 0.922 | 0.001 | 1.000 | down |
| RP4-555D20.3 | SEPSECS | 0.940 | 0.001 | 1.000 | down |
| RP4-555D20.3 | CABP1 | 0.935 | 0.001 | 1.000 | down |
| RP11-507K2.3 | CRELD1 | 0.902 | 0.002 | 1.000 | down |
| RP11-507K2.3 | ATP2B4 | 0.979 | 0.000 | 1.000 | down |
| DQ594967 | SEPSECS | 0.913 | 0.002 | 1.000 | down |
| DQ594967 | TSPAN7 | 0.958 | 0.000 | 1.000 | down |
| TOB1-AS1 (1) | CRELD1 | 0.948 | 0.000 | 1.000 | down |
| AP000255.6 | SEPSECS | 0.957 | 0.000 | 1.000 | down |
| AP000255.6 | TSPAN7 | 0.917 | 0.001 | 1.000 | down |
| AP000255.6 | CABP1 | 0.935 | 0.001 | 1.000 | down |
| AP000255.6 | RPS6KA5 | 0.909 | 0.002 | 1.000 | down |
| AK000953 | TENM4 | 0.915 | 0.001 | 1.000 | down |
| AK000953 | SPTB | 0.914 | 0.002 | 1.000 | down |
| AK000953 | RPS6KA5 | 0.938 | 0.001 | 1.000 | down |
| RP4-798A10.7 | CRELD1 | 0.928 | 0.001 | 1.000 | down |
| RP4-798A10.7 | ATP2B4 | 0.953 | 0.000 | 1.000 | down |
| RP4-798A10.7 | ERC2 | 0.950 | 0.000 | 1.000 | down |
| RP4-798A10.7 | PTPRN2 | 0.905 | 0.002 | 1.000 | down |
| RP4-798A10.7 | MAP2 | 0.922 | 0.001 | 1.000 | down |
| RP4-798A10.7 | DPYSL5 | 0.917 | 0.001 | 1.000 | down |
| CTD-2377O17.1 | ATP2B4 | 0.948 | 0.000 | 1.000 | down |
| CTD-2377O17.1 | SPTB | 0.909 | 0.002 | 1.000 | down |
| CTD-2377O17.1 | SEPSECS | 0.931 | 0.001 | 1.000 | down |
| CTD-2377O17.1 | TSPAN7 | 0.943 | 0.000 | 1.000 | down |
| CTD-2377O17.1 | CABP1 | 0.982 | 0.000 | 1.000 | down |

| | | | | | |
|---|---|---|---|---|---|
| LINC00630 | ERC2 | 0.930 | 0.001 | 1.000 | down |
| LINC00630 | MAP2 | 0.911 | 0.002 | 1.000 | down |
| G049105 | HSP90AA1 | 0.905 | 0.002 | 1.000 | down |
| RP11-254F7.2 | ATP2B4 | 0.910 | 0.002 | 1.000 | down |
| RP11-254F7.2 | ERC2 | 0.946 | 0.000 | 1.000 | down |
| RP11-254F7.2 | CALB2 | 0.905 | 0.002 | 1.000 | down |
| RP11-254F7.2 | SEPSECS | 0.975 | 0.000 | 1.000 | down |
| RP11-254F7.2 | TSPAN7 | 0.940 | 0.001 | 1.000 | down |
| G031140 | PHB | 0.914 | 0.002 | 1.000 | down |
| uc.76 | ARHGAP32 | 0.952 | 0.000 | 1.000 | down |
| PRR36 | AP1G2 | 0.907 | 0.002 | 1.000 | down |
| LL22NC03-86G7.1 | CALB2 | 0.906 | 0.002 | 1.000 | down |
| LL22NC03-86G7.1 | DPYSL3 | 0.927 | 0.001 | 1.000 | down |
| RP11-332H14.1 | SPTB | 0.925 | 0.001 | 1.000 | down |
| RP11-332H14.1 | SEPSECS | 0.916 | 0.001 | 1.000 | down |
| RP11-332H14.1 | TSPAN7 | 0.950 | 0.000 | 1.000 | down |
| RP11-332H14.1 | CABP1 | 0.986 | 0.000 | 1.000 | down |
| ULK4P1 | MCTP2 | 0.937 | 0.001 | 1.000 | down |
| G066437 | DAB1 | 0.947 | 0.000 | 1.000 | down |
| G066437 | ERC2 | 0.903 | 0.002 | 1.000 | down |
| G066437 | BTBD9 | 0.909 | 0.002 | 1.000 | down |
| G066437 | PPP2R1A | 0.920 | 0.001 | 1.000 | down |
| G057325 | TENM4 | 0.905 | 0.002 | 1.000 | down |
| G057325 | ACTN1 | 0.959 | 0.000 | 1.000 | down |
| ENO1-AS1 | HSP90AA1 | 0.948 | 0.000 | 1.000 | down |
| ENO1-AS1 | CHRNA3 | 0.952 | 0.000 | 1.000 | down |
| ENO1-AS1 | ERC2 | 0.902 | 0.002 | 1.000 | down |
| ENO1-AS1 | SEPSECS | 0.901 | 0.002 | 1.000 | down |
| RP1-261G23.7 | ATP2B4 | 0.912 | 0.002 | 1.000 | down |
| RP1-261G23.7 | TSPAN7 | 0.916 | 0.001 | 1.000 | down |
| RP1-261G23.7 | CABP1 | 0.971 | 0.000 | 1.000 | down |
| G023087 | ATP2B4 | 0.931 | 0.001 | 1.000 | down |
| G023087 | SEPSECS | 0.904 | 0.002 | 1.000 | down |
| G023087 | TSPAN7 | 0.905 | 0.002 | 1.000 | down |
| G023087 | CABP1 | 0.951 | 0.000 | 1.000 | down |
| RP11-1275H24.1 | TSPAN7 | 0.958 | 0.000 | 1.000 | down |
| RP11-1275H24.1 | CABP1 | 0.940 | 0.001 | 1.000 | down |
| RP1-223E5.4 | CRELD1 | 0.950 | 0.000 | 1.000 | down |
| RP1-223E5.4 | ATP2B4 | 0.913 | 0.002 | 1.000 | down |
| RP1-223E5.4 | ERC2 | 0.944 | 0.000 | 1.000 | down |
| RP1-223E5.4 | PTPRN2 | 0.923 | 0.001 | 1.000 | down |
| RP1-223E5.4 | DPYSL5 | 0.924 | 0.001 | 1.000 | down |
| LINC01152 | OLFM1 | 0.940 | 0.001 | 1.000 | down |
| G037470 | CABP1 | 0.947 | 0.000 | 1.000 | down |
| G037470 | CHRNA7 | 0.913 | 0.002 | 1.000 | down |
| RP11-680F8.1 | SIPA1L2 | 0.929 | 0.001 | 1.000 | down |
| RP11-680F8.1 | ALDOC | 0.985 | 0.000 | 1.000 | down |
| DGCR5 | HSP90AA1 | 0.944 | 0.000 | 1.000 | down |
| DGCR5 | ERC2 | 0.920 | 0.001 | 1.000 | down |
| DGCR5 | PPP2R1A | 0.934 | 0.001 | 1.000 | down |
| RP11-413M3.4 | BTBD9 | 0.965 | 0.000 | 1.000 | down |
| FAM66D | DPYSL3 | 0.983 | 0.000 | 1.000 | down |
| LINC00463 | CALB2 | 0.918 | 0.001 | 1.000 | down |
| G010838 | ARHGAP32 | 0.937 | 0.001 | 1.000 | down |
| G010838 | ATP2B4 | 0.924 | 0.001 | 1.000 | down |
| G010838 | ERC2 | 0.936 | 0.001 | 1.000 | down |
| G010838 | PTPRN2 | 0.942 | 0.000 | 1.000 | down |
| G010838 | MAP2 | 0.937 | 0.001 | 1.000 | down |
| G027543 | HSP90AA1 | 0.941 | 0.000 | 1.000 | down |
| G027543 | CHRNA3 | 0.902 | 0.002 | 1.000 | down |
| G027543 | CALB2 | 0.928 | 0.001 | 1.000 | down |
| RP11-573G6.8 | CRELD1 | 0.916 | 0.001 | 1.000 | down |
| RP11-573G6.8 | ARHGAP32 | 0.910 | 0.002 | 1.000 | down |
| RP11-573G6.8 | ATP2B4 | 0.923 | 0.001 | 1.000 | down |
| RP11-573G6.8 | ERC2 | 0.976 | 0.000 | 1.000 | down |

| | | | | | |
|---|---|---|---|---|---|
| RP11-573G6.8 | PTPRN2 | 0.969 | 0.000 | 1.000 | down |
| RP11-573G6.8 | MAP2 | 0.961 | 0.000 | 1.000 | down |
| RP11-573G6.8 | DPYSL5 | 0.955 | 0.000 | 1.000 | down |
| XLOC_007467 | FILIP1 | 0.952 | 0.000 | 1.000 | down |
| XLOC_007467 | SEPSECS | 0.915 | 0.001 | 1.000 | down |
| RP11-107N15.1 | ELFN1 | 0.947 | 0.000 | 1.000 | down |
| RP11-107N15.1 | CABP1 | 0.918 | 0.001 | 1.000 | down |
| LL09NC01-139C3.1 | CRELD1 | 0.953 | 0.000 | 1.000 | down |
| LL09NC01-139C3.1 | ARHGAP32 | 0.904 | 0.002 | 1.000 | down |
| LL09NC01-139C3.1 | ATP2B4 | 0.923 | 0.001 | 1.000 | down |
| LL09NC01-139C3.1 | ERC2 | 0.972 | 0.000 | 1.000 | down |
| LL09NC01-139C3.1 | CALB2 | 0.904 | 0.002 | 1.000 | down |
| LL09NC01-139C3.1 | PTPRN2 | 0.941 | 0.000 | 1.000 | down |
| LL09NC01-139C3.1 | SEPSECS | 0.921 | 0.001 | 1.000 | down |
| LL09NC01-139C3.1 | MAP2 | 0.931 | 0.001 | 1.000 | down |
| LL09NC01-139C3.1 | DPYSL5 | 0.924 | 0.001 | 1.000 | down |
| G061722 | CHRNA3 | 0.907 | 0.002 | 1.000 | down |
| G046707 | SOS1 | 0.901 | 0.002 | 1.000 | down |
| G046707 | AP1G2 | 0.904 | 0.002 | 1.000 | down |
| LOC100288846 | ATP2B4 | 0.980 | 0.000 | 1.000 | down |
| LOC100288846 | ERC2 | 0.902 | 0.002 | 1.000 | down |
| LOC100288846 | SEPSECS | 0.937 | 0.001 | 1.000 | down |
| LOC100288846 | TSPAN7 | 0.949 | 0.000 | 1.000 | down |
| LOC100288846 | CABP1 | 0.967 | 0.000 | 1.000 | down |
| RP11-798M19.6 | ADGRL2 | 0.902 | 0.002 | 1.000 | down |
| RP11-798M19.6 | SIPA1L2 | 0.934 | 0.001 | 1.000 | down |
| RP11-798M19.6 | GPSM2 | 0.936 | 0.001 | 1.000 | down |
| RP11-798M19.6 | DPYSL3 | 0.925 | 0.001 | 1.000 | down |
| G068576 | FXYD6 | 0.943 | 0.000 | 1.000 | down |
| RP11-872D17.4 | EXOC7 | 0.900 | 0.002 | 1.000 | down |
| G036568 | MCTP2 | 0.946 | 0.000 | 1.000 | down |
| CXADRP3 | DAB1 | 0.914 | 0.001 | 1.000 | down |
| CXADRP3 | PTCH1 | 0.961 | 0.000 | 1.000 | down |
| CXADRP3 | ERC2 | 0.921 | 0.001 | 1.000 | down |
| CXADRP3 | BTBD9 | 0.908 | 0.002 | 1.000 | down |
| CXADRP3 | PPP2R1A | 0.906 | 0.002 | 1.000 | down |
| UBAC2-AS1 | ATP2B4 | 0.914 | 0.001 | 1.000 | down |
| UBAC2-AS1 | SEPSECS | 0.916 | 0.001 | 1.000 | down |
| UBAC2-AS1 | TSPAN7 | 0.928 | 0.001 | 1.000 | down |
| UBAC2-AS1 | CABP1 | 0.905 | 0.002 | 1.000 | down |
| CTD-2527I21.7 | SOS1 | 0.922 | 0.001 | 1.000 | down |
| CTD-2527I21.7 | CHRNB3 | 0.946 | 0.000 | 1.000 | down |
| XLOC_005289 | EXOC7 | 0.930 | 0.001 | 1.000 | down |
| RP11-177G23.2 | CHRNA7 | 0.959 | 0.000 | 1.000 | down |
| G000672 | SEPSECS | 0.913 | 0.002 | 1.000 | down |
| CTD-2382E5.2 | SOS1 | 0.960 | 0.000 | 1.000 | down |
| CTD-2382E5.2 | CHRNB3 | 0.938 | 0.001 | 1.000 | down |
| CTD-2382E5.2 | CHRNA3 | 0.931 | 0.001 | 1.000 | down |
| EZR-AS1 | ADGRL2 | 0.917 | 0.001 | 1.000 | down |
| RP1-86C11.7 | CRELD1 | 0.954 | 0.000 | 1.000 | down |
| RP1-86C11.7 | SEPSECS | 0.900 | 0.002 | 1.000 | down |
| RP1-86C11.7 | CABP1 | 0.913 | 0.002 | 1.000 | down |
| G081642 | SOS1 | 0.925 | 0.001 | 1.000 | down |
| G081642 | EXOC7 | 0.951 | 0.000 | 1.000 | down |
| G081642 | CHRNA3 | 0.924 | 0.001 | 1.000 | down |
| G058460 | OLFM1 | 0.970 | 0.000 | 1.000 | down |
| CTD-2653B5.1 | GPSM2 | 0.917 | 0.001 | 1.000 | down |
| CTD-2653B5.1 | PPP2R1A | 0.903 | 0.002 | 1.000 | down |
| G047565 | OLFM1 | 0.909 | 0.002 | 1.000 | down |
| G047565 | BTBD9 | 0.928 | 0.001 | 1.000 | down |
| G064611 | MCTP2 | 0.906 | 0.002 | 1.000 | down |
| G064611 | FXYD6 | 0.932 | 0.001 | 1.000 | down |
| FLJ20021 | CRELD1 | 0.949 | 0.000 | 1.000 | down |
| FLJ20021 | ATP2B4 | 0.908 | 0.002 | 1.000 | down |
| FLJ20021 | FILIP1 | 0.902 | 0.002 | 1.000 | down |

| | | | | | |
|---|---|---|---|---|---|
| FLJ20021 | SEPSECS | 0.925 | 0.001 | 1.000 | down |
| FLJ20021 | CABP1 | 0.917 | 0.001 | 1.000 | down |
| FLJ20021 | RPS6KA5 | 0.920 | 0.001 | 1.000 | down |
| uc.247 | UNC13C | 0.942 | 0.000 | 1.000 | down |
| RP11-109D9.4 | CRELD1 | 0.906 | 0.002 | 1.000 | down |
| RP11-109D9.4 | ATP2B4 | 0.901 | 0.002 | 1.000 | down |
| RP11-109D9.4 | PTCH1 | 0.912 | 0.002 | 1.000 | down |
| RP11-109D9.4 | SEPSECS | 0.934 | 0.001 | 1.000 | down |
| RP11-110G21.1 | CALB2 | 0.918 | 0.001 | 1.000 | down |
| CTC-444N24.13 | CRELD1 | 0.976 | 0.000 | 1.000 | down |
| CTC-444N24.13 | ATP2B4 | 0.933 | 0.001 | 1.000 | down |
| CTC-444N24.13 | DPYSL3 | 0.905 | 0.002 | 1.000 | down |
| RP11-339B21.10 | NRP2 | 0.918 | 0.001 | 1.000 | down |
| RP11-339B21.10 | SPTB | 0.904 | 0.002 | 1.000 | down |
| RP11-339B21.10 | RPS6KA5 | 0.913 | 0.002 | 1.000 | down |
| LINC00449 | ARHGAP32 | 0.933 | 0.001 | 1.000 | down |
| LINC00449 | ATP2B4 | 0.944 | 0.000 | 1.000 | down |
| LINC00449 | PTCH1 | 0.952 | 0.000 | 1.000 | down |
| RP11-61L19.2 | CRELD1 | 0.925 | 0.001 | 1.000 | down |
| RP11-61L19.2 | ELFN1 | 0.909 | 0.002 | 1.000 | down |
| RP11-61L19.2 | ATP2B4 | 0.978 | 0.000 | 1.000 | down |
| RP11-61L19.2 | CABP1 | 0.940 | 0.001 | 1.000 | down |
| LOC101929066 | ARHGAP32 | 0.945 | 0.000 | 1.000 | down |
| LOC101929066 | ATP2B4 | 0.968 | 0.000 | 1.000 | down |
| G054532 | DAB1 | 0.918 | 0.001 | 1.000 | down |
| G054532 | DNAJB1 | 0.949 | 0.000 | 1.000 | down |
| RP11-85A1.3 | CRELD1 | 0.918 | 0.001 | 1.000 | down |
| RP11-85A1.3 | ATP2B4 | 0.932 | 0.001 | 1.000 | down |
| RP11-85A1.3 | CABP1 | 0.932 | 0.001 | 1.000 | down |
| G061596 | ATP2B4 | 0.907 | 0.002 | 1.000 | down |
| G061596 | ERC2 | 0.923 | 0.001 | 1.000 | down |
| G061596 | PTPRN2 | 0.916 | 0.001 | 1.000 | down |
| AK125078 | SOS1 | 0.919 | 0.001 | 1.000 | down |
| AK125078 | SIPA1L2 | 0.918 | 0.001 | 1.000 | down |
| AK125078 | CHRNA3 | 0.902 | 0.002 | 1.000 | down |
| DHRS4-AS1 (2) | ARHGAP32 | 0.931 | 0.001 | 1.000 | down |
| BX248253 | NRP2 | 0.951 | 0.000 | 1.000 | down |
| BX248253 | SPTB | 0.909 | 0.002 | 1.000 | down |
| BX248253 | SEPSECS | 0.916 | 0.001 | 1.000 | down |
| RP11-344P13.6 | CRELD1 | 0.972 | 0.000 | 1.000 | down |
| RP11-344P13.6 | ATP2B4 | 0.921 | 0.001 | 1.000 | down |
| RP11-344P13.6 | DPYSL3 | 0.923 | 0.001 | 1.000 | down |
| LOC101927822 | HSPA8 | 0.906 | 0.002 | 1.000 | down |
| LOC101927822 | CRH | 0.948 | 0.000 | 1.000 | down |
| G070405 | CRELD1 | 0.922 | 0.001 | 1.000 | down |
| G070405 | ELFN1 | 0.914 | 0.002 | 1.000 | down |
| RP11-565F19.2 | TSPAN7 | 0.955 | 0.000 | 1.000 | down |
| RP11-565F19.2 | CABP1 | 0.915 | 0.001 | 1.000 | down |
| G054079 | PTCH1 | 0.900 | 0.002 | 1.000 | down |
| ANKRD20A12P | CRELD1 | 0.919 | 0.001 | 1.000 | down |
| ANKRD20A12P | HSP90AA1 | 0.917 | 0.001 | 1.000 | down |
| ANKRD20A12P | ADGRL2 | 0.915 | 0.001 | 1.000 | down |
| ANKRD20A12P | DPYSL3 | 0.950 | 0.000 | 1.000 | down |
| RP11-53I6.3 | ERC2 | 0.908 | 0.002 | 1.000 | down |
| RP11-53I6.3 | SPTB | 0.919 | 0.001 | 1.000 | down |
| RP11-53I6.3 | SEPSECS | 0.953 | 0.000 | 1.000 | down |
| RP11-53I6.3 | TSPAN7 | 0.928 | 0.001 | 1.000 | down |
| RP11-53I6.3 | CABP1 | 0.929 | 0.001 | 1.000 | down |
| RP11-483I13.5 | ARHGAP32 | 0.905 | 0.002 | 1.000 | down |
| RP11-483I13.5 | PTPRN2 | 0.915 | 0.001 | 1.000 | down |
| RP11-314A20.5 | DNAJB1 | 0.914 | 0.001 | 1.000 | down |
| RP11-314A20.5 | ERC2 | 0.915 | 0.001 | 1.000 | down |
| RP11-314A20.5 | CALB2 | 0.953 | 0.000 | 1.000 | down |
| AK055974 | TSPAN7 | 0.916 | 0.001 | 1.000 | down |
| AK055974 | CABP1 | 0.918 | 0.001 | 1.000 | down |

| | | | | | |
|---|---|---|---|---|---|
| G060549 | CABP1 | 0.904 | 0.002 | 1.000 | down |
| G060549 | CHRNA7 | 0.921 | 0.001 | 1.000 | down |
| RP11-474O21.5 | SPTB | 0.912 | 0.002 | 1.000 | down |
| G046528 | SPTB | 0.902 | 0.002 | 1.000 | down |
| CRYBB2P1 | AP1G2 | 0.917 | 0.001 | 1.000 | down |
| RP11-196G18.24 | ATP2B4 | 0.915 | 0.001 | 1.000 | down |
| RP11-196G18.24 | ERC2 | 0.959 | 0.000 | 1.000 | down |
| RP11-196G18.24 | CALB2 | 0.915 | 0.001 | 1.000 | down |
| RP11-196G18.24 | SEPSECS | 0.913 | 0.002 | 1.000 | down |
| RP11-196G18.24 | TSPAN7 | 0.921 | 0.001 | 1.000 | down |
| RP11-196G18.24 | MAP2 | 0.906 | 0.002 | 1.000 | down |
| LOC101927811 | DAB1 | 0.925 | 0.001 | 1.000 | down |
| LOC101927811 | DNAJB1 | 0.941 | 0.001 | 1.000 | down |
| LOC101927811 | PTCH1 | 0.911 | 0.002 | 1.000 | down |
| LOC101927811 | SEPSECS | 0.905 | 0.002 | 1.000 | down |
| CTD-2623N2.11 | ATP2B4 | 0.941 | 0.000 | 1.000 | down |
| CTD-2623N2.11 | ERC2 | 0.902 | 0.002 | 1.000 | down |
| CTD-2623N2.11 | PTPRN2 | 0.901 | 0.002 | 1.000 | down |
| CTD-2623N2.11 | CABP1 | 0.946 | 0.000 | 1.000 | down |
| RP11-254F7.1 | CHRNB3 | 0.905 | 0.002 | 1.000 | down |
| uc.412 | HSPA8 | 0.912 | 0.002 | 1.000 | down |
| uc.412 | DAB1 | 0.910 | 0.002 | 1.000 | down |
| uc.412 | DNAJB1 | 0.913 | 0.002 | 1.000 | down |
| uc.412 | CRH | 0.959 | 0.000 | 1.000 | down |
| CTC-350I8.1 | EXOC7 | 0.920 | 0.001 | 1.000 | down |
| G023340 | ERC2 | 0.962 | 0.000 | 1.000 | down |
| G023340 | CALB2 | 0.922 | 0.001 | 1.000 | down |
| G023340 | PTPRN2 | 0.980 | 0.000 | 1.000 | down |
| G023340 | MAP2 | 0.952 | 0.000 | 1.000 | down |
| G023340 | DPYSL5 | 0.966 | 0.000 | 1.000 | down |
| H1FX-AS1 | SOS1 | 0.908 | 0.002 | 1.000 | down |
| H1FX-AS1 | CHRNB3 | 0.904 | 0.002 | 1.000 | down |
| KB-318B8.7 | GPSM2 | 0.912 | 0.002 | 1.000 | down |
| G063514 | TENM4 | 0.925 | 0.001 | 1.000 | down |
| G063514 | AP1G2 | 0.927 | 0.001 | 1.000 | down |
| G063514 | NRP2 | 0.919 | 0.001 | 1.000 | down |
| G063514 | SPTB | 0.968 | 0.000 | 1.000 | down |
| RP6-109B7.4 | ARHGAP32 | 0.926 | 0.001 | 1.000 | down |
| G070622 | PHB | 0.911 | 0.002 | 1.000 | down |
| RP11-30L3.2 | BTBD9 | 0.958 | 0.000 | 1.000 | down |
| RP11-30L3.2 | PPP2R1A | 0.936 | 0.001 | 1.000 | down |
| G014106 | CRELD1 | 0.907 | 0.002 | 1.000 | down |
| G014106 | ADGRL2 | 0.937 | 0.001 | 1.000 | down |
| G014106 | DAB1 | 0.929 | 0.001 | 1.000 | down |
| G014106 | BTBD9 | 0.903 | 0.002 | 1.000 | down |
| G014106 | PPP2R1A | 0.902 | 0.002 | 1.000 | down |
| RP1-232L24.3 | ATP2B4 | 0.926 | 0.001 | 1.000 | down |
| RP1-232L24.3 | SEPSECS | 0.960 | 0.000 | 1.000 | down |
| RP1-232L24.3 | TSPAN7 | 0.952 | 0.000 | 1.000 | down |
| RP1-232L24.3 | CABP1 | 0.950 | 0.000 | 1.000 | down |
| RP11-362L22.1 | HSPA8 | 0.912 | 0.002 | 1.000 | down |
| RP11-362L22.1 | FXYD6 | 0.914 | 0.001 | 1.000 | down |
| RP11-467P9.1 | ATP2B4 | 0.940 | 0.001 | 1.000 | down |
| RP11-467P9.1 | SEPSECS | 0.949 | 0.000 | 1.000 | down |
| XXbac-B476C20.9 | ATP2B4 | 0.936 | 0.001 | 1.000 | down |
| XXbac-B476C20.9 | PTCH1 | 0.917 | 0.001 | 1.000 | down |
| G002695 | CRELD1 | 0.944 | 0.000 | 1.000 | down |
| G002695 | ELFN1 | 0.917 | 0.001 | 1.000 | down |
| G021588 | ATP2B4 | 0.929 | 0.001 | 1.000 | down |
| G021588 | PTCH1 | 0.935 | 0.001 | 1.000 | down |
| G021588 | SEPSECS | 0.926 | 0.001 | 1.000 | down |
| G006578 | CRELD1 | 0.940 | 0.001 | 1.000 | down |
| G006578 | DPYSL5 | 0.910 | 0.002 | 1.000 | down |
| G006578 | DPYSL3 | 0.931 | 0.001 | 1.000 | down |
| RP11-371A19.2 | ATP2B4 | 0.945 | 0.000 | 1.000 | down |

| | | | | | |
|---|---|---|---|---|---|
| RP11-371A19.2 | TSPAN7 | 0.928 | 0.001 | 1.000 | down |
| RP11-371A19.2 | CABP1 | 0.965 | 0.000 | 1.000 | down |
| RP11-95P2.3 | TENM4 | 0.925 | 0.001 | 1.000 | down |
| RP11-95P2.3 | SPTB | 0.928 | 0.001 | 1.000 | down |
| RP11-95P2.3 | SEPSECS | 0.957 | 0.000 | 1.000 | down |
| RP11-118K6.3 | NRP2 | 0.909 | 0.002 | 1.000 | down |
| RP11-118K6.3 | CHRNA7 | 0.909 | 0.002 | 1.000 | down |
| G061238 | HSPA8 | 0.911 | 0.002 | 1.000 | down |
| G061238 | BTBD9 | 0.953 | 0.000 | 1.000 | down |
| RP11-895M11.3 | CHRNA7 | 0.908 | 0.002 | 1.000 | down |
| LOC101928489 | CRELD1 | 0.944 | 0.000 | 1.000 | down |
| RP11-355F22.1 | PHB | 0.903 | 0.002 | 1.000 | down |
| RP11-355F22.1 | CALB2 | 0.926 | 0.001 | 1.000 | down |
| JARID2-AS1 | DNAJB1 | 0.915 | 0.001 | 1.000 | down |
| G036604 | FXYD6 | 0.913 | 0.002 | 1.000 | down |
| G075319 | CRELD1 | 0.936 | 0.001 | 1.000 | down |
| G075319 | FILIP1 | 0.927 | 0.001 | 1.000 | down |
| G075319 | GPSM2 | 0.925 | 0.001 | 1.000 | down |
| G075319 | RPS6KA5 | 0.900 | 0.002 | 1.000 | down |
| G075319 | DPYSL3 | 0.952 | 0.000 | 1.000 | down |
| XLOC_010219 | EXOC7 | 0.907 | 0.002 | 1.000 | down |
| XLOC_010219 | MCTP2 | 0.954 | 0.000 | 1.000 | down |
| GSE61474_XLOC_0676( | RPS6KA5 | 0.982 | 0.000 | 1.000 | down |
| LOC283140 | DNAJB1 | 0.959 | 0.000 | 1.000 | down |
| LOC283140 | PTCH1 | 0.903 | 0.002 | 1.000 | down |
| DNAJC9-AS1 | CHRNA3 | 0.947 | 0.000 | 1.000 | down |
| RP11-820I16.4 | CRELD1 | 0.909 | 0.002 | 1.000 | down |
| RP11-820I16.4 | ATP2B4 | 0.962 | 0.000 | 1.000 | down |
| RP11-820I16.4 | CABP1 | 0.938 | 0.001 | 1.000 | down |
| RP11-79P5.9 | CRELD1 | 0.928 | 0.001 | 1.000 | down |
| RP11-79P5.9 | PTPRN2 | 0.905 | 0.002 | 1.000 | down |
| RP11-79P5.9 | DPYSL5 | 0.918 | 0.001 | 1.000 | down |
| RP11-79P5.9 | DPYSL3 | 0.936 | 0.001 | 1.000 | down |
| G057611 | HSPA8 | 0.905 | 0.002 | 1.000 | down |
| G057611 | DNAJB1 | 0.914 | 0.001 | 1.000 | down |
| RP11-166B2.3 | CRELD1 | 0.936 | 0.001 | 1.000 | down |
| RP11-166B2.3 | DAB1 | 0.913 | 0.002 | 1.000 | down |
| RP11-166B2.3 | ERC2 | 0.924 | 0.001 | 1.000 | down |
| RP11-166B2.3 | FILIP1 | 0.943 | 0.000 | 1.000 | down |
| RP11-166B2.3 | GPSM2 | 0.920 | 0.001 | 1.000 | down |
| RP11-168O10.6 | EXOC7 | 0.922 | 0.001 | 1.000 | down |
| RP11-168O10.6 | AP1G2 | 0.926 | 0.001 | 1.000 | down |
| LOC100270746 | CRELD1 | 0.918 | 0.001 | 1.000 | down |
| LOC100270746 | HSP90AA1 | 0.908 | 0.002 | 1.000 | down |
| G075703 | ELFN1 | 0.905 | 0.002 | 1.000 | down |
| G075703 | CABP1 | 0.912 | 0.002 | 1.000 | down |
| G075703 | CHRNA7 | 0.915 | 0.001 | 1.000 | down |
| LINC00858 | CABP1 | 0.935 | 0.001 | 1.000 | down |
| LINC00858 | CHRNA7 | 0.923 | 0.001 | 1.000 | down |
| BC047644 | DAB1 | 0.937 | 0.001 | 1.000 | down |
| BC047644 | ERC2 | 0.928 | 0.001 | 1.000 | down |
| BC047644 | GPSM2 | 0.944 | 0.000 | 1.000 | down |
| BC047644 | CALB2 | 0.912 | 0.002 | 1.000 | down |
| BC047644 | MAP2 | 0.908 | 0.002 | 1.000 | down |
| BC047644 | BTBD9 | 0.919 | 0.001 | 1.000 | down |
| BC047644 | PPP2R1A | 0.963 | 0.000 | 1.000 | down |
| G022520 | MCTP2 | 0.910 | 0.002 | 1.000 | down |
| G022520 | OLFM1 | 0.926 | 0.001 | 1.000 | down |
| RP11-820L6.1 | SOS1 | 0.926 | 0.001 | 1.000 | down |
| RP11-820L6.1 | CHRNA3 | 0.958 | 0.000 | 1.000 | down |
| LINC01604 | CRELD1 | 0.903 | 0.002 | 1.000 | down |
| LINC01604 | ARHGAP32 | 0.914 | 0.002 | 1.000 | down |
| LINC01604 | ATP2B4 | 0.973 | 0.000 | 1.000 | down |
| LINC01604 | ERC2 | 0.906 | 0.002 | 1.000 | down |
| LINC01604 | PTPRN2 | 0.913 | 0.002 | 1.000 | down |

| | | | | | |
|---|---|---|---|---|---|
| LINC01604 | CABP1 | 0.931 | 0.001 | 1.000 | down |
| LOC102723385 | PTCH1 | 0.910 | 0.002 | 1.000 | down |
| RP11-803B1.8 | TSPAN7 | 0.907 | 0.002 | 1.000 | down |
| RP11-803B1.8 | CABP1 | 0.925 | 0.001 | 1.000 | down |
| G005619 | TENM4 | 0.928 | 0.001 | 1.000 | down |
| G005619 | NRP2 | 0.942 | 0.000 | 1.000 | down |
| G063721 | CRH | 0.904 | 0.002 | 1.000 | down |
| XLOC_004894 | EXOC7 | 0.937 | 0.001 | 1.000 | down |
| XLOC_004894 | CHRNA3 | 0.907 | 0.002 | 1.000 | down |
| G015529 | NRP2 | 0.904 | 0.002 | 1.000 | down |
| G015529 | SPTB | 0.943 | 0.000 | 1.000 | down |
| G015529 | SEPSECS | 0.939 | 0.001 | 1.000 | down |
| G015529 | CABP1 | 0.925 | 0.001 | 1.000 | down |
| CTD-2517O10.6 | ELFN1 | 0.902 | 0.002 | 1.000 | down |
| CTD-2517O10.6 | SPTB | 0.909 | 0.002 | 1.000 | down |
| CTD-2517O10.6 | CABP1 | 0.948 | 0.000 | 1.000 | down |
| CTD-2517O10.6 | CHRNA7 | 0.950 | 0.000 | 1.000 | down |
| G001497 | ERC2 | 0.950 | 0.000 | 1.000 | down |
| G001497 | CALB2 | 0.967 | 0.000 | 1.000 | down |
| G001497 | MAP2 | 0.915 | 0.001 | 1.000 | down |
| RP11-439C15.4 | CRELD1 | 0.957 | 0.000 | 1.000 | down |
| RP11-439C15.4 | ATP2B4 | 0.915 | 0.001 | 1.000 | down |
| RP11-439C15.4 | ERC2 | 0.959 | 0.000 | 1.000 | down |
| RP11-439C15.4 | PTPRN2 | 0.936 | 0.001 | 1.000 | down |
| RP11-439C15.4 | MAP2 | 0.926 | 0.001 | 1.000 | down |
| RP11-439C15.4 | DPYSL5 | 0.958 | 0.000 | 1.000 | down |
| RP11-439C15.4 | DPYSL3 | 0.914 | 0.001 | 1.000 | down |
| TOB1-AS1 (2) | ATP2B4 | 0.914 | 0.002 | 1.000 | down |
| TOB1-AS1 (2) | CABP1 | 0.921 | 0.001 | 1.000 | down |
| TOB1-AS1 (2) | CHRNA7 | 0.931 | 0.001 | 1.000 | down |
| LOC101927358 | DAB1 | 0.904 | 0.002 | 1.000 | down |
| BC040628 | TSPAN7 | 0.917 | 0.001 | 1.000 | down |
| RP11-196G18.22 | ATP2B4 | 0.913 | 0.002 | 1.000 | down |
| RP11-196G18.22 | ERC2 | 0.957 | 0.000 | 1.000 | down |
| RP11-196G18.22 | CALB2 | 0.908 | 0.002 | 1.000 | down |
| RP11-196G18.22 | PTPRN2 | 0.946 | 0.000 | 1.000 | down |
| RP11-196G18.22 | MAP2 | 0.918 | 0.001 | 1.000 | down |
| RP11-196G18.22 | DPYSL5 | 0.917 | 0.001 | 1.000 | down |
| CTC-248O19.1 | ERC2 | 0.911 | 0.002 | 1.000 | down |
| CTC-248O19.1 | CALB2 | 0.938 | 0.001 | 1.000 | down |
| CTC-248O19.1 | PTPRN2 | 0.918 | 0.001 | 1.000 | down |
| CTC-248O19.1 | MAP2 | 0.918 | 0.001 | 1.000 | down |
| RP11-379F4.9 | CRELD1 | 0.917 | 0.001 | 1.000 | down |
| RP11-379F4.9 | ARHGAP32 | 0.942 | 0.000 | 1.000 | down |
| RP11-379F4.9 | ATP2B4 | 0.960 | 0.000 | 1.000 | down |
| RP11-379F4.9 | ERC2 | 0.940 | 0.001 | 1.000 | down |
| RP11-379F4.9 | PTPRN2 | 0.951 | 0.000 | 1.000 | down |
| RP11-379F4.9 | MAP2 | 0.929 | 0.001 | 1.000 | down |
| RP11-379F4.9 | DPYSL5 | 0.922 | 0.001 | 1.000 | down |
| G085202 | ATP2B4 | 0.921 | 0.001 | 1.000 | down |
| G085202 | SEPSECS | 0.916 | 0.001 | 1.000 | down |
| G085202 | TSPAN7 | 0.929 | 0.001 | 1.000 | down |
| LINC00670 | CHRNA7 | 0.947 | 0.000 | 1.000 | down |
| ATP2B4 | XLOC_012303 | -0.927 | 0.001 | -1.000 | down |
| ATP2B4 | LOC100505736 | -0.918 | 0.001 | -1.000 | down |
| ATP2B4 | G060633 | -0.925 | 0.001 | -1.000 | down |
| ATP2B4 | G029454 | -0.945 | 0.000 | -1.000 | down |
| ATP2B4 | G068163 | -0.928 | 0.001 | -1.000 | down |
| ATP2B4 | G021277 | -0.921 | 0.001 | -1.000 | down |
| ATP2B4 | G080487 | -0.915 | 0.001 | -1.000 | down |
| ATP2B4 | XLOC_008025 | -0.966 | 0.000 | -1.000 | down |
| ATP2B4 | XLOC_l2_011118 | -0.913 | 0.002 | -1.000 | down |
| ATP2B4 | LINC00379 | -0.910 | 0.002 | -1.000 | down |
| ATP2B4 | G072062 | -0.944 | 0.000 | -1.000 | down |

| | | | | | |
|---|---|---|---|---|---|
| ATP2B4 | RP11-717D12.1 | -0.967 | 0.000 | -1.000 | down |
| ATP2B4 | G006360 | -0.988 | 0.000 | -1.000 | down |
| ATP2B4 | LINC01149 | -0.929 | 0.001 | -1.000 | down |
| ATP2B4 | G016657 | -0.947 | 0.000 | -1.000 | down |
| ATP2B4 | G015181 | -0.912 | 0.002 | -1.000 | down |
| ATP2B4 | G037029 | -0.952 | 0.000 | -1.000 | down |
| ATP2B4 | G046492 | -0.922 | 0.001 | -1.000 | down |
| ATP2B4 | G072257 | -0.912 | 0.002 | -1.000 | down |
| ATP2B4 | CTD- | 0.964 | 0.000 | 1.000 | down |
| ATP2B4 | RP11-525A16.4 | 0.918 | 0.001 | 1.000 | down |
| ATP2B4 | RP11-350N15.5 | 0.965 | 0.000 | 1.000 | down |
| ATP2B4 | RP4-549L20.3 | 0.915 | 0.001 | 1.000 | down |
| ATP2B4 | BRWD1-IT2 | 0.947 | 0.000 | 1.000 | down |
| ATP2B4 | RP11-490O6.2 | 0.957 | 0.000 | 1.000 | down |
| ATP2B4 | ZFX-AS1 | 0.914 | 0.001 | 1.000 | down |
| ATP2B4 | RP11- | 0.967 | 0.000 | 1.000 | down |
| ATP2B4 | CTD- | 0.908 | 0.002 | 1.000 | down |
| ATP2B4 | CTD-2588C8.8 | 0.902 | 0.002 | 1.000 | down |
| ATP2B4 | G047225 | 0.916 | 0.001 | 1.000 | down |
| ATP2B4 | RP1-309F20.3 | 0.919 | 0.001 | 1.000 | down |
| ATP2B4 | RP11- | 0.905 | 0.002 | 1.000 | down |
| ATP2B4 | XLOC_008875 | 0.926 | 0.001 | 1.000 | down |
| ATP2B4 | RP11-191L17.1 | 0.959 | 0.000 | 1.000 | down |
| ATP2B4 | RP11-165A20.3 | 0.938 | 0.001 | 1.000 | down |
| ATP2B4 | G000481 | 0.917 | 0.001 | 1.000 | down |
| ATP2B4 | RP11-285F16.1 | 0.964 | 0.000 | 1.000 | down |
| ATP2B4 | RP11-351D16.3 | 0.952 | 0.000 | 1.000 | down |
| ATP2B4 | AK123758 | 0.935 | 0.001 | 1.000 | down |
| ATP2B4 | RP4-683L5.1 | 0.936 | 0.001 | 1.000 | down |
| ATP2B4 | RP11-353K11.1 | 0.966 | 0.000 | 1.000 | down |
| ATP2B4 | HRAT5 | 0.922 | 0.001 | 1.000 | down |
| ATP2B4 | RP11-312B8.1 | 0.977 | 0.000 | 1.000 | down |
| ATP2B4 | GAS5-AS1 | 0.929 | 0.001 | 1.000 | down |
| ATP2B4 | RP3-410B11.1 | 0.921 | 0.001 | 1.000 | down |
| ATP2B4 | G089005 | 0.941 | 0.000 | 1.000 | down |
| ATP2B4 | LOC101929741 | 0.904 | 0.002 | 1.000 | down |
| ATP2B4 | G079654 | 0.956 | 0.000 | 1.000 | down |
| ATP2B4 | TEX26-AS1 | 0.957 | 0.000 | 1.000 | down |
| ATP2B4 | RP11-496H1.2 | 0.916 | 0.001 | 1.000 | down |
| ATP2B4 | FRMD6-AS1 | 0.935 | 0.001 | 1.000 | down |
| ATP2B4 | AL355711 | 0.948 | 0.000 | 1.000 | down |
| ATP2B4 | CTD- | 0.967 | 0.000 | 1.000 | down |
| ATP2B4 | BC070391 | 0.935 | 0.001 | 1.000 | down |
| ATP2B4 | G041948 | 0.937 | 0.001 | 1.000 | down |
| ATP2B4 | G034470 | 0.939 | 0.001 | 1.000 | down |
| ATP2B4 | G073576 | 0.943 | 0.000 | 1.000 | down |
| ATP2B4 | RP11-424N24.2 | 0.910 | 0.002 | 1.000 | down |
| ATP2B4 | XLOC_011052 | 0.947 | 0.000 | 1.000 | down |
| ATP2B4 | RP1-202O8.3 | 0.952 | 0.000 | 1.000 | down |
| ATP2B4 | G019632 | 0.902 | 0.002 | 1.000 | down |
| ATP2B4 | G038972 | 0.948 | 0.000 | 1.000 | down |
| ATP2B4 | LINC-PINT (2) | 0.945 | 0.000 | 1.000 | down |
| ATP2B4 | RP11-505K9.1 | 0.957 | 0.000 | 1.000 | down |
| ATP2B4 | RP11-138E2.1 | 0.930 | 0.001 | 1.000 | down |
| ATP2B4 | G003798 | 0.964 | 0.000 | 1.000 | down |
| ATP2B4 | LOC148709 | 0.901 | 0.002 | 1.000 | down |
| ATP2B4 | G044581 | 0.955 | 0.000 | 1.000 | down |
| ATP2B4 | RP5-884G6.2 | 0.934 | 0.001 | 1.000 | down |
| ATP2B4 | CTD-2383M3.1 | 0.914 | 0.001 | 1.000 | down |
| ATP2B4 | AC092667.2 | 0.935 | 0.001 | 1.000 | down |
| ATP2B4 | RP11-383C5.5 | 0.965 | 0.000 | 1.000 | down |
| ATP2B4 | LOC644656 | 0.924 | 0.001 | 1.000 | down |
| ATP2B4 | G001739 | 0.960 | 0.000 | 1.000 | down |
| ATP2B4 | RP11-22N19.2 | 0.954 | 0.000 | 1.000 | down |
| ATP2B4 | LOC101928402 | 0.912 | 0.002 | 1.000 | down |

| | | | | | |
|---|---|---|---|---|---|
| ATP2B4 | GFOD1 | 0.938 | 0.001 | 1.000 | down |
| ATP2B4 | LOC101928417 | 0.926 | 0.001 | 1.000 | down |
| ATP2B4 | RP4-616B8.4 | 0.937 | 0.001 | 1.000 | down |
| ATP2B4 | RP4-555D20.3 | 0.934 | 0.001 | 1.000 | down |
| ATP2B4 | RP11-507K2.3 | 0.979 | 0.000 | 1.000 | down |
| ATP2B4 | RP4-798A10.7 | 0.953 | 0.000 | 1.000 | down |
| ATP2B4 | CTD- | 0.948 | 0.000 | 1.000 | down |
| ATP2B4 | RP11-254F7.2 | 0.910 | 0.002 | 1.000 | down |
| ATP2B4 | RP1-261G23.7 | 0.912 | 0.002 | 1.000 | down |
| ATP2B4 | G023087 | 0.931 | 0.001 | 1.000 | down |
| ATP2B4 | RP1-223E5.4 | 0.913 | 0.002 | 1.000 | down |
| ATP2B4 | G010838 | 0.924 | 0.001 | 1.000 | down |
| ATP2B4 | RP11-573G6.8 | 0.923 | 0.001 | 1.000 | down |
| ATP2B4 | LL09NC01-139C3.1 | 0.923 | 0.001 | 1.000 | down |
| ATP2B4 | LOC100288846 | 0.980 | 0.000 | 1.000 | down |
| ATP2B4 | UBAC2-AS1 | 0.914 | 0.001 | 1.000 | down |
| ATP2B4 | FLJ20021 | 0.908 | 0.002 | 1.000 | down |
| ATP2B4 | RP11-109D9.4 | 0.901 | 0.002 | 1.000 | down |
| ATP2B4 | CTC- | 0.933 | 0.001 | 1.000 | down |
| ATP2B4 | LINC00449 | 0.944 | 0.000 | 1.000 | down |
| ATP2B4 | RP11-61L19.2 | 0.978 | 0.000 | 1.000 | down |
| ATP2B4 | LOC101929066 | 0.968 | 0.000 | 1.000 | down |
| ATP2B4 | RP11-85A1.3 | 0.932 | 0.001 | 1.000 | down |
| ATP2B4 | G061596 | 0.907 | 0.002 | 1.000 | down |
| ATP2B4 | RP11-344P13.6 | 0.921 | 0.001 | 1.000 | down |
| ATP2B4 | RP11- | 0.915 | 0.001 | 1.000 | down |
| ATP2B4 | CTD- | 0.941 | 0.000 | 1.000 | down |
| ATP2B4 | RP1-232L24.3 | 0.926 | 0.001 | 1.000 | down |
| ATP2B4 | RP11-467P9.1 | 0.940 | 0.001 | 1.000 | down |
| ATP2B4 | XXbac-B476C20.9 | 0.936 | 0.001 | 1.000 | down |
| ATP2B4 | G021588 | 0.929 | 0.001 | 1.000 | down |
| ATP2B4 | RP11-371A19.2 | 0.945 | 0.000 | 1.000 | down |
| ATP2B4 | RP11-820I16.4 | 0.962 | 0.000 | 1.000 | down |
| ATP2B4 | LINC01604 | 0.973 | 0.000 | 1.000 | down |
| ATP2B4 | RP11-439C15.4 | 0.915 | 0.001 | 1.000 | down |
| ATP2B4 | TOB1-AS1 (2) | 0.914 | 0.002 | 1.000 | down |
| ATP2B4 | RP11- | 0.913 | 0.002 | 1.000 | down |
| ATP2B4 | RP11-379F4.9 | 0.960 | 0.000 | 1.000 | down |
| ATP2B4 | G085202 | 0.921 | 0.001 | 1.000 | down |
| PTCH1 | G060633 | -0.909 | 0.002 | -1.000 | down |
| PTCH1 | G061035 | -0.906 | 0.002 | -1.000 | down |
| PTCH1 | RP4-761J14.9 | -0.921 | 0.001 | -1.000 | down |
| PTCH1 | AP001625.6 | -0.946 | 0.000 | -1.000 | down |
| PTCH1 | XLOC_008025 | -0.940 | 0.001 | -1.000 | down |
| PTCH1 | XLOC_l2_011118 | -0.925 | 0.001 | -1.000 | down |
| PTCH1 | G081302 | -0.955 | 0.000 | -1.000 | down |
| PTCH1 | GSE61474_XLOC_028076 | -0.911 | 0.002 | -1.000 | down |
| PTCH1 | G063659 | -0.934 | 0.001 | -1.000 | down |
| PTCH1 | G072890 | -0.930 | 0.001 | -1.000 | down |
| PTCH1 | LOC101927691 | -0.971 | 0.000 | -1.000 | down |
| PTCH1 | RMRPP3 | -0.960 | 0.000 | -1.000 | down |
| PTCH1 | G058409 | -0.914 | 0.002 | -1.000 | down |
| PTCH1 | G009329 | -0.914 | 0.001 | -1.000 | down |
| PTCH1 | G015181 | -0.956 | 0.000 | -1.000 | down |
| PTCH1 | G051586 | -0.923 | 0.001 | -1.000 | down |
| PTCH1 | G010533 | -0.989 | 0.000 | -1.000 | down |
| PTCH1 | XLOC_003048 | -0.928 | 0.001 | -1.000 | down |
| PTCH1 | CTD- | 0.914 | 0.001 | 1.000 | down |
| PTCH1 | LOC729083 | 0.916 | 0.001 | 1.000 | down |
| PTCH1 | RP11-285F16.1 | 0.908 | 0.002 | 1.000 | down |
| PTCH1 | G079654 | 0.915 | 0.001 | 1.000 | down |

| | | | | | |
|---|---|---|---|---|---|
| PTCH1 | TEX26-AS1 | 0.914 | 0.002 | 1.000 | down |
| PTCH1 | G009018 | 0.997 | 0.000 | 1.000 | down |
| PTCH1 | RP11-1055B8.8 | 0.916 | 0.001 | 1.000 | down |
| PTCH1 | LINC-PINT (1) | 0.916 | 0.001 | 1.000 | down |
| PTCH1 | RP11-299H21.1 | 0.908 | 0.002 | 1.000 | down |
| PTCH1 | XLOC_011052 | 0.910 | 0.002 | 1.000 | down |
| PTCH1 | LINC-PINT (2) | 0.902 | 0.002 | 1.000 | down |
| PTCH1 | G066606 | 0.900 | 0.002 | 1.000 | down |
| PTCH1 | G019668 | 0.915 | 0.001 | 1.000 | down |
| PTCH1 | AC005220.3 | 0.916 | 0.001 | 1.000 | down |
| PTCH1 | CXADRP3 | 0.961 | 0.000 | 1.000 | down |
| PTCH1 | RP11-109D9.4 | 0.912 | 0.002 | 1.000 | down |
| PTCH1 | LINC00449 | 0.952 | 0.000 | 1.000 | down |
| PTCH1 | G054079 | 0.900 | 0.002 | 1.000 | down |
| PTCH1 | LOC101927811 | 0.911 | 0.002 | 1.000 | down |
| PTCH1 | XXbac-B476C20.9 | 0.917 | 0.001 | 1.000 | down |
| PTCH1 | G021588 | 0.935 | 0.001 | 1.000 | down |
| PTCH1 | LOC283140 | 0.903 | 0.002 | 1.000 | down |
| PTCH1 | LOC102723385 | 0.910 | 0.002 | 1.000 | down |
| ERC2 | G060633 | -0.944 | 0.000 | -1.000 | down |
| ERC2 | G011310 | -0.922 | 0.001 | -1.000 | down |
| ERC2 | AP000438.2 | -0.944 | 0.000 | -1.000 | down |
| ERC2 | uc.49 | -0.915 | 0.001 | -1.000 | down |
| ERC2 | G006360 | -0.900 | 0.002 | -1.000 | down |
| ERC2 | G073689 | -0.944 | 0.000 | -1.000 | down |
| ERC2 | G054530 | -0.916 | 0.001 | -1.000 | down |
| ERC2 | EML2-AS1 | -0.967 | 0.000 | -1.000 | down |
| ERC2 | XLOC_012740 | -0.942 | 0.000 | -1.000 | down |
| ERC2 | G051586 | -0.919 | 0.001 | -1.000 | down |
| ERC2 | RP5-1112D6.7 | -0.916 | 0.001 | -1.000 | down |
| ERC2 | RP11-461K13.1 | -0.924 | 0.001 | -1.000 | down |
| ERC2 | RP11-602N24.3 | 0.903 | 0.002 | 1.000 | down |
| ERC2 | AL110181 | 0.942 | 0.000 | 1.000 | down |
| ERC2 | TIPARP-AS1 | 0.900 | 0.002 | 1.000 | down |
| ERC2 | RP11-363E6.3 | 0.946 | 0.000 | 1.000 | down |
| ERC2 | CTD- | 0.952 | 0.000 | 1.000 | down |
| ERC2 | LINC00652 | 0.970 | 0.000 | 1.000 | down |
| ERC2 | RP11-303E16.7 | 0.913 | 0.002 | 1.000 | down |
| ERC2 | RP11-350N15.5 | 0.932 | 0.001 | 1.000 | down |
| ERC2 | RP4-549L20.3 | 0.904 | 0.002 | 1.000 | down |
| ERC2 | RP11-686O6.2 | 0.935 | 0.001 | 1.000 | down |
| ERC2 | G061591 | 0.967 | 0.000 | 1.000 | down |
| ERC2 | RP1-40E16.12 | 0.982 | 0.000 | 1.000 | down |
| ERC2 | GBAT2 | 0.970 | 0.000 | 1.000 | down |
| ERC2 | ZFX-AS1 | 0.946 | 0.000 | 1.000 | down |
| ERC2 | TM4SF1-AS1 | 0.942 | 0.000 | 1.000 | down |
| ERC2 | LINC00662 | 0.932 | 0.001 | 1.000 | down |
| ERC2 | RP11-297J22.1 | 0.938 | 0.001 | 1.000 | down |
| ERC2 | RP11-179B2.2 | 0.960 | 0.000 | 1.000 | down |
| ERC2 | RP11-191L17.1 | 0.965 | 0.000 | 1.000 | down |
| ERC2 | AC099754.1 | 0.922 | 0.001 | 1.000 | down |
| ERC2 | RP11-285F16.1 | 0.937 | 0.001 | 1.000 | down |
| ERC2 | MGC2889 | 0.927 | 0.001 | 1.000 | down |
| ERC2 | RP4-683L5.1 | 0.917 | 0.001 | 1.000 | down |
| ERC2 | GAS5-AS1 | 0.930 | 0.001 | 1.000 | down |
| ERC2 | RP11-204E9.1 | 0.909 | 0.002 | 1.000 | down |
| ERC2 | G085410 | 0.942 | 0.000 | 1.000 | down |
| ERC2 | DHRS4-AS1 | 0.926 | 0.001 | 1.000 | down |
| ERC2 | G089005 | 0.948 | 0.000 | 1.000 | down |
| ERC2 | OLMALINC | 0.915 | 0.001 | 1.000 | down |
| ERC2 | RP11-496H1.2 | 0.955 | 0.000 | 1.000 | down |
| ERC2 | MAPT-IT1 | 0.917 | 0.001 | 1.000 | down |
| ERC2 | G073576 | 0.961 | 0.000 | 1.000 | down |
| ERC2 | XLOC_013222 | 0.944 | 0.000 | 1.000 | down |

| | | | | | |
|---|---|---|---|---|---|
| ERC2 | XLOC_011052 | 0.960 | 0.000 | 1.000 | down |
| ERC2 | TNRC18P1 | 0.927 | 0.001 | 1.000 | down |
| ERC2 | G065067 | 0.940 | 0.001 | 1.000 | down |
| ERC2 | RP11-138E2.1 | 0.931 | 0.001 | 1.000 | down |
| ERC2 | G044581 | 0.908 | 0.002 | 1.000 | down |
| ERC2 | AC099850.1 | 0.926 | 0.001 | 1.000 | down |
| ERC2 | BC042589 | 0.944 | 0.000 | 1.000 | down |
| ERC2 | LINC00634 | 0.961 | 0.000 | 1.000 | down |
| ERC2 | AC092667.2 | 0.903 | 0.002 | 1.000 | down |
| ERC2 | RP11-22N19.2 | 0.951 | 0.000 | 1.000 | down |
| ERC2 | RP11-51J9.5 | 0.923 | 0.001 | 1.000 | down |
| ERC2 | RP4-616B8.4 | 0.921 | 0.001 | 1.000 | down |
| ERC2 | RP4-555D20.3 | 0.922 | 0.001 | 1.000 | down |
| ERC2 | RP4-798A10.7 | 0.950 | 0.000 | 1.000 | down |
| ERC2 | LINC00630 | 0.930 | 0.001 | 1.000 | down |
| ERC2 | RP11-254F7.2 | 0.946 | 0.000 | 1.000 | down |
| ERC2 | G066437 | 0.903 | 0.002 | 1.000 | down |
| ERC2 | ENO1-AS1 | 0.902 | 0.002 | 1.000 | down |
| ERC2 | RP1-223E5.4 | 0.944 | 0.000 | 1.000 | down |
| ERC2 | DGCR5 | 0.920 | 0.001 | 1.000 | down |
| ERC2 | G010838 | 0.936 | 0.001 | 1.000 | down |
| ERC2 | RP11-573G6.8 | 0.976 | 0.000 | 1.000 | down |
| ERC2 | LL09NC01-139C3.1 | 0.972 | 0.000 | 1.000 | down |
| ERC2 | LOC100288846 | 0.902 | 0.002 | 1.000 | down |
| ERC2 | CXADRP3 | 0.921 | 0.001 | 1.000 | down |
| ERC2 | G061596 | 0.923 | 0.001 | 1.000 | down |
| ERC2 | RP11-53I6.3 | 0.908 | 0.002 | 1.000 | down |
| ERC2 | RP11-314A20.5 | 0.915 | 0.001 | 1.000 | down |
| ERC2 | RP11- | 0.959 | 0.000 | 1.000 | down |
| ERC2 | CTD- | 0.902 | 0.002 | 1.000 | down |
| ERC2 | G023340 | 0.962 | 0.000 | 1.000 | down |
| ERC2 | RP11-166B2.3 | 0.924 | 0.001 | 1.000 | down |
| ERC2 | BC047644 | 0.928 | 0.001 | 1.000 | down |
| ERC2 | LINC01604 | 0.906 | 0.002 | 1.000 | down |
| ERC2 | G001497 | 0.950 | 0.000 | 1.000 | down |
| ERC2 | RP11-439C15.4 | 0.959 | 0.000 | 1.000 | down |
| ERC2 | RP11- | 0.957 | 0.000 | 1.000 | down |
| ERC2 | CTC-248O19.1 | 0.911 | 0.002 | 1.000 | down |
| ERC2 | RP11-379F4.9 | 0.940 | 0.001 | 1.000 | down |
| FILIP1 | RP1-296L11.1 | -0.905 | 0.002 | -1.000 | down |
| FILIP1 | G060633 | -0.935 | 0.001 | -1.000 | down |
| FILIP1 | G045736 | -0.920 | 0.001 | -1.000 | down |
| FILIP1 | G061888 | -0.904 | 0.002 | -1.000 | down |
| FILIP1 | RP11-303E16.7 | 0.908 | 0.002 | 1.000 | down |
| FILIP1 | LINC00662 | 0.907 | 0.002 | 1.000 | down |
| FILIP1 | G018162 | 0.965 | 0.000 | 1.000 | down |
| FILIP1 | AC002467.7 | 0.960 | 0.000 | 1.000 | down |
| FILIP1 | RP11-174G6.5 | 0.920 | 0.001 | 1.000 | down |
| FILIP1 | G034470 | 0.918 | 0.001 | 1.000 | down |
| FILIP1 | LOC730102 | 0.907 | 0.002 | 1.000 | down |
| FILIP1 | G044581 | 0.912 | 0.002 | 1.000 | down |
| FILIP1 | BC042589 | 0.941 | 0.000 | 1.000 | down |
| FILIP1 | AC005220.3 | 0.927 | 0.001 | 1.000 | down |
| FILIP1 | RP11-22N19.2 | 0.906 | 0.002 | 1.000 | down |
| FILIP1 | XLOC_007467 | 0.952 | 0.000 | 1.000 | down |
| FILIP1 | FLJ20021 | 0.902 | 0.002 | 1.000 | down |
| FILIP1 | G075319 | 0.927 | 0.001 | 1.000 | down |
| FILIP1 | RP11-166B2.3 | 0.943 | 0.000 | 1.000 | down |
| GPSM2 | G011310 | -0.953 | 0.000 | -1.000 | down |
| GPSM2 | G008916 | -0.912 | 0.002 | -1.000 | down |
| GPSM2 | XLOC_006150 | -0.921 | 0.001 | -1.000 | down |
| GPSM2 | G073689 | -0.913 | 0.002 | -1.000 | down |
| GPSM2 | G003431 | -0.908 | 0.002 | -1.000 | down |
| GPSM2 | AC025442.3 | -0.934 | 0.001 | -1.000 | down |

| | | | | | |
|---|---|---|---|---|---|
| GPSM2 | G013465 | -0.982 | 0.000 | -1.000 | down |
| GPSM2 | AL110181 | 0.904 | 0.002 | 1.000 | down |
| GPSM2 | TIPARP-AS1 | 0.952 | 0.000 | 1.000 | down |
| GPSM2 | CTD-2023N9.3 | 0.912 | 0.002 | 1.000 | down |
| GPSM2 | RP11-356J5.12 | 0.908 | 0.002 | 1.000 | down |
| GPSM2 | GBAT2 | 0.902 | 0.002 | 1.000 | down |
| GPSM2 | RP11-297J22.1 | 0.903 | 0.002 | 1.000 | down |
| GPSM2 | AC144652.1 | 0.968 | 0.000 | 1.000 | down |
| GPSM2 | OLMALINC | 0.913 | 0.002 | 1.000 | down |
| GPSM2 | LINC00408 | 0.931 | 0.001 | 1.000 | down |
| GPSM2 | G023041 | 0.908 | 0.002 | 1.000 | down |
| GPSM2 | G053697 | 0.916 | 0.001 | 1.000 | down |
| GPSM2 | LOC730102 | 0.956 | 0.000 | 1.000 | down |
| GPSM2 | LINC00634 | 0.948 | 0.000 | 1.000 | down |
| GPSM2 | RP11-51J9.5 | 0.941 | 0.000 | 1.000 | down |
| GPSM2 | RP11- | 0.936 | 0.001 | 1.000 | down |
| GPSM2 | CTD-2653B5.1 | 0.917 | 0.001 | 1.000 | down |
| GPSM2 | KB-318B8.7 | 0.912 | 0.002 | 1.000 | down |
| GPSM2 | G075319 | 0.925 | 0.001 | 1.000 | down |
| GPSM2 | RP11-166B2.3 | 0.920 | 0.001 | 1.000 | down |
| GPSM2 | BC047644 | 0.944 | 0.000 | 1.000 | down |
| CALB2 | RP11-70P17.1 | -0.947 | 0.000 | -1.000 | down |
| CALB2 | G011310 | -0.943 | 0.000 | -1.000 | down |
| CALB2 | G071640 | -0.903 | 0.002 | -1.000 | down |
| CALB2 | uc.49 | -0.927 | 0.001 | -1.000 | down |
| CALB2 | G073689 | -0.936 | 0.001 | -1.000 | down |
| CALB2 | EML2-AS1 | -0.910 | 0.002 | -1.000 | down |
| CALB2 | XLOC_012740 | -0.901 | 0.002 | -1.000 | down |
| CALB2 | CTD- | -0.926 | 0.001 | -1.000 | down |
| CALB2 | G089283 | -0.928 | 0.001 | -1.000 | down |
| CALB2 | RP5-1112D6.7 | -0.937 | 0.001 | -1.000 | down |
| CALB2 | RP11-461K13.1 | -0.906 | 0.002 | -1.000 | down |
| CALB2 | IFT74-AS1 | 0.916 | 0.001 | 1.000 | down |
| CALB2 | RP11-602N24.3 | 0.917 | 0.001 | 1.000 | down |
| CALB2 | AL110181 | 0.957 | 0.000 | 1.000 | down |
| CALB2 | RP11-363E6.3 | 0.942 | 0.000 | 1.000 | down |
| CALB2 | LINC00652 | 0.957 | 0.000 | 1.000 | down |
| CALB2 | G061591 | 0.927 | 0.001 | 1.000 | down |
| CALB2 | RP1-40E16.12 | 0.949 | 0.000 | 1.000 | down |
| CALB2 | GBAT2 | 0.943 | 0.000 | 1.000 | down |
| CALB2 | TM4SF1-AS1 | 0.906 | 0.002 | 1.000 | down |
| CALB2 | LINC00662 | 0.904 | 0.002 | 1.000 | down |
| CALB2 | RP11-179B2.2 | 0.905 | 0.002 | 1.000 | down |
| CALB2 | G017886 | 0.909 | 0.002 | 1.000 | down |
| CALB2 | AC099754.1 | 0.922 | 0.001 | 1.000 | down |
| CALB2 | MGC2889 | 0.954 | 0.000 | 1.000 | down |
| CALB2 | G065067 | 0.934 | 0.001 | 1.000 | down |
| CALB2 | NIPBL-AS1 | 0.916 | 0.001 | 1.000 | down |
| CALB2 | LINC00634 | 0.905 | 0.002 | 1.000 | down |
| CALB2 | G078120 | 0.905 | 0.002 | 1.000 | down |
| CALB2 | RP11-254F7.2 | 0.905 | 0.002 | 1.000 | down |
| CALB2 | LL22NC03-86G7.1 | 0.906 | 0.002 | 1.000 | down |
| CALB2 | LINC00463 | 0.918 | 0.001 | 1.000 | down |
| CALB2 | G027543 | 0.928 | 0.001 | 1.000 | down |
| CALB2 | LL09NC01-139C3.1 | 0.904 | 0.002 | 1.000 | down |
| CALB2 | RP11-110G21.1 | 0.918 | 0.001 | 1.000 | down |
| CALB2 | RP11-314A20.5 | 0.953 | 0.000 | 1.000 | down |
| CALB2 | RP11- | 0.915 | 0.001 | 1.000 | down |
| CALB2 | G023340 | 0.922 | 0.001 | 1.000 | down |
| CALB2 | RP11-355F22.1 | 0.926 | 0.001 | 1.000 | down |
| CALB2 | BC047644 | 0.912 | 0.002 | 1.000 | down |
| CALB2 | G001497 | 0.967 | 0.000 | 1.000 | down |
| CALB2 | RP11- | 0.908 | 0.002 | 1.000 | down |

| | | | | | |
|---|---|---|---|---|---|
| CALB2 | CTC-248O19.1 | 0.938 | 0.001 | 1.000 | down |
| SLC17A8 | FAM138E | 0.914 | 0.002 | 1.000 | down |
| SPTB | PPIEL | -0.903 | 0.002 | -1.000 | down |
| SPTB | XLOC_l2_009332 | -0.917 | 0.001 | -1.000 | down |
| SPTB | G087508 | -0.914 | 0.002 | -1.000 | down |
| SPTB | G050702 | -0.933 | 0.001 | -1.000 | down |
| SPTB | ZFAT-AS1 | -0.915 | 0.001 | -1.000 | down |
| SPTB | RP11-18D7.2 | -0.904 | 0.002 | -1.000 | down |
| SPTB | AK093193 | -0.949 | 0.000 | -1.000 | down |
| SPTB | LOC101927164 | -0.947 | 0.000 | -1.000 | down |
| SPTB | CTD- | -0.940 | 0.001 | -1.000 | down |
| SPTB | XLOC_013958 | -0.956 | 0.000 | -1.000 | down |
| SPTB | BX648501 | -0.931 | 0.001 | -1.000 | down |
| SPTB | RP11-374P20.4 | -0.905 | 0.002 | -1.000 | down |
| SPTB | G045736 | -0.939 | 0.001 | -1.000 | down |
| SPTB | RP11-30O15.1 | -0.920 | 0.001 | -1.000 | down |
| SPTB | G067215 | -0.920 | 0.001 | -1.000 | down |
| SPTB | G037029 | -0.900 | 0.002 | -1.000 | down |
| SPTB | G010088 | -0.909 | 0.002 | -1.000 | down |
| SPTB | uc.50 | -0.902 | 0.002 | -1.000 | down |
| SPTB | RP11-651L5.2 | -0.904 | 0.002 | -1.000 | down |
| SPTB | G071259 | -0.915 | 0.001 | -1.000 | down |
| SPTB | G068890 | -0.904 | 0.002 | -1.000 | down |
| SPTB | G059595 | -0.908 | 0.002 | -1.000 | down |
| SPTB | FAM74A4 | 0.922 | 0.001 | 1.000 | down |
| SPTB | RP11-255E6.6 | 0.912 | 0.002 | 1.000 | down |
| SPTB | RP11-744I24.3 | 0.962 | 0.000 | 1.000 | down |
| SPTB | XLOC_006779 | 0.912 | 0.002 | 1.000 | down |
| SPTB | GSE61474_XLOC_054139 | 0.942 | 0.000 | 1.000 | down |
| SPTB | LOC100130417 | 0.947 | 0.000 | 1.000 | down |
| SPTB | USP27X-AS1 | 0.960 | 0.000 | 1.000 | down |
| SPTB | RP11-490O6.2 | 0.908 | 0.002 | 1.000 | down |
| SPTB | G002862 | 0.916 | 0.001 | 1.000 | down |
| SPTB | RP1-309F20.3 | 0.902 | 0.002 | 1.000 | down |
| SPTB | G050681 | 0.974 | 0.000 | 1.000 | down |
| SPTB | RP3-410B11.1 | 0.904 | 0.002 | 1.000 | down |
| SPTB | AC093391.2 | 0.952 | 0.000 | 1.000 | down |
| SPTB | XLOC_006928 | 0.956 | 0.000 | 1.000 | down |
| SPTB | RP11-128P10.1 | 0.957 | 0.000 | 1.000 | down |
| SPTB | RP11-174G6.5 | 0.925 | 0.001 | 1.000 | down |
| SPTB | AL355711 | 0.911 | 0.002 | 1.000 | down |
| SPTB | RP1-202O8.3 | 0.912 | 0.002 | 1.000 | down |
| SPTB | RP13-131K19.2 | 0.921 | 0.001 | 1.000 | down |
| SPTB | FOXN3-AS1 | 0.957 | 0.000 | 1.000 | down |
| SPTB | IDH1-AS1 | 0.921 | 0.001 | 1.000 | down |
| SPTB | RP11-505K9.1 | 0.905 | 0.002 | 1.000 | down |
| SPTB | LOC148709 | 0.914 | 0.001 | 1.000 | down |
| SPTB | GYG2P1 | 0.938 | 0.001 | 1.000 | down |
| SPTB | FGF13-AS1 | 0.940 | 0.001 | 1.000 | down |
| SPTB | AK000953 | 0.914 | 0.002 | 1.000 | down |
| SPTB | CTD- | 0.909 | 0.002 | 1.000 | down |
| SPTB | RP11-332H14.1 | 0.925 | 0.001 | 1.000 | down |
| SPTB | RP11- | 0.904 | 0.002 | 1.000 | down |
| SPTB | BX248253 | 0.909 | 0.002 | 1.000 | down |
| SPTB | RP11-53I6.3 | 0.919 | 0.001 | 1.000 | down |
| SPTB | RP11-474O21.5 | 0.912 | 0.002 | 1.000 | down |
| SPTB | G046528 | 0.902 | 0.002 | 1.000 | down |
| SPTB | G063514 | 0.968 | 0.000 | 1.000 | down |
| SPTB | RP11-95P2.3 | 0.928 | 0.001 | 1.000 | down |
| SPTB | G015529 | 0.943 | 0.000 | 1.000 | down |
| SPTB | CTD- | 0.909 | 0.002 | 1.000 | down |
| PTPRN2 | AP000438.2 | -0.921 | 0.001 | -1.000 | down |
| PTPRN2 | RP11-466A19.1 | -0.931 | 0.001 | -1.000 | down |

| | | | | | |
|---|---|---|---|---|---|
| PTPRN2 | EML2-AS1 | -0.987 | 0.000 | -1.000 | down |
| PTPRN2 | XLOC_012740 | -0.911 | 0.002 | -1.000 | down |
| PTPRN2 | LOC100506834 | -0.928 | 0.001 | -1.000 | down |
| PTPRN2 | EF565122 | -0.932 | 0.001 | -1.000 | down |
| PTPRN2 | RP11-602N24.3 | 0.931 | 0.001 | 1.000 | down |
| PTPRN2 | AL110181 | 0.906 | 0.002 | 1.000 | down |
| PTPRN2 | RP11-363E6.3 | 0.917 | 0.001 | 1.000 | down |
| PTPRN2 | CTD- | 0.913 | 0.002 | 1.000 | down |
| PTPRN2 | RP11-350N15.5 | 0.928 | 0.001 | 1.000 | down |
| PTPRN2 | RP11-686O6.2 | 0.957 | 0.000 | 1.000 | down |
| PTPRN2 | G061591 | 0.984 | 0.000 | 1.000 | down |
| PTPRN2 | RP1-40E16.12 | 0.974 | 0.000 | 1.000 | down |
| PTPRN2 | GBAT2 | 0.958 | 0.000 | 1.000 | down |
| PTPRN2 | ZFX-AS1 | 0.945 | 0.000 | 1.000 | down |
| PTPRN2 | AC138655.6 | 0.920 | 0.001 | 1.000 | down |
| PTPRN2 | RP11-297J22.1 | 0.946 | 0.000 | 1.000 | down |
| PTPRN2 | RP11-179B2.2 | 0.962 | 0.000 | 1.000 | down |
| PTPRN2 | RP11-191L17.1 | 0.922 | 0.001 | 1.000 | down |
| PTPRN2 | AC099754.1 | 0.905 | 0.002 | 1.000 | down |
| PTPRN2 | RP4-683L5.1 | 0.907 | 0.002 | 1.000 | down |
| PTPRN2 | GAS5-AS1 | 0.919 | 0.001 | 1.000 | down |
| PTPRN2 | G085410 | 0.934 | 0.001 | 1.000 | down |
| PTPRN2 | G089005 | 0.911 | 0.002 | 1.000 | down |
| PTPRN2 | RP11-496H1.2 | 0.944 | 0.000 | 1.000 | down |
| PTPRN2 | G073576 | 0.948 | 0.000 | 1.000 | down |
| PTPRN2 | XLOC_013222 | 0.910 | 0.002 | 1.000 | down |
| PTPRN2 | TNRC18P1 | 0.933 | 0.001 | 1.000 | down |
| PTPRN2 | G065067 | 0.947 | 0.000 | 1.000 | down |
| PTPRN2 | RP11-138E2.1 | 0.903 | 0.002 | 1.000 | down |
| PTPRN2 | AC099850.1 | 0.943 | 0.000 | 1.000 | down |
| PTPRN2 | LINC00634 | 0.928 | 0.001 | 1.000 | down |
| PTPRN2 | AC092667.2 | 0.900 | 0.002 | 1.000 | down |
| PTPRN2 | RP4-798A10.7 | 0.905 | 0.002 | 1.000 | down |
| PTPRN2 | RP1-223E5.4 | 0.923 | 0.001 | 1.000 | down |
| PTPRN2 | G010838 | 0.942 | 0.000 | 1.000 | down |
| PTPRN2 | RP11-573G6.8 | 0.969 | 0.000 | 1.000 | down |
| PTPRN2 | LL09NC01-139C3.1 | 0.941 | 0.000 | 1.000 | down |
| PTPRN2 | G061596 | 0.916 | 0.001 | 1.000 | down |
| PTPRN2 | RP11-483I13.5 | 0.915 | 0.001 | 1.000 | down |
| PTPRN2 | CTD- | 0.901 | 0.002 | 1.000 | down |
| PTPRN2 | G023340 | 0.980 | 0.000 | 1.000 | down |
| PTPRN2 | RP11-79P5.9 | 0.905 | 0.002 | 1.000 | down |
| PTPRN2 | LINC01604 | 0.913 | 0.002 | 1.000 | down |
| PTPRN2 | RP11-439C15.4 | 0.936 | 0.001 | 1.000 | down |
| PTPRN2 | RP11- | 0.946 | 0.000 | 1.000 | down |
| PTPRN2 | CTC-248O19.1 | 0.918 | 0.001 | 1.000 | down |
| PTPRN2 | RP11-379F4.9 | 0.951 | 0.000 | 1.000 | down |
| SEPSECS | XLOC_012303 | -0.914 | 0.001 | -1.000 | down |
| SEPSECS | G087508 | -0.901 | 0.002 | -1.000 | down |
| SEPSECS | G050702 | -0.903 | 0.002 | -1.000 | down |
| SEPSECS | AK093193 | -0.973 | 0.000 | -1.000 | down |
| SEPSECS | RP11-424I19.2 | -0.923 | 0.001 | -1.000 | down |
| SEPSECS | G006360 | -0.917 | 0.001 | -1.000 | down |
| SEPSECS | XLOC_013958 | -0.954 | 0.000 | -1.000 | down |
| SEPSECS | G045736 | -0.971 | 0.000 | -1.000 | down |
| SEPSECS | G016657 | -0.922 | 0.001 | -1.000 | down |
| SEPSECS | G067215 | -0.906 | 0.002 | -1.000 | down |
| SEPSECS | G037029 | -0.954 | 0.000 | -1.000 | down |
| SEPSECS | G046492 | -0.907 | 0.002 | -1.000 | down |
| SEPSECS | G059595 | -0.917 | 0.001 | -1.000 | down |
| SEPSECS | IFT74-AS1 | 0.912 | 0.002 | 1.000 | down |
| SEPSECS | RP11-255E6.6 | 0.930 | 0.001 | 1.000 | down |
| SEPSECS | RP11-744I24.3 | 0.948 | 0.000 | 1.000 | down |

| | | | | | |
|---|---|---|---|---|---|
| SEPSECS | GSE61474_XLOC_054139 | 0.947 | 0.000 | 1.000 | down |
| SEPSECS | LOC100130417 | 0.935 | 0.001 | 1.000 | down |
| SEPSECS | RP11-525A16.4 | 0.956 | 0.000 | 1.000 | down |
| SEPSECS | XLOC_008005 | 0.917 | 0.001 | 1.000 | down |
| SEPSECS | USP27X-AS1 | 0.909 | 0.002 | 1.000 | down |
| SEPSECS | BRWD1-IT2 | 0.917 | 0.001 | 1.000 | down |
| SEPSECS | RP11-490O6.2 | 0.953 | 0.000 | 1.000 | down |
| SEPSECS | PSMG3-AS1 | 0.942 | 0.000 | 1.000 | down |
| SEPSECS | TM4SF1-AS1 | 0.906 | 0.002 | 1.000 | down |
| SEPSECS | CTD-2588C8.8 | 0.902 | 0.002 | 1.000 | down |
| SEPSECS | RP1-309F20.3 | 0.935 | 0.001 | 1.000 | down |
| SEPSECS | RP11-252A24.3 | 0.906 | 0.002 | 1.000 | down |
| SEPSECS | RP11-191L17.1 | 0.946 | 0.000 | 1.000 | down |
| SEPSECS | RP11-165A20.3 | 0.906 | 0.002 | 1.000 | down |
| SEPSECS | RP11-285F16.1 | 0.944 | 0.000 | 1.000 | down |
| SEPSECS | G050681 | 0.928 | 0.001 | 1.000 | down |
| SEPSECS | RP11-351D16.3 | 0.901 | 0.002 | 1.000 | down |
| SEPSECS | AK123758 | 0.913 | 0.002 | 1.000 | down |
| SEPSECS | RP11-312B8.1 | 0.923 | 0.001 | 1.000 | down |
| SEPSECS | DHRS4-AS1 | 0.928 | 0.001 | 1.000 | down |
| SEPSECS | RP3-410B11.1 | 0.917 | 0.001 | 1.000 | down |
| SEPSECS | G089005 | 0.901 | 0.002 | 1.000 | down |
| SEPSECS | G079654 | 0.940 | 0.001 | 1.000 | down |
| SEPSECS | XLOC_006928 | 0.928 | 0.001 | 1.000 | down |
| SEPSECS | RP11-174G6.5 | 0.979 | 0.000 | 1.000 | down |
| SEPSECS | AL355711 | 0.941 | 0.001 | 1.000 | down |
| SEPSECS | CTD- | 0.932 | 0.001 | 1.000 | down |
| SEPSECS | RP11-424N24.2 | 0.967 | 0.000 | 1.000 | down |
| SEPSECS | XLOC_011052 | 0.947 | 0.000 | 1.000 | down |
| SEPSECS | RP11-502I4.3 | 0.911 | 0.002 | 1.000 | down |
| SEPSECS | RP1-202O8.3 | 0.952 | 0.000 | 1.000 | down |
| SEPSECS | RP13-131K19.2 | 0.918 | 0.001 | 1.000 | down |
| SEPSECS | FOXN3-AS1 | 0.950 | 0.000 | 1.000 | down |
| SEPSECS | RP11-290L1.2 | 0.916 | 0.001 | 1.000 | down |
| SEPSECS | IDH1-AS1 | 0.903 | 0.002 | 1.000 | down |
| SEPSECS | G038972 | 0.917 | 0.001 | 1.000 | down |
| SEPSECS | RP11-505K9.1 | 0.956 | 0.000 | 1.000 | down |
| SEPSECS | RP11-792A8.4 | 0.916 | 0.001 | 1.000 | down |
| SEPSECS | RP11-775D22.3 | 0.908 | 0.002 | 1.000 | down |
| SEPSECS | RP11-2E11.5 | 0.905 | 0.002 | 1.000 | down |
| SEPSECS | LOC148709 | 0.903 | 0.002 | 1.000 | down |
| SEPSECS | G044581 | 0.927 | 0.001 | 1.000 | down |
| SEPSECS | BC042589 | 0.917 | 0.001 | 1.000 | down |
| SEPSECS | AC092667.2 | 0.916 | 0.001 | 1.000 | down |
| SEPSECS | RP11-383C5.5 | 0.924 | 0.001 | 1.000 | down |
| SEPSECS | AC005220.3 | 0.927 | 0.001 | 1.000 | down |
| SEPSECS | LOC101928402 | 0.913 | 0.002 | 1.000 | down |
| SEPSECS | RP4-555D20.3 | 0.940 | 0.001 | 1.000 | down |
| SEPSECS | DQ594967 | 0.913 | 0.002 | 1.000 | down |
| SEPSECS | AP000255.6 | 0.957 | 0.000 | 1.000 | down |
| SEPSECS | CTD- | 0.931 | 0.001 | 1.000 | down |
| SEPSECS | RP11-254F7.2 | 0.975 | 0.000 | 1.000 | down |
| SEPSECS | RP11-332H14.1 | 0.916 | 0.001 | 1.000 | down |
| SEPSECS | ENO1-AS1 | 0.901 | 0.002 | 1.000 | down |
| SEPSECS | G023087 | 0.904 | 0.002 | 1.000 | down |
| SEPSECS | XLOC_007467 | 0.915 | 0.001 | 1.000 | down |
| SEPSECS | LL09NC01-139C3.1 | 0.921 | 0.001 | 1.000 | down |
| SEPSECS | LOC100288846 | 0.937 | 0.001 | 1.000 | down |
| SEPSECS | UBAC2-AS1 | 0.916 | 0.001 | 1.000 | down |
| SEPSECS | G000672 | 0.913 | 0.002 | 1.000 | down |
| SEPSECS | RP1-86C11.7 | 0.900 | 0.002 | 1.000 | down |
| SEPSECS | FLJ20021 | 0.925 | 0.001 | 1.000 | down |
| SEPSECS | RP11-109D9.4 | 0.934 | 0.001 | 1.000 | down |

| | | | | | |
|---|---|---|---|---|---|
| SEPSECS | BX248253 | 0.916 | 0.001 | 1.000 | down |
| SEPSECS | RP11-53I6.3 | 0.953 | 0.000 | 1.000 | down |
| SEPSECS | RP11- | 0.913 | 0.002 | 1.000 | down |
| SEPSECS | LOC101927811 | 0.905 | 0.002 | 1.000 | down |
| SEPSECS | RP1-232L24.3 | 0.960 | 0.000 | 1.000 | down |
| SEPSECS | RP11-467P9.1 | 0.949 | 0.000 | 1.000 | down |
| SEPSECS | G021588 | 0.926 | 0.001 | 1.000 | down |
| SEPSECS | RP11-95P2.3 | 0.957 | 0.000 | 1.000 | down |
| SEPSECS | G015529 | 0.939 | 0.001 | 1.000 | down |
| SEPSECS | G085202 | 0.916 | 0.001 | 1.000 | down |
| ALDOC | G057625 | -0.917 | 0.001 | -1.000 | down |
| ALDOC | G036779 | -0.907 | 0.002 | -1.000 | down |
| ALDOC | AC003682.16 | -0.964 | 0.000 | -1.000 | down |
| ALDOC | G022034 | -0.971 | 0.000 | -1.000 | down |
| ALDOC | G011388 | -0.939 | 0.001 | -1.000 | down |
| ALDOC | G002461 | -0.938 | 0.001 | -1.000 | down |
| ALDOC | AF086132 | -0.944 | 0.000 | -1.000 | down |
| ALDOC | SIPA1L2 | 0.906 | 0.002 | 1.000 | down |
| ALDOC | G017670 | 0.902 | 0.002 | 1.000 | down |
| ALDOC | RP11-680F8.1 | 0.985 | 0.000 | 1.000 | down |
| TSPAN7 | G029454 | -0.901 | 0.002 | -1.000 | down |
| TSPAN7 | G053606 | -0.907 | 0.002 | -1.000 | down |
| TSPAN7 | G006738 | -0.905 | 0.002 | -1.000 | down |
| TSPAN7 | G063174 | -0.918 | 0.001 | -1.000 | down |
| TSPAN7 | LINC00379 | -0.915 | 0.001 | -1.000 | down |
| TSPAN7 | G070831 | -0.900 | 0.002 | -1.000 | down |
| TSPAN7 | AK093193 | -0.930 | 0.001 | -1.000 | down |
| TSPAN7 | XLOC_013958 | -0.945 | 0.000 | -1.000 | down |
| TSPAN7 | G016657 | -0.916 | 0.001 | -1.000 | down |
| TSPAN7 | RP11-30O15.1 | -0.927 | 0.001 | -1.000 | down |
| TSPAN7 | G037029 | -0.938 | 0.001 | -1.000 | down |
| TSPAN7 | G010088 | -0.925 | 0.001 | -1.000 | down |
| TSPAN7 | G046492 | -0.916 | 0.001 | -1.000 | down |
| TSPAN7 | CTD- | -0.905 | 0.002 | -1.000 | down |
| TSPAN7 | IFT74-AS1 | 0.910 | 0.002 | 1.000 | down |
| TSPAN7 | RP11-744I24.3 | 0.963 | 0.000 | 1.000 | down |
| TSPAN7 | LOC100130417 | 0.978 | 0.000 | 1.000 | down |
| TSPAN7 | OR2L1P | 0.929 | 0.001 | 1.000 | down |
| TSPAN7 | RP11-525A16.4 | 0.911 | 0.002 | 1.000 | down |
| TSPAN7 | USP27X-AS1 | 0.927 | 0.001 | 1.000 | down |
| TSPAN7 | G070136 | 0.939 | 0.001 | 1.000 | down |
| TSPAN7 | BRWD1-IT2 | 0.913 | 0.002 | 1.000 | down |
| TSPAN7 | RP11-490O6.2 | 0.939 | 0.001 | 1.000 | down |
| TSPAN7 | PSMG3-AS1 | 0.913 | 0.002 | 1.000 | down |
| TSPAN7 | CTD- | 0.938 | 0.001 | 1.000 | down |
| TSPAN7 | CTD-2588C8.8 | 0.901 | 0.002 | 1.000 | down |
| TSPAN7 | RP1-309F20.3 | 0.906 | 0.002 | 1.000 | down |
| TSPAN7 | RP11- | 0.918 | 0.001 | 1.000 | down |
| TSPAN7 | RP11-191L17.1 | 0.906 | 0.002 | 1.000 | down |
| TSPAN7 | RP11-165A20.3 | 0.930 | 0.001 | 1.000 | down |
| TSPAN7 | G000481 | 0.923 | 0.001 | 1.000 | down |
| TSPAN7 | RP11-285F16.1 | 0.927 | 0.001 | 1.000 | down |
| TSPAN7 | G050681 | 0.912 | 0.002 | 1.000 | down |
| TSPAN7 | HRAT5 | 0.938 | 0.001 | 1.000 | down |
| TSPAN7 | GAS5-AS1 | 0.911 | 0.002 | 1.000 | down |
| TSPAN7 | RP3-410B11.1 | 0.949 | 0.000 | 1.000 | down |
| TSPAN7 | G079654 | 0.905 | 0.002 | 1.000 | down |
| TSPAN7 | RP11-510M2.2 | 0.912 | 0.002 | 1.000 | down |
| TSPAN7 | FRMD6-AS1 | 0.921 | 0.001 | 1.000 | down |
| TSPAN7 | RP11-174G6.5 | 0.907 | 0.002 | 1.000 | down |
| TSPAN7 | AL355711 | 0.912 | 0.002 | 1.000 | down |
| TSPAN7 | CTD- | 0.914 | 0.001 | 1.000 | down |
| TSPAN7 | XLOC_011052 | 0.903 | 0.002 | 1.000 | down |
| TSPAN7 | RP1-202O8.3 | 0.966 | 0.000 | 1.000 | down |
| TSPAN7 | RP13-131K19.2 | 0.904 | 0.002 | 1.000 | down |

| | | | | | |
|---|---|---|---|---|---|
| TSPAN7 | FOXN3-AS1 | 0.982 | 0.000 | 1.000 | down |
| TSPAN7 | G019632 | 0.918 | 0.001 | 1.000 | down |
| TSPAN7 | G038972 | 0.932 | 0.001 | 1.000 | down |
| TSPAN7 | RP11-505K9.1 | 0.974 | 0.000 | 1.000 | down |
| TSPAN7 | RP11-2E11.5 | 0.949 | 0.000 | 1.000 | down |
| TSPAN7 | LOC148709 | 0.912 | 0.002 | 1.000 | down |
| TSPAN7 | AC092667.2 | 0.938 | 0.001 | 1.000 | down |
| TSPAN7 | LOC101928402 | 0.942 | 0.000 | 1.000 | down |
| TSPAN7 | GFOD1 | 0.906 | 0.002 | 1.000 | down |
| TSPAN7 | RP4-616B8.4 | 0.903 | 0.002 | 1.000 | down |
| TSPAN7 | DQ594967 | 0.958 | 0.000 | 1.000 | down |
| TSPAN7 | AP000255.6 | 0.917 | 0.001 | 1.000 | down |
| TSPAN7 | CTD- | 0.943 | 0.000 | 1.000 | down |
| TSPAN7 | RP11-254F7.2 | 0.940 | 0.001 | 1.000 | down |
| TSPAN7 | RP11-332H14.1 | 0.950 | 0.000 | 1.000 | down |
| TSPAN7 | RP1-261G23.7 | 0.916 | 0.001 | 1.000 | down |
| TSPAN7 | G023087 | 0.905 | 0.002 | 1.000 | down |
| TSPAN7 | RP11- | 0.958 | 0.000 | 1.000 | down |
| TSPAN7 | LOC100288846 | 0.949 | 0.000 | 1.000 | down |
| TSPAN7 | UBAC2-AS1 | 0.928 | 0.001 | 1.000 | down |
| TSPAN7 | RP11-565F19.2 | 0.955 | 0.000 | 1.000 | down |
| TSPAN7 | RP11-53I6.3 | 0.928 | 0.001 | 1.000 | down |
| TSPAN7 | AK055974 | 0.916 | 0.001 | 1.000 | down |
| TSPAN7 | RP11- | 0.921 | 0.001 | 1.000 | down |
| TSPAN7 | RP1-232L24.3 | 0.952 | 0.000 | 1.000 | down |
| TSPAN7 | RP11-371A19.2 | 0.928 | 0.001 | 1.000 | down |
| TSPAN7 | RP11-803B1.8 | 0.907 | 0.002 | 1.000 | down |
| TSPAN7 | BC040628 | 0.917 | 0.001 | 1.000 | down |
| TSPAN7 | G085202 | 0.929 | 0.001 | 1.000 | down |
| MAP2 | AP000438.2 | -0.972 | 0.000 | -1.000 | down |
| MAP2 | G045594 | -0.905 | 0.002 | -1.000 | down |
| MAP2 | uc.49 | -0.909 | 0.002 | -1.000 | down |
| MAP2 | XLOC_009306 | -0.904 | 0.002 | -1.000 | down |
| MAP2 | G020793 | -0.911 | 0.002 | -1.000 | down |
| MAP2 | EML2-AS1 | -0.962 | 0.000 | -1.000 | down |
| MAP2 | XLOC_012740 | -0.923 | 0.001 | -1.000 | down |
| MAP2 | G051586 | -0.906 | 0.002 | -1.000 | down |
| MAP2 | RP11-602N24.3 | 0.901 | 0.002 | 1.000 | down |
| MAP2 | G039106 | 0.905 | 0.002 | 1.000 | down |
| MAP2 | CTD- | 0.930 | 0.001 | 1.000 | down |
| MAP2 | LINC00652 | 0.933 | 0.001 | 1.000 | down |
| MAP2 | RP11-350N15.5 | 0.911 | 0.002 | 1.000 | down |
| MAP2 | G061591 | 0.977 | 0.000 | 1.000 | down |
| MAP2 | RP1-40E16.12 | 0.960 | 0.000 | 1.000 | down |
| MAP2 | GBAT2 | 0.958 | 0.000 | 1.000 | down |
| MAP2 | ZFX-AS1 | 0.945 | 0.000 | 1.000 | down |
| MAP2 | RP11-297J22.1 | 0.966 | 0.000 | 1.000 | down |
| MAP2 | RP11-179B2.2 | 0.936 | 0.001 | 1.000 | down |
| MAP2 | RP11-191L17.1 | 0.908 | 0.002 | 1.000 | down |
| MAP2 | G085410 | 0.924 | 0.001 | 1.000 | down |
| MAP2 | G089005 | 0.910 | 0.002 | 1.000 | down |
| MAP2 | RP11-496H1.2 | 0.937 | 0.001 | 1.000 | down |
| MAP2 | G073576 | 0.926 | 0.001 | 1.000 | down |
| MAP2 | XLOC_011052 | 0.914 | 0.002 | 1.000 | down |
| MAP2 | TNRC18P1 | 0.947 | 0.000 | 1.000 | down |
| MAP2 | G065067 | 0.924 | 0.001 | 1.000 | down |
| MAP2 | AC099850.1 | 0.912 | 0.002 | 1.000 | down |
| MAP2 | LINC00634 | 0.930 | 0.001 | 1.000 | down |
| MAP2 | RP11-51J9.5 | 0.900 | 0.002 | 1.000 | down |
| MAP2 | RP4-798A10.7 | 0.922 | 0.001 | 1.000 | down |
| MAP2 | LINC00630 | 0.911 | 0.002 | 1.000 | down |
| MAP2 | G010838 | 0.937 | 0.001 | 1.000 | down |
| MAP2 | RP11-573G6.8 | 0.961 | 0.000 | 1.000 | down |
| MAP2 | LL09NC01-139C3.1 | 0.931 | 0.001 | 1.000 | down |

| | | | | | |
|---|---|---|---|---|---|
| MAP2 | RP11- | 0.906 | 0.002 | 1.000 | down |
| MAP2 | G023340 | 0.952 | 0.000 | 1.000 | down |
| MAP2 | BC047644 | 0.908 | 0.002 | 1.000 | down |
| MAP2 | G001497 | 0.915 | 0.001 | 1.000 | down |
| MAP2 | RP11-439C15.4 | 0.926 | 0.001 | 1.000 | down |
| MAP2 | RP11- | 0.918 | 0.001 | 1.000 | down |
| MAP2 | CTC-248O19.1 | 0.918 | 0.001 | 1.000 | down |
| MAP2 | RP11-379F4.9 | 0.929 | 0.001 | 1.000 | down |
| CABP1 | XLOC_012303 | -0.925 | 0.001 | -1.000 | down |
| CABP1 | G087508 | -0.932 | 0.001 | -1.000 | down |
| CABP1 | G080487 | -0.922 | 0.001 | -1.000 | down |
| CABP1 | LINC00379 | -0.909 | 0.002 | -1.000 | down |
| CABP1 | G070831 | -0.954 | 0.000 | -1.000 | down |
| CABP1 | RP11-717D12.1 | -0.952 | 0.000 | -1.000 | down |
| CABP1 | AK093193 | -0.915 | 0.001 | -1.000 | down |
| CABP1 | CTD- | -0.910 | 0.002 | -1.000 | down |
| CABP1 | SNHG7 | -0.924 | 0.001 | -1.000 | down |
| CABP1 | G006360 | -0.916 | 0.001 | -1.000 | down |
| CABP1 | XLOC_013958 | -0.983 | 0.000 | -1.000 | down |
| CABP1 | G063524 | -0.916 | 0.001 | -1.000 | down |
| CABP1 | RP11-374P20.4 | -0.947 | 0.000 | -1.000 | down |
| CABP1 | G045736 | -0.917 | 0.001 | -1.000 | down |
| CABP1 | G016657 | -0.979 | 0.000 | -1.000 | down |
| CABP1 | RP11-30O15.1 | -0.921 | 0.001 | -1.000 | down |
| CABP1 | G067215 | -0.948 | 0.000 | -1.000 | down |
| CABP1 | G037029 | -0.981 | 0.000 | -1.000 | down |
| CABP1 | G010088 | -0.931 | 0.001 | -1.000 | down |
| CABP1 | G046492 | -0.902 | 0.002 | -1.000 | down |
| CABP1 | XLOC_000048 | -0.928 | 0.001 | -1.000 | down |
| CABP1 | G071259 | -0.963 | 0.000 | -1.000 | down |
| CABP1 | LOC100130417 | 0.962 | 0.000 | 1.000 | down |
| CABP1 | RP4-549L20.3 | 0.927 | 0.001 | 1.000 | down |
| CABP1 | USP27X-AS1 | 0.903 | 0.002 | 1.000 | down |
| CABP1 | BRWD1-IT2 | 0.964 | 0.000 | 1.000 | down |
| CABP1 | RP11-490O6.2 | 0.986 | 0.000 | 1.000 | down |
| CABP1 | RP11- | 0.938 | 0.001 | 1.000 | down |
| CABP1 | CTD- | 0.901 | 0.002 | 1.000 | down |
| CABP1 | CTD-2588C8.8 | 0.912 | 0.002 | 1.000 | down |
| CABP1 | RP1-309F20.3 | 0.980 | 0.000 | 1.000 | down |
| CABP1 | RP11- | 0.946 | 0.000 | 1.000 | down |
| CABP1 | RP11-191L17.1 | 0.929 | 0.001 | 1.000 | down |
| CABP1 | RP11-165A20.3 | 0.976 | 0.000 | 1.000 | down |
| CABP1 | G000481 | 0.972 | 0.000 | 1.000 | down |
| CABP1 | RP11-285F16.1 | 0.936 | 0.001 | 1.000 | down |
| CABP1 | G050681 | 0.942 | 0.000 | 1.000 | down |
| CABP1 | RP11-351D16.3 | 0.965 | 0.000 | 1.000 | down |
| CABP1 | AK123758 | 0.916 | 0.001 | 1.000 | down |
| CABP1 | RP4-683L5.1 | 0.908 | 0.002 | 1.000 | down |
| CABP1 | HRAT5 | 0.957 | 0.000 | 1.000 | down |
| CABP1 | RP11-312B8.1 | 0.920 | 0.001 | 1.000 | down |
| CABP1 | GAS5-AS1 | 0.918 | 0.001 | 1.000 | down |
| CABP1 | RP3-410B11.1 | 0.984 | 0.000 | 1.000 | down |
| CABP1 | G000619 | 0.940 | 0.001 | 1.000 | down |
| CABP1 | LOC101929741 | 0.935 | 0.001 | 1.000 | down |
| CABP1 | G079654 | 0.943 | 0.000 | 1.000 | down |
| CABP1 | RP11-510M2.2 | 0.900 | 0.002 | 1.000 | down |
| CABP1 | XLOC_006928 | 0.905 | 0.002 | 1.000 | down |
| CABP1 | RP11-2E11.10 | 0.931 | 0.001 | 1.000 | down |
| CABP1 | FRMD6-AS1 | 0.964 | 0.000 | 1.000 | down |
| CABP1 | RP11-174G6.5 | 0.927 | 0.001 | 1.000 | down |
| CABP1 | AL355711 | 0.983 | 0.000 | 1.000 | down |
| CABP1 | CTD- | 0.975 | 0.000 | 1.000 | down |
| CABP1 | BC070391 | 0.913 | 0.002 | 1.000 | down |
| CABP1 | RP1-202O8.3 | 0.976 | 0.000 | 1.000 | down |
| CABP1 | RP13-131K19.2 | 0.901 | 0.002 | 1.000 | down |

| | | | | | |
|---|---|---|---|---|---|
| CABP1 | FOXN3-AS1 | 0.973 | 0.000 | 1.000 | down |
| CABP1 | GSE61474_XLOC_038226 | 0.958 | 0.000 | 1.000 | down |
| CABP1 | G038972 | 0.969 | 0.000 | 1.000 | down |
| CABP1 | RP11-505K9.1 | 0.971 | 0.000 | 1.000 | down |
| CABP1 | RP11-138E2.1 | 0.903 | 0.002 | 1.000 | down |
| CABP1 | G003798 | 0.933 | 0.001 | 1.000 | down |
| CABP1 | AK026244 | 0.933 | 0.001 | 1.000 | down |
| CABP1 | RP11-2E11.5 | 0.933 | 0.001 | 1.000 | down |
| CABP1 | LOC148709 | 0.949 | 0.000 | 1.000 | down |
| CABP1 | G044581 | 0.916 | 0.001 | 1.000 | down |
| CABP1 | PVRL3-AS1 | 0.906 | 0.002 | 1.000 | down |
| CABP1 | CTD-2383M3.1 | 0.932 | 0.001 | 1.000 | down |
| CABP1 | AC092667.2 | 0.961 | 0.000 | 1.000 | down |
| CABP1 | RP11-376P6.3 | 0.955 | 0.000 | 1.000 | down |
| CABP1 | RP11-383C5.5 | 0.967 | 0.000 | 1.000 | down |
| CABP1 | G001739 | 0.906 | 0.002 | 1.000 | down |
| CABP1 | LOC101928402 | 0.933 | 0.001 | 1.000 | down |
| CABP1 | GFOD1 | 0.944 | 0.000 | 1.000 | down |
| CABP1 | LOC101928417 | 0.937 | 0.001 | 1.000 | down |
| CABP1 | RP4-555D20.3 | 0.935 | 0.001 | 1.000 | down |
| CABP1 | AP000255.6 | 0.935 | 0.001 | 1.000 | down |
| CABP1 | CTD- | 0.982 | 0.000 | 1.000 | down |
| CABP1 | RP11-332H14.1 | 0.986 | 0.000 | 1.000 | down |
| CABP1 | RP1-261G23.7 | 0.971 | 0.000 | 1.000 | down |
| CABP1 | G023087 | 0.951 | 0.000 | 1.000 | down |
| CABP1 | RP11- | 0.940 | 0.001 | 1.000 | down |
| CABP1 | G037470 | 0.947 | 0.000 | 1.000 | down |
| CABP1 | RP11-107N15.1 | 0.918 | 0.001 | 1.000 | down |
| CABP1 | LOC100288846 | 0.967 | 0.000 | 1.000 | down |
| CABP1 | UBAC2-AS1 | 0.905 | 0.002 | 1.000 | down |
| CABP1 | RP1-86C11.7 | 0.913 | 0.002 | 1.000 | down |
| CABP1 | FLJ20021 | 0.917 | 0.001 | 1.000 | down |
| CABP1 | RP11-61L19.2 | 0.940 | 0.001 | 1.000 | down |
| CABP1 | RP11-85A1.3 | 0.932 | 0.001 | 1.000 | down |
| CABP1 | RP11-565F19.2 | 0.915 | 0.001 | 1.000 | down |
| CABP1 | RP11-53I6.3 | 0.929 | 0.001 | 1.000 | down |
| CABP1 | AK055974 | 0.918 | 0.001 | 1.000 | down |
| CABP1 | G060549 | 0.904 | 0.002 | 1.000 | down |
| CABP1 | CTD- | 0.946 | 0.000 | 1.000 | down |
| CABP1 | RP1-232L24.3 | 0.950 | 0.000 | 1.000 | down |
| CABP1 | RP11-371A19.2 | 0.965 | 0.000 | 1.000 | down |
| CABP1 | RP11-820I16.4 | 0.938 | 0.001 | 1.000 | down |
| CABP1 | G075703 | 0.912 | 0.002 | 1.000 | down |
| CABP1 | LINC00858 | 0.935 | 0.001 | 1.000 | down |
| CABP1 | LINC01604 | 0.931 | 0.001 | 1.000 | down |
| CABP1 | RP11-803B1.8 | 0.925 | 0.001 | 1.000 | down |
| CABP1 | G015529 | 0.925 | 0.001 | 1.000 | down |
| CABP1 | CTD- | 0.948 | 0.000 | 1.000 | down |
| CABP1 | TOB1-AS1 (2) | 0.921 | 0.001 | 1.000 | down |
| DPYSL5 | LOC100499484-C9ORF174 | -0.902 | 0.002 | -1.000 | down |
| DPYSL5 | AP000438.2 | -0.933 | 0.001 | -1.000 | down |
| DPYSL5 | RP11-466A19.1 | -0.946 | 0.000 | -1.000 | down |
| DPYSL5 | XLOC_009306 | -0.914 | 0.001 | -1.000 | down |
| DPYSL5 | EML2-AS1 | -0.983 | 0.000 | -1.000 | down |
| DPYSL5 | XLOC_012740 | -0.908 | 0.002 | -1.000 | down |
| DPYSL5 | G013465 | -0.905 | 0.002 | -1.000 | down |
| DPYSL5 | G029241 | -0.904 | 0.002 | -1.000 | down |
| DPYSL5 | LOC100506834 | -0.939 | 0.001 | -1.000 | down |
| DPYSL5 | EF565122 | -0.959 | 0.000 | -1.000 | down |
| DPYSL5 | RP11-461K13.1 | -0.915 | 0.001 | -1.000 | down |
| DPYSL5 | RP11-602N24.3 | 0.919 | 0.001 | 1.000 | down |
| DPYSL5 | AL110181 | 0.913 | 0.002 | 1.000 | down |
| DPYSL5 | RP11-303E16.7 | 0.902 | 0.002 | 1.000 | down |

| | | | | | |
|---|---|---|---|---|---|
| DPYSL5 | RP11-686O6.2 | 0.959 | 0.000 | 1.000 | down |
| DPYSL5 | G061591 | 0.975 | 0.000 | 1.000 | down |
| DPYSL5 | RP1-40E16.12 | 0.969 | 0.000 | 1.000 | down |
| DPYSL5 | GBAT2 | 0.967 | 0.000 | 1.000 | down |
| DPYSL5 | RP11-297J22.1 | 0.977 | 0.000 | 1.000 | down |
| DPYSL5 | RP11-179B2.2 | 0.927 | 0.001 | 1.000 | down |
| DPYSL5 | RP11-191L17.1 | 0.904 | 0.002 | 1.000 | down |
| DPYSL5 | RP4-683L5.1 | 0.907 | 0.002 | 1.000 | down |
| DPYSL5 | AC144652.1 | 0.932 | 0.001 | 1.000 | down |
| DPYSL5 | G085410 | 0.927 | 0.001 | 1.000 | down |
| DPYSL5 | G089005 | 0.919 | 0.001 | 1.000 | down |
| DPYSL5 | G073576 | 0.951 | 0.000 | 1.000 | down |
| DPYSL5 | XLOC_013222 | 0.913 | 0.002 | 1.000 | down |
| DPYSL5 | TNRC18P1 | 0.941 | 0.000 | 1.000 | down |
| DPYSL5 | RP11-138E2.1 | 0.901 | 0.002 | 1.000 | down |
| DPYSL5 | LINC00634 | 0.958 | 0.000 | 1.000 | down |
| DPYSL5 | RP4-798A10.7 | 0.917 | 0.001 | 1.000 | down |
| DPYSL5 | RP1-223E5.4 | 0.924 | 0.001 | 1.000 | down |
| DPYSL5 | RP11-573G6.8 | 0.955 | 0.000 | 1.000 | down |
| DPYSL5 | LL09NC01-139C3.1 | 0.924 | 0.001 | 1.000 | down |
| DPYSL5 | G023340 | 0.966 | 0.000 | 1.000 | down |
| DPYSL5 | G006578 | 0.910 | 0.002 | 1.000 | down |
| DPYSL5 | RP11-79P5.9 | 0.918 | 0.001 | 1.000 | down |
| DPYSL5 | RP11-439C15.4 | 0.958 | 0.000 | 1.000 | down |
| DPYSL5 | RP11- | 0.917 | 0.001 | 1.000 | down |
| DPYSL5 | RP11-379F4.9 | 0.922 | 0.001 | 1.000 | down |
| RPS6KA5 | SLC7A11-AS1 | -0.903 | 0.002 | -1.000 | down |
| RPS6KA5 | XLOC_l2_009332 | -0.969 | 0.000 | -1.000 | down |
| RPS6KA5 | G045736 | -0.935 | 0.001 | -1.000 | down |
| RPS6KA5 | XLOC_006124 | -0.909 | 0.002 | -1.000 | down |
| RPS6KA5 | G041587 | -0.949 | 0.000 | -1.000 | down |
| RPS6KA5 | RP11-303E16.7 | 0.917 | 0.001 | 1.000 | down |
| RPS6KA5 | XLOC_008005 | 0.950 | 0.000 | 1.000 | down |
| RPS6KA5 | RP1-309F20.3 | 0.915 | 0.001 | 1.000 | down |
| RPS6KA5 | AC002467.7 | 0.908 | 0.002 | 1.000 | down |
| RPS6KA5 | AC093391.2 | 0.923 | 0.001 | 1.000 | down |
| RPS6KA5 | RP11-174G6.5 | 0.904 | 0.002 | 1.000 | down |
| RPS6KA5 | G082866 | 0.931 | 0.001 | 1.000 | down |
| RPS6KA5 | AP000255.6 | 0.909 | 0.002 | 1.000 | down |
| RPS6KA5 | AK000953 | 0.938 | 0.001 | 1.000 | down |
| RPS6KA5 | FLJ20021 | 0.920 | 0.001 | 1.000 | down |
| RPS6KA5 | RP11- | 0.913 | 0.002 | 1.000 | down |
| RPS6KA5 | G075319 | 0.900 | 0.002 | 1.000 | down |
| RPS6KA5 | GSE61474_XLOC_067608 | 0.982 | 0.000 | 1.000 | down |
| DPYSL3 | LOC100499484-C9ORF174 | -0.964 | 0.000 | -1.000 | down |
| DPYSL3 | SLC7A11-AS1 | -0.987 | 0.000 | -1.000 | down |
| DPYSL3 | RP11-466A19.1 | -0.967 | 0.000 | -1.000 | down |
| DPYSL3 | G003431 | -0.946 | 0.000 | -1.000 | down |
| DPYSL3 | G013465 | -0.943 | 0.000 | -1.000 | down |
| DPYSL3 | G029241 | -0.914 | 0.002 | -1.000 | down |
| DPYSL3 | EF565122 | -0.966 | 0.000 | -1.000 | down |
| DPYSL3 | CTD-2015A6.1 | -0.936 | 0.001 | -1.000 | down |
| DPYSL3 | G013179 | -0.926 | 0.001 | -1.000 | down |
| DPYSL3 | RP11-461K13.1 | -0.921 | 0.001 | -1.000 | down |
| DPYSL3 | G041587 | -0.920 | 0.001 | -1.000 | down |
| DPYSL3 | AL110181 | 0.904 | 0.002 | 1.000 | down |
| DPYSL3 | RP11-303E16.7 | 0.935 | 0.001 | 1.000 | down |
| DPYSL3 | RP11-686O6.2 | 0.937 | 0.001 | 1.000 | down |
| DPYSL3 | RP11-356J5.12 | 0.936 | 0.001 | 1.000 | down |
| DPYSL3 | GBAT2 | 0.900 | 0.002 | 1.000 | down |
| DPYSL3 | AC144652.1 | 0.949 | 0.000 | 1.000 | down |

| | | | | | |
|---|---|---|---|---|---|
| DPYSL3 | LINC00408 | 0.932 | 0.001 | 1.000 | down |
| DPYSL3 | G059716 | 0.927 | 0.001 | 1.000 | down |
| DPYSL3 | LINC00634 | 0.926 | 0.001 | 1.000 | down |
| DPYSL3 | LL22NC03-86G7.1 | 0.927 | 0.001 | 1.000 | down |
| DPYSL3 | FAM66D | 0.983 | 0.000 | 1.000 | down |
| DPYSL3 | RP11- | 0.925 | 0.001 | 1.000 | down |
| DPYSL3 | CTC- | 0.905 | 0.002 | 1.000 | down |
| DPYSL3 | RP11-344P13.6 | 0.923 | 0.001 | 1.000 | down |
| DPYSL3 | ANKRD20A12 | 0.950 | 0.000 | 1.000 | down |
| DPYSL3 | G006578 | 0.931 | 0.001 | 1.000 | down |
| DPYSL3 | G075319 | 0.952 | 0.000 | 1.000 | down |
| DPYSL3 | RP11-79P5.9 | 0.936 | 0.001 | 1.000 | down |
| DPYSL3 | RP11-439C15.4 | 0.914 | 0.001 | 1.000 | down |
| OLFM1 | CTC-462L7.1 | -0.933 | 0.001 | -1.000 | down |
| OLFM1 | AK124233 | -0.924 | 0.001 | -1.000 | down |
| OLFM1 | RP4-816N1.7 | -0.912 | 0.002 | -1.000 | down |
| OLFM1 | AQP7P3 | -0.946 | 0.000 | -1.000 | down |
| OLFM1 | AC074363.1 | -0.900 | 0.002 | -1.000 | down |
| OLFM1 | G082315 | -0.935 | 0.001 | -1.000 | down |
| OLFM1 | AX747064 | -0.977 | 0.000 | -1.000 | down |
| OLFM1 | LOC727896 | -0.964 | 0.000 | -1.000 | down |
| OLFM1 | CTD- | -0.905 | 0.002 | -1.000 | down |
| OLFM1 | G068469 | -0.972 | 0.000 | -1.000 | down |
| OLFM1 | AK024936 | -0.946 | 0.000 | -1.000 | down |
| OLFM1 | G090511 | 0.933 | 0.001 | 1.000 | down |
| OLFM1 | G017670 | 0.927 | 0.001 | 1.000 | down |
| OLFM1 | AC073321.4 | 0.936 | 0.001 | 1.000 | down |
| OLFM1 | XLOC_003595 | 0.969 | 0.000 | 1.000 | down |
| OLFM1 | RP11-382D12.2 | 0.937 | 0.001 | 1.000 | down |
| OLFM1 | LOC102724661 | 0.930 | 0.001 | 1.000 | down |
| OLFM1 | LINC01152 | 0.940 | 0.001 | 1.000 | down |
| OLFM1 | G058460 | 0.970 | 0.000 | 1.000 | down |
| OLFM1 | G047565 | 0.909 | 0.002 | 1.000 | down |
| OLFM1 | G022520 | 0.926 | 0.001 | 1.000 | down |
| BTBD9 | G038103 | -0.927 | 0.001 | -1.000 | down |
| BTBD9 | RP11-498C9.12 | -0.907 | 0.002 | -1.000 | down |
| BTBD9 | AP001625.6 | -0.923 | 0.001 | -1.000 | down |
| BTBD9 | RP11-669E14.6 | -0.924 | 0.001 | -1.000 | down |
| BTBD9 | NEAT1 | -0.978 | 0.000 | -1.000 | down |
| BTBD9 | G087395 | -0.924 | 0.001 | -1.000 | down |
| BTBD9 | uc.49 | -0.913 | 0.002 | -1.000 | down |
| BTBD9 | G065302 | -0.920 | 0.001 | -1.000 | down |
| BTBD9 | G072890 | -0.946 | 0.000 | -1.000 | down |
| BTBD9 | RMRPP3 | -0.932 | 0.001 | -1.000 | down |
| BTBD9 | uc.474 | -0.953 | 0.000 | -1.000 | down |
| BTBD9 | GOLGA8B | -0.942 | 0.000 | -1.000 | down |
| BTBD9 | G080747 | -0.945 | 0.000 | -1.000 | down |
| BTBD9 | LOC101926941 | -0.920 | 0.001 | -1.000 | down |
| BTBD9 | G009199 | -0.938 | 0.001 | -1.000 | down |
| BTBD9 | DLG3-AS1 | -0.906 | 0.002 | -1.000 | down |
| BTBD9 | XLOC_003048 | -0.959 | 0.000 | -1.000 | down |
| BTBD9 | RP11-355B11.2 | 0.973 | 0.000 | 1.000 | down |
| BTBD9 | SH3GL1P2 | 0.902 | 0.002 | 1.000 | down |
| BTBD9 | RP11-46J23.1 | 0.945 | 0.000 | 1.000 | down |
| BTBD9 | RP11-396C23.4 | 0.941 | 0.000 | 1.000 | down |
| BTBD9 | CTC-232P5.3 | 0.960 | 0.000 | 1.000 | down |
| BTBD9 | MGC2889 | 0.913 | 0.002 | 1.000 | down |
| BTBD9 | uc.85 | 0.903 | 0.002 | 1.000 | down |
| BTBD9 | OLMALINC | 0.934 | 0.001 | 1.000 | down |
| BTBD9 | CCNT2-AS1 | 0.906 | 0.002 | 1.000 | down |
| BTBD9 | G023041 | 0.901 | 0.002 | 1.000 | down |
| BTBD9 | NIPBL-AS1 | 0.909 | 0.002 | 1.000 | down |
| BTBD9 | G066437 | 0.909 | 0.002 | 1.000 | down |
| BTBD9 | RP11-413M3.4 | 0.965 | 0.000 | 1.000 | down |

| | | | | | |
|---|---|---|---|---|---|
| BTBD9 | CXADRP3 | 0.908 | 0.002 | 1.000 | down |
| BTBD9 | G047565 | 0.928 | 0.001 | 1.000 | down |
| BTBD9 | RP11-30L3.2 | 0.958 | 0.000 | 1.000 | down |
| BTBD9 | G014106 | 0.903 | 0.002 | 1.000 | down |
| BTBD9 | G061238 | 0.953 | 0.000 | 1.000 | down |
| BTBD9 | BC047644 | 0.919 | 0.001 | 1.000 | down |
| CHRNA7 | G071732 | -0.909 | 0.002 | -1.000 | down |
| CHRNA7 | G010088 | -0.917 | 0.001 | -1.000 | down |
| CHRNA7 | RP11-551L14.6 | -0.926 | 0.001 | -1.000 | down |
| CHRNA7 | RP11-651L5.2 | -0.911 | 0.002 | -1.000 | down |
| CHRNA7 | G068224 | -0.952 | 0.000 | -1.000 | down |
| CHRNA7 | SPTY2D1-AS1 | -0.960 | 0.000 | -1.000 | down |
| CHRNA7 | RP11- | 0.914 | 0.001 | 1.000 | down |
| CHRNA7 | G000481 | 0.902 | 0.002 | 1.000 | down |
| CHRNA7 | G000619 | 0.903 | 0.002 | 1.000 | down |
| CHRNA7 | FRMD6-AS1 | 0.911 | 0.002 | 1.000 | down |
| CHRNA7 | G081251 | 0.940 | 0.001 | 1.000 | down |
| CHRNA7 | AK026244 | 0.948 | 0.000 | 1.000 | down |
| CHRNA7 | LOC148709 | 0.918 | 0.001 | 1.000 | down |
| CHRNA7 | RP11-376P6.3 | 0.917 | 0.001 | 1.000 | down |
| CHRNA7 | G037470 | 0.913 | 0.002 | 1.000 | down |
| CHRNA7 | RP11-177G23.2 | 0.959 | 0.000 | 1.000 | down |
| CHRNA7 | G060549 | 0.921 | 0.001 | 1.000 | down |
| CHRNA7 | RP11-118K6.3 | 0.909 | 0.002 | 1.000 | down |
| CHRNA7 | RP11- | 0.908 | 0.002 | 1.000 | down |
| CHRNA7 | G075703 | 0.915 | 0.001 | 1.000 | down |
| CHRNA7 | LINC00858 | 0.923 | 0.001 | 1.000 | down |
| CHRNA7 | CTD- | 0.950 | 0.000 | 1.000 | down |
| CHRNA7 | TOB1-AS1 (2) | 0.931 | 0.001 | 1.000 | down |
| CHRNA7 | LINC00670 | 0.947 | 0.000 | 1.000 | down |
| PPP2R1A | RP11-70P17.1 | -0.907 | 0.002 | -1.000 | down |
| PPP2R1A | G060633 | -0.905 | 0.002 | -1.000 | down |
| PPP2R1A | G011310 | -0.946 | 0.000 | -1.000 | down |
| PPP2R1A | G008916 | -0.941 | 0.000 | -1.000 | down |
| PPP2R1A | AP000438.2 | -0.957 | 0.000 | -1.000 | down |
| PPP2R1A | AP001625.6 | -0.924 | 0.001 | -1.000 | down |
| PPP2R1A | NEAT1 | -0.905 | 0.002 | -1.000 | down |
| PPP2R1A | PMCHL2 | -0.919 | 0.001 | -1.000 | down |
| PPP2R1A | uc.49 | -0.926 | 0.001 | -1.000 | down |
| PPP2R1A | G073689 | -0.963 | 0.000 | -1.000 | down |
| PPP2R1A | G054530 | -0.934 | 0.001 | -1.000 | down |
| PPP2R1A | XLOC_012740 | -0.919 | 0.001 | -1.000 | down |
| PPP2R1A | LOC101926941 | -0.901 | 0.002 | -1.000 | down |
| PPP2R1A | TIPARP-AS1 | 0.963 | 0.000 | 1.000 | down |
| PPP2R1A | G039106 | 0.909 | 0.002 | 1.000 | down |
| PPP2R1A | LINC00652 | 0.945 | 0.000 | 1.000 | down |
| PPP2R1A | G061591 | 0.924 | 0.001 | 1.000 | down |
| PPP2R1A | RP1-40E16.12 | 0.920 | 0.001 | 1.000 | down |
| PPP2R1A | GBAT2 | 0.945 | 0.000 | 1.000 | down |
| PPP2R1A | RP11-297J22.1 | 0.938 | 0.001 | 1.000 | down |
| PPP2R1A | RP11-344E13.4 | 0.911 | 0.002 | 1.000 | down |
| PPP2R1A | MGC2889 | 0.933 | 0.001 | 1.000 | down |
| PPP2R1A | RP11-204E9.1 | 0.913 | 0.002 | 1.000 | down |
| PPP2R1A | OLMALINC | 0.961 | 0.000 | 1.000 | down |
| PPP2R1A | G023041 | 0.921 | 0.001 | 1.000 | down |
| PPP2R1A | NIPBL-AS1 | 0.934 | 0.001 | 1.000 | down |
| PPP2R1A | LINC00634 | 0.932 | 0.001 | 1.000 | down |
| PPP2R1A | RP11-51J9.5 | 0.966 | 0.000 | 1.000 | down |
| PPP2R1A | G066437 | 0.920 | 0.001 | 1.000 | down |
| PPP2R1A | DGCR5 | 0.934 | 0.001 | 1.000 | down |
| PPP2R1A | CXADRP3 | 0.906 | 0.002 | 1.000 | down |
| PPP2R1A | CTD-2653B5.1 | 0.903 | 0.002 | 1.000 | down |
| PPP2R1A | RP11-30L3.2 | 0.936 | 0.001 | 1.000 | down |
| PPP2R1A | G014106 | 0.902 | 0.002 | 1.000 | down |
| PPP2R1A | BC047644 | 0.963 | 0.000 | 1.000 | down |

| | | | | | |
|---|---|---|---|---|---|
| FXYD6 | RP11-674P19.2 | -0.956 | 0.000 | -1.000 | down |
| FXYD6 | G089550 | -0.945 | 0.000 | -1.000 | down |
| FXYD6 | RP4-816N1.7 | -0.961 | 0.000 | -1.000 | down |
| FXYD6 | LIPE-AS1 | -0.911 | 0.002 | -1.000 | down |
| FXYD6 | G000948 | -0.954 | 0.000 | -1.000 | down |
| FXYD6 | G033096 | -0.951 | 0.000 | -1.000 | down |
| FXYD6 | RP11-49K24.8 | 0.914 | 0.001 | 1.000 | down |
| FXYD6 | G086828 | 0.907 | 0.002 | 1.000 | down |
| FXYD6 | LOC102724661 | 0.923 | 0.001 | 1.000 | down |
| FXYD6 | G068576 | 0.943 | 0.000 | 1.000 | down |
| FXYD6 | G064611 | 0.932 | 0.001 | 1.000 | down |
| FXYD6 | RP11-362L22.1 | 0.914 | 0.001 | 1.000 | down |
| FXYD6 | G036604 | 0.913 | 0.002 | 1.000 | down |

**Supplementary Table 10. Dysregulated lncRNA-mitochondrial mRNAs co-expression network in propofol exposed organoids**

| lncRNAs (source genes) | Mitochondial mRNAs (target genes) | Correlation coefficient | P value | Positive/Negative Correlation Coefficient | Regulation |
|---|---|---|---|---|---|
| G037251 | PTRH2 | -0.911 | 0.002 | -1.000 | up |
| G000844 | SLC25A19 | -0.911 | 0.002 | -1.000 | up |
| G003271 | MT-ND1 | -0.916 | 0.001 | -1.000 | up |
| XLOC_012303 | IBA57 | -0.952 | 0.000 | -1.000 | up |
| XLOC_012303 | NBR1 | -0.927 | 0.001 | -1.000 | up |
| XLOC_012303 | CMPK2 | -0.911 | 0.002 | -1.000 | up |
| HCG18 | MT-ND1 | -0.961 | 0.000 | -1.000 | up |
| HCG18 | NBR1 | -0.908 | 0.002 | -1.000 | up |
| HCG18 | MSRA | -0.924 | 0.001 | -1.000 | up |
| HCG18 | FOXO3 | -0.950 | 0.000 | -1.000 | up |
| RP11-308N19.1 | SATB1 | -0.928 | 0.001 | -1.000 | up |
| RP1-296L11.1 | PTRH2 | -0.900 | 0.002 | -1.000 | up |
| G042193 | PKM | -0.901 | 0.002 | -1.000 | up |
| LOC100499484-C9ORF174 | PNPLA8 | -0.929 | 0.001 | -1.000 | up |
| G056910 | PKM | -0.912 | 0.002 | -1.000 | up |
| G056910 | UCP2 | -0.923 | 0.001 | -1.000 | up |
| SLC7A11-AS1 | ATAD3B | -0.912 | 0.002 | -1.000 | up |
| SLC7A11-AS1 | PNPLA8 | -0.929 | 0.001 | -1.000 | up |
| LINC00473 | NDUFAF7 | -0.905 | 0.002 | -1.000 | up |
| LINC00473 | PGS1 | -0.944 | 0.000 | -1.000 | up |
| CTC-462L7.1 | BNIP3L | -0.931 | 0.001 | -1.000 | up |
| RP11-70P17.1 | ATAD3B | -0.903 | 0.002 | -1.000 | up |
| G076532 | SATB1 | -0.915 | 0.001 | -1.000 | up |
| ITGA9-AS1 | PUS1 | -0.913 | 0.002 | -1.000 | up |
| ITGA9-AS1 | CKMT1A | -0.920 | 0.001 | -1.000 | up |
| PPIEL | SLC25A37 | -0.951 | 0.000 | -1.000 | up |
| PPIEL | WARS2 | -0.909 | 0.002 | -1.000 | up |
| G075352 | MSRA | -0.907 | 0.002 | -1.000 | up |
| G038103 | UCP2 | -0.920 | 0.001 | -1.000 | up |
| G074385 | LYRM7 | -0.930 | 0.001 | -1.000 | up |
| G038095 | MSRA | -0.915 | 0.001 | -1.000 | up |
| G038095 | FOXO3 | -0.936 | 0.001 | -1.000 | up |
| G066551 | BNIP3L | -0.926 | 0.001 | -1.000 | up |
| G066551 | OGG1 | -0.920 | 0.001 | -1.000 | up |
| XLOC_l2_00933 | AURKAIP1 | -0.920 | 0.001 | -1.000 | up |
| XLOC_l2_00933 | NBR1 | -0.909 | 0.002 | -1.000 | up |
| XLOC_l2_00933 | CMPK2 | -0.976 | 0.000 | -1.000 | up |
| Alpha_250_dna | PMPCA | -0.909 | 0.002 | -1.000 | up |
| Alpha_250_dna | TRAP1 | -0.919 | 0.001 | -1.000 | up |
| Alpha_250_dna | OGG1 | -0.901 | 0.002 | -1.000 | up |
| LOC102723439 | HSPA1B | -0.955 | 0.000 | -1.000 | up |
| G060633 | OGG1 | -0.911 | 0.002 | -1.000 | up |
| G029454 | IBA57 | -0.930 | 0.001 | -1.000 | up |
| G020731 | WARS2 | -0.920 | 0.001 | -1.000 | up |
| IGBP1-AS2 | PTRH2 | -0.905 | 0.002 | -1.000 | up |
| IGBP1-AS2 | PKM | -0.922 | 0.001 | -1.000 | up |
| IGBP1-AS2 | DNAJC19 | -0.901 | 0.002 | -1.000 | up |
| IGBP1-AS2 | NBR1 | -0.909 | 0.002 | -1.000 | up |
| G011310 | PMPCA | -0.903 | 0.002 | -1.000 | up |
| G011310 | PUS1 | -0.927 | 0.001 | -1.000 | up |
| G011310 | TRAP1 | -0.921 | 0.001 | -1.000 | up |
| G011310 | ATAD3B | -0.947 | 0.000 | -1.000 | up |
| G011310 | PNPLA8 | -0.916 | 0.001 | -1.000 | up |
| G011310 | OGG1 | -0.921 | 0.001 | -1.000 | up |
| ASAP1-IT1 | PMPCA | -0.930 | 0.001 | -1.000 | up |
| ASAP1-IT1 | AURKAIP1 | -0.904 | 0.002 | -1.000 | up |
| ASAP1-IT1 | TRAP1 | -0.960 | 0.000 | -1.000 | up |

| | | | | | |
|---|---|---|---|---|---|
| ASAP1-IT1 | HSPA1B | -0.910 | 0.002 | -1.000 | up |
| ASAP1-IT1 | PPRC1 | -0.926 | 0.001 | -1.000 | up |
| AK124233 | BNIP3L | -0.949 | 0.000 | -1.000 | up |
| G061035 | PDK1 | -0.926 | 0.001 | -1.000 | up |
| RP11-3K16.2 | PMPCA | -0.909 | 0.002 | -1.000 | up |
| RP11-3K16.2 | PTRH2 | -0.934 | 0.001 | -1.000 | up |
| RP11-3K16.2 | PUS1 | -0.911 | 0.002 | -1.000 | up |
| RP11-3K16.2 | CKMT1A | -0.936 | 0.001 | -1.000 | up |
| RP11-3K16.2 | CKMT1B | -0.947 | 0.000 | -1.000 | up |
| G008916 | PUS1 | -0.916 | 0.001 | -1.000 | up |
| G008916 | PGS1 | -0.929 | 0.001 | -1.000 | up |
| G008916 | OGG1 | -0.938 | 0.001 | -1.000 | up |
| RP11-187C18.2 | HSPE1 | -0.916 | 0.001 | -1.000 | up |
| G053606 | NBR1 | -0.926 | 0.001 | -1.000 | up |
| G087508 | LYRM7 | -0.956 | 0.000 | -1.000 | up |
| G087508 | CMPK2 | -0.901 | 0.002 | -1.000 | up |
| G068163 | IBA57 | -0.921 | 0.001 | -1.000 | up |
| LACTB2-AS1 | ABCF2 | -0.914 | 0.001 | -1.000 | up |
| G049993 | HSPE1 | -0.909 | 0.002 | -1.000 | up |
| G049993 | AURKAIP1 | -0.948 | 0.000 | -1.000 | up |
| G073552 | WARS2 | -0.917 | 0.001 | -1.000 | up |
| AQP7P3 | NDUFAF7 | -0.903 | 0.002 | -1.000 | up |
| AQP7P3 | BNIP3L | -0.908 | 0.002 | -1.000 | up |
| GSE61474_XLOC_020343 | PHB | -0.963 | 0.000 | -1.000 | up |
| GSE61474_XLOC_020343 | PGS1 | -0.908 | 0.002 | -1.000 | up |
| GSE61474_XLOC_020343 | CKMT1B | -0.912 | 0.002 | -1.000 | up |
| G009303 | MRPL12 | -0.964 | 0.000 | -1.000 | up |
| G057625 | ALDOC | -0.917 | 0.001 | -1.000 | up |
| CHKB-CPT1B | HSPA1B | -0.925 | 0.001 | -1.000 | up |
| G080487 | IBA57 | -0.961 | 0.000 | -1.000 | up |
| G080487 | CMPK2 | -0.914 | 0.001 | -1.000 | up |
| AP000438.2 | ATAD3B | -0.901 | 0.002 | -1.000 | up |
| AC074363.1 | PHB | -0.904 | 0.002 | -1.000 | up |
| AC074363.1 | NDUFAF4 | -0.902 | 0.002 | -1.000 | up |
| G046678 | HSPE1 | -0.922 | 0.001 | -1.000 | up |
| G046678 | PMPCA | -0.910 | 0.002 | -1.000 | up |
| G046678 | AURKAIP1 | -0.950 | 0.000 | -1.000 | up |
| G046678 | SLC25A37 | -0.905 | 0.002 | -1.000 | up |
| G046678 | HSPA1B | -0.917 | 0.001 | -1.000 | up |
| G046678 | WARS2 | -0.906 | 0.002 | -1.000 | up |
| G046678 | NBR1 | -0.902 | 0.002 | -1.000 | up |
| PHF2P2 | CKMT1B | -0.904 | 0.002 | -1.000 | up |
| G081487 | AURKAIP1 | -0.909 | 0.002 | -1.000 | up |
| G081487 | TRAP1 | -0.944 | 0.000 | -1.000 | up |
| G081487 | HSPA1B | -0.958 | 0.000 | -1.000 | up |
| G081487 | PPRC1 | -0.914 | 0.001 | -1.000 | up |
| G081487 | PDK1 | -0.903 | 0.002 | -1.000 | up |
| AC016682.1 | WARS2 | -0.915 | 0.001 | -1.000 | up |
| AC016682.1 | NBR1 | -0.911 | 0.002 | -1.000 | up |
| AC016682.1 | FOXO3 | -0.923 | 0.001 | -1.000 | up |
| RP11-466A19.1 | ATAD3B | -0.909 | 0.002 | -1.000 | up |
| RP11-466A19.1 | PNPLA8 | -0.937 | 0.001 | -1.000 | up |
| AP001625.6 | PUS1 | -0.925 | 0.001 | -1.000 | up |
| AP001625.6 | PDK1 | -0.917 | 0.001 | -1.000 | up |
| RP11-484O2.1 | AURKAIP1 | -0.929 | 0.001 | -1.000 | up |
| RP11-484O2.1 | UCP2 | -0.904 | 0.002 | -1.000 | up |
| RP11-484O2.1 | WARS2 | -0.908 | 0.002 | -1.000 | up |
| XLOC_008025 | PTRH2 | -0.913 | 0.002 | -1.000 | up |
| XLOC_008025 | IBA57 | -0.924 | 0.001 | -1.000 | up |
| G050702 | SLC25A37 | -0.906 | 0.002 | -1.000 | up |
| G050702 | WARS2 | -0.950 | 0.000 | -1.000 | up |
| G050702 | NBR1 | -0.901 | 0.002 | -1.000 | up |

| | | | | | |
|---|---|---|---|---|---|
| RP11-587H10.1 | PMPCA | -0.901 | 0.002 | -1.000 | up |
| RP11-737O24.3 | HSPE1 | -0.940 | 0.001 | -1.000 | up |
| RP11-737O24.3 | PMPCA | -0.947 | 0.000 | -1.000 | up |
| RP11-737O24.3 | PTRH2 | -0.929 | 0.001 | -1.000 | up |
| RP11-737O24.3 | OGG1 | -0.946 | 0.000 | -1.000 | up |
| XLOC_003405 | MRPL12 | -0.912 | 0.002 | -1.000 | up |
| XLOC_003405 | PPRC1 | -0.903 | 0.002 | -1.000 | up |
| XLOC_003405 | PDK1 | -0.935 | 0.001 | -1.000 | up |
| G015131 | MSRA | -0.954 | 0.000 | -1.000 | up |
| G015131 | FOXO3 | -0.941 | 0.001 | -1.000 | up |
| G009235 | PKM | -0.918 | 0.001 | -1.000 | up |
| G063995 | WARS2 | -0.903 | 0.002 | -1.000 | up |
| AADACL2-AS1 | CKMT1A | -0.915 | 0.001 | -1.000 | up |
| AADACL2-AS1 | CKMT1B | -0.905 | 0.002 | -1.000 | up |
| RP11-669E14.6 | PMPCA | -0.913 | 0.002 | -1.000 | up |
| RP11-669E14.6 | PUS1 | -0.969 | 0.000 | -1.000 | up |
| RP11-669E14.6 | TRAP1 | -0.909 | 0.002 | -1.000 | up |
| RP11-669E14.6 | PGS1 | -0.912 | 0.002 | -1.000 | up |
| RP11-669E14.6 | PPRC1 | -0.924 | 0.001 | -1.000 | up |
| RP11-669E14.6 | CKMT1B | -0.911 | 0.002 | -1.000 | up |
| NEAT1 | PUS1 | -0.932 | 0.001 | -1.000 | up |
| XLOC_012579 | UCP2 | -0.952 | 0.000 | -1.000 | up |
| XLOC_l2_01111 | IBA57 | -0.901 | 0.002 | -1.000 | up |
| XLOC_l2_01111 | PDK1 | -0.910 | 0.002 | -1.000 | up |
| RN7SKP264 | HSPE1 | -0.913 | 0.002 | -1.000 | up |
| AX747064 | BNIP3L | -0.967 | 0.000 | -1.000 | up |
| RP11-219C20.3 | PHB | -0.906 | 0.002 | -1.000 | up |
| RP11-219C20.3 | HSPE1 | -0.928 | 0.001 | -1.000 | up |
| RP11-219C20.3 | NDUFAF4 | -0.918 | 0.001 | -1.000 | up |
| G070831 | NBR1 | -0.909 | 0.002 | -1.000 | up |
| ZFAT-AS1 | SLC25A37 | -0.963 | 0.000 | -1.000 | up |
| ZFAT-AS1 | PKM | -0.912 | 0.002 | -1.000 | up |
| ZFAT-AS1 | WARS2 | -0.956 | 0.000 | -1.000 | up |
| G040015 | NDUFAF4 | -0.929 | 0.001 | -1.000 | up |
| G040015 | CCDC58 | -0.907 | 0.002 | -1.000 | up |
| G024041 | CCDC58 | -0.907 | 0.002 | -1.000 | up |
| G087265 | NDUFAF4 | -0.905 | 0.002 | -1.000 | up |
| G087265 | MT-ND1 | -0.942 | 0.000 | -1.000 | up |
| G072062 | PTRH2 | -0.921 | 0.001 | -1.000 | up |
| G072062 | IBA57 | -0.930 | 0.001 | -1.000 | up |
| Z69720.2 | HSPE1 | -0.948 | 0.000 | -1.000 | up |
| Z69720.2 | NDUFAF4 | -0.942 | 0.000 | -1.000 | up |
| Z69720.2 | EARS2 | -0.919 | 0.001 | -1.000 | up |
| Z69720.2 | MSRA | -0.942 | 0.000 | -1.000 | up |
| CTC-261N6.1 | MRPL12 | -0.932 | 0.001 | -1.000 | up |
| CTC-261N6.1 | TRAP1 | -0.920 | 0.001 | -1.000 | up |
| CTC-261N6.1 | PPRC1 | -0.978 | 0.000 | -1.000 | up |
| G081302 | HSPA1B | -0.934 | 0.001 | -1.000 | up |
| G081302 | PDK1 | -0.940 | 0.001 | -1.000 | up |
| BC034456 | HSPE1 | -0.940 | 0.001 | -1.000 | up |
| RP11-18D7.2 | PKM | -0.936 | 0.001 | -1.000 | up |
| RP11-18D7.2 | DNAJC19 | -0.904 | 0.002 | -1.000 | up |
| G071732 | SATB1 | -0.924 | 0.001 | -1.000 | up |
| G071640 | PPRC1 | -0.925 | 0.001 | -1.000 | up |
| G071640 | LYRM7 | -0.957 | 0.000 | -1.000 | up |
| AC107218.3 | PHB | -0.902 | 0.002 | -1.000 | up |
| AC107218.3 | NDUFAF7 | -0.909 | 0.002 | -1.000 | up |
| CTB-102L5.8 | MRPL12 | -0.935 | 0.001 | -1.000 | up |
| G047392 | PPRC1 | -0.908 | 0.002 | -1.000 | up |
| RP11-717D12.1 | LYRM7 | -0.919 | 0.001 | -1.000 | up |
| RP11-717D12.1 | IBA57 | -0.969 | 0.000 | -1.000 | up |
| AK093193 | SLC25A37 | -0.937 | 0.001 | -1.000 | up |
| AK093193 | WARS2 | -0.974 | 0.000 | -1.000 | up |
| AK093193 | NBR1 | -0.961 | 0.000 | -1.000 | up |
| LOC101927164 | MT-ND1 | -0.905 | 0.002 | -1.000 | up |

| | | | | | |
|---|---|---|---|---|---|
| LOC101927164 | NBR1 | -0.915 | 0.001 | -1.000 | up |
| G045594 | SLC25A19 | -0.929 | 0.001 | -1.000 | up |
| CTD-2050E21.1 | SLC25A37 | -0.933 | 0.001 | -1.000 | up |
| CTD-2050E21.1 | PKM | -0.940 | 0.001 | -1.000 | up |
| CTD-2050E21.1 | DNAJC19 | -0.950 | 0.000 | -1.000 | up |
| CTD-2050E21.1 | WARS2 | -0.910 | 0.002 | -1.000 | up |
| CTD-2050E21.1 | NBR1 | -0.943 | 0.000 | -1.000 | up |
| LOC727896 | EARS2 | -0.941 | 0.000 | -1.000 | up |
| LOC727896 | BNIP3L | -0.948 | 0.000 | -1.000 | up |
| G036779 | ALDOC | -0.907 | 0.002 | -1.000 | up |
| AC003682.16 | ALDOC | -0.964 | 0.000 | -1.000 | up |
| AC003682.16 | MSRA | -0.909 | 0.002 | -1.000 | up |
| CRAT37 | CCDC58 | -0.919 | 0.001 | -1.000 | up |
| SNHG7 | IBA57 | -0.905 | 0.002 | -1.000 | up |
| SNHG7 | NBR1 | -0.934 | 0.001 | -1.000 | up |
| G046027 | MSRA | -0.920 | 0.001 | -1.000 | up |
| RP1-124C6.1 | MRPL12 | -0.942 | 0.000 | -1.000 | up |
| RP1-124C6.1 | PPRC1 | -0.916 | 0.001 | -1.000 | up |
| G022034 | ALDOC | -0.971 | 0.000 | -1.000 | up |
| G022034 | BNIP3L | -0.902 | 0.002 | -1.000 | up |
| uc.49 | PMPCA | -0.948 | 0.000 | -1.000 | up |
| uc.49 | PUS1 | -0.973 | 0.000 | -1.000 | up |
| uc.49 | PGS1 | -0.927 | 0.001 | -1.000 | up |
| uc.49 | ATAD3B | -0.934 | 0.001 | -1.000 | up |
| uc.49 | OGG1 | -0.910 | 0.002 | -1.000 | up |
| G043880 | UCP2 | -0.937 | 0.001 | -1.000 | up |
| RP11-424I19.2 | WARS2 | -0.903 | 0.002 | -1.000 | up |
| RP11-424I19.2 | NBR1 | -0.926 | 0.001 | -1.000 | up |
| RP11-424I19.2 | FOXO3 | -0.909 | 0.002 | -1.000 | up |
| G006360 | IBA57 | -0.966 | 0.000 | -1.000 | up |
| XLOC_013958 | IBA57 | -0.920 | 0.001 | -1.000 | up |
| XLOC_013958 | NBR1 | -0.964 | 0.000 | -1.000 | up |
| XLOC_013958 | CMPK2 | -0.912 | 0.002 | -1.000 | up |
| XLOC_009306 | SLC25A19 | -0.904 | 0.002 | -1.000 | up |
| G060178 | WARS2 | -0.925 | 0.001 | -1.000 | up |
| G063524 | IBA57 | -0.910 | 0.002 | -1.000 | up |
| G063524 | CMPK2 | -0.929 | 0.001 | -1.000 | up |
| G071944 | MRPL12 | -0.968 | 0.000 | -1.000 | up |
| G011388 | ALDOC | -0.939 | 0.001 | -1.000 | up |
| G011388 | BNIP3L | -0.933 | 0.001 | -1.000 | up |
| G011388 | MSRA | -0.921 | 0.001 | -1.000 | up |
| XLOC_003837 | AURKAIP1 | -0.923 | 0.001 | -1.000 | up |
| XLOC_003837 | MRPL12 | -0.933 | 0.001 | -1.000 | up |
| XLOC_003837 | TRAP1 | -0.901 | 0.002 | -1.000 | up |
| XLOC_003837 | PPRC1 | -0.951 | 0.000 | -1.000 | up |
| BX648501 | SLC25A37 | -0.903 | 0.002 | -1.000 | up |
| BX648501 | PKM | -0.930 | 0.001 | -1.000 | up |
| BX648501 | DNAJC19 | -0.921 | 0.001 | -1.000 | up |
| BX648501 | WARS2 | -0.905 | 0.002 | -1.000 | up |
| BX648501 | NBR1 | -0.941 | 0.000 | -1.000 | up |
| G073689 | PMPCA | -0.918 | 0.001 | -1.000 | up |
| G073689 | ATAD3B | -0.907 | 0.002 | -1.000 | up |
| G073689 | OGG1 | -0.981 | 0.000 | -1.000 | up |
| G036156 | PHB | -0.934 | 0.001 | -1.000 | up |
| G036156 | NDUFAF4 | -0.971 | 0.000 | -1.000 | up |
| G036156 | CKMT1B | -0.916 | 0.001 | -1.000 | up |
| G032508 | SATB1 | -0.957 | 0.000 | -1.000 | up |
| CTD-3118D11.2 | NDUFAF7 | -0.926 | 0.001 | -1.000 | up |
| RP11-374P20.4 | NBR1 | -0.917 | 0.001 | -1.000 | up |
| G063659 | PTRH2 | -0.952 | 0.000 | -1.000 | up |
| G063659 | CKMT1A | -0.908 | 0.002 | -1.000 | up |
| G068469 | BNIP3L | -0.949 | 0.000 | -1.000 | up |
| G020793 | SLC25A19 | -0.912 | 0.002 | -1.000 | up |
| G054530 | OGG1 | -0.974 | 0.000 | -1.000 | up |
| CTD-2023M8.1 | PUS1 | -0.943 | 0.000 | -1.000 | up |

| | | | | | |
|---|---|---|---|---|---|
| CTD-2023M8.1 | PGS1 | -0.920 | 0.001 | -1.000 | up |
| CTD-2023M8.1 | CKMT1A | -0.947 | 0.000 | -1.000 | up |
| CTD-2023M8.1 | CKMT1B | -0.966 | 0.000 | -1.000 | up |
| G045736 | AURKAIP1 | -0.902 | 0.002 | -1.000 | up |
| G045736 | WARS2 | -0.936 | 0.001 | -1.000 | up |
| G045736 | NBR1 | -0.945 | 0.000 | -1.000 | up |
| G045736 | CMPK2 | -0.927 | 0.001 | -1.000 | up |
| EML2-AS1 | ATAD3B | -0.914 | 0.001 | -1.000 | up |
| EML2-AS1 | PNPLA8 | -0.945 | 0.000 | -1.000 | up |
| LINC01149 | IBA57 | -0.917 | 0.001 | -1.000 | up |
| G030124 | PTRH2 | -0.901 | 0.002 | -1.000 | up |
| G030124 | PGS1 | -0.934 | 0.001 | -1.000 | up |
| G030124 | CKMT1A | -0.951 | 0.000 | -1.000 | up |
| G065302 | HSPA1B | -0.958 | 0.000 | -1.000 | up |
| XLOC_006124 | CMPK2 | -0.932 | 0.001 | -1.000 | up |
| NBPF8 | AURKAIP1 | -0.913 | 0.002 | -1.000 | up |
| NBPF8 | MRPL12 | -0.955 | 0.000 | -1.000 | up |
| NBPF8 | PDK1 | -0.933 | 0.001 | -1.000 | up |
| LOC101927691 | PTRH2 | -0.912 | 0.002 | -1.000 | up |
| LOC101927691 | PDK1 | -0.913 | 0.002 | -1.000 | up |
| LINC00930 | MRPL12 | -0.975 | 0.000 | -1.000 | up |
| LINC00930 | PPRC1 | -0.904 | 0.002 | -1.000 | up |
| AK025321 | PTRH2 | -0.900 | 0.002 | -1.000 | up |
| G002461 | ALDOC | -0.938 | 0.001 | -1.000 | up |
| G002461 | MSRA | -0.929 | 0.001 | -1.000 | up |
| G002461 | FOXO3 | -0.907 | 0.002 | -1.000 | up |
| LOC101926908 | AURKAIP1 | -0.920 | 0.001 | -1.000 | up |
| LOC101926908 | PDK1 | -0.920 | 0.001 | -1.000 | up |
| G003431 | ATAD3B | -0.920 | 0.001 | -1.000 | up |
| G003431 | PNPLA8 | -0.902 | 0.002 | -1.000 | up |
| G001795 | PTRH2 | -0.904 | 0.002 | -1.000 | up |
| G016657 | IBA57 | -0.967 | 0.000 | -1.000 | up |
| G016657 | NBR1 | -0.922 | 0.001 | -1.000 | up |
| AP000344.4 | HSPE1 | -0.921 | 0.001 | -1.000 | up |
| AP000344.4 | NBR1 | -0.908 | 0.002 | -1.000 | up |
| AP000344.4 | MSRA | -0.921 | 0.001 | -1.000 | up |
| AP000344.4 | FOXO3 | -0.942 | 0.000 | -1.000 | up |
| RMRPP3 | PDK1 | -0.928 | 0.001 | -1.000 | up |
| XLOC_012740 | PMPCA | -0.935 | 0.001 | -1.000 | up |
| XLOC_012740 | PTRH2 | -0.903 | 0.002 | -1.000 | up |
| XLOC_012740 | PUS1 | -0.982 | 0.000 | -1.000 | up |
| XLOC_012740 | PGS1 | -0.918 | 0.001 | -1.000 | up |
| XLOC_012740 | ATAD3B | -0.958 | 0.000 | -1.000 | up |
| XLOC_012740 | PNPLA8 | -0.902 | 0.002 | -1.000 | up |
| XLOC_012740 | OGG1 | -0.904 | 0.002 | -1.000 | up |
| uc.474 | PUS1 | -0.906 | 0.002 | -1.000 | up |
| XLOC_006485 | PTRH2 | -0.935 | 0.001 | -1.000 | up |
| XLOC_006485 | UCP2 | -0.910 | 0.002 | -1.000 | up |
| TMLHE-AS1 | SLC25A37 | -0.905 | 0.002 | -1.000 | up |
| TMLHE-AS1 | HSPA1B | -0.959 | 0.000 | -1.000 | up |
| TMLHE-AS1 | UCP2 | -0.914 | 0.002 | -1.000 | up |
| RP1-65P5.3 | AURKAIP1 | -0.902 | 0.002 | -1.000 | up |
| RP1-65P5.3 | MRPL12 | -0.934 | 0.001 | -1.000 | up |
| RP1-65P5.3 | PDK1 | -0.939 | 0.001 | -1.000 | up |
| RP11-30O15.1 | NBR1 | -0.940 | 0.001 | -1.000 | up |
| G058409 | SLC25A37 | -0.947 | 0.000 | -1.000 | up |
| G058409 | HSPA1B | -0.950 | 0.000 | -1.000 | up |
| G009329 | SLC25A37 | -0.903 | 0.002 | -1.000 | up |
| G009329 | UCP2 | -0.907 | 0.002 | -1.000 | up |
| G009329 | PDK1 | -0.901 | 0.002 | -1.000 | up |
| G067215 | IBA57 | -0.906 | 0.002 | -1.000 | up |
| G067215 | NBR1 | -0.935 | 0.001 | -1.000 | up |
| G067215 | CMPK2 | -0.967 | 0.000 | -1.000 | up |
| RP11-10J18.3 | MSRA | -0.902 | 0.002 | -1.000 | up |
| G020470 | MSRA | -0.907 | 0.002 | -1.000 | up |

| | | | | | |
|---|---|---|---|---|---|
| LINC00685 | AURKAIP1 | -0.914 | 0.001 | -1.000 | up |
| LINC00685 | SLC25A37 | -0.901 | 0.002 | -1.000 | up |
| LINC00685 | PDK1 | -0.955 | 0.000 | -1.000 | up |
| G015181 | PTRH2 | -0.901 | 0.002 | -1.000 | up |
| G037029 | LYRM7 | -0.924 | 0.001 | -1.000 | up |
| G037029 | IBA57 | -0.961 | 0.000 | -1.000 | up |
| G037029 | NBR1 | -0.921 | 0.001 | -1.000 | up |
| G082713 | PMPCA | -0.909 | 0.002 | -1.000 | up |
| G082713 | TRAP1 | -0.912 | 0.002 | -1.000 | up |
| G010088 | SLC25A37 | -0.944 | 0.000 | -1.000 | up |
| G010088 | DNAJC19 | -0.953 | 0.000 | -1.000 | up |
| XLOC_008270 | MRPL12 | -0.971 | 0.000 | -1.000 | up |
| G060726 | UCP2 | -0.969 | 0.000 | -1.000 | up |
| G046492 | IBA57 | -0.905 | 0.002 | -1.000 | up |
| G046492 | NBR1 | -0.919 | 0.001 | -1.000 | up |
| AC026188.1 | HSPE1 | -0.926 | 0.001 | -1.000 | up |
| AC026188.1 | NDUFAF4 | -0.908 | 0.002 | -1.000 | up |
| G052905 | HSPE1 | -0.924 | 0.001 | -1.000 | up |
| G052905 | AURKAIP1 | -0.901 | 0.002 | -1.000 | up |
| RP11-551L14.6 | PTRH2 | -0.927 | 0.001 | -1.000 | up |
| RP11-551L14.6 | IBA57 | -0.916 | 0.001 | -1.000 | up |
| uc.50 | AURKAIP1 | -0.923 | 0.001 | -1.000 | up |
| uc.50 | NBR1 | -0.930 | 0.001 | -1.000 | up |
| uc.50 | CMPK2 | -0.929 | 0.001 | -1.000 | up |
| uc.50 | FOXO3 | -0.938 | 0.001 | -1.000 | up |
| RP4-760C5.3 | MT-ND1 | -0.933 | 0.001 | -1.000 | up |
| CTD-3193O13.1 | PMPCA | -0.930 | 0.001 | -1.000 | up |
| CTD-3193O13.1 | TRAP1 | -0.904 | 0.002 | -1.000 | up |
| CTD-3193O13.1 | LYRM7 | -0.936 | 0.001 | -1.000 | up |
| G072257 | IBA57 | -0.926 | 0.001 | -1.000 | up |
| AF086132 | ALDOC | -0.944 | 0.000 | -1.000 | up |
| AF086132 | BNIP3L | -0.912 | 0.002 | -1.000 | up |
| AF086132 | MSRA | -0.946 | 0.000 | -1.000 | up |
| AF086132 | FOXO3 | -0.919 | 0.001 | -1.000 | up |
| CTC-250I14.3 | NDUFAF7 | -0.914 | 0.001 | -1.000 | up |
| CTC-250I14.3 | CCDC58 | -0.913 | 0.002 | -1.000 | up |
| G051586 | PMPCA | -0.920 | 0.001 | -1.000 | up |
| G051586 | PTRH2 | -0.947 | 0.000 | -1.000 | up |
| G051586 | OGG1 | -0.942 | 0.000 | -1.000 | up |
| RP11-126O1.4 | HSPA1B | -0.956 | 0.000 | -1.000 | up |
| RP11-651L5.2 | SLC25A37 | -0.918 | 0.001 | -1.000 | up |
| RP11-651L5.2 | PKM | -0.905 | 0.002 | -1.000 | up |
| RP11-651L5.2 | DNAJC19 | -0.967 | 0.000 | -1.000 | up |
| XLOC_000048 | NBR1 | -0.912 | 0.002 | -1.000 | up |
| G053358 | PPRC1 | -0.903 | 0.002 | -1.000 | up |
| BCRP3 | MRPL12 | -0.914 | 0.001 | -1.000 | up |
| LOC105373609 | LYRM7 | -0.911 | 0.002 | -1.000 | up |
| G009199 | HSPA1B | -0.900 | 0.002 | -1.000 | up |
| G033096 | ABCF2 | -0.915 | 0.001 | -1.000 | up |
| G089283 | PMPCA | -0.973 | 0.000 | -1.000 | up |
| G089283 | TRAP1 | -0.921 | 0.001 | -1.000 | up |
| G089283 | PPRC1 | -0.932 | 0.001 | -1.000 | up |
| G089283 | LYRM7 | -0.901 | 0.002 | -1.000 | up |
| XLOC_007868 | EARS2 | -0.972 | 0.000 | -1.000 | up |
| AC025442.3 | NDUFAF7 | -0.924 | 0.001 | -1.000 | up |
| AC025442.3 | PGS1 | -0.950 | 0.000 | -1.000 | up |
| G013465 | PGS1 | -0.923 | 0.001 | -1.000 | up |
| G013465 | ATAD3B | -0.970 | 0.000 | -1.000 | up |
| G013465 | PNPLA8 | -0.952 | 0.000 | -1.000 | up |
| G071259 | IBA57 | -0.910 | 0.002 | -1.000 | up |
| G071259 | NBR1 | -0.919 | 0.001 | -1.000 | up |
| G071259 | CMPK2 | -0.926 | 0.001 | -1.000 | up |
| RP5-1112D6.7 | PMPCA | -0.910 | 0.002 | -1.000 | up |
| RP5-1112D6.7 | ATAD3B | -0.930 | 0.001 | -1.000 | up |
| RP5-1112D6.7 | LYRM7 | -0.958 | 0.000 | -1.000 | up |

| | | | | | |
|---|---|---|---|---|---|
| RP5-1112D6.7 | PNPLA8 | -0.926 | 0.001 | -1.000 | up |
| MIMT1 | MRPL12 | -0.910 | 0.002 | -1.000 | up |
| MIMT1 | PPRC1 | -0.923 | 0.001 | -1.000 | up |
| G068890 | MT-ND1 | -0.965 | 0.000 | -1.000 | up |
| G068890 | PKM | -0.903 | 0.002 | -1.000 | up |
| G068890 | NBR1 | -0.906 | 0.002 | -1.000 | up |
| G068890 | FOXO3 | -0.907 | 0.002 | -1.000 | up |
| G010941 | CCDC58 | -0.917 | 0.001 | -1.000 | up |
| SPTY2D1-AS1 | DNAJC19 | -0.907 | 0.002 | -1.000 | up |
| SPTY2D1-AS1 | SATB1 | -0.929 | 0.001 | -1.000 | up |
| G031271 | PUS1 | -0.930 | 0.001 | -1.000 | up |
| G029241 | PUS1 | -0.951 | 0.000 | -1.000 | up |
| G029241 | PGS1 | -0.963 | 0.000 | -1.000 | up |
| G029241 | ATAD3A | -0.975 | 0.000 | -1.000 | up |
| G029241 | ATAD3B | -0.969 | 0.000 | -1.000 | up |
| G029241 | CKMT1B | -0.950 | 0.000 | -1.000 | up |
| G029241 | PNPLA8 | -0.926 | 0.001 | -1.000 | up |
| AK022848 | MT-ND1 | -0.918 | 0.001 | -1.000 | up |
| G051433 | AURKAIP1 | -0.965 | 0.000 | -1.000 | up |
| NBPF25P | AURKAIP1 | -0.927 | 0.001 | -1.000 | up |
| DLG3-AS1 | HSPE1 | -0.918 | 0.001 | -1.000 | up |
| DLG3-AS1 | PMPCA | -0.923 | 0.001 | -1.000 | up |
| DLG3-AS1 | PTRH2 | -0.944 | 0.000 | -1.000 | up |
| DLG3-AS1 | PUS1 | -0.928 | 0.001 | -1.000 | up |
| DLG3-AS1 | OGG1 | -0.935 | 0.001 | -1.000 | up |
| EF565122 | ATAD3B | -0.912 | 0.002 | -1.000 | up |
| EF565122 | PNPLA8 | -0.940 | 0.001 | -1.000 | up |
| G088912 | PHB | -0.903 | 0.002 | -1.000 | up |
| G088912 | PGS1 | -0.932 | 0.001 | -1.000 | up |
| G084746 | MSRA | -0.904 | 0.002 | -1.000 | up |
| G084746 | FOXO3 | -0.908 | 0.002 | -1.000 | up |
| G013179 | LYRM7 | -0.924 | 0.001 | -1.000 | up |
| G013179 | PNPLA8 | -0.919 | 0.001 | -1.000 | up |
| GSE61474_XLOC_028262 | UCP2 | -0.951 | 0.000 | -1.000 | up |
| RN7SL176P | AURKAIP1 | -0.910 | 0.002 | -1.000 | up |
| RN7SL176P | PDK1 | -0.907 | 0.002 | -1.000 | up |
| G074298 | PPRC1 | -0.959 | 0.000 | -1.000 | up |
| RP11-354H21.2 | UCP2 | -0.925 | 0.001 | -1.000 | up |
| G010533 | PTRH2 | -0.931 | 0.001 | -1.000 | up |
| G010533 | HSPA1B | -0.910 | 0.002 | -1.000 | up |
| G010533 | PDK1 | -0.948 | 0.000 | -1.000 | up |
| G010224 | PUS1 | -0.927 | 0.001 | -1.000 | up |
| G010224 | PGS1 | -0.937 | 0.001 | -1.000 | up |
| G010224 | ATAD3A | -0.938 | 0.001 | -1.000 | up |
| G010224 | CKMT1A | -0.933 | 0.001 | -1.000 | up |
| G010224 | CKMT1B | -0.979 | 0.000 | -1.000 | up |
| G030745 | HSPE1 | -0.908 | 0.002 | -1.000 | up |
| AK024936 | BNIP3L | -0.914 | 0.001 | -1.000 | up |
| XLOC_009429 | TRAP1 | -0.901 | 0.002 | -1.000 | up |
| XLOC_009429 | HSPA1B | -0.967 | 0.000 | -1.000 | up |
| RP11-461K13.1 | ATAD3B | -0.934 | 0.001 | -1.000 | up |
| RP11-461K13.1 | LYRM7 | -0.932 | 0.001 | -1.000 | up |
| RP11-461K13.1 | PNPLA8 | -0.989 | 0.000 | -1.000 | up |
| G041587 | CMPK2 | -0.952 | 0.000 | -1.000 | up |
| G059595 | PMPCA | -0.958 | 0.000 | -1.000 | up |
| G059595 | AURKAIP1 | -0.925 | 0.001 | -1.000 | up |
| G059595 | TRAP1 | -0.947 | 0.000 | -1.000 | up |
| G059595 | SLC25A37 | -0.920 | 0.001 | -1.000 | up |
| G059595 | HSPA1B | -0.937 | 0.001 | -1.000 | up |
| G059595 | PPRC1 | -0.927 | 0.001 | -1.000 | up |
| G059595 | WARS2 | -0.920 | 0.001 | -1.000 | up |
| G059595 | NBR1 | -0.900 | 0.002 | -1.000 | up |
| PHB | GSE61474_XLOC_020343 | -0.963 | 0.000 | -1.000 | up |

| | | | | | |
|---|---|---|---|---|---|
| PHB | AC074363.1 | -0.904 | 0.002 | -1.000 | up |
| PHB | RP11-219C20.3 | -0.906 | 0.002 | -1.000 | up |
| PHB | AC107218.3 | -0.902 | 0.002 | -1.000 | up |
| PHB | G036156 | -0.934 | 0.001 | -1.000 | up |
| PHB | G088912 | -0.903 | 0.002 | -1.000 | up |
| PHB | G079009 | 0.914 | 0.002 | 1.000 | up |
| PHB | FAM66A | 0.921 | 0.001 | 1.000 | up |
| PHB | RP11-475I24.3 | 0.922 | 0.001 | 1.000 | up |
| PHB | G061579 | 0.914 | 0.001 | 1.000 | up |
| PHB | G031140 | 0.914 | 0.002 | 1.000 | up |
| PHB | G070622 | 0.911 | 0.002 | 1.000 | up |
| PHB | RP11-355F22.1 | 0.903 | 0.002 | 1.000 | up |
| HSPE1 | RP11-187C18.2 | -0.916 | 0.001 | -1.000 | up |
| HSPE1 | G049993 | -0.909 | 0.002 | -1.000 | up |
| HSPE1 | G046678 | -0.922 | 0.001 | -1.000 | up |
| HSPE1 | RP11-737O24.3 | -0.940 | 0.001 | -1.000 | up |
| HSPE1 | RN7SKP264 | -0.913 | 0.002 | -1.000 | up |
| HSPE1 | RP11-219C20.3 | -0.928 | 0.001 | -1.000 | up |
| HSPE1 | Z69720.2 | -0.948 | 0.000 | -1.000 | up |
| HSPE1 | BC034456 | -0.940 | 0.001 | -1.000 | up |
| HSPE1 | AP000344.4 | -0.921 | 0.001 | -1.000 | up |
| HSPE1 | AC026188.1 | -0.926 | 0.001 | -1.000 | up |
| HSPE1 | G052905 | -0.924 | 0.001 | -1.000 | up |
| HSPE1 | DLG3-AS1 | -0.918 | 0.001 | -1.000 | up |
| HSPE1 | G030745 | -0.908 | 0.002 | -1.000 | up |
| HSPE1 | GSE61474_XLOC_054139 | 0.926 | 0.001 | 1.000 | up |
| HSPE1 | GS1-433O24.1 | 0.923 | 0.001 | 1.000 | up |
| HSPE1 | G017718 | 0.901 | 0.002 | 1.000 | up |
| HSPE1 | AC073321.4 | 0.919 | 0.001 | 1.000 | up |
| HSPE1 | RP11-290L1.2 | 0.920 | 0.001 | 1.000 | up |
| HSPE1 | IDH1-AS1 | 0.904 | 0.002 | 1.000 | up |
| HSPE1 | G063668 | 0.909 | 0.002 | 1.000 | up |
| HSPE1 | ENO1-AS1 | 0.927 | 0.001 | 1.000 | up |
| HSPE1 | CTD-2382E5.2 | 0.951 | 0.000 | 1.000 | up |
| HSPE1 | AK125078 | 0.943 | 0.000 | 1.000 | up |
| HSPE1 | RP11-820L6.1 | 0.923 | 0.001 | 1.000 | up |
| HSPE1 | G005619 | 0.951 | 0.000 | 1.000 | up |
| PMPCA | Alpha_250_dna | -0.909 | 0.002 | -1.000 | up |
| PMPCA | G011310 | -0.903 | 0.002 | -1.000 | up |
| PMPCA | ASAP1-IT1 | -0.930 | 0.001 | -1.000 | up |
| PMPCA | RP11-3K16.2 | -0.909 | 0.002 | -1.000 | up |
| PMPCA | G046678 | -0.910 | 0.002 | -1.000 | up |
| PMPCA | RP11-587H10.1 | -0.901 | 0.002 | -1.000 | up |
| PMPCA | RP11-737O24.3 | -0.947 | 0.000 | -1.000 | up |
| PMPCA | RP11-669E14.6 | -0.913 | 0.002 | -1.000 | up |
| PMPCA | uc.49 | -0.948 | 0.000 | -1.000 | up |
| PMPCA | G073689 | -0.918 | 0.001 | -1.000 | up |
| PMPCA | XLOC_012740 | -0.935 | 0.001 | -1.000 | up |
| PMPCA | G082713 | -0.909 | 0.002 | -1.000 | up |
| PMPCA | CTD-3193O13.1 | -0.930 | 0.001 | -1.000 | up |
| PMPCA | G051586 | -0.920 | 0.001 | -1.000 | up |
| PMPCA | G089283 | -0.973 | 0.000 | -1.000 | up |
| PMPCA | RP5-1112D6.7 | -0.910 | 0.002 | -1.000 | up |
| PMPCA | DLG3-AS1 | -0.923 | 0.001 | -1.000 | up |
| PMPCA | G059595 | -0.958 | 0.000 | -1.000 | up |
| PMPCA | IFT74-AS1 | 0.917 | 0.001 | 1.000 | up |
| PMPCA | AL110181 | 0.941 | 0.000 | 1.000 | up |
| PMPCA | RP11-255E6.6 | 0.923 | 0.001 | 1.000 | up |
| PMPCA | LINC00652 | 0.918 | 0.001 | 1.000 | up |
| PMPCA | RP1-40E16.12 | 0.919 | 0.001 | 1.000 | up |
| PMPCA | DLG5-AS1 (1) | 0.907 | 0.002 | 1.000 | up |
| PMPCA | SH3GL1P2 | 0.917 | 0.001 | 1.000 | up |
| PMPCA | TM4SF1-AS1 | 0.914 | 0.001 | 1.000 | up |
| PMPCA | LINC00662 | 0.926 | 0.001 | 1.000 | up |

| | | | | | |
|---|---|---|---|---|---|
| PMPCA | RP11-179B2.2 | 0.911 | 0.002 | 1.000 | up |
| PMPCA | G064655 | 0.920 | 0.001 | 1.000 | up |
| PMPCA | RP11-715J22.6 | 0.963 | 0.000 | 1.000 | up |
| PMPCA | AC099754.1 | 0.945 | 0.000 | 1.000 | up |
| PMPCA | RP11-396C23.4 | 0.924 | 0.001 | 1.000 | up |
| PMPCA | RP11-285F16.1 | 0.902 | 0.002 | 1.000 | up |
| PMPCA | CTC-232P5.3 | 0.928 | 0.001 | 1.000 | up |
| PMPCA | RP5-991G20.4 | 0.934 | 0.001 | 1.000 | up |
| PMPCA | MGC2889 | 0.963 | 0.000 | 1.000 | up |
| PMPCA | G085410 | 0.920 | 0.001 | 1.000 | up |
| PMPCA | OLMALINC | 0.923 | 0.001 | 1.000 | up |
| PMPCA | G052526 | 0.905 | 0.002 | 1.000 | up |
| PMPCA | CCNT2-AS1 | 0.966 | 0.000 | 1.000 | up |
| PMPCA | XLOC_011052 | 0.925 | 0.001 | 1.000 | up |
| PMPCA | RP11-775D22.3 | 0.927 | 0.001 | 1.000 | up |
| PMPCA | G078120 | 0.940 | 0.001 | 1.000 | up |
| PMPCA | RP11-254F7.2 | 0.932 | 0.001 | 1.000 | up |
| PMPCA | G066437 | 0.939 | 0.001 | 1.000 | up |
| PMPCA | ENO1-AS1 | 0.944 | 0.000 | 1.000 | up |
| PMPCA | DGCR5 | 0.937 | 0.001 | 1.000 | up |
| PMPCA | LINC00463 | 0.910 | 0.002 | 1.000 | up |
| PMPCA | G027543 | 0.944 | 0.000 | 1.000 | up |
| PMPCA | LL09NC01-139C3.1 | 0.928 | 0.001 | 1.000 | up |
| PMPCA | G068576 | 0.936 | 0.001 | 1.000 | up |
| PMPCA | XLOC_004632 | 0.904 | 0.002 | 1.000 | up |
| PMPCA | CXADRP3 | 0.932 | 0.001 | 1.000 | up |
| PMPCA | RP11-110G21.1 | 0.978 | 0.000 | 1.000 | up |
| PMPCA | BX248253 | 0.956 | 0.000 | 1.000 | up |
| PMPCA | uc.349 | 0.948 | 0.000 | 1.000 | up |
| PMPCA | G003603 | 0.942 | 0.000 | 1.000 | up |
| PMPCA | RP11-53I6.3 | 0.952 | 0.000 | 1.000 | up |
| PMPCA | RP11-314A20.5 | 0.954 | 0.000 | 1.000 | up |
| PMPCA | RP11-196G18.24 | 0.923 | 0.001 | 1.000 | up |
| PMPCA | LOC101927811 | 0.922 | 0.001 | 1.000 | up |
| PMPCA | G014106 | 0.927 | 0.001 | 1.000 | up |
| PMPCA | RP11-355F22.1 | 0.937 | 0.001 | 1.000 | up |
| PMPCA | LOC283140 | 0.907 | 0.002 | 1.000 | up |
| PMPCA | DNAJC9-AS1 | 0.907 | 0.002 | 1.000 | up |
| PMPCA | CTB-181F24.1 | 0.909 | 0.002 | 1.000 | up |
| PMPCA | BC047644 | 0.908 | 0.002 | 1.000 | up |
| PMPCA | LOC102723385 | 0.965 | 0.000 | 1.000 | up |
| PMPCA | G063721 | 0.913 | 0.002 | 1.000 | up |
| PMPCA | G001497 | 0.949 | 0.000 | 1.000 | up |
| PMPCA | CTC-248O19.1 | 0.927 | 0.001 | 1.000 | up |
| SLC25A19 | G000844 | -0.911 | 0.002 | -1.000 | up |
| SLC25A19 | G045594 | -0.929 | 0.001 | -1.000 | up |
| SLC25A19 | XLOC_009306 | -0.904 | 0.002 | -1.000 | up |
| SLC25A19 | G020793 | -0.912 | 0.002 | -1.000 | up |
| SLC25A19 | TAF1A-AS1 | 0.923 | 0.001 | 1.000 | up |
| SLC25A19 | G048385 | 0.963 | 0.000 | 1.000 | up |
| SLC25A19 | SIX3-AS1 | 0.933 | 0.001 | 1.000 | up |
| SLC25A19 | LOC101928489 | 0.921 | 0.001 | 1.000 | up |
| NDUFAF4 | AC074363.1 | -0.902 | 0.002 | -1.000 | up |
| NDUFAF4 | RP11-219C20.3 | -0.918 | 0.001 | -1.000 | up |
| NDUFAF4 | G040015 | -0.929 | 0.001 | -1.000 | up |
| NDUFAF4 | G087265 | -0.905 | 0.002 | -1.000 | up |
| NDUFAF4 | Z69720.2 | -0.942 | 0.000 | -1.000 | up |
| NDUFAF4 | G036156 | -0.971 | 0.000 | -1.000 | up |
| NDUFAF4 | AC026188.1 | -0.908 | 0.002 | -1.000 | up |
| NDUFAF4 | RP11-44N22.3 | 0.902 | 0.002 | 1.000 | up |
| NDUFAF4 | BC047090 | 0.911 | 0.002 | 1.000 | up |
| NDUFAF4 | XLOC_013660 | 0.933 | 0.001 | 1.000 | up |
| NDUFAF4 | XLOC_004894 | 0.902 | 0.002 | 1.000 | up |
| NDUFAF7 | LINC00473 | -0.905 | 0.002 | -1.000 | up |
| NDUFAF7 | AQP7P3 | -0.903 | 0.002 | -1.000 | up |

| | | | | | |
|---|---|---|---|---|---|
| NDUFAF7 | AC107218.3 | -0.909 | 0.002 | -1.000 | up |
| NDUFAF7 | CTD-3118D11.2 | -0.926 | 0.001 | -1.000 | up |
| NDUFAF7 | CTC-250I14.3 | -0.914 | 0.001 | -1.000 | up |
| NDUFAF7 | AC025442.3 | -0.924 | 0.001 | -1.000 | up |
| NDUFAF7 | RP11-762I7.4 | 0.907 | 0.002 | 1.000 | up |
| NDUFAF7 | RP11-344E13.4 | 0.900 | 0.002 | 1.000 | up |
| NDUFAF7 | G014329 | 0.927 | 0.001 | 1.000 | up |
| NDUFAF7 | G066319 | 0.929 | 0.001 | 1.000 | up |
| NDUFAF7 | FAM66A | 0.935 | 0.001 | 1.000 | up |
| NDUFAF7 | RP11-475I24.3 | 0.934 | 0.001 | 1.000 | up |
| NDUFAF7 | LINC00408 | 0.938 | 0.001 | 1.000 | up |
| AURKAIP1 | XLOC_l2_009332 | -0.920 | 0.001 | -1.000 | up |
| AURKAIP1 | ASAP1-IT1 | -0.904 | 0.002 | -1.000 | up |
| AURKAIP1 | G049993 | -0.948 | 0.000 | -1.000 | up |
| AURKAIP1 | G046678 | -0.950 | 0.000 | -1.000 | up |
| AURKAIP1 | G081487 | -0.909 | 0.002 | -1.000 | up |
| AURKAIP1 | RP11-484O2.1 | -0.929 | 0.001 | -1.000 | up |
| AURKAIP1 | XLOC_003837 | -0.923 | 0.001 | -1.000 | up |
| AURKAIP1 | G045736 | -0.902 | 0.002 | -1.000 | up |
| AURKAIP1 | NBPF8 | -0.913 | 0.002 | -1.000 | up |
| AURKAIP1 | LOC101926908 | -0.920 | 0.001 | -1.000 | up |
| AURKAIP1 | RP1-65P5.3 | -0.902 | 0.002 | -1.000 | up |
| AURKAIP1 | LINC00685 | -0.914 | 0.001 | -1.000 | up |
| AURKAIP1 | G052905 | -0.901 | 0.002 | -1.000 | up |
| AURKAIP1 | uc.50 | -0.923 | 0.001 | -1.000 | up |
| AURKAIP1 | G051433 | -0.965 | 0.000 | -1.000 | up |
| AURKAIP1 | NBPF25P | -0.927 | 0.001 | -1.000 | up |
| AURKAIP1 | RN7SL176P | -0.910 | 0.002 | -1.000 | up |
| AURKAIP1 | G059595 | -0.925 | 0.001 | -1.000 | up |
| AURKAIP1 | RP11-255E6.6 | 0.941 | 0.000 | 1.000 | up |
| AURKAIP1 | G018162 | 0.904 | 0.002 | 1.000 | up |
| AURKAIP1 | CTD-2005H7.1 | 0.926 | 0.001 | 1.000 | up |
| AURKAIP1 | G029844 | 0.903 | 0.002 | 1.000 | up |
| AURKAIP1 | IDH1-AS1 | 0.913 | 0.002 | 1.000 | up |
| AURKAIP1 | LOC100379224 | 0.920 | 0.001 | 1.000 | up |
| AURKAIP1 | AC005220.3 | 0.912 | 0.002 | 1.000 | up |
| AURKAIP1 | G054532 | 0.935 | 0.001 | 1.000 | up |
| AURKAIP1 | AK125078 | 0.925 | 0.001 | 1.000 | up |
| AURKAIP1 | BX248253 | 0.928 | 0.001 | 1.000 | up |
| AURKAIP1 | LOC101927811 | 0.929 | 0.001 | 1.000 | up |
| AURKAIP1 | RP11-118K6.3 | 0.904 | 0.002 | 1.000 | up |
| AURKAIP1 | G005619 | 0.926 | 0.001 | 1.000 | up |
| AURKAIP1 | G015529 | 0.935 | 0.001 | 1.000 | up |
| CCDC58 | G040015 | -0.907 | 0.002 | -1.000 | up |
| CCDC58 | G024041 | -0.907 | 0.002 | -1.000 | up |
| CCDC58 | CRAT37 | -0.919 | 0.001 | -1.000 | up |
| CCDC58 | CTC-250I14.3 | -0.913 | 0.002 | -1.000 | up |
| CCDC58 | G010941 | -0.917 | 0.001 | -1.000 | up |
| CCDC58 | BC047090 | 0.924 | 0.001 | 1.000 | up |
| CCDC58 | XLOC_013660 | 0.902 | 0.002 | 1.000 | up |
| CCDC58 | ITFG1-AS1 | 0.946 | 0.000 | 1.000 | up |
| CCDC58 | RP11-820L6.1 | 0.922 | 0.001 | 1.000 | up |
| EARS2 | Z69720.2 | -0.919 | 0.001 | -1.000 | up |
| EARS2 | LOC727896 | -0.941 | 0.000 | -1.000 | up |
| EARS2 | XLOC_007868 | -0.972 | 0.000 | -1.000 | up |
| EARS2 | RP11-689C9.1 | 0.929 | 0.001 | 1.000 | up |
| EARS2 | BC047090 | 0.900 | 0.002 | 1.000 | up |
| EARS2 | G057990 | 0.958 | 0.000 | 1.000 | up |
| EARS2 | BC112270 | 0.915 | 0.001 | 1.000 | up |
| EARS2 | uc.95 | 0.938 | 0.001 | 1.000 | up |
| EARS2 | RP11-872D17.4 | 0.923 | 0.001 | 1.000 | up |
| EARS2 | XLOC_004894 | 0.929 | 0.001 | 1.000 | up |
| EARS2 | G007298 | 0.948 | 0.000 | 1.000 | up |
| MRPL12 | G009303 | -0.964 | 0.000 | -1.000 | up |
| MRPL12 | XLOC_003405 | -0.912 | 0.002 | -1.000 | up |

| | | | | | |
|---|---|---|---|---|---|
| MRPL12 | CTC-261N6.1 | -0.932 | 0.001 | -1.000 | up |
| MRPL12 | CTB-102L5.8 | -0.935 | 0.001 | -1.000 | up |
| MRPL12 | RP1-124C6.1 | -0.942 | 0.000 | -1.000 | up |
| MRPL12 | G071944 | -0.968 | 0.000 | -1.000 | up |
| MRPL12 | XLOC_003837 | -0.933 | 0.001 | -1.000 | up |
| MRPL12 | NBPF8 | -0.955 | 0.000 | -1.000 | up |
| MRPL12 | LINC00930 | -0.975 | 0.000 | -1.000 | up |
| MRPL12 | RP1-65P5.3 | -0.934 | 0.001 | -1.000 | up |
| MRPL12 | XLOC_008270 | -0.971 | 0.000 | -1.000 | up |
| MRPL12 | BCRP3 | -0.914 | 0.001 | -1.000 | up |
| MRPL12 | MIMT1 | -0.910 | 0.002 | -1.000 | up |
| MRPL12 | G077330 | 0.911 | 0.002 | 1.000 | up |
| MRPL12 | CTD-2005H7.1 | 0.915 | 0.001 | 1.000 | up |
| MRPL12 | LOC100379224 | 0.947 | 0.000 | 1.000 | up |
| MRPL12 | G006537 | 0.905 | 0.002 | 1.000 | up |
| MRPL12 | LOC101927822 | 0.933 | 0.001 | 1.000 | up |
| MRPL12 | uc.412 | 0.942 | 0.000 | 1.000 | up |
| MRPL12 | JARID2-AS1 | 0.953 | 0.000 | 1.000 | up |
| PTRH2 | G037251 | -0.911 | 0.002 | -1.000 | up |
| PTRH2 | RP1-296L11.1 | -0.900 | 0.002 | -1.000 | up |
| PTRH2 | IGBP1-AS2 | -0.905 | 0.002 | -1.000 | up |
| PTRH2 | RP11-3K16.2 | -0.934 | 0.001 | -1.000 | up |
| PTRH2 | XLOC_008025 | -0.913 | 0.002 | -1.000 | up |
| PTRH2 | RP11-737O24.3 | -0.929 | 0.001 | -1.000 | up |
| PTRH2 | G072062 | -0.921 | 0.001 | -1.000 | up |
| PTRH2 | G063659 | -0.952 | 0.000 | -1.000 | up |
| PTRH2 | G030124 | -0.901 | 0.002 | -1.000 | up |
| PTRH2 | LOC101927691 | -0.912 | 0.002 | -1.000 | up |
| PTRH2 | AK025321 | -0.900 | 0.002 | -1.000 | up |
| PTRH2 | G001795 | -0.904 | 0.002 | -1.000 | up |
| PTRH2 | XLOC_012740 | -0.903 | 0.002 | -1.000 | up |
| PTRH2 | XLOC_006485 | -0.935 | 0.001 | -1.000 | up |
| PTRH2 | G015181 | -0.901 | 0.002 | -1.000 | up |
| PTRH2 | RP11-551L14.6 | -0.927 | 0.001 | -1.000 | up |
| PTRH2 | G051586 | -0.947 | 0.000 | -1.000 | up |
| PTRH2 | DLG3-AS1 | -0.944 | 0.000 | -1.000 | up |
| PTRH2 | G010533 | -0.931 | 0.001 | -1.000 | up |
| PTRH2 | RP11-255E6.6 | 0.930 | 0.001 | 1.000 | up |
| PTRH2 | RP11-196G18.23 | 0.917 | 0.001 | 1.000 | up |
| PTRH2 | RP11-285F16.1 | 0.909 | 0.002 | 1.000 | up |
| PTRH2 | RP5-1092A3.4 | 0.914 | 0.002 | 1.000 | up |
| PTRH2 | XLOC_006554 | 0.920 | 0.001 | 1.000 | up |
| PTRH2 | G085410 | 0.943 | 0.000 | 1.000 | up |
| PTRH2 | G089005 | 0.905 | 0.002 | 1.000 | up |
| PTRH2 | G079654 | 0.925 | 0.001 | 1.000 | up |
| PTRH2 | G009018 | 0.904 | 0.002 | 1.000 | up |
| PTRH2 | G081251 | 0.909 | 0.002 | 1.000 | up |
| PTRH2 | G074033 | 0.907 | 0.002 | 1.000 | up |
| PTRH2 | XLOC_011052 | 0.918 | 0.001 | 1.000 | up |
| PTRH2 | RP11-502I4.3 | 0.902 | 0.002 | 1.000 | up |
| PTRH2 | IDH1-AS1 | 0.921 | 0.001 | 1.000 | up |
| PTRH2 | PVRL3-AS1 | 0.959 | 0.000 | 1.000 | up |
| PTRH2 | RP4-555D20.3 | 0.920 | 0.001 | 1.000 | up |
| PTRH2 | TOB1-AS1 (1) | 0.928 | 0.001 | 1.000 | up |
| PTRH2 | G066437 | 0.903 | 0.002 | 1.000 | up |
| PTRH2 | ENO1-AS1 | 0.936 | 0.001 | 1.000 | up |
| PTRH2 | LL09NC01-139C3.1 | 0.919 | 0.001 | 1.000 | up |
| PTRH2 | G061722 | 0.904 | 0.002 | 1.000 | up |
| PTRH2 | CXADRP3 | 0.902 | 0.002 | 1.000 | up |
| PTRH2 | CTD-2382E5.2 | 0.942 | 0.000 | 1.000 | up |
| PTRH2 | EZR-AS1 | 0.938 | 0.001 | 1.000 | up |
| PTRH2 | RP1-86C11.7 | 0.920 | 0.001 | 1.000 | up |
| PTRH2 | RP11-109D9.4 | 0.940 | 0.001 | 1.000 | up |
| PTRH2 | LINC00449 | 0.902 | 0.002 | 1.000 | up |
| PTRH2 | AK125078 | 0.921 | 0.001 | 1.000 | up |

| | | | | | |
|---|---|---|---|---|---|
| PTRH2 | BX248253 | 0.912 | 0.002 | 1.000 | up |
| PTRH2 | G070405 | 0.945 | 0.000 | 1.000 | up |
| PTRH2 | G054079 | 0.903 | 0.002 | 1.000 | up |
| PTRH2 | RP11-53I6.3 | 0.904 | 0.002 | 1.000 | up |
| PTRH2 | G014106 | 0.929 | 0.001 | 1.000 | up |
| PTRH2 | G002695 | 0.926 | 0.001 | 1.000 | up |
| PTRH2 | G021588 | 0.937 | 0.001 | 1.000 | up |
| PTRH2 | LOC101928489 | 0.912 | 0.002 | 1.000 | up |
| PTRH2 | G075703 | 0.952 | 0.000 | 1.000 | up |
| PTRH2 | RP11-820L6.1 | 0.951 | 0.000 | 1.000 | up |
| PTRH2 | LOC102723385 | 0.913 | 0.002 | 1.000 | up |
| PUS1 | ITGA9-AS1 | -0.913 | 0.002 | -1.000 | up |
| PUS1 | G011310 | -0.927 | 0.001 | -1.000 | up |
| PUS1 | RP11-3K16.2 | -0.911 | 0.002 | -1.000 | up |
| PUS1 | G008916 | -0.916 | 0.001 | -1.000 | up |
| PUS1 | AP001625.6 | -0.925 | 0.001 | -1.000 | up |
| PUS1 | RP11-669E14.6 | -0.969 | 0.000 | -1.000 | up |
| PUS1 | NEAT1 | -0.932 | 0.001 | -1.000 | up |
| PUS1 | uc.49 | -0.973 | 0.000 | -1.000 | up |
| PUS1 | CTD-2023M8.1 | -0.943 | 0.000 | -1.000 | up |
| PUS1 | XLOC_012740 | -0.982 | 0.000 | -1.000 | up |
| PUS1 | uc.474 | -0.906 | 0.002 | -1.000 | up |
| PUS1 | G031271 | -0.930 | 0.001 | -1.000 | up |
| PUS1 | G029241 | -0.951 | 0.000 | -1.000 | up |
| PUS1 | DLG3-AS1 | -0.928 | 0.001 | -1.000 | up |
| PUS1 | G010224 | -0.927 | 0.001 | -1.000 | up |
| PUS1 | RP11-602N24.3 | 0.944 | 0.000 | 1.000 | up |
| PUS1 | AL110181 | 0.937 | 0.001 | 1.000 | up |
| PUS1 | TIPARP-AS1 | 0.913 | 0.002 | 1.000 | up |
| PUS1 | G061591 | 0.918 | 0.001 | 1.000 | up |
| PUS1 | RP1-40E16.12 | 0.928 | 0.001 | 1.000 | up |
| PUS1 | GBAT2 | 0.909 | 0.002 | 1.000 | up |
| PUS1 | G075198 | 0.900 | 0.002 | 1.000 | up |
| PUS1 | DLG5-AS1 (1) | 0.908 | 0.002 | 1.000 | up |
| PUS1 | LINC00662 | 0.924 | 0.001 | 1.000 | up |
| PUS1 | RP11-46J23.1 | 0.902 | 0.002 | 1.000 | up |
| PUS1 | RP11-715J22.6 | 0.934 | 0.001 | 1.000 | up |
| PUS1 | AC099754.1 | 0.937 | 0.001 | 1.000 | up |
| PUS1 | G079009 | 0.903 | 0.002 | 1.000 | up |
| PUS1 | RP11-396C23.4 | 0.956 | 0.000 | 1.000 | up |
| PUS1 | MGC2889 | 0.951 | 0.000 | 1.000 | up |
| PUS1 | CTD-2005H7.1 | 0.909 | 0.002 | 1.000 | up |
| PUS1 | G042247 | 0.915 | 0.001 | 1.000 | up |
| PUS1 | AC144652.1 | 0.904 | 0.002 | 1.000 | up |
| PUS1 | G085410 | 0.949 | 0.000 | 1.000 | up |
| PUS1 | OLMALINC | 0.956 | 0.000 | 1.000 | up |
| PUS1 | MAPT-IT1 | 0.927 | 0.001 | 1.000 | up |
| PUS1 | CCNT2-AS1 | 0.968 | 0.000 | 1.000 | up |
| PUS1 | G023041 | 0.902 | 0.002 | 1.000 | up |
| PUS1 | TNRC18P1 | 0.907 | 0.002 | 1.000 | up |
| PUS1 | NIPBL-AS1 | 0.910 | 0.002 | 1.000 | up |
| PUS1 | G053697 | 0.970 | 0.000 | 1.000 | up |
| PUS1 | LOC730102 | 0.920 | 0.001 | 1.000 | up |
| PUS1 | G010782 | 0.922 | 0.001 | 1.000 | up |
| PUS1 | LINC00634 | 0.926 | 0.001 | 1.000 | up |
| PUS1 | G066437 | 0.946 | 0.000 | 1.000 | up |
| PUS1 | DGCR5 | 0.916 | 0.001 | 1.000 | up |
| PUS1 | CXADRP3 | 0.920 | 0.001 | 1.000 | up |
| PUS1 | EZR-AS1 | 0.931 | 0.001 | 1.000 | up |
| PUS1 | ANKRD20A12P | 0.904 | 0.002 | 1.000 | up |
| PUS1 | G003603 | 0.908 | 0.002 | 1.000 | up |
| PUS1 | KB-318B8.7 | 0.965 | 0.000 | 1.000 | up |
| PUS1 | G014106 | 0.989 | 0.000 | 1.000 | up |
| PUS1 | RP11-355F22.1 | 0.929 | 0.001 | 1.000 | up |
| PUS1 | RP11-166B2.3 | 0.921 | 0.001 | 1.000 | up |

| | | | | | |
|---|---|---|---|---|---|
| PUS1 | BC047644 | 0.969 | 0.000 | 1.000 | up |
| PUS1 | GLIDR | 0.948 | 0.000 | 1.000 | up |
| PUS1 | LOC101927358 | 0.922 | 0.001 | 1.000 | up |
| PUS1 | CTC-248O19.1 | 0.907 | 0.002 | 1.000 | up |
| TRAP1 | Alpha_250_dna | -0.919 | 0.001 | -1.000 | up |
| TRAP1 | G011310 | -0.921 | 0.001 | -1.000 | up |
| TRAP1 | ASAP1-IT1 | -0.960 | 0.000 | -1.000 | up |
| TRAP1 | G081487 | -0.944 | 0.000 | -1.000 | up |
| TRAP1 | RP11-669E14.6 | -0.909 | 0.002 | -1.000 | up |
| TRAP1 | CTC-261N6.1 | -0.920 | 0.001 | -1.000 | up |
| TRAP1 | XLOC_003837 | -0.901 | 0.002 | -1.000 | up |
| TRAP1 | G082713 | -0.912 | 0.002 | -1.000 | up |
| TRAP1 | CTD-3193O13.1 | -0.904 | 0.002 | -1.000 | up |
| TRAP1 | G089283 | -0.921 | 0.001 | -1.000 | up |
| TRAP1 | XLOC_009429 | -0.901 | 0.002 | -1.000 | up |
| TRAP1 | G059595 | -0.947 | 0.000 | -1.000 | up |
| TRAP1 | IFT74-AS1 | 0.928 | 0.001 | 1.000 | up |
| TRAP1 | SH3GL1P2 | 0.962 | 0.000 | 1.000 | up |
| TRAP1 | G064655 | 0.932 | 0.001 | 1.000 | up |
| TRAP1 | RP11-715J22.6 | 0.913 | 0.002 | 1.000 | up |
| TRAP1 | RP5-991G20.4 | 0.974 | 0.000 | 1.000 | up |
| TRAP1 | MGC2889 | 0.952 | 0.000 | 1.000 | up |
| TRAP1 | CCNT2-AS1 | 0.929 | 0.001 | 1.000 | up |
| TRAP1 | RP11-775D22.3 | 0.931 | 0.001 | 1.000 | up |
| TRAP1 | G078120 | 0.918 | 0.001 | 1.000 | up |
| TRAP1 | LINC00463 | 0.907 | 0.002 | 1.000 | up |
| TRAP1 | G068576 | 0.908 | 0.002 | 1.000 | up |
| TRAP1 | RP11-110G21.1 | 0.925 | 0.001 | 1.000 | up |
| TRAP1 | RP11-314A20.5 | 0.958 | 0.000 | 1.000 | up |
| TRAP1 | LOC101927811 | 0.904 | 0.002 | 1.000 | up |
| TRAP1 | LOC283140 | 0.918 | 0.001 | 1.000 | up |
| PGS1 | LINC00473 | -0.944 | 0.000 | -1.000 | up |
| PGS1 | G008916 | -0.929 | 0.001 | -1.000 | up |
| PGS1 | GSE61474_XLOC_020343 | -0.908 | 0.002 | -1.000 | up |
| PGS1 | RP11-669E14.6 | -0.912 | 0.002 | -1.000 | up |
| PGS1 | uc.49 | -0.927 | 0.001 | -1.000 | up |
| PGS1 | CTD-2023M8.1 | -0.920 | 0.001 | -1.000 | up |
| PGS1 | G030124 | -0.934 | 0.001 | -1.000 | up |
| PGS1 | XLOC_012740 | -0.918 | 0.001 | -1.000 | up |
| PGS1 | AC025442.3 | -0.950 | 0.000 | -1.000 | up |
| PGS1 | G013465 | -0.923 | 0.001 | -1.000 | up |
| PGS1 | G029241 | -0.963 | 0.000 | -1.000 | up |
| PGS1 | G088912 | -0.932 | 0.001 | -1.000 | up |
| PGS1 | G010224 | -0.937 | 0.001 | -1.000 | up |
| PGS1 | AL110181 | 0.906 | 0.002 | 1.000 | up |
| PGS1 | GBAT2 | 0.903 | 0.002 | 1.000 | up |
| PGS1 | RP11-715J22.6 | 0.906 | 0.002 | 1.000 | up |
| PGS1 | G079009 | 0.912 | 0.002 | 1.000 | up |
| PGS1 | RP11-396C23.4 | 0.914 | 0.001 | 1.000 | up |
| PGS1 | G014329 | 0.922 | 0.001 | 1.000 | up |
| PGS1 | G066319 | 0.920 | 0.001 | 1.000 | up |
| PGS1 | RP11-475I24.3 | 0.960 | 0.000 | 1.000 | up |
| PGS1 | AC144652.1 | 0.900 | 0.002 | 1.000 | up |
| PGS1 | G085410 | 0.914 | 0.001 | 1.000 | up |
| PGS1 | OLMALINC | 0.919 | 0.001 | 1.000 | up |
| PGS1 | LINC00408 | 0.952 | 0.000 | 1.000 | up |
| PGS1 | G053697 | 0.975 | 0.000 | 1.000 | up |
| PGS1 | G010782 | 0.937 | 0.001 | 1.000 | up |
| PGS1 | G031140 | 0.902 | 0.002 | 1.000 | up |
| PGS1 | LL22NC03-86G7.1 | 0.925 | 0.001 | 1.000 | up |
| PGS1 | DGCR5 | 0.904 | 0.002 | 1.000 | up |
| PGS1 | RP11-798M19.6 | 0.941 | 0.000 | 1.000 | up |
| PGS1 | ANKRD20A12P | 0.928 | 0.001 | 1.000 | up |
| PGS1 | KB-318B8.7 | 0.943 | 0.000 | 1.000 | up |

| | | | | | |
|---|---|---|---|---|---|
| PGS1 | G014106 | 0.958 | 0.000 | 1.000 | up |
| PGS1 | RP11-355F22.1 | 0.926 | 0.001 | 1.000 | up |
| PGS1 | BC047644 | 0.908 | 0.002 | 1.000 | up |
| PGS1 | GLIDR | 0.952 | 0.000 | 1.000 | up |
| SLC25A37 | PPIEL | -0.951 | 0.000 | -1.000 | up |
| SLC25A37 | G046678 | -0.905 | 0.002 | -1.000 | up |
| SLC25A37 | G050702 | -0.906 | 0.002 | -1.000 | up |
| SLC25A37 | ZFAT-AS1 | -0.963 | 0.000 | -1.000 | up |
| SLC25A37 | AK093193 | -0.937 | 0.001 | -1.000 | up |
| SLC25A37 | CTD-2050E21.1 | -0.933 | 0.001 | -1.000 | up |
| SLC25A37 | BX648501 | -0.903 | 0.002 | -1.000 | up |
| SLC25A37 | TMLHE-AS1 | -0.905 | 0.002 | -1.000 | up |
| SLC25A37 | G058409 | -0.947 | 0.000 | -1.000 | up |
| SLC25A37 | G009329 | -0.903 | 0.002 | -1.000 | up |
| SLC25A37 | LINC00685 | -0.901 | 0.002 | -1.000 | up |
| SLC25A37 | G010088 | -0.944 | 0.000 | -1.000 | up |
| SLC25A37 | RP11-651L5.2 | -0.918 | 0.001 | -1.000 | up |
| SLC25A37 | G059595 | -0.920 | 0.001 | -1.000 | up |
| SLC25A37 | RP11-255E6.6 | 0.929 | 0.001 | 1.000 | up |
| SLC25A37 | RP11-744I24.3 | 0.907 | 0.002 | 1.000 | up |
| SLC25A37 | LOC100130417 | 0.902 | 0.002 | 1.000 | up |
| SLC25A37 | USP27X-AS1 | 0.933 | 0.001 | 1.000 | up |
| SLC25A37 | G070136 | 0.920 | 0.001 | 1.000 | up |
| SLC25A37 | G002862 | 0.914 | 0.001 | 1.000 | up |
| SLC25A37 | CCDC144NL-AS1 | 0.944 | 0.000 | 1.000 | up |
| SLC25A37 | G050681 | 0.929 | 0.001 | 1.000 | up |
| SLC25A37 | G079888 | 0.921 | 0.001 | 1.000 | up |
| SLC25A37 | XLOC_006928 | 0.942 | 0.000 | 1.000 | up |
| SLC25A37 | RP11-299H21.1 | 0.923 | 0.001 | 1.000 | up |
| SLC25A37 | RP13-131K19.2 | 0.958 | 0.000 | 1.000 | up |
| SLC25A37 | FOXN3-AS1 | 0.934 | 0.001 | 1.000 | up |
| SLC25A37 | INAFM1 | 0.901 | 0.002 | 1.000 | up |
| SLC25A37 | RP11-775D22.3 | 0.901 | 0.002 | 1.000 | up |
| SLC25A37 | LOC148709 | 0.928 | 0.001 | 1.000 | up |
| SLC25A37 | GYG2P1 | 0.905 | 0.002 | 1.000 | up |
| SLC25A37 | PRR36 | 0.939 | 0.001 | 1.000 | up |
| SLC25A37 | RP11-1275H24.1 | 0.908 | 0.002 | 1.000 | up |
| SLC25A37 | CRYBB2P1 | 0.947 | 0.000 | 1.000 | up |
| SLC25A37 | LOC101927811 | 0.909 | 0.002 | 1.000 | up |
| SLC25A37 | G063514 | 0.912 | 0.002 | 1.000 | up |
| SLC25A37 | JARID2-AS1 | 0.923 | 0.001 | 1.000 | up |
| SLC25A37 | LOC283140 | 0.903 | 0.002 | 1.000 | up |
| SLC25A37 | G015529 | 0.950 | 0.000 | 1.000 | up |
| ATAD3A | G029241 | -0.975 | 0.000 | -1.000 | up |
| ATAD3A | G010224 | -0.938 | 0.001 | -1.000 | up |
| ATAD3A | RP11-602N24.3 | 0.931 | 0.001 | 1.000 | up |
| ATAD3A | DLG5-AS1 (1) | 0.902 | 0.002 | 1.000 | up |
| ATAD3A | AC138655.6 | 0.903 | 0.002 | 1.000 | up |
| ATAD3A | RP11-475I24.3 | 0.962 | 0.000 | 1.000 | up |
| ATAD3A | AC144652.1 | 0.908 | 0.002 | 1.000 | up |
| ATAD3A | G087757 | 0.902 | 0.002 | 1.000 | up |
| ATAD3A | G053697 | 0.975 | 0.000 | 1.000 | up |
| ATAD3A | G010782 | 0.947 | 0.000 | 1.000 | up |
| ATAD3A | LL22NC03-86G7.1 | 0.942 | 0.000 | 1.000 | up |
| ATAD3A | FAM66D | 0.908 | 0.002 | 1.000 | up |
| ATAD3A | ANKRD20A12P | 0.910 | 0.002 | 1.000 | up |
| ATAD3A | KB-318B8.7 | 0.974 | 0.000 | 1.000 | up |
| ATAD3A | RP11-355F22.1 | 0.919 | 0.001 | 1.000 | up |
| ATAD3A | GLIDR | 0.929 | 0.001 | 1.000 | up |
| ATAD3B | SLC7A11-AS1 | -0.912 | 0.002 | -1.000 | up |
| ATAD3B | RP11-70P17.1 | -0.903 | 0.002 | -1.000 | up |
| ATAD3B | G011310 | -0.947 | 0.000 | -1.000 | up |
| ATAD3B | AP000438.2 | -0.901 | 0.002 | -1.000 | up |
| ATAD3B | RP11-466A19.1 | -0.909 | 0.002 | -1.000 | up |
| ATAD3B | uc.49 | -0.934 | 0.001 | -1.000 | up |

| | | | | | |
|---|---|---|---|---|---|
| ATAD3B | G073689 | -0.907 | 0.002 | -1.000 | up |
| ATAD3B | EML2-AS1 | -0.914 | 0.001 | -1.000 | up |
| ATAD3B | G003431 | -0.920 | 0.001 | -1.000 | up |
| ATAD3B | XLOC_012740 | -0.958 | 0.000 | -1.000 | up |
| ATAD3B | G013465 | -0.970 | 0.000 | -1.000 | up |
| ATAD3B | RP5-1112D6.7 | -0.930 | 0.001 | -1.000 | up |
| ATAD3B | G029241 | -0.969 | 0.000 | -1.000 | up |
| ATAD3B | EF565122 | -0.912 | 0.002 | -1.000 | up |
| ATAD3B | RP11-461K13.1 | -0.934 | 0.001 | -1.000 | up |
| ATAD3B | RP11-602N24.3 | 0.954 | 0.000 | 1.000 | up |
| ATAD3B | AL110181 | 0.980 | 0.000 | 1.000 | up |
| ATAD3B | TIPARP-AS1 | 0.905 | 0.002 | 1.000 | up |
| ATAD3B | RP11-363E6.3 | 0.917 | 0.001 | 1.000 | up |
| ATAD3B | AK055712 | 0.925 | 0.001 | 1.000 | up |
| ATAD3B | RP11-303E16.7 | 0.926 | 0.001 | 1.000 | up |
| ATAD3B | CTD-2023N9.3 | 0.932 | 0.001 | 1.000 | up |
| ATAD3B | RP11-686O6.2 | 0.941 | 0.000 | 1.000 | up |
| ATAD3B | G061591 | 0.950 | 0.000 | 1.000 | up |
| ATAD3B | RP11-356J5.12 | 0.968 | 0.000 | 1.000 | up |
| ATAD3B | RP1-40E16.12 | 0.972 | 0.000 | 1.000 | up |
| ATAD3B | GBAT2 | 0.955 | 0.000 | 1.000 | up |
| ATAD3B | AL137474 | 0.937 | 0.001 | 1.000 | up |
| ATAD3B | AC138655.6 | 0.912 | 0.002 | 1.000 | up |
| ATAD3B | LINC00662 | 0.941 | 0.000 | 1.000 | up |
| ATAD3B | RP11-297J22.1 | 0.917 | 0.001 | 1.000 | up |
| ATAD3B | RP11-179B2.2 | 0.915 | 0.001 | 1.000 | up |
| ATAD3B | AC099754.1 | 0.937 | 0.001 | 1.000 | up |
| ATAD3B | MGC2889 | 0.924 | 0.001 | 1.000 | up |
| ATAD3B | RP4-683L5.1 | 0.901 | 0.002 | 1.000 | up |
| ATAD3B | AC144652.1 | 0.979 | 0.000 | 1.000 | up |
| ATAD3B | G085410 | 0.940 | 0.001 | 1.000 | up |
| ATAD3B | G089005 | 0.903 | 0.002 | 1.000 | up |
| ATAD3B | OLMALINC | 0.907 | 0.002 | 1.000 | up |
| ATAD3B | LINC00408 | 0.909 | 0.002 | 1.000 | up |
| ATAD3B | MAPT-IT1 | 0.946 | 0.000 | 1.000 | up |
| ATAD3B | G073576 | 0.910 | 0.002 | 1.000 | up |
| ATAD3B | XLOC_013222 | 0.938 | 0.001 | 1.000 | up |
| ATAD3B | G082866 | 0.905 | 0.002 | 1.000 | up |
| ATAD3B | TNRC18P1 | 0.914 | 0.001 | 1.000 | up |
| ATAD3B | RP11-138E2.1 | 0.923 | 0.001 | 1.000 | up |
| ATAD3B | G053697 | 0.963 | 0.000 | 1.000 | up |
| ATAD3B | LOC730102 | 0.908 | 0.002 | 1.000 | up |
| ATAD3B | G059716 | 0.930 | 0.001 | 1.000 | up |
| ATAD3B | BC042589 | 0.922 | 0.001 | 1.000 | up |
| ATAD3B | LINC00634 | 0.978 | 0.000 | 1.000 | up |
| ATAD3B | RP11-22N19.2 | 0.902 | 0.002 | 1.000 | up |
| ATAD3B | LL22NC03-86G7.1 | 0.948 | 0.000 | 1.000 | up |
| ATAD3B | RP1-223E5.4 | 0.946 | 0.000 | 1.000 | up |
| ATAD3B | FAM66D | 0.933 | 0.001 | 1.000 | up |
| ATAD3B | LL09NC01-139C3.1 | 0.920 | 0.001 | 1.000 | up |
| ATAD3B | RP11-798M19.6 | 0.911 | 0.002 | 1.000 | up |
| ATAD3B | ANKRD20A12P | 0.936 | 0.001 | 1.000 | up |
| ATAD3B | G023340 | 0.950 | 0.000 | 1.000 | up |
| ATAD3B | KB-318B8.7 | 0.974 | 0.000 | 1.000 | up |
| ATAD3B | G014106 | 0.934 | 0.001 | 1.000 | up |
| ATAD3B | G006578 | 0.934 | 0.001 | 1.000 | up |
| ATAD3B | RP11-355F22.1 | 0.922 | 0.001 | 1.000 | up |
| ATAD3B | G075319 | 0.943 | 0.000 | 1.000 | up |
| ATAD3B | RP11-166B2.3 | 0.951 | 0.000 | 1.000 | up |
| ATAD3B | BC047644 | 0.949 | 0.000 | 1.000 | up |
| ATAD3B | GLIDR | 0.919 | 0.001 | 1.000 | up |
| ATAD3B | RP11-439C15.4 | 0.932 | 0.001 | 1.000 | up |
| ATAD3B | LOC101927358 | 0.901 | 0.002 | 1.000 | up |
| MT-ND1 | G003271 | -0.916 | 0.001 | -1.000 | up |
| MT-ND1 | HCG18 | -0.961 | 0.000 | -1.000 | up |

| | | | | | |
|---|---|---|---|---|---|
| MT-ND1 | G087265 | -0.942 | 0.000 | -1.000 | up |
| MT-ND1 | LOC101927164 | -0.905 | 0.002 | -1.000 | up |
| MT-ND1 | RP4-760C5.3 | -0.933 | 0.001 | -1.000 | up |
| MT-ND1 | G068890 | -0.965 | 0.000 | -1.000 | up |
| MT-ND1 | AK022848 | -0.918 | 0.001 | -1.000 | up |
| MT-ND1 | FAM74A4 | 0.952 | 0.000 | 1.000 | up |
| MT-ND1 | RP11-872D17.4 | 0.935 | 0.001 | 1.000 | up |
| MT-ND1 | G081642 | 0.950 | 0.000 | 1.000 | up |
| MT-ND1 | CTC-350I8.1 | 0.932 | 0.001 | 1.000 | up |
| MT-ND1 | XLOC_004894 | 0.964 | 0.000 | 1.000 | up |
| HSPA1B | LOC102723439 | -0.955 | 0.000 | -1.000 | up |
| HSPA1B | ASAP1-IT1 | -0.910 | 0.002 | -1.000 | up |
| HSPA1B | CHKB-CPT1B | -0.925 | 0.001 | -1.000 | up |
| HSPA1B | G046678 | -0.917 | 0.001 | -1.000 | up |
| HSPA1B | G081487 | -0.958 | 0.000 | -1.000 | up |
| HSPA1B | G081302 | -0.934 | 0.001 | -1.000 | up |
| HSPA1B | G065302 | -0.958 | 0.000 | -1.000 | up |
| HSPA1B | TMLHE-AS1 | -0.959 | 0.000 | -1.000 | up |
| HSPA1B | G058409 | -0.950 | 0.000 | -1.000 | up |
| HSPA1B | RP11-126O1.4 | -0.956 | 0.000 | -1.000 | up |
| HSPA1B | G009199 | -0.900 | 0.002 | -1.000 | up |
| HSPA1B | G010533 | -0.910 | 0.002 | -1.000 | up |
| HSPA1B | XLOC_009429 | -0.967 | 0.000 | -1.000 | up |
| HSPA1B | G059595 | -0.937 | 0.001 | -1.000 | up |
| HSPA1B | IFT74-AS1 | 0.930 | 0.001 | 1.000 | up |
| HSPA1B | G021652 | 0.929 | 0.001 | 1.000 | up |
| HSPA1B | SH3GL1P2 | 0.957 | 0.000 | 1.000 | up |
| HSPA1B | TM4SF1-AS1 | 0.939 | 0.001 | 1.000 | up |
| HSPA1B | G064655 | 0.959 | 0.000 | 1.000 | up |
| HSPA1B | RP11-252A24.3 | 0.944 | 0.000 | 1.000 | up |
| HSPA1B | CTC-232P5.3 | 0.911 | 0.002 | 1.000 | up |
| HSPA1B | RP5-991G20.4 | 0.952 | 0.000 | 1.000 | up |
| HSPA1B | MGC2889 | 0.914 | 0.001 | 1.000 | up |
| HSPA1B | RP11-299H21.1 | 0.914 | 0.002 | 1.000 | up |
| HSPA1B | CCNT2-AS1 | 0.902 | 0.002 | 1.000 | up |
| HSPA1B | RP11-775D22.3 | 0.985 | 0.000 | 1.000 | up |
| HSPA1B | G066606 | 0.934 | 0.001 | 1.000 | up |
| HSPA1B | AC005220.3 | 0.929 | 0.001 | 1.000 | up |
| HSPA1B | GPR50-AS1 | 0.934 | 0.001 | 1.000 | up |
| HSPA1B | G078120 | 0.929 | 0.001 | 1.000 | up |
| HSPA1B | G066437 | 0.915 | 0.001 | 1.000 | up |
| HSPA1B | CXADRP3 | 0.949 | 0.000 | 1.000 | up |
| HSPA1B | RP11-110G21.1 | 0.902 | 0.002 | 1.000 | up |
| HSPA1B | G054532 | 0.955 | 0.000 | 1.000 | up |
| HSPA1B | CTC-359M8.1 | 0.945 | 0.000 | 1.000 | up |
| HSPA1B | RP11-314A20.5 | 0.926 | 0.001 | 1.000 | up |
| HSPA1B | CRYBB2P1 | 0.902 | 0.002 | 1.000 | up |
| HSPA1B | LOC101927811 | 0.968 | 0.000 | 1.000 | up |
| HSPA1B | G064225 | 0.915 | 0.001 | 1.000 | up |
| HSPA1B | LOC283140 | 0.979 | 0.000 | 1.000 | up |
| HSPA1B | G057611 | 0.942 | 0.000 | 1.000 | up |
| HSPA1B | LOC102723385 | 0.921 | 0.001 | 1.000 | up |
| HSPA1B | G063721 | 0.902 | 0.002 | 1.000 | up |
| PPRC1 | ASAP1-IT1 | -0.926 | 0.001 | -1.000 | up |
| PPRC1 | G081487 | -0.914 | 0.001 | -1.000 | up |
| PPRC1 | XLOC_003405 | -0.903 | 0.002 | -1.000 | up |
| PPRC1 | RP11-669E14.6 | -0.924 | 0.001 | -1.000 | up |
| PPRC1 | CTC-261N6.1 | -0.978 | 0.000 | -1.000 | up |
| PPRC1 | G071640 | -0.925 | 0.001 | -1.000 | up |
| PPRC1 | G047392 | -0.908 | 0.002 | -1.000 | up |
| PPRC1 | RP1-124C6.1 | -0.916 | 0.001 | -1.000 | up |
| PPRC1 | XLOC_003837 | -0.951 | 0.000 | -1.000 | up |
| PPRC1 | LINC00930 | -0.904 | 0.002 | -1.000 | up |
| PPRC1 | G053358 | -0.903 | 0.002 | -1.000 | up |
| PPRC1 | G089283 | -0.932 | 0.001 | -1.000 | up |

| | | | | | |
|---|---|---|---|---|---|
| PPRC1 | MIMT1 | -0.923 | 0.001 | -1.000 | up |
| PPRC1 | G074298 | -0.959 | 0.000 | -1.000 | up |
| PPRC1 | G059595 | -0.927 | 0.001 | -1.000 | up |
| PPRC1 | RPL23AP7 | 0.928 | 0.001 | 1.000 | up |
| PPRC1 | DLG5-AS1 (1) | 0.918 | 0.001 | 1.000 | up |
| PPRC1 | SH3GL1P2 | 0.939 | 0.001 | 1.000 | up |
| PPRC1 | G064655 | 0.923 | 0.001 | 1.000 | up |
| PPRC1 | RP11-715J22.6 | 0.957 | 0.000 | 1.000 | up |
| PPRC1 | AC099754.1 | 0.904 | 0.002 | 1.000 | up |
| PPRC1 | RP11-396C23.4 | 0.924 | 0.001 | 1.000 | up |
| PPRC1 | RP5-991G20.4 | 0.958 | 0.000 | 1.000 | up |
| PPRC1 | MGC2889 | 0.905 | 0.002 | 1.000 | up |
| PPRC1 | G082153 | 0.939 | 0.001 | 1.000 | up |
| PPRC1 | CTD-2005H7.1 | 0.918 | 0.001 | 1.000 | up |
| PPRC1 | G079888 | 0.936 | 0.001 | 1.000 | up |
| PPRC1 | G052526 | 0.914 | 0.001 | 1.000 | up |
| PPRC1 | G041225 | 0.931 | 0.001 | 1.000 | up |
| PPRC1 | CCNT2-AS1 | 0.961 | 0.000 | 1.000 | up |
| PPRC1 | XLOC_004632 | 0.967 | 0.000 | 1.000 | up |
| PPRC1 | BX248253 | 0.907 | 0.002 | 1.000 | up |
| PPRC1 | LOC101927822 | 0.907 | 0.002 | 1.000 | up |
| PPRC1 | G003603 | 0.924 | 0.001 | 1.000 | up |
| PPRC1 | RP11-314A20.5 | 0.914 | 0.002 | 1.000 | up |
| PPRC1 | uc.412 | 0.934 | 0.001 | 1.000 | up |
| PPRC1 | RP11-355F22.1 | 0.909 | 0.002 | 1.000 | up |
| PPRC1 | JARID2-AS1 | 0.902 | 0.002 | 1.000 | up |
| PPRC1 | CTB-181F24.1 | 0.912 | 0.002 | 1.000 | up |
| PPRC1 | LOC102723385 | 0.939 | 0.001 | 1.000 | up |
| PPRC1 | G063721 | 0.929 | 0.001 | 1.000 | up |
| ABCF2 | LACTB2-AS1 | -0.914 | 0.001 | -1.000 | up |
| ABCF2 | G033096 | -0.915 | 0.001 | -1.000 | up |
| ABCF2 | RP11-602N24.3 | 0.910 | 0.002 | 1.000 | up |
| ABCF2 | RP11-49K24.8 | 0.932 | 0.001 | 1.000 | up |
| ABCF2 | GSE61474_XLOC_042356 | 0.914 | 0.001 | 1.000 | up |
| ABCF2 | G018241 | 0.918 | 0.001 | 1.000 | up |
| ABCF2 | PLBD1-AS1 | 0.909 | 0.002 | 1.000 | up |
| ABCF2 | DLG5-AS1 (1) | 0.928 | 0.001 | 1.000 | up |
| ABCF2 | AC138655.6 | 0.915 | 0.001 | 1.000 | up |
| ABCF2 | G079009 | 0.926 | 0.001 | 1.000 | up |
| ABCF2 | G014329 | 0.925 | 0.001 | 1.000 | up |
| ABCF2 | G010782 | 0.926 | 0.001 | 1.000 | up |
| ABCF2 | G036604 | 0.922 | 0.001 | 1.000 | up |
| ABCF2 | CTC-248O19.1 | 0.944 | 0.000 | 1.000 | up |
| IFT74-AS1 | PMPCA | 0.917 | 0.001 | 1.000 | down |
| IFT74-AS1 | TRAP1 | 0.928 | 0.001 | 1.000 | down |
| IFT74-AS1 | HSPA1B | 0.930 | 0.001 | 1.000 | down |
| IFT74-AS1 | OGG1 | 0.912 | 0.002 | 1.000 | down |
| FAM74A4 | MT-ND1 | 0.952 | 0.000 | 1.000 | down |
| FAM74A4 | NBR1 | 0.951 | 0.000 | 1.000 | down |
| FAM74A4 | FOXO3 | 0.949 | 0.000 | 1.000 | down |
| RP11-602N24.3 | PUS1 | 0.944 | 0.000 | 1.000 | down |
| RP11-602N24.3 | ATAD3A | 0.931 | 0.001 | 1.000 | down |
| RP11-602N24.3 | ATAD3B | 0.954 | 0.000 | 1.000 | down |
| RP11-602N24.3 | ABCF2 | 0.910 | 0.002 | 1.000 | down |
| RP11-602N24.3 | PNPLA8 | 0.902 | 0.002 | 1.000 | down |
| AL110181 | PMPCA | 0.941 | 0.000 | 1.000 | down |
| AL110181 | PUS1 | 0.937 | 0.001 | 1.000 | down |
| AL110181 | PGS1 | 0.906 | 0.002 | 1.000 | down |
| AL110181 | ATAD3B | 0.980 | 0.000 | 1.000 | down |
| AL110181 | LYRM7 | 0.936 | 0.001 | 1.000 | down |
| AL110181 | PNPLA8 | 0.982 | 0.000 | 1.000 | down |
| AL110181 | OGG1 | 0.919 | 0.001 | 1.000 | down |
| TIPARP-AS1 | PUS1 | 0.913 | 0.002 | 1.000 | down |
| TIPARP-AS1 | ATAD3B | 0.905 | 0.002 | 1.000 | down |

| | | | | | |
|---|---|---|---|---|---|
| TIPARP-AS1 | OGG1 | 0.924 | 0.001 | 1.000 | down |
| RP11-255E6.6 | PMPCA | 0.923 | 0.001 | 1.000 | down |
| RP11-255E6.6 | AURKAIP1 | 0.941 | 0.000 | 1.000 | down |
| RP11-255E6.6 | PTRH2 | 0.930 | 0.001 | 1.000 | down |
| RP11-255E6.6 | SLC25A37 | 0.929 | 0.001 | 1.000 | down |
| RP11-255E6.6 | NBR1 | 0.928 | 0.001 | 1.000 | down |
| RP11-255E6.6 | PDK1 | 0.922 | 0.001 | 1.000 | down |
| RP11-744I24.3 | SLC25A37 | 0.907 | 0.002 | 1.000 | down |
| RP11-744I24.3 | WARS2 | 0.936 | 0.001 | 1.000 | down |
| RP11-744I24.3 | NBR1 | 0.952 | 0.000 | 1.000 | down |
| XLOC_006779 | PKM | 0.918 | 0.001 | 1.000 | down |
| XLOC_006779 | DNAJC19 | 0.945 | 0.000 | 1.000 | down |
| GSE61474_XLOC_054139 | HSPE1 | 0.926 | 0.001 | 1.000 | down |
| GSE61474_XLOC_054139 | WARS2 | 0.912 | 0.002 | 1.000 | down |
| GSE61474_XLOC_054139 | NBR1 | 0.966 | 0.000 | 1.000 | down |
| GSE61474_XLOC_054139 | FOXO3 | 0.930 | 0.001 | 1.000 | down |
| RP11-363E6.3 | ATAD3B | 0.917 | 0.001 | 1.000 | down |
| RP11-363E6.3 | LYRM7 | 0.969 | 0.000 | 1.000 | down |
| RP11-363E6.3 | PNPLA8 | 0.952 | 0.000 | 1.000 | down |
| AK055712 | ATAD3B | 0.925 | 0.001 | 1.000 | down |
| AK055712 | LYRM7 | 0.959 | 0.000 | 1.000 | down |
| AK055712 | PNPLA8 | 0.926 | 0.001 | 1.000 | down |
| RP11-49K24.8 | ABCF2 | 0.932 | 0.001 | 1.000 | down |
| LOC100130417 | SLC25A37 | 0.902 | 0.002 | 1.000 | down |
| LOC100130417 | DNAJC19 | 0.930 | 0.001 | 1.000 | down |
| LOC100130417 | NBR1 | 0.944 | 0.000 | 1.000 | down |
| RPL23AP7 | PPRC1 | 0.928 | 0.001 | 1.000 | down |
| RPL23AP7 | PDK1 | 0.916 | 0.001 | 1.000 | down |
| RP11-44N22.3 | NDUFAF4 | 0.902 | 0.002 | 1.000 | down |
| RP11-44N22.3 | CKMT1B | 0.929 | 0.001 | 1.000 | down |
| CTD-2302E22.4 | IBA57 | 0.914 | 0.001 | 1.000 | down |
| CTD-2302E22.4 | OGG1 | 0.901 | 0.002 | 1.000 | down |
| PSMG3-AS1 (1) | PKM | 0.909 | 0.002 | 1.000 | down |
| PSMG3-AS1 (1) | DNAJC19 | 0.941 | 0.000 | 1.000 | down |
| GS1-433O24.1 | HSPE1 | 0.923 | 0.001 | 1.000 | down |
| GSE61474_XLOC_042356 | ABCF2 | 0.914 | 0.001 | 1.000 | down |
| RP11-196G18.23 | PTRH2 | 0.917 | 0.001 | 1.000 | down |
| RP11-196G18.23 | CKMT1A | 0.911 | 0.002 | 1.000 | down |
| LINC00652 | PMPCA | 0.918 | 0.001 | 1.000 | down |
| LINC00652 | OGG1 | 0.985 | 0.000 | 1.000 | down |
| RP11-525A16.4 | NBR1 | 0.924 | 0.001 | 1.000 | down |
| G017718 | HSPE1 | 0.901 | 0.002 | 1.000 | down |
| RP11-689C9.1 | EARS2 | 0.929 | 0.001 | 1.000 | down |
| RP11-303E16.7 | ATAD3B | 0.926 | 0.001 | 1.000 | down |
| RP11-303E16.7 | LYRM7 | 0.911 | 0.002 | 1.000 | down |
| RP11-303E16.7 | PNPLA8 | 0.976 | 0.000 | 1.000 | down |
| XLOC_008005 | NBR1 | 0.916 | 0.001 | 1.000 | down |
| XLOC_008005 | CMPK2 | 0.948 | 0.000 | 1.000 | down |
| XLOC_008005 | FOXO3 | 0.914 | 0.001 | 1.000 | down |
| CTD-2023N9.3 | ATAD3B | 0.932 | 0.001 | 1.000 | down |
| RP11-350N15.5 | IBA57 | 0.936 | 0.001 | 1.000 | down |
| RP11-762I7.4 | NDUFAF7 | 0.907 | 0.002 | 1.000 | down |
| RP4-549L20.3 | LYRM7 | 0.966 | 0.000 | 1.000 | down |
| RP4-549L20.3 | PNPLA8 | 0.934 | 0.001 | 1.000 | down |
| RP4-549L20.3 | IBA57 | 0.915 | 0.001 | 1.000 | down |
| RP11-686O6.2 | ATAD3B | 0.941 | 0.000 | 1.000 | down |
| RP11-686O6.2 | LYRM7 | 0.950 | 0.000 | 1.000 | down |
| RP11-686O6.2 | PNPLA8 | 0.979 | 0.000 | 1.000 | down |
| G061591 | PUS1 | 0.918 | 0.001 | 1.000 | down |
| G061591 | ATAD3B | 0.950 | 0.000 | 1.000 | down |

| | | | | | |
|---|---|---|---|---|---|
| G061591 | PNPLA8 | 0.922 | 0.001 | 1.000 | down |
| RP11-356J5.12 | ATAD3B | 0.968 | 0.000 | 1.000 | down |
| RP11-356J5.12 | LYRM7 | 0.919 | 0.001 | 1.000 | down |
| RP11-356J5.12 | PNPLA8 | 0.971 | 0.000 | 1.000 | down |
| USP27X-AS1 | SLC25A37 | 0.933 | 0.001 | 1.000 | down |
| USP27X-AS1 | PKM | 0.922 | 0.001 | 1.000 | down |
| USP27X-AS1 | DNAJC19 | 0.934 | 0.001 | 1.000 | down |
| USP27X-AS1 | WARS2 | 0.942 | 0.000 | 1.000 | down |
| USP27X-AS1 | NBR1 | 0.955 | 0.000 | 1.000 | down |
| G018241 | ABCF2 | 0.918 | 0.001 | 1.000 | down |
| G070136 | SLC25A37 | 0.920 | 0.001 | 1.000 | down |
| G070136 | DNAJC19 | 0.951 | 0.000 | 1.000 | down |
| RP1-40E16.12 | PMPCA | 0.919 | 0.001 | 1.000 | down |
| RP1-40E16.12 | PUS1 | 0.928 | 0.001 | 1.000 | down |
| RP1-40E16.12 | ATAD3B | 0.972 | 0.000 | 1.000 | down |
| RP1-40E16.12 | LYRM7 | 0.918 | 0.001 | 1.000 | down |
| RP1-40E16.12 | PNPLA8 | 0.965 | 0.000 | 1.000 | down |
| RP1-40E16.12 | OGG1 | 0.924 | 0.001 | 1.000 | down |
| BRWD1-IT2 | IBA57 | 0.966 | 0.000 | 1.000 | down |
| BRWD1-IT2 | NBR1 | 0.936 | 0.001 | 1.000 | down |
| G021652 | HSPA1B | 0.929 | 0.001 | 1.000 | down |
| GBAT2 | PUS1 | 0.909 | 0.002 | 1.000 | down |
| GBAT2 | PGS1 | 0.903 | 0.002 | 1.000 | down |
| GBAT2 | ATAD3B | 0.955 | 0.000 | 1.000 | down |
| GBAT2 | PNPLA8 | 0.948 | 0.000 | 1.000 | down |
| GBAT2 | OGG1 | 0.950 | 0.000 | 1.000 | down |
| RP11-490O6.2 | LYRM7 | 0.913 | 0.002 | 1.000 | down |
| RP11-490O6.2 | IBA57 | 0.969 | 0.000 | 1.000 | down |
| RP11-490O6.2 | NBR1 | 0.931 | 0.001 | 1.000 | down |
| AL137474 | ATAD3B | 0.937 | 0.001 | 1.000 | down |
| AL137474 | LYRM7 | 0.933 | 0.001 | 1.000 | down |
| AL137474 | PNPLA8 | 0.936 | 0.001 | 1.000 | down |
| BC047090 | NDUFAF4 | 0.911 | 0.002 | 1.000 | down |
| BC047090 | CCDC58 | 0.924 | 0.001 | 1.000 | down |
| BC047090 | EARS2 | 0.900 | 0.002 | 1.000 | down |
| RP11-1017G21.6 | IBA57 | 0.965 | 0.000 | 1.000 | down |
| PLBD1-AS1 | ABCF2 | 0.909 | 0.002 | 1.000 | down |
| G002862 | SLC25A37 | 0.914 | 0.001 | 1.000 | down |
| G002862 | PKM | 0.918 | 0.001 | 1.000 | down |
| G002862 | WARS2 | 0.938 | 0.001 | 1.000 | down |
| G002862 | NBR1 | 0.908 | 0.002 | 1.000 | down |
| G033752 | UCP2 | 0.903 | 0.002 | 1.000 | down |
| G075198 | PUS1 | 0.900 | 0.002 | 1.000 | down |
| PSMG3-AS1 (2) | NBR1 | 0.931 | 0.001 | 1.000 | down |
| DLG5-AS1 (1) | PMPCA | 0.907 | 0.002 | 1.000 | down |
| DLG5-AS1 (1) | PUS1 | 0.908 | 0.002 | 1.000 | down |
| DLG5-AS1 (1) | ATAD3A | 0.902 | 0.002 | 1.000 | down |
| DLG5-AS1 (1) | PPRC1 | 0.918 | 0.001 | 1.000 | down |
| DLG5-AS1 (1) | ABCF2 | 0.928 | 0.001 | 1.000 | down |
| DLG5-AS1 (1) | CKMT1B | 0.922 | 0.001 | 1.000 | down |
| SH3GL1P2 | PMPCA | 0.917 | 0.001 | 1.000 | down |
| SH3GL1P2 | TRAP1 | 0.962 | 0.000 | 1.000 | down |
| SH3GL1P2 | HSPA1B | 0.957 | 0.000 | 1.000 | down |
| SH3GL1P2 | PPRC1 | 0.939 | 0.001 | 1.000 | down |
| AC138655.6 | ATAD3A | 0.903 | 0.002 | 1.000 | down |
| AC138655.6 | ATAD3B | 0.912 | 0.002 | 1.000 | down |
| AC138655.6 | ABCF2 | 0.915 | 0.001 | 1.000 | down |
| AC138655.6 | LYRM7 | 0.936 | 0.001 | 1.000 | down |
| TM4SF1-AS1 | PMPCA | 0.914 | 0.001 | 1.000 | down |
| TM4SF1-AS1 | HSPA1B | 0.939 | 0.001 | 1.000 | down |
| TM4SF1-AS1 | OGG1 | 0.940 | 0.001 | 1.000 | down |
| TPTE2P3 | PKM | 0.949 | 0.000 | 1.000 | down |
| TPTE2P3 | DNAJC19 | 0.930 | 0.001 | 1.000 | down |
| LINC00662 | PMPCA | 0.926 | 0.001 | 1.000 | down |
| LINC00662 | PUS1 | 0.924 | 0.001 | 1.000 | down |

| | | | | | |
|---|---|---|---|---|---|
| LINC00662 | ATAD3B | 0.941 | 0.000 | 1.000 | down |
| LINC00662 | LYRM7 | 0.915 | 0.001 | 1.000 | down |
| LINC00662 | PNPLA8 | 0.937 | 0.001 | 1.000 | down |
| RP11-297J22.1 | ATAD3B | 0.917 | 0.001 | 1.000 | down |
| RP11-179B2.2 | PMPCA | 0.911 | 0.002 | 1.000 | down |
| RP11-179B2.2 | ATAD3B | 0.915 | 0.001 | 1.000 | down |
| RP11-179B2.2 | LYRM7 | 0.944 | 0.000 | 1.000 | down |
| RP11-179B2.2 | PNPLA8 | 0.918 | 0.001 | 1.000 | down |
| G064655 | PMPCA | 0.920 | 0.001 | 1.000 | down |
| G064655 | TRAP1 | 0.932 | 0.001 | 1.000 | down |
| G064655 | HSPA1B | 0.959 | 0.000 | 1.000 | down |
| G064655 | PPRC1 | 0.923 | 0.001 | 1.000 | down |
| G064655 | PDK1 | 0.912 | 0.002 | 1.000 | down |
| CTD-2588C8.8 | IBA57 | 0.909 | 0.002 | 1.000 | down |
| CTD-2588C8.8 | NBR1 | 0.932 | 0.001 | 1.000 | down |
| G057990 | EARS2 | 0.958 | 0.000 | 1.000 | down |
| G047225 | IBA57 | 0.944 | 0.000 | 1.000 | down |
| G047225 | CMPK2 | 0.902 | 0.002 | 1.000 | down |
| RP1-309F20.3 | IBA57 | 0.952 | 0.000 | 1.000 | down |
| RP1-309F20.3 | NBR1 | 0.915 | 0.001 | 1.000 | down |
| RP1-309F20.3 | CMPK2 | 0.949 | 0.000 | 1.000 | down |
| RP11-643M14.1 | DNAJC19 | 0.915 | 0.001 | 1.000 | down |
| RP11-643M14.1 | IBA57 | 0.932 | 0.001 | 1.000 | down |
| RP11-344E13.4 | NDUFAF7 | 0.900 | 0.002 | 1.000 | down |
| SCAMP1-AS1 | PKM | 0.916 | 0.001 | 1.000 | down |
| G017670 | ALDOC | 0.902 | 0.002 | 1.000 | down |
| G017670 | BNIP3L | 0.969 | 0.000 | 1.000 | down |
| RP11-21M24.2 | UCP2 | 0.958 | 0.000 | 1.000 | down |
| RP11-252A24.3 | HSPA1B | 0.944 | 0.000 | 1.000 | down |
| RP11-252A24.3 | OGG1 | 0.940 | 0.001 | 1.000 | down |
| XLOC_013660 | NDUFAF4 | 0.933 | 0.001 | 1.000 | down |
| XLOC_013660 | CCDC58 | 0.902 | 0.002 | 1.000 | down |
| AC073321.4 | HSPE1 | 0.919 | 0.001 | 1.000 | down |
| AC073321.4 | BNIP3L | 0.926 | 0.001 | 1.000 | down |
| XLOC_008875 | IBA57 | 0.919 | 0.001 | 1.000 | down |
| G077330 | MRPL12 | 0.911 | 0.002 | 1.000 | down |
| RP11-46J23.1 | PUS1 | 0.902 | 0.002 | 1.000 | down |
| RP11-715J22.6 | PMPCA | 0.963 | 0.000 | 1.000 | down |
| RP11-715J22.6 | PUS1 | 0.934 | 0.001 | 1.000 | down |
| RP11-715J22.6 | TRAP1 | 0.913 | 0.002 | 1.000 | down |
| RP11-715J22.6 | PGS1 | 0.906 | 0.002 | 1.000 | down |
| RP11-715J22.6 | PPRC1 | 0.957 | 0.000 | 1.000 | down |
| RP11-715J22.6 | CKMT1B | 0.947 | 0.000 | 1.000 | down |
| RP11-277L2.3 | MSRA | 0.957 | 0.000 | 1.000 | down |
| RP11-277L2.3 | FOXO3 | 0.946 | 0.000 | 1.000 | down |
| RP11-191L17.1 | LYRM7 | 0.913 | 0.002 | 1.000 | down |
| RP11-191L17.1 | PNPLA8 | 0.934 | 0.001 | 1.000 | down |
| RP11-191L17.1 | IBA57 | 0.934 | 0.001 | 1.000 | down |
| RP11-191L17.1 | OGG1 | 0.922 | 0.001 | 1.000 | down |
| CCDC144NL- | SLC25A37 | 0.944 | 0.000 | 1.000 | down |
| CCDC144NL- | PKM | 0.929 | 0.001 | 1.000 | down |
| CCDC144NL- | DNAJC19 | 0.924 | 0.001 | 1.000 | down |
| WWC2-AS2 | MSRA | 0.947 | 0.000 | 1.000 | down |
| WWC2-AS2 | FOXO3 | 0.936 | 0.001 | 1.000 | down |
| AC099754.1 | PMPCA | 0.945 | 0.000 | 1.000 | down |
| AC099754.1 | PUS1 | 0.937 | 0.001 | 1.000 | down |
| AC099754.1 | ATAD3B | 0.937 | 0.001 | 1.000 | down |
| AC099754.1 | PPRC1 | 0.904 | 0.002 | 1.000 | down |
| AC099754.1 | LYRM7 | 0.948 | 0.000 | 1.000 | down |
| AC099754.1 | PNPLA8 | 0.911 | 0.002 | 1.000 | down |
| RP11-165A20.3 | IBA57 | 0.967 | 0.000 | 1.000 | down |
| RP11-165A20.3 | NBR1 | 0.918 | 0.001 | 1.000 | down |
| G018162 | AURKAIP1 | 0.904 | 0.002 | 1.000 | down |
| G079009 | PHB | 0.914 | 0.002 | 1.000 | down |
| G079009 | PUS1 | 0.903 | 0.002 | 1.000 | down |

| | | | | | |
|---|---|---|---|---|---|
| G079009 | PGS1 | 0.912 | 0.002 | 1.000 | down |
| G079009 | ABCF2 | 0.926 | 0.001 | 1.000 | down |
| G079009 | CKMT1B | 0.923 | 0.001 | 1.000 | down |
| RP11-396C23.4 | PMPCA | 0.924 | 0.001 | 1.000 | down |
| RP11-396C23.4 | PUS1 | 0.956 | 0.000 | 1.000 | down |
| RP11-396C23.4 | PGS1 | 0.914 | 0.001 | 1.000 | down |
| RP11-396C23.4 | PPRC1 | 0.924 | 0.001 | 1.000 | down |
| RP11-396C23.4 | CKMT1B | 0.907 | 0.002 | 1.000 | down |
| G000481 | IBA57 | 0.950 | 0.000 | 1.000 | down |
| RP11-285F16.1 | PMPCA | 0.902 | 0.002 | 1.000 | down |
| RP11-285F16.1 | PTRH2 | 0.909 | 0.002 | 1.000 | down |
| RP11-285F16.1 | LYRM7 | 0.906 | 0.002 | 1.000 | down |
| RP11-285F16.1 | IBA57 | 0.938 | 0.001 | 1.000 | down |
| RP11-285F16.1 | NBR1 | 0.909 | 0.002 | 1.000 | down |
| G050681 | SLC25A37 | 0.929 | 0.001 | 1.000 | down |
| G050681 | PKM | 0.907 | 0.002 | 1.000 | down |
| G050681 | DNAJC19 | 0.925 | 0.001 | 1.000 | down |
| G050681 | WARS2 | 0.929 | 0.001 | 1.000 | down |
| G050681 | NBR1 | 0.977 | 0.000 | 1.000 | down |
| G048723 | CKMT1B | 0.925 | 0.001 | 1.000 | down |
| CTC-232P5.3 | PMPCA | 0.928 | 0.001 | 1.000 | down |
| CTC-232P5.3 | HSPA1B | 0.911 | 0.002 | 1.000 | down |
| CTC-232P5.3 | OGG1 | 0.923 | 0.001 | 1.000 | down |
| RP5-991G20.4 | PMPCA | 0.934 | 0.001 | 1.000 | down |
| RP5-991G20.4 | TRAP1 | 0.974 | 0.000 | 1.000 | down |
| RP5-991G20.4 | HSPA1B | 0.952 | 0.000 | 1.000 | down |
| RP5-991G20.4 | PPRC1 | 0.958 | 0.000 | 1.000 | down |
| RP5-1092A3.4 | PTRH2 | 0.914 | 0.002 | 1.000 | down |
| RP5-1092A3.4 | LYRM7 | 0.906 | 0.002 | 1.000 | down |
| RP5-1092A3.4 | IBA57 | 0.904 | 0.002 | 1.000 | down |
| BC112270 | EARS2 | 0.915 | 0.001 | 1.000 | down |
| G014329 | NDUFAF7 | 0.927 | 0.001 | 1.000 | down |
| G014329 | PGS1 | 0.922 | 0.001 | 1.000 | down |
| G014329 | ABCF2 | 0.925 | 0.001 | 1.000 | down |
| RP11-351D16.3 | LYRM7 | 0.935 | 0.001 | 1.000 | down |
| RP11-351D16.3 | IBA57 | 0.966 | 0.000 | 1.000 | down |
| MGC2889 | PMPCA | 0.963 | 0.000 | 1.000 | down |
| MGC2889 | PUS1 | 0.951 | 0.000 | 1.000 | down |
| MGC2889 | TRAP1 | 0.952 | 0.000 | 1.000 | down |
| MGC2889 | ATAD3B | 0.924 | 0.001 | 1.000 | down |
| MGC2889 | HSPA1B | 0.914 | 0.001 | 1.000 | down |
| MGC2889 | PPRC1 | 0.905 | 0.002 | 1.000 | down |
| MGC2889 | OGG1 | 0.931 | 0.001 | 1.000 | down |
| AK123758 | IBA57 | 0.939 | 0.001 | 1.000 | down |
| AK123758 | NBR1 | 0.932 | 0.001 | 1.000 | down |
| RP4-683L5.1 | ATAD3B | 0.901 | 0.002 | 1.000 | down |
| RP4-683L5.1 | LYRM7 | 0.937 | 0.001 | 1.000 | down |
| RP4-683L5.1 | PNPLA8 | 0.941 | 0.001 | 1.000 | down |
| RP4-683L5.1 | IBA57 | 0.934 | 0.001 | 1.000 | down |
| RP11-353K11.1 | IBA57 | 0.960 | 0.000 | 1.000 | down |
| G082153 | PPRC1 | 0.939 | 0.001 | 1.000 | down |
| HRAT5 | LYRM7 | 0.903 | 0.002 | 1.000 | down |
| HRAT5 | IBA57 | 0.929 | 0.001 | 1.000 | down |
| RP11-209D14.2 | LYRM7 | 0.924 | 0.001 | 1.000 | down |
| RP11-209D14.2 | PNPLA8 | 0.921 | 0.001 | 1.000 | down |
| AC002467.7 | PNPLA8 | 0.902 | 0.002 | 1.000 | down |
| AC002467.7 | CMPK2 | 0.924 | 0.001 | 1.000 | down |
| G015380 | IBA57 | 0.914 | 0.001 | 1.000 | down |
| G015380 | CMPK2 | 0.942 | 0.000 | 1.000 | down |
| CTD-2005H7.1 | AURKAIP1 | 0.926 | 0.001 | 1.000 | down |
| CTD-2005H7.1 | MRPL12 | 0.915 | 0.001 | 1.000 | down |
| CTD-2005H7.1 | PUS1 | 0.909 | 0.002 | 1.000 | down |
| CTD-2005H7.1 | PPRC1 | 0.918 | 0.001 | 1.000 | down |
| CTD-2005H7.1 | PDK1 | 0.941 | 0.000 | 1.000 | down |
| TAF1A-AS1 | SLC25A19 | 0.923 | 0.001 | 1.000 | down |

| | | | | | |
|---|---|---|---|---|---|
| uc.85 | UCP2 | 0.904 | 0.002 | 1.000 | down |
| XLOC_001189 | IBA57 | 0.920 | 0.001 | 1.000 | down |
| XLOC_003595 | BNIP3L | 0.918 | 0.001 | 1.000 | down |
| LINC00883 | BNIP3L | 0.935 | 0.001 | 1.000 | down |
| LINC00883 | OGG1 | 0.913 | 0.002 | 1.000 | down |
| G066319 | NDUFAF7 | 0.929 | 0.001 | 1.000 | down |
| G066319 | PGS1 | 0.920 | 0.001 | 1.000 | down |
| XLOC_006554 | PTRH2 | 0.920 | 0.001 | 1.000 | down |
| G051725 | SATB1 | 0.961 | 0.000 | 1.000 | down |
| FAM66A | PHB | 0.921 | 0.001 | 1.000 | down |
| FAM66A | NDUFAF7 | 0.935 | 0.001 | 1.000 | down |
| G042247 | PUS1 | 0.915 | 0.001 | 1.000 | down |
| G042247 | CKMT1A | 0.912 | 0.002 | 1.000 | down |
| RP11-475I24.3 | PHB | 0.922 | 0.001 | 1.000 | down |
| RP11-475I24.3 | NDUFAF7 | 0.934 | 0.001 | 1.000 | down |
| RP11-475I24.3 | PGS1 | 0.960 | 0.000 | 1.000 | down |
| RP11-475I24.3 | ATAD3A | 0.962 | 0.000 | 1.000 | down |
| RP11-475I24.3 | CKMT1B | 0.904 | 0.002 | 1.000 | down |
| RP11-312B8.1 | IBA57 | 0.970 | 0.000 | 1.000 | down |
| RP11-312B8.1 | NBR1 | 0.909 | 0.002 | 1.000 | down |
| GAS5-AS1 | LYRM7 | 0.954 | 0.000 | 1.000 | down |
| GAS5-AS1 | PNPLA8 | 0.914 | 0.001 | 1.000 | down |
| GAS5-AS1 | IBA57 | 0.912 | 0.002 | 1.000 | down |
| AC144652.1 | PUS1 | 0.904 | 0.002 | 1.000 | down |
| AC144652.1 | PGS1 | 0.900 | 0.002 | 1.000 | down |
| AC144652.1 | ATAD3A | 0.908 | 0.002 | 1.000 | down |
| AC144652.1 | ATAD3B | 0.979 | 0.000 | 1.000 | down |
| AC144652.1 | PNPLA8 | 0.960 | 0.000 | 1.000 | down |
| RP11-204E9.1 | OGG1 | 0.950 | 0.000 | 1.000 | down |
| G085410 | PMPCA | 0.920 | 0.001 | 1.000 | down |
| G085410 | PTRH2 | 0.943 | 0.000 | 1.000 | down |
| G085410 | PUS1 | 0.949 | 0.000 | 1.000 | down |
| G085410 | PGS1 | 0.914 | 0.001 | 1.000 | down |
| G085410 | ATAD3B | 0.940 | 0.001 | 1.000 | down |
| G085410 | CKMT1A | 0.903 | 0.002 | 1.000 | down |
| G085410 | PNPLA8 | 0.908 | 0.002 | 1.000 | down |
| G085410 | OGG1 | 0.912 | 0.002 | 1.000 | down |
| DHRS4-AS1 (1) | OGG1 | 0.959 | 0.000 | 1.000 | down |
| RP3-410B11.1 | IBA57 | 0.946 | 0.000 | 1.000 | down |
| RP3-410B11.1 | NBR1 | 0.920 | 0.001 | 1.000 | down |
| G079888 | SLC25A37 | 0.921 | 0.001 | 1.000 | down |
| G079888 | PPRC1 | 0.936 | 0.001 | 1.000 | down |
| RP11-326A19.3 | PDK1 | 0.912 | 0.002 | 1.000 | down |
| G089005 | PTRH2 | 0.905 | 0.002 | 1.000 | down |
| G089005 | ATAD3B | 0.903 | 0.002 | 1.000 | down |
| G089005 | PNPLA8 | 0.919 | 0.001 | 1.000 | down |
| G089005 | IBA57 | 0.913 | 0.002 | 1.000 | down |
| G089005 | OGG1 | 0.913 | 0.002 | 1.000 | down |
| G048385 | SLC25A19 | 0.963 | 0.000 | 1.000 | down |
| OLMALINC | PMPCA | 0.923 | 0.001 | 1.000 | down |
| OLMALINC | PUS1 | 0.956 | 0.000 | 1.000 | down |
| OLMALINC | PGS1 | 0.919 | 0.001 | 1.000 | down |
| OLMALINC | ATAD3B | 0.907 | 0.002 | 1.000 | down |
| OLMALINC | OGG1 | 0.953 | 0.000 | 1.000 | down |
| AC093391.2 | WARS2 | 0.925 | 0.001 | 1.000 | down |
| AC093391.2 | NBR1 | 0.901 | 0.002 | 1.000 | down |
| LOC101929741 | IBA57 | 0.942 | 0.000 | 1.000 | down |
| LOC101929741 | NBR1 | 0.911 | 0.002 | 1.000 | down |
| G079654 | PTRH2 | 0.925 | 0.001 | 1.000 | down |
| G079654 | IBA57 | 0.948 | 0.000 | 1.000 | down |
| G079654 | NBR1 | 0.927 | 0.001 | 1.000 | down |
| TEX26-AS1 | IBA57 | 0.914 | 0.001 | 1.000 | down |
| RP11-496H1.2 | LYRM7 | 0.902 | 0.002 | 1.000 | down |
| G010185 | SATB1 | 0.908 | 0.002 | 1.000 | down |
| XLOC_006928 | SLC25A37 | 0.942 | 0.000 | 1.000 | down |

| | | | | | |
|---|---|---|---|---|---|
| XLOC_006928 | WARS2 | 0.974 | 0.000 | 1.000 | down |
| XLOC_006928 | NBR1 | 0.943 | 0.000 | 1.000 | down |
| XLOC_006928 | CMPK2 | 0.908 | 0.002 | 1.000 | down |
| G009018 | PTRH2 | 0.904 | 0.002 | 1.000 | down |
| G009018 | PDK1 | 0.900 | 0.002 | 1.000 | down |
| SUCLA2-AS1 | CKMT1A | 0.915 | 0.001 | 1.000 | down |
| G029844 | AURKAIP1 | 0.903 | 0.002 | 1.000 | down |
| RP11-2E11.10 | LYRM7 | 0.971 | 0.000 | 1.000 | down |
| RP11-2E11.10 | IBA57 | 0.908 | 0.002 | 1.000 | down |
| FRMD6-AS1 | IBA57 | 0.959 | 0.000 | 1.000 | down |
| FRMD6-AS1 | NBR1 | 0.909 | 0.002 | 1.000 | down |
| G052526 | PMPCA | 0.905 | 0.002 | 1.000 | down |
| G052526 | PPRC1 | 0.914 | 0.001 | 1.000 | down |
| G052526 | CKMT1B | 0.900 | 0.002 | 1.000 | down |
| LINC00408 | NDUFAF7 | 0.938 | 0.001 | 1.000 | down |
| LINC00408 | PGS1 | 0.952 | 0.000 | 1.000 | down |
| LINC00408 | ATAD3B | 0.909 | 0.002 | 1.000 | down |
| RP11-128P10.1 | PKM | 0.925 | 0.001 | 1.000 | down |
| RP11-128P10.1 | WARS2 | 0.906 | 0.002 | 1.000 | down |
| RP11-128P10.1 | NBR1 | 0.952 | 0.000 | 1.000 | down |
| RP11-382D12.2 | BNIP3L | 0.954 | 0.000 | 1.000 | down |
| RP11-174G6.5 | WARS2 | 0.932 | 0.001 | 1.000 | down |
| RP11-174G6.5 | NBR1 | 0.951 | 0.000 | 1.000 | down |
| RP11-174G6.5 | CMPK2 | 0.918 | 0.001 | 1.000 | down |
| AL355711 | IBA57 | 0.971 | 0.000 | 1.000 | down |
| AL355711 | NBR1 | 0.942 | 0.000 | 1.000 | down |
| AL355711 | CMPK2 | 0.914 | 0.001 | 1.000 | down |
| CTD-2256P15.2 | IBA57 | 0.980 | 0.000 | 1.000 | down |
| CTD-2256P15.2 | NBR1 | 0.929 | 0.001 | 1.000 | down |
| G053260 | CKMT1B | 0.910 | 0.002 | 1.000 | down |
| BC070391 | LYRM7 | 0.915 | 0.001 | 1.000 | down |
| BC070391 | PNPLA8 | 0.908 | 0.002 | 1.000 | down |
| BC070391 | IBA57 | 0.942 | 0.000 | 1.000 | down |
| SIX3-AS1 | SLC25A19 | 0.933 | 0.001 | 1.000 | down |
| G034470 | IBA57 | 0.935 | 0.001 | 1.000 | down |
| MAPT-IT1 | PUS1 | 0.927 | 0.001 | 1.000 | down |
| MAPT-IT1 | ATAD3B | 0.946 | 0.000 | 1.000 | down |
| MAPT-IT1 | PNPLA8 | 0.914 | 0.002 | 1.000 | down |
| LINC01132 | PKM | 0.978 | 0.000 | 1.000 | down |
| LINC01132 | DNAJC19 | 0.939 | 0.001 | 1.000 | down |
| G041225 | PPRC1 | 0.931 | 0.001 | 1.000 | down |
| G073576 | ATAD3B | 0.910 | 0.002 | 1.000 | down |
| G073576 | LYRM7 | 0.904 | 0.002 | 1.000 | down |
| G073576 | PNPLA8 | 0.952 | 0.000 | 1.000 | down |
| G073576 | IBA57 | 0.917 | 0.001 | 1.000 | down |
| XLOC_013222 | ATAD3B | 0.938 | 0.001 | 1.000 | down |
| XLOC_013222 | LYRM7 | 0.950 | 0.000 | 1.000 | down |
| XLOC_013222 | PNPLA8 | 0.967 | 0.000 | 1.000 | down |
| RP11-299H21.1 | SLC25A37 | 0.923 | 0.001 | 1.000 | down |
| RP11-299H21.1 | HSPA1B | 0.914 | 0.002 | 1.000 | down |
| RP11-299H21.1 | WARS2 | 0.902 | 0.002 | 1.000 | down |
| RP11-299H21.1 | PDK1 | 0.936 | 0.001 | 1.000 | down |
| ITFG1-AS1 | CCDC58 | 0.946 | 0.000 | 1.000 | down |
| G081251 | PTRH2 | 0.909 | 0.002 | 1.000 | down |
| G081251 | NBR1 | 0.900 | 0.002 | 1.000 | down |
| G081251 | SATB1 | 0.930 | 0.001 | 1.000 | down |
| G074033 | PTRH2 | 0.907 | 0.002 | 1.000 | down |
| G074033 | CKMT1A | 0.901 | 0.002 | 1.000 | down |
| CTD- | OGG1 | 0.911 | 0.002 | 1.000 | down |
| RP11-424N24.2 | NBR1 | 0.968 | 0.000 | 1.000 | down |
| RP11-424N24.2 | FOXO3 | 0.910 | 0.002 | 1.000 | down |
| CCNT2-AS1 | PMPCA | 0.966 | 0.000 | 1.000 | down |
| CCNT2-AS1 | PUS1 | 0.968 | 0.000 | 1.000 | down |
| CCNT2-AS1 | TRAP1 | 0.929 | 0.001 | 1.000 | down |
| CCNT2-AS1 | HSPA1B | 0.902 | 0.002 | 1.000 | down |

| | | | | | |
|---|---|---|---|---|---|
| CCNT2-AS1 | PPRC1 | 0.961 | 0.000 | 1.000 | down |
| CCNT2-AS1 | CKMT1B | 0.920 | 0.001 | 1.000 | down |
| XLOC_011052 | PMPCA | 0.925 | 0.001 | 1.000 | down |
| XLOC_011052 | PTRH2 | 0.918 | 0.001 | 1.000 | down |
| XLOC_011052 | IBA57 | 0.902 | 0.002 | 1.000 | down |
| XLOC_011052 | NBR1 | 0.905 | 0.002 | 1.000 | down |
| XLOC_011052 | OGG1 | 0.945 | 0.000 | 1.000 | down |
| G061579 | PHB | 0.914 | 0.001 | 1.000 | down |
| RP11-502I4.3 | PTRH2 | 0.902 | 0.002 | 1.000 | down |
| RP11-502I4.3 | NBR1 | 0.920 | 0.001 | 1.000 | down |
| G023041 | PUS1 | 0.902 | 0.002 | 1.000 | down |
| G023041 | OGG1 | 0.922 | 0.001 | 1.000 | down |
| RP1-202O8.3 | IBA57 | 0.955 | 0.000 | 1.000 | down |
| RP1-202O8.3 | NBR1 | 0.949 | 0.000 | 1.000 | down |
| G087757 | ATAD3A | 0.902 | 0.002 | 1.000 | down |
| G087757 | CKMT1A | 0.907 | 0.002 | 1.000 | down |
| RP13-131K19.2 | SLC25A37 | 0.958 | 0.000 | 1.000 | down |
| RP13-131K19.2 | DNAJC19 | 0.930 | 0.001 | 1.000 | down |
| RP13-131K19.2 | WARS2 | 0.911 | 0.002 | 1.000 | down |
| RP13-131K19.2 | NBR1 | 0.940 | 0.001 | 1.000 | down |
| G082866 | ATAD3B | 0.905 | 0.002 | 1.000 | down |
| G082866 | LYRM7 | 0.956 | 0.000 | 1.000 | down |
| G082866 | PNPLA8 | 0.961 | 0.000 | 1.000 | down |
| G013286 | MSRA | 0.928 | 0.001 | 1.000 | down |
| G013286 | FOXO3 | 0.901 | 0.002 | 1.000 | down |
| FOXN3-AS1 | SLC25A37 | 0.934 | 0.001 | 1.000 | down |
| FOXN3-AS1 | DNAJC19 | 0.905 | 0.002 | 1.000 | down |
| FOXN3-AS1 | WARS2 | 0.929 | 0.001 | 1.000 | down |
| FOXN3-AS1 | IBA57 | 0.909 | 0.002 | 1.000 | down |
| FOXN3-AS1 | NBR1 | 0.953 | 0.000 | 1.000 | down |
| RP11-290L1.2 | HSPE1 | 0.920 | 0.001 | 1.000 | down |
| RP11-290L1.2 | NBR1 | 0.935 | 0.001 | 1.000 | down |
| RP11-290L1.2 | MSRA | 0.907 | 0.002 | 1.000 | down |
| RP11-290L1.2 | FOXO3 | 0.942 | 0.000 | 1.000 | down |
| TNRC18P1 | PUS1 | 0.907 | 0.002 | 1.000 | down |
| TNRC18P1 | ATAD3B | 0.914 | 0.001 | 1.000 | down |
| GSE61474_XLOC_038226 | LYRM7 | 0.967 | 0.000 | 1.000 | down |
| GSE61474_XLOC_038226 | IBA57 | 0.920 | 0.001 | 1.000 | down |
| IDH1-AS1 | HSPE1 | 0.904 | 0.002 | 1.000 | down |
| IDH1-AS1 | AURKAIP1 | 0.913 | 0.002 | 1.000 | down |
| IDH1-AS1 | PTRH2 | 0.921 | 0.001 | 1.000 | down |
| IDH1-AS1 | NBR1 | 0.959 | 0.000 | 1.000 | down |
| G065067 | LYRM7 | 0.903 | 0.002 | 1.000 | down |
| INAFM1 | SLC25A37 | 0.901 | 0.002 | 1.000 | down |
| INAFM1 | WARS2 | 0.919 | 0.001 | 1.000 | down |
| NIPBL-AS1 | PUS1 | 0.910 | 0.002 | 1.000 | down |
| G038972 | LYRM7 | 0.947 | 0.000 | 1.000 | down |
| G038972 | IBA57 | 0.954 | 0.000 | 1.000 | down |
| LINC-PINT (2) | IBA57 | 0.942 | 0.000 | 1.000 | down |
| LOC100379224 | AURKAIP1 | 0.920 | 0.001 | 1.000 | down |
| LOC100379224 | MRPL12 | 0.947 | 0.000 | 1.000 | down |
| LOC100379224 | PDK1 | 0.934 | 0.001 | 1.000 | down |
| RP11-505K9.1 | IBA57 | 0.950 | 0.000 | 1.000 | down |
| RP11-505K9.1 | NBR1 | 0.928 | 0.001 | 1.000 | down |
| RP11-792A8.4 | NBR1 | 0.908 | 0.002 | 1.000 | down |
| RP11-792A8.4 | MSRA | 0.917 | 0.001 | 1.000 | down |
| RP11-792A8.4 | FOXO3 | 0.946 | 0.000 | 1.000 | down |
| G063668 | HSPE1 | 0.909 | 0.002 | 1.000 | down |
| G063668 | BNIP3L | 0.943 | 0.000 | 1.000 | down |
| G063668 | MSRA | 0.916 | 0.001 | 1.000 | down |
| RP11-138E2.1 | ATAD3B | 0.923 | 0.001 | 1.000 | down |
| RP11-138E2.1 | LYRM7 | 0.942 | 0.000 | 1.000 | down |
| RP11-138E2.1 | PNPLA8 | 0.944 | 0.000 | 1.000 | down |

| | | | | | |
|---|---|---|---|---|---|
| RP11-138E2.1 | IBA57 | 0.919 | 0.001 | 1.000 | down |
| RP11-775D22.3 | PMPCA | 0.927 | 0.001 | 1.000 | down |
| RP11-775D22.3 | TRAP1 | 0.931 | 0.001 | 1.000 | down |
| RP11-775D22.3 | SLC25A37 | 0.901 | 0.002 | 1.000 | down |
| RP11-775D22.3 | HSPA1B | 0.985 | 0.000 | 1.000 | down |
| RP11-775D22.3 | OGG1 | 0.915 | 0.001 | 1.000 | down |
| G003798 | IBA57 | 0.967 | 0.000 | 1.000 | down |
| G053697 | PUS1 | 0.970 | 0.000 | 1.000 | down |
| G053697 | PGS1 | 0.975 | 0.000 | 1.000 | down |
| G053697 | ATAD3A | 0.975 | 0.000 | 1.000 | down |
| G053697 | ATAD3B | 0.963 | 0.000 | 1.000 | down |
| G053697 | CKMT1A | 0.903 | 0.002 | 1.000 | down |
| G053697 | CKMT1B | 0.962 | 0.000 | 1.000 | down |
| G006537 | MRPL12 | 0.905 | 0.002 | 1.000 | down |
| G066606 | HSPA1B | 0.934 | 0.001 | 1.000 | down |
| LOC730102 | PUS1 | 0.920 | 0.001 | 1.000 | down |
| LOC730102 | ATAD3B | 0.908 | 0.002 | 1.000 | down |
| AK026244 | DNAJC19 | 0.916 | 0.001 | 1.000 | down |
| AK026244 | IBA57 | 0.915 | 0.001 | 1.000 | down |
| AK026244 | NBR1 | 0.901 | 0.002 | 1.000 | down |
| G059716 | ATAD3B | 0.930 | 0.001 | 1.000 | down |
| G059716 | PNPLA8 | 0.938 | 0.001 | 1.000 | down |
| RP11-2E11.5 | NBR1 | 0.903 | 0.002 | 1.000 | down |
| LOC148709 | SLC25A37 | 0.928 | 0.001 | 1.000 | down |
| LOC148709 | DNAJC19 | 0.920 | 0.001 | 1.000 | down |
| LOC148709 | IBA57 | 0.924 | 0.001 | 1.000 | down |
| LOC148709 | NBR1 | 0.925 | 0.001 | 1.000 | down |
| G044581 | PNPLA8 | 0.902 | 0.002 | 1.000 | down |
| G044581 | IBA57 | 0.950 | 0.000 | 1.000 | down |
| RP11-567G11.1 | NBR1 | 0.916 | 0.001 | 1.000 | down |
| RP11-567G11.1 | FOXO3 | 0.946 | 0.000 | 1.000 | down |
| RP5-884G6.2 | IBA57 | 0.936 | 0.001 | 1.000 | down |
| PVRL3-AS1 | PTRH2 | 0.959 | 0.000 | 1.000 | down |
| PVRL3-AS1 | LYRM7 | 0.911 | 0.002 | 1.000 | down |
| PVRL3-AS1 | CKMT1A | 0.911 | 0.002 | 1.000 | down |
| G010782 | PUS1 | 0.922 | 0.001 | 1.000 | down |
| G010782 | PGS1 | 0.937 | 0.001 | 1.000 | down |
| G010782 | ATAD3A | 0.947 | 0.000 | 1.000 | down |
| G010782 | ABCF2 | 0.926 | 0.001 | 1.000 | down |
| G010782 | CKMT1B | 0.948 | 0.000 | 1.000 | down |
| BC042589 | ATAD3B | 0.922 | 0.001 | 1.000 | down |
| BC042589 | PNPLA8 | 0.938 | 0.001 | 1.000 | down |
| BC042589 | OGG1 | 0.942 | 0.000 | 1.000 | down |
| LINC00634 | PUS1 | 0.926 | 0.001 | 1.000 | down |
| LINC00634 | ATAD3B | 0.978 | 0.000 | 1.000 | down |
| LINC00634 | PNPLA8 | 0.967 | 0.000 | 1.000 | down |
| LINC00634 | OGG1 | 0.912 | 0.002 | 1.000 | down |
| CTD-2383M3.1 | LYRM7 | 0.946 | 0.000 | 1.000 | down |
| CTD-2383M3.1 | IBA57 | 0.931 | 0.001 | 1.000 | down |
| AC092667.2 | LYRM7 | 0.947 | 0.000 | 1.000 | down |
| AC092667.2 | IBA57 | 0.934 | 0.001 | 1.000 | down |
| GYG2P1 | SLC25A37 | 0.905 | 0.002 | 1.000 | down |
| GYG2P1 | PKM | 0.950 | 0.000 | 1.000 | down |
| GYG2P1 | DNAJC19 | 0.920 | 0.001 | 1.000 | down |
| GYG2P1 | WARS2 | 0.911 | 0.002 | 1.000 | down |
| GYG2P1 | NBR1 | 0.947 | 0.000 | 1.000 | down |
| RP11-376P6.3 | IBA57 | 0.943 | 0.000 | 1.000 | down |
| RP11-376P6.3 | NBR1 | 0.903 | 0.002 | 1.000 | down |
| FGF13-AS1 | PKM | 0.922 | 0.001 | 1.000 | down |
| FGF13-AS1 | WARS2 | 0.902 | 0.002 | 1.000 | down |
| FGF13-AS1 | NBR1 | 0.949 | 0.000 | 1.000 | down |
| RP11-383C5.5 | IBA57 | 0.985 | 0.000 | 1.000 | down |
| RP11-383C5.5 | NBR1 | 0.914 | 0.002 | 1.000 | down |
| RP11-383C5.5 | CMPK2 | 0.908 | 0.002 | 1.000 | down |
| G001739 | IBA57 | 0.979 | 0.000 | 1.000 | down |

| | | | | | |
|---|---|---|---|---|---|
| AC005220.3 | AURKAIP1 | 0.912 | 0.002 | 1.000 | down |
| AC005220.3 | HSPA1B | 0.929 | 0.001 | 1.000 | down |
| AC005220.3 | PDK1 | 0.935 | 0.001 | 1.000 | down |
| AC005220.3 | OGG1 | 0.909 | 0.002 | 1.000 | down |
| RP11-22N19.2 | ATAD3B | 0.902 | 0.002 | 1.000 | down |
| RP11-22N19.2 | PNPLA8 | 0.932 | 0.001 | 1.000 | down |
| RP11-22N19.2 | IBA57 | 0.929 | 0.001 | 1.000 | down |
| RP11-51J9.5 | OGG1 | 0.936 | 0.001 | 1.000 | down |
| uc.95 | EARS2 | 0.938 | 0.001 | 1.000 | down |
| LOC101928402 | IBA57 | 0.916 | 0.001 | 1.000 | down |
| LOC101928402 | NBR1 | 0.917 | 0.001 | 1.000 | down |
| GFOD1 | IBA57 | 0.950 | 0.000 | 1.000 | down |
| LOC101928417 | IBA57 | 0.963 | 0.000 | 1.000 | down |
| GPR50-AS1 | HSPA1B | 0.934 | 0.001 | 1.000 | down |
| GPR50-AS1 | PDK1 | 0.935 | 0.001 | 1.000 | down |
| G078120 | PMPCA | 0.940 | 0.001 | 1.000 | down |
| G078120 | TRAP1 | 0.918 | 0.001 | 1.000 | down |
| G078120 | HSPA1B | 0.929 | 0.001 | 1.000 | down |
| G078120 | OGG1 | 0.954 | 0.000 | 1.000 | down |
| RP4-555D20.3 | PTRH2 | 0.920 | 0.001 | 1.000 | down |
| RP4-555D20.3 | LYRM7 | 0.921 | 0.001 | 1.000 | down |
| RP4-555D20.3 | PNPLA8 | 0.930 | 0.001 | 1.000 | down |
| RP4-555D20.3 | IBA57 | 0.929 | 0.001 | 1.000 | down |
| RP4-555D20.3 | NBR1 | 0.911 | 0.002 | 1.000 | down |
| RP11-507K2.3 | IBA57 | 0.961 | 0.000 | 1.000 | down |
| TOB1-AS1 (1) | PTRH2 | 0.928 | 0.001 | 1.000 | down |
| TOB1-AS1 (1) | CKMT1A | 0.929 | 0.001 | 1.000 | down |
| AP000255.6 | PNPLA8 | 0.912 | 0.002 | 1.000 | down |
| AP000255.6 | NBR1 | 0.927 | 0.001 | 1.000 | down |
| AK000953 | NBR1 | 0.908 | 0.002 | 1.000 | down |
| AK000953 | CMPK2 | 0.928 | 0.001 | 1.000 | down |
| RP4-798A10.7 | IBA57 | 0.914 | 0.002 | 1.000 | down |
| CTD-2377O17.1 | IBA57 | 0.966 | 0.000 | 1.000 | down |
| CTD-2377O17.1 | NBR1 | 0.940 | 0.001 | 1.000 | down |
| RP11-254F7.2 | PMPCA | 0.932 | 0.001 | 1.000 | down |
| RP11-254F7.2 | NBR1 | 0.925 | 0.001 | 1.000 | down |
| RP11-254F7.2 | OGG1 | 0.959 | 0.000 | 1.000 | down |
| G031140 | PHB | 0.914 | 0.002 | 1.000 | down |
| G031140 | PGS1 | 0.902 | 0.002 | 1.000 | down |
| PRR36 | SLC25A37 | 0.939 | 0.001 | 1.000 | down |
| LL22NC03- | PGS1 | 0.925 | 0.001 | 1.000 | down |
| LL22NC03- | ATAD3A | 0.942 | 0.000 | 1.000 | down |
| LL22NC03- | ATAD3B | 0.948 | 0.000 | 1.000 | down |
| LL22NC03- | PNPLA8 | 0.938 | 0.001 | 1.000 | down |
| RP11-332H14.1 | LYRM7 | 0.925 | 0.001 | 1.000 | down |
| RP11-332H14.1 | IBA57 | 0.925 | 0.001 | 1.000 | down |
| RP11-332H14.1 | NBR1 | 0.919 | 0.001 | 1.000 | down |
| G066437 | PMPCA | 0.939 | 0.001 | 1.000 | down |
| G066437 | PTRH2 | 0.903 | 0.002 | 1.000 | down |
| G066437 | PUS1 | 0.946 | 0.000 | 1.000 | down |
| G066437 | HSPA1B | 0.915 | 0.001 | 1.000 | down |
| G066437 | OGG1 | 0.951 | 0.000 | 1.000 | down |
| ENO1-AS1 | HSPE1 | 0.927 | 0.001 | 1.000 | down |
| ENO1-AS1 | PMPCA | 0.944 | 0.000 | 1.000 | down |
| ENO1-AS1 | PTRH2 | 0.936 | 0.001 | 1.000 | down |
| ENO1-AS1 | OGG1 | 0.948 | 0.000 | 1.000 | down |
| RP1-261G23.7 | IBA57 | 0.948 | 0.000 | 1.000 | down |
| RP1-261G23.7 | NBR1 | 0.909 | 0.002 | 1.000 | down |
| G023087 | IBA57 | 0.953 | 0.000 | 1.000 | down |
| G023087 | NBR1 | 0.932 | 0.001 | 1.000 | down |
| RP11-1275H24.1 | SLC25A37 | 0.908 | 0.002 | 1.000 | down |
| RP11-1275H24.1 | LYRM7 | 0.935 | 0.001 | 1.000 | down |
| RP1-223E5.4 | ATAD3B | 0.946 | 0.000 | 1.000 | down |
| RP1-223E5.4 | LYRM7 | 0.951 | 0.000 | 1.000 | down |
| RP1-223E5.4 | PNPLA8 | 0.965 | 0.000 | 1.000 | down |

| | | | | | |
|---|---|---|---|---|---|
| G037470 | IBA57 | 0.926 | 0.001 | 1.000 | down |
| RP11-680F8.1 | ALDOC | 0.985 | 0.000 | 1.000 | down |
| DGCR5 | PMPCA | 0.937 | 0.001 | 1.000 | down |
| DGCR5 | PUS1 | 0.916 | 0.001 | 1.000 | down |
| DGCR5 | PGS1 | 0.904 | 0.002 | 1.000 | down |
| DGCR5 | OGG1 | 0.977 | 0.000 | 1.000 | down |
| FAM66D | ATAD3A | 0.908 | 0.002 | 1.000 | down |
| FAM66D | ATAD3B | 0.933 | 0.001 | 1.000 | down |
| FAM66D | PNPLA8 | 0.952 | 0.000 | 1.000 | down |
| LINC00463 | PMPCA | 0.910 | 0.002 | 1.000 | down |
| LINC00463 | TRAP1 | 0.907 | 0.002 | 1.000 | down |
| G027543 | PMPCA | 0.944 | 0.000 | 1.000 | down |
| G027543 | OGG1 | 0.933 | 0.001 | 1.000 | down |
| RP11-573G6.8 | PNPLA8 | 0.923 | 0.001 | 1.000 | down |
| RP11-573G6.8 | OGG1 | 0.917 | 0.001 | 1.000 | down |
| RP11-107N15.1 | IBA57 | 0.935 | 0.001 | 1.000 | down |
| LL09NC01- | PMPCA | 0.928 | 0.001 | 1.000 | down |
| LL09NC01- | PTRH2 | 0.919 | 0.001 | 1.000 | down |
| LL09NC01- | ATAD3B | 0.920 | 0.001 | 1.000 | down |
| LL09NC01- | LYRM7 | 0.904 | 0.002 | 1.000 | down |
| LL09NC01- | PNPLA8 | 0.935 | 0.001 | 1.000 | down |
| LL09NC01- | OGG1 | 0.944 | 0.000 | 1.000 | down |
| G061722 | PTRH2 | 0.904 | 0.002 | 1.000 | down |
| G046707 | PKM | 0.936 | 0.001 | 1.000 | down |
| LOC100288846 | IBA57 | 0.971 | 0.000 | 1.000 | down |
| RP11-798M19.6 | PGS1 | 0.941 | 0.000 | 1.000 | down |
| RP11-798M19.6 | ATAD3B | 0.911 | 0.002 | 1.000 | down |
| G068576 | PMPCA | 0.936 | 0.001 | 1.000 | down |
| G068576 | TRAP1 | 0.908 | 0.002 | 1.000 | down |
| G068576 | OGG1 | 0.906 | 0.002 | 1.000 | down |
| RP11-872D17.4 | EARS2 | 0.923 | 0.001 | 1.000 | down |
| RP11-872D17.4 | MT-ND1 | 0.935 | 0.001 | 1.000 | down |
| XLOC_004632 | PMPCA | 0.904 | 0.002 | 1.000 | down |
| XLOC_004632 | PPRC1 | 0.967 | 0.000 | 1.000 | down |
| G036568 | UCP2 | 0.913 | 0.002 | 1.000 | down |
| CXADRP3 | PMPCA | 0.932 | 0.001 | 1.000 | down |
| CXADRP3 | PTRH2 | 0.902 | 0.002 | 1.000 | down |
| CXADRP3 | PUS1 | 0.920 | 0.001 | 1.000 | down |
| CXADRP3 | HSPA1B | 0.949 | 0.000 | 1.000 | down |
| CXADRP3 | PDK1 | 0.934 | 0.001 | 1.000 | down |
| CXADRP3 | OGG1 | 0.928 | 0.001 | 1.000 | down |
| UBAC2-AS1 | NBR1 | 0.919 | 0.001 | 1.000 | down |
| RP11-53B2.6 | PKM | 0.923 | 0.001 | 1.000 | down |
| RP11-53B2.6 | DNAJC19 | 0.934 | 0.001 | 1.000 | down |
| RP11-53B2.6 | NBR1 | 0.916 | 0.001 | 1.000 | down |
| DDX39B-AS1 | DNAJC19 | 0.966 | 0.000 | 1.000 | down |
| RP11-177G23.2 | PKM | 0.925 | 0.001 | 1.000 | down |
| RP11-177G23.2 | DNAJC19 | 0.928 | 0.001 | 1.000 | down |
| CTD-2382E5.2 | HSPE1 | 0.951 | 0.000 | 1.000 | down |
| CTD-2382E5.2 | PTRH2 | 0.942 | 0.000 | 1.000 | down |
| CTD-2382E5.2 | CKMT1A | 0.907 | 0.002 | 1.000 | down |
| EZR-AS1 | PTRH2 | 0.938 | 0.001 | 1.000 | down |
| EZR-AS1 | PUS1 | 0.931 | 0.001 | 1.000 | down |
| EZR-AS1 | CKMT1A | 0.978 | 0.000 | 1.000 | down |
| EZR-AS1 | CKMT1B | 0.944 | 0.000 | 1.000 | down |
| RP1-86C11.7 | PTRH2 | 0.920 | 0.001 | 1.000 | down |
| RP1-86C11.7 | IBA57 | 0.915 | 0.001 | 1.000 | down |
| RP1-86C11.7 | CMPK2 | 0.928 | 0.001 | 1.000 | down |
| G081642 | MT-ND1 | 0.950 | 0.000 | 1.000 | down |
| G058460 | BNIP3L | 0.937 | 0.001 | 1.000 | down |
| G047565 | OGG1 | 0.910 | 0.002 | 1.000 | down |
| RP11-793H13.11 | OGG1 | 0.944 | 0.000 | 1.000 | down |
| FLJ20021 | IBA57 | 0.923 | 0.001 | 1.000 | down |
| FLJ20021 | NBR1 | 0.912 | 0.002 | 1.000 | down |
| FLJ20021 | CMPK2 | 0.949 | 0.000 | 1.000 | down |

| | | | | | |
|---|---|---|---|---|---|
| RP11-109D9.4 | PTRH2 | 0.940 | 0.001 | 1.000 | down |
| RP11-109D9.4 | NBR1 | 0.923 | 0.001 | 1.000 | down |
| RP11-109D9.4 | OGG1 | 0.903 | 0.002 | 1.000 | down |
| RP11-110G21.1 | PMPCA | 0.978 | 0.000 | 1.000 | down |
| RP11-110G21.1 | TRAP1 | 0.925 | 0.001 | 1.000 | down |
| RP11-110G21.1 | HSPA1B | 0.902 | 0.002 | 1.000 | down |
| RP11-110G21.1 | OGG1 | 0.952 | 0.000 | 1.000 | down |
| CTC-444N24.13 | IBA57 | 0.942 | 0.000 | 1.000 | down |
| LINC00449 | PTRH2 | 0.902 | 0.002 | 1.000 | down |
| LINC00449 | IBA57 | 0.902 | 0.002 | 1.000 | down |
| RP11-61L19.2 | IBA57 | 0.984 | 0.000 | 1.000 | down |
| LOC101929066 | IBA57 | 0.930 | 0.001 | 1.000 | down |
| G054532 | AURKAIP1 | 0.935 | 0.001 | 1.000 | down |
| G054532 | HSPA1B | 0.955 | 0.000 | 1.000 | down |
| G054532 | PDK1 | 0.960 | 0.000 | 1.000 | down |
| RP11-85A1.3 | PNPLA8 | 0.913 | 0.002 | 1.000 | down |
| RP11-85A1.3 | IBA57 | 0.953 | 0.000 | 1.000 | down |
| G061596 | LYRM7 | 0.947 | 0.000 | 1.000 | down |
| AK125078 | HSPE1 | 0.943 | 0.000 | 1.000 | down |
| AK125078 | AURKAIP1 | 0.925 | 0.001 | 1.000 | down |
| AK125078 | PTRH2 | 0.921 | 0.001 | 1.000 | down |
| AK125078 | CKMT1A | 0.930 | 0.001 | 1.000 | down |
| BX248253 | PMPCA | 0.956 | 0.000 | 1.000 | down |
| BX248253 | AURKAIP1 | 0.928 | 0.001 | 1.000 | down |
| BX248253 | PTRH2 | 0.912 | 0.002 | 1.000 | down |
| BX248253 | PPRC1 | 0.907 | 0.002 | 1.000 | down |
| BX248253 | LYRM7 | 0.909 | 0.002 | 1.000 | down |
| BX248253 | NBR1 | 0.911 | 0.002 | 1.000 | down |
| uc.349 | PMPCA | 0.948 | 0.000 | 1.000 | down |
| uc.349 | OGG1 | 0.915 | 0.001 | 1.000 | down |
| RP11-344P13.6 | PNPLA8 | 0.921 | 0.001 | 1.000 | down |
| RP11-344P13.6 | IBA57 | 0.933 | 0.001 | 1.000 | down |
| CTC-359M8.1 | HSPA1B | 0.945 | 0.000 | 1.000 | down |
| uc.69 | SATB1 | 0.947 | 0.000 | 1.000 | down |
| LOC101927822 | MRPL12 | 0.933 | 0.001 | 1.000 | down |
| LOC101927822 | PPRC1 | 0.907 | 0.002 | 1.000 | down |
| G070405 | PTRH2 | 0.945 | 0.000 | 1.000 | down |
| RP11-565F19.2 | LYRM7 | 0.920 | 0.001 | 1.000 | down |
| G054079 | PTRH2 | 0.903 | 0.002 | 1.000 | down |
| G054079 | UCP2 | 0.921 | 0.001 | 1.000 | down |
| ANKRD20A12P | PUS1 | 0.904 | 0.002 | 1.000 | down |
| ANKRD20A12P | PGS1 | 0.928 | 0.001 | 1.000 | down |
| ANKRD20A12P | ATAD3A | 0.910 | 0.002 | 1.000 | down |
| ANKRD20A12P | ATAD3B | 0.936 | 0.001 | 1.000 | down |
| ANKRD20A12P | CKMT1A | 0.920 | 0.001 | 1.000 | down |
| ANKRD20A12P | CKMT1B | 0.922 | 0.001 | 1.000 | down |
| ANKRD20A12P | PNPLA8 | 0.925 | 0.001 | 1.000 | down |
| G003603 | PMPCA | 0.942 | 0.000 | 1.000 | down |
| G003603 | PUS1 | 0.908 | 0.002 | 1.000 | down |
| G003603 | PPRC1 | 0.924 | 0.001 | 1.000 | down |
| G003603 | LYRM7 | 0.902 | 0.002 | 1.000 | down |
| RP11-53I6.3 | PMPCA | 0.952 | 0.000 | 1.000 | down |
| RP11-53I6.3 | PTRH2 | 0.904 | 0.002 | 1.000 | down |
| RP11-53I6.3 | LYRM7 | 0.946 | 0.000 | 1.000 | down |
| RP11-53I6.3 | PNPLA8 | 0.906 | 0.002 | 1.000 | down |
| RP11-53I6.3 | NBR1 | 0.931 | 0.001 | 1.000 | down |
| RP11-314A20.5 | PMPCA | 0.954 | 0.000 | 1.000 | down |
| RP11-314A20.5 | TRAP1 | 0.958 | 0.000 | 1.000 | down |
| RP11-314A20.5 | HSPA1B | 0.926 | 0.001 | 1.000 | down |
| RP11-314A20.5 | PPRC1 | 0.914 | 0.002 | 1.000 | down |
| RP11-314A20.5 | OGG1 | 0.900 | 0.002 | 1.000 | down |
| AK055974 | LYRM7 | 0.945 | 0.000 | 1.000 | down |
| G060549 | DNAJC19 | 0.933 | 0.001 | 1.000 | down |
| RP11-474O21.5 | NBR1 | 0.932 | 0.001 | 1.000 | down |
| RP11-474O21.5 | CMPK2 | 0.923 | 0.001 | 1.000 | down |

| | | | | | |
|---|---|---|---|---|---|
| G046528 | NBR1 | 0.922 | 0.001 | 1.000 | down |
| CRYBB2P1 | SLC25A37 | 0.947 | 0.000 | 1.000 | down |
| CRYBB2P1 | HSPA1B | 0.902 | 0.002 | 1.000 | down |
| CRYBB2P1 | WARS2 | 0.921 | 0.001 | 1.000 | down |
| RP11-196G18.24 | PMPCA | 0.923 | 0.001 | 1.000 | down |
| RP11-196G18.24 | LYRM7 | 0.913 | 0.002 | 1.000 | down |
| RP11-196G18.24 | OGG1 | 0.912 | 0.002 | 1.000 | down |
| LOC101927811 | PMPCA | 0.922 | 0.001 | 1.000 | down |
| LOC101927811 | AURKAIP1 | 0.929 | 0.001 | 1.000 | down |
| LOC101927811 | TRAP1 | 0.904 | 0.002 | 1.000 | down |
| LOC101927811 | SLC25A37 | 0.909 | 0.002 | 1.000 | down |
| LOC101927811 | HSPA1B | 0.968 | 0.000 | 1.000 | down |
| LOC101927811 | PDK1 | 0.958 | 0.000 | 1.000 | down |
| CTD-2623N2.11 | LYRM7 | 0.949 | 0.000 | 1.000 | down |
| CTD-2623N2.11 | PNPLA8 | 0.917 | 0.001 | 1.000 | down |
| CTD-2623N2.11 | IBA57 | 0.946 | 0.000 | 1.000 | down |
| G064225 | HSPA1B | 0.915 | 0.001 | 1.000 | down |
| G064225 | UCP2 | 0.963 | 0.000 | 1.000 | down |
| uc.412 | MRPL12 | 0.942 | 0.000 | 1.000 | down |
| uc.412 | PPRC1 | 0.934 | 0.001 | 1.000 | down |
| uc.412 | PDK1 | 0.901 | 0.002 | 1.000 | down |
| CTC-350I8.1 | MT-ND1 | 0.932 | 0.001 | 1.000 | down |
| G023340 | ATAD3B | 0.950 | 0.000 | 1.000 | down |
| G023340 | LYRM7 | 0.931 | 0.001 | 1.000 | down |
| G023340 | PNPLA8 | 0.944 | 0.000 | 1.000 | down |
| H1FX-AS1 | UCP2 | 0.944 | 0.000 | 1.000 | down |
| KB-318B8.7 | PUS1 | 0.965 | 0.000 | 1.000 | down |
| KB-318B8.7 | PGS1 | 0.943 | 0.000 | 1.000 | down |
| KB-318B8.7 | ATAD3A | 0.974 | 0.000 | 1.000 | down |
| KB-318B8.7 | ATAD3B | 0.974 | 0.000 | 1.000 | down |
| KB-318B8.7 | CKMT1B | 0.926 | 0.001 | 1.000 | down |
| KB-318B8.7 | PNPLA8 | 0.907 | 0.002 | 1.000 | down |
| G063514 | SLC25A37 | 0.912 | 0.002 | 1.000 | down |
| G063514 | WARS2 | 0.944 | 0.000 | 1.000 | down |
| G063514 | NBR1 | 0.937 | 0.001 | 1.000 | down |
| G070622 | PHB | 0.911 | 0.002 | 1.000 | down |
| G014106 | PMPCA | 0.927 | 0.001 | 1.000 | down |
| G014106 | PTRH2 | 0.929 | 0.001 | 1.000 | down |
| G014106 | PUS1 | 0.989 | 0.000 | 1.000 | down |
| G014106 | PGS1 | 0.958 | 0.000 | 1.000 | down |
| G014106 | ATAD3B | 0.934 | 0.001 | 1.000 | down |
| G014106 | CKMT1A | 0.927 | 0.001 | 1.000 | down |
| G014106 | CKMT1B | 0.914 | 0.001 | 1.000 | down |
| RP1-232L24.3 | IBA57 | 0.920 | 0.001 | 1.000 | down |
| RP1-232L24.3 | NBR1 | 0.913 | 0.002 | 1.000 | down |
| RP11-467P9.1 | IBA57 | 0.915 | 0.001 | 1.000 | down |
| RP11-467P9.1 | NBR1 | 0.919 | 0.001 | 1.000 | down |
| RP11-467P9.1 | OGG1 | 0.905 | 0.002 | 1.000 | down |
| G002695 | PTRH2 | 0.926 | 0.001 | 1.000 | down |
| G002695 | IBA57 | 0.904 | 0.002 | 1.000 | down |
| G021588 | PTRH2 | 0.937 | 0.001 | 1.000 | down |
| G021588 | IBA57 | 0.907 | 0.002 | 1.000 | down |
| G021588 | NBR1 | 0.934 | 0.001 | 1.000 | down |
| G006578 | ATAD3B | 0.934 | 0.001 | 1.000 | down |
| G006578 | LYRM7 | 0.915 | 0.001 | 1.000 | down |
| G006578 | PNPLA8 | 0.951 | 0.000 | 1.000 | down |
| RP11-371A19.2 | LYRM7 | 0.938 | 0.001 | 1.000 | down |
| RP11-371A19.2 | IBA57 | 0.954 | 0.000 | 1.000 | down |
| RP11-95P2.3 | WARS2 | 0.936 | 0.001 | 1.000 | down |
| RP11-95P2.3 | NBR1 | 0.975 | 0.000 | 1.000 | down |
| RP11-95P2.3 | FOXO3 | 0.919 | 0.001 | 1.000 | down |
| RP11-118K6.3 | AURKAIP1 | 0.904 | 0.002 | 1.000 | down |
| RP11-118K6.3 | SATB1 | 0.961 | 0.000 | 1.000 | down |
| LOC101928489 | SLC25A19 | 0.921 | 0.001 | 1.000 | down |
| LOC101928489 | PTRH2 | 0.912 | 0.002 | 1.000 | down |

| | | | | | |
|---|---|---|---|---|---|
| RP11-355F22.1 | PHB | 0.903 | 0.002 | 1.000 | down |
| RP11-355F22.1 | PMPCA | 0.937 | 0.001 | 1.000 | down |
| RP11-355F22.1 | PUS1 | 0.929 | 0.001 | 1.000 | down |
| RP11-355F22.1 | PGS1 | 0.926 | 0.001 | 1.000 | down |
| RP11-355F22.1 | ATAD3A | 0.919 | 0.001 | 1.000 | down |
| RP11-355F22.1 | ATAD3B | 0.922 | 0.001 | 1.000 | down |
| RP11-355F22.1 | PPRC1 | 0.909 | 0.002 | 1.000 | down |
| RP11-355F22.1 | CKMT1B | 0.930 | 0.001 | 1.000 | down |
| JARID2-AS1 | MRPL12 | 0.953 | 0.000 | 1.000 | down |
| JARID2-AS1 | SLC25A37 | 0.923 | 0.001 | 1.000 | down |
| JARID2-AS1 | PPRC1 | 0.902 | 0.002 | 1.000 | down |
| G036604 | ABCF2 | 0.922 | 0.001 | 1.000 | down |
| G075319 | ATAD3B | 0.943 | 0.000 | 1.000 | down |
| G075319 | PNPLA8 | 0.952 | 0.000 | 1.000 | down |
| XLOC_010219 | UCP2 | 0.915 | 0.001 | 1.000 | down |
| GSE61474_XLOC_067608 | PNPLA8 | 0.926 | 0.001 | 1.000 | down |
| GSE61474_XLOC_067608 | CMPK2 | 0.957 | 0.000 | 1.000 | down |
| RP11-330O11.3 | LYRM7 | 0.904 | 0.002 | 1.000 | down |
| LOC283140 | PMPCA | 0.907 | 0.002 | 1.000 | down |
| LOC283140 | TRAP1 | 0.918 | 0.001 | 1.000 | down |
| LOC283140 | SLC25A37 | 0.903 | 0.002 | 1.000 | down |
| LOC283140 | HSPA1B | 0.979 | 0.000 | 1.000 | down |
| LOC283140 | PDK1 | 0.922 | 0.001 | 1.000 | down |
| DNAJC9-AS1 | PMPCA | 0.907 | 0.002 | 1.000 | down |
| DNAJC9-AS1 | NBR1 | 0.913 | 0.002 | 1.000 | down |
| RP11-820I16.4 | IBA57 | 0.979 | 0.000 | 1.000 | down |
| G021721 | OGG1 | 0.935 | 0.001 | 1.000 | down |
| G026062 | UCP2 | 0.920 | 0.001 | 1.000 | down |
| RP11-79P5.9 | PNPLA8 | 0.936 | 0.001 | 1.000 | down |
| G057611 | HSPA1B | 0.942 | 0.000 | 1.000 | down |
| CTB-181F24.1 | PMPCA | 0.909 | 0.002 | 1.000 | down |
| CTB-181F24.1 | PPRC1 | 0.912 | 0.002 | 1.000 | down |
| CTB-181F24.1 | LYRM7 | 0.973 | 0.000 | 1.000 | down |
| RP11-166B2.3 | PUS1 | 0.921 | 0.001 | 1.000 | down |
| RP11-166B2.3 | ATAD3B | 0.951 | 0.000 | 1.000 | down |
| RP11-166B2.3 | PNPLA8 | 0.942 | 0.000 | 1.000 | down |
| G075703 | PTRH2 | 0.952 | 0.000 | 1.000 | down |
| G075703 | IBA57 | 0.901 | 0.002 | 1.000 | down |
| G075703 | NBR1 | 0.925 | 0.001 | 1.000 | down |
| BC047644 | PMPCA | 0.908 | 0.002 | 1.000 | down |
| BC047644 | PUS1 | 0.969 | 0.000 | 1.000 | down |
| BC047644 | PGS1 | 0.908 | 0.002 | 1.000 | down |
| BC047644 | ATAD3B | 0.949 | 0.000 | 1.000 | down |
| BC047644 | OGG1 | 0.911 | 0.002 | 1.000 | down |
| RP11-820L6.1 | HSPE1 | 0.923 | 0.001 | 1.000 | down |
| RP11-820L6.1 | CCDC58 | 0.922 | 0.001 | 1.000 | down |
| RP11-820L6.1 | PTRH2 | 0.951 | 0.000 | 1.000 | down |
| LINC01604 | IBA57 | 0.963 | 0.000 | 1.000 | down |
| LOC102723385 | PMPCA | 0.965 | 0.000 | 1.000 | down |
| LOC102723385 | PTRH2 | 0.913 | 0.002 | 1.000 | down |
| LOC102723385 | HSPA1B | 0.921 | 0.001 | 1.000 | down |
| LOC102723385 | PPRC1 | 0.939 | 0.001 | 1.000 | down |
| RP11-803B1.8 | LYRM7 | 0.979 | 0.000 | 1.000 | down |
| G005619 | HSPE1 | 0.951 | 0.000 | 1.000 | down |
| G005619 | AURKAIP1 | 0.926 | 0.001 | 1.000 | down |
| G063721 | PMPCA | 0.913 | 0.002 | 1.000 | down |
| G063721 | HSPA1B | 0.902 | 0.002 | 1.000 | down |
| G063721 | PPRC1 | 0.929 | 0.001 | 1.000 | down |
| XLOC_004894 | NDUFAF4 | 0.902 | 0.002 | 1.000 | down |
| XLOC_004894 | EARS2 | 0.929 | 0.001 | 1.000 | down |
| XLOC_004894 | MT-ND1 | 0.964 | 0.000 | 1.000 | down |
| G015529 | AURKAIP1 | 0.935 | 0.001 | 1.000 | down |
| G015529 | SLC25A37 | 0.950 | 0.000 | 1.000 | down |

| | | | | | |
|---|---|---|---|---|---|
| G015529 | WARS2 | 0.929 | 0.001 | 1.000 | down |
| G015529 | NBR1 | 0.947 | 0.000 | 1.000 | down |
| CTD-2517O10.6 | NBR1 | 0.901 | 0.002 | 1.000 | down |
| G001497 | PMPCA | 0.949 | 0.000 | 1.000 | down |
| G001497 | OGG1 | 0.968 | 0.000 | 1.000 | down |
| GLIDR | PUS1 | 0.948 | 0.000 | 1.000 | down |
| GLIDR | PGS1 | 0.952 | 0.000 | 1.000 | down |
| GLIDR | ATAD3A | 0.929 | 0.001 | 1.000 | down |
| GLIDR | ATAD3B | 0.919 | 0.001 | 1.000 | down |
| RP11-439C15.4 | ATAD3B | 0.932 | 0.001 | 1.000 | down |
| RP11-439C15.4 | PNPLA8 | 0.959 | 0.000 | 1.000 | down |
| TOB1-AS1 (2) | DNAJC19 | 0.915 | 0.001 | 1.000 | down |
| TOB1-AS1 (2) | IBA57 | 0.922 | 0.001 | 1.000 | down |
| G007298 | EARS2 | 0.948 | 0.000 | 1.000 | down |
| LOC101927358 | PUS1 | 0.922 | 0.001 | 1.000 | down |
| LOC101927358 | ATAD3B | 0.901 | 0.002 | 1.000 | down |
| BC040628 | DNAJC19 | 0.914 | 0.001 | 1.000 | down |
| RP11-196G18.22 | LYRM7 | 0.943 | 0.000 | 1.000 | down |
| RP11-196G18.22 | PNPLA8 | 0.931 | 0.001 | 1.000 | down |
| CTC-248O19.1 | PMPCA | 0.927 | 0.001 | 1.000 | down |
| CTC-248O19.1 | PUS1 | 0.907 | 0.002 | 1.000 | down |
| CTC-248O19.1 | ABCF2 | 0.944 | 0.000 | 1.000 | down |
| RP11-379F4.9 | IBA57 | 0.930 | 0.001 | 1.000 | down |
| G014762 | PKM | 0.959 | 0.000 | 1.000 | down |
| G014762 | DNAJC19 | 0.970 | 0.000 | 1.000 | down |
| G085202 | NBR1 | 0.902 | 0.002 | 1.000 | down |
| LINC00670 | SATB1 | 0.916 | 0.001 | 1.000 | down |
| ALDOC | G057625 | -0.917 | 0.001 | -1.000 | down |
| ALDOC | G036779 | -0.907 | 0.002 | -1.000 | down |
| ALDOC | AC003682.16 | -0.964 | 0.000 | -1.000 | down |
| ALDOC | G022034 | -0.971 | 0.000 | -1.000 | down |
| ALDOC | G011388 | -0.939 | 0.001 | -1.000 | down |
| ALDOC | G002461 | -0.938 | 0.001 | -1.000 | down |
| ALDOC | AF086132 | -0.944 | 0.000 | -1.000 | down |
| ALDOC | G017670 | 0.902 | 0.002 | 1.000 | down |
| ALDOC | RP11-680F8.1 | 0.985 | 0.000 | 1.000 | down |
| PKM | G042193 | -0.901 | 0.002 | -1.000 | down |
| PKM | G056910 | -0.912 | 0.002 | -1.000 | down |
| PKM | IGBP1-AS2 | -0.922 | 0.001 | -1.000 | down |
| PKM | G009235 | -0.918 | 0.001 | -1.000 | down |
| PKM | ZFAT-AS1 | -0.912 | 0.002 | -1.000 | down |
| PKM | RP11-18D7.2 | -0.936 | 0.001 | -1.000 | down |
| PKM | CTD-2050E21.1 | -0.940 | 0.001 | -1.000 | down |
| PKM | BX648501 | -0.930 | 0.001 | -1.000 | down |
| PKM | RP11-651L5.2 | -0.905 | 0.002 | -1.000 | down |
| PKM | G068890 | -0.903 | 0.002 | -1.000 | down |
| PKM | XLOC_006779 | 0.918 | 0.001 | 1.000 | down |
| PKM | PSMG3-AS1 (1) | 0.909 | 0.002 | 1.000 | down |
| PKM | USP27X-AS1 | 0.922 | 0.001 | 1.000 | down |
| PKM | G002862 | 0.918 | 0.001 | 1.000 | down |
| PKM | TPTE2P3 | 0.949 | 0.000 | 1.000 | down |
| PKM | SCAMP1-AS1 | 0.916 | 0.001 | 1.000 | down |
| PKM | CCDC144NL-AS1 | 0.929 | 0.001 | 1.000 | down |
| PKM | G050681 | 0.907 | 0.002 | 1.000 | down |
| PKM | RP11-128P10.1 | 0.925 | 0.001 | 1.000 | down |
| PKM | LINC01132 | 0.978 | 0.000 | 1.000 | down |
| PKM | GYG2P1 | 0.950 | 0.000 | 1.000 | down |
| PKM | FGF13-AS1 | 0.922 | 0.001 | 1.000 | down |
| PKM | G046707 | 0.936 | 0.001 | 1.000 | down |
| PKM | RP11-53B2.6 | 0.923 | 0.001 | 1.000 | down |
| PKM | RP11-177G23.2 | 0.925 | 0.001 | 1.000 | down |
| PKM | G014762 | 0.959 | 0.000 | 1.000 | down |
| DNAJC19 | IGBP1-AS2 | -0.901 | 0.002 | -1.000 | down |
| DNAJC19 | RP11-18D7.2 | -0.904 | 0.002 | -1.000 | down |
| DNAJC19 | CTD-2050E21.1 | -0.950 | 0.000 | -1.000 | down |

| | | | | | |
|---|---|---|---|---|---|
| DNAJC19 | BX648501 | -0.921 | 0.001 | -1.000 | down |
| DNAJC19 | G010088 | -0.953 | 0.000 | -1.000 | down |
| DNAJC19 | RP11-651L5.2 | -0.967 | 0.000 | -1.000 | down |
| DNAJC19 | SPTY2D1-AS1 | -0.907 | 0.002 | -1.000 | down |
| DNAJC19 | XLOC_006779 | 0.945 | 0.000 | 1.000 | down |
| DNAJC19 | LOC100130417 | 0.930 | 0.001 | 1.000 | down |
| DNAJC19 | PSMG3-AS1 (1) | 0.941 | 0.000 | 1.000 | down |
| DNAJC19 | USP27X-AS1 | 0.934 | 0.001 | 1.000 | down |
| DNAJC19 | G070136 | 0.951 | 0.000 | 1.000 | down |
| DNAJC19 | TPTE2P3 | 0.930 | 0.001 | 1.000 | down |
| DNAJC19 | RP11-643M14.1 | 0.915 | 0.001 | 1.000 | down |
| DNAJC19 | CCDC144NL-AS1 | 0.924 | 0.001 | 1.000 | down |
| DNAJC19 | G050681 | 0.925 | 0.001 | 1.000 | down |
| DNAJC19 | LINC01132 | 0.939 | 0.001 | 1.000 | down |
| DNAJC19 | RP13-131K19.2 | 0.930 | 0.001 | 1.000 | down |
| DNAJC19 | FOXN3-AS1 | 0.905 | 0.002 | 1.000 | down |
| DNAJC19 | AK026244 | 0.916 | 0.001 | 1.000 | down |
| DNAJC19 | LOC148709 | 0.920 | 0.001 | 1.000 | down |
| DNAJC19 | GYG2P1 | 0.920 | 0.001 | 1.000 | down |
| DNAJC19 | RP11-53B2.6 | 0.934 | 0.001 | 1.000 | down |
| DNAJC19 | DDX39B-AS1 | 0.966 | 0.000 | 1.000 | down |
| DNAJC19 | RP11-177G23.2 | 0.928 | 0.001 | 1.000 | down |
| DNAJC19 | G060549 | 0.933 | 0.001 | 1.000 | down |
| DNAJC19 | TOB1-AS1 (2) | 0.915 | 0.001 | 1.000 | down |
| DNAJC19 | BC040628 | 0.914 | 0.001 | 1.000 | down |
| DNAJC19 | G014762 | 0.970 | 0.000 | 1.000 | down |
| UCP2 | G056910 | -0.923 | 0.001 | -1.000 | down |
| UCP2 | G038103 | -0.920 | 0.001 | -1.000 | down |
| UCP2 | RP11-484O2.1 | -0.904 | 0.002 | -1.000 | down |
| UCP2 | XLOC_012579 | -0.952 | 0.000 | -1.000 | down |
| UCP2 | G043880 | -0.937 | 0.001 | -1.000 | down |
| UCP2 | XLOC_006485 | -0.910 | 0.002 | -1.000 | down |
| UCP2 | TMLHE-AS1 | -0.914 | 0.002 | -1.000 | down |
| UCP2 | G009329 | -0.907 | 0.002 | -1.000 | down |
| UCP2 | G060726 | -0.969 | 0.000 | -1.000 | down |
| UCP2 | GSE61474_XLOC_028262 | -0.951 | 0.000 | -1.000 | down |
| UCP2 | RP11-354H21.2 | -0.925 | 0.001 | -1.000 | down |
| UCP2 | G033752 | 0.903 | 0.002 | 1.000 | down |
| UCP2 | RP11-21M24.2 | 0.958 | 0.000 | 1.000 | down |
| UCP2 | uc.85 | 0.904 | 0.002 | 1.000 | down |
| UCP2 | G036568 | 0.913 | 0.002 | 1.000 | down |
| UCP2 | G054079 | 0.921 | 0.001 | 1.000 | down |
| UCP2 | G064225 | 0.963 | 0.000 | 1.000 | down |
| UCP2 | H1FX-AS1 | 0.944 | 0.000 | 1.000 | down |
| UCP2 | XLOC_010219 | 0.915 | 0.001 | 1.000 | down |
| UCP2 | G026062 | 0.920 | 0.001 | 1.000 | down |
| LYRM7 | G074385 | -0.930 | 0.001 | -1.000 | down |
| LYRM7 | G087508 | -0.956 | 0.000 | -1.000 | down |
| LYRM7 | G071640 | -0.957 | 0.000 | -1.000 | down |
| LYRM7 | RP11-717D12.1 | -0.919 | 0.001 | -1.000 | down |
| LYRM7 | G037029 | -0.924 | 0.001 | -1.000 | down |
| LYRM7 | CTD-3193O13.1 | -0.936 | 0.001 | -1.000 | down |
| LYRM7 | LOC105373609 | -0.911 | 0.002 | -1.000 | down |
| LYRM7 | G089283 | -0.901 | 0.002 | -1.000 | down |
| LYRM7 | RP5-1112D6.7 | -0.958 | 0.000 | -1.000 | down |
| LYRM7 | G013179 | -0.924 | 0.001 | -1.000 | down |
| LYRM7 | RP11-461K13.1 | -0.932 | 0.001 | -1.000 | down |
| LYRM7 | AL110181 | 0.936 | 0.001 | 1.000 | down |
| LYRM7 | RP11-363E6.3 | 0.969 | 0.000 | 1.000 | down |
| LYRM7 | AK055712 | 0.959 | 0.000 | 1.000 | down |
| LYRM7 | RP11-303E16.7 | 0.911 | 0.002 | 1.000 | down |
| LYRM7 | RP4-549L20.3 | 0.966 | 0.000 | 1.000 | down |
| LYRM7 | RP11-686O6.2 | 0.950 | 0.000 | 1.000 | down |
| LYRM7 | RP11-356J5.12 | 0.919 | 0.001 | 1.000 | down |

| | | | | | |
|---|---|---|---|---|---|
| LYRM7 | RP1-40E16.12 | 0.918 | 0.001 | 1.000 | down |
| LYRM7 | RP11-490O6.2 | 0.913 | 0.002 | 1.000 | down |
| LYRM7 | AL137474 | 0.933 | 0.001 | 1.000 | down |
| LYRM7 | AC138655.6 | 0.936 | 0.001 | 1.000 | down |
| LYRM7 | LINC00662 | 0.915 | 0.001 | 1.000 | down |
| LYRM7 | RP11-179B2.2 | 0.944 | 0.000 | 1.000 | down |
| LYRM7 | RP11-191L17.1 | 0.913 | 0.002 | 1.000 | down |
| LYRM7 | AC099754.1 | 0.948 | 0.000 | 1.000 | down |
| LYRM7 | RP11-285F16.1 | 0.906 | 0.002 | 1.000 | down |
| LYRM7 | RP5-1092A3.4 | 0.906 | 0.002 | 1.000 | down |
| LYRM7 | RP11-351D16.3 | 0.935 | 0.001 | 1.000 | down |
| LYRM7 | RP4-683L5.1 | 0.937 | 0.001 | 1.000 | down |
| LYRM7 | HRAT5 | 0.903 | 0.002 | 1.000 | down |
| LYRM7 | RP11-209D14.2 | 0.924 | 0.001 | 1.000 | down |
| LYRM7 | GAS5-AS1 | 0.954 | 0.000 | 1.000 | down |
| LYRM7 | RP11-496H1.2 | 0.902 | 0.002 | 1.000 | down |
| LYRM7 | RP11-2E11.10 | 0.971 | 0.000 | 1.000 | down |
| LYRM7 | BC070391 | 0.915 | 0.001 | 1.000 | down |
| LYRM7 | G073576 | 0.904 | 0.002 | 1.000 | down |
| LYRM7 | XLOC_013222 | 0.950 | 0.000 | 1.000 | down |
| LYRM7 | G082866 | 0.956 | 0.000 | 1.000 | down |
| LYRM7 | GSE61474_XLOC_038226 | 0.967 | 0.000 | 1.000 | down |
| LYRM7 | G065067 | 0.903 | 0.002 | 1.000 | down |
| LYRM7 | G038972 | 0.947 | 0.000 | 1.000 | down |
| LYRM7 | RP11-138E2.1 | 0.942 | 0.000 | 1.000 | down |
| LYRM7 | PVRL3-AS1 | 0.911 | 0.002 | 1.000 | down |
| LYRM7 | CTD-2383M3.1 | 0.946 | 0.000 | 1.000 | down |
| LYRM7 | AC092667.2 | 0.947 | 0.000 | 1.000 | down |
| LYRM7 | RP4-555D20.3 | 0.921 | 0.001 | 1.000 | down |
| LYRM7 | RP11-332H14.1 | 0.925 | 0.001 | 1.000 | down |
| LYRM7 | RP11-1275H24.1 | 0.935 | 0.001 | 1.000 | down |
| LYRM7 | RP1-223E5.4 | 0.951 | 0.000 | 1.000 | down |
| LYRM7 | LL09NC01-139C3.1 | 0.904 | 0.002 | 1.000 | down |
| LYRM7 | G061596 | 0.947 | 0.000 | 1.000 | down |
| LYRM7 | BX248253 | 0.909 | 0.002 | 1.000 | down |
| LYRM7 | RP11-565F19.2 | 0.920 | 0.001 | 1.000 | down |
| LYRM7 | G003603 | 0.902 | 0.002 | 1.000 | down |
| LYRM7 | RP11-53I6.3 | 0.946 | 0.000 | 1.000 | down |
| LYRM7 | AK055974 | 0.945 | 0.000 | 1.000 | down |
| LYRM7 | RP11-196G18.24 | 0.913 | 0.002 | 1.000 | down |
| LYRM7 | CTD-2623N2.11 | 0.949 | 0.000 | 1.000 | down |
| LYRM7 | G023340 | 0.931 | 0.001 | 1.000 | down |
| LYRM7 | G006578 | 0.915 | 0.001 | 1.000 | down |
| LYRM7 | RP11-371A19.2 | 0.938 | 0.001 | 1.000 | down |
| LYRM7 | RP11-330O11.3 | 0.904 | 0.002 | 1.000 | down |
| LYRM7 | CTB-181F24.1 | 0.973 | 0.000 | 1.000 | down |
| LYRM7 | RP11-803B1.8 | 0.979 | 0.000 | 1.000 | down |
| LYRM7 | RP11-196G18.22 | 0.943 | 0.000 | 1.000 | down |
| WARS2 | PPIEL | -0.909 | 0.002 | -1.000 | down |
| WARS2 | G020731 | -0.920 | 0.001 | -1.000 | down |
| WARS2 | G073552 | -0.917 | 0.001 | -1.000 | down |
| WARS2 | G046678 | -0.906 | 0.002 | -1.000 | down |
| WARS2 | AC016682.1 | -0.915 | 0.001 | -1.000 | down |
| WARS2 | RP11-484O2.1 | -0.908 | 0.002 | -1.000 | down |
| WARS2 | G050702 | -0.950 | 0.000 | -1.000 | down |
| WARS2 | G063995 | -0.903 | 0.002 | -1.000 | down |
| WARS2 | ZFAT-AS1 | -0.956 | 0.000 | -1.000 | down |
| WARS2 | AK093193 | -0.974 | 0.000 | -1.000 | down |
| WARS2 | CTD-2050E21.1 | -0.910 | 0.002 | -1.000 | down |
| WARS2 | RP11-424I19.2 | -0.903 | 0.002 | -1.000 | down |
| WARS2 | G060178 | -0.925 | 0.001 | -1.000 | down |
| WARS2 | BX648501 | -0.905 | 0.002 | -1.000 | down |
| WARS2 | G045736 | -0.936 | 0.001 | -1.000 | down |
| WARS2 | G059595 | -0.920 | 0.001 | -1.000 | down |

| | | | | | |
|---|---|---|---|---|---|
| WARS2 | RP11-744I24.3 | 0.936 | 0.001 | 1.000 | down |
| WARS2 | GSE61474_XLOC_054139 | 0.912 | 0.002 | 1.000 | down |
| WARS2 | USP27X-AS1 | 0.942 | 0.000 | 1.000 | down |
| WARS2 | G002862 | 0.938 | 0.001 | 1.000 | down |
| WARS2 | G050681 | 0.929 | 0.001 | 1.000 | down |
| WARS2 | AC093391.2 | 0.925 | 0.001 | 1.000 | down |
| WARS2 | XLOC_006928 | 0.974 | 0.000 | 1.000 | down |
| WARS2 | RP11-128P10.1 | 0.906 | 0.002 | 1.000 | down |
| WARS2 | RP11-174G6.5 | 0.932 | 0.001 | 1.000 | down |
| WARS2 | RP11-299H21.1 | 0.902 | 0.002 | 1.000 | down |
| WARS2 | RP13-131K19.2 | 0.911 | 0.002 | 1.000 | down |
| WARS2 | FOXN3-AS1 | 0.929 | 0.001 | 1.000 | down |
| WARS2 | INAFM1 | 0.919 | 0.001 | 1.000 | down |
| WARS2 | GYG2P1 | 0.911 | 0.002 | 1.000 | down |
| WARS2 | FGF13-AS1 | 0.902 | 0.002 | 1.000 | down |
| WARS2 | CRYBB2P1 | 0.921 | 0.001 | 1.000 | down |
| WARS2 | G063514 | 0.944 | 0.000 | 1.000 | down |
| WARS2 | RP11-95P2.3 | 0.936 | 0.001 | 1.000 | down |
| WARS2 | G015529 | 0.929 | 0.001 | 1.000 | down |
| CKMT1A | ITGA9-AS1 | -0.920 | 0.001 | -1.000 | down |
| CKMT1A | RP11-3K16.2 | -0.936 | 0.001 | -1.000 | down |
| CKMT1A | AADACL2-AS1 | -0.915 | 0.001 | -1.000 | down |
| CKMT1A | G063659 | -0.908 | 0.002 | -1.000 | down |
| CKMT1A | CTD-2023M8.1 | -0.947 | 0.000 | -1.000 | down |
| CKMT1A | G030124 | -0.951 | 0.000 | -1.000 | down |
| CKMT1A | G010224 | -0.933 | 0.001 | -1.000 | down |
| CKMT1A | RP11-196G18.23 | 0.911 | 0.002 | 1.000 | down |
| CKMT1A | G042247 | 0.912 | 0.002 | 1.000 | down |
| CKMT1A | G085410 | 0.903 | 0.002 | 1.000 | down |
| CKMT1A | SUCLA2-AS1 | 0.915 | 0.001 | 1.000 | down |
| CKMT1A | G074033 | 0.901 | 0.002 | 1.000 | down |
| CKMT1A | G087757 | 0.907 | 0.002 | 1.000 | down |
| CKMT1A | G053697 | 0.903 | 0.002 | 1.000 | down |
| CKMT1A | PVRL3-AS1 | 0.911 | 0.002 | 1.000 | down |
| CKMT1A | TOB1-AS1 (1) | 0.929 | 0.001 | 1.000 | down |
| CKMT1A | CTD-2382E5.2 | 0.907 | 0.002 | 1.000 | down |
| CKMT1A | EZR-AS1 | 0.978 | 0.000 | 1.000 | down |
| CKMT1A | AK125078 | 0.930 | 0.001 | 1.000 | down |
| CKMT1A | ANKRD20A12P | 0.920 | 0.001 | 1.000 | down |
| CKMT1A | G014106 | 0.927 | 0.001 | 1.000 | down |
| CKMT1B | RP11-3K16.2 | -0.947 | 0.000 | -1.000 | down |
| CKMT1B | GSE61474_XLOC_020343 | -0.912 | 0.002 | -1.000 | down |
| CKMT1B | PHF2P2 | -0.904 | 0.002 | -1.000 | down |
| CKMT1B | AADACL2-AS1 | -0.905 | 0.002 | -1.000 | down |
| CKMT1B | RP11-669E14.6 | -0.911 | 0.002 | -1.000 | down |
| CKMT1B | G036156 | -0.916 | 0.001 | -1.000 | down |
| CKMT1B | CTD-2023M8.1 | -0.966 | 0.000 | -1.000 | down |
| CKMT1B | G029241 | -0.950 | 0.000 | -1.000 | down |
| CKMT1B | G010224 | -0.979 | 0.000 | -1.000 | down |
| CKMT1B | RP11-44N22.3 | 0.929 | 0.001 | 1.000 | down |
| CKMT1B | DLG5-AS1 (1) | 0.922 | 0.001 | 1.000 | down |
| CKMT1B | RP11-715J22.6 | 0.947 | 0.000 | 1.000 | down |
| CKMT1B | G079009 | 0.923 | 0.001 | 1.000 | down |
| CKMT1B | RP11-396C23.4 | 0.907 | 0.002 | 1.000 | down |
| CKMT1B | G048723 | 0.925 | 0.001 | 1.000 | down |
| CKMT1B | RP11-475I24.3 | 0.904 | 0.002 | 1.000 | down |
| CKMT1B | G052526 | 0.900 | 0.002 | 1.000 | down |
| CKMT1B | G053260 | 0.910 | 0.002 | 1.000 | down |
| CKMT1B | CCNT2-AS1 | 0.920 | 0.001 | 1.000 | down |
| CKMT1B | G053697 | 0.962 | 0.000 | 1.000 | down |
| CKMT1B | G010782 | 0.948 | 0.000 | 1.000 | down |
| CKMT1B | EZR-AS1 | 0.944 | 0.000 | 1.000 | down |
| CKMT1B | ANKRD20A12P | 0.922 | 0.001 | 1.000 | down |

| | | | | | |
|---|---|---|---|---|---|
| CKMT1B | KB-318B8.7 | 0.926 | 0.001 | 1.000 | down |
| CKMT1B | G014106 | 0.914 | 0.001 | 1.000 | down |
| CKMT1B | RP11-355F22.1 | 0.930 | 0.001 | 1.000 | down |
| BNIP3L | CTC-462L7.1 | -0.931 | 0.001 | -1.000 | down |
| BNIP3L | G066551 | -0.926 | 0.001 | -1.000 | down |
| BNIP3L | AK124233 | -0.949 | 0.000 | -1.000 | down |
| BNIP3L | AQP7P3 | -0.908 | 0.002 | -1.000 | down |
| BNIP3L | AX747064 | -0.967 | 0.000 | -1.000 | down |
| BNIP3L | LOC727896 | -0.948 | 0.000 | -1.000 | down |
| BNIP3L | G022034 | -0.902 | 0.002 | -1.000 | down |
| BNIP3L | G011388 | -0.933 | 0.001 | -1.000 | down |
| BNIP3L | G068469 | -0.949 | 0.000 | -1.000 | down |
| BNIP3L | AF086132 | -0.912 | 0.002 | -1.000 | down |
| BNIP3L | AK024936 | -0.914 | 0.001 | -1.000 | down |
| BNIP3L | G017670 | 0.969 | 0.000 | 1.000 | down |
| BNIP3L | AC073321.4 | 0.926 | 0.001 | 1.000 | down |
| BNIP3L | XLOC_003595 | 0.918 | 0.001 | 1.000 | down |
| BNIP3L | LINC00883 | 0.935 | 0.001 | 1.000 | down |
| BNIP3L | RP11-382D12.2 | 0.954 | 0.000 | 1.000 | down |
| BNIP3L | G063668 | 0.943 | 0.000 | 1.000 | down |
| BNIP3L | G058460 | 0.937 | 0.001 | 1.000 | down |
| PNPLA8 | LOC100499484-C9ORF174 | -0.929 | 0.001 | -1.000 | down |
| PNPLA8 | SLC7A11-AS1 | -0.929 | 0.001 | -1.000 | down |
| PNPLA8 | G011310 | -0.916 | 0.001 | -1.000 | down |
| PNPLA8 | RP11-466A19.1 | -0.937 | 0.001 | -1.000 | down |
| PNPLA8 | EML2-AS1 | -0.945 | 0.000 | -1.000 | down |
| PNPLA8 | G003431 | -0.902 | 0.002 | -1.000 | down |
| PNPLA8 | XLOC_012740 | -0.902 | 0.002 | -1.000 | down |
| PNPLA8 | G013465 | -0.952 | 0.000 | -1.000 | down |
| PNPLA8 | RP5-1112D6.7 | -0.926 | 0.001 | -1.000 | down |
| PNPLA8 | G029241 | -0.926 | 0.001 | -1.000 | down |
| PNPLA8 | EF565122 | -0.940 | 0.001 | -1.000 | down |
| PNPLA8 | G013179 | -0.919 | 0.001 | -1.000 | down |
| PNPLA8 | RP11-461K13.1 | -0.989 | 0.000 | -1.000 | down |
| PNPLA8 | RP11-602N24.3 | 0.902 | 0.002 | 1.000 | down |
| PNPLA8 | AL110181 | 0.982 | 0.000 | 1.000 | down |
| PNPLA8 | RP11-363E6.3 | 0.952 | 0.000 | 1.000 | down |
| PNPLA8 | AK055712 | 0.926 | 0.001 | 1.000 | down |
| PNPLA8 | RP11-303E16.7 | 0.976 | 0.000 | 1.000 | down |
| PNPLA8 | RP4-549L20.3 | 0.934 | 0.001 | 1.000 | down |
| PNPLA8 | RP11-686O6.2 | 0.979 | 0.000 | 1.000 | down |
| PNPLA8 | G061591 | 0.922 | 0.001 | 1.000 | down |
| PNPLA8 | RP11-356J5.12 | 0.971 | 0.000 | 1.000 | down |
| PNPLA8 | RP1-40E16.12 | 0.965 | 0.000 | 1.000 | down |
| PNPLA8 | GBAT2 | 0.948 | 0.000 | 1.000 | down |
| PNPLA8 | AL137474 | 0.936 | 0.001 | 1.000 | down |
| PNPLA8 | LINC00662 | 0.937 | 0.001 | 1.000 | down |
| PNPLA8 | RP11-179B2.2 | 0.918 | 0.001 | 1.000 | down |
| PNPLA8 | RP11-191L17.1 | 0.934 | 0.001 | 1.000 | down |
| PNPLA8 | AC099754.1 | 0.911 | 0.002 | 1.000 | down |
| PNPLA8 | RP4-683L5.1 | 0.941 | 0.001 | 1.000 | down |
| PNPLA8 | RP11-209D14.2 | 0.921 | 0.001 | 1.000 | down |
| PNPLA8 | AC002467.7 | 0.902 | 0.002 | 1.000 | down |
| PNPLA8 | GAS5-AS1 | 0.914 | 0.001 | 1.000 | down |
| PNPLA8 | AC144652.1 | 0.960 | 0.000 | 1.000 | down |
| PNPLA8 | G085410 | 0.908 | 0.002 | 1.000 | down |
| PNPLA8 | G089005 | 0.919 | 0.001 | 1.000 | down |
| PNPLA8 | BC070391 | 0.908 | 0.002 | 1.000 | down |
| PNPLA8 | MAPT-IT1 | 0.914 | 0.002 | 1.000 | down |
| PNPLA8 | G073576 | 0.952 | 0.000 | 1.000 | down |
| PNPLA8 | XLOC_013222 | 0.967 | 0.000 | 1.000 | down |
| PNPLA8 | G082866 | 0.961 | 0.000 | 1.000 | down |
| PNPLA8 | RP11-138E2.1 | 0.944 | 0.000 | 1.000 | down |
| PNPLA8 | G059716 | 0.938 | 0.001 | 1.000 | down |

| | | | | | |
|---|---|---|---|---|---|
| PNPLA8 | G044581 | 0.902 | 0.002 | 1.000 | down |
| PNPLA8 | BC042589 | 0.938 | 0.001 | 1.000 | down |
| PNPLA8 | LINC00634 | 0.967 | 0.000 | 1.000 | down |
| PNPLA8 | RP11-22N19.2 | 0.932 | 0.001 | 1.000 | down |
| PNPLA8 | RP4-555D20.3 | 0.930 | 0.001 | 1.000 | down |
| PNPLA8 | AP000255.6 | 0.912 | 0.002 | 1.000 | down |
| PNPLA8 | LL22NC03-86G7.1 | 0.938 | 0.001 | 1.000 | down |
| PNPLA8 | RP1-223E5.4 | 0.965 | 0.000 | 1.000 | down |
| PNPLA8 | FAM66D | 0.952 | 0.000 | 1.000 | down |
| PNPLA8 | RP11-573G6.8 | 0.923 | 0.001 | 1.000 | down |
| PNPLA8 | LL09NC01-139C3.1 | 0.935 | 0.001 | 1.000 | down |
| PNPLA8 | RP11-85A1.3 | 0.913 | 0.002 | 1.000 | down |
| PNPLA8 | RP11-344P13.6 | 0.921 | 0.001 | 1.000 | down |
| PNPLA8 | ANKRD20A12P | 0.925 | 0.001 | 1.000 | down |
| PNPLA8 | RP11-53I6.3 | 0.906 | 0.002 | 1.000 | down |
| PNPLA8 | CTD-2623N2.11 | 0.917 | 0.001 | 1.000 | down |
| PNPLA8 | G023340 | 0.944 | 0.000 | 1.000 | down |
| PNPLA8 | KB-318B8.7 | 0.907 | 0.002 | 1.000 | down |
| PNPLA8 | G006578 | 0.951 | 0.000 | 1.000 | down |
| PNPLA8 | G075319 | 0.952 | 0.000 | 1.000 | down |
| PNPLA8 | GSE61474_XLOC_067608 | 0.926 | 0.001 | 1.000 | down |
| PNPLA8 | RP11-79P5.9 | 0.936 | 0.001 | 1.000 | down |
| PNPLA8 | RP11-166B2.3 | 0.942 | 0.000 | 1.000 | down |
| PNPLA8 | RP11-439C15.4 | 0.959 | 0.000 | 1.000 | down |
| PNPLA8 | RP11-196G18.22 | 0.931 | 0.001 | 1.000 | down |
| IBA57 | XLOC_012303 | -0.952 | 0.000 | -1.000 | down |
| IBA57 | G029454 | -0.930 | 0.001 | -1.000 | down |
| IBA57 | G068163 | -0.921 | 0.001 | -1.000 | down |
| IBA57 | G080487 | -0.961 | 0.000 | -1.000 | down |
| IBA57 | XLOC_008025 | -0.924 | 0.001 | -1.000 | down |
| IBA57 | XLOC_l2_011118 | -0.901 | 0.002 | -1.000 | down |
| IBA57 | G072062 | -0.930 | 0.001 | -1.000 | down |
| IBA57 | RP11-717D12.1 | -0.969 | 0.000 | -1.000 | down |
| IBA57 | SNHG7 | -0.905 | 0.002 | -1.000 | down |
| IBA57 | G006360 | -0.966 | 0.000 | -1.000 | down |
| IBA57 | XLOC_013958 | -0.920 | 0.001 | -1.000 | down |
| IBA57 | G063524 | -0.910 | 0.002 | -1.000 | down |
| IBA57 | LINC01149 | -0.917 | 0.001 | -1.000 | down |
| IBA57 | G016657 | -0.967 | 0.000 | -1.000 | down |
| IBA57 | G067215 | -0.906 | 0.002 | -1.000 | down |
| IBA57 | G037029 | -0.961 | 0.000 | -1.000 | down |
| IBA57 | G046492 | -0.905 | 0.002 | -1.000 | down |
| IBA57 | RP11-551L14.6 | -0.916 | 0.001 | -1.000 | down |
| IBA57 | G072257 | -0.926 | 0.001 | -1.000 | down |
| IBA57 | G071259 | -0.910 | 0.002 | -1.000 | down |
| IBA57 | CTD-2302E22.4 | 0.914 | 0.001 | 1.000 | down |
| IBA57 | RP11-350N15.5 | 0.936 | 0.001 | 1.000 | down |
| IBA57 | RP4-549L20.3 | 0.915 | 0.001 | 1.000 | down |
| IBA57 | BRWD1-IT2 | 0.966 | 0.000 | 1.000 | down |
| IBA57 | RP11-490O6.2 | 0.969 | 0.000 | 1.000 | down |
| IBA57 | RP11-1017G21.6 | 0.965 | 0.000 | 1.000 | down |
| IBA57 | CTD-2588C8.8 | 0.909 | 0.002 | 1.000 | down |
| IBA57 | G047225 | 0.944 | 0.000 | 1.000 | down |
| IBA57 | RP1-309F20.3 | 0.952 | 0.000 | 1.000 | down |
| IBA57 | RP11-643M14.1 | 0.932 | 0.001 | 1.000 | down |
| IBA57 | XLOC_008875 | 0.919 | 0.001 | 1.000 | down |
| IBA57 | RP11-191L17.1 | 0.934 | 0.001 | 1.000 | down |
| IBA57 | RP11-165A20.3 | 0.967 | 0.000 | 1.000 | down |
| IBA57 | G000481 | 0.950 | 0.000 | 1.000 | down |
| IBA57 | RP11-285F16.1 | 0.938 | 0.001 | 1.000 | down |
| IBA57 | RP5-1092A3.4 | 0.904 | 0.002 | 1.000 | down |
| IBA57 | RP11-351D16.3 | 0.966 | 0.000 | 1.000 | down |
| IBA57 | AK123758 | 0.939 | 0.001 | 1.000 | down |
| IBA57 | RP4-683L5.1 | 0.934 | 0.001 | 1.000 | down |

| | | | | | |
|---|---|---|---|---|---|
| IBA57 | RP11-353K11.1 | 0.960 | 0.000 | 1.000 | down |
| IBA57 | HRAT5 | 0.929 | 0.001 | 1.000 | down |
| IBA57 | G015380 | 0.914 | 0.001 | 1.000 | down |
| IBA57 | XLOC_001189 | 0.920 | 0.001 | 1.000 | down |
| IBA57 | RP11-312B8.1 | 0.970 | 0.000 | 1.000 | down |
| IBA57 | GAS5-AS1 | 0.912 | 0.002 | 1.000 | down |
| IBA57 | RP3-410B11.1 | 0.946 | 0.000 | 1.000 | down |
| IBA57 | G089005 | 0.913 | 0.002 | 1.000 | down |
| IBA57 | LOC101929741 | 0.942 | 0.000 | 1.000 | down |
| IBA57 | G079654 | 0.948 | 0.000 | 1.000 | down |
| IBA57 | TEX26-AS1 | 0.914 | 0.001 | 1.000 | down |
| IBA57 | RP11-2E11.10 | 0.908 | 0.002 | 1.000 | down |
| IBA57 | FRMD6-AS1 | 0.959 | 0.000 | 1.000 | down |
| IBA57 | AL355711 | 0.971 | 0.000 | 1.000 | down |
| IBA57 | CTD-2256P15.2 | 0.980 | 0.000 | 1.000 | down |
| IBA57 | BC070391 | 0.942 | 0.000 | 1.000 | down |
| IBA57 | G034470 | 0.935 | 0.001 | 1.000 | down |
| IBA57 | G073576 | 0.917 | 0.001 | 1.000 | down |
| IBA57 | XLOC_011052 | 0.902 | 0.002 | 1.000 | down |
| IBA57 | RP1-202O8.3 | 0.955 | 0.000 | 1.000 | down |
| IBA57 | FOXN3-AS1 | 0.909 | 0.002 | 1.000 | down |
| IBA57 | GSE61474_XLOC_038226 | 0.920 | 0.001 | 1.000 | down |
| IBA57 | G038972 | 0.954 | 0.000 | 1.000 | down |
| IBA57 | LINC-PINT (2) | 0.942 | 0.000 | 1.000 | down |
| IBA57 | RP11-505K9.1 | 0.950 | 0.000 | 1.000 | down |
| IBA57 | RP11-138E2.1 | 0.919 | 0.001 | 1.000 | down |
| IBA57 | G003798 | 0.967 | 0.000 | 1.000 | down |
| IBA57 | AK026244 | 0.915 | 0.001 | 1.000 | down |
| IBA57 | LOC148709 | 0.924 | 0.001 | 1.000 | down |
| IBA57 | G044581 | 0.950 | 0.000 | 1.000 | down |
| IBA57 | RP5-884G6.2 | 0.936 | 0.001 | 1.000 | down |
| IBA57 | CTD-2383M3.1 | 0.931 | 0.001 | 1.000 | down |
| IBA57 | AC092667.2 | 0.934 | 0.001 | 1.000 | down |
| IBA57 | RP11-376P6.3 | 0.943 | 0.000 | 1.000 | down |
| IBA57 | RP11-383C5.5 | 0.985 | 0.000 | 1.000 | down |
| IBA57 | G001739 | 0.979 | 0.000 | 1.000 | down |
| IBA57 | RP11-22N19.2 | 0.929 | 0.001 | 1.000 | down |
| IBA57 | LOC101928402 | 0.916 | 0.001 | 1.000 | down |
| IBA57 | GFOD1 | 0.950 | 0.000 | 1.000 | down |
| IBA57 | LOC101928417 | 0.963 | 0.000 | 1.000 | down |
| IBA57 | RP4-555D20.3 | 0.929 | 0.001 | 1.000 | down |
| IBA57 | RP11-507K2.3 | 0.961 | 0.000 | 1.000 | down |
| IBA57 | RP4-798A10.7 | 0.914 | 0.002 | 1.000 | down |
| IBA57 | CTD-2377O17.1 | 0.966 | 0.000 | 1.000 | down |
| IBA57 | RP11-332H14.1 | 0.925 | 0.001 | 1.000 | down |
| IBA57 | RP1-261G23.7 | 0.948 | 0.000 | 1.000 | down |
| IBA57 | G023087 | 0.953 | 0.000 | 1.000 | down |
| IBA57 | G037470 | 0.926 | 0.001 | 1.000 | down |
| IBA57 | RP11-107N15.1 | 0.935 | 0.001 | 1.000 | down |
| IBA57 | LOC100288846 | 0.971 | 0.000 | 1.000 | down |
| IBA57 | RP1-86C11.7 | 0.915 | 0.001 | 1.000 | down |
| IBA57 | FLJ20021 | 0.923 | 0.001 | 1.000 | down |
| IBA57 | CTC-444N24.13 | 0.942 | 0.000 | 1.000 | down |
| IBA57 | LINC00449 | 0.902 | 0.002 | 1.000 | down |
| IBA57 | RP11-61L19.2 | 0.984 | 0.000 | 1.000 | down |
| IBA57 | LOC101929066 | 0.930 | 0.001 | 1.000 | down |
| IBA57 | RP11-85A1.3 | 0.953 | 0.000 | 1.000 | down |
| IBA57 | RP11-344P13.6 | 0.933 | 0.001 | 1.000 | down |
| IBA57 | CTD-2623N2.11 | 0.946 | 0.000 | 1.000 | down |
| IBA57 | RP1-232L24.3 | 0.920 | 0.001 | 1.000 | down |
| IBA57 | RP11-467P9.1 | 0.915 | 0.001 | 1.000 | down |
| IBA57 | G002695 | 0.904 | 0.002 | 1.000 | down |
| IBA57 | G021588 | 0.907 | 0.002 | 1.000 | down |
| IBA57 | RP11-371A19.2 | 0.954 | 0.000 | 1.000 | down |

| | | | | | |
|---|---|---|---|---|---|
| IBA57 | RP11-820I16.4 | 0.979 | 0.000 | 1.000 | down |
| IBA57 | G075703 | 0.901 | 0.002 | 1.000 | down |
| IBA57 | LINC01604 | 0.963 | 0.000 | 1.000 | down |
| IBA57 | TOB1-AS1 (2) | 0.922 | 0.001 | 1.000 | down |
| IBA57 | RP11-379F4.9 | 0.930 | 0.001 | 1.000 | down |
| NBR1 | XLOC_012303 | -0.927 | 0.001 | -1.000 | down |
| NBR1 | HCG18 | -0.908 | 0.002 | -1.000 | down |
| NBR1 | XLOC_l2_009332 | -0.909 | 0.002 | -1.000 | down |
| NBR1 | IGBP1-AS2 | -0.909 | 0.002 | -1.000 | down |
| NBR1 | G053606 | -0.926 | 0.001 | -1.000 | down |
| NBR1 | G046678 | -0.902 | 0.002 | -1.000 | down |
| NBR1 | AC016682.1 | -0.911 | 0.002 | -1.000 | down |
| NBR1 | G050702 | -0.901 | 0.002 | -1.000 | down |
| NBR1 | G070831 | -0.909 | 0.002 | -1.000 | down |
| NBR1 | AK093193 | -0.961 | 0.000 | -1.000 | down |
| NBR1 | LOC101927164 | -0.915 | 0.001 | -1.000 | down |
| NBR1 | CTD-2050E21.1 | -0.943 | 0.000 | -1.000 | down |
| NBR1 | SNHG7 | -0.934 | 0.001 | -1.000 | down |
| NBR1 | RP11-424I19.2 | -0.926 | 0.001 | -1.000 | down |
| NBR1 | XLOC_013958 | -0.964 | 0.000 | -1.000 | down |
| NBR1 | BX648501 | -0.941 | 0.000 | -1.000 | down |
| NBR1 | RP11-374P20.4 | -0.917 | 0.001 | -1.000 | down |
| NBR1 | G045736 | -0.945 | 0.000 | -1.000 | down |
| NBR1 | G016657 | -0.922 | 0.001 | -1.000 | down |
| NBR1 | AP000344.4 | -0.908 | 0.002 | -1.000 | down |
| NBR1 | RP11-30O15.1 | -0.940 | 0.001 | -1.000 | down |
| NBR1 | G067215 | -0.935 | 0.001 | -1.000 | down |
| NBR1 | G037029 | -0.921 | 0.001 | -1.000 | down |
| NBR1 | G046492 | -0.919 | 0.001 | -1.000 | down |
| NBR1 | uc.50 | -0.930 | 0.001 | -1.000 | down |
| NBR1 | XLOC_000048 | -0.912 | 0.002 | -1.000 | down |
| NBR1 | G071259 | -0.919 | 0.001 | -1.000 | down |
| NBR1 | G068890 | -0.906 | 0.002 | -1.000 | down |
| NBR1 | G059595 | -0.900 | 0.002 | -1.000 | down |
| NBR1 | FAM74A4 | 0.951 | 0.000 | 1.000 | down |
| NBR1 | RP11-255E6.6 | 0.928 | 0.001 | 1.000 | down |
| NBR1 | RP11-744I24.3 | 0.952 | 0.000 | 1.000 | down |
| NBR1 | GSE61474_XLOC_054139 | 0.966 | 0.000 | 1.000 | down |
| NBR1 | LOC100130417 | 0.944 | 0.000 | 1.000 | down |
| NBR1 | RP11-525A16.4 | 0.924 | 0.001 | 1.000 | down |
| NBR1 | XLOC_008005 | 0.916 | 0.001 | 1.000 | down |
| NBR1 | USP27X-AS1 | 0.955 | 0.000 | 1.000 | down |
| NBR1 | BRWD1-IT2 | 0.936 | 0.001 | 1.000 | down |
| NBR1 | RP11-490O6.2 | 0.931 | 0.001 | 1.000 | down |
| NBR1 | G002862 | 0.908 | 0.002 | 1.000 | down |
| NBR1 | PSMG3-AS1 (2) | 0.931 | 0.001 | 1.000 | down |
| NBR1 | CTD-2588C8.8 | 0.932 | 0.001 | 1.000 | down |
| NBR1 | RP1-309F20.3 | 0.915 | 0.001 | 1.000 | down |
| NBR1 | RP11-165A20.3 | 0.918 | 0.001 | 1.000 | down |
| NBR1 | RP11-285F16.1 | 0.909 | 0.002 | 1.000 | down |
| NBR1 | G050681 | 0.977 | 0.000 | 1.000 | down |
| NBR1 | AK123758 | 0.932 | 0.001 | 1.000 | down |
| NBR1 | RP11-312B8.1 | 0.909 | 0.002 | 1.000 | down |
| NBR1 | RP3-410B11.1 | 0.920 | 0.001 | 1.000 | down |
| NBR1 | AC093391.2 | 0.901 | 0.002 | 1.000 | down |
| NBR1 | LOC101929741 | 0.911 | 0.002 | 1.000 | down |
| NBR1 | G079654 | 0.927 | 0.001 | 1.000 | down |
| NBR1 | XLOC_006928 | 0.943 | 0.000 | 1.000 | down |
| NBR1 | FRMD6-AS1 | 0.909 | 0.002 | 1.000 | down |
| NBR1 | RP11-128P10.1 | 0.952 | 0.000 | 1.000 | down |
| NBR1 | RP11-174G6.5 | 0.951 | 0.000 | 1.000 | down |
| NBR1 | AL355711 | 0.942 | 0.000 | 1.000 | down |
| NBR1 | CTD-2256P15.2 | 0.929 | 0.001 | 1.000 | down |
| NBR1 | G081251 | 0.900 | 0.002 | 1.000 | down |

| | | | | | |
|---|---|---|---|---|---|
| NBR1 | RP11-424N24.2 | 0.968 | 0.000 | 1.000 | down |
| NBR1 | XLOC_011052 | 0.905 | 0.002 | 1.000 | down |
| NBR1 | RP11-502I4.3 | 0.920 | 0.001 | 1.000 | down |
| NBR1 | RP1-202O8.3 | 0.949 | 0.000 | 1.000 | down |
| NBR1 | RP13-131K19.2 | 0.940 | 0.001 | 1.000 | down |
| NBR1 | FOXN3-AS1 | 0.953 | 0.000 | 1.000 | down |
| NBR1 | RP11-290L1.2 | 0.935 | 0.001 | 1.000 | down |
| NBR1 | IDH1-AS1 | 0.959 | 0.000 | 1.000 | down |
| NBR1 | RP11-505K9.1 | 0.928 | 0.001 | 1.000 | down |
| NBR1 | RP11-792A8.4 | 0.908 | 0.002 | 1.000 | down |
| NBR1 | AK026244 | 0.901 | 0.002 | 1.000 | down |
| NBR1 | RP11-2E11.5 | 0.903 | 0.002 | 1.000 | down |
| NBR1 | LOC148709 | 0.925 | 0.001 | 1.000 | down |
| NBR1 | RP11-567G11.1 | 0.916 | 0.001 | 1.000 | down |
| NBR1 | GYG2P1 | 0.947 | 0.000 | 1.000 | down |
| NBR1 | RP11-376P6.3 | 0.903 | 0.002 | 1.000 | down |
| NBR1 | FGF13-AS1 | 0.949 | 0.000 | 1.000 | down |
| NBR1 | RP11-383C5.5 | 0.914 | 0.002 | 1.000 | down |
| NBR1 | LOC101928402 | 0.917 | 0.001 | 1.000 | down |
| NBR1 | RP4-555D20.3 | 0.911 | 0.002 | 1.000 | down |
| NBR1 | AP000255.6 | 0.927 | 0.001 | 1.000 | down |
| NBR1 | AK000953 | 0.908 | 0.002 | 1.000 | down |
| NBR1 | CTD-2377O17.1 | 0.940 | 0.001 | 1.000 | down |
| NBR1 | RP11-254F7.2 | 0.925 | 0.001 | 1.000 | down |
| NBR1 | RP11-332H14.1 | 0.919 | 0.001 | 1.000 | down |
| NBR1 | RP1-261G23.7 | 0.909 | 0.002 | 1.000 | down |
| NBR1 | G023087 | 0.932 | 0.001 | 1.000 | down |
| NBR1 | UBAC2-AS1 | 0.919 | 0.001 | 1.000 | down |
| NBR1 | RP11-53B2.6 | 0.916 | 0.001 | 1.000 | down |
| NBR1 | FLJ20021 | 0.912 | 0.002 | 1.000 | down |
| NBR1 | RP11-109D9.4 | 0.923 | 0.001 | 1.000 | down |
| NBR1 | BX248253 | 0.911 | 0.002 | 1.000 | down |
| NBR1 | RP11-53I6.3 | 0.931 | 0.001 | 1.000 | down |
| NBR1 | RP11-474O21.5 | 0.932 | 0.001 | 1.000 | down |
| NBR1 | G046528 | 0.922 | 0.001 | 1.000 | down |
| NBR1 | G063514 | 0.937 | 0.001 | 1.000 | down |
| NBR1 | RP1-232L24.3 | 0.913 | 0.002 | 1.000 | down |
| NBR1 | RP11-467P9.1 | 0.919 | 0.001 | 1.000 | down |
| NBR1 | G021588 | 0.934 | 0.001 | 1.000 | down |
| NBR1 | RP11-95P2.3 | 0.975 | 0.000 | 1.000 | down |
| NBR1 | DNAJC9-AS1 | 0.913 | 0.002 | 1.000 | down |
| NBR1 | G075703 | 0.925 | 0.001 | 1.000 | down |
| NBR1 | G015529 | 0.947 | 0.000 | 1.000 | down |
| NBR1 | CTD-2517O10.6 | 0.901 | 0.002 | 1.000 | down |
| NBR1 | G085202 | 0.902 | 0.002 | 1.000 | down |
| PDK1 | G061035 | -0.926 | 0.001 | -1.000 | down |
| PDK1 | G081487 | -0.903 | 0.002 | -1.000 | down |
| PDK1 | AP001625.6 | -0.917 | 0.001 | -1.000 | down |
| PDK1 | XLOC_003405 | -0.935 | 0.001 | -1.000 | down |
| PDK1 | XLOC_l2_011118 | -0.910 | 0.002 | -1.000 | down |
| PDK1 | G081302 | -0.940 | 0.001 | -1.000 | down |
| PDK1 | NBPF8 | -0.933 | 0.001 | -1.000 | down |
| PDK1 | LOC101927691 | -0.913 | 0.002 | -1.000 | down |
| PDK1 | LOC101926908 | -0.920 | 0.001 | -1.000 | down |
| PDK1 | RMRPP3 | -0.928 | 0.001 | -1.000 | down |
| PDK1 | RP1-65P5.3 | -0.939 | 0.001 | -1.000 | down |
| PDK1 | G009329 | -0.901 | 0.002 | -1.000 | down |
| PDK1 | LINC00685 | -0.955 | 0.000 | -1.000 | down |
| PDK1 | RN7SL176P | -0.907 | 0.002 | -1.000 | down |
| PDK1 | G010533 | -0.948 | 0.000 | -1.000 | down |
| PDK1 | RP11-255E6.6 | 0.922 | 0.001 | 1.000 | down |
| PDK1 | RPL23AP7 | 0.916 | 0.001 | 1.000 | down |
| PDK1 | G064655 | 0.912 | 0.002 | 1.000 | down |
| PDK1 | CTD-2005H7.1 | 0.941 | 0.000 | 1.000 | down |
| PDK1 | RP11-326A19.3 | 0.912 | 0.002 | 1.000 | down |

| | | | | | |
|---|---|---|---|---|---|
| PDK1 | G009018 | 0.900 | 0.002 | 1.000 | down |
| PDK1 | RP11-299H21.1 | 0.936 | 0.001 | 1.000 | down |
| PDK1 | LOC100379224 | 0.934 | 0.001 | 1.000 | down |
| PDK1 | AC005220.3 | 0.935 | 0.001 | 1.000 | down |
| PDK1 | GPR50-AS1 | 0.935 | 0.001 | 1.000 | down |
| PDK1 | CXADRP3 | 0.934 | 0.001 | 1.000 | down |
| PDK1 | G054532 | 0.960 | 0.000 | 1.000 | down |
| PDK1 | LOC101927811 | 0.958 | 0.000 | 1.000 | down |
| PDK1 | uc.412 | 0.901 | 0.002 | 1.000 | down |
| PDK1 | LOC283140 | 0.922 | 0.001 | 1.000 | down |
| MSRA | HCG18 | -0.924 | 0.001 | -1.000 | down |
| MSRA | G075352 | -0.907 | 0.002 | -1.000 | down |
| MSRA | G038095 | -0.915 | 0.001 | -1.000 | down |
| MSRA | G015131 | -0.954 | 0.000 | -1.000 | down |
| MSRA | Z69720.2 | -0.942 | 0.000 | -1.000 | down |
| MSRA | AC003682.16 | -0.909 | 0.002 | -1.000 | down |
| MSRA | G046027 | -0.920 | 0.001 | -1.000 | down |
| MSRA | G011388 | -0.921 | 0.001 | -1.000 | down |
| MSRA | G002461 | -0.929 | 0.001 | -1.000 | down |
| MSRA | AP000344.4 | -0.921 | 0.001 | -1.000 | down |
| MSRA | RP11-10J18.3 | -0.902 | 0.002 | -1.000 | down |
| MSRA | G020470 | -0.907 | 0.002 | -1.000 | down |
| MSRA | AF086132 | -0.946 | 0.000 | -1.000 | down |
| MSRA | G084746 | -0.904 | 0.002 | -1.000 | down |
| MSRA | RP11-277L2.3 | 0.957 | 0.000 | 1.000 | down |
| MSRA | WWC2-AS2 | 0.947 | 0.000 | 1.000 | down |
| MSRA | G013286 | 0.928 | 0.001 | 1.000 | down |
| MSRA | RP11-290L1.2 | 0.907 | 0.002 | 1.000 | down |
| MSRA | RP11-792A8.4 | 0.917 | 0.001 | 1.000 | down |
| MSRA | G063668 | 0.916 | 0.001 | 1.000 | down |
| OGG1 | G066551 | -0.920 | 0.001 | -1.000 | down |
| OGG1 | Alpha_250_dna | -0.901 | 0.002 | -1.000 | down |
| OGG1 | G060633 | -0.911 | 0.002 | -1.000 | down |
| OGG1 | G011310 | -0.921 | 0.001 | -1.000 | down |
| OGG1 | G008916 | -0.938 | 0.001 | -1.000 | down |
| OGG1 | RP11-737O24.3 | -0.946 | 0.000 | -1.000 | down |
| OGG1 | uc.49 | -0.910 | 0.002 | -1.000 | down |
| OGG1 | G073689 | -0.981 | 0.000 | -1.000 | down |
| OGG1 | G054530 | -0.974 | 0.000 | -1.000 | down |
| OGG1 | XLOC_012740 | -0.904 | 0.002 | -1.000 | down |
| OGG1 | G051586 | -0.942 | 0.000 | -1.000 | down |
| OGG1 | DLG3-AS1 | -0.935 | 0.001 | -1.000 | down |
| OGG1 | IFT74-AS1 | 0.912 | 0.002 | 1.000 | down |
| OGG1 | AL110181 | 0.919 | 0.001 | 1.000 | down |
| OGG1 | TIPARP-AS1 | 0.924 | 0.001 | 1.000 | down |
| OGG1 | CTD-2302E22.4 | 0.901 | 0.002 | 1.000 | down |
| OGG1 | LINC00652 | 0.985 | 0.000 | 1.000 | down |
| OGG1 | RP1-40E16.12 | 0.924 | 0.001 | 1.000 | down |
| OGG1 | GBAT2 | 0.950 | 0.000 | 1.000 | down |
| OGG1 | TM4SF1-AS1 | 0.940 | 0.001 | 1.000 | down |
| OGG1 | RP11-252A24.3 | 0.940 | 0.001 | 1.000 | down |
| OGG1 | RP11-191L17.1 | 0.922 | 0.001 | 1.000 | down |
| OGG1 | CTC-232P5.3 | 0.923 | 0.001 | 1.000 | down |
| OGG1 | MGC2889 | 0.931 | 0.001 | 1.000 | down |
| OGG1 | LINC00883 | 0.913 | 0.002 | 1.000 | down |
| OGG1 | RP11-204E9.1 | 0.950 | 0.000 | 1.000 | down |
| OGG1 | G085410 | 0.912 | 0.002 | 1.000 | down |
| OGG1 | DHRS4-AS1 (1) | 0.959 | 0.000 | 1.000 | down |
| OGG1 | G089005 | 0.913 | 0.002 | 1.000 | down |
| OGG1 | OLMALINC | 0.953 | 0.000 | 1.000 | down |
| OGG1 | CTD-2517M22.16 | 0.911 | 0.002 | 1.000 | down |
| OGG1 | XLOC_011052 | 0.945 | 0.000 | 1.000 | down |
| OGG1 | G023041 | 0.922 | 0.001 | 1.000 | down |
| OGG1 | RP11-775D22.3 | 0.915 | 0.001 | 1.000 | down |
| OGG1 | BC042589 | 0.942 | 0.000 | 1.000 | down |

| | | | | | |
|---|---|---|---|---|---|
| OGG1 | LINC00634 | 0.912 | 0.002 | 1.000 | down |
| OGG1 | AC005220.3 | 0.909 | 0.002 | 1.000 | down |
| OGG1 | RP11-51J9.5 | 0.936 | 0.001 | 1.000 | down |
| OGG1 | G078120 | 0.954 | 0.000 | 1.000 | down |
| OGG1 | RP11-254F7.2 | 0.959 | 0.000 | 1.000 | down |
| OGG1 | G066437 | 0.951 | 0.000 | 1.000 | down |
| OGG1 | ENO1-AS1 | 0.948 | 0.000 | 1.000 | down |
| OGG1 | DGCR5 | 0.977 | 0.000 | 1.000 | down |
| OGG1 | G027543 | 0.933 | 0.001 | 1.000 | down |
| OGG1 | RP11-573G6.8 | 0.917 | 0.001 | 1.000 | down |
| OGG1 | LL09NC01-139C3.1 | 0.944 | 0.000 | 1.000 | down |
| OGG1 | G068576 | 0.906 | 0.002 | 1.000 | down |
| OGG1 | CXADRP3 | 0.928 | 0.001 | 1.000 | down |
| OGG1 | G047565 | 0.910 | 0.002 | 1.000 | down |
| OGG1 | RP11-793H13.11 | 0.944 | 0.000 | 1.000 | down |
| OGG1 | RP11-109D9.4 | 0.903 | 0.002 | 1.000 | down |
| OGG1 | RP11-110G21.1 | 0.952 | 0.000 | 1.000 | down |
| OGG1 | uc.349 | 0.915 | 0.001 | 1.000 | down |
| OGG1 | RP11-314A20.5 | 0.900 | 0.002 | 1.000 | down |
| OGG1 | RP11-196G18.24 | 0.912 | 0.002 | 1.000 | down |
| OGG1 | RP11-467P9.1 | 0.905 | 0.002 | 1.000 | down |
| OGG1 | G021721 | 0.935 | 0.001 | 1.000 | down |
| OGG1 | BC047644 | 0.911 | 0.002 | 1.000 | down |
| OGG1 | G001497 | 0.968 | 0.000 | 1.000 | down |
| CMPK2 | XLOC_012303 | -0.911 | 0.002 | -1.000 | down |
| CMPK2 | XLOC_l2_009332 | -0.976 | 0.000 | -1.000 | down |
| CMPK2 | G087508 | -0.901 | 0.002 | -1.000 | down |
| CMPK2 | G080487 | -0.914 | 0.001 | -1.000 | down |
| CMPK2 | XLOC_013958 | -0.912 | 0.002 | -1.000 | down |
| CMPK2 | G063524 | -0.929 | 0.001 | -1.000 | down |
| CMPK2 | G045736 | -0.927 | 0.001 | -1.000 | down |
| CMPK2 | XLOC_006124 | -0.932 | 0.001 | -1.000 | down |
| CMPK2 | G067215 | -0.967 | 0.000 | -1.000 | down |
| CMPK2 | uc.50 | -0.929 | 0.001 | -1.000 | down |
| CMPK2 | G071259 | -0.926 | 0.001 | -1.000 | down |
| CMPK2 | G041587 | -0.952 | 0.000 | -1.000 | down |
| CMPK2 | XLOC_008005 | 0.948 | 0.000 | 1.000 | down |
| CMPK2 | G047225 | 0.902 | 0.002 | 1.000 | down |
| CMPK2 | RP1-309F20.3 | 0.949 | 0.000 | 1.000 | down |
| CMPK2 | AC002467.7 | 0.924 | 0.001 | 1.000 | down |
| CMPK2 | G015380 | 0.942 | 0.000 | 1.000 | down |
| CMPK2 | XLOC_006928 | 0.908 | 0.002 | 1.000 | down |
| CMPK2 | RP11-174G6.5 | 0.918 | 0.001 | 1.000 | down |
| CMPK2 | AL355711 | 0.914 | 0.001 | 1.000 | down |
| CMPK2 | RP11-383C5.5 | 0.908 | 0.002 | 1.000 | down |
| CMPK2 | AK000953 | 0.928 | 0.001 | 1.000 | down |
| CMPK2 | RP1-86C11.7 | 0.928 | 0.001 | 1.000 | down |
| CMPK2 | FLJ20021 | 0.949 | 0.000 | 1.000 | down |
| CMPK2 | RP11-474O21.5 | 0.923 | 0.001 | 1.000 | down |
| CMPK2 | GSE61474_XLOC_067608 | 0.957 | 0.000 | 1.000 | down |
| FOXO3 | HCG18 | -0.950 | 0.000 | -1.000 | down |
| FOXO3 | G038095 | -0.936 | 0.001 | -1.000 | down |
| FOXO3 | AC016682.1 | -0.923 | 0.001 | -1.000 | down |
| FOXO3 | G015131 | -0.941 | 0.001 | -1.000 | down |
| FOXO3 | RP11-424I19.2 | -0.909 | 0.002 | -1.000 | down |
| FOXO3 | G002461 | -0.907 | 0.002 | -1.000 | down |
| FOXO3 | AP000344.4 | -0.942 | 0.000 | -1.000 | down |
| FOXO3 | uc.50 | -0.938 | 0.001 | -1.000 | down |
| FOXO3 | AF086132 | -0.919 | 0.001 | -1.000 | down |
| FOXO3 | G068890 | -0.907 | 0.002 | -1.000 | down |
| FOXO3 | G084746 | -0.908 | 0.002 | -1.000 | down |
| FOXO3 | FAM74A4 | 0.949 | 0.000 | 1.000 | down |
| FOXO3 | GSE61474_XLOC_054139 | 0.930 | 0.001 | 1.000 | down |

| | | | | | |
|---|---|---|---|---|---|
| FOXO3 | XLOC_008005 | 0.914 | 0.001 | 1.000 | down |
| FOXO3 | RP11-277L2.3 | 0.946 | 0.000 | 1.000 | down |
| FOXO3 | WWC2-AS2 | 0.936 | 0.001 | 1.000 | down |
| FOXO3 | RP11-424N24.2 | 0.910 | 0.002 | 1.000 | down |
| FOXO3 | G013286 | 0.901 | 0.002 | 1.000 | down |
| FOXO3 | RP11-290L1.2 | 0.942 | 0.000 | 1.000 | down |
| FOXO3 | RP11-792A8.4 | 0.946 | 0.000 | 1.000 | down |
| FOXO3 | RP11-567G11.1 | 0.946 | 0.000 | 1.000 | down |
| FOXO3 | RP11-95P2.3 | 0.919 | 0.001 | 1.000 | down |
| SATB1 | RP11-308N19.1 | -0.928 | 0.001 | -1.000 | down |
| SATB1 | G076532 | -0.915 | 0.001 | -1.000 | down |
| SATB1 | G071732 | -0.924 | 0.001 | -1.000 | down |
| SATB1 | G032508 | -0.957 | 0.000 | -1.000 | down |
| SATB1 | SPTY2D1-AS1 | -0.929 | 0.001 | -1.000 | down |
| SATB1 | G051725 | 0.961 | 0.000 | 1.000 | down |
| SATB1 | G010185 | 0.908 | 0.002 | 1.000 | down |
| SATB1 | G081251 | 0.930 | 0.001 | 1.000 | down |
| SATB1 | uc.69 | 0.947 | 0.000 | 1.000 | down |
| SATB1 | RP11-118K6.3 | 0.961 | 0.000 | 1.000 | down |
| SATB1 | LINC00670 | 0.916 | 0.001 | 1.000 | down |

**Supplementary Table 11. Dysregulated lncRNA-inflammation mRNAs coexpressing network in anesthetics exposed organoids**

| LncRNAs (source genes) | Inflammatory mRNAs (target genes) | Correlation coefficient | P value | Positive/Negative Correlation Coefficient | Regulation |
|---|---|---|---|---|---|
| G037251 | NLRP2 | -0.943 | 0.000 | -1.000 | up |
| NADK2-AS1 | FKBP4 | -0.937 | 0.001 | -1.000 | up |
| G003271 | SLC26A6 | -0.908 | 0.002 | -1.000 | up |
| HCG18 | IGF2R | -0.917 | 0.001 | -1.000 | up |
| RP11-308N19.1 | CMKLR1 | -0.929 | 0.001 | -1.000 | up |
| G042193 | NLRP10 | -0.902 | 0.002 | -1.000 | up |
| G056910 | NLRP10 | -0.935 | 0.001 | -1.000 | up |
| G056910 | XBP1 | -0.923 | 0.001 | -1.000 | up |
| RP11-21M24.3 | C1QTNF9B | 0.937 | 0.001 | 1.000 | up |
| LINC00473 | ATP6V0A2 | -0.938 | 0.001 | -1.000 | up |
| RP11-674P19.2 | ATP6V0A2 | -0.975 | 0.000 | -1.000 | up |
| RP11-70P17.1 | ATP6V0A2 | -0.946 | 0.000 | -1.000 | up |
| G038095 | IGF2R | -0.930 | 0.001 | -1.000 | up |
| Alpha_250_dna | FKBP4 | -0.961 | 0.000 | -1.000 | up |
| Alpha_250_dna | ADAM15 | -0.906 | 0.002 | -1.000 | up |
| LOC102723439 | FKBP4 | -0.908 | 0.002 | -1.000 | up |
| G038394 | MALAT1 | -0.917 | 0.001 | -1.000 | up |
| G089550 | F13A1 | -0.971 | 0.000 | -1.000 | up |
| G089550 | GATA4 | -0.979 | 0.000 | -1.000 | up |
| G089550 | C1QTNF9B | 0.932 | 0.001 | 1.000 | up |
| G053736 | PON2 | -0.911 | 0.002 | -1.000 | up |
| G020731 | XBP1 | -0.948 | 0.000 | -1.000 | up |
| G020731 | SCYL1 | -0.947 | 0.000 | -1.000 | up |
| IGBP1-AS2 | NLRP2 | -0.907 | 0.002 | -1.000 | up |
| G011310 | ADAM15 | -0.933 | 0.001 | -1.000 | up |
| ASAP1-IT1 | FKBP4 | -0.953 | 0.000 | -1.000 | up |
| ASAP1-IT1 | ADAM15 | -0.927 | 0.001 | -1.000 | up |
| ASAP1-IT1 | NBR1 | 0.906 | 0.002 | 1.000 | up |
| AK124233 | RGS1 | -0.937 | 0.001 | -1.000 | up |
| RP11-3K16.2 | NLRP2 | -0.948 | 0.000 | -1.000 | up |
| RP11-3K16.2 | NBR1 | 0.904 | 0.002 | 1.000 | up |
| RP11-187C18.2 | C1QTNF9B | 0.916 | 0.001 | 1.000 | up |
| G061982 | SLC26A6 | -0.932 | 0.001 | -1.000 | up |
| G076011 | PON2 | -0.936 | 0.001 | -1.000 | up |
| G053606 | MALAT1 | -0.921 | 0.001 | -1.000 | up |
| XLOC_006150 | FKBP4 | -0.907 | 0.002 | -1.000 | up |
| G006738 | MALAT1 | -0.955 | 0.000 | -1.000 | up |
| G087508 | NBR1 | 0.917 | 0.001 | 1.000 | up |
| G068163 | MALAT1 | -0.917 | 0.001 | -1.000 | up |
| G068163 | PON2 | -0.920 | 0.001 | -1.000 | up |
| LACTB2-AS1 | NLRP2 | -0.922 | 0.001 | -1.000 | up |
| RP3-416H24.4 | PON2 | -0.973 | 0.000 | -1.000 | up |
| RP3-416H24.4 | UCP2 | 0.936 | 0.001 | 1.000 | up |
| G073552 | SCYL1 | -0.950 | 0.000 | -1.000 | up |
| G073552 | HSP90AB1 | -0.914 | 0.002 | -1.000 | up |
| G058275 | RGS1 | -0.969 | 0.000 | -1.000 | up |
| G058275 | MPO | 0.916 | 0.001 | 1.000 | up |
| RP4-761J14.9 | NELL1 | 0.906 | 0.002 | 1.000 | up |
| RP4-816N1.7 | ATP6V0A2 | -0.940 | 0.001 | -1.000 | up |
| AQP7P3 | ATP6V0A2 | -0.902 | 0.002 | -1.000 | up |
| RP5-1065P14.2 | RGS1 | -0.915 | 0.001 | -1.000 | up |
| G033744 | NLRP2 | -0.959 | 0.000 | -1.000 | up |
| G021277 | NELL1 | 0.936 | 0.001 | 1.000 | up |
| G021277 | UCP2 | 0.949 | 0.000 | 1.000 | up |
| AC074363.1 | NFAT5 | -0.902 | 0.002 | -1.000 | up |
| AC074363.1 | NLRP2 | -0.910 | 0.002 | -1.000 | up |
| G046678 | SCYL1 | -0.904 | 0.002 | -1.000 | up |
| G046678 | HSP90AB1 | -0.942 | 0.000 | -1.000 | up |

| | | | | | |
|---|---|---|---|---|---|
| G031832 | C1QTNF9B | 0.922 | 0.001 | 1.000 | up |
| G081487 | FKBP4 | -0.928 | 0.001 | -1.000 | up |
| G081487 | HSP90AB1 | -0.908 | 0.002 | -1.000 | up |
| G081487 | NBR1 | 0.921 | 0.001 | 1.000 | up |
| G055386 | MALAT1 | -0.950 | 0.000 | -1.000 | up |
| AC016682.1 | XBP1 | -0.954 | 0.000 | -1.000 | up |
| AC016682.1 | SCYL1 | -0.918 | 0.001 | -1.000 | up |
| AC016682.1 | HSP90AB1 | -0.902 | 0.002 | -1.000 | up |
| RP11-484O2.1 | XBP1 | -0.942 | 0.000 | -1.000 | up |
| RP11-484O2.1 | SCYL1 | -0.910 | 0.002 | -1.000 | up |
| RP11-484O2.1 | HSP90AB1 | -0.944 | 0.000 | -1.000 | up |
| G050702 | SCYL1 | -0.930 | 0.001 | -1.000 | up |
| G050702 | HSP90AB1 | -0.940 | 0.001 | -1.000 | up |
| PSG10P | MALAT1 | -0.959 | 0.000 | -1.000 | up |
| RP11-587H10.1 | GATA4 | -0.947 | 0.000 | -1.000 | up |
| RP11-587H10.1 | NLRP2 | -0.942 | 0.000 | -1.000 | up |
| G082315 | C1QTNF9B | 0.925 | 0.001 | 1.000 | up |
| RP11-737O24.3 | ADAM15 | -0.914 | 0.001 | -1.000 | up |
| RP11-737O24.3 | NLRP2 | -0.946 | 0.000 | -1.000 | up |
| XLOC_003405 | NBR1 | 0.945 | 0.000 | 1.000 | up |
| G015131 | IGF2R | -0.915 | 0.001 | -1.000 | up |
| G009235 | XBP1 | -0.946 | 0.000 | -1.000 | up |
| G009235 | SCYL1 | -0.909 | 0.002 | -1.000 | up |
| G063995 | SCYL1 | -0.939 | 0.001 | -1.000 | up |
| G063995 | HSP90AB1 | -0.922 | 0.001 | -1.000 | up |
| RP11-669E14.6 | ADAM15 | -0.910 | 0.002 | -1.000 | up |
| RP11-669E14.6 | NBR1 | 0.929 | 0.001 | 1.000 | up |
| NEAT1 | ATP6V0A2 | -0.911 | 0.002 | -1.000 | up |
| G012913 | NELL1 | 0.911 | 0.002 | 1.000 | up |
| G012913 | NBR1 | 0.914 | 0.002 | 1.000 | up |
| RN7SKP264 | RGS1 | -0.939 | 0.001 | -1.000 | up |
| RN7SKP264 | MPO | 0.910 | 0.002 | 1.000 | up |
| LINC00379 | MALAT1 | -0.916 | 0.001 | -1.000 | up |
| RP11-514F8.2 | SLC26A6 | -0.917 | 0.001 | -1.000 | up |
| RP11-514F8.2 | MALAT1 | -0.936 | 0.001 | -1.000 | up |
| PMCHL2 | ATP6V0A2 | -0.946 | 0.000 | -1.000 | up |
| RP11-219C20.3 | RGS1 | -0.922 | 0.001 | -1.000 | up |
| RP11-219C20.3 | MPO | 0.967 | 0.000 | 1.000 | up |
| RP11-219C20.3 | VCAM1 | 0.920 | 0.001 | 1.000 | up |
| G070831 | MALAT1 | -0.960 | 0.000 | -1.000 | up |
| ZFAT-AS1 | XBP1 | -0.959 | 0.000 | -1.000 | up |
| ZFAT-AS1 | SCYL1 | -0.976 | 0.000 | -1.000 | up |
| ZFAT-AS1 | HSP90AB1 | -0.978 | 0.000 | -1.000 | up |
| G040015 | NLRP2 | -0.928 | 0.001 | -1.000 | up |
| G024041 | MALAT1 | -0.951 | 0.000 | -1.000 | up |
| G024041 | PON2 | -0.912 | 0.002 | -1.000 | up |
| G087265 | SCYL1 | -0.917 | 0.001 | -1.000 | up |
| Z69720.2 | RGS1 | -0.919 | 0.001 | -1.000 | up |
| CTC-261N6.1 | NBR1 | 0.964 | 0.000 | 1.000 | up |
| BC034456 | RGS1 | -0.964 | 0.000 | -1.000 | up |
| BC034456 | MPO | 0.926 | 0.001 | 1.000 | up |
| XLOC_006182 | NELL1 | 0.934 | 0.001 | 1.000 | up |
| RP11-18D7.2 | MALAT1 | -0.911 | 0.002 | -1.000 | up |
| G071640 | NBR1 | 0.948 | 0.000 | 1.000 | up |
| G082861 | PON2 | -0.957 | 0.000 | -1.000 | up |
| G000948 | ATP6V0A2 | -0.959 | 0.000 | -1.000 | up |
| G014972 | NBR1 | 0.904 | 0.002 | 1.000 | up |
| LOC727896 | HSPA4 | -0.908 | 0.002 | -1.000 | up |
| CRAT37 | NFAT5 | -0.962 | 0.000 | -1.000 | up |
| SNHG7 | MALAT1 | -0.952 | 0.000 | -1.000 | up |
| G046027 | RGS1 | -0.933 | 0.001 | -1.000 | up |
| G022034 | NFAT5 | -0.903 | 0.002 | -1.000 | up |
| G062685 | HSPA4 | -0.976 | 0.000 | -1.000 | up |
| G062685 | SLC26A6 | -0.971 | 0.000 | -1.000 | up |
| G080055 | MALAT1 | -0.914 | 0.002 | -1.000 | up |

| | | | | | |
|---|---|---|---|---|---|
| uc.49 | ATP6V0A2 | -0.933 | 0.001 | -1.000 | up |
| uc.49 | ADAM15 | -0.952 | 0.000 | -1.000 | up |
| uc.49 | NLRP2 | -0.923 | 0.001 | -1.000 | up |
| XLOC_013958 | MALAT1 | -0.955 | 0.000 | -1.000 | up |
| XLOC_005405 | NFAT5 | -0.906 | 0.002 | -1.000 | up |
| G011388 | HSPA4 | -0.913 | 0.002 | -1.000 | up |
| XLOC_003837 | NBR1 | 0.924 | 0.001 | 1.000 | up |
| BX648501 | XBP1 | -0.935 | 0.001 | -1.000 | up |
| BX648501 | SCYL1 | -0.930 | 0.001 | -1.000 | up |
| G060010 | HSPA4 | -0.902 | 0.002 | -1.000 | up |
| G060010 | VCAM1 | 0.922 | 0.001 | 1.000 | up |
| G073689 | ADAM15 | -0.941 | 0.000 | -1.000 | up |
| G072364 | UCP2 | 0.936 | 0.001 | 1.000 | up |
| CYP2D7 | XBP1 | -0.905 | 0.002 | -1.000 | up |
| CYP2D7 | SCYL1 | -0.902 | 0.002 | -1.000 | up |
| RP11-374P20.4 | MALAT1 | -0.968 | 0.000 | -1.000 | up |
| G020793 | UCP2 | 0.951 | 0.000 | 1.000 | up |
| LINC01149 | MALAT1 | -0.918 | 0.001 | -1.000 | up |
| LINC01149 | NELL1 | 0.901 | 0.002 | 1.000 | up |
| LINC01149 | UCP2 | 0.927 | 0.001 | 1.000 | up |
| LOC101927285 | XBP1 | -0.919 | 0.001 | -1.000 | up |
| AK025321 | NFAT5 | -0.903 | 0.002 | -1.000 | up |
| G002461 | IGF2R | -0.960 | 0.000 | -1.000 | up |
| LOC101926908 | FKBP4 | -0.948 | 0.000 | -1.000 | up |
| G001795 | NLRP2 | -0.926 | 0.001 | -1.000 | up |
| G001795 | NELL1 | 0.973 | 0.000 | 1.000 | up |
| G001795 | UCP2 | 0.924 | 0.001 | 1.000 | up |
| G016657 | MALAT1 | -0.954 | 0.000 | -1.000 | up |
| XLOC_012740 | ADAM15 | -0.929 | 0.001 | -1.000 | up |
| XLOC_012740 | NBR1 | 0.920 | 0.001 | 1.000 | up |
| uc.474 | ATP6V0A2 | -0.912 | 0.002 | -1.000 | up |
| TMLHE-AS1 | XBP1 | -0.921 | 0.001 | -1.000 | up |
| TMLHE-AS1 | SCYL1 | -0.905 | 0.002 | -1.000 | up |
| TMLHE-AS1 | HSP90AB1 | -0.923 | 0.001 | -1.000 | up |
| G060484 | PON2 | -0.945 | 0.000 | -1.000 | up |
| G060484 | NELL1 | 0.917 | 0.001 | 1.000 | up |
| G060484 | UCP2 | 0.961 | 0.000 | 1.000 | up |
| RP11-45M22.5 | CMKLR1 | -0.905 | 0.002 | -1.000 | up |
| RP11-30O15.1 | MALAT1 | -0.967 | 0.000 | -1.000 | up |
| RP11-768G7.2 | F13A1 | -0.900 | 0.002 | -1.000 | up |
| G058409 | NELL1 | 0.901 | 0.002 | 1.000 | up |
| G009329 | XBP1 | -0.902 | 0.002 | -1.000 | up |
| LOC642846 | F13A1 | -0.976 | 0.000 | -1.000 | up |
| LOC642846 | GATA4 | -0.971 | 0.000 | -1.000 | up |
| RP11-10J18.3 | RGS1 | -0.940 | 0.001 | -1.000 | up |
| LINC00685 | NBR1 | 0.916 | 0.001 | 1.000 | up |
| G015181 | UCP2 | 0.935 | 0.001 | 1.000 | up |
| G082713 | MPO | 0.928 | 0.001 | 1.000 | up |
| G082713 | VCAM1 | 0.920 | 0.001 | 1.000 | up |
| G046492 | MALAT1 | -0.910 | 0.002 | -1.000 | up |
| G046492 | NELL1 | 0.909 | 0.002 | 1.000 | up |
| AC026188.1 | C1QTNF9B | 0.933 | 0.001 | 1.000 | up |
| RP4-760C5.3 | SCYL1 | -0.907 | 0.002 | -1.000 | up |
| CTD- | ADAM15 | -0.918 | 0.001 | -1.000 | up |
| CTD- | NBR1 | 0.915 | 0.001 | 1.000 | up |
| G003928 | FKBP4 | -0.930 | 0.001 | -1.000 | up |
| G072257 | MALAT1 | -0.925 | 0.001 | -1.000 | up |
| AF086132 | RGS1 | -0.938 | 0.001 | -1.000 | up |
| AF086132 | IGF2R | -0.906 | 0.002 | -1.000 | up |
| CTC-250I14.3 | NFAT5 | -0.968 | 0.000 | -1.000 | up |
| G051586 | ADAM15 | -0.904 | 0.002 | -1.000 | up |
| G051586 | NLRP2 | -0.916 | 0.001 | -1.000 | up |
| RP11-126O1.4 | FKBP4 | -0.958 | 0.000 | -1.000 | up |
| RP11-651L5.2 | NBR1 | 0.905 | 0.002 | 1.000 | up |

| | | | | | |
|---|---|---|---|---|---|
| GSE61474_XLOC_042934 | C1QTNF9B | 0.934 | 0.001 | 1.000 | up |
| AK025564 | MALAT1 | -0.935 | 0.001 | -1.000 | up |
| XLOC_007237 | NELL1 | 0.914 | 0.001 | 1.000 | up |
| XLOC_000048 | MALAT1 | -0.943 | 0.000 | -1.000 | up |
| LOC101926941 | ATP6V0A2 | -0.964 | 0.000 | -1.000 | up |
| G053358 | GATA4 | -0.902 | 0.002 | -1.000 | up |
| G053358 | C1QTNF9B | 0.947 | 0.000 | 1.000 | up |
| BCRP3 | XBP1 | -0.904 | 0.002 | -1.000 | up |
| BCRP3 | SCYL1 | -0.931 | 0.001 | -1.000 | up |
| BCRP3 | HSP90AB1 | -0.964 | 0.000 | -1.000 | up |
| G033096 | GATA4 | -0.910 | 0.002 | -1.000 | up |
| G033096 | NLRP2 | -0.949 | 0.000 | -1.000 | up |
| G089283 | ADAM15 | -0.963 | 0.000 | -1.000 | up |
| G089283 | NLRP2 | -0.936 | 0.001 | -1.000 | up |
| G089283 | NBR1 | 0.941 | 0.000 | 1.000 | up |
| XLOC_007868 | HSPA4 | -0.943 | 0.000 | -1.000 | up |
| AC025442.3 | ATP6V0A2 | -0.911 | 0.002 | -1.000 | up |
| G071259 | MALAT1 | -0.960 | 0.000 | -1.000 | up |
| RP5-1112D6.7 | ADAM15 | -0.917 | 0.001 | -1.000 | up |
| RP5-1112D6.7 | NBR1 | 0.942 | 0.000 | 1.000 | up |
| MINOS1P1 | C1QTNF9B | 0.945 | 0.000 | 1.000 | up |
| MIMT1 | NBR1 | 0.922 | 0.001 | 1.000 | up |
| G079644 | NELL1 | 0.943 | 0.000 | 1.000 | up |
| G068890 | XBP1 | -0.932 | 0.001 | -1.000 | up |
| G068890 | SCYL1 | -0.918 | 0.001 | -1.000 | up |
| G010941 | NFAT5 | -0.921 | 0.001 | -1.000 | up |
| G031271 | NBR1 | 0.901 | 0.002 | 1.000 | up |
| AK022848 | XBP1 | -0.932 | 0.001 | -1.000 | up |
| G051433 | HSP90AB1 | -0.901 | 0.002 | -1.000 | up |
| DLG3-AS1 | ADAM15 | -0.904 | 0.002 | -1.000 | up |
| DLG3-AS1 | NLRP2 | -0.917 | 0.001 | -1.000 | up |
| G088912 | RGS1 | -0.909 | 0.002 | -1.000 | up |
| RN7SL176P | HSP90AB1 | -0.913 | 0.002 | -1.000 | up |
| G074298 | NLRP2 | -0.909 | 0.002 | -1.000 | up |
| G074298 | NBR1 | 0.965 | 0.000 | 1.000 | up |
| G022519 | HSPA4 | -0.935 | 0.001 | -1.000 | up |
| G022519 | VCAM1 | 0.935 | 0.001 | 1.000 | up |
| AK024936 | ATP6V0A2 | -0.942 | 0.000 | -1.000 | up |
| XLOC_009429 | ADAM15 | -0.916 | 0.001 | -1.000 | up |
| G059595 | ADAM15 | -0.942 | 0.000 | -1.000 | up |
| G059595 | HSP90AB1 | -0.917 | 0.001 | -1.000 | up |
| G059595 | NBR1 | 0.947 | 0.000 | 1.000 | up |
| IFT74-AS1 | ADAM15 | 0.943 | 0.000 | 1.000 | down |
| FAM74A4 | IGF2R | 0.903 | 0.002 | 1.000 | down |
| AL110181 | ADAM15 | 0.930 | 0.001 | 1.000 | down |
| AL110181 | NBR1 | -0.918 | 0.001 | -1.000 | down |
| TIPARP-AS1 | ADAM15 | 0.903 | 0.002 | 1.000 | down |
| RP11-255E6.6 | NBR1 | -0.956 | 0.000 | -1.000 | down |
| XLOC_010525 | NFAT5 | 0.944 | 0.000 | 1.000 | down |
| RP11-744I24.3 | SCYL1 | 0.912 | 0.002 | 1.000 | down |
| XLOC_003724 | C1QTNF9B | -0.946 | 0.000 | -1.000 | down |
| RP11-363E6.3 | ADAM15 | 0.902 | 0.002 | 1.000 | down |
| RP11-363E6.3 | NBR1 | -0.908 | 0.002 | -1.000 | down |
| AK055712 | NBR1 | -0.901 | 0.002 | -1.000 | down |
| RP11-230C9.2 | SLC26A6 | 0.923 | 0.001 | 1.000 | down |
| RP11-230C9.2 | NELL1 | -0.924 | 0.001 | -1.000 | down |
| LOC100130417 | MALAT1 | 0.947 | 0.000 | 1.000 | down |
| RPL23AP7 | NBR1 | -0.982 | 0.000 | -1.000 | down |
| G090511 | F13A1 | 0.943 | 0.000 | 1.000 | down |
| G090511 | C1QTNF9B | -0.943 | 0.000 | -1.000 | down |
| RP11-44N22.3 | NLRP2 | 0.915 | 0.001 | 1.000 | down |
| OR2L1P | NELL1 | -0.905 | 0.002 | -1.000 | down |
| PSMG3-AS1 | NELL1 | -0.915 | 0.001 | -1.000 | down |
| GS1-433O24.1 | FKBP4 | 0.906 | 0.002 | 1.000 | down |

| | | | | | |
|---|---|---|---|---|---|
| LINC00652 | ADAM15 | 0.945 | 0.000 | 1.000 | down |
| G017718 | MPO | -0.951 | 0.000 | -1.000 | down |
| G017718 | VCAM1 | -0.931 | 0.001 | -1.000 | down |
| RP11-689C9.1 | HSPA4 | 0.983 | 0.000 | 1.000 | down |
| RP11-689C9.1 | SLC26A6 | 0.939 | 0.001 | 1.000 | down |
| G020528 | NFAT5 | 0.931 | 0.001 | 1.000 | down |
| RP11-762I7.4 | NFAT5 | 0.918 | 0.001 | 1.000 | down |
| RP11-624D11.2 | HSPA4 | 0.928 | 0.001 | 1.000 | down |
| FAM230C | NLRP2 | 0.922 | 0.001 | 1.000 | down |
| USP27X-AS1 | XBP1 | 0.918 | 0.001 | 1.000 | down |
| RP1-40E16.12 | ADAM15 | 0.915 | 0.001 | 1.000 | down |
| BRWD1-IT2 | MALAT1 | 0.955 | 0.000 | 1.000 | down |
| RP11-355B11.2 | NLRP2 | 0.903 | 0.002 | 1.000 | down |
| G021652 | FKBP4 | 0.945 | 0.000 | 1.000 | down |
| G047503 | UCP2 | -0.913 | 0.002 | -1.000 | down |
| G002862 | XBP1 | 0.985 | 0.000 | 1.000 | down |
| G002862 | SCYL1 | 0.975 | 0.000 | 1.000 | down |
| G002862 | HSP90AB1 | 0.911 | 0.002 | 1.000 | down |
| PSMG3-AS1 | SLC26A6 | 0.918 | 0.001 | 1.000 | down |
| DLG5-AS1 (1) | NLRP2 | 0.925 | 0.001 | 1.000 | down |
| XLOC_011336 | NELL1 | -0.930 | 0.001 | -1.000 | down |
| XLOC_011336 | UCP2 | -0.938 | 0.001 | -1.000 | down |
| G049752 | ADAM15 | 0.900 | 0.002 | 1.000 | down |
| G049752 | NELL1 | -0.901 | 0.002 | -1.000 | down |
| SH3GL1P2 | ADAM15 | 0.944 | 0.000 | 1.000 | down |
| LOC729083 | UCP2 | -0.904 | 0.002 | -1.000 | down |
| TM4SF1-AS1 | ADAM15 | 0.944 | 0.000 | 1.000 | down |
| TPTE2P3 | NLRP10 | 0.947 | 0.000 | 1.000 | down |
| CTD- | MALAT1 | 0.908 | 0.002 | 1.000 | down |
| CTD- | NELL1 | -0.955 | 0.000 | -1.000 | down |
| RP11-599B13.7 | SLC26A6 | 0.916 | 0.001 | 1.000 | down |
| RP11-599B13.7 | PON2 | 0.905 | 0.002 | 1.000 | down |
| LINC00662 | ADAM15 | 0.923 | 0.001 | 1.000 | down |
| LINC00662 | NBR1 | -0.947 | 0.000 | -1.000 | down |
| RP11-179B2.2 | NBR1 | -0.903 | 0.002 | -1.000 | down |
| G064655 | ADAM15 | 0.939 | 0.001 | 1.000 | down |
| G064655 | NBR1 | -0.963 | 0.000 | -1.000 | down |
| G017886 | MPO | -0.929 | 0.001 | -1.000 | down |
| G017886 | VCAM1 | -0.986 | 0.000 | -1.000 | down |
| RP11-182N22.8 | NLRP10 | 0.936 | 0.001 | 1.000 | down |
| RP11-182N22.8 | MALAT1 | 0.921 | 0.001 | 1.000 | down |
| RP11-182N22.8 | PON2 | 0.952 | 0.000 | 1.000 | down |
| CTD-2588C8.8 | MALAT1 | 0.926 | 0.001 | 1.000 | down |
| G057990 | F13A1 | 0.947 | 0.000 | 1.000 | down |
| RP11-643M14.1 | MALAT1 | 0.942 | 0.000 | 1.000 | down |
| RP11-344E13.4 | ATP6V0A2 | 0.965 | 0.000 | 1.000 | down |
| G017670 | RGS1 | 0.934 | 0.001 | 1.000 | down |
| G017670 | MPO | -0.901 | 0.002 | -1.000 | down |
| RP11-252A24.3 | ADAM15 | 0.907 | 0.002 | 1.000 | down |
| CTC-336P14.1 | NELL1 | -0.928 | 0.001 | -1.000 | down |
| RP11-46J23.1 | ATP6V0A2 | 0.903 | 0.002 | 1.000 | down |
| RP11-46J23.1 | NLRP2 | 0.915 | 0.001 | 1.000 | down |
| RP11-715J22.6 | ADAM15 | 0.931 | 0.001 | 1.000 | down |
| RP11-715J22.6 | NLRP2 | 0.952 | 0.000 | 1.000 | down |
| RP11-715J22.6 | NBR1 | -0.946 | 0.000 | -1.000 | down |
| G037472 | SLC26A6 | 0.966 | 0.000 | 1.000 | down |
| DLG5-AS1 (2) | SLC26A6 | 0.959 | 0.000 | 1.000 | down |
| CCDC144NL-AS1 | NLRP10 | 0.903 | 0.002 | 1.000 | down |
| WWC2-AS2 | IGF2R | 0.902 | 0.002 | 1.000 | down |
| AC099754.1 | ADAM15 | 0.936 | 0.001 | 1.000 | down |
| AC099754.1 | NBR1 | -0.965 | 0.000 | -1.000 | down |
| RP11-165A20.3 | MALAT1 | 0.918 | 0.001 | 1.000 | down |
| G079009 | ATP6V0A2 | 0.917 | 0.001 | 1.000 | down |
| G079009 | NLRP2 | 0.913 | 0.002 | 1.000 | down |

| | | | | | |
|---|---|---|---|---|---|
| RP11-396C23.4 | ATP6V0A2 | 0.919 | 0.001 | 1.000 | down |
| RP11-396C23.4 | ADAM15 | 0.918 | 0.001 | 1.000 | down |
| RP11-396C23.4 | NLRP2 | 0.933 | 0.001 | 1.000 | down |
| G000481 | MALAT1 | 0.964 | 0.000 | 1.000 | down |
| G050681 | MALAT1 | 0.927 | 0.001 | 1.000 | down |
| CTC-232P5.3 | ADAM15 | 0.938 | 0.001 | 1.000 | down |
| CTC-232P5.3 | NLRP2 | 0.909 | 0.002 | 1.000 | down |
| G047031 | ATP6V0A2 | 0.913 | 0.002 | 1.000 | down |
| RP5-991G20.4 | ADAM15 | 0.943 | 0.000 | 1.000 | down |
| RP5-991G20.4 | NBR1 | -0.933 | 0.001 | -1.000 | down |
| RP5-1092A3.4 | MALAT1 | 0.910 | 0.002 | 1.000 | down |
| BC112270 | HSPA4 | 0.917 | 0.001 | 1.000 | down |
| BC112270 | SLC26A6 | 0.953 | 0.000 | 1.000 | down |
| G014329 | ATP6V0A2 | 0.944 | 0.000 | 1.000 | down |
| MGC2889 | ADAM15 | 0.984 | 0.000 | 1.000 | down |
| MGC2889 | NBR1 | -0.926 | 0.001 | -1.000 | down |
| AK123758 | MALAT1 | 0.915 | 0.001 | 1.000 | down |
| G086828 | F13A1 | 0.960 | 0.000 | 1.000 | down |
| G086828 | GATA4 | 0.906 | 0.002 | 1.000 | down |
| G082153 | NBR1 | -0.927 | 0.001 | -1.000 | down |
| HRAT5 | MALAT1 | 0.960 | 0.000 | 1.000 | down |
| CTD-2005H7.1 | NBR1 | -0.960 | 0.000 | -1.000 | down |
| TAF1A-AS1 | NFAT5 | 0.904 | 0.002 | 1.000 | down |
| G066319 | NFAT5 | 0.940 | 0.001 | 1.000 | down |
| FAM66A | ATP6V0A2 | 0.922 | 0.001 | 1.000 | down |
| RP11-475I24.3 | ATP6V0A2 | 0.925 | 0.001 | 1.000 | down |
| AC006273.4 | CMKLR1 | 0.922 | 0.001 | 1.000 | down |
| CTB-113I20.2 | MALAT1 | 0.922 | 0.001 | 1.000 | down |
| RP5-968P14.2 | HSPA4 | 0.925 | 0.001 | 1.000 | down |
| RP5-968P14.2 | SLC26A6 | 0.958 | 0.000 | 1.000 | down |
| G085410 | NLRP2 | 0.906 | 0.002 | 1.000 | down |
| uc.446 | MPO | -0.909 | 0.002 | -1.000 | down |
| uc.446 | VCAM1 | -0.916 | 0.001 | -1.000 | down |
| DHRS4-AS1 (1) | SLC26A6 | 0.906 | 0.002 | 1.000 | down |
| RP3-410B11.1 | MALAT1 | 0.967 | 0.000 | 1.000 | down |
| G079888 | HSP90AB1 | 0.909 | 0.002 | 1.000 | down |
| G079888 | NBR1 | -0.919 | 0.001 | -1.000 | down |
| G019879 | SLC26A6 | 0.965 | 0.000 | 1.000 | down |
| OLMALINC | ATP6V0A2 | 0.908 | 0.002 | 1.000 | down |
| OLMALINC | ADAM15 | 0.930 | 0.001 | 1.000 | down |
| G000619 | MALAT1 | 0.979 | 0.000 | 1.000 | down |
| LOC101929741 | MALAT1 | 0.963 | 0.000 | 1.000 | down |
| RP11-510M2.2 | MALAT1 | 0.943 | 0.000 | 1.000 | down |
| RP11-510M2.2 | NELL1 | -0.919 | 0.001 | -1.000 | down |
| G029844 | FKBP4 | 0.961 | 0.000 | 1.000 | down |
| FRMD6-AS1 | MALAT1 | 0.914 | 0.001 | 1.000 | down |
| G052526 | NLRP2 | 0.957 | 0.000 | 1.000 | down |
| RP11-382D12.2 | HSPA4 | 0.907 | 0.002 | 1.000 | down |
| TCR-alpha | UCP2 | -0.919 | 0.001 | -1.000 | down |
| AL355711 | MALAT1 | 0.925 | 0.001 | 1.000 | down |
| CTD-2256P15.2 | MALAT1 | 0.924 | 0.001 | 1.000 | down |
| TUB-AS1 | XBP1 | 0.908 | 0.002 | 1.000 | down |
| TUB-AS1 | SCYL1 | 0.921 | 0.001 | 1.000 | down |
| SIX3-AS1 | PON2 | 0.926 | 0.001 | 1.000 | down |
| MAPT-IT1 | NBR1 | -0.906 | 0.002 | -1.000 | down |
| LINC01132 | NLRP10 | 0.946 | 0.000 | 1.000 | down |
| G041225 | NLRP2 | 0.905 | 0.002 | 1.000 | down |
| G045191 | CMKLR1 | 0.903 | 0.002 | 1.000 | down |
| LINC-PINT (1) | NLRP10 | 0.912 | 0.002 | 1.000 | down |
| G081251 | MALAT1 | 0.915 | 0.001 | 1.000 | down |
| G074033 | NLRP2 | 0.907 | 0.002 | 1.000 | down |
| G074033 | NBR1 | -0.906 | 0.002 | -1.000 | down |
| CTD- | SLC26A6 | 0.919 | 0.001 | 1.000 | down |
| TPI1P3 | C1QTNF9B | -0.926 | 0.001 | -1.000 | down |
| CCNT2-AS1 | ADAM15 | 0.954 | 0.000 | 1.000 | down |

| | | | | | |
|---|---|---|---|---|---|
| CCNT2-AS1 | NLRP2 | 0.948 | 0.000 | 1.000 | down |
| CCNT2-AS1 | NBR1 | -0.969 | 0.000 | -1.000 | down |
| XLOC_011052 | ADAM15 | 0.910 | 0.002 | 1.000 | down |
| RP11-502I4.3 | SLC26A6 | 0.913 | 0.002 | 1.000 | down |
| RP11-502I4.3 | MALAT1 | 0.909 | 0.002 | 1.000 | down |
| RP11-502I4.3 | NLRP2 | 0.902 | 0.002 | 1.000 | down |
| G023041 | FKBP4 | 0.929 | 0.001 | 1.000 | down |
| RP1-202O8.3 | MALAT1 | 0.941 | 0.000 | 1.000 | down |
| XLOC_000308 | MPO | -0.904 | 0.002 | -1.000 | down |
| XLOC_000308 | VCAM1 | -0.933 | 0.001 | -1.000 | down |
| G013286 | HSPA4 | 0.941 | 0.000 | 1.000 | down |
| G013286 | IGF2R | 0.960 | 0.000 | 1.000 | down |
| RP11-290L1.2 | SLC26A6 | 0.907 | 0.002 | 1.000 | down |
| RP11-290L1.2 | IGF2R | 0.901 | 0.002 | 1.000 | down |
| INAFM1 | XBP1 | 0.911 | 0.002 | 1.000 | down |
| G019632 | MALAT1 | 0.904 | 0.002 | 1.000 | down |
| G019632 | NELL1 | -0.946 | 0.000 | -1.000 | down |
| G019632 | UCP2 | -0.904 | 0.002 | -1.000 | down |
| NIPBL-AS1 | ATP6V0A2 | 0.956 | 0.000 | 1.000 | down |
| NIPBL-AS1 | ADAM15 | 0.917 | 0.001 | 1.000 | down |
| KB-1592A4.14 | MPO | -0.919 | 0.001 | -1.000 | down |
| KB-1592A4.14 | VCAM1 | -0.901 | 0.002 | -1.000 | down |
| LOC100506083 | SLC26A6 | 0.913 | 0.002 | 1.000 | down |
| RP11-792A8.4 | HSPA4 | 0.907 | 0.002 | 1.000 | down |
| RP11-792A8.4 | IGF2R | 0.937 | 0.001 | 1.000 | down |
| G063668 | HSPA4 | 0.905 | 0.002 | 1.000 | down |
| G063668 | RGS1 | 0.918 | 0.001 | 1.000 | down |
| G063668 | MPO | -0.928 | 0.001 | -1.000 | down |
| G063668 | VCAM1 | -0.923 | 0.001 | -1.000 | down |
| RP11-775D22.3 | ADAM15 | 0.948 | 0.000 | 1.000 | down |
| RP11-775D22.3 | NBR1 | -0.902 | 0.002 | -1.000 | down |
| G066606 | ADAM15 | 0.907 | 0.002 | 1.000 | down |
| AK026244 | MALAT1 | 0.913 | 0.002 | 1.000 | down |
| RP11-2E11.5 | MALAT1 | 0.951 | 0.000 | 1.000 | down |
| RP11-2E11.5 | NELL1 | -0.918 | 0.001 | -1.000 | down |
| RP11-567G11.1 | IGF2R | 0.909 | 0.002 | 1.000 | down |
| RP5-884G6.2 | MALAT1 | 0.916 | 0.001 | 1.000 | down |
| RP5-884G6.2 | PON2 | 0.931 | 0.001 | 1.000 | down |
| PVRL3-AS1 | NBR1 | -0.923 | 0.001 | -1.000 | down |
| G010782 | NLRP2 | 0.906 | 0.002 | 1.000 | down |
| AC092667.2 | MALAT1 | 0.914 | 0.001 | 1.000 | down |
| GYG2P1 | NLRP10 | 0.917 | 0.001 | 1.000 | down |
| GYG2P1 | XBP1 | 0.910 | 0.002 | 1.000 | down |
| RP11-376P6.3 | MALAT1 | 0.915 | 0.001 | 1.000 | down |
| RP1-97J1.2 | PON2 | 0.903 | 0.002 | 1.000 | down |
| LOC102724661 | F13A1 | 0.923 | 0.001 | 1.000 | down |
| LINC01023 | MALAT1 | 0.928 | 0.001 | 1.000 | down |
| uc.95 | HSPA4 | 0.947 | 0.000 | 1.000 | down |
| uc.95 | SLC26A6 | 0.905 | 0.002 | 1.000 | down |
| LOC101928402 | MALAT1 | 0.920 | 0.001 | 1.000 | down |
| LOC101928417 | MALAT1 | 0.938 | 0.001 | 1.000 | down |
| G078120 | ADAM15 | 0.968 | 0.000 | 1.000 | down |
| CTD- | MALAT1 | 0.959 | 0.000 | 1.000 | down |
| G049105 | HSPA4 | 0.900 | 0.002 | 1.000 | down |
| RP11-254F7.2 | ADAM15 | 0.926 | 0.001 | 1.000 | down |
| PRR36 | XBP1 | 0.940 | 0.001 | 1.000 | down |
| PRR36 | SCYL1 | 0.936 | 0.001 | 1.000 | down |
| PRR36 | HSP90AB1 | 0.955 | 0.000 | 1.000 | down |
| RP11-332H14.1 | MALAT1 | 0.949 | 0.000 | 1.000 | down |
| G066437 | ADAM15 | 0.937 | 0.001 | 1.000 | down |
| G057325 | VCAM1 | -0.924 | 0.001 | -1.000 | down |
| ENO1-AS1 | ADAM15 | 0.910 | 0.002 | 1.000 | down |
| ENO1-AS1 | NLRP2 | 0.937 | 0.001 | 1.000 | down |
| RP1-261G23.7 | MALAT1 | 0.953 | 0.000 | 1.000 | down |
| G023087 | MALAT1 | 0.937 | 0.001 | 1.000 | down |

| | | | | | |
|---|---|---|---|---|---|
| RP11- | NBR1 | -0.900 | 0.002 | -1.000 | down |
| LINC01152 | F13A1 | 0.909 | 0.002 | 1.000 | down |
| RP11-680F8.1 | NFAT5 | 0.902 | 0.002 | 1.000 | down |
| DGCR5 | ATP6V0A2 | 0.902 | 0.002 | 1.000 | down |
| DGCR5 | ADAM15 | 0.930 | 0.001 | 1.000 | down |
| RP11-413M3.4 | ATP6V0A2 | 0.907 | 0.002 | 1.000 | down |
| LINC00463 | ADAM15 | 0.911 | 0.002 | 1.000 | down |
| LINC00463 | GATA4 | 0.931 | 0.001 | 1.000 | down |
| G027543 | ADAM15 | 0.927 | 0.001 | 1.000 | down |
| G027543 | NLRP2 | 0.911 | 0.002 | 1.000 | down |
| G027543 | VCAM1 | -0.938 | 0.001 | -1.000 | down |
| RP11-107N15.1 | MALAT1 | 0.904 | 0.002 | 1.000 | down |
| LL09NC01-139C3.1 | ADAM15 | 0.906 | 0.002 | 1.000 | down |
| G061722 | MALAT1 | 0.932 | 0.001 | 1.000 | down |
| G061722 | NLRP2 | 0.913 | 0.002 | 1.000 | down |
| G046707 | NLRP10 | 0.949 | 0.000 | 1.000 | down |
| G068576 | F13A1 | 0.921 | 0.001 | 1.000 | down |
| G068576 | ATP6V0A2 | 0.901 | 0.002 | 1.000 | down |
| G068576 | ADAM15 | 0.945 | 0.000 | 1.000 | down |
| G068576 | GATA4 | 0.939 | 0.001 | 1.000 | down |
| G068576 | NLRP2 | 0.918 | 0.001 | 1.000 | down |
| RP11-872D17.4 | HSPA4 | 0.909 | 0.002 | 1.000 | down |
| XLOC_004632 | NLRP2 | 0.910 | 0.002 | 1.000 | down |
| XLOC_004632 | NBR1 | -0.906 | 0.002 | -1.000 | down |
| CXADRP3 | ADAM15 | 0.940 | 0.001 | 1.000 | down |
| CXADRP3 | NBR1 | -0.917 | 0.001 | -1.000 | down |
| UBAC2-AS1 | SLC26A6 | 0.900 | 0.002 | 1.000 | down |
| UBAC2-AS1 | MALAT1 | 0.937 | 0.001 | 1.000 | down |
| UBAC2-AS1 | NELL1 | -0.920 | 0.001 | -1.000 | down |
| RP11-53B2.6 | NLRP10 | 0.906 | 0.002 | 1.000 | down |
| RP11-53B2.6 | MALAT1 | 0.904 | 0.002 | 1.000 | down |
| RP11-53B2.6 | PON2 | 0.907 | 0.002 | 1.000 | down |
| RP11-53B2.6 | NELL1 | -0.922 | 0.001 | -1.000 | down |
| XLOC_005289 | SLC26A6 | 0.929 | 0.001 | 1.000 | down |
| DDX39B-AS1 | NELL1 | -0.958 | 0.000 | -1.000 | down |
| RP11-177G23.2 | NLRP10 | 0.901 | 0.002 | 1.000 | down |
| RP11-177G23.2 | MALAT1 | 0.905 | 0.002 | 1.000 | down |
| G000672 | SLC26A6 | 0.961 | 0.000 | 1.000 | down |
| EZR-AS1 | NBR1 | -0.904 | 0.002 | -1.000 | down |
| G047565 | ADAM15 | 0.906 | 0.002 | 1.000 | down |
| G064611 | F13A1 | 0.902 | 0.002 | 1.000 | down |
| RP11-110G21.1 | ADAM15 | 0.976 | 0.000 | 1.000 | down |
| RP11-110G21.1 | GATA4 | 0.906 | 0.002 | 1.000 | down |
| RP11-110G21.1 | NLRP2 | 0.939 | 0.001 | 1.000 | down |
| LINC00449 | NELL1 | -0.921 | 0.001 | -1.000 | down |
| LINC00449 | UCP2 | -0.940 | 0.001 | -1.000 | down |
| G054532 | FKBP4 | 0.929 | 0.001 | 1.000 | down |
| G061596 | NBR1 | -0.924 | 0.001 | -1.000 | down |
| DHRS4-AS1 (2) | SLC26A6 | 0.909 | 0.002 | 1.000 | down |
| DHRS4-AS1 (2) | PON2 | 0.929 | 0.001 | 1.000 | down |
| DHRS4-AS1 (2) | UCP2 | -0.947 | 0.000 | -1.000 | down |
| BX248253 | ADAM15 | 0.915 | 0.001 | 1.000 | down |
| BX248253 | NLRP2 | 0.909 | 0.002 | 1.000 | down |
| BX248253 | NBR1 | -0.957 | 0.000 | -1.000 | down |
| uc.349 | ADAM15 | 0.942 | 0.000 | 1.000 | down |
| uc.349 | NLRP2 | 0.951 | 0.000 | 1.000 | down |
| uc.349 | NELL1 | -0.921 | 0.001 | -1.000 | down |
| CTC-359M8.1 | FKBP4 | 0.935 | 0.001 | 1.000 | down |
| RP11-565F19.2 | NELL1 | -0.912 | 0.002 | -1.000 | down |
| RP11-565F19.2 | NBR1 | -0.901 | 0.002 | -1.000 | down |
| G003603 | ADAM15 | 0.940 | 0.001 | 1.000 | down |
| G003603 | NLRP2 | 0.916 | 0.001 | 1.000 | down |
| G003603 | NBR1 | -0.965 | 0.000 | -1.000 | down |
| RP11-53I6.3 | ADAM15 | 0.920 | 0.001 | 1.000 | down |

| | | | | | |
|---|---|---|---|---|---|
| RP11-53I6.3 | NBR1 | -0.944 | 0.000 | -1.000 | down |
| RP11-483I13.5 | NFAT5 | 0.924 | 0.001 | 1.000 | down |
| RP11-314A20.5 | ADAM15 | 0.977 | 0.000 | 1.000 | down |
| RP11-314A20.5 | NBR1 | -0.942 | 0.000 | -1.000 | down |
| AK055974 | NBR1 | -0.933 | 0.001 | -1.000 | down |
| G060549 | MALAT1 | 0.956 | 0.000 | 1.000 | down |
| CRYBB2P1 | XBP1 | 0.949 | 0.000 | 1.000 | down |
| CRYBB2P1 | SCYL1 | 0.952 | 0.000 | 1.000 | down |
| CRYBB2P1 | HSP90AB1 | 0.969 | 0.000 | 1.000 | down |
| RP11- | ADAM15 | 0.930 | 0.001 | 1.000 | down |
| RP11- | NBR1 | -0.909 | 0.002 | -1.000 | down |
| LOC101927811 | ADAM15 | 0.922 | 0.001 | 1.000 | down |
| LOC101927811 | NBR1 | -0.940 | 0.001 | -1.000 | down |
| uc.412 | NBR1 | -0.935 | 0.001 | -1.000 | down |
| G000727 | NBR1 | -0.936 | 0.001 | -1.000 | down |
| H1FX-AS1 | XBP1 | 0.914 | 0.001 | 1.000 | down |
| GSE61474_XLOC_052497 | SLC26A6 | 0.917 | 0.001 | 1.000 | down |
| GSE61474_XLOC_052497 | MALAT1 | 0.926 | 0.001 | 1.000 | down |
| G063514 | HSP90AB1 | 0.925 | 0.001 | 1.000 | down |
| RP6-109B7.4 | PON2 | 0.924 | 0.001 | 1.000 | down |
| RP6-109B7.4 | UCP2 | -0.936 | 0.001 | -1.000 | down |
| RP11-30L3.2 | ATP6V0A2 | 0.914 | 0.001 | 1.000 | down |
| RP11-30L3.2 | ADAM15 | 0.916 | 0.001 | 1.000 | down |
| G006089 | GATA4 | 0.935 | 0.001 | 1.000 | down |
| G014106 | ADAM15 | 0.907 | 0.002 | 1.000 | down |
| G014106 | NLRP2 | 0.918 | 0.001 | 1.000 | down |
| RP11-362L22.1 | F13A1 | 0.909 | 0.002 | 1.000 | down |
| RP11-362L22.1 | GATA4 | 0.905 | 0.002 | 1.000 | down |
| XXbac-B476C20.9 | NELL1 | -0.920 | 0.001 | -1.000 | down |
| RP11-895M11.3 | MALAT1 | 0.939 | 0.001 | 1.000 | down |
| RP11-355F22.1 | ADAM15 | 0.916 | 0.001 | 1.000 | down |
| RP11-355F22.1 | NLRP2 | 0.919 | 0.001 | 1.000 | down |
| JARID2-AS1 | HSP90AB1 | 0.949 | 0.000 | 1.000 | down |
| XLOC_010219 | F13A1 | 0.913 | 0.002 | 1.000 | down |
| RP11-330O11.3 | NBR1 | -0.937 | 0.001 | -1.000 | down |
| LOC283140 | ADAM15 | 0.928 | 0.001 | 1.000 | down |
| LOC283140 | NBR1 | -0.924 | 0.001 | -1.000 | down |
| DNAJC9-AS1 | MALAT1 | 0.925 | 0.001 | 1.000 | down |
| DNAJC9-AS1 | NLRP2 | 0.929 | 0.001 | 1.000 | down |
| G065507 | UCP2 | -0.901 | 0.002 | -1.000 | down |
| G021721 | HSPA4 | 0.914 | 0.002 | 1.000 | down |
| CTB-181F24.1 | NBR1 | -0.966 | 0.000 | -1.000 | down |
| RP11-168O10.6 | XBP1 | 0.935 | 0.001 | 1.000 | down |
| RP11-168O10.6 | SCYL1 | 0.969 | 0.000 | 1.000 | down |
| RP11-168O10.6 | HSP90AB1 | 0.918 | 0.001 | 1.000 | down |
| G075703 | MALAT1 | 0.911 | 0.002 | 1.000 | down |
| LINC00858 | MALAT1 | 0.950 | 0.000 | 1.000 | down |
| BC047644 | ADAM15 | 0.932 | 0.001 | 1.000 | down |
| RN7SL832P | NFAT5 | 0.935 | 0.001 | 1.000 | down |
| RP11-820L6.1 | MALAT1 | 0.911 | 0.002 | 1.000 | down |
| RP11-820L6.1 | NLRP2 | 0.902 | 0.002 | 1.000 | down |
| LOC102723385 | ADAM15 | 0.948 | 0.000 | 1.000 | down |
| LOC102723385 | NLRP2 | 0.954 | 0.000 | 1.000 | down |
| LOC102723385 | NELL1 | -0.918 | 0.001 | -1.000 | down |
| LOC102723385 | NBR1 | -0.977 | 0.000 | -1.000 | down |
| RP11-803B1.8 | NBR1 | -0.908 | 0.002 | -1.000 | down |
| G005619 | RGS1 | 0.905 | 0.002 | 1.000 | down |
| G063721 | ADAM15 | 0.921 | 0.001 | 1.000 | down |
| G063721 | NLRP2 | 0.901 | 0.002 | 1.000 | down |
| G063721 | NBR1 | -0.943 | 0.000 | -1.000 | down |
| G015529 | NBR1 | -0.910 | 0.002 | -1.000 | down |
| CTD- | MALAT1 | 0.943 | 0.000 | 1.000 | down |

| | | | | | |
|---|---|---|---|---|---|
| G001497 | ADAM15 | 0.962 | 0.000 | 1.000 | down |
| GLIDR | ATP6V0A2 | 0.968 | 0.000 | 1.000 | down |
| TOB1-AS1 (2) | MALAT1 | 0.940 | 0.001 | 1.000 | down |
| TOB1-AS1 (2) | NELL1 | -0.904 | 0.002 | -1.000 | down |
| G007298 | HSPA4 | 0.944 | 0.000 | 1.000 | down |
| G007298 | F13A1 | 0.935 | 0.001 | 1.000 | down |
| G007298 | SLC26A6 | 0.902 | 0.002 | 1.000 | down |
| LOC101927358 | NBR1 | -0.923 | 0.001 | -1.000 | down |
| CTC-248O19.1 | ADAM15 | 0.936 | 0.001 | 1.000 | down |
| CTC-248O19.1 | NLRP2 | 0.908 | 0.002 | 1.000 | down |
| HSPA4 | LOC727896 | -0.908 | 0.002 | -1.000 | up |
| HSPA4 | G062685 | -0.976 | 0.000 | -1.000 | up |
| HSPA4 | G011388 | -0.913 | 0.002 | -1.000 | up |
| HSPA4 | G060010 | -0.902 | 0.002 | -1.000 | up |
| HSPA4 | XLOC_007868 | -0.943 | 0.000 | -1.000 | up |
| HSPA4 | G022519 | -0.935 | 0.001 | -1.000 | up |
| HSPA4 | RP11-689C9.1 | 0.983 | 0.000 | 1.000 | up |
| HSPA4 | RP11-624D11.2 | 0.928 | 0.001 | 1.000 | up |
| HSPA4 | BC112270 | 0.917 | 0.001 | 1.000 | up |
| HSPA4 | RP5-968P14.2 | 0.925 | 0.001 | 1.000 | up |
| HSPA4 | RP11-382D12.2 | 0.907 | 0.002 | 1.000 | up |
| HSPA4 | G013286 | 0.941 | 0.000 | 1.000 | up |
| HSPA4 | RP11-792A8.4 | 0.907 | 0.002 | 1.000 | up |
| HSPA4 | G063668 | 0.905 | 0.002 | 1.000 | up |
| HSPA4 | uc.95 | 0.947 | 0.000 | 1.000 | up |
| HSPA4 | G049105 | 0.900 | 0.002 | 1.000 | up |
| HSPA4 | RP11-872D17.4 | 0.909 | 0.002 | 1.000 | up |
| HSPA4 | G021721 | 0.914 | 0.002 | 1.000 | up |
| HSPA4 | G007298 | 0.944 | 0.000 | 1.000 | up |
| F13A1 | G089550 | -0.971 | 0.000 | -1.000 | up |
| F13A1 | RP11-768G7.2 | -0.900 | 0.002 | -1.000 | up |
| F13A1 | LOC642846 | -0.976 | 0.000 | -1.000 | up |
| F13A1 | G090511 | 0.943 | 0.000 | 1.000 | up |
| F13A1 | G057990 | 0.947 | 0.000 | 1.000 | up |
| F13A1 | G086828 | 0.960 | 0.000 | 1.000 | up |
| F13A1 | LOC102724661 | 0.923 | 0.001 | 1.000 | up |
| F13A1 | LINC01152 | 0.909 | 0.002 | 1.000 | up |
| F13A1 | G068576 | 0.921 | 0.001 | 1.000 | up |
| F13A1 | G064611 | 0.902 | 0.002 | 1.000 | up |
| F13A1 | RP11-362L22.1 | 0.909 | 0.002 | 1.000 | up |
| F13A1 | XLOC_010219 | 0.913 | 0.002 | 1.000 | up |
| F13A1 | G007298 | 0.935 | 0.001 | 1.000 | up |
| RGS1 | AK124233 | -0.937 | 0.001 | -1.000 | up |
| RGS1 | G058275 | -0.969 | 0.000 | -1.000 | up |
| RGS1 | RP5-1065P14.2 | -0.915 | 0.001 | -1.000 | up |
| RGS1 | RN7SKP264 | -0.939 | 0.001 | -1.000 | up |
| RGS1 | RP11-219C20.3 | -0.922 | 0.001 | -1.000 | up |
| RGS1 | Z69720.2 | -0.919 | 0.001 | -1.000 | up |
| RGS1 | BC034456 | -0.964 | 0.000 | -1.000 | up |
| RGS1 | G046027 | -0.933 | 0.001 | -1.000 | up |
| RGS1 | RP11-10J18.3 | -0.940 | 0.001 | -1.000 | up |
| RGS1 | AF086132 | -0.938 | 0.001 | -1.000 | up |
| RGS1 | G088912 | -0.909 | 0.002 | -1.000 | up |
| RGS1 | G017670 | 0.934 | 0.001 | 1.000 | up |
| RGS1 | G063668 | 0.918 | 0.001 | 1.000 | up |
| RGS1 | G005619 | 0.905 | 0.002 | 1.000 | up |
| NLRP10 | G042193 | -0.902 | 0.002 | -1.000 | up |
| NLRP10 | G056910 | -0.935 | 0.001 | -1.000 | up |
| NLRP10 | TPTE2P3 | 0.947 | 0.000 | 1.000 | up |
| NLRP10 | RP11-182N22.8 | 0.936 | 0.001 | 1.000 | up |
| NLRP10 | CCDC144NL-AS1 | 0.903 | 0.002 | 1.000 | up |
| NLRP10 | LINC01132 | 0.946 | 0.000 | 1.000 | up |
| NLRP10 | LINC-PINT (1) | 0.912 | 0.002 | 1.000 | up |
| NLRP10 | GYG2P1 | 0.917 | 0.001 | 1.000 | up |
| NLRP10 | G046707 | 0.949 | 0.000 | 1.000 | up |

| | | | | | |
|---|---|---|---|---|---|
| NLRP10 | RP11-53B2.6 | 0.906 | 0.002 | 1.000 | up |
| NLRP10 | RP11-177G23.2 | 0.901 | 0.002 | 1.000 | up |
| SLC26A6 | G003271 | -0.908 | 0.002 | -1.000 | up |
| SLC26A6 | G061982 | -0.932 | 0.001 | -1.000 | up |
| SLC26A6 | RP11-514F8.2 | -0.917 | 0.001 | -1.000 | up |
| SLC26A6 | G062685 | -0.971 | 0.000 | -1.000 | up |
| SLC26A6 | RP11-230C9.2 | 0.923 | 0.001 | 1.000 | up |
| SLC26A6 | RP11-689C9.1 | 0.939 | 0.001 | 1.000 | up |
| SLC26A6 | PSMG3-AS1 (2) | 0.918 | 0.001 | 1.000 | up |
| SLC26A6 | RP11-599B13.7 | 0.916 | 0.001 | 1.000 | up |
| SLC26A6 | G037472 | 0.966 | 0.000 | 1.000 | up |
| SLC26A6 | DLG5-AS1 (2) | 0.959 | 0.000 | 1.000 | up |
| SLC26A6 | BC112270 | 0.953 | 0.000 | 1.000 | up |
| SLC26A6 | RP5-968P14.2 | 0.958 | 0.000 | 1.000 | up |
| SLC26A6 | DHRS4-AS1 (1) | 0.906 | 0.002 | 1.000 | up |
| SLC26A6 | G019879 | 0.965 | 0.000 | 1.000 | up |
| SLC26A6 | CTD-2517M22.16 | 0.919 | 0.001 | 1.000 | up |
| SLC26A6 | RP11-502I4.3 | 0.913 | 0.002 | 1.000 | up |
| SLC26A6 | RP11-290L1.2 | 0.907 | 0.002 | 1.000 | up |
| SLC26A6 | LOC100506083 | 0.913 | 0.002 | 1.000 | up |
| SLC26A6 | uc.95 | 0.905 | 0.002 | 1.000 | up |
| SLC26A6 | UBAC2-AS1 | 0.900 | 0.002 | 1.000 | up |
| SLC26A6 | XLOC_005289 | 0.929 | 0.001 | 1.000 | up |
| SLC26A6 | G000672 | 0.961 | 0.000 | 1.000 | up |
| SLC26A6 | DHRS4-AS1 (2) | 0.909 | 0.002 | 1.000 | up |
| SLC26A6 | GSE61474_XLOC_052497 | 0.917 | 0.001 | 1.000 | up |
| SLC26A6 | G007298 | 0.902 | 0.002 | 1.000 | up |
| XBP1 | G056910 | -0.923 | 0.001 | -1.000 | up |
| XBP1 | G020731 | -0.948 | 0.000 | -1.000 | up |
| XBP1 | AC016682.1 | -0.954 | 0.000 | -1.000 | up |
| XBP1 | RP11-484O2.1 | -0.942 | 0.000 | -1.000 | up |
| XBP1 | G009235 | -0.946 | 0.000 | -1.000 | up |
| XBP1 | ZFAT-AS1 | -0.959 | 0.000 | -1.000 | up |
| XBP1 | BX648501 | -0.935 | 0.001 | -1.000 | up |
| XBP1 | CYP2D7 | -0.905 | 0.002 | -1.000 | up |
| XBP1 | LOC101927285 | -0.919 | 0.001 | -1.000 | up |
| XBP1 | TMLHE-AS1 | -0.921 | 0.001 | -1.000 | up |
| XBP1 | G009329 | -0.902 | 0.002 | -1.000 | up |
| XBP1 | BCRP3 | -0.904 | 0.002 | -1.000 | up |
| XBP1 | G068890 | -0.932 | 0.001 | -1.000 | up |
| XBP1 | AK022848 | -0.932 | 0.001 | -1.000 | up |
| XBP1 | USP27X-AS1 | 0.918 | 0.001 | 1.000 | up |
| XBP1 | G002862 | 0.985 | 0.000 | 1.000 | up |
| XBP1 | TUB-AS1 | 0.908 | 0.002 | 1.000 | up |
| XBP1 | INAFM1 | 0.911 | 0.002 | 1.000 | up |
| XBP1 | GYG2P1 | 0.910 | 0.002 | 1.000 | up |
| XBP1 | PRR36 | 0.940 | 0.001 | 1.000 | up |
| XBP1 | CRYBB2P1 | 0.949 | 0.000 | 1.000 | up |
| XBP1 | H1FX-AS1 | 0.914 | 0.001 | 1.000 | up |
| XBP1 | RP11-168O10.6 | 0.935 | 0.001 | 1.000 | up |
| MALAT1 | G038394 | -0.917 | 0.001 | -1.000 | up |
| MALAT1 | G053606 | -0.921 | 0.001 | -1.000 | up |
| MALAT1 | G006738 | -0.955 | 0.000 | -1.000 | up |
| MALAT1 | G068163 | -0.917 | 0.001 | -1.000 | up |
| MALAT1 | G055386 | -0.950 | 0.000 | -1.000 | up |
| MALAT1 | PSG10P | -0.959 | 0.000 | -1.000 | up |
| MALAT1 | LINC00379 | -0.916 | 0.001 | -1.000 | up |
| MALAT1 | RP11-514F8.2 | -0.936 | 0.001 | -1.000 | up |
| MALAT1 | G070831 | -0.960 | 0.000 | -1.000 | up |
| MALAT1 | G024041 | -0.951 | 0.000 | -1.000 | up |
| MALAT1 | RP11-18D7.2 | -0.911 | 0.002 | -1.000 | up |
| MALAT1 | SNHG7 | -0.952 | 0.000 | -1.000 | up |
| MALAT1 | G080055 | -0.914 | 0.002 | -1.000 | up |
| MALAT1 | XLOC_013958 | -0.955 | 0.000 | -1.000 | up |

| | | | | | |
|---|---|---|---|---|---|
| MALAT1 | RP11-374P20.4 | -0.968 | 0.000 | -1.000 | up |
| MALAT1 | LINC01149 | -0.918 | 0.001 | -1.000 | up |
| MALAT1 | G016657 | -0.954 | 0.000 | -1.000 | up |
| MALAT1 | RP11-30O15.1 | -0.967 | 0.000 | -1.000 | up |
| MALAT1 | G046492 | -0.910 | 0.002 | -1.000 | up |
| MALAT1 | G072257 | -0.925 | 0.001 | -1.000 | up |
| MALAT1 | AK025564 | -0.935 | 0.001 | -1.000 | up |
| MALAT1 | XLOC_000048 | -0.943 | 0.000 | -1.000 | up |
| MALAT1 | G071259 | -0.960 | 0.000 | -1.000 | up |
| MALAT1 | LOC100130417 | 0.947 | 0.000 | 1.000 | up |
| MALAT1 | BRWD1-IT2 | 0.955 | 0.000 | 1.000 | up |
| MALAT1 | CTD-2033A16.3 | 0.908 | 0.002 | 1.000 | up |
| MALAT1 | RP11-182N22.8 | 0.921 | 0.001 | 1.000 | up |
| MALAT1 | CTD-2588C8.8 | 0.926 | 0.001 | 1.000 | up |
| MALAT1 | RP11-643M14.1 | 0.942 | 0.000 | 1.000 | up |
| MALAT1 | RP11-165A20.3 | 0.918 | 0.001 | 1.000 | up |
| MALAT1 | G000481 | 0.964 | 0.000 | 1.000 | up |
| MALAT1 | G050681 | 0.927 | 0.001 | 1.000 | up |
| MALAT1 | RP5-1092A3.4 | 0.910 | 0.002 | 1.000 | up |
| MALAT1 | AK123758 | 0.915 | 0.001 | 1.000 | up |
| MALAT1 | HRAT5 | 0.960 | 0.000 | 1.000 | up |
| MALAT1 | CTB-113I20.2 | 0.922 | 0.001 | 1.000 | up |
| MALAT1 | RP3-410B11.1 | 0.967 | 0.000 | 1.000 | up |
| MALAT1 | G000619 | 0.979 | 0.000 | 1.000 | up |
| MALAT1 | LOC101929741 | 0.963 | 0.000 | 1.000 | up |
| MALAT1 | RP11-510M2.2 | 0.943 | 0.000 | 1.000 | up |
| MALAT1 | FRMD6-AS1 | 0.914 | 0.001 | 1.000 | up |
| MALAT1 | AL355711 | 0.925 | 0.001 | 1.000 | up |
| MALAT1 | CTD-2256P15.2 | 0.924 | 0.001 | 1.000 | up |
| MALAT1 | G081251 | 0.915 | 0.001 | 1.000 | up |
| MALAT1 | RP11-502I4.3 | 0.909 | 0.002 | 1.000 | up |
| MALAT1 | RP1-202O8.3 | 0.941 | 0.000 | 1.000 | up |
| MALAT1 | G019632 | 0.904 | 0.002 | 1.000 | up |
| MALAT1 | AK026244 | 0.913 | 0.002 | 1.000 | up |
| MALAT1 | RP11-2E11.5 | 0.951 | 0.000 | 1.000 | up |
| MALAT1 | RP5-884G6.2 | 0.916 | 0.001 | 1.000 | up |
| MALAT1 | AC092667.2 | 0.914 | 0.001 | 1.000 | up |
| MALAT1 | RP11-376P6.3 | 0.915 | 0.001 | 1.000 | up |
| MALAT1 | LINC01023 | 0.928 | 0.001 | 1.000 | up |
| MALAT1 | LOC101928402 | 0.920 | 0.001 | 1.000 | up |
| MALAT1 | LOC101928417 | 0.938 | 0.001 | 1.000 | up |
| MALAT1 | CTD-2377O17.1 | 0.959 | 0.000 | 1.000 | up |
| MALAT1 | RP11-332H14.1 | 0.949 | 0.000 | 1.000 | up |
| MALAT1 | RP1-261G23.7 | 0.953 | 0.000 | 1.000 | up |
| MALAT1 | G023087 | 0.937 | 0.001 | 1.000 | up |
| MALAT1 | RP11-107N15.1 | 0.904 | 0.002 | 1.000 | up |
| MALAT1 | G061722 | 0.932 | 0.001 | 1.000 | up |
| MALAT1 | UBAC2-AS1 | 0.937 | 0.001 | 1.000 | up |
| MALAT1 | RP11-53B2.6 | 0.904 | 0.002 | 1.000 | up |
| MALAT1 | RP11-177G23.2 | 0.905 | 0.002 | 1.000 | up |
| MALAT1 | G060549 | 0.956 | 0.000 | 1.000 | up |
| MALAT1 | GSE61474_XLOC_052497 | 0.926 | 0.001 | 1.000 | up |
| MALAT1 | RP11-895M11.3 | 0.939 | 0.001 | 1.000 | up |
| MALAT1 | DNAJC9-AS1 | 0.925 | 0.001 | 1.000 | up |
| MALAT1 | G075703 | 0.911 | 0.002 | 1.000 | up |
| MALAT1 | LINC00858 | 0.950 | 0.000 | 1.000 | up |
| MALAT1 | RP11-820L6.1 | 0.911 | 0.002 | 1.000 | up |
| MALAT1 | CTD-2517O10.6 | 0.943 | 0.000 | 1.000 | up |
| MALAT1 | TOB1-AS1 (2) | 0.940 | 0.001 | 1.000 | up |
| FKBP4 | NADK2-AS1 | -0.937 | 0.001 | -1.000 | up |
| FKBP4 | Alpha_250_dna | -0.961 | 0.000 | -1.000 | up |
| FKBP4 | LOC102723439 | -0.908 | 0.002 | -1.000 | up |
| FKBP4 | ASAP1-IT1 | -0.953 | 0.000 | -1.000 | up |
| FKBP4 | XLOC_006150 | -0.907 | 0.002 | -1.000 | up |

| | | | | | |
|---|---|---|---|---|---|
| FKBP4 | G081487 | -0.928 | 0.001 | -1.000 | up |
| FKBP4 | LOC101926908 | -0.948 | 0.000 | -1.000 | up |
| FKBP4 | G003928 | -0.930 | 0.001 | -1.000 | up |
| FKBP4 | RP11-126O1.4 | -0.958 | 0.000 | -1.000 | up |
| FKBP4 | GS1-433O24.1 | 0.906 | 0.002 | 1.000 | up |
| FKBP4 | G021652 | 0.945 | 0.000 | 1.000 | up |
| FKBP4 | G029844 | 0.961 | 0.000 | 1.000 | up |
| FKBP4 | G023041 | 0.929 | 0.001 | 1.000 | up |
| FKBP4 | G054532 | 0.929 | 0.001 | 1.000 | up |
| FKBP4 | CTC-359M8.1 | 0.935 | 0.001 | 1.000 | up |
| ATP6V0A2 | LINC00473 | -0.938 | 0.001 | -1.000 | up |
| ATP6V0A2 | RP11-674P19.2 | -0.975 | 0.000 | -1.000 | up |
| ATP6V0A2 | RP11-70P17.1 | -0.946 | 0.000 | -1.000 | up |
| ATP6V0A2 | RP4-816N1.7 | -0.940 | 0.001 | -1.000 | up |
| ATP6V0A2 | AQP7P3 | -0.902 | 0.002 | -1.000 | up |
| ATP6V0A2 | NEAT1 | -0.911 | 0.002 | -1.000 | up |
| ATP6V0A2 | PMCHL2 | -0.946 | 0.000 | -1.000 | up |
| ATP6V0A2 | G000948 | -0.959 | 0.000 | -1.000 | up |
| ATP6V0A2 | uc.49 | -0.933 | 0.001 | -1.000 | up |
| ATP6V0A2 | uc.474 | -0.912 | 0.002 | -1.000 | up |
| ATP6V0A2 | LOC101926941 | -0.964 | 0.000 | -1.000 | up |
| ATP6V0A2 | AC025442.3 | -0.911 | 0.002 | -1.000 | up |
| ATP6V0A2 | AK024936 | -0.942 | 0.000 | -1.000 | up |
| ATP6V0A2 | RP11-344E13.4 | 0.965 | 0.000 | 1.000 | up |
| ATP6V0A2 | RP11-46J23.1 | 0.903 | 0.002 | 1.000 | up |
| ATP6V0A2 | G079009 | 0.917 | 0.001 | 1.000 | up |
| ATP6V0A2 | RP11-396C23.4 | 0.919 | 0.001 | 1.000 | up |
| ATP6V0A2 | G047031 | 0.913 | 0.002 | 1.000 | up |
| ATP6V0A2 | G014329 | 0.944 | 0.000 | 1.000 | up |
| ATP6V0A2 | FAM66A | 0.922 | 0.001 | 1.000 | up |
| ATP6V0A2 | RP11-475I24.3 | 0.925 | 0.001 | 1.000 | up |
| ATP6V0A2 | OLMALINC | 0.908 | 0.002 | 1.000 | up |
| ATP6V0A2 | NIPBL-AS1 | 0.956 | 0.000 | 1.000 | up |
| ATP6V0A2 | DGCR5 | 0.902 | 0.002 | 1.000 | up |
| ATP6V0A2 | RP11-413M3.4 | 0.907 | 0.002 | 1.000 | up |
| ATP6V0A2 | G068576 | 0.901 | 0.002 | 1.000 | up |
| ATP6V0A2 | RP11-30L3.2 | 0.914 | 0.001 | 1.000 | up |
| ATP6V0A2 | GLIDR | 0.968 | 0.000 | 1.000 | up |
| CMKLR1 | RP11-308N19.1 | -0.929 | 0.001 | -1.000 | up |
| CMKLR1 | RP11-45M22.5 | -0.905 | 0.002 | -1.000 | up |
| CMKLR1 | AC006273.4 | 0.922 | 0.001 | 1.000 | up |
| CMKLR1 | G045191 | 0.903 | 0.002 | 1.000 | up |
| SCYL1 | G020731 | -0.947 | 0.000 | -1.000 | up |
| SCYL1 | G073552 | -0.950 | 0.000 | -1.000 | up |
| SCYL1 | G046678 | -0.904 | 0.002 | -1.000 | up |
| SCYL1 | AC016682.1 | -0.918 | 0.001 | -1.000 | up |
| SCYL1 | RP11-484O2.1 | -0.910 | 0.002 | -1.000 | up |
| SCYL1 | G050702 | -0.930 | 0.001 | -1.000 | up |
| SCYL1 | G009235 | -0.909 | 0.002 | -1.000 | up |
| SCYL1 | G063995 | -0.939 | 0.001 | -1.000 | up |
| SCYL1 | ZFAT-AS1 | -0.976 | 0.000 | -1.000 | up |
| SCYL1 | G087265 | -0.917 | 0.001 | -1.000 | up |
| SCYL1 | BX648501 | -0.930 | 0.001 | -1.000 | up |
| SCYL1 | CYP2D7 | -0.902 | 0.002 | -1.000 | up |
| SCYL1 | TMLHE-AS1 | -0.905 | 0.002 | -1.000 | up |
| SCYL1 | RP4-760C5.3 | -0.907 | 0.002 | -1.000 | up |
| SCYL1 | BCRP3 | -0.931 | 0.001 | -1.000 | up |
| SCYL1 | G068890 | -0.918 | 0.001 | -1.000 | up |
| SCYL1 | RP11-744I24.3 | 0.912 | 0.002 | 1.000 | up |
| SCYL1 | G002862 | 0.975 | 0.000 | 1.000 | up |
| SCYL1 | TUB-AS1 | 0.921 | 0.001 | 1.000 | up |
| SCYL1 | PRR36 | 0.936 | 0.001 | 1.000 | up |
| SCYL1 | CRYBB2P1 | 0.952 | 0.000 | 1.000 | up |
| SCYL1 | RP11-168O10.6 | 0.969 | 0.000 | 1.000 | up |
| ADAM15 | Alpha_250_dna | -0.906 | 0.002 | -1.000 | up |

| | | | | | |
|---|---|---|---|---|---|
| ADAM15 | G011310 | -0.933 | 0.001 | -1.000 | up |
| ADAM15 | ASAP1-IT1 | -0.927 | 0.001 | -1.000 | up |
| ADAM15 | RP11-737O24.3 | -0.914 | 0.001 | -1.000 | up |
| ADAM15 | RP11-669E14.6 | -0.910 | 0.002 | -1.000 | up |
| ADAM15 | uc.49 | -0.952 | 0.000 | -1.000 | up |
| ADAM15 | G073689 | -0.941 | 0.000 | -1.000 | up |
| ADAM15 | XLOC_012740 | -0.929 | 0.001 | -1.000 | up |
| ADAM15 | CTD-3193O13.1 | -0.918 | 0.001 | -1.000 | up |
| ADAM15 | G051586 | -0.904 | 0.002 | -1.000 | up |
| ADAM15 | G089283 | -0.963 | 0.000 | -1.000 | up |
| ADAM15 | RP5-1112D6.7 | -0.917 | 0.001 | -1.000 | up |
| ADAM15 | DLG3-AS1 | -0.904 | 0.002 | -1.000 | up |
| ADAM15 | XLOC_009429 | -0.916 | 0.001 | -1.000 | up |
| ADAM15 | G059595 | -0.942 | 0.000 | -1.000 | up |
| ADAM15 | IFT74-AS1 | 0.943 | 0.000 | 1.000 | up |
| ADAM15 | AL110181 | 0.930 | 0.001 | 1.000 | up |
| ADAM15 | TIPARP-AS1 | 0.903 | 0.002 | 1.000 | up |
| ADAM15 | RP11-363E6.3 | 0.902 | 0.002 | 1.000 | up |
| ADAM15 | LINC00652 | 0.945 | 0.000 | 1.000 | up |
| ADAM15 | RP1-40E16.12 | 0.915 | 0.001 | 1.000 | up |
| ADAM15 | G049752 | 0.900 | 0.002 | 1.000 | up |
| ADAM15 | SH3GL1P2 | 0.944 | 0.000 | 1.000 | up |
| ADAM15 | TM4SF1-AS1 | 0.944 | 0.000 | 1.000 | up |
| ADAM15 | LINC00662 | 0.923 | 0.001 | 1.000 | up |
| ADAM15 | G064655 | 0.939 | 0.001 | 1.000 | up |
| ADAM15 | RP11-252A24.3 | 0.907 | 0.002 | 1.000 | up |
| ADAM15 | RP11-715J22.6 | 0.931 | 0.001 | 1.000 | up |
| ADAM15 | AC099754.1 | 0.936 | 0.001 | 1.000 | up |
| ADAM15 | RP11-396C23.4 | 0.918 | 0.001 | 1.000 | up |
| ADAM15 | CTC-232P5.3 | 0.938 | 0.001 | 1.000 | up |
| ADAM15 | RP5-991G20.4 | 0.943 | 0.000 | 1.000 | up |
| ADAM15 | MGC2889 | 0.984 | 0.000 | 1.000 | up |
| ADAM15 | OLMALINC | 0.930 | 0.001 | 1.000 | up |
| ADAM15 | CCNT2-AS1 | 0.954 | 0.000 | 1.000 | up |
| ADAM15 | XLOC_011052 | 0.910 | 0.002 | 1.000 | up |
| ADAM15 | NIPBL-AS1 | 0.917 | 0.001 | 1.000 | up |
| ADAM15 | RP11-775D22.3 | 0.948 | 0.000 | 1.000 | up |
| ADAM15 | G066606 | 0.907 | 0.002 | 1.000 | up |
| ADAM15 | G078120 | 0.968 | 0.000 | 1.000 | up |
| ADAM15 | RP11-254F7.2 | 0.926 | 0.001 | 1.000 | up |
| ADAM15 | G066437 | 0.937 | 0.001 | 1.000 | up |
| ADAM15 | ENO1-AS1 | 0.910 | 0.002 | 1.000 | up |
| ADAM15 | DGCR5 | 0.930 | 0.001 | 1.000 | up |
| ADAM15 | LINC00463 | 0.911 | 0.002 | 1.000 | up |
| ADAM15 | G027543 | 0.927 | 0.001 | 1.000 | up |
| ADAM15 | LL09NC01-139C3.1 | 0.906 | 0.002 | 1.000 | up |
| ADAM15 | G068576 | 0.945 | 0.000 | 1.000 | up |
| ADAM15 | CXADRP3 | 0.940 | 0.001 | 1.000 | up |
| ADAM15 | G047565 | 0.906 | 0.002 | 1.000 | up |
| ADAM15 | RP11-110G21.1 | 0.976 | 0.000 | 1.000 | up |
| ADAM15 | BX248253 | 0.915 | 0.001 | 1.000 | up |
| ADAM15 | uc.349 | 0.942 | 0.000 | 1.000 | up |
| ADAM15 | G003603 | 0.940 | 0.001 | 1.000 | up |
| ADAM15 | RP11-53I6.3 | 0.920 | 0.001 | 1.000 | up |
| ADAM15 | RP11-314A20.5 | 0.977 | 0.000 | 1.000 | up |
| ADAM15 | RP11-196G18.24 | 0.930 | 0.001 | 1.000 | up |
| ADAM15 | LOC101927811 | 0.922 | 0.001 | 1.000 | up |
| ADAM15 | RP11-30L3.2 | 0.916 | 0.001 | 1.000 | up |
| ADAM15 | G014106 | 0.907 | 0.002 | 1.000 | up |
| ADAM15 | RP11-355F22.1 | 0.916 | 0.001 | 1.000 | up |
| ADAM15 | LOC283140 | 0.928 | 0.001 | 1.000 | up |
| ADAM15 | BC047644 | 0.932 | 0.001 | 1.000 | up |
| ADAM15 | LOC102723385 | 0.948 | 0.000 | 1.000 | up |
| ADAM15 | G063721 | 0.921 | 0.001 | 1.000 | up |
| ADAM15 | G001497 | 0.962 | 0.000 | 1.000 | up |

| | | | | | |
|---|---|---|---|---|---|
| ADAM15 | CTC-248O19.1 | 0.936 | 0.001 | 1.000 | up |
| HSP90AB1 | G073552 | -0.914 | 0.002 | -1.000 | up |
| HSP90AB1 | G046678 | -0.942 | 0.000 | -1.000 | up |
| HSP90AB1 | G081487 | -0.908 | 0.002 | -1.000 | up |
| HSP90AB1 | AC016682.1 | -0.902 | 0.002 | -1.000 | up |
| HSP90AB1 | RP11-484O2.1 | -0.944 | 0.000 | -1.000 | up |
| HSP90AB1 | G050702 | -0.940 | 0.001 | -1.000 | up |
| HSP90AB1 | G063995 | -0.922 | 0.001 | -1.000 | up |
| HSP90AB1 | ZFAT-AS1 | -0.978 | 0.000 | -1.000 | up |
| HSP90AB1 | TMLHE-AS1 | -0.923 | 0.001 | -1.000 | up |
| HSP90AB1 | BCRP3 | -0.964 | 0.000 | -1.000 | up |
| HSP90AB1 | G051433 | -0.901 | 0.002 | -1.000 | up |
| HSP90AB1 | RN7SL176P | -0.913 | 0.002 | -1.000 | up |
| HSP90AB1 | G059595 | -0.917 | 0.001 | -1.000 | up |
| HSP90AB1 | G002862 | 0.911 | 0.002 | 1.000 | up |
| HSP90AB1 | G079888 | 0.909 | 0.002 | 1.000 | up |
| HSP90AB1 | PRR36 | 0.955 | 0.000 | 1.000 | up |
| HSP90AB1 | CRYBB2P1 | 0.969 | 0.000 | 1.000 | up |
| HSP90AB1 | G063514 | 0.925 | 0.001 | 1.000 | up |
| HSP90AB1 | JARID2-AS1 | 0.949 | 0.000 | 1.000 | up |
| HSP90AB1 | RP11-168O10.6 | 0.918 | 0.001 | 1.000 | up |
| NFAT5 | AC074363.1 | -0.902 | 0.002 | -1.000 | up |
| NFAT5 | CRAT37 | -0.962 | 0.000 | -1.000 | up |
| NFAT5 | G022034 | -0.903 | 0.002 | -1.000 | up |
| NFAT5 | XLOC_005405 | -0.906 | 0.002 | -1.000 | up |
| NFAT5 | AK025321 | -0.903 | 0.002 | -1.000 | up |
| NFAT5 | CTC-250I14.3 | -0.968 | 0.000 | -1.000 | up |
| NFAT5 | G010941 | -0.921 | 0.001 | -1.000 | up |
| NFAT5 | XLOC_010525 | 0.944 | 0.000 | 1.000 | up |
| NFAT5 | G020528 | 0.931 | 0.001 | 1.000 | up |
| NFAT5 | RP11-762I7.4 | 0.918 | 0.001 | 1.000 | up |
| NFAT5 | TAF1A-AS1 | 0.904 | 0.002 | 1.000 | up |
| NFAT5 | G066319 | 0.940 | 0.001 | 1.000 | up |
| NFAT5 | RP11-680F8.1 | 0.902 | 0.002 | 1.000 | up |
| NFAT5 | RP11-483I13.5 | 0.924 | 0.001 | 1.000 | up |
| NFAT5 | RN7SL832P | 0.935 | 0.001 | 1.000 | up |
| GATA4 | G089550 | -0.979 | 0.000 | -1.000 | up |
| GATA4 | RP11-587H10.1 | -0.947 | 0.000 | -1.000 | up |
| GATA4 | LOC642846 | -0.971 | 0.000 | -1.000 | up |
| GATA4 | G053358 | -0.902 | 0.002 | -1.000 | up |
| GATA4 | G033096 | -0.910 | 0.002 | -1.000 | up |
| GATA4 | G086828 | 0.906 | 0.002 | 1.000 | up |
| GATA4 | LINC00463 | 0.931 | 0.001 | 1.000 | up |
| GATA4 | G068576 | 0.939 | 0.001 | 1.000 | up |
| GATA4 | RP11-110G21.1 | 0.906 | 0.002 | 1.000 | up |
| GATA4 | G006089 | 0.935 | 0.001 | 1.000 | up |
| GATA4 | RP11-362L22.1 | 0.905 | 0.002 | 1.000 | up |
| NLRP2 | G037251 | -0.943 | 0.000 | -1.000 | up |
| NLRP2 | IGBP1-AS2 | -0.907 | 0.002 | -1.000 | up |
| NLRP2 | RP11-3K16.2 | -0.948 | 0.000 | -1.000 | up |
| NLRP2 | LACTB2-AS1 | -0.922 | 0.001 | -1.000 | up |
| NLRP2 | G033744 | -0.959 | 0.000 | -1.000 | up |
| NLRP2 | AC074363.1 | -0.910 | 0.002 | -1.000 | up |
| NLRP2 | RP11-587H10.1 | -0.942 | 0.000 | -1.000 | up |
| NLRP2 | RP11-737O24.3 | -0.946 | 0.000 | -1.000 | up |
| NLRP2 | G040015 | -0.928 | 0.001 | -1.000 | up |
| NLRP2 | uc.49 | -0.923 | 0.001 | -1.000 | up |
| NLRP2 | G001795 | -0.926 | 0.001 | -1.000 | up |
| NLRP2 | G051586 | -0.916 | 0.001 | -1.000 | up |
| NLRP2 | G033096 | -0.949 | 0.000 | -1.000 | up |
| NLRP2 | G089283 | -0.936 | 0.001 | -1.000 | up |
| NLRP2 | DLG3-AS1 | -0.917 | 0.001 | -1.000 | up |
| NLRP2 | G074298 | -0.909 | 0.002 | -1.000 | up |
| NLRP2 | RP11-44N22.3 | 0.915 | 0.001 | 1.000 | up |
| NLRP2 | FAM230C | 0.922 | 0.001 | 1.000 | up |

| | | | | | |
|---|---|---|---|---|---|
| NLRP2 | RP11-355B11.2 | 0.903 | 0.002 | 1.000 | up |
| NLRP2 | DLG5-AS1 (1) | 0.925 | 0.001 | 1.000 | up |
| NLRP2 | RP11-46J23.1 | 0.915 | 0.001 | 1.000 | up |
| NLRP2 | RP11-715J22.6 | 0.952 | 0.000 | 1.000 | up |
| NLRP2 | G079009 | 0.913 | 0.002 | 1.000 | up |
| NLRP2 | RP11-396C23.4 | 0.933 | 0.001 | 1.000 | up |
| NLRP2 | CTC-232P5.3 | 0.909 | 0.002 | 1.000 | up |
| NLRP2 | G085410 | 0.906 | 0.002 | 1.000 | up |
| NLRP2 | G052526 | 0.957 | 0.000 | 1.000 | up |
| NLRP2 | G041225 | 0.905 | 0.002 | 1.000 | up |
| NLRP2 | G074033 | 0.907 | 0.002 | 1.000 | up |
| NLRP2 | CCNT2-AS1 | 0.948 | 0.000 | 1.000 | up |
| NLRP2 | RP11-502I4.3 | 0.902 | 0.002 | 1.000 | up |
| NLRP2 | G010782 | 0.906 | 0.002 | 1.000 | up |
| NLRP2 | ENO1-AS1 | 0.937 | 0.001 | 1.000 | up |
| NLRP2 | G027543 | 0.911 | 0.002 | 1.000 | up |
| NLRP2 | G061722 | 0.913 | 0.002 | 1.000 | up |
| NLRP2 | G068576 | 0.918 | 0.001 | 1.000 | up |
| NLRP2 | XLOC_004632 | 0.910 | 0.002 | 1.000 | up |
| NLRP2 | RP11-110G21.1 | 0.939 | 0.001 | 1.000 | up |
| NLRP2 | BX248253 | 0.909 | 0.002 | 1.000 | up |
| NLRP2 | uc.349 | 0.951 | 0.000 | 1.000 | up |
| NLRP2 | G003603 | 0.916 | 0.001 | 1.000 | up |
| NLRP2 | G014106 | 0.918 | 0.001 | 1.000 | up |
| NLRP2 | RP11-355F22.1 | 0.919 | 0.001 | 1.000 | up |
| NLRP2 | DNAJC9-AS1 | 0.929 | 0.001 | 1.000 | up |
| NLRP2 | RP11-820L6.1 | 0.902 | 0.002 | 1.000 | up |
| NLRP2 | LOC102723385 | 0.954 | 0.000 | 1.000 | up |
| NLRP2 | G063721 | 0.901 | 0.002 | 1.000 | up |
| NLRP2 | CTC-248O19.1 | 0.908 | 0.002 | 1.000 | up |
| PON2 | G053736 | -0.911 | 0.002 | -1.000 | up |
| PON2 | G076011 | -0.936 | 0.001 | -1.000 | up |
| PON2 | G068163 | -0.920 | 0.001 | -1.000 | up |
| PON2 | RP3-416H24.4 | -0.973 | 0.000 | -1.000 | up |
| PON2 | G024041 | -0.912 | 0.002 | -1.000 | up |
| PON2 | G082861 | -0.957 | 0.000 | -1.000 | up |
| PON2 | G060484 | -0.945 | 0.000 | -1.000 | up |
| PON2 | RP11-599B13.7 | 0.905 | 0.002 | 1.000 | up |
| PON2 | RP11-182N22.8 | 0.952 | 0.000 | 1.000 | up |
| PON2 | SIX3-AS1 | 0.926 | 0.001 | 1.000 | up |
| PON2 | RP5-884G6.2 | 0.931 | 0.001 | 1.000 | up |
| PON2 | RP1-97J1.2 | 0.903 | 0.002 | 1.000 | up |
| PON2 | RP11-53B2.6 | 0.907 | 0.002 | 1.000 | up |
| PON2 | DHRS4-AS1 (2) | 0.929 | 0.001 | 1.000 | up |
| PON2 | RP6-109B7.4 | 0.924 | 0.001 | 1.000 | up |
| IGF2R | HCG18 | -0.917 | 0.001 | -1.000 | up |
| IGF2R | G038095 | -0.930 | 0.001 | -1.000 | up |
| IGF2R | G015131 | -0.915 | 0.001 | -1.000 | up |
| IGF2R | G002461 | -0.960 | 0.000 | -1.000 | up |
| IGF2R | AF086132 | -0.906 | 0.002 | -1.000 | up |
| IGF2R | FAM74A4 | 0.903 | 0.002 | 1.000 | up |
| IGF2R | WWC2-AS2 | 0.902 | 0.002 | 1.000 | up |
| IGF2R | G013286 | 0.960 | 0.000 | 1.000 | up |
| IGF2R | RP11-290L1.2 | 0.901 | 0.002 | 1.000 | up |
| IGF2R | RP11-792A8.4 | 0.937 | 0.001 | 1.000 | up |
| IGF2R | RP11-567G11.1 | 0.909 | 0.002 | 1.000 | up |
| NELL1 | RP4-761J14.9 | 0.906 | 0.002 | 1.000 | down |
| NELL1 | G021277 | 0.936 | 0.001 | 1.000 | down |
| NELL1 | G012913 | 0.911 | 0.002 | 1.000 | down |
| NELL1 | XLOC_006182 | 0.934 | 0.001 | 1.000 | down |
| NELL1 | LINC01149 | 0.901 | 0.002 | 1.000 | down |
| NELL1 | G001795 | 0.973 | 0.000 | 1.000 | down |
| NELL1 | G060484 | 0.917 | 0.001 | 1.000 | down |
| NELL1 | G058409 | 0.901 | 0.002 | 1.000 | down |
| NELL1 | G046492 | 0.909 | 0.002 | 1.000 | down |

| | | | | | |
|---|---|---|---|---|---|
| NELL1 | XLOC_007237 | 0.914 | 0.001 | 1.000 | down |
| NELL1 | G079644 | 0.943 | 0.000 | 1.000 | down |
| NELL1 | RP11-230C9.2 | -0.924 | 0.001 | -1.000 | down |
| NELL1 | OR2L1P | -0.905 | 0.002 | -1.000 | down |
| NELL1 | PSMG3-AS1 (1) | -0.915 | 0.001 | -1.000 | down |
| NELL1 | XLOC_011336 | -0.930 | 0.001 | -1.000 | down |
| NELL1 | G049752 | -0.901 | 0.002 | -1.000 | down |
| NELL1 | CTD-2033A16.3 | -0.955 | 0.000 | -1.000 | down |
| NELL1 | CTC-336P14.1 | -0.928 | 0.001 | -1.000 | down |
| NELL1 | RP11-510M2.2 | -0.919 | 0.001 | -1.000 | down |
| NELL1 | G019632 | -0.946 | 0.000 | -1.000 | down |
| NELL1 | RP11-2E11.5 | -0.918 | 0.001 | -1.000 | down |
| NELL1 | UBAC2-AS1 | -0.920 | 0.001 | -1.000 | down |
| NELL1 | RP11-53B2.6 | -0.922 | 0.001 | -1.000 | down |
| NELL1 | DDX39B-AS1 | -0.958 | 0.000 | -1.000 | down |
| NELL1 | LINC00449 | -0.921 | 0.001 | -1.000 | down |
| NELL1 | uc.349 | -0.921 | 0.001 | -1.000 | down |
| NELL1 | RP11-565F19.2 | -0.912 | 0.002 | -1.000 | down |
| NELL1 | XXbac-B476C20.9 | -0.920 | 0.001 | -1.000 | down |
| NELL1 | LOC102723385 | -0.918 | 0.001 | -1.000 | down |
| NELL1 | TOB1-AS1 (2) | -0.904 | 0.002 | -1.000 | down |
| MPO | G058275 | 0.916 | 0.001 | 1.000 | down |
| MPO | RN7SKP264 | 0.910 | 0.002 | 1.000 | down |
| MPO | RP11-219C20.3 | 0.967 | 0.000 | 1.000 | down |
| MPO | BC034456 | 0.926 | 0.001 | 1.000 | down |
| MPO | G082713 | 0.928 | 0.001 | 1.000 | down |
| MPO | G017718 | -0.951 | 0.000 | -1.000 | down |
| MPO | G017886 | -0.929 | 0.001 | -1.000 | down |
| MPO | G017670 | -0.901 | 0.002 | -1.000 | down |
| MPO | uc.446 | -0.909 | 0.002 | -1.000 | down |
| MPO | XLOC_000308 | -0.904 | 0.002 | -1.000 | down |
| MPO | KB-1592A4.14 | -0.919 | 0.001 | -1.000 | down |
| MPO | G063668 | -0.928 | 0.001 | -1.000 | down |
| VCAM1 | RP11-219C20.3 | 0.920 | 0.001 | 1.000 | down |
| VCAM1 | G060010 | 0.922 | 0.001 | 1.000 | down |
| VCAM1 | G082713 | 0.920 | 0.001 | 1.000 | down |
| VCAM1 | G022519 | 0.935 | 0.001 | 1.000 | down |
| VCAM1 | G017718 | -0.931 | 0.001 | -1.000 | down |
| VCAM1 | G017886 | -0.986 | 0.000 | -1.000 | down |
| VCAM1 | uc.446 | -0.916 | 0.001 | -1.000 | down |
| VCAM1 | XLOC_000308 | -0.933 | 0.001 | -1.000 | down |
| VCAM1 | KB-1592A4.14 | -0.901 | 0.002 | -1.000 | down |
| VCAM1 | G063668 | -0.923 | 0.001 | -1.000 | down |
| VCAM1 | G057325 | -0.924 | 0.001 | -1.000 | down |
| VCAM1 | G027543 | -0.938 | 0.001 | -1.000 | down |
| UCP2 | RP3-416H24.4 | 0.936 | 0.001 | 1.000 | down |
| UCP2 | G021277 | 0.949 | 0.000 | 1.000 | down |
| UCP2 | G072364 | 0.936 | 0.001 | 1.000 | down |
| UCP2 | G020793 | 0.951 | 0.000 | 1.000 | down |
| UCP2 | LINC01149 | 0.927 | 0.001 | 1.000 | down |
| UCP2 | G001795 | 0.924 | 0.001 | 1.000 | down |
| UCP2 | G060484 | 0.961 | 0.000 | 1.000 | down |
| UCP2 | G015181 | 0.935 | 0.001 | 1.000 | down |
| UCP2 | G047503 | -0.913 | 0.002 | -1.000 | down |
| UCP2 | XLOC_011336 | -0.938 | 0.001 | -1.000 | down |
| UCP2 | LOC729083 | -0.904 | 0.002 | -1.000 | down |
| UCP2 | TCR-alpha | -0.919 | 0.001 | -1.000 | down |
| UCP2 | G019632 | -0.904 | 0.002 | -1.000 | down |
| UCP2 | LINC00449 | -0.940 | 0.001 | -1.000 | down |
| UCP2 | DHRS4-AS1 (2) | -0.947 | 0.000 | -1.000 | down |
| UCP2 | RP6-109B7.4 | -0.936 | 0.001 | -1.000 | down |
| UCP2 | G065507 | -0.901 | 0.002 | -1.000 | down |
| NBR1 | ASAP1-IT1 | 0.906 | 0.002 | 1.000 | down |
| NBR1 | RP11-3K16.2 | 0.904 | 0.002 | 1.000 | down |
| NBR1 | G087508 | 0.917 | 0.001 | 1.000 | down |

| | | | | | |
|---|---|---|---|---|---|
| NBR1 | G081487 | 0.921 | 0.001 | 1.000 | down |
| NBR1 | XLOC_003405 | 0.945 | 0.000 | 1.000 | down |
| NBR1 | RP11-669E14.6 | 0.929 | 0.001 | 1.000 | down |
| NBR1 | G012913 | 0.914 | 0.002 | 1.000 | down |
| NBR1 | CTC-261N6.1 | 0.964 | 0.000 | 1.000 | down |
| NBR1 | G071640 | 0.948 | 0.000 | 1.000 | down |
| NBR1 | G014972 | 0.904 | 0.002 | 1.000 | down |
| NBR1 | XLOC_003837 | 0.924 | 0.001 | 1.000 | down |
| NBR1 | XLOC_012740 | 0.920 | 0.001 | 1.000 | down |
| NBR1 | LINC00685 | 0.916 | 0.001 | 1.000 | down |
| NBR1 | CTD-3193O13.1 | 0.915 | 0.001 | 1.000 | down |
| NBR1 | RP11-651L5.2 | 0.905 | 0.002 | 1.000 | down |
| NBR1 | G089283 | 0.941 | 0.000 | 1.000 | down |
| NBR1 | RP5-1112D6.7 | 0.942 | 0.000 | 1.000 | down |
| NBR1 | MIMT1 | 0.922 | 0.001 | 1.000 | down |
| NBR1 | G031271 | 0.901 | 0.002 | 1.000 | down |
| NBR1 | G074298 | 0.965 | 0.000 | 1.000 | down |
| NBR1 | G059595 | 0.947 | 0.000 | 1.000 | down |
| NBR1 | AL110181 | -0.918 | 0.001 | -1.000 | down |
| NBR1 | RP11-255E6.6 | -0.956 | 0.000 | -1.000 | down |
| NBR1 | RP11-363E6.3 | -0.908 | 0.002 | -1.000 | down |
| NBR1 | AK055712 | -0.901 | 0.002 | -1.000 | down |
| NBR1 | RPL23AP7 | -0.982 | 0.000 | -1.000 | down |
| NBR1 | LINC00662 | -0.947 | 0.000 | -1.000 | down |
| NBR1 | RP11-179B2.2 | -0.903 | 0.002 | -1.000 | down |
| NBR1 | G064655 | -0.963 | 0.000 | -1.000 | down |
| NBR1 | RP11-715J22.6 | -0.946 | 0.000 | -1.000 | down |
| NBR1 | AC099754.1 | -0.965 | 0.000 | -1.000 | down |
| NBR1 | RP5-991G20.4 | -0.933 | 0.001 | -1.000 | down |
| NBR1 | MGC2889 | -0.926 | 0.001 | -1.000 | down |
| NBR1 | G082153 | -0.927 | 0.001 | -1.000 | down |
| NBR1 | CTD-2005H7.1 | -0.960 | 0.000 | -1.000 | down |
| NBR1 | G079888 | -0.919 | 0.001 | -1.000 | down |
| NBR1 | MAPT-IT1 | -0.906 | 0.002 | -1.000 | down |
| NBR1 | G074033 | -0.906 | 0.002 | -1.000 | down |
| NBR1 | CCNT2-AS1 | -0.969 | 0.000 | -1.000 | down |
| NBR1 | RP11-775D22.3 | -0.902 | 0.002 | -1.000 | down |
| NBR1 | PVRL3-AS1 | -0.923 | 0.001 | -1.000 | down |
| NBR1 | RP11-1275H24.1 | -0.900 | 0.002 | -1.000 | down |
| NBR1 | XLOC_004632 | -0.906 | 0.002 | -1.000 | down |
| NBR1 | CXADRP3 | -0.917 | 0.001 | -1.000 | down |
| NBR1 | EZR-AS1 | -0.904 | 0.002 | -1.000 | down |
| NBR1 | G061596 | -0.924 | 0.001 | -1.000 | down |
| NBR1 | BX248253 | -0.957 | 0.000 | -1.000 | down |
| NBR1 | RP11-565F19.2 | -0.901 | 0.002 | -1.000 | down |
| NBR1 | G003603 | -0.965 | 0.000 | -1.000 | down |
| NBR1 | RP11-53I6.3 | -0.944 | 0.000 | -1.000 | down |
| NBR1 | RP11-314A20.5 | -0.942 | 0.000 | -1.000 | down |
| NBR1 | AK055974 | -0.933 | 0.001 | -1.000 | down |
| NBR1 | RP11-196G18.24 | -0.909 | 0.002 | -1.000 | down |
| NBR1 | LOC101927811 | -0.940 | 0.001 | -1.000 | down |
| NBR1 | uc.412 | -0.935 | 0.001 | -1.000 | down |
| NBR1 | G000727 | -0.936 | 0.001 | -1.000 | down |
| NBR1 | RP11-330O11.3 | -0.937 | 0.001 | -1.000 | down |
| NBR1 | LOC283140 | -0.924 | 0.001 | -1.000 | down |
| NBR1 | CTB-181F24.1 | -0.966 | 0.000 | -1.000 | down |
| NBR1 | LOC102723385 | -0.977 | 0.000 | -1.000 | down |
| NBR1 | RP11-803B1.8 | -0.908 | 0.002 | -1.000 | down |
| NBR1 | G063721 | -0.943 | 0.000 | -1.000 | down |
| NBR1 | G015529 | -0.910 | 0.002 | -1.000 | down |
| NBR1 | LOC101927358 | -0.923 | 0.001 | -1.000 | down |
| C1QTNF9B | RP11-21M24.3 | 0.937 | 0.001 | 1.000 | down |
| C1QTNF9B | G089550 | 0.932 | 0.001 | 1.000 | down |
| C1QTNF9B | RP11-187C18.2 | 0.916 | 0.001 | 1.000 | down |
| C1QTNF9B | G031832 | 0.922 | 0.001 | 1.000 | down |

| | | | | | |
|---|---|---|---|---|---|
| C1QTNF9B | G082315 | 0.925 | 0.001 | 1.000 | down |
| C1QTNF9B | AC026188.1 | 0.933 | 0.001 | 1.000 | down |
| C1QTNF9B | GSE61474_XLOC_042934 | 0.934 | 0.001 | 1.000 | down |
| C1QTNF9B | G053358 | 0.947 | 0.000 | 1.000 | down |
| C1QTNF9B | MINOS1P1 | 0.945 | 0.000 | 1.000 | down |
| C1QTNF9B | XLOC_003724 | -0.946 | 0.000 | -1.000 | down |
| C1QTNF9B | G090511 | -0.943 | 0.000 | -1.000 | down |
| C1QTNF9B | TPI1P3 | -0.926 | 0.001 | -1.000 | down |

**Supplementary Table 12. General anesthetic-induced dysregulated coding genes (mRNAs) in serum from pediatric patients < 4 years old**

| Gene ID | P value | Fold change | Regulation |
|---|---|---|---|
| FAM217A | 0.002 | 1.201 | Down |
| TMEM81 | 0.047 | 1.204 | Down |
| VPS11 | 0.020 | 1.204 | Down |
| B4GALNT2 | 0.003 | 1.204 | Up |
| CNR2 | 0.024 | 1.206 | Down |
| CATG00000040684.1 | 0.011 | 1.207 | Down |
| NBEAL2 | 0.002 | 1.207 | Up |
| LCN6 | 0.044 | 1.207 | Up |
| ZNF277 | 0.023 | 1.209 | Down |
| SPOCD1 | 0.003 | 1.209 | Down |
| SLC25A32 | 0.012 | 1.209 | Up |
| SLC25A25 | 0.013 | 1.210 | Down |
| HK3 | 0.042 | 1.212 | Up |
| ZNF552 | 0.012 | 1.215 | Down |
| MON1B | 0.001 | 1.217 | Down |
| DPY19L1 | 0.004 | 1.222 | Down |
| TKFC | 0.005 | 1.223 | Down |
| EIF2B2 | 0.014 | 1.225 | Down |
| TPTE2 | 0.017 | 1.225 | Down |
| PTPRJ | 0.028 | 1.226 | Up |
| UBQLN3 | 0.047 | 1.228 | Up |
| DBNDD1 | 0.031 | 1.229 | Down |
| PGLYRP2 | 0.017 | 1.229 | Up |
| SYT2 | 0.046 | 1.232 | Up |
| PGPEP1 | 0.043 | 1.232 | Up |
| LILRA6 | 0.029 | 1.233 | Down |
| RWDD4 | 0.017 | 1.235 | Up |
| FEM1B | 0.044 | 1.235 | Up |
| MYH6 | 0.026 | 1.236 | Up |
| UBE4B | 0.046 | 1.237 | Down |
| CATG00000074958.1 | 0.029 | 1.239 | Up |
| SP7 | 0.044 | 1.240 | Down |
| SART3 | 0.041 | 1.241 | Up |
| FGF19 | 0.042 | 1.243 | Up |
| FBXW8 | 0.035 | 1.244 | Up |
| MPP1 | 0.015 | 1.244 | Up |
| DUSP19 | 0.040 | 1.245 | Down |
| RABL2B | 0.026 | 1.247 | Down |
| CORO2B | 0.048 | 1.248 | Up |
| S1PR3 | 0.045 | 1.249 | Up |
| CCNH | 0.006 | 1.250 | Up |
| TMPRSS11A | 0.049 | 1.251 | Down |
| TMEM143 | 0.031 | 1.253 | Up |
| ADORA2A | 0.010 | 1.254 | Down |
| VXN | 0.026 | 1.255 | Up |
| INTS5 | 0.044 | 1.256 | Up |
| CISD1 | 0.022 | 1.256 | Down |
| REG3G | 0.049 | 1.259 | Up |
| NARF | 0.046 | 1.262 | Up |
| MCEMP1 | 0.037 | 1.263 | Up |
| MAPRE2 | 0.038 | 1.266 | Up |
| ANKS4B | 0.001 | 1.266 | Down |
| IFI27L2 | 0.034 | 1.266 | Up |
| CATG00000056197.1 | 0.050 | 1.267 | Down |
| GSC2 | 0.018 | 1.272 | Up |
| POLE4 | 0.050 | 1.272 | Down |
| EIF2B4 | 0.029 | 1.273 | Down |
| SPZ1 | 0.004 | 1.274 | Up |
| ACSS3 | 0.002 | 1.277 | Up |
| CAT | 0.038 | 1.277 | Up |

| | | | |
|---|---|---|---|
| INTS4 | 0.024 | 1.277 | Down |
| CATG00000025826.1 | 0.037 | 1.278 | Up |
| KCNMB1 | 0.022 | 1.278 | Up |
| RAB42 | 0.021 | 1.278 | Down |
| C2orf15 | 0.022 | 1.279 | Down |
| CATG00000102848.1 | 0.002 | 1.282 | Down |
| RSL1D1 | 0.023 | 1.283 | Up |
| GUCY1B1 | 0.020 | 1.283 | Up |
| SERPINF2 | 0.038 | 1.283 | Down |
| PLA1A | 0.010 | 1.284 | Down |
| CATG00000067666.1 | 0.021 | 1.284 | Up |
| AKAP1 | 0.001 | 1.287 | Up |
| CLIP3 | 0.034 | 1.289 | Up |
| TMTC1 | 0.029 | 1.292 | Down |
| ANKMY1 | 0.030 | 1.293 | Up |
| GAL3ST1 | 0.016 | 1.293 | Down |
| TACSTD2 | 0.035 | 1.293 | Up |
| CPNE8 | 0.049 | 1.297 | Down |
| ICOSLG | 0.033 | 1.297 | Up |
| KRT1 | 0.033 | 1.298 | Up |
| ARMH1 | 0.020 | 1.298 | Up |
| CATG00000038552.1 | 0.012 | 1.299 | Down |
| MED27 | 0.008 | 1.300 | Up |
| TMEM39B | 0.016 | 1.300 | Down |
| PGLYRP1 | 0.006 | 1.303 | Down |
| CATG00000030338.1 | 0.043 | 1.304 | Down |
| FCAR | 0.022 | 1.306 | Down |
| MT1G | 0.013 | 1.307 | Up |
| FAM160A1 | 0.044 | 1.308 | Up |
| HSD17B1 | 0.017 | 1.310 | Up |
| AL161784.1 | 0.045 | 1.310 | Up |
| TRIM31 | 0.045 | 1.311 | Up |
| CELF6 | 0.005 | 1.312 | Down |
| ZNF497 | 0.029 | 1.313 | Down |
| DGKA | 0.013 | 1.313 | Down |
| KIAA1551 | 0.028 | 1.313 | Up |
| FGFBP3 | 0.044 | 1.317 | Down |
| PTGES3L-AARSD1 | 0.008 | 1.318 | Down |
| IL23A | 0.026 | 1.318 | Up |
| BTN3A2 | 0.049 | 1.318 | Up |
| TACR2 | 0.019 | 1.320 | Down |
| CNDP2 | 0.032 | 1.320 | Up |
| SLC5A11 | 0.008 | 1.321 | Up |
| LRRC25 | 0.011 | 1.321 | Up |
| ATG7 | 0.017 | 1.321 | Down |
| GP2 | 0.029 | 1.322 | Down |
| PHYHIP | 0.020 | 1.322 | Down |
| CHRM1 | 0.036 | 1.322 | Down |
| AL590822.1 | 0.033 | 1.324 | Down |
| TEDC1 | 0.040 | 1.325 | Up |
| CATG00000025525.1 | 0.002 | 1.325 | Down |
| CYREN | 0.006 | 1.325 | Down |
| RPS28 | 0.030 | 1.326 | Down |
| KAZN | 0.020 | 1.326 | Down |
| BZW2 | 0.046 | 1.327 | Up |
| CATG00000100722.1 | 0.045 | 1.328 | Up |
| CCDC13 | 0.033 | 1.329 | Down |
| CATG00000031702.1 | 0.007 | 1.330 | Down |
| PUM2 | 0.044 | 1.335 | Down |
| KCTD13 | 0.037 | 1.336 | Up |
| KNG1 | 0.037 | 1.336 | Up |
| TMEM179 | 0.012 | 1.337 | Down |
| KLF2 | 0.030 | 1.338 | Up |
| ADCY3 | 0.002 | 1.339 | Up |
| SORT1 | 0.041 | 1.340 | Up |

| | | | |
|---|---|---|---|
| EMID1 | 0.039 | 1.342 | Up |
| SSNA1 | 0.028 | 1.343 | Up |
| DLL3 | 0.008 | 1.344 | Down |
| MTMR7 | 0.020 | 1.344 | Down |
| SNU13 | 0.033 | 1.346 | Down |
| MAF1 | 0.029 | 1.348 | Up |
| ACTN1 | 0.036 | 1.349 | Up |
| ENY2 | 0.017 | 1.351 | Down |
| SLC13A1 | 0.048 | 1.357 | Down |
| CES2 | 0.020 | 1.357 | Down |
| LACTB | 0.038 | 1.358 | Down |
| ADRM1 | 0.011 | 1.358 | Up |
| SLMAP | 0.048 | 1.358 | Up |
| HBA1 | 0.031 | 1.359 | Down |
| ATP5PD | 0.013 | 1.360 | Up |
| ZNF17 | 0.036 | 1.361 | Down |
| TMEM225 | 0.004 | 1.361 | Down |
| HNRNPK | 0.047 | 1.362 | Up |
| CATG00000005195. | 0.023 | 1.362 | Down |
| GIMD1 | 0.030 | 1.362 | Up |
| CASP3 | 0.041 | 1.363 | Down |
| CATG00000028328. | 0.030 | 1.364 | Up |
| NDST1 | 0.017 | 1.367 | Up |
| CBFB | 0.035 | 1.369 | Up |
| WRNIP1 | 0.015 | 1.370 | Up |
| DBT | 0.024 | 1.372 | Down |
| GABRB2 | 0.027 | 1.372 | Down |
| CATG00000049995. | 0.012 | 1.373 | Down |
| AL160272.2 | 0.000 | 1.375 | Up |
| CATG00000100036. | 0.046 | 1.376 | Down |
| NFYB | 0.048 | 1.377 | Down |
| NT5C1A | 0.047 | 1.378 | Up |
| STX16 | 0.026 | 1.378 | Up |
| TMEM155 | 0.025 | 1.383 | Up |
| SLC25A40 | 0.033 | 1.384 | Up |
| MIS12 | 0.048 | 1.385 | Down |
| ARID3B | 0.005 | 1.386 | Down |
| CTBP1 | 0.035 | 1.387 | Up |
| MASP2 | 0.003 | 1.390 | Down |
| NGDN | 0.007 | 1.390 | Up |
| CYP1A1 | 0.012 | 1.392 | Up |
| WDR62 | 0.043 | 1.392 | Up |
| SHOC2 | 0.006 | 1.393 | Up |
| TUBB4A | 0.038 | 1.394 | Down |
| GLOD4 | 0.001 | 1.397 | Down |
| KLB | 0.024 | 1.398 | Down |
| SGCG | 0.037 | 1.399 | Down |
| TBX10 | 0.016 | 1.399 | Down |
| SRSF1 | 0.041 | 1.400 | Down |
| ZMYND8 | 0.030 | 1.400 | Up |
| RGL3 | 0.010 | 1.401 | Down |
| ALG1L2 | 0.008 | 1.402 | Up |
| RPL3L | 0.007 | 1.402 | Up |
| FAM216A | 0.048 | 1.403 | Up |
| DCDC2C | 0.001 | 1.404 | Up |
| CATG00000010465. | 0.023 | 1.404 | Down |
| EMP2 | 0.002 | 1.406 | Down |
| CNTNAP2 | 0.013 | 1.406 | Down |
| ZNF878 | 0.000 | 1.408 | Up |
| SLC35A4 | 0.020 | 1.408 | Down |
| PPIP5K2 | 0.037 | 1.408 | Down |
| ZNF582 | 0.016 | 1.408 | Down |
| CATG00000099157. | 0.048 | 1.408 | Up |
| GSG1L2 | 0.024 | 1.409 | Down |
| SGCZ | 0.050 | 1.409 | Down |

| | | | |
|---|---|---|---|
| TMC4 | 0.015 | 1.411 | Down |
| USP14 | 0.044 | 1.411 | Up |
| C16orf86 | 0.049 | 1.411 | Down |
| TBC1D12 | 0.030 | 1.412 | Up |
| CAPZA1 | 0.000 | 1.413 | Up |
| UNC13A | 0.020 | 1.413 | Down |
| OR2D3 | 0.045 | 1.413 | Down |
| TUBB2A | 0.022 | 1.415 | Up |
| SEC14L6 | 0.016 | 1.416 | Up |
| CATG00000037285.1 | 0.048 | 1.416 | Down |
| MAP4K5 | 0.020 | 1.417 | Up |
| OR4D11 | 0.027 | 1.417 | Up |
| ANO6 | 0.019 | 1.417 | Up |
| PRMT1 | 0.019 | 1.420 | Up |
| RNF216 | 0.018 | 1.420 | Up |
| ZNF667 | 0.043 | 1.422 | Up |
| CCL18 | 0.029 | 1.422 | Down |
| CATG00000002739.1 | 0.020 | 1.422 | Up |
| NKX2-5 | 0.047 | 1.424 | Up |
| PLAT | 0.023 | 1.427 | Down |
| ARHGAP45 | 0.022 | 1.427 | Up |
| TPH1 | 0.020 | 1.427 | Up |
| SLC26A9 | 0.002 | 1.428 | Up |
| ATPAF1 | 0.037 | 1.428 | Down |
| ATP6V0E2 | 0.034 | 1.430 | Down |
| ANG | 0.036 | 1.430 | Down |
| TSC22D4 | 0.021 | 1.430 | Up |
| TPMT | 0.025 | 1.433 | Up |
| XDH | 0.009 | 1.435 | Down |
| ZNF174 | 0.018 | 1.436 | Down |
| SLC28A1 | 0.037 | 1.437 | Down |
| FAM217B | 0.046 | 1.439 | Up |
| DHX29 | 0.002 | 1.439 | Down |
| C6orf58 | 0.002 | 1.442 | Down |
| AC009163.4 | 0.035 | 1.442 | Down |
| UPF1 | 0.038 | 1.442 | Up |
| SPINK13 | 0.041 | 1.442 | Up |
| COL26A1 | 0.036 | 1.443 | Up |
| TUBA1C | 0.031 | 1.443 | Up |
| DIXDC1 | 0.024 | 1.448 | Up |
| ZNF835 | 0.038 | 1.450 | Up |
| POMGNT2 | 0.029 | 1.450 | Up |
| UCK1 | 0.003 | 1.450 | Up |
| HCFC1 | 0.006 | 1.450 | Down |
| GPRC6A | 0.013 | 1.450 | Down |
| PCDHA4 | 0.028 | 1.452 | Up |
| XKRX | 0.042 | 1.454 | Down |
| CATG00000042334.1 | 0.012 | 1.455 | Up |
| KRT86 | 0.046 | 1.456 | Down |
| PSG7 | 0.040 | 1.456 | Up |
| ZIM2 | 0.039 | 1.456 | Down |
| HAO2 | 0.018 | 1.458 | Down |
| DEFB124 | 0.002 | 1.458 | Down |
| DDR1 | 0.022 | 1.460 | Down |
| GIF | 0.017 | 1.460 | Down |
| RESP18 | 0.040 | 1.461 | Up |
| METTL24 | 0.022 | 1.462 | Up |
| FAM43A | 0.000 | 1.463 | Down |
| NT5DC4 | 0.037 | 1.463 | Down |
| FBXL18 | 0.006 | 1.464 | Up |
| TIGAR | 0.007 | 1.464 | Up |
| SETD1B | 0.040 | 1.464 | Up |
| THAP8 | 0.040 | 1.464 | Up |
| GPRIN3 | 0.010 | 1.465 | Up |
| SHC2 | 0.015 | 1.466 | Up |

| | | | |
|---|---|---|---|
| NUBP2 | 0.041 | 1.466 | Up |
| SLC22A18 | 0.048 | 1.467 | Up |
| WBP11 | 0.045 | 1.468 | Up |
| NMUR2 | 0.014 | 1.469 | Down |
| CYP26B1 | 0.034 | 1.469 | Up |
| CRISP3 | 0.046 | 1.470 | Down |
| LEF1 | 0.040 | 1.471 | Down |
| ORMDL3 | 0.042 | 1.472 | Up |
| PNPO | 0.014 | 1.472 | Down |
| SEZ6L2 | 0.042 | 1.472 | Up |
| RHOG | 0.023 | 1.472 | Up |
| BHLHE41 | 0.041 | 1.473 | Down |
| RHOXF1 | 0.019 | 1.473 | Down |
| KAT2A | 0.013 | 1.474 | Up |
| S100A16 | 0.021 | 1.475 | Up |
| CTNNAL1 | 0.005 | 1.476 | Down |
| CATG00000002574.1 | 0.024 | 1.476 | Up |
| RGL4 | 0.009 | 1.477 | Up |
| CYP2A6 | 0.007 | 1.478 | Up |
| PCSK6 | 0.016 | 1.478 | Down |
| KCNAB2 | 0.048 | 1.478 | Up |
| ZNF200 | 0.014 | 1.478 | Up |
| DHRSX | 0.018 | 1.479 | Up |
| PLCXD2 | 0.019 | 1.480 | Up |
| RPE65 | 0.050 | 1.481 | Down |
| KAAG1 | 0.041 | 1.482 | Down |
| MARCH9 | 0.033 | 1.483 | Up |
| FAM214B | 0.035 | 1.483 | Down |
| DPM1 | 0.049 | 1.484 | Down |
| ALKBH6 | 0.042 | 1.484 | Up |
| TMC8 | 0.041 | 1.486 | Up |
| PEX7 | 0.035 | 1.486 | Up |
| CATG00000071025.1 | 0.040 | 1.487 | Up |
| YRDC | 0.026 | 1.487 | Up |
| SLC29A2 | 0.043 | 1.488 | Up |
| SYCP2 | 0.032 | 1.490 | Up |
| NEURL1B | 0.005 | 1.493 | Up |
| TGFBI | 0.014 | 1.493 | Up |
| EDF1 | 0.041 | 1.494 | Up |
| TAT | 0.008 | 1.496 | Up |
| MVP | 0.035 | 1.497 | Up |
| CAP2 | 0.003 | 1.497 | Down |
| EID1 | 0.007 | 1.497 | Down |
| SUMF2 | 0.012 | 1.498 | Up |
| AMZ1 | 0.039 | 1.499 | Up |
| NEURL3 | 0.016 | 1.500 | Up |
| REXO1L1 | 0.037 | 1.500 | Up |
| GZMM | 0.019 | 1.501 | Down |
| CD79A | 0.008 | 1.501 | Up |
| SDCBP | 0.009 | 1.501 | Up |
| MOGAT3 | 0.021 | 1.502 | Up |
| MYLIP | 0.041 | 1.503 | Up |
| PCOLCE | 0.005 | 1.504 | Down |
| RAP1GAP2 | 0.033 | 1.504 | Up |
| AC090427.1 | 0.027 | 1.504 | Down |
| CATG00000107174.1 | 0.023 | 1.504 | Up |
| KRTAP10-9 | 0.031 | 1.506 | Down |
| TJP2 | 0.037 | 1.506 | Up |
| CATG00000041439.1 | 0.030 | 1.507 | Down |
| DDOST | 0.041 | 1.510 | Up |
| TEX45 | 0.041 | 1.510 | Up |
| IL10RB | 0.006 | 1.510 | Down |
| CSNK1D | 0.030 | 1.511 | Up |
| ACTL8 | 0.032 | 1.512 | Down |
| C5orf67 | 0.021 | 1.515 | Up |

| | | | |
|---|---|---|---|
| CLK2 | 0.035 | 1.515 | Up |
| WDR7 | 0.046 | 1.516 | Up |
| TAZ | 0.028 | 1.519 | Up |
| STX4 | 0.012 | 1.519 | Up |
| CFAP161 | 0.029 | 1.519 | Up |
| SEPT12 | 0.019 | 1.522 | Up |
| TNPO3 | 0.008 | 1.522 | Down |
| ATIC | 0.014 | 1.522 | Up |
| NGEF | 0.040 | 1.523 | Up |
| CLDN25 | 0.032 | 1.523 | Down |
| PARP10 | 0.020 | 1.525 | Up |
| MRPS11 | 0.038 | 1.525 | Up |
| CATG00000058112.1 | 0.045 | 1.526 | Up |
| FIG4 | 0.015 | 1.526 | Down |
| KIAA1324L | 0.011 | 1.527 | Down |
| MFRP | 0.037 | 1.527 | Down |
| EVC | 0.003 | 1.527 | Up |
| SLC20A1 | 0.043 | 1.528 | Up |
| HMGB3 | 0.039 | 1.528 | Down |
| ZFYVE27 | 0.046 | 1.530 | Up |
| PCNA | 0.027 | 1.531 | Down |
| CATG00000067519.1 | 0.007 | 1.535 | Up |
| PRAC2 | 0.033 | 1.535 | Down |
| ASH2L | 0.015 | 1.535 | Down |
| CASR | 0.042 | 1.537 | Down |
| HIST1H2BK | 0.037 | 1.537 | Down |
| ZNF285 | 0.030 | 1.540 | Down |
| CATG00000052071.1 | 0.045 | 1.540 | Down |
| EHD4 | 0.015 | 1.540 | Up |
| SNRPD3 | 0.043 | 1.541 | Up |
| GLYR1 | 0.004 | 1.541 | Up |
| IFT140 | 0.042 | 1.542 | Down |
| MGAM | 0.009 | 1.543 | Down |
| ATG4D | 0.039 | 1.544 | Up |
| NEMF | 0.022 | 1.545 | Up |
| BIRC7 | 0.043 | 1.545 | Up |
| CATG00000071657.1 | 0.004 | 1.547 | Up |
| CATG00000031258.1 | 0.006 | 1.548 | Up |
| ROR2 | 0.003 | 1.548 | Down |
| CATG00000062929.1 | 0.004 | 1.549 | Up |
| TMEM255A | 0.041 | 1.549 | Down |
| EIF3G | 0.050 | 1.552 | Up |
| DGAT2 | 0.009 | 1.553 | Up |
| MANEAL | 0.033 | 1.553 | Up |
| BSN | 0.013 | 1.555 | Up |
| SSH3 | 0.018 | 1.555 | Down |
| SULT1A2 | 0.016 | 1.557 | Down |
| NXNL2 | 0.015 | 1.558 | Up |
| ACTRT2 | 0.036 | 1.563 | Up |
| KCNA4 | 0.049 | 1.563 | Up |
| SMIM32 | 0.030 | 1.564 | Up |
| IQCA1L | 0.026 | 1.565 | Up |
| TRIML1 | 0.008 | 1.566 | Up |
| IFITM2 | 0.004 | 1.566 | Up |
| PSEN2 | 0.021 | 1.566 | Up |
| OSBPL11 | 0.018 | 1.567 | Down |
| TRIM29 | 0.028 | 1.567 | Up |
| GABBR1 | 0.008 | 1.569 | Down |
| EEF2K | 0.011 | 1.569 | Up |
| RASL10B | 0.003 | 1.569 | Up |
| CEP170B | 0.027 | 1.570 | Up |
| SOBP | 0.033 | 1.570 | Up |
| BCL2L2 | 0.013 | 1.571 | Down |
| HINFP | 0.038 | 1.572 | Down |
| SKA3 | 0.003 | 1.573 | Up |

| | | | |
|---|---|---|---|
| S1PR4 | 0.040 | 1.574 | Down |
| NREP | 0.026 | 1.574 | Down |
| PNLIPRP3 | 0.046 | 1.574 | Down |
| TMSB15B | 0.027 | 1.576 | Down |
| SETBP1 | 0.026 | 1.577 | Up |
| TUBGCP2 | 0.050 | 1.578 | Down |
| CXorf57 | 0.027 | 1.578 | Up |
| RIN1 | 0.024 | 1.579 | Down |
| PKM | 0.034 | 1.579 | Up |
| CTSC | 0.030 | 1.580 | Up |
| MT1F | 0.022 | 1.581 | Up |
| CATG00000019495.1 | 0.008 | 1.581 | Up |
| TOM1 | 0.041 | 1.581 | Up |
| SLC26A4 | 0.037 | 1.581 | Down |
| AIF1L | 0.035 | 1.582 | Down |
| PLA2G15 | 0.030 | 1.582 | Down |
| CATG00000059127.1 | 0.043 | 1.582 | Up |
| LRRC8A | 0.050 | 1.582 | Down |
| KRBA2 | 0.022 | 1.585 | Up |
| ST8SIA2 | 0.022 | 1.586 | Up |
| CATG00000053992.1 | 0.031 | 1.587 | Up |
| UNC45B | 0.043 | 1.589 | Up |
| C10orf53 | 0.001 | 1.589 | Up |
| ST6GALNAC4 | 0.015 | 1.590 | Up |
| CALHM2 | 0.048 | 1.591 | Up |
| ELFN1 | 0.015 | 1.593 | Down |
| CD164L2 | 0.029 | 1.593 | Up |
| FAM126B | 0.015 | 1.593 | Down |
| HIST1H2BN | 0.040 | 1.596 | Up |
| ACP6 | 0.022 | 1.598 | Down |
| PCDHGA4 | 0.016 | 1.598 | Up |
| GPR155 | 0.000 | 1.599 | Down |
| CD36 | 0.017 | 1.600 | Up |
| TCEA1 | 0.010 | 1.600 | Up |
| GAP43 | 0.044 | 1.600 | Down |
| SHANK3 | 0.042 | 1.603 | Down |
| SCARB1 | 0.019 | 1.604 | Down |
| KRTAP5-4 | 0.018 | 1.606 | Down |
| PKD1 | 0.024 | 1.607 | Up |
| CATG00000077030.1 | 0.047 | 1.607 | Up |
| IGFBP2 | 0.033 | 1.607 | Up |
| NDC80 | 0.017 | 1.608 | Down |
| NCAPG2 | 0.036 | 1.609 | Down |
| CDC42BPB | 0.014 | 1.609 | Up |
| TMEM202 | 0.034 | 1.610 | Up |
| RAB33A | 0.014 | 1.612 | Down |
| VSIG1 | 0.028 | 1.613 | Up |
| OR1G1 | 0.004 | 1.614 | Down |
| HEPH | 0.045 | 1.615 | Down |
| DEFB134 | 0.017 | 1.615 | Down |
| C9orf153 | 0.015 | 1.615 | Up |
| ZCRB1 | 0.044 | 1.616 | Down |
| SOX15 | 0.033 | 1.616 | Up |
| RAB6B | 0.038 | 1.617 | Up |
| ZNF639 | 0.041 | 1.617 | Up |
| BEND2 | 0.016 | 1.617 | Down |
| VPS4A | 0.008 | 1.617 | Up |
| RBPJL | 0.015 | 1.618 | Down |
| ACP4 | 0.037 | 1.618 | Down |
| RELN | 0.019 | 1.618 | Up |
| RDH5 | 0.047 | 1.619 | Up |
| PDZK1IP1 | 0.000 | 1.619 | Up |
| ESRRB | 0.016 | 1.620 | Up |
| MREG | 0.040 | 1.621 | Up |
| ZNF365 | 0.025 | 1.621 | Up |

| | | | |
|---|---|---|---|
| INMT | 0.023 | 1.622 | Down |
| TGM5 | 0.005 | 1.623 | Up |
| GMPR | 0.020 | 1.624 | Down |
| RPL26 | 0.028 | 1.624 | Up |
| ABCA2 | 0.021 | 1.625 | Up |
| ADAMTS18 | 0.044 | 1.628 | Up |
| EVX2 | 0.033 | 1.628 | Down |
| ANKRD49 | 0.011 | 1.629 | Up |
| CATG00000100646.1 | 0.045 | 1.630 | Up |
| IL12RB1 | 0.030 | 1.630 | Down |
| CENPS | 0.020 | 1.632 | Down |
| DCUN1D5 | 0.038 | 1.633 | Up |
| PRPS1L1 | 0.013 | 1.634 | Down |
| TAS2R42 | 0.039 | 1.634 | Down |
| NEURL4 | 0.033 | 1.634 | Down |
| DEPDC7 | 0.049 | 1.634 | Up |
| API5 | 0.031 | 1.635 | Up |
| AC015989.2 | 0.048 | 1.637 | Down |
| KAT14 | 0.005 | 1.637 | Down |
| ACR | 0.043 | 1.637 | Down |
| SPEG | 0.028 | 1.638 | Down |
| ACKR1 | 0.022 | 1.639 | Down |
| NCKAP5L | 0.038 | 1.639 | Up |
| WAPL | 0.024 | 1.639 | Up |
| CATG00000109000.1 | 0.035 | 1.640 | Up |
| FCMR | 0.006 | 1.640 | Up |
| PPT1 | 0.026 | 1.641 | Up |
| AC019294.1 | 0.018 | 1.641 | Up |
| CASP9 | 0.020 | 1.641 | Up |
| SULT1A2 | 0.049 | 1.644 | Up |
| FAM166B | 0.002 | 1.644 | Up |
| ITPKB | 0.041 | 1.644 | Up |
| PPP2CB | 0.044 | 1.645 | Down |
| CATG00000090147.1 | 0.024 | 1.645 | Up |
| SLC35C2 | 0.029 | 1.647 | Up |
| ZNF746 | 0.011 | 1.647 | Up |
| CBLC | 0.012 | 1.648 | Up |
| NLN | 0.003 | 1.650 | Down |
| SLC35G1 | 0.035 | 1.650 | Down |
| HTN1 | 0.017 | 1.650 | Up |
| LRRC26 | 0.049 | 1.651 | Down |
| ACBD5 | 0.049 | 1.651 | Down |
| CUL7 | 0.048 | 1.652 | Up |
| KRTAP5-2 | 0.009 | 1.652 | Up |
| KRTAP10-2 | 0.049 | 1.653 | Down |
| PLA2G6 | 0.026 | 1.655 | Up |
| PMEL | 0.048 | 1.659 | Up |
| RGS5 | 0.008 | 1.659 | Down |
| ANKS1A | 0.049 | 1.660 | Up |
| CTSL | 0.016 | 1.660 | Up |
| CATG00000095962.1 | 0.007 | 1.662 | Up |
| ZBTB42 | 0.020 | 1.663 | Down |
| TSKU | 0.043 | 1.663 | Up |
| VCX | 0.048 | 1.665 | Down |
| HUG1 | 0.016 | 1.665 | Up |
| AQP10 | 0.034 | 1.666 | Up |
| EHD3 | 0.007 | 1.666 | Up |
| RHBDD3 | 0.043 | 1.667 | Up |
| FBXL16 | 0.013 | 1.667 | Up |
| CYP51A1 | 0.025 | 1.669 | Up |
| OR8S1 | 0.032 | 1.670 | Up |
| NRIP2 | 0.041 | 1.670 | Down |
| ADPRHL1 | 0.000 | 1.672 | Up |
| PHLDB1 | 0.001 | 1.674 | Down |
| CATG00000113805.1 | 0.036 | 1.675 | Up |

| | | | |
|---|---|---|---|
| CST7 | 0.006 | 1.675 | Up |
| SCLT1 | 0.032 | 1.675 | Up |
| AP001094.1 | 0.010 | 1.675 | Down |
| HAUS5 | 0.020 | 1.676 | Down |
| THEM5 | 0.038 | 1.678 | Up |
| BAG3 | 0.012 | 1.678 | Up |
| IMP4 | 0.013 | 1.679 | Up |
| SLC25A43 | 0.023 | 1.680 | Up |
| BLOC1S4 | 0.011 | 1.680 | Up |
| PRLH | 0.008 | 1.680 | Up |
| TYROBP | 0.031 | 1.680 | Down |
| HNRNPM | 0.031 | 1.681 | Up |
| MRI1 | 0.000 | 1.681 | Up |
| TMC1 | 0.003 | 1.681 | Down |
| UBE2R2 | 0.002 | 1.682 | Down |
| CATG00000013555.1 | 0.018 | 1.682 | Up |
| ACSS1 | 0.031 | 1.682 | Up |
| CATG00000096869.1 | 0.032 | 1.682 | Up |
| MZT2B | 0.033 | 1.687 | Down |
| CATG00000060044.1 | 0.045 | 1.688 | Up |
| CATG00000036987.1 | 0.017 | 1.689 | Up |
| PLCXD1 | 0.049 | 1.689 | Up |
| CD46 | 0.038 | 1.690 | Up |
| PRKD2 | 0.033 | 1.691 | Up |
| PQLC1 | 0.047 | 1.692 | Down |
| VOPP1 | 0.040 | 1.692 | Down |
| CSF1R | 0.045 | 1.693 | Up |
| PDPR | 0.043 | 1.694 | Up |
| C7orf25 | 0.016 | 1.694 | Down |
| SLC51B | 0.006 | 1.694 | Up |
| C16orf58 | 0.030 | 1.695 | Up |
| ZMYM3 | 0.026 | 1.696 | Up |
| NOP58 | 0.008 | 1.696 | Up |
| WDR18 | 0.048 | 1.697 | Up |
| CES3 | 0.001 | 1.697 | Down |
| PRELID3B | 0.048 | 1.699 | Up |
| SAMD1 | 0.030 | 1.703 | Up |
| GGCT | 0.023 | 1.705 | Up |
| MED20 | 0.024 | 1.706 | Down |
| ERH | 0.016 | 1.707 | Down |
| RNF103-CHMP3 | 0.027 | 1.707 | Down |
| MIPOL1 | 0.036 | 1.708 | Up |
| CYB561D2 | 0.015 | 1.710 | Up |
| TMEM268 | 0.048 | 1.710 | Up |
| VAMP8 | 0.009 | 1.713 | Up |
| NEK6 | 0.030 | 1.713 | Down |
| GTF2IRD1 | 0.030 | 1.714 | Down |
| APLP2 | 0.041 | 1.715 | Down |
| CATG00000079682.1 | 0.008 | 1.717 | Up |
| SERPINB12 | 0.039 | 1.718 | Down |
| SP8 | 0.009 | 1.720 | Up |
| CATG00000106826.1 | 0.007 | 1.720 | Up |
| BTN3A1 | 0.017 | 1.720 | Down |
| ABHD16A | 0.013 | 1.721 | Up |
| FUT11 | 0.033 | 1.725 | Down |
| RAD1 | 0.027 | 1.725 | Down |
| ZNF208 | 0.044 | 1.726 | Up |
| NDUFB10 | 0.028 | 1.726 | Up |
| GUCA1B | 0.013 | 1.727 | Up |
| AKAP3 | 0.033 | 1.731 | Up |
| MAP7 | 0.037 | 1.732 | Down |
| RTEL1 | 0.012 | 1.736 | Up |
| TSLP | 0.006 | 1.736 | Down |
| EOGT | 0.037 | 1.737 | Up |
| F10 | 0.045 | 1.737 | Up |

| | | | |
|---|---|---|---|
| DDX25 | 0.008 | 1.737 | Up |
| ASCL4 | 0.043 | 1.738 | Down |
| CATG00000038465.1 | 0.049 | 1.739 | Up |
| GOT1L1 | 0.008 | 1.740 | Up |
| LRRC36 | 0.040 | 1.740 | Down |
| SLC44A3 | 0.023 | 1.741 | Down |
| ZNF337 | 0.022 | 1.742 | Down |
| CAMSAP3 | 0.003 | 1.743 | Down |
| TPRA1 | 0.043 | 1.746 | Down |
| GNAZ | 0.023 | 1.746 | Up |
| COL11A2 | 0.007 | 1.747 | Up |
| AL035078.4 | 0.045 | 1.748 | Up |
| IL2RG | 0.013 | 1.748 | Up |
| PATE3 | 0.044 | 1.751 | Down |
| CAPN3 | 0.015 | 1.752 | Up |
| PCDHAC1 | 0.037 | 1.754 | Up |
| SLC26A8 | 0.040 | 1.756 | Up |
| TMIGD2 | 0.046 | 1.757 | Up |
| HLA-DRA | 0.038 | 1.758 | Up |
| MRGPRE | 0.047 | 1.759 | Down |
| RTEL1 | 0.026 | 1.761 | Up |
| HAS1 | 0.008 | 1.762 | Up |
| TRIM60 | 0.039 | 1.762 | Down |
| DNAH8 | 0.038 | 1.762 | Up |
| CATG00000041177.1 | 0.018 | 1.764 | Up |
| TIGIT | 0.015 | 1.764 | Down |
| KIAA1210 | 0.046 | 1.768 | Down |
| AC007773.3 | 0.026 | 1.773 | Up |
| PYCR1 | 0.033 | 1.774 | Up |
| USP31 | 0.033 | 1.774 | Down |
| LDHA | 0.017 | 1.775 | Up |
| KIF21B | 0.027 | 1.776 | Up |
| VANGL1 | 0.020 | 1.776 | Down |
| CATG00000001978.1 | 0.029 | 1.777 | Down |
| ZNF30 | 0.011 | 1.777 | Up |
| AGPAT2 | 0.006 | 1.778 | Up |
| CATG00000110136.1 | 0.043 | 1.779 | Down |
| WDR46 | 0.002 | 1.780 | Up |
| TPTE | 0.019 | 1.780 | Up |
| OR6C65 | 0.005 | 1.782 | Down |
| HTN3 | 0.012 | 1.782 | Down |
| AC011448.1 | 0.004 | 1.783 | Down |
| SLC2A5 | 0.019 | 1.784 | Down |
| CATG00000055416.1 | 0.027 | 1.784 | Down |
| PPIL2 | 0.006 | 1.785 | Up |
| PTCD3 | 0.011 | 1.785 | Down |
| ST20-MTHFS | 0.029 | 1.786 | Down |
| RWDD2A | 0.022 | 1.787 | Up |
| RNF40 | 0.003 | 1.788 | Up |
| TRIR | 0.015 | 1.788 | Up |
| AOX1 | 0.028 | 1.789 | Up |
| ZNF584 | 0.048 | 1.792 | Up |
| IDI1 | 0.004 | 1.793 | Up |
| KLHDC4 | 0.032 | 1.793 | Up |
| RANBP3 | 0.039 | 1.795 | Up |
| VAPB | 0.017 | 1.796 | Down |
| LMO3 | 0.025 | 1.796 | Down |
| CATG00000093881.1 | 0.044 | 1.799 | Up |
| NOTO | 0.040 | 1.800 | Down |
| CATG00000039609.1 | 0.018 | 1.800 | Up |
| GP6 | 0.010 | 1.800 | Up |
| CPE | 0.010 | 1.801 | Down |
| OR5B21 | 0.047 | 1.803 | Down |
| CATG00000008640.1 | 0.049 | 1.804 | Down |
| ARHGEF15 | 0.006 | 1.805 | Up |

| | | | |
|---|---|---|---|
| GATD1 | 0.017 | 1.806 | Down |
| MTFP1 | 0.003 | 1.807 | Down |
| CATG00000044153.1 | 0.006 | 1.807 | Down |
| DCTN2 | 0.001 | 1.809 | Down |
| MYD88 | 0.029 | 1.809 | Up |
| NR2C2AP | 0.043 | 1.811 | Down |
| OXA1L | 0.013 | 1.811 | Down |
| CDC25C | 0.011 | 1.812 | Down |
| MRPL33 | 0.007 | 1.813 | Down |
| CATG00000109944.1 | 0.014 | 1.814 | Up |
| PCDHA3 | 0.034 | 1.819 | Up |
| REG4 | 0.035 | 1.819 | Up |
| SCN2A | 0.024 | 1.819 | Up |
| RNF141 | 0.035 | 1.820 | Down |
| EXOSC8 | 0.025 | 1.821 | Up |
| CATG00000044521.1 | 0.011 | 1.824 | Up |
| AC112205.1 | 0.026 | 1.825 | Down |
| PDYN | 0.032 | 1.825 | Up |
| AKAP11 | 0.021 | 1.826 | Down |
| AKT2 | 0.002 | 1.827 | Up |
| PLOD1 | 0.009 | 1.830 | Up |
| RIT2 | 0.014 | 1.831 | Down |
| FAM120A | 0.037 | 1.831 | Up |
| OSGIN1 | 0.023 | 1.831 | Up |
| DHRS9 | 0.020 | 1.833 | Up |
| ZNF565 | 0.034 | 1.833 | Up |
| SGCA | 0.007 | 1.838 | Down |
| BPIFA2 | 0.048 | 1.839 | Down |
| GJB5 | 0.033 | 1.840 | Up |
| TBCD | 0.040 | 1.842 | Down |
| CATG00000094618.1 | 0.007 | 1.846 | Down |
| DNAH1 | 0.028 | 1.847 | Down |
| GDAP1L1 | 0.000 | 1.848 | Up |
| QTRT1 | 0.024 | 1.849 | Down |
| SLC6A17 | 0.001 | 1.850 | Down |
| RPL17 | 0.029 | 1.850 | Up |
| LYSMD1 | 0.035 | 1.850 | Down |
| ERCC5 | 0.032 | 1.854 | Up |
| PILRB | 0.013 | 1.854 | Up |
| CATG00000074987.1 | 0.016 | 1.856 | Up |
| KIAA0355 | 0.042 | 1.856 | Up |
| CATG00000075641.1 | 0.001 | 1.857 | Up |
| SLC15A1 | 0.007 | 1.857 | Up |
| FGFR4 | 0.025 | 1.857 | Up |
| FAM243B | 0.013 | 1.859 | Down |
| ZNF48 | 0.025 | 1.860 | Down |
| RGS8 | 0.040 | 1.863 | Up |
| MED4 | 0.039 | 1.863 | Down |
| ABCD4 | 0.024 | 1.864 | Down |
| UVRAG | 0.018 | 1.865 | Up |
| CTAG1B | 0.017 | 1.865 | Up |
| MMRN1 | 0.001 | 1.868 | Up |
| CATG00000039479.1 | 0.027 | 1.868 | Down |
| WDFY4 | 0.023 | 1.871 | Up |
| DNASE2 | 0.003 | 1.871 | Down |
| SPRN | 0.030 | 1.871 | Down |
| SC5D | 0.020 | 1.872 | Up |
| AC118553.2 | 0.042 | 1.872 | Down |
| CATG00000055392.1 | 0.050 | 1.873 | Up |
| KIF16B | 0.047 | 1.873 | Down |
| PIPOX | 0.036 | 1.874 | Up |
| GOLGA3 | 0.009 | 1.874 | Up |
| ACAP1 | 0.002 | 1.875 | Up |
| EEFSEC | 0.000 | 1.876 | Up |
| HOOK1 | 0.023 | 1.877 | Up |

| | | | |
|---|---|---|---|
| AC018867.2 | 0.048 | 1.878 | Down |
| HAO1 | 0.041 | 1.879 | Down |
| CATG00000102233.1 | 0.019 | 1.880 | Down |
| BCL11A | 0.013 | 1.880 | Down |
| SHC3 | 0.042 | 1.880 | Down |
| CATG00000018214.1 | 0.013 | 1.881 | Up |
| CCDC196 | 0.044 | 1.883 | Down |
| NEO1 | 0.034 | 1.883 | Up |
| ST7L | 0.023 | 1.884 | Up |
| CATG00000092640.1 | 0.023 | 1.884 | Up |
| PRXL2C | 0.048 | 1.888 | Down |
| CATG00000026180.1 | 0.003 | 1.888 | Up |
| PCDHGA3 | 0.022 | 1.889 | Up |
| PIP | 0.024 | 1.890 | Down |
| LCN15 | 0.035 | 1.891 | Up |
| OR5W2 | 0.018 | 1.891 | Down |
| TP53RK | 0.037 | 1.892 | Up |
| PLIN3 | 0.033 | 1.894 | Down |
| DUSP14 | 0.037 | 1.897 | Down |
| CATG00000033583.1 | 0.038 | 1.897 | Down |
| RAG1 | 0.044 | 1.900 | Down |
| ZNF670-ZNF695 | 0.002 | 1.901 | Up |
| AC037459.1 | 0.019 | 1.901 | Up |
| PPP1R7 | 0.007 | 1.902 | Down |
| HAUS2 | 0.024 | 1.904 | Down |
| CATG00000041282.1 | 0.042 | 1.904 | Down |
| CPSF4L | 0.021 | 1.904 | Up |
| IGF1 | 0.010 | 1.904 | Down |
| NHP2 | 0.032 | 1.905 | Up |
| FAM135A | 0.010 | 1.908 | Up |
| KCNN3 | 0.039 | 1.910 | Down |
| CBLN2 | 0.028 | 1.913 | Down |
| PID1 | 0.049 | 1.914 | Up |
| ERBIN | 0.039 | 1.914 | Down |
| OR2C1 | 0.041 | 1.915 | Up |
| OR7G2 | 0.041 | 1.919 | Down |
| FOXD2 | 0.008 | 1.919 | Down |
| TRMT6 | 0.008 | 1.923 | Up |
| SIMC1 | 0.049 | 1.924 | Up |
| AZIN2 | 0.032 | 1.924 | Up |
| PRMT8 | 0.005 | 1.925 | Down |
| PGAP2 | 0.042 | 1.925 | Up |
| HYPK | 0.042 | 1.926 | Down |
| XAGE3 | 0.048 | 1.927 | Down |
| TANC2 | 0.000 | 1.928 | Up |
| GLI4 | 0.002 | 1.928 | Up |
| TUBA4A | 0.044 | 1.928 | Up |
| OR8J3 | 0.026 | 1.932 | Down |
| TAS2R60 | 0.030 | 1.932 | Down |
| SIGLEC12 | 0.014 | 1.932 | Down |
| CHN2 | 0.001 | 1.935 | Down |
| ESRRG | 0.040 | 1.936 | Down |
| EPN3 | 0.043 | 1.940 | Down |
| SLC22A23 | 0.017 | 1.941 | Down |
| TMEM109 | 0.007 | 1.943 | Up |
| TRMT10B | 0.016 | 1.947 | Down |
| CATG00000090636.1 | 0.002 | 1.948 | Up |
| PHOX2B | 0.017 | 1.949 | Up |
| CATG00000019706.1 | 0.012 | 1.950 | Up |
| PLK4 | 0.033 | 1.950 | Up |
| GAPVD1 | 0.004 | 1.951 | Up |
| C9orf135 | 0.035 | 1.952 | Up |
| TBCEL | 0.035 | 1.952 | Down |
| ECHS1 | 0.014 | 1.952 | Down |
| HIST1H2BC | 0.041 | 1.956 | Up |

| | | | |
|---|---|---|---|
| PDCL3 | 0.022 | 1.956 | Up |
| ATP6V1B2 | 0.003 | 1.957 | Up |
| IFT80 | 0.007 | 1.960 | Up |
| APOA4 | 0.048 | 1.962 | Down |
| HCAR1 | 0.022 | 1.963 | Down |
| FXYD3 | 0.035 | 1.965 | Up |
| CATG00000033941.1 | 0.030 | 1.965 | Down |
| HS6ST3 | 0.023 | 1.966 | Up |
| SPRED3 | 0.005 | 1.966 | Up |
| EPHA2 | 0.014 | 1.967 | Up |
| PTPN11 | 0.015 | 1.968 | Up |
| RNF169 | 0.022 | 1.969 | Up |
| EXOC3L2 | 0.008 | 1.974 | Up |
| C17orf50 | 0.005 | 1.978 | Down |
| CSTF1 | 0.000 | 1.978 | Down |
| KRBA1 | 0.005 | 1.979 | Up |
| AGFG2 | 0.042 | 1.986 | Down |
| CATG00000094958.1 | 0.031 | 1.986 | Down |
| ANTXRL | 0.032 | 1.989 | Up |
| CYP4A22 | 0.019 | 1.989 | Down |
| MRPL30 | 0.006 | 1.990 | Down |
| CATG00000043838.1 | 0.046 | 1.990 | Up |
| FCN2 | 0.002 | 1.991 | Up |
| SLC7A14 | 0.037 | 1.993 | Down |
| ARHGAP23 | 0.020 | 1.994 | Down |
| INTS10 | 0.039 | 2.000 | Up |
| MVB12A | 0.036 | 2.000 | Down |
| AL353354.2 | 0.047 | 2.001 | Down |
| SLC22A15 | 0.037 | 2.004 | Down |
| GAS2L3 | 0.022 | 2.007 | Down |
| UHRF2 | 0.003 | 2.007 | Down |
| MYH7B | 0.027 | 2.007 | Up |
| TMEM206 | 0.022 | 2.014 | Up |
| PCDHGA9 | 0.042 | 2.016 | Down |
| ANTXR2 | 0.009 | 2.018 | Down |
| EFL1 | 0.031 | 2.019 | Down |
| CATG00000047496.1 | 0.013 | 2.023 | Up |
| ME1 | 0.009 | 2.024 | Down |
| APOL1 | 0.007 | 2.025 | Up |
| PRDX4 | 0.036 | 2.027 | Down |
| CD226 | 0.002 | 2.028 | Up |
| ZBTB26 | 0.012 | 2.028 | Down |
| PCDHA2 | 0.005 | 2.031 | Up |
| IRGQ | 0.024 | 2.031 | Down |
| CATG00000044399.1 | 0.007 | 2.033 | Down |
| C20orf203 | 0.013 | 2.033 | Down |
| CENPI | 0.029 | 2.034 | Down |
| CATG00000020173.1 | 0.033 | 2.035 | Up |
| HIGD1C | 0.012 | 2.035 | Down |
| SLC4A4 | 0.009 | 2.035 | Up |
| WT1 | 0.047 | 2.035 | Up |
| CCDC74A | 0.018 | 2.038 | Down |
| CKMT1B | 0.023 | 2.038 | Down |
| ZNF624 | 0.015 | 2.039 | Down |
| CHMP3 | 0.029 | 2.040 | Up |
| FRAT2 | 0.035 | 2.040 | Down |
| IL23R | 0.002 | 2.043 | Down |
| WNT11 | 0.001 | 2.043 | Up |
| DHRS4L2 | 0.036 | 2.047 | Down |
| TBC1D30 | 0.042 | 2.047 | Down |
| FAM153C | 0.017 | 2.047 | Down |
| PRKCI | 0.001 | 2.048 | Up |
| HTD2 | 0.016 | 2.050 | Down |
| STAT3 | 0.005 | 2.050 | Up |
| CATG00000043259.1 | 0.004 | 2.053 | Down |

| | | | |
|---|---|---|---|
| PSMA2 | 0.029 | 2.054 | Down |
| XK | 0.038 | 2.058 | Down |
| SLC39A8 | 0.036 | 2.062 | Down |
| CATG00000060386.1 | 0.003 | 2.063 | Up |
| CYP27B1 | 0.028 | 2.070 | Down |
| POLM | 0.017 | 2.073 | Up |
| RBMX | 0.013 | 2.074 | Up |
| GJE1 | 0.034 | 2.076 | Down |
| NUP188 | 0.034 | 2.080 | Up |
| NCOA1 | 0.012 | 2.081 | Down |
| ZNF567 | 0.025 | 2.086 | Down |
| C3orf33 | 0.007 | 2.087 | Down |
| TMEM8B | 0.002 | 2.096 | Down |
| CATG00000038360.1 | 0.034 | 2.097 | Down |
| CEP126 | 0.004 | 2.098 | Up |
| QSOX1 | 0.030 | 2.099 | Down |
| TPX2 | 0.049 | 2.101 | Down |
| H2AFZ | 0.003 | 2.103 | Up |
| AES | 0.046 | 2.104 | Up |
| SELENON | 0.025 | 2.104 | Down |
| PGBD2 | 0.025 | 2.107 | Up |
| PGAP1 | 0.029 | 2.111 | Down |
| FN3K | 0.033 | 2.119 | Up |
| ZNF496 | 0.015 | 2.120 | Up |
| SLC9A2 | 0.011 | 2.123 | Up |
| NDUFA7 | 0.012 | 2.123 | Down |
| GSPT1 | 0.019 | 2.125 | Down |
| FTH1 | 0.036 | 2.126 | Down |
| PLAC8L1 | 0.020 | 2.129 | Up |
| KCTD8 | 0.000 | 2.129 | Up |
| KLRD1 | 0.004 | 2.130 | Up |
| CATG00000053945.1 | 0.022 | 2.132 | Up |
| PTCHD1 | 0.022 | 2.137 | Down |
| GPR15 | 0.022 | 2.137 | Down |
| DHPS | 0.040 | 2.140 | Down |
| TANK | 0.013 | 2.146 | Up |
| DMKN | 0.026 | 2.151 | Up |
| AC010877.1 | 0.040 | 2.153 | Down |
| SPON2 | 0.042 | 2.161 | Down |
| APC2 | 0.014 | 2.162 | Down |
| DSG3 | 0.011 | 2.165 | Down |
| ETV1 | 0.015 | 2.165 | Up |
| CFI | 0.040 | 2.169 | Down |
| UBE2G2 | 0.019 | 2.169 | Down |
| NBPF9 | 0.015 | 2.171 | Down |
| CEBPG | 0.010 | 2.172 | Up |
| MFSD6 | 0.034 | 2.174 | Down |
| REEP4 | 0.043 | 2.175 | Down |
| AC104389.5 | 0.018 | 2.175 | Up |
| TCAP | 0.049 | 2.176 | Up |
| SMARCA5 | 0.038 | 2.176 | Up |
| TCHH | 0.049 | 2.177 | Down |
| C19orf85 | 0.042 | 2.178 | Down |
| AFMID | 0.039 | 2.178 | Down |
| ARHGAP8 | 0.036 | 2.191 | Down |
| CATG00000040081.1 | 0.022 | 2.194 | Down |
| MFF | 0.007 | 2.197 | Up |
| BCAN | 0.031 | 2.200 | Up |
| ANKRD36B | 0.014 | 2.205 | Up |
| ZNF28 | 0.001 | 2.210 | Down |
| WFDC11 | 0.013 | 2.211 | Up |
| DNAJC17 | 0.044 | 2.212 | Up |
| OR2A2 | 0.013 | 2.214 | Down |
| ZNF131 | 0.030 | 2.215 | Down |
| HIST1H4A | 0.040 | 2.215 | Up |

| | | | |
|---|---|---|---|
| GC | 0.044 | 2.215 | Down |
| NLRP6 | 0.034 | 2.220 | Up |
| INSL3 | 0.009 | 2.220 | Down |
| COCH | 0.022 | 2.222 | Down |
| YIPF4 | 0.029 | 2.223 | Down |
| CACNB2 | 0.031 | 2.226 | Down |
| IL17RC | 0.015 | 2.228 | Down |
| CPSF2 | 0.025 | 2.231 | Up |
| ZNF726 | 0.029 | 2.239 | Up |
| MEGF9 | 0.038 | 2.241 | Up |
| KNL1 | 0.016 | 2.244 | Down |
| CIDEA | 0.008 | 2.244 | Up |
| TFAP2C | 0.039 | 2.249 | Down |
| BYSL | 0.015 | 2.251 | Down |
| ZNF638 | 0.048 | 2.252 | Down |
| SERINC1 | 0.030 | 2.254 | Down |
| SOWAHC | 0.024 | 2.260 | Down |
| C4orf3 | 0.049 | 2.263 | Up |
| TFF1 | 0.009 | 2.263 | Down |
| PRH1-PRR4 | 0.041 | 2.264 | Down |
| PGP | 0.029 | 2.268 | Down |
| ALG1L | 0.037 | 2.272 | Down |
| PARD6B | 0.030 | 2.273 | Up |
| MMADHC | 0.030 | 2.276 | Down |
| EXD2 | 0.041 | 2.277 | Down |
| CCDC124 | 0.024 | 2.278 | Down |
| ATF7IP2 | 0.045 | 2.282 | Down |
| PTPN7 | 0.031 | 2.283 | Down |
| CATG00000028059.1 | 0.011 | 2.286 | Down |
| SRPK2 | 0.017 | 2.286 | Down |
| CATG00000110256.1 | 0.030 | 2.287 | Down |
| OTUD4 | 0.011 | 2.289 | Down |
| DDTL | 0.019 | 2.292 | Up |
| AGO4 | 0.009 | 2.295 | Up |
| CATG00000056958.1 | 0.027 | 2.295 | Down |
| FRMPD2 | 0.028 | 2.296 | Up |
| FAM198B | 0.036 | 2.298 | Up |
| RPL26 | 0.047 | 2.305 | Up |
| OR4K5 | 0.000 | 2.310 | Down |
| CATG00000071673.1 | 0.039 | 2.312 | Down |
| CBLB | 0.007 | 2.317 | Down |
| C16orf72 | 0.030 | 2.318 | Down |
| CATG00000020284.1 | 0.021 | 2.323 | Up |
| NANOG | 0.000 | 2.323 | Down |
| KRTAP10-10 | 0.002 | 2.327 | Down |
| SAA1 | 0.029 | 2.334 | Up |
| COQ7 | 0.037 | 2.341 | Up |
| PGPEP1L | 0.028 | 2.341 | Down |
| CATG00000096107.1 | 0.022 | 2.344 | Up |
| AGAP8 | 0.043 | 2.350 | Up |
| KIF13A | 0.022 | 2.350 | Down |
| CATG00000107439.1 | 0.023 | 2.351 | Up |
| FABP2 | 0.045 | 2.355 | Up |
| PURB | 0.027 | 2.362 | Down |
| TMED2 | 0.027 | 2.363 | Down |
| GLYATL2 | 0.003 | 2.366 | Down |
| TMEM161B | 0.028 | 2.367 | Up |
| DPH5 | 0.014 | 2.371 | Up |
| ARMCX5 | 0.024 | 2.372 | Down |
| CEP170 | 0.032 | 2.378 | Down |
| CLIP2 | 0.028 | 2.379 | Down |
| LMNB1 | 0.014 | 2.395 | Down |
| HIST1H3C | 0.037 | 2.401 | Up |
| DNAJC30 | 0.020 | 2.401 | Down |
| KRTAP19-5 | 0.040 | 2.402 | Down |

| | | | |
|---|---|---|---|
| MXD3 | 0.007 | 2.403 | Down |
| CATG00000107414.1 | 0.016 | 2.405 | Down |
| GPR183 | 0.041 | 2.405 | Down |
| RTP2 | 0.028 | 2.405 | Down |
| C3orf38 | 0.010 | 2.407 | Down |
| EIF4E | 0.011 | 2.412 | Down |
| NAAA | 0.003 | 2.414 | Down |
| TRIM54 | 0.049 | 2.414 | Down |
| CCNG2 | 0.012 | 2.418 | Down |
| SAMD12 | 0.049 | 2.419 | Up |
| CLEC10A | 0.030 | 2.424 | Up |
| GBF1 | 0.049 | 2.428 | Up |
| RGR | 0.024 | 2.433 | Up |
| CATG00000016282.1 | 0.006 | 2.436 | Up |
| METTL7A | 0.020 | 2.446 | Up |
| SEPT6 | 0.009 | 2.450 | Up |
| SLIT2 | 0.044 | 2.454 | Down |
| RPP30 | 0.017 | 2.454 | Down |
| CATG00000040193.1 | 0.038 | 2.458 | Down |
| STMN4 | 0.018 | 2.459 | Up |
| CATG00000079193.1 | 0.040 | 2.460 | Up |
| HECA | 0.033 | 2.464 | Down |
| LRRC18 | 0.010 | 2.466 | Up |
| CD99L2 | 0.028 | 2.467 | Up |
| FAM170B | 0.006 | 2.467 | Down |
| TTLL9 | 0.016 | 2.484 | Down |
| WFDC12 | 0.041 | 2.490 | Up |
| C10orf99 | 0.050 | 2.496 | Down |
| MAP2K2 | 0.015 | 2.498 | Down |
| ABHD17C | 0.018 | 2.499 | Down |
| ZNF141 | 0.017 | 2.500 | Up |
| KLF13 | 0.014 | 2.506 | Down |
| SLC36A2 | 0.029 | 2.507 | Down |
| DBX2 | 0.010 | 2.509 | Down |
| CALB2 | 0.027 | 2.509 | Up |
| ZNF718 | 0.022 | 2.510 | Down |
| ZNF275 | 0.027 | 2.511 | Down |
| PTGFR | 0.038 | 2.511 | Down |
| PSG8 | 0.004 | 2.515 | Up |
| TTC17 | 0.004 | 2.519 | Down |
| CPXM2 | 0.006 | 2.524 | Up |
| RNASE11 | 0.048 | 2.536 | Down |
| MCOLN2 | 0.042 | 2.548 | Down |
| FBXO11 | 0.010 | 2.549 | Down |
| ROR1 | 0.009 | 2.560 | Down |
| TMEFF2 | 0.018 | 2.561 | Down |
| SLC25A53 | 0.006 | 2.564 | Down |
| FBXO25 | 0.023 | 2.567 | Down |
| BAZ1B | 0.027 | 2.568 | Up |
| CALML5 | 0.017 | 2.574 | Up |
| PLK5 | 0.032 | 2.583 | Down |
| NXF3 | 0.041 | 2.585 | Down |
| CATG00000083549.1 | 0.026 | 2.585 | Up |
| EIF2S1 | 0.002 | 2.590 | Down |
| COPZ2 | 0.020 | 2.590 | Up |
| CCDC38 | 0.043 | 2.590 | Down |
| OTUD6B | 0.000 | 2.590 | Down |
| AC005549.3 | 0.045 | 2.594 | Down |
| INTS3 | 0.039 | 2.600 | Down |
| ZP4 | 0.009 | 2.608 | Down |
| KCNJ9 | 0.025 | 2.611 | Up |
| TNNC2 | 0.009 | 2.613 | Down |
| SIRT4 | 0.030 | 2.617 | Down |
| GRIN2A | 0.001 | 2.620 | Down |
| TRAF3IP2 | 0.016 | 2.621 | Down |

| | | | |
|---|---|---|---|
| PTMA | 0.038 | 2.634 | Up |
| ZBED4 | 0.026 | 2.657 | Down |
| IRF2 | 0.009 | 2.660 | Up |
| DEFA1 | 0.035 | 2.660 | Down |
| OR9Q1 | 0.018 | 2.664 | Up |
| SCYGR8 | 0.030 | 2.666 | Down |
| CAPN13 | 0.006 | 2.666 | Down |
| SAXO1 | 0.007 | 2.674 | Down |
| HDAC4 | 0.040 | 2.700 | Down |
| AP1S1 | 0.010 | 2.705 | Down |
| FAM237A | 0.040 | 2.706 | Up |
| ECD | 0.048 | 2.706 | Down |
| ZNF551 | 0.013 | 2.709 | Up |
| ATP1A1 | 0.003 | 2.709 | Down |
| NRSN1 | 0.025 | 2.712 | Down |
| GLIS2 | 0.002 | 2.714 | Down |
| ROBO2 | 0.023 | 2.719 | Up |
| THAP11 | 0.031 | 2.726 | Down |
| SERINC5 | 0.026 | 2.727 | Down |
| JMJD1C | 0.042 | 2.732 | Down |
| MRM3 | 0.015 | 2.737 | Down |
| PPY | 0.015 | 2.737 | Down |
| MRPL57 | 0.037 | 2.741 | Up |
| IRX5 | 0.008 | 2.741 | Down |
| SLC38A1 | 0.022 | 2.745 | Up |
| OR4E2 | 0.036 | 2.746 | Down |
| PSMD14 | 0.049 | 2.748 | Down |
| RELCH | 0.001 | 2.752 | Down |
| ANGPTL6 | 0.019 | 2.761 | Down |
| ZFX | 0.020 | 2.762 | Down |
| ZBTB1 | 0.021 | 2.763 | Down |
| CATG00000066043.1 | 0.037 | 2.765 | Down |
| SLC3A2 | 0.030 | 2.772 | Down |
| SLC5A1 | 0.021 | 2.779 | Down |
| CINP | 0.001 | 2.781 | Down |
| CATG00000079533.1 | 0.008 | 2.789 | Down |
| HADH | 0.035 | 2.791 | Down |
| FLJ20306 | 0.046 | 2.795 | Up |
| COA5 | 0.011 | 2.800 | Down |
| C1orf35 | 0.009 | 2.801 | Up |
| HIST1H2AC | 0.010 | 2.801 | Up |
| GJD4 | 0.042 | 2.803 | Up |
| CR769776.1 | 0.017 | 2.803 | Down |
| UBXN4 | 0.013 | 2.804 | Up |
| RTN4RL1 | 0.033 | 2.806 | Down |
| SNCG | 0.022 | 2.812 | Down |
| LTA4H | 0.026 | 2.814 | Down |
| CATG00000002338.1 | 0.005 | 2.816 | Up |
| OR51E1 | 0.019 | 2.820 | Down |
| VRK2 | 0.030 | 2.822 | Down |
| CATG00000038184.1 | 0.042 | 2.827 | Down |
| ARL6 | 0.032 | 2.834 | Up |
| CATG00000053260.1 | 0.035 | 2.841 | Down |
| COMMD5 | 0.023 | 2.845 | Up |
| BIRC8 | 0.036 | 2.857 | Down |
| MAGED2 | 0.042 | 2.862 | Up |
| STX2 | 0.029 | 2.863 | Down |
| CATG00000101230.1 | 0.003 | 2.876 | Up |
| LIPI | 0.016 | 2.880 | Down |
| AL136531.1 | 0.039 | 2.881 | Up |
| OAZ2 | 0.030 | 2.892 | Down |
| KRTCAP3 | 0.039 | 2.902 | Down |
| SET | 0.021 | 2.902 | Down |
| OTUB2 | 0.027 | 2.905 | Down |
| NDUFA10 | 0.007 | 2.915 | Down |

| | | | |
|---|---|---|---|
| SH3RF3 | 0.005 | 2.917 | Down |
| ARHGAP17 | 0.001 | 2.920 | Down |
| FSTL1 | 0.019 | 2.927 | Down |
| PTPRN2 | 0.019 | 2.928 | Down |
| AL121845.3 | 0.029 | 2.937 | Up |
| ANAPC5 | 0.049 | 2.941 | Down |
| ROM1 | 0.016 | 2.941 | Up |
| RAB28 | 0.020 | 2.944 | Down |
| HSD17B13 | 0.006 | 2.946 | Down |
| SUOX | 0.008 | 2.948 | Down |
| RRP8 | 0.044 | 2.959 | Up |
| HTR7 | 0.009 | 2.963 | Down |
| CATG00000022091.1 | 0.023 | 2.972 | Down |
| TBR1 | 0.009 | 2.977 | Up |
| NAV1 | 0.035 | 2.983 | Up |
| UQCRQ | 0.019 | 3.000 | Down |
| CPNE7 | 0.026 | 3.007 | Down |
| CHST1 | 0.003 | 3.012 | Down |
| MXD4 | 0.028 | 3.015 | Down |
| MTRNR2L11 | 0.038 | 3.015 | Up |
| NT5C2 | 0.041 | 3.022 | Down |
| TSTD3 | 0.041 | 3.025 | Down |
| SPRY3 | 0.037 | 3.031 | Down |
| MED18 | 0.023 | 3.031 | Up |
| ATP6V1G2-DDX39E | 0.035 | 3.032 | Down |
| ITGBL1 | 0.025 | 3.036 | Down |
| HLA-B | 0.009 | 3.037 | Down |
| HEMGN | 0.023 | 3.043 | Down |
| ARNT | 0.025 | 3.044 | Down |
| RAPGEF5 | 0.012 | 3.046 | Down |
| FDFT1 | 0.016 | 3.050 | Up |
| MPO | 0.039 | 3.060 | Down |
| CATG00000079740.1 | 0.007 | 3.060 | Up |
| AP5S1 | 0.029 | 3.064 | Down |
| CHMP7 | 0.011 | 3.070 | Down |
| TSHZ3 | 0.024 | 3.072 | Down |
| PCBD2 | 0.001 | 3.075 | Down |
| RHAG | 0.029 | 3.078 | Down |
| PPARA | 0.014 | 3.085 | Up |
| VN1R1 | 0.046 | 3.098 | Up |
| PRKCA | 0.015 | 3.098 | Down |
| SLC6A16 | 0.010 | 3.103 | Up |
| PRKCQ | 0.020 | 3.104 | Down |
| CATG00000057844.1 | 0.032 | 3.109 | Down |
| RPL29 | 0.028 | 3.112 | Down |
| CATG00000002792.1 | 0.009 | 3.126 | Up |
| C11orf58 | 0.025 | 3.128 | Down |
| SYT17 | 0.042 | 3.128 | Down |
| SLC35F6 | 0.006 | 3.138 | Up |
| KCNJ8 | 0.022 | 3.147 | Down |
| AL596220.1 | 0.021 | 3.160 | Down |
| CBWD2 | 0.023 | 3.167 | Up |
| CATG00000050657.1 | 0.017 | 3.179 | Up |
| MT-CYB | 0.024 | 3.182 | Up |
| KRBOX1 | 0.026 | 3.186 | Down |
| FIS1 | 0.012 | 3.188 | Down |
| PRSS55 | 0.027 | 3.199 | Down |
| TPP1 | 0.006 | 3.200 | Down |
| XPO5 | 0.001 | 3.202 | Down |
| TRARG1 | 0.009 | 3.204 | Down |
| MTTP | 0.045 | 3.208 | Down |
| ZNF727 | 0.006 | 3.212 | Up |
| ZDHHC9 | 0.011 | 3.214 | Down |
| OVCH2 | 0.029 | 3.249 | Up |
| TBCCD1 | 0.047 | 3.260 | Down |

| | | | |
|---|---|---|---|
| SLC25A14 | 0.023 | 3.276 | Up |
| MMAA | 0.018 | 3.276 | Down |
| NMU | 0.001 | 3.279 | Down |
| CATG00000022646.1 | 0.019 | 3.283 | Down |
| PKP2 | 0.021 | 3.293 | Up |
| RXYLT1 | 0.001 | 3.298 | Up |
| SEC24B | 0.015 | 3.302 | Down |
| COQ8A | 0.018 | 3.328 | Down |
| GJC1 | 0.021 | 3.332 | Down |
| AC005477.1 | 0.013 | 3.359 | Down |
| PKD2 | 0.013 | 3.360 | Down |
| PREX1 | 0.012 | 3.362 | Down |
| ZNF664 | 0.005 | 3.369 | Up |
| MPHOSPH6 | 0.023 | 3.380 | Down |
| ZNF599 | 0.031 | 3.383 | Down |
| MRPS16 | 0.027 | 3.397 | Up |
| GPR137C | 0.004 | 3.400 | Down |
| MAU2 | 0.022 | 3.404 | Down |
| RPS24 | 0.021 | 3.409 | Up |
| OTUD6A | 0.030 | 3.411 | Up |
| CFAP97D2 | 0.036 | 3.411 | Down |
| PHKG2 | 0.042 | 3.417 | Down |
| GRK7 | 0.037 | 3.419 | Up |
| JMJD4 | 0.016 | 3.423 | Down |
| CATG00000068829.1 | 0.045 | 3.428 | Up |
| BICDL2 | 0.007 | 3.436 | Down |
| RPL30 | 0.046 | 3.438 | Down |
| MACF1 | 0.036 | 3.442 | Down |
| CATG00000058915.1 | 0.018 | 3.455 | Up |
| COPS9 | 0.004 | 3.466 | Down |
| FRZB | 0.005 | 3.474 | Down |
| JAM2 | 0.026 | 3.493 | Up |
| S100G | 0.001 | 3.512 | Down |
| STX19 | 0.021 | 3.533 | Down |
| ATR | 0.017 | 3.540 | Down |
| ADD1 | 0.017 | 3.556 | Down |
| TMEM217 | 0.043 | 3.558 | Up |
| CATG00000056954.1 | 0.026 | 3.559 | Up |
| RPS10-NUDT3 | 0.030 | 3.559 | Up |
| MPZL1 | 0.010 | 3.584 | Down |
| SLC30A2 | 0.024 | 3.587 | Down |
| ATOH7 | 0.009 | 3.610 | Down |
| CASD1 | 0.013 | 3.622 | Down |
| CCDC63 | 0.029 | 3.632 | Down |
| SNX11 | 0.031 | 3.633 | Down |
| IFIT1B | 0.017 | 3.636 | Down |
| SERF1A | 0.037 | 3.639 | Up |
| ALDH6A1 | 0.040 | 3.640 | Up |
| LNX2 | 0.013 | 3.641 | Down |
| RNASEH2A | 0.023 | 3.644 | Up |
| UFC1 | 0.043 | 3.645 | Up |
| CATG00000035216.1 | 0.012 | 3.650 | Up |
| IRAK1BP1 | 0.016 | 3.656 | Up |
| SSR1 | 0.003 | 3.659 | Down |
| HIST3H2A | 0.031 | 3.674 | Down |
| OCLM | 0.034 | 3.704 | Down |
| ANKRD28 | 0.048 | 3.704 | Up |
| PQBP1 | 0.004 | 3.705 | Down |
| HBA2 | 0.047 | 3.727 | Up |
| CATG00000092121.1 | 0.004 | 3.735 | Up |
| TM4SF19 | 0.036 | 3.749 | Up |
| NAPB | 0.039 | 3.752 | Down |
| BRD3 | 0.024 | 3.763 | Down |
| POGK | 0.008 | 3.766 | Up |
| TMEM242 | 0.016 | 3.767 | Down |

| | | | |
|---|---|---|---|
| SPATA12 | 0.014 | 3.768 | Down |
| CATG00000004584.1 | 0.032 | 3.773 | Up |
| INSL4 | 0.017 | 3.791 | Down |
| FAM171A2 | 0.010 | 3.795 | Down |
| RBM39 | 0.006 | 3.806 | Down |
| NPVF | 0.006 | 3.808 | Down |
| KRT40 | 0.033 | 3.810 | Down |
| ZRANB2 | 0.047 | 3.810 | Down |
| AMMECR1 | 0.010 | 3.827 | Down |
| GNAI1 | 0.003 | 3.846 | Down |
| CLDN6 | 0.003 | 3.847 | Up |
| PRSS21 | 0.039 | 3.863 | Down |
| KPNA1 | 0.006 | 3.878 | Down |
| CATG00000023238.1 | 0.008 | 3.880 | Down |
| SHB | 0.029 | 3.887 | Down |
| ATRX | 0.013 | 3.895 | Down |
| FBXO27 | 0.021 | 3.903 | Up |
| ZBTB12 | 0.038 | 3.904 | Down |
| NUP43 | 0.015 | 3.915 | Up |
| IL13 | 0.031 | 3.924 | Down |
| HNRNPA1 | 0.045 | 3.961 | Up |
| PROKR1 | 0.018 | 3.974 | Down |
| IZUMO2 | 0.040 | 4.000 | Down |
| TSTD1 | 0.031 | 4.009 | Down |
| NDUFB1 | 0.049 | 4.027 | Down |
| JPT2 | 0.019 | 4.029 | Up |
| CATG00000060946.1 | 0.001 | 4.035 | Up |
| LIN52 | 0.025 | 4.043 | Down |
| BMP7 | 0.004 | 4.045 | Down |
| ETV3L | 0.044 | 4.048 | Down |
| CCDC61 | 0.038 | 4.061 | Up |
| OR4D10 | 0.010 | 4.061 | Down |
| CPED1 | 0.011 | 4.065 | Down |
| FCRL2 | 0.010 | 4.068 | Down |
| H3F3C | 0.021 | 4.070 | Up |
| CALHM1 | 0.010 | 4.095 | Up |
| SCNN1D | 0.012 | 4.103 | Up |
| BRDT | 0.022 | 4.117 | Down |
| PTDSS1 | 0.020 | 4.135 | Down |
| HPS6 | 0.025 | 4.147 | Down |
| HASPIN | 0.040 | 4.161 | Up |
| ERVV-2 | 0.004 | 4.173 | Up |
| LECT2 | 0.005 | 4.177 | Up |
| NAA16 | 0.035 | 4.199 | Up |
| SIGLEC11 | 0.038 | 4.218 | Up |
| CATG00000008569.1 | 0.002 | 4.222 | Down |
| PF4V1 | 0.013 | 4.230 | Up |
| GIMAP8 | 0.008 | 4.280 | Up |
| CREB3L2 | 0.019 | 4.283 | Down |
| EIF4E1B | 0.004 | 4.284 | Down |
| AC132872.2 | 0.036 | 4.301 | Up |
| CSN1S1 | 0.047 | 4.302 | Down |
| CATG00000098834.1 | 0.001 | 4.314 | Up |
| TDG | 0.036 | 4.314 | Down |
| LIMS1 | 0.021 | 4.321 | Up |
| CATG00000062558.1 | 0.006 | 4.330 | Up |
| BAG1 | 0.045 | 4.343 | Down |
| MT-ND3 | 0.024 | 4.353 | Up |
| BHLHE22 | 0.033 | 4.359 | Down |
| RPS29 | 0.020 | 4.370 | Up |
| BTF3 | 0.034 | 4.371 | Up |
| USH2A | 0.032 | 4.371 | Down |
| CATG00000113188.1 | 0.009 | 4.384 | Down |
| ANP32B | 0.039 | 4.384 | Up |
| R3HDML | 0.005 | 4.419 | Up |

| | | | |
|---|---|---|---|
| C2orf80 | 0.025 | 4.432 | Down |
| RBM38 | 0.010 | 4.468 | Up |
| AC079210.1 | 0.018 | 4.469 | Up |
| KIAA0586 | 0.034 | 4.475 | Down |
| HBEGF | 0.013 | 4.482 | Down |
| TXNRD1 | 0.008 | 4.486 | Down |
| DCAF4L1 | 0.004 | 4.509 | Down |
| HILPDA | 0.023 | 4.513 | Up |
| BRD3OS | 0.014 | 4.527 | Down |
| FAM177B | 0.023 | 4.527 | Up |
| BOD1 | 0.002 | 4.528 | Down |
| VEGFC | 0.006 | 4.529 | Down |
| HOXB4 | 0.003 | 4.535 | Down |
| RBM14-RBM4 | 0.001 | 4.536 | Down |
| TOR1A | 0.030 | 4.547 | Down |
| SLC4A5 | 0.015 | 4.554 | Up |
| CLEC1A | 0.004 | 4.578 | Down |
| C12orf60 | 0.043 | 4.583 | Down |
| CAPZA3 | 0.015 | 4.593 | Down |
| GOLGA7 | 0.003 | 4.601 | Down |
| MAP1LC3C | 0.006 | 4.631 | Down |
| AL645922.1 | 0.012 | 4.641 | Up |
| KRT2 | 0.023 | 4.644 | Down |
| CATG00000114024.1 | 0.022 | 4.668 | Up |
| RYR2 | 0.040 | 4.670 | Down |
| DDX55 | 0.016 | 4.682 | Up |
| PJA1 | 0.046 | 4.699 | Down |
| DEFB129 | 0.025 | 4.701 | Down |
| TFAP2E | 0.010 | 4.724 | Down |
| AC110615.1 | 0.032 | 4.738 | Down |
| CATG00000040303.1 | 0.002 | 4.747 | Up |
| DRD1 | 0.005 | 4.761 | Down |
| TSFM | 0.013 | 4.770 | Up |
| AX747191 | 0.011 | 4.780 | Up |
| LGALS8 | 0.006 | 4.782 | Down |
| MAGEB3 | 0.012 | 4.786 | Up |
| CATG00000005569.1 | 0.025 | 4.830 | Up |
| OR52E5 | 0.003 | 4.837 | Down |
| TMA7 | 0.027 | 4.842 | Up |
| ARGLU1 | 0.012 | 4.847 | Down |
| TMA7 | 0.034 | 4.859 | Up |
| THSD7A | 0.005 | 4.860 | Down |
| IMMP2L | 0.010 | 4.864 | Down |
| SLC12A7 | 0.018 | 4.898 | Down |
| MNDA | 0.002 | 4.906 | Down |
| FUZ | 0.039 | 4.906 | Down |
| ENOSF1 | 0.022 | 4.912 | Down |
| CATG00000038121.1 | 0.023 | 4.921 | Down |
| CATG00000027660.1 | 0.019 | 4.925 | Up |
| NFRKB | 0.050 | 4.957 | Down |
| PRLHR | 0.021 | 5.008 | Down |
| PNPT1 | 0.015 | 5.051 | Up |
| RBMS2 | 0.000 | 5.129 | Down |
| ALKBH4 | 0.013 | 5.134 | Up |
| MFSD14C | 0.014 | 5.145 | Up |
| CYB561D1 | 0.018 | 5.155 | Up |
| STOML3 | 0.015 | 5.163 | Down |
| CATG00000027020.1 | 0.010 | 5.183 | Up |
| ZNF440 | 0.022 | 5.198 | Up |
| ZNF3 | 0.013 | 5.208 | Up |
| MUT | 0.012 | 5.212 | Down |
| RPS4X | 0.009 | 5.215 | Up |
| MARCKSL1 | 0.022 | 5.252 | Up |
| RPL22 | 0.010 | 5.253 | Down |
| P2RY8 | 0.015 | 5.273 | Up |

| | | | |
|---|---|---|---|
| IRX2 | 0.011 | 5.278 | Down |
| ZNF517 | 0.047 | 5.284 | Up |
| LRRC39 | 0.001 | 5.291 | Down |
| ELMOD3 | 0.020 | 5.327 | Up |
| CPA6 | 0.006 | 5.331 | Down |
| OR8G1 | 0.031 | 5.340 | Up |
| CATG00000020281.1 | 0.023 | 5.353 | Up |
| CNGB1 | 0.007 | 5.438 | Up |
| AC135983.2 | 0.012 | 5.452 | Up |
| RDX | 0.026 | 5.461 | Down |
| GFI1B | 0.012 | 5.472 | Up |
| CDK2AP2 | 0.004 | 5.481 | Down |
| PRTFDC1 | 0.008 | 5.569 | Down |
| SMIM33 | 0.032 | 5.638 | Up |
| NDUFA11 | 0.036 | 5.641 | Up |
| IL17D | 0.031 | 5.653 | Down |
| AC132186.1 | 0.008 | 5.695 | Up |
| RPL26 | 0.014 | 5.699 | Up |
| ZNF641 | 0.047 | 5.732 | Down |
| LARP4 | 0.016 | 5.771 | Down |
| HTRA4 | 0.025 | 5.785 | Up |
| TOP3A | 0.022 | 5.787 | Up |
| IPP | 0.012 | 5.800 | Up |
| ZNF765 | 0.032 | 5.850 | Down |
| ACOT4 | 0.011 | 5.858 | Up |
| CD8A | 0.026 | 5.885 | Up |
| IRS2 | 0.006 | 5.936 | Down |
| EFEMP2 | 0.004 | 5.937 | Up |
| ZNF526 | 0.012 | 5.954 | Up |
| PTCD1 | 0.007 | 5.968 | Up |
| ODF2L | 0.021 | 5.988 | Down |
| ZNF600 | 0.005 | 6.013 | Up |
| EGR1 | 0.001 | 6.040 | Down |
| STAT6 | 0.006 | 6.043 | Down |
| COMMD2 | 0.013 | 6.101 | Up |
| NUDT11 | 0.014 | 6.121 | Down |
| POLR2E | 0.011 | 6.134 | Up |
| GDE1 | 0.021 | 6.135 | Down |
| RPL17-C18orf32 | 0.014 | 6.142 | Up |
| KIAA1658 | 0.027 | 6.155 | Down |
| SOCS6 | 0.049 | 6.171 | Down |
| CATG00000040897.1 | 0.018 | 6.196 | Up |
| CATG00000068089.1 | 0.001 | 6.219 | Up |
| PCLAF | 0.007 | 6.258 | Up |
| PPP1R3E | 0.046 | 6.269 | Down |
| OR10H1 | 0.014 | 6.313 | Up |
| PCCB | 0.004 | 6.326 | Up |
| BHMT2 | 0.007 | 6.419 | Up |
| CCL8 | 0.006 | 6.435 | Down |
| CATG00000071558.1 | 0.009 | 6.545 | Up |
| DDX52 | 0.008 | 6.551 | Up |
| MTDH | 0.004 | 6.580 | Up |
| PLEKHA8 | 0.049 | 6.595 | Down |
| CATG00000112207.1 | 0.009 | 6.615 | Up |
| QDPR | 0.021 | 6.672 | Down |
| PGAM5 | 0.004 | 6.685 | Up |
| MPLKIP | 0.003 | 6.726 | Down |
| AC016586.1 | 0.007 | 6.740 | Up |
| CATG00000040980.1 | 0.009 | 6.767 | Up |
| OTP | 0.013 | 6.786 | Down |
| HIST3H3 | 0.002 | 6.792 | Up |
| MARVELD3 | 0.008 | 6.847 | Up |
| RPS19 | 0.012 | 6.868 | Up |
| YY2 | 0.008 | 6.896 | Up |
| CATG00000049183.1 | 0.009 | 6.976 | Up |

| | | | |
|---|---|---|---|
| CATG00000063098.1 | 0.007 | 7.035 | Up |
| AC015802.6 | 0.024 | 7.049 | Up |
| CATG00000086904.1 | 0.037 | 7.061 | Down |
| TRIM13 | 0.008 | 7.063 | Up |
| CATG00000083857.1 | 0.020 | 7.103 | Down |
| SRRM4 | 0.004 | 7.132 | Down |
| FBXO6 | 0.010 | 7.175 | Up |
| ANKS3 | 0.008 | 7.241 | Up |
| IDE | 0.006 | 7.280 | Down |
| CPNE3 | 0.045 | 7.281 | Down |
| EXOC8 | 0.001 | 7.309 | Down |
| CHMP4C | 0.029 | 7.314 | Down |
| MTRNR2L5 | 0.035 | 7.336 | Up |
| ZNF69 | 0.007 | 7.342 | Up |
| VHL | 0.011 | 7.374 | Up |
| AX747379 | 0.005 | 7.398 | Up |
| OPA3 | 0.005 | 7.428 | Up |
| AHSG | 0.005 | 7.554 | Up |
| ZSWIM1 | 0.010 | 7.603 | Up |
| FOXC2 | 0.029 | 7.654 | Down |
| AL135998.1 | 0.013 | 7.673 | Up |
| ACER1 | 0.007 | 7.710 | Up |
| TMEM169 | 0.009 | 7.736 | Down |
| SNUPN | 0.016 | 7.752 | Up |
| BC150535 | 0.016 | 7.820 | Up |
| CATG00000032923.1 | 0.012 | 7.821 | Up |
| UQCR11 | 0.000 | 7.903 | Up |
| Z98049.1 | 0.007 | 7.904 | Up |
| RPL37A | 0.002 | 7.905 | Up |
| USP33 | 0.002 | 7.923 | Up |
| MCAM | 0.023 | 7.936 | Down |
| AC093157.1 | 0.007 | 7.984 | Up |
| ALOX15B | 0.012 | 7.997 | Up |
| CBX6 | 0.019 | 8.244 | Down |
| SCYGR4 | 0.002 | 8.378 | Up |
| NEK2 | 0.018 | 8.586 | Down |
| TRAM2 | 0.024 | 8.822 | Down |
| CATG00000063086.1 | 0.003 | 8.848 | Up |
| TXNL4A | 0.023 | 8.947 | Down |
| ZNF587 | 0.049 | 8.958 | Down |
| MTRNR2L1 | 0.042 | 9.061 | Up |
| EXOC1L | 0.003 | 9.089 | Down |
| CYP4B1 | 0.023 | 9.126 | Down |
| AC025278.1 | 0.009 | 9.161 | Up |
| PRPSAP2 | 0.026 | 9.231 | Up |
| CCDC73 | 0.004 | 9.409 | Down |
| HOXC13 | 0.041 | 9.622 | Down |
| RPL26L1 | 0.003 | 9.808 | Up |
| ZNF559-ZNF177 | 0.047 | 10.275 | Up |
| CHURC1 | 0.035 | 10.561 | Up |
| HIST1H3I | 0.003 | 10.705 | Up |
| TRMT13 | 0.021 | 11.384 | Down |
| DCAF12L2 | 0.005 | 11.406 | Down |
| PTMA | 0.003 | 11.416 | Up |
| FZD5 | 0.047 | 11.462 | Down |
| ZNF587B | 0.004 | 12.534 | Up |
| RPL31 | 0.011 | 12.603 | Up |
| CTSD | 0.037 | 13.706 | Up |
| HBG2 | 0.006 | 15.163 | Up |
| MT-CO3 | 0.017 | 16.032 | Up |
| PPBP | 0.019 | 16.490 | Up |
| SLC28A3 | 0.001 | 16.609 | Down |
| FTL | 0.003 | 19.556 | Up |
| RPS25 | 0.003 | 19.869 | Up |
| RPS21 | 0.002 | 21.240 | Up |

| | | | |
|---|---|---|---|
| RPS3A | 0.002 | 29.890 | Up |
| RPL17 | 0.001 | 31.828 | Up |

**Supplementary Table 13. General anesthetic-induced dysregulated long noncoding genes (lncRNAs) in serum from pediatric patients <4 years old**

| Gene ID | P value | Fold change | Regulation |
|---|---|---|---|
| CATG00000082103.1 | 0.019 | 38.186 | Up |
| G030772 | 0.026 | 18.972 | Up |
| CATG00000016957.1 | 0.003 | 17.047 | Up |
| CRAT | 0.034 | 16.752 | Up |
| XLOC_002730 | 0.013 | 13.969 | Up |
| AC090236.2 | 0.010 | 13.484 | Up |
| AC008915.2 | 0.010 | 13.331 | Up |
| LINC02175 | 0.010 | 12.100 | Up |
| CATG00000072356.1 | 0.005 | 11.990 | Up |
| CATG00000056335.1 | 0.008 | 11.560 | Up |
| AC007114.1 | 0.009 | 11.461 | Up |
| AC022960.1 | 0.012 | 11.460 | Up |
| NKAP | 0.011 | 11.343 | Up |
| LINC01224 | 0.011 | 11.308 | Up |
| GRHPR | 0.023 | 11.093 | Up |
| LINC01786 | 0.007 | 11.070 | Up |
| AL589935.1 | 0.009 | 11.037 | Up |
| NIPAL3 | 0.008 | 10.984 | Up |
| CATG00000083353.1 | 0.008 | 10.961 | Up |
| AC114489.1 | 0.009 | 10.830 | Up |
| AC020917.2 | 0.004 | 10.718 | Up |
| AC159540.1 | 0.001 | 10.463 | Up |
| AC005323.1 | 0.003 | 10.361 | Up |
| AC105052.4 | 0.007 | 10.278 | Up |
| CATG00000048676.1 | 0.008 | 10.193 | Up |
| GJA10 | 0.014 | 10.058 | Up |
| G006392 | 0.008 | 9.854 | Up |
| AC092718.6 | 0.001 | 9.750 | Up |
| STK31 | 0.008 | 9.659 | Up |
| AL358876.2 | 0.011 | 9.512 | Up |
| G089593 | 0.041 | 9.506 | Up |
| AC037486.1 | 0.011 | 9.306 | Up |
| NUBP2 | 0.008 | 9.301 | Up |
| AX747485 | 0.007 | 9.288 | Up |
| AC104982.1 | 0.007 | 9.261 | Up |
| AL359636.1 | 0.008 | 9.177 | Up |
| LINC02157 | 0.014 | 9.171 | Up |
| GNAS | 0.033 | 9.146 | Up |
| AL162725.2 | 0.008 | 9.104 | Up |
| MNS1 | 0.015 | 9.097 | Up |
| AC010618.2 | 0.011 | 9.053 | Up |
| AL136295.2 | 0.005 | 9.051 | Up |
| CATG00000064485.1 | 0.006 | 8.945 | Up |
| CATG00000031425.1 | 0.003 | 8.926 | Up |
| CATG00000043391.1 | 0.009 | 8.925 | Up |
| AL121972.1 | 0.007 | 8.904 | Up |
| XLOC_007446 | 0.014 | 8.902 | Up |
| LINC01923 | 0.013 | 8.845 | Up |
| FRY-AS1 | 0.008 | 8.716 | Up |
| AP2A1 | 0.003 | 8.691 | Up |
| AC022558.1 | 0.010 | 8.676 | Up |
| IQCH-AS1 | 0.030 | 8.653 | Up |
| CATG00000026162.1 | 0.018 | 8.594 | Up |
| AC007922.2 | 0.013 | 8.590 | Up |
| AC027796.1 | 0.005 | 8.572 | Up |
| CATG00000085243.1 | 0.004 | 8.550 | Up |
| HAND2-AS1 | 0.007 | 8.491 | Up |
| CARMN | 0.009 | 8.448 | Up |
| LINC01503 | 0.008 | 8.447 | Up |
| AC009996.1 | 0.010 | 8.447 | Up |

| | | | |
|---|---|---|---|
| AP006248.4 | 0.013 | 8.428 | Up |
| CATG00000078660.1 | 0.015 | 8.377 | Up |
| CATG00000072758.1 | 0.010 | 8.319 | Up |
| SLC25A16 | 0.029 | 8.237 | Up |
| CSNK1G2-AS1 | 0.007 | 8.168 | Up |
| AC020656.1 | 0.011 | 8.158 | Up |
| AC011524.1 | 0.005 | 8.114 | Up |
| AC009229.2 | 0.013 | 8.109 | Up |
| CATG00000116767.1 | 0.005 | 8.035 | Up |
| CATG00000013919.1 | 0.017 | 7.997 | Up |
| AC022211.3 | 0.018 | 7.959 | Up |
| AC007036.2 | 0.008 | 7.920 | Up |
| AL590640.1 | 0.007 | 7.817 | Up |
| AL592435.1 | 0.006 | 7.803 | Up |
| AC137735.2 | 0.013 | 7.682 | Up |
| CATG00000014052.1 | 0.001 | 7.669 | Up |
| LUADT1 | 0.011 | 7.630 | Up |
| CATG00000044616.1 | 0.014 | 7.592 | Up |
| AP005328.2 | 0.008 | 7.547 | Up |
| AC092634.8 | 0.021 | 7.543 | Up |
| AC069185.1 | 0.012 | 7.532 | Up |
| AC083806.2 | 0.010 | 7.466 | Up |
| AL031722.1 | 0.013 | 7.434 | Up |
| APAF1 | 0.009 | 7.409 | Up |
| AC131097.1 | 0.023 | 7.390 | Up |
| CCNG1 | 0.009 | 7.379 | Up |
| RAB30-AS1 | 0.008 | 7.320 | Up |
| AC145207.7 | 0.009 | 7.298 | Up |
| COMMD9 | 0.006 | 7.291 | Up |
| AC007620.2 | 0.002 | 7.267 | Up |
| BIG-lncRNA-780 | 0.003 | 7.259 | Up |
| XLOC_009952 | 0.037 | 7.228 | Up |
| XLOC_l2_004592 | 0.009 | 7.217 | Up |
| C19orf70 | 0.011 | 7.216 | Up |
| AC067930.1 | 0.028 | 7.159 | Up |
| AL596325.2 | 0.010 | 7.149 | Up |
| AC073842.2 | 0.002 | 7.101 | Up |
| GABPB1-AS1 | 0.014 | 7.092 | Up |
| CATG00000057476.1 | 0.003 | 7.075 | Up |
| AP000897.1 | 0.009 | 7.061 | Up |
| CATG00000081428.1 | 0.008 | 7.048 | Up |
| AC020663.3 | 0.007 | 7.008 | Up |
| TRMT2B-AS1 | 0.007 | 7.007 | Up |
| AL031731.1 | 0.013 | 7.000 | Up |
| CATG00000053365.1 | 0.003 | 6.974 | Up |
| FLJ00277 | 0.002 | 6.973 | Up |
| AC090952.2 | 0.006 | 6.928 | Up |
| CATG00000038167.1 | 0.009 | 6.918 | Up |
| AP002336.2 | 0.017 | 6.902 | Up |
| RP11-14N7.2 | 0.026 | 6.896 | Up |
| ADAM33 | 0.018 | 6.896 | Up |
| NARF-IT1 | 0.005 | 6.882 | Up |
| GDF5OS | 0.012 | 6.872 | Up |
| AC108136.1 | 0.009 | 6.860 | Up |
| AKT3-IT1 | 0.002 | 6.849 | Up |
| NARF-AS1 | 0.004 | 6.847 | Up |
| DSCAM-AS1 | 0.025 | 6.835 | Up |
| AC004253.1 | 0.011 | 6.793 | Up |
| LINC01363 | 0.008 | 6.785 | Up |
| REV3L | 0.025 | 6.773 | Up |
| G052734 | 0.011 | 6.748 | Up |
| AC019185.2 | 0.028 | 6.716 | Up |
| AL109935.1 | 0.002 | 6.713 | Up |
| AC104135.1 | 0.015 | 6.696 | Up |
| LOC101929259 | 0.007 | 6.664 | Up |

| | | | |
|---|---|---|---|
| CATG00000094945.1 | 0.025 | 6.653 | Up |
| AC022535.1 | 0.006 | 6.639 | Up |
| AL590723.1 | 0.009 | 6.636 | Up |
| BIG-lncRNA-1361 | 0.003 | 6.626 | Up |
| BACH1-AS1 | 0.011 | 6.597 | Up |
| NFYC-AS1 | 0.017 | 6.564 | Up |
| AL136115.2 | 0.020 | 6.549 | Up |
| LINC00649 | 0.009 | 6.543 | Up |
| AL136084.2 | 0.005 | 6.514 | Up |
| LUCAT1 | 0.011 | 6.478 | Up |
| AC024075.3 | 0.008 | 6.459 | Up |
| AC069366.2 | 0.014 | 6.449 | Up |
| AC135050.1 | 0.023 | 6.445 | Up |
| CATG00000078672.1 | 0.007 | 6.424 | Up |
| RASSF8-AS1 | 0.011 | 6.408 | Up |
| CASC6 | 0.014 | 6.394 | Up |
| CATG00000009864.1 | 0.006 | 6.391 | Up |
| SEC62-AS1 | 0.002 | 6.382 | Up |
| MIATNB | 0.015 | 6.335 | Up |
| U52111.1 | 0.011 | 6.329 | Up |
| AC127496.1 | 0.013 | 6.321 | Up |
| AL161665.1 | 0.009 | 6.305 | Up |
| CATG00000110610.1 | 0.007 | 6.301 | Up |
| CATG00000054890.1 | 0.020 | 6.300 | Up |
| AL355388.2 | 0.013 | 6.297 | Up |
| LINC02447 | 0.007 | 6.292 | Up |
| RP11-216M21.7 | 0.005 | 6.288 | Up |
| CATG00000107113.1 | 0.013 | 6.281 | Up |
| LINC01285 | 0.003 | 6.240 | Up |
| PLEKHM3 | 0.004 | 6.238 | Up |
| CATG00000115220.1 | 0.028 | 6.206 | Up |
| AC005726.5 | 0.016 | 6.187 | Up |
| POLH-AS1 | 0.013 | 6.075 | Up |
| AL031714.1 | 0.026 | 6.067 | Up |
| CATG00000066195.1 | 0.005 | 6.056 | Up |
| LINC00685 | 0.008 | 6.037 | Up |
| CATG00000011925.1 | 0.004 | 6.034 | Up |
| EAF1-AS1 | 0.007 | 6.024 | Up |
| AC097103.2 | 0.019 | 6.006 | Up |
| AL157938.2 | 0.008 | 6.004 | Up |
| AL139424.3 | 0.016 | 6.000 | Up |
| AC027237.4 | 0.013 | 5.998 | Up |
| AP003171.3 | 0.023 | 5.998 | Up |
| LINC01228 | 0.019 | 5.984 | Up |
| EIF3J-DT | 0.022 | 5.980 | Up |
| CATG00000112963.1 | 0.009 | 5.966 | Up |
| AC020651.1 | 0.010 | 5.953 | Up |
| AC016907.2 | 0.012 | 5.948 | Up |
| AL049747.1 | 0.009 | 5.945 | Up |
| L29074.1 | 0.009 | 5.939 | Up |
| AC083798.2 | 0.010 | 5.927 | Up |
| AC113615.1 | 0.003 | 5.923 | Up |
| CATG00000107441.1 | 0.031 | 5.885 | Up |
| CATG00000000618.1 | 0.038 | 5.880 | Up |
| LINC00856 | 0.046 | 5.872 | Up |
| CATG00000054135.1 | 0.022 | 5.871 | Up |
| LINC01809 | 0.011 | 5.844 | Up |
| LINC01513 | 0.018 | 5.842 | Up |
| AC087721.1 | 0.010 | 5.814 | Up |
| CATG00000024710.1 | 0.010 | 5.783 | Up |
| AC016747.1 | 0.038 | 5.780 | Up |
| BIG-lncRNA-1026 | 0.007 | 5.779 | Up |
| AC011461.1 | 0.016 | 5.778 | Up |
| CATG00000061552.1 | 0.002 | 5.768 | Up |
| AC068888.2 | 0.009 | 5.761 | Up |

| | | | |
|---|---|---|---|
| EWSAT1 | 0.010 | 5.743 | Up |
| AC018362.2 | 0.015 | 5.736 | Up |
| CATG00000094248.1 | 0.036 | 5.715 | Up |
| AC005906.2 | 0.001 | 5.706 | Up |
| PROX1-AS1 | 0.010 | 5.697 | Up |
| AC092068.1 | 0.015 | 5.696 | Up |
| MIR100HG | 0.017 | 5.691 | Up |
| SNHG20 | 0.011 | 5.686 | Up |
| AC234582.1 | 0.028 | 5.676 | Up |
| FAM228B | 0.012 | 5.656 | Up |
| CATG00000054584.1 | 0.001 | 5.655 | Up |
| DNAH9 | 0.013 | 5.654 | Up |
| CATG00000109288.1 | 0.017 | 5.653 | Up |
| AL139254.1 | 0.005 | 5.644 | Up |
| LINC02337 | 0.011 | 5.624 | Up |
| AL357497.1 | 0.010 | 5.621 | Up |
| MAFG-DT | 0.005 | 5.606 | Up |
| LINC00937 | 0.008 | 5.601 | Up |
| ADIRF-AS1 | 0.011 | 5.578 | Up |
| AC117402.1 | 0.009 | 5.566 | Up |
| SLC8A1-AS1 | 0.018 | 5.550 | Up |
| CATG00000117920.1 | 0.012 | 5.544 | Up |
| AC009034.1 | 0.005 | 5.531 | Up |
| G076416 | 0.018 | 5.525 | Up |
| AC103740.1 | 0.007 | 5.524 | Up |
| CATG00000058866.1 | 0.011 | 5.522 | Up |
| CATG00000052162.1 | 0.013 | 5.517 | Up |
| AC018521.2 | 0.013 | 5.516 | Up |
| AL157396.1 | 0.001 | 5.513 | Up |
| MEG8 | 0.019 | 5.509 | Up |
| CATG00000024798.1 | 0.010 | 5.486 | Up |
| LA16c-4G1.3 | 0.010 | 5.479 | Up |
| CATG00000052772.1 | 0.049 | 5.475 | Up |
| AL139099.2 | 0.012 | 5.453 | Up |
| RP4-604K5.2 | 0.010 | 5.449 | Up |
| AC009276.1 | 0.013 | 5.398 | Up |
| ADAM17 | 0.011 | 5.396 | Up |
| AL139220.2 | 0.014 | 5.392 | Up |
| AC036214.2 | 0.012 | 5.392 | Up |
| TMA7 | 0.009 | 5.389 | Up |
| AL450326.1 | 0.014 | 5.382 | Up |
| CATG00000050705.1 | 0.020 | 5.376 | Up |
| BC047644 | 0.007 | 5.375 | Up |
| AL445472.1 | 0.049 | 5.357 | Up |
| CATG00000041225.1 | 0.006 | 5.338 | Up |
| AC008870.1 | 0.014 | 5.310 | Up |
| CATG00000109888.1 | 0.015 | 5.305 | Up |
| G061543 | 0.015 | 5.304 | Up |
| CLK3 | 0.008 | 5.300 | Up |
| BIG-lncRNA-1563 | 0.018 | 5.291 | Up |
| CATG00000038550.1 | 0.016 | 5.285 | Up |
| AC090377.1 | 0.010 | 5.284 | Up |
| LINC00663 | 0.009 | 5.283 | Up |
| AC022517.1 | 0.016 | 5.279 | Up |
| VPS33B-DT | 0.037 | 5.278 | Up |
| AC087477.5 | 0.039 | 5.275 | Up |
| BIG-lncRNA-1581 | 0.001 | 5.269 | Up |
| AL683887.1 | 0.017 | 5.268 | Up |
| CATG00000032467.1 | 0.008 | 5.267 | Up |
| MFSD8 | 0.009 | 5.258 | Up |
| FTX | 0.011 | 5.253 | Up |
| AC019205.1 | 0.003 | 5.252 | Up |
| AC137932.3 | 0.010 | 5.247 | Up |
| AC005954.1 | 0.014 | 5.245 | Up |
| DANCR | 0.045 | 5.235 | Up |

| | | | |
|---|---|---|---|
| CATG00000041279.1 | 0.012 | 5.229 | Up |
| LINC00863 | 0.009 | 5.218 | Up |
| OPRM1 | 0.005 | 5.218 | Up |
| PDZD2 | 0.014 | 5.215 | Up |
| AC007066.2 | 0.010 | 5.210 | Up |
| CATG00000048536.1 | 0.009 | 5.185 | Up |
| CMC1 | 0.019 | 5.144 | Up |
| G083480 | 0.033 | 5.143 | Up |
| CATG00000043846.1 | 0.017 | 5.142 | Up |
| CHPT1 | 0.011 | 5.141 | Up |
| AC067852.3 | 0.018 | 5.134 | Up |
| CATG00000073817.1 | 0.015 | 5.130 | Up |
| AC091932.1 | 0.034 | 5.119 | Up |
| CATG00000001138.1 | 0.004 | 5.117 | Up |
| LINC01973 | 0.012 | 5.109 | Up |
| LINC00273 | 0.043 | 5.099 | Up |
| UBL7-AS1 | 0.011 | 5.098 | Up |
| AP001033.2 | 0.017 | 5.083 | Up |
| AL079307.1 | 0.004 | 5.070 | Up |
| AC020913.3 | 0.019 | 5.067 | Up |
| BIG-lncRNA-1113 | 0.015 | 5.063 | Up |
| IDE | 0.018 | 5.046 | Up |
| G024116 | 0.011 | 5.044 | Up |
| AC024614.4 | 0.010 | 5.034 | Up |
| ANP32A | 0.015 | 5.034 | Up |
| AP001469.3 | 0.005 | 5.031 | Up |
| LINC00578 | 0.018 | 5.029 | Up |
| XLOC_013763 | 0.001 | 5.024 | Up |
| AL034376.1 | 0.025 | 5.019 | Up |
| AC008982.2 | 0.020 | 4.971 | Up |
| AC022140.1 | 0.026 | 4.967 | Up |
| IKBKB | 0.009 | 4.957 | Up |
| AC006115.1 | 0.015 | 4.954 | Up |
| AC020907.4 | 0.021 | 4.954 | Up |
| THUMPD3-AS1 | 0.029 | 4.946 | Up |
| AC011379.1 | 0.006 | 4.928 | Up |
| CATG00000001041.1 | 0.004 | 4.926 | Up |
| SYCE2 | 0.026 | 4.919 | Up |
| AC001226.1 | 0.025 | 4.912 | Up |
| AL163051.1 | 0.012 | 4.911 | Up |
| G002838 | 0.030 | 4.911 | Up |
| AP005717.1 | 0.020 | 4.905 | Up |
| AC023024.1 | 0.033 | 4.888 | Up |
| LINC00958 | 0.006 | 4.880 | Up |
| LINC00235 | 0.018 | 4.869 | Up |
| NAV1 | 0.018 | 4.862 | Up |
| ZNF532 | 0.012 | 4.860 | Up |
| AL359232.1 | 0.016 | 4.845 | Up |
| CATG00000061817.1 | 0.029 | 4.830 | Up |
| AGAP1-IT1 | 0.033 | 4.820 | Up |
| AL162719.1 | 0.014 | 4.815 | Up |
| AL355482.1 | 0.038 | 4.814 | Up |
| G007505 | 0.039 | 4.813 | Up |
| AL583810.1 | 0.027 | 4.811 | Up |
| GPR82 | 0.015 | 4.803 | Up |
| RP11-395P13.6 | 0.021 | 4.799 | Up |
| AC122129.1 | 0.019 | 4.797 | Up |
| AC090630.1 | 0.016 | 4.796 | Up |
| CATG00000039158.1 | 0.002 | 4.789 | Up |
| CATG00000016630.1 | 0.009 | 4.787 | Up |
| CEP89 | 0.017 | 4.787 | Up |
| AC021491.1 | 0.029 | 4.781 | Up |
| AP001059.2 | 0.027 | 4.769 | Up |
| G003293 | 0.013 | 4.760 | Up |
| AL139317.3 | 0.017 | 4.756 | Up |

| | | | |
|---|---|---|---|
| AP003680.1 | 0.025 | 4.755 | Up |
| XLOC_011448 | 0.019 | 4.752 | Up |
| AP000679.1 | 0.016 | 4.752 | Up |
| AL157931.1 | 0.007 | 4.750 | Up |
| AL031666.2 | 0.039 | 4.747 | Up |
| RPS17 | 0.030 | 4.736 | Up |
| PCBP1-AS1 | 0.005 | 4.730 | Up |
| AC108718.1 | 0.007 | 4.729 | Up |
| LOC729609 | 0.010 | 4.727 | Up |
| AP003692.1 | 0.016 | 4.717 | Up |
| RUNX1 | 0.023 | 4.709 | Up |
| TAF1D | 0.009 | 4.705 | Up |
| G063090 | 0.004 | 4.682 | Up |
| AC121333.1 | 0.004 | 4.681 | Up |
| CEP57L1 | 0.004 | 4.676 | Up |
| AC017091.1 | 0.017 | 4.673 | Up |
| AL596087.2 | 0.014 | 4.664 | Up |
| AC005899.5 | 0.016 | 4.658 | Up |
| CATG00000026306.1 | 0.022 | 4.643 | Up |
| AMT | 0.044 | 4.640 | Up |
| GSE61474_XLOC_015976 | 0.003 | 4.627 | Up |
| RBSN | 0.022 | 4.601 | Up |
| WWC3-AS1 | 0.033 | 4.567 | Up |
| TMEM78 | 0.026 | 4.563 | Up |
| MIR3976HG | 0.009 | 4.539 | Up |
| AC010273.2 | 0.021 | 4.518 | Up |
| AL662907.3 | 0.013 | 4.512 | Up |
| CATG00000070753.1 | 0.022 | 4.506 | Up |
| AC131009.2 | 0.013 | 4.497 | Up |
| AC068631.1 | 0.015 | 4.490 | Up |
| AC126175.1 | 0.012 | 4.490 | Up |
| PWRN1 | 0.014 | 4.482 | Up |
| AC136944.2 | 0.012 | 4.468 | Up |
| AL138756.1 | 0.011 | 4.461 | Up |
| AC008760.2 | 0.018 | 4.450 | Up |
| AC087430.1 | 0.023 | 4.448 | Up |
| AC084357.2 | 0.011 | 4.446 | Up |
| AC139100.2 | 0.005 | 4.446 | Up |
| AC138207.2 | 0.028 | 4.441 | Up |
| KCNG1 | 0.028 | 4.432 | Up |
| AC006504.5 | 0.006 | 4.428 | Up |
| AC009133.3 | 0.013 | 4.402 | Up |
| LINC00909 | 0.034 | 4.381 | Up |
| AC004232.2 | 0.023 | 4.376 | Up |
| CATG00000038859.1 | 0.022 | 4.345 | Up |
| LINC02377 | 0.009 | 4.338 | Up |
| TTTY14 | 0.018 | 4.334 | Up |
| CATG00000003009.1 | 0.021 | 4.330 | Up |
| AC009720.1 | 0.012 | 4.322 | Up |
| PCAT1 | 0.007 | 4.317 | Up |
| BIG-lncRNA-1681 | 0.003 | 4.316 | Up |
| AC004973.1 | 0.010 | 4.307 | Up |
| AC092068.2 | 0.027 | 4.266 | Up |
| TIMM50 | 0.011 | 4.261 | Up |
| SSTR5-AS1 | 0.018 | 4.253 | Up |
| AC025187.1 | 0.001 | 4.249 | Up |
| BIG-lncRNA-788 | 0.007 | 4.243 | Up |
| AC026336.2 | 0.003 | 4.193 | Up |
| B3GALNT1 | 0.026 | 4.162 | Up |
| KRTAP5-AS1 | 0.004 | 4.158 | Up |
| AC010551.3 | 0.003 | 4.153 | Up |
| AC080038.1 | 0.024 | 4.152 | Up |
| AC011405.1 | 0.007 | 4.152 | Up |
| MUC20-OT1 | 0.047 | 4.149 | Up |
| KCTD21-AS1 | 0.025 | 4.146 | Up |

| | | | |
|---|---|---|---|
| AC087463.3 | 0.036 | 4.146 | Up |
| AC012313.8 | 0.029 | 4.145 | Up |
| CATG00000047589.1 | 0.022 | 4.139 | Up |
| LINC02112 | 0.001 | 4.120 | Up |
| AC010422.2 | 0.018 | 4.120 | Up |
| AC020911.1 | 0.013 | 4.077 | Up |
| G078783 | 0.012 | 4.063 | Up |
| AC092153.1 | 0.039 | 4.056 | Up |
| CATG00000040018.1 | 0.016 | 4.055 | Up |
| AL078590.3 | 0.032 | 4.027 | Up |
| CATG00000005937.1 | 0.013 | 4.023 | Up |
| DTX2P1-UPK3BP1-PMS2P1 | 0.005 | 3.998 | Up |
| AC092757.2 | 0.007 | 3.993 | Up |
| LINC01355 | 0.019 | 3.990 | Up |
| AL031280.1 | 0.035 | 3.981 | Up |
| AL022341.2 | 0.046 | 3.979 | Up |
| ZBED3-AS1 | 0.004 | 3.972 | Up |
| AC016816.1 | 0.009 | 3.962 | Up |
| AC032044.1 | 0.027 | 3.956 | Up |
| NCBP2-AS1 | 0.023 | 3.956 | Up |
| SCRN3 | 0.010 | 3.934 | Up |
| DLEU1 | 0.012 | 3.933 | Up |
| AC008937.3 | 0.018 | 3.928 | Up |
| AC015987.1 | 0.021 | 3.919 | Up |
| BIG-lncRNA-1358 | 0.003 | 3.917 | Up |
| AP001094.2 | 0.021 | 3.914 | Up |
| JAKMIP3 | 0.022 | 3.907 | Up |
| LINC02268 | 0.006 | 3.895 | Up |
| AL121901.1 | 0.027 | 3.894 | Up |
| XLOC_009511 | 0.028 | 3.889 | Up |
| CATG00000071724.1 | 0.020 | 3.882 | Up |
| SOS1-IT1 | 0.016 | 3.882 | Up |
| AC132872.3 | 0.024 | 3.880 | Up |
| BIG-lncRNA-1322 | 0.004 | 3.877 | Up |
| LINC01152 | 0.002 | 3.866 | Up |
| EXOSC3 | 0.019 | 3.860 | Up |
| AC004231.1 | 0.008 | 3.850 | Up |
| CATG00000054106.1 | 0.021 | 3.845 | Up |
| BIG-lncRNA-660 | 0.009 | 3.844 | Up |
| PARP14 | 0.029 | 3.843 | Up |
| AC010636.2 | 0.022 | 3.835 | Up |
| ZNF382 | 0.006 | 3.826 | Up |
| DCP1B | 0.023 | 3.825 | Up |
| CATG00000062140.1 | 0.002 | 3.814 | Up |
| AC010542.4 | 0.022 | 3.802 | Up |
| CATG00000006648.1 | 0.029 | 3.789 | Up |
| AC087473.1 | 0.023 | 3.784 | Up |
| G073197 | 0.024 | 3.764 | Up |
| AP001496.4 | 0.011 | 3.750 | Up |
| AL590235.1 | 0.029 | 3.741 | Up |
| AC010737.1 | 0.021 | 3.736 | Up |
| C1orf61 | 0.023 | 3.724 | Up |
| BIG-lncRNA-781 | 0.004 | 3.721 | Up |
| SCAMP1 | 0.011 | 3.714 | Up |
| CATG00000092959.1 | 0.004 | 3.701 | Up |
| AC084706.1 | 0.007 | 3.690 | Up |
| CHKB | 0.011 | 3.689 | Up |
| AC021683.3 | 0.009 | 3.684 | Up |
| LINC02250 | 0.017 | 3.668 | Up |
| CATG00000017109.1 | 0.001 | 3.666 | Up |
| AC078850.1 | 0.013 | 3.664 | Up |
| CATG00000053098.1 | 0.026 | 3.651 | Up |
| MIR99AHG | 0.038 | 3.646 | Up |
| CEP83 | 0.011 | 3.645 | Up |
| AC011825.4 | 0.016 | 3.635 | Up |

| | | | |
|---|---|---|---|
| AL359237.1 | 0.025 | 3.628 | Up |
| ENTPD1-AS1 | 0.003 | 3.625 | Up |
| AC013652.1 | 0.009 | 3.595 | Up |
| SNHG5 | 0.035 | 3.585 | Up |
| AL133523.1 | 0.017 | 3.585 | Up |
| AC036222.1 | 0.017 | 3.576 | Up |
| GPSM2 | 0.025 | 3.551 | Up |
| AC083880.1 | 0.036 | 3.548 | Up |
| CATG00000045568.1 | 0.027 | 3.545 | Up |
| AC093655.1 | 0.034 | 3.530 | Up |
| HCG9 | 0.032 | 3.527 | Up |
| CATG00000039519.1 | 0.021 | 3.505 | Up |
| AC112907.1 | 0.027 | 3.500 | Up |
| SNHG6 | 0.034 | 3.495 | Up |
| AC013643.2 | 0.027 | 3.490 | Up |
| AC021915.2 | 0.011 | 3.484 | Up |
| AC107375.1 | 0.044 | 3.481 | Up |
| LINC00511 | 0.041 | 3.468 | Up |
| AL358214.1 | 0.011 | 3.461 | Up |
| LINC00355 | 0.001 | 3.456 | Up |
| CATG00000013305.1 | 0.017 | 3.451 | Up |
| LINC01322 | 0.027 | 3.436 | Up |
| ITCH-AS1 | 0.017 | 3.431 | Up |
| CATG00000084124.1 | 0.011 | 3.405 | Up |
| ZNF397 | 0.022 | 3.401 | Up |
| PLCE1-AS2 | 0.003 | 3.401 | Up |
| AC098859.2 | 0.028 | 3.398 | Up |
| REV3L-IT1 | 0.034 | 3.391 | Up |
| BIG-lncRNA-202 | 0.011 | 3.369 | Up |
| LINC01480 | 0.012 | 3.368 | Up |
| AL133284.1 | 0.010 | 3.364 | Up |
| G054876 | 0.025 | 3.363 | Up |
| G038959 | 0.017 | 3.362 | Up |
| XLOC_011682 | 0.026 | 3.357 | Up |
| CATG00000044067.1 | 0.028 | 3.356 | Up |
| CATG00000096859.1 | 0.009 | 3.352 | Up |
| SEMA6A-AS2 | 0.017 | 3.350 | Up |
| AC008125.1 | 0.031 | 3.349 | Up |
| CATG00000111728.1 | 0.033 | 3.346 | Up |
| LINC01249 | 0.023 | 3.339 | Up |
| XLOC_006972 | 0.031 | 3.338 | Up |
| RBFADN | 0.043 | 3.336 | Up |
| ERVK-28 | 0.040 | 3.334 | Up |
| ZMYM3 | 0.035 | 3.333 | Up |
| ARHGEF26-AS1 | 0.027 | 3.323 | Up |
| CATG00000090511.1 | 0.005 | 3.314 | Up |
| LHFPL3-AS1 | 0.005 | 3.309 | Up |
| CATG00000115245.1 | 0.014 | 3.303 | Up |
| CARD8 | 0.034 | 3.294 | Up |
| KC6 | 0.012 | 3.288 | Up |
| AC004448.2 | 0.017 | 3.287 | Up |
| PODNL1 | 0.021 | 3.265 | Up |
| CATG00000025547.1 | 0.023 | 3.254 | Up |
| AC096711.1 | 0.020 | 3.238 | Up |
| AC104534.1 | 0.014 | 3.233 | Up |
| AC130456.2 | 0.015 | 3.233 | Up |
| GOLGA8IP | 0.049 | 3.221 | Up |
| G025830 | 0.047 | 3.212 | Up |
| XLOC_009426 | 0.002 | 3.208 | Up |
| LINC01307 | 0.004 | 3.206 | Up |
| G013478 | 0.030 | 3.173 | Up |
| AC074286.1 | 0.015 | 3.171 | Up |
| LINC00868 | 0.013 | 3.161 | Up |
| DSG1-AS1 | 0.032 | 3.152 | Up |
| AL607028.1 | 0.038 | 3.138 | Up |

| | | | |
|---|---|---|---|
| AC023813.3 | 0.017 | 3.134 | Up |
| LINC00884 | 0.040 | 3.129 | Up |
| BIG-lncRNA-783 | 0.005 | 3.127 | Up |
| AC093283.1 | 0.018 | 3.126 | Up |
| AC010536.3 | 0.020 | 3.124 | Up |
| AC011676.4 | 0.010 | 3.119 | Up |
| AL022322.1 | 0.012 | 3.112 | Up |
| SMCR5 | 0.011 | 3.108 | Up |
| CATG00000071063.1 | 0.008 | 3.103 | Up |
| G056794 | 0.009 | 3.102 | Up |
| AP001020.1 | 0.024 | 3.097 | Up |
| CATG00000010537.1 | 0.001 | 3.091 | Up |
| TTN-AS1 | 0.006 | 3.075 | Up |
| AC026992.1 | 0.008 | 3.073 | Up |
| SIRLNT | 0.037 | 3.068 | Up |
| LINC00299 | 0.015 | 3.061 | Up |
| LINC02363 | 0.008 | 3.041 | Up |
| C12orf56 | 0.038 | 3.030 | Up |
| CATG00000021649.1 | 0.002 | 3.027 | Up |
| AC025569.1 | 0.039 | 3.026 | Up |
| LINC01002 | 0.039 | 3.023 | Up |
| AP001271.1 | 0.023 | 3.020 | Up |
| AL591767.1 | 0.026 | 3.019 | Up |
| LINC02432 | 0.043 | 3.013 | Up |
| AC055717.2 | 0.011 | 3.009 | Up |
| AC010210.1 | 0.015 | 3.000 | Up |
| BIG-lncRNA-784 | 0.003 | 2.994 | Up |
| AC138207.7 | 0.027 | 2.986 | Up |
| AL590705.1 | 0.014 | 2.985 | Up |
| MFSD14C | 0.045 | 2.976 | Up |
| AC004951.1 | 0.032 | 2.974 | Up |
| AC244153.1 | 0.029 | 2.973 | Up |
| DSCR9 | 0.046 | 2.972 | Up |
| LINC01702 | 0.050 | 2.966 | Up |
| AC022893.1 | 0.036 | 2.954 | Up |
| PCAT14 | 0.041 | 2.943 | Up |
| AC012594.1 | 0.039 | 2.942 | Up |
| ZNF561-AS1 | 0.037 | 2.942 | Up |
| LINC00276 | 0.043 | 2.925 | Up |
| AL162457.1 | 0.034 | 2.920 | Up |
| AP002754.2 | 0.000 | 2.917 | Up |
| AKAP14 | 0.036 | 2.910 | Up |
| LINC01841 | 0.040 | 2.906 | Up |
| LINC00662 | 0.040 | 2.903 | Up |
| AC093525.4 | 0.015 | 2.894 | Up |
| ZNF888 | 0.037 | 2.891 | Up |
| CATG00000047301.1 | 0.011 | 2.890 | Up |
| AC091180.2 | 0.044 | 2.887 | Up |
| AC108471.2 | 0.042 | 2.880 | Up |
| Z69720.1 | 0.032 | 2.880 | Up |
| AC008737.3 | 0.024 | 2.878 | Up |
| LINC02552 | 0.017 | 2.875 | Up |
| NRG1 | 0.039 | 2.870 | Up |
| AC012358.1 | 0.003 | 2.869 | Up |
| AL359962.2 | 0.018 | 2.866 | Up |
| AC092802.3 | 0.046 | 2.863 | Up |
| AC127024.2 | 0.049 | 2.859 | Up |
| G079559 | 0.006 | 2.857 | Up |
| AC104041.1 | 0.019 | 2.850 | Up |
| AC092902.5 | 0.035 | 2.850 | Up |
| AL390786.1 | 0.033 | 2.848 | Up |
| LINC02577 | 0.049 | 2.845 | Up |
| CATG00000091077.1 | 0.043 | 2.843 | Up |
| AC026191.1 | 0.019 | 2.841 | Up |
| AC002464.1 | 0.037 | 2.834 | Up |

| | | | |
|---|---|---|---|
| AC007861.2 | 0.040 | 2.815 | Up |
| CASC9 | 0.014 | 2.810 | Up |
| AC099778.1 | 0.011 | 2.804 | Up |
| LINC01798 | 0.042 | 2.800 | Up |
| AL157944.1 | 0.007 | 2.798 | Up |
| AP000235.1 | 0.041 | 2.797 | Up |
| LINC02093 | 0.015 | 2.780 | Up |
| AC073320.1 | 0.012 | 2.778 | Up |
| AC124798.1 | 0.030 | 2.778 | Up |
| LINC01821 | 0.032 | 2.778 | Up |
| G007290 | 0.000 | 2.767 | Up |
| AC114400.1 | 0.043 | 2.762 | Up |
| LINC00996 | 0.017 | 2.759 | Up |
| AC008728.1 | 0.040 | 2.756 | Up |
| AC092069.1 | 0.027 | 2.751 | Up |
| BIG-lncRNA-1619 | 0.030 | 2.749 | Up |
| AC005884.1 | 0.036 | 2.745 | Up |
| PSMA3-AS1 | 0.041 | 2.745 | Up |
| GRIK1-AS1 | 0.013 | 2.740 | Up |
| ZNF385B | 0.029 | 2.738 | Up |
| ASMTL-AS1 | 0.017 | 2.726 | Up |
| AC073529.1 | 0.043 | 2.725 | Up |
| AC097461.1 | 0.046 | 2.720 | Up |
| KCNMB2-AS1 | 0.033 | 2.719 | Up |
| CAV2 | 0.037 | 2.718 | Up |
| AC073534.1 | 0.006 | 2.712 | Up |
| TNRC6A | 0.036 | 2.710 | Up |
| AL356277.3 | 0.005 | 2.710 | Up |
| CATG00000020199.1 | 0.049 | 2.709 | Up |
| LOC338963 | 0.005 | 2.702 | Up |
| GMDS-DT | 0.013 | 2.701 | Up |
| OBSCN | 0.002 | 2.698 | Up |
| CATG00000006995.1 | 0.016 | 2.694 | Up |
| HDAC10 | 0.022 | 2.688 | Up |
| AC008738.2 | 0.033 | 2.687 | Up |
| AC107463.1 | 0.000 | 2.682 | Up |
| AC102797.1 | 0.043 | 2.673 | Up |
| TBX2-AS1 | 0.011 | 2.671 | Up |
| AC068152.1 | 0.046 | 2.649 | Up |
| AP000640.2 | 0.023 | 2.639 | Up |
| C15orf53 | 0.047 | 2.638 | Up |
| LINC01569 | 0.038 | 2.635 | Up |
| AC009654.1 | 0.049 | 2.632 | Up |
| LINC00635 | 0.011 | 2.631 | Up |
| ADAMTS9-AS2 | 0.038 | 2.630 | Up |
| BIG-lncRNA-661 | 0.014 | 2.628 | Up |
| BIG-lncRNA-990 | 0.003 | 2.627 | Up |
| AC026316.2 | 0.033 | 2.622 | Up |
| AC145350.2 | 0.002 | 2.621 | Up |
| AC012629.2 | 0.034 | 2.617 | Up |
| LINC00907 | 0.023 | 2.613 | Up |
| AL162311.3 | 0.034 | 2.612 | Up |
| ERVMER61-1 | 0.004 | 2.611 | Up |
| AC106798.1 | 0.007 | 2.609 | Up |
| AC008574.1 | 0.014 | 2.608 | Up |
| CATG00000061776.1 | 0.045 | 2.605 | Up |
| SLC7A7 | 0.003 | 2.603 | Up |
| CYYR1-AS1 | 0.026 | 2.600 | Up |
| CATG00000008928.1 | 0.026 | 2.589 | Up |
| BIG-lncRNA-1621 | 0.038 | 2.583 | Up |
| XLOC_005552 | 0.003 | 2.582 | Up |
| AC109588.1 | 0.043 | 2.582 | Up |
| XLOC_005888 | 0.026 | 2.582 | Up |
| AC027097.2 | 0.035 | 2.580 | Up |
| CATG00000037022.1 | 0.040 | 2.579 | Up |

| | | | |
|---|---|---|---|
| CATG00000041223.1 | 0.050 | 2.567 | Up |
| CATG00000091713.1 | 0.030 | 2.563 | Up |
| CATG00000013271.1 | 0.018 | 2.562 | Up |
| LINC02253 | 0.016 | 2.562 | Up |
| CATG00000042143.1 | 0.029 | 2.552 | Up |
| AL450311.1 | 0.050 | 2.551 | Up |
| AC068875.1 | 0.023 | 2.551 | Up |
| AL589743.4 | 0.044 | 2.551 | Up |
| LINC00917 | 0.009 | 2.545 | Up |
| AC087500.1 | 0.020 | 2.529 | Up |
| LINC01146 | 0.020 | 2.529 | Up |
| BIG-lncRNA-972 | 0.020 | 2.522 | Up |
| AL583810.2 | 0.039 | 2.518 | Up |
| CYP2U1 | 0.029 | 2.511 | Up |
| CATG00000012014.1 | 0.011 | 2.507 | Up |
| AC020719.1 | 0.040 | 2.503 | Up |
| AC108681.1 | 0.038 | 2.503 | Up |
| CATG00000004404.1 | 0.047 | 2.498 | Up |
| AL133268.2 | 0.024 | 2.495 | Up |
| AC016550.3 | 0.040 | 2.489 | Up |
| SAMMSON | 0.010 | 2.483 | Up |
| AC022639.1 | 0.026 | 2.458 | Up |
| AC131056.5 | 0.041 | 2.453 | Up |
| TMEM51-AS1 | 0.018 | 2.450 | Up |
| CATG00000049817.1 | 0.006 | 2.448 | Up |
| AC024257.4 | 0.028 | 2.447 | Up |
| G064015 | 0.012 | 2.446 | Up |
| G002611 | 0.024 | 2.445 | Up |
| STK3 | 0.040 | 2.442 | Up |
| AC044839.1 | 0.013 | 2.438 | Up |
| AL160408.4 | 0.028 | 2.424 | Up |
| AC140912.1 | 0.022 | 2.423 | Up |
| AC005972.3 | 0.008 | 2.416 | Up |
| CELF2-AS1 | 0.047 | 2.415 | Up |
| AC069277.1 | 0.018 | 2.410 | Up |
| AC005324.1 | 0.040 | 2.409 | Up |
| LINC01748 | 0.027 | 2.399 | Up |
| LINC00891 | 0.001 | 2.394 | Up |
| ERLNC1 | 0.049 | 2.392 | Up |
| AC016152.1 | 0.019 | 2.385 | Up |
| NDUFA2 | 0.003 | 2.378 | Up |
| GSE61474_XLOC_057016 | 0.031 | 2.375 | Up |
| XLOC_009637 | 0.007 | 2.373 | Up |
| C16orf95 | 0.034 | 2.370 | Up |
| G023595 | 0.013 | 2.367 | Up |
| AC116353.4 | 0.013 | 2.360 | Up |
| AC092132.1 | 0.019 | 2.352 | Up |
| AL592429.2 | 0.005 | 2.351 | Up |
| CYP1B1-AS1 | 0.023 | 2.345 | Up |
| PIGB | 0.030 | 2.336 | Up |
| DNAJC3-DT | 0.029 | 2.334 | Up |
| HCG22 | 0.038 | 2.326 | Up |
| CD55 | 0.034 | 2.325 | Up |
| AC007834.1 | 0.017 | 2.319 | Up |
| CATG00000004175.1 | 0.016 | 2.317 | Up |
| AC111149.2 | 0.033 | 2.316 | Up |
| AL031316.1 | 0.002 | 2.315 | Up |
| AL391095.3 | 0.028 | 2.311 | Up |
| AL357793.1 | 0.032 | 2.310 | Up |
| AC137834.2 | 0.003 | 2.307 | Up |
| AP002358.1 | 0.038 | 2.307 | Up |
| AC233309.1 | 0.024 | 2.302 | Up |
| AL133372.2 | 0.025 | 2.298 | Up |
| LINC00941 | 0.009 | 2.296 | Up |
| AL031663.3 | 0.027 | 2.295 | Up |

| | | | |
|---|---|---|---|
| BC048141 | 0.046 | 2.289 | Up |
| AC103858.1 | 0.010 | 2.282 | Up |
| EFCAB6-AS1 | 0.037 | 2.278 | Up |
| AL117335.1 | 0.035 | 2.278 | Up |
| LINC02155 | 0.016 | 2.273 | Up |
| FAM167A-AS1 | 0.008 | 2.270 | Up |
| MIR137HG | 0.030 | 2.269 | Up |
| MYT1L-AS1 | 0.003 | 2.266 | Up |
| AL023806.2 | 0.050 | 2.264 | Up |
| AL512631.2 | 0.045 | 2.263 | Up |
| ALKBH3-AS1 | 0.038 | 2.258 | Up |
| Z84485.1 | 0.020 | 2.256 | Up |
| AC138627.1 | 0.012 | 2.254 | Up |
| AC104170.1 | 0.001 | 2.254 | Up |
| AC018638.6 | 0.005 | 2.244 | Up |
| AC093732.2 | 0.040 | 2.240 | Up |
| FIRRE | 0.010 | 2.239 | Up |
| G031056 | 0.037 | 2.229 | Up |
| AC104393.1 | 0.037 | 2.228 | Up |
| AL049629.1 | 0.010 | 2.226 | Up |
| AC008103.1 | 0.003 | 2.224 | Up |
| CATG00000046972.1 | 0.031 | 2.224 | Up |
| AC068722.2 | 0.048 | 2.224 | Up |
| LINC00466 | 0.009 | 2.222 | Up |
| AC027287.2 | 0.027 | 2.219 | Up |
| CATG00000095582.1 | 0.034 | 2.217 | Up |
| CATG00000118162.1 | 0.039 | 2.208 | Up |
| CATG00000078081.1 | 0.018 | 2.199 | Up |
| CATG00000013090.1 | 0.017 | 2.198 | Up |
| AL078612.2 | 0.026 | 2.197 | Up |
| AC091563.1 | 0.006 | 2.194 | Up |
| AC092017.4 | 0.007 | 2.192 | Up |
| XLOC_008866 | 0.041 | 2.190 | Up |
| Z94160.1 | 0.016 | 2.188 | Up |
| G029743 | 0.003 | 2.185 | Up |
| AL451137.1 | 0.015 | 2.179 | Up |
| FLJ13224 | 0.023 | 2.178 | Up |
| CATG00000048273.1 | 0.004 | 2.169 | Up |
| LOC100505915 | 0.024 | 2.161 | Up |
| AL138781.2 | 0.034 | 2.158 | Up |
| AL035420.2 | 0.018 | 2.156 | Up |
| AL591485.1 | 0.036 | 2.155 | Up |
| XLOC_006442 | 0.030 | 2.155 | Up |
| LINC02116 | 0.047 | 2.154 | Up |
| AP002026.1 | 0.012 | 2.151 | Up |
| CATG00000057773.1 | 0.044 | 2.140 | Up |
| FRYL | 0.021 | 2.140 | Up |
| LINC00424 | 0.007 | 2.139 | Up |
| AC137932.1 | 0.048 | 2.138 | Up |
| SEPT5 | 0.036 | 2.138 | Up |
| AL592043.1 | 0.018 | 2.136 | Up |
| CATG00000022496.1 | 0.002 | 2.136 | Up |
| LIX1L-AS1 | 0.035 | 2.136 | Up |
| TMC6 | 0.035 | 2.129 | Up |
| KCNMA1-AS2 | 0.007 | 2.128 | Up |
| BAIAP2-DT | 0.012 | 2.127 | Up |
| G028058 | 0.007 | 2.127 | Up |
| LINC01219 | 0.006 | 2.124 | Up |
| LINC00426 | 0.008 | 2.124 | Up |
| AC113386.1 | 0.043 | 2.124 | Up |
| AC128685.1 | 0.031 | 2.121 | Up |
| LINC01568 | 0.045 | 2.121 | Up |
| AC004969.1 | 0.021 | 2.120 | Up |
| TLR8-AS1 | 0.029 | 2.118 | Up |
| G017655 | 0.030 | 2.118 | Up |

| | | | |
|---|---|---|---|
| AC023824.4 | 0.001 | 2.118 | Up |
| AL162431.1 | 0.022 | 2.117 | Up |
| MEF2C-AS1 | 0.018 | 2.116 | Up |
| FP325330.3 | 0.009 | 2.113 | Up |
| TET2-AS1 | 0.010 | 2.112 | Up |
| TEX41 | 0.009 | 2.108 | Up |
| LINC01473 | 0.023 | 2.105 | Up |
| AC022031.2 | 0.013 | 2.104 | Up |
| AL133387.3 | 0.026 | 2.104 | Up |
| LACTB2-AS1 | 0.008 | 2.103 | Up |
| GS1-600G8.3 | 0.010 | 2.099 | Up |
| ELMO1 | 0.029 | 2.098 | Up |
| CATG00000110068.1 | 0.032 | 2.096 | Up |
| G040022 | 0.023 | 2.095 | Up |
| AC006557.3 | 0.016 | 2.091 | Up |
| AC110491.3 | 0.014 | 2.089 | Up |
| IDH2-DT | 0.043 | 2.089 | Up |
| AC244517.11 | 0.027 | 2.087 | Up |
| AP003721.4 | 0.017 | 2.086 | Up |
| XLOC_010574 | 0.035 | 2.080 | Up |
| GNG12-AS1 | 0.019 | 2.080 | Up |
| LINC00836 | 0.033 | 2.078 | Up |
| XLOC_004700 | 0.011 | 2.077 | Up |
| AC107871.2 | 0.049 | 2.075 | Up |
| RBM12B | 0.009 | 2.072 | Up |
| G090268 | 0.050 | 2.072 | Up |
| SMIM35 | 0.005 | 2.071 | Up |
| AL020996.1 | 0.049 | 2.069 | Up |
| AC136475.8 | 0.027 | 2.065 | Up |
| AC018755.4 | 0.049 | 2.065 | Up |
| CATG00000057360.1 | 0.042 | 2.064 | Up |
| G035514 | 0.013 | 2.064 | Up |
| LINC01641 | 0.046 | 2.062 | Up |
| AL121987.1 | 0.008 | 2.062 | Up |
| SLC4A1 | 0.033 | 2.061 | Up |
| VARS2 | 0.037 | 2.060 | Up |
| ATXN8OS | 0.008 | 2.057 | Up |
| AL355306.2 | 0.006 | 2.057 | Up |
| AL139246.3 | 0.041 | 2.056 | Up |
| Z82249.1 | 0.017 | 2.054 | Up |
| XLOC_009873 | 0.048 | 2.054 | Up |
| GARS-DT | 0.036 | 2.053 | Up |
| IDI2-AS1 | 0.002 | 2.053 | Up |
| GPR78 | 0.045 | 2.052 | Up |
| LINC00661 | 0.026 | 2.050 | Up |
| AC140847.2 | 0.046 | 2.041 | Up |
| CATG00000107547.1 | 0.030 | 2.041 | Up |
| AC113383.1 | 0.016 | 2.040 | Up |
| ITGB3BP | 0.014 | 2.039 | Up |
| AC083973.1 | 0.024 | 2.038 | Up |
| AL669970.2 | 0.046 | 2.038 | Up |
| LCN15 | 0.016 | 2.034 | Up |
| XLOC_003682 | 0.004 | 2.033 | Up |
| AC004233.2 | 0.041 | 2.032 | Up |
| NUBP1 | 0.028 | 2.029 | Up |
| AL022315.1 | 0.049 | 2.022 | Up |
| SLC6A20 | 0.043 | 2.021 | Up |
| AC239798.4 | 0.014 | 2.019 | Up |
| LAX1 | 0.001 | 2.018 | Up |
| CYTOR | 0.003 | 2.017 | Up |
| Z97987.1 | 0.036 | 2.013 | Up |
| AP000442.2 | 0.007 | 2.009 | Up |
| LINC00523 | 0.045 | 2.009 | Up |
| RHOXF1-AS1 | 0.009 | 2.008 | Up |
| PCED1B-AS1 | 0.038 | 2.006 | Up |

| | | | |
|---|---|---|---|
| AC010524.1 | 0.009 | 2.004 | Up |
| AC022509.4 | 0.037 | 2.000 | Up |
| AL451105.2 | 0.015 | 1.999 | Up |
| LINC00570 | 0.042 | 1.999 | Up |
| GAS2 | 0.026 | 1.988 | Up |
| PGM5P4-AS1 | 0.043 | 1.986 | Up |
| CATG00000043382.1 | 0.038 | 1.986 | Up |
| CATG00000101238.1 | 0.019 | 1.986 | Up |
| AC018946.1 | 0.046 | 1.984 | Up |
| G009433 | 0.014 | 1.981 | Up |
| G038394 | 0.023 | 1.977 | Up |
| XLOC_006825 | 0.022 | 1.972 | Up |
| AC148477.1 | 0.019 | 1.971 | Up |
| SMC5-AS1 | 0.050 | 1.970 | Up |
| AC003666.1 | 0.023 | 1.970 | Up |
| LINC01697 | 0.013 | 1.970 | Up |
| LINC00906 | 0.030 | 1.967 | Up |
| CATG00000032517.1 | 0.045 | 1.963 | Up |
| XLOC_000199 | 0.027 | 1.961 | Up |
| AC008555.5 | 0.033 | 1.960 | Up |
| LINC01433 | 0.042 | 1.957 | Up |
| CATG00000017734.1 | 0.003 | 1.957 | Up |
| AC112236.2 | 0.018 | 1.953 | Up |
| KCNIP2-AS1 | 0.038 | 1.950 | Up |
| AC044784.2 | 0.025 | 1.950 | Up |
| AC016526.2 | 0.030 | 1.949 | Up |
| SNX5 | 0.035 | 1.945 | Up |
| AC087463.4 | 0.001 | 1.945 | Up |
| AC091138.1 | 0.017 | 1.944 | Up |
| LINC00469 | 0.019 | 1.937 | Up |
| AL133551.1 | 0.021 | 1.937 | Up |
| S1PR3 | 0.015 | 1.937 | Up |
| AC067752.1 | 0.044 | 1.936 | Up |
| AC138123.1 | 0.017 | 1.933 | Up |
| LRTOMT | 0.048 | 1.930 | Up |
| LINC00686 | 0.007 | 1.930 | Up |
| CATG00000098333.1 | 0.019 | 1.928 | Up |
| LINC00841 | 0.037 | 1.926 | Up |
| G037500 | 0.014 | 1.923 | Up |
| POT1-AS1 | 0.036 | 1.922 | Up |
| CATG00000113515.1 | 0.025 | 1.921 | Up |
| AC008443.2 | 0.001 | 1.919 | Up |
| IPO5P1 | 0.046 | 1.917 | Up |
| AC007126.1 | 0.042 | 1.915 | Up |
| PPARGC1A | 0.003 | 1.910 | Up |
| AC103808.2 | 0.014 | 1.908 | Up |
| G023452 | 0.003 | 1.907 | Up |
| CD81-AS1 | 0.007 | 1.905 | Up |
| LINC01913 | 0.016 | 1.905 | Up |
| HOXB-AS1 | 0.018 | 1.904 | Up |
| AC138305.1 | 0.039 | 1.901 | Up |
| H19 | 0.005 | 1.899 | Up |
| BAALC-AS1 | 0.007 | 1.898 | Up |
| CLHC1 | 0.002 | 1.898 | Up |
| CATG00000084862.1 | 0.003 | 1.897 | Up |
| HCG25 | 0.001 | 1.897 | Up |
| AC105118.1 | 0.037 | 1.894 | Up |
| CATG00000055418.1 | 0.016 | 1.894 | Up |
| LINC01677 | 0.010 | 1.893 | Up |
| PCA3 | 0.001 | 1.892 | Up |
| CATG00000005696.1 | 0.003 | 1.890 | Up |
| XLOC_006623 | 0.005 | 1.890 | Up |
| IFT140 | 0.018 | 1.887 | Up |
| LINC02056 | 0.005 | 1.885 | Up |
| NEDD4L | 0.020 | 1.885 | Up |

| | | | |
|---|---|---|---|
| XLOC_007585 | 0.021 | 1.884 | Up |
| LINC01505 | 0.016 | 1.884 | Up |
| AF121898.1 | 0.033 | 1.884 | Up |
| JRK | 0.014 | 1.883 | Up |
| CLMAT3 | 0.003 | 1.879 | Up |
| RPL35A | 0.027 | 1.879 | Up |
| AC009704.1 | 0.026 | 1.878 | Up |
| G027616 | 0.039 | 1.876 | Up |
| XLOC_001343 | 0.041 | 1.875 | Up |
| AC135895.1 | 0.009 | 1.874 | Up |
| CATG00000005835.1 | 0.015 | 1.874 | Up |
| XLOC_002300 | 0.017 | 1.873 | Up |
| CATG00000018531.1 | 0.017 | 1.873 | Up |
| AC079906.1 | 0.019 | 1.871 | Up |
| LINC01562 | 0.015 | 1.868 | Up |
| PPP2R2B | 0.002 | 1.868 | Up |
| AC240565.1 | 0.034 | 1.865 | Up |
| SNRPD1 | 0.029 | 1.865 | Up |
| AC104777.1 | 0.032 | 1.864 | Up |
| XLOC_012590 | 0.022 | 1.863 | Up |
| G053330 | 0.048 | 1.863 | Up |
| NAGPA | 0.043 | 1.862 | Up |
| RNF185-AS1 | 0.012 | 1.862 | Up |
| XLOC_008653 | 0.043 | 1.862 | Up |
| G053569 | 0.022 | 1.862 | Up |
| TMEM256-PLSCR3 | 0.013 | 1.859 | Up |
| LINC00486 | 0.043 | 1.858 | Up |
| ZNF542P | 0.021 | 1.855 | Up |
| AL355300.1 | 0.021 | 1.855 | Up |
| AL031430.1 | 0.014 | 1.853 | Up |
| AC106799.3 | 0.016 | 1.853 | Up |
| CATG00000052995.1 | 0.021 | 1.852 | Up |
| AP3S1 | 0.019 | 1.852 | Up |
| G071172 | 0.047 | 1.846 | Up |
| AC116612.1 | 0.049 | 1.843 | Up |
| AC108002.1 | 0.026 | 1.843 | Up |
| AP000873.3 | 0.042 | 1.842 | Up |
| AC099520.1 | 0.044 | 1.842 | Up |
| AC068418.2 | 0.047 | 1.841 | Up |
| GAS5 | 0.001 | 1.839 | Up |
| LOC729739 | 0.050 | 1.838 | Up |
| LOC644172 | 0.019 | 1.837 | Up |
| AC004687.1 | 0.023 | 1.837 | Up |
| G004875 | 0.026 | 1.835 | Up |
| LINC01134 | 0.039 | 1.831 | Up |
| AC136489.1 | 0.028 | 1.831 | Up |
| AC246817.1 | 0.007 | 1.830 | Up |
| AC119424.1 | 0.021 | 1.829 | Up |
| G077834 | 0.046 | 1.829 | Up |
| AC060809.1 | 0.020 | 1.825 | Up |
| AC004803.1 | 0.026 | 1.825 | Up |
| MLIP-AS1 | 0.030 | 1.824 | Up |
| DNTTIP2 | 0.009 | 1.823 | Up |
| AC008568.1 | 0.028 | 1.822 | Up |
| AC004014.1 | 0.033 | 1.821 | Up |
| TSNAX-DISC1 | 0.036 | 1.821 | Up |
| AL360014.1 | 0.050 | 1.819 | Up |
| AC005842.1 | 0.029 | 1.817 | Up |
| AC021549.1 | 0.043 | 1.815 | Up |
| LINC00112 | 0.004 | 1.814 | Up |
| GSE61474_XLOC_032135 | 0.012 | 1.814 | Up |
| OSTN-AS1 | 0.036 | 1.813 | Up |
| CATG00000054576.1 | 0.011 | 1.812 | Up |
| TEN1-CDK3 | 0.026 | 1.812 | Up |
| RAD51-AS1 | 0.026 | 1.806 | Up |

| | | | |
|---|---|---|---|
| AC099681.3 | 0.035 | 1.806 | Up |
| MZF1-AS1 | 0.027 | 1.805 | Up |
| DEPP1 | 0.038 | 1.801 | Up |
| AC016168.2 | 0.020 | 1.800 | Up |
| AC006111.3 | 0.050 | 1.799 | Up |
| LINC00877 | 0.008 | 1.798 | Up |
| DXO | 0.023 | 1.797 | Up |
| AC018832.1 | 0.015 | 1.796 | Up |
| AC023128.1 | 0.007 | 1.795 | Up |
| AC104257.1 | 0.035 | 1.795 | Up |
| G018858 | 0.013 | 1.791 | Up |
| CATG00000027674.1 | 0.008 | 1.788 | Up |
| LINC00618 | 0.040 | 1.787 | Up |
| GSN-AS1 | 0.042 | 1.783 | Up |
| RALY-AS1 | 0.023 | 1.782 | Up |
| LINC01515 | 0.015 | 1.781 | Up |
| G070086 | 0.017 | 1.780 | Up |
| RASAL2-AS1 | 0.036 | 1.779 | Up |
| AC108488.1 | 0.017 | 1.778 | Up |
| LINC00908 | 0.025 | 1.778 | Up |
| PMS2 | 0.009 | 1.776 | Up |
| LINC01105 | 0.018 | 1.775 | Up |
| AC124944.1 | 0.014 | 1.774 | Up |
| CATG00000058583.1 | 0.046 | 1.774 | Up |
| CCDC26 | 0.002 | 1.773 | Up |
| LINC00698 | 0.016 | 1.772 | Up |
| AC144450.1 | 0.010 | 1.771 | Up |
| AC020659.1 | 0.011 | 1.771 | Up |
| AC005329.1 | 0.007 | 1.766 | Up |
| AC087645.2 | 0.040 | 1.764 | Up |
| GSE61474_XLOC_015884 | 0.029 | 1.764 | Up |
| G038783 | 0.027 | 1.764 | Up |
| CHKB-DT | 0.019 | 1.763 | Up |
| CTD-2201I18.1 | 0.030 | 1.763 | Up |
| AC027682.4 | 0.044 | 1.762 | Up |
| PXN-AS1 | 0.022 | 1.762 | Up |
| AC074366.1 | 0.015 | 1.761 | Up |
| AC103702.2 | 0.024 | 1.761 | Up |
| CYTH2 | 0.000 | 1.760 | Up |
| ATP6V0E2-AS1 | 0.017 | 1.760 | Up |
| RPL7 | 0.042 | 1.758 | Up |
| PRELID3B | 0.022 | 1.758 | Up |
| CSPG4P1Y | 0.031 | 1.748 | Up |
| MIR7515HG | 0.012 | 1.745 | Up |
| LIPE-AS1 | 0.017 | 1.744 | Up |
| ERVK13-1 | 0.009 | 1.743 | Up |
| AL022725.1 | 0.027 | 1.743 | Up |
| AC246817.2 | 0.025 | 1.742 | Up |
| LINC02551 | 0.006 | 1.742 | Up |
| AL365277.1 | 0.037 | 1.741 | Up |
| LINC01012 | 0.045 | 1.739 | Up |
| MAGI2-AS3 | 0.035 | 1.738 | Up |
| LINC01854 | 0.003 | 1.738 | Up |
| AP001107.6 | 0.013 | 1.736 | Up |
| ZFAS1 | 0.020 | 1.736 | Up |
| LINC01095 | 0.035 | 1.734 | Up |
| AL590683.1 | 0.024 | 1.733 | Up |
| DCTN1-AS1 | 0.041 | 1.731 | Up |
| LINC00968 | 0.025 | 1.730 | Up |
| NTMT1 | 0.040 | 1.730 | Up |
| C13orf42 | 0.019 | 1.728 | Up |
| AC114488.2 | 0.009 | 1.727 | Up |
| COL4A6 | 0.048 | 1.724 | Up |
| AC104066.2 | 0.027 | 1.720 | Up |
| AL691520.1 | 0.035 | 1.719 | Up |

| | | | |
|---|---|---|---|
| SALRNA1 | 0.028 | 1.718 | Up |
| XLOC_013934 | 0.023 | 1.715 | Up |
| BAG1 | 0.048 | 1.714 | Up |
| CATG00000040757.1 | 0.026 | 1.714 | Up |
| G007481 | 0.041 | 1.712 | Up |
| NAV2-AS3 | 0.005 | 1.712 | Up |
| LINC02354 | 0.036 | 1.712 | Up |
| AC026688.2 | 0.008 | 1.710 | Up |
| ELK4 | 0.020 | 1.707 | Up |
| AC008676.1 | 0.018 | 1.702 | Up |
| AL356056.1 | 0.003 | 1.701 | Up |
| LINC01749 | 0.021 | 1.698 | Up |
| AC010880.1 | 0.033 | 1.697 | Up |
| MIR205HG | 0.001 | 1.696 | Up |
| VPS53 | 0.009 | 1.696 | Up |
| AC016876.1 | 0.021 | 1.695 | Up |
| AL359094.1 | 0.019 | 1.693 | Up |
| GSE61474_XLOC_007281 | 0.023 | 1.693 | Up |
| RTL8A | 0.015 | 1.690 | Up |
| AC009299.3 | 0.049 | 1.690 | Up |
| AC073569.2 | 0.044 | 1.689 | Up |
| MFGE8 | 0.003 | 1.689 | Up |
| HNF4A-AS1 | 0.004 | 1.688 | Up |
| CATG00000095007.1 | 0.025 | 1.686 | Up |
| LINC00519 | 0.012 | 1.686 | Up |
| ZNF503-AS2 | 0.034 | 1.684 | Up |
| ATP1B3-AS1 | 0.015 | 1.684 | Up |
| LINC00964 | 0.024 | 1.682 | Up |
| CATG00000038433.1 | 0.005 | 1.681 | Up |
| G076420 | 0.009 | 1.680 | Up |
| AC004540.1 | 0.040 | 1.679 | Up |
| POLN | 0.014 | 1.679 | Up |
| AP000911.1 | 0.025 | 1.679 | Up |
| XLOC_005522 | 0.028 | 1.677 | Up |
| XLOC_l2_012931 | 0.036 | 1.676 | Up |
| PTOV1-AS1 | 0.001 | 1.675 | Up |
| TSPOAP1-AS1 | 0.050 | 1.674 | Up |
| AL359475.1 | 0.013 | 1.673 | Up |
| G049463 | 0.038 | 1.673 | Up |
| CATG00000061635.1 | 0.040 | 1.672 | Up |
| CATG00000029387.1 | 0.011 | 1.671 | Up |
| LINC01301 | 0.018 | 1.667 | Up |
| LINC01082 | 0.025 | 1.667 | Up |
| AC000032.1 | 0.043 | 1.664 | Up |
| G001272 | 0.011 | 1.664 | Up |
| LINC00313 | 0.024 | 1.662 | Up |
| G007194 | 0.010 | 1.662 | Up |
| ASMT | 0.046 | 1.661 | Up |
| FAM95B1 | 0.031 | 1.660 | Up |
| AC005786.3 | 0.050 | 1.659 | Up |
| AL022345.4 | 0.014 | 1.658 | Up |
| DNAJC9-AS1 | 0.042 | 1.657 | Up |
| MFAP4 | 0.011 | 1.656 | Up |
| XLOC_003601 | 0.025 | 1.655 | Up |
| AL096828.3 | 0.049 | 1.654 | Up |
| PHF19 | 0.015 | 1.652 | Up |
| AC022784.1 | 0.001 | 1.651 | Up |
| AC084125.4 | 0.049 | 1.651 | Up |
| AC092447.5 | 0.041 | 1.650 | Up |
| LINC02440 | 0.048 | 1.647 | Up |
| LINC00454 | 0.029 | 1.647 | Up |
| AC063979.2 | 0.036 | 1.647 | Up |
| AL592309.2 | 0.012 | 1.646 | Up |
| AP000880.1 | 0.000 | 1.645 | Up |
| AC092436.4 | 0.048 | 1.645 | Up |

| | | | |
|---|---|---|---|
| RMDN2-AS1 | 0.021 | 1.643 | Up |
| DANT2 | 0.031 | 1.643 | Up |
| AC022007.1 | 0.023 | 1.642 | Up |
| AP003390.2 | 0.004 | 1.641 | Up |
| LINC02130 | 0.048 | 1.641 | Up |
| ATPAF1 | 0.002 | 1.641 | Up |
| HLA-DQB1-AS1 | 0.014 | 1.639 | Up |
| ARID3A | 0.039 | 1.638 | Up |
| G057325 | 0.032 | 1.637 | Up |
| AC132938.3 | 0.033 | 1.637 | Up |
| CATG00000064929.1 | 0.040 | 1.636 | Up |
| XLOC_004624 | 0.041 | 1.635 | Up |
| GAS6-AS1 | 0.016 | 1.634 | Up |
| G077476 | 0.017 | 1.631 | Up |
| GSE61474_XLOC_032653 | 0.008 | 1.630 | Up |
| CAPS2 | 0.035 | 1.624 | Up |
| AC079610.1 | 0.030 | 1.624 | Up |
| CATG00000028512.1 | 0.041 | 1.624 | Up |
| LINC00404 | 0.028 | 1.624 | Up |
| AC011352.1 | 0.013 | 1.623 | Up |
| AC134978.1 | 0.028 | 1.623 | Up |
| FOXP4-AS1 | 0.020 | 1.623 | Up |
| SPZ1 | 0.025 | 1.623 | Up |
| THRA1/BTR | 0.018 | 1.621 | Up |
| AC004974.1 | 0.004 | 1.621 | Up |
| LINC01865 | 0.027 | 1.620 | Up |
| AC103974.1 | 0.043 | 1.620 | Up |
| RP11-782C8.2 | 0.049 | 1.619 | Up |
| AC111198.1 | 0.030 | 1.618 | Up |
| NTRK3 | 0.032 | 1.617 | Up |
| AP000146.1 | 0.012 | 1.617 | Up |
| CATG00000022086.1 | 0.030 | 1.616 | Up |
| EVX1-AS | 0.032 | 1.615 | Up |
| LINC00221 | 0.031 | 1.615 | Up |
| G029855 | 0.013 | 1.614 | Up |
| AC010267.1 | 0.032 | 1.614 | Up |
| COL4A2 | 0.022 | 1.613 | Up |
| MYOZ3 | 0.014 | 1.612 | Up |
| KRI1 | 0.029 | 1.609 | Up |
| AC093072.1 | 0.035 | 1.609 | Up |
| GS1-24F4.2 | 0.004 | 1.608 | Up |
| ZNF385D-AS2 | 0.023 | 1.607 | Up |
| XLOC_002124 | 0.040 | 1.605 | Up |
| LINC00535 | 0.031 | 1.604 | Up |
| AL589923.1 | 0.018 | 1.603 | Up |
| AC105053.1 | 0.025 | 1.601 | Up |
| C9orf135-DT | 0.042 | 1.600 | Up |
| PTPRG-AS1 | 0.018 | 1.598 | Up |
| AF111169.3 | 0.019 | 1.597 | Up |
| DALRD3 | 0.010 | 1.596 | Up |
| AC010531.5 | 0.016 | 1.595 | Up |
| TM2D1 | 0.045 | 1.594 | Up |
| GPRC5D-AS1 | 0.047 | 1.593 | Up |
| FAM182A | 0.046 | 1.591 | Up |
| AC007728.2 | 0.048 | 1.590 | Up |
| CATG00000033949.1 | 0.029 | 1.588 | Up |
| AL365184.1 | 0.016 | 1.587 | Up |
| AC078785.1 | 0.023 | 1.587 | Up |
| AP000915.1 | 0.034 | 1.586 | Up |
| MRPS30-DT | 0.044 | 1.586 | Up |
| DHRS7 | 0.002 | 1.585 | Up |
| AP001010.1 | 0.042 | 1.585 | Up |
| ITFG2 | 0.009 | 1.584 | Up |
| AC091100.1 | 0.014 | 1.583 | Up |
| G041680 | 0.037 | 1.581 | Up |

| | | | |
|---|---|---|---|
| GSE61474_XLOC_065085 | 0.041 | 1.581 | Up |
| CATG00000048939.1 | 0.009 | 1.580 | Up |
| LINC01300 | 0.021 | 1.580 | Up |
| AC124057.1 | 0.041 | 1.579 | Up |
| AC091133.2 | 0.018 | 1.577 | Up |
| LINC00491 | 0.039 | 1.576 | Up |
| G074854 | 0.041 | 1.573 | Up |
| CD27-AS1 | 0.044 | 1.572 | Up |
| LINC02154 | 0.048 | 1.570 | Up |
| CATG00000112861.1 | 0.020 | 1.570 | Up |
| AC078950.1 | 0.046 | 1.569 | Up |
| LINC00482 | 0.047 | 1.568 | Up |
| CROCCP2 | 0.034 | 1.568 | Up |
| TXNDC12-AS1 | 0.040 | 1.566 | Up |
| FAM74A1 | 0.002 | 1.566 | Up |
| AC018647.1 | 0.043 | 1.564 | Up |
| AC138207.4 | 0.030 | 1.564 | Up |
| DPH6 | 0.011 | 1.563 | Up |
| LINC02137 | 0.045 | 1.563 | Up |
| AC005293.1 | 0.049 | 1.563 | Up |
| AL359399.1 | 0.036 | 1.563 | Up |
| CATG00000018991.1 | 0.004 | 1.562 | Up |
| CATG00000059503.1 | 0.043 | 1.561 | Up |
| BCL2L12 | 0.014 | 1.560 | Up |
| MPPE1 | 0.000 | 1.557 | Up |
| LINC00939 | 0.017 | 1.555 | Up |
| G029540 | 0.014 | 1.555 | Up |
| PIN1 | 0.029 | 1.555 | Up |
| TP53TG1 | 0.004 | 1.555 | Up |
| AC008164.1 | 0.022 | 1.554 | Up |
| LINC01120 | 0.022 | 1.554 | Up |
| G079121 | 0.041 | 1.554 | Up |
| LINC00350 | 0.006 | 1.552 | Up |
| AP003557.1 | 0.036 | 1.551 | Up |
| LINC01866 | 0.037 | 1.551 | Up |
| CATG00000057892.1 | 0.034 | 1.551 | Up |
| LUNAR1 | 0.049 | 1.550 | Up |
| BACE1 | 0.039 | 1.547 | Up |
| G048412 | 0.003 | 1.547 | Up |
| FAM230B | 0.028 | 1.546 | Up |
| AP000553.2 | 0.012 | 1.546 | Up |
| AL807752.3 | 0.034 | 1.546 | Up |
| AL049569.1 | 0.002 | 1.546 | Up |
| AL049557.1 | 0.034 | 1.544 | Up |
| LINC02386 | 0.024 | 1.542 | Up |
| LOC442132 | 0.026 | 1.540 | Up |
| SNRK-AS1 | 0.038 | 1.540 | Up |
| AC113133.1 | 0.047 | 1.539 | Up |
| ATG10-IT1 | 0.003 | 1.539 | Up |
| AC105105.1 | 0.004 | 1.538 | Up |
| MCM3AP-AS1 | 0.027 | 1.534 | Up |
| LINC00114 | 0.006 | 1.533 | Up |
| SPINT1-AS1 | 0.041 | 1.532 | Up |
| AP001107.8 | 0.016 | 1.530 | Up |
| BIG-lncRNA-1398 | 0.024 | 1.530 | Up |
| AL136988.1 | 0.042 | 1.526 | Up |
| AP000785.1 | 0.035 | 1.526 | Up |
| TTC28-AS1 | 0.010 | 1.524 | Up |
| AC022201.2 | 0.021 | 1.524 | Up |
| NR2F2-AS1 | 0.038 | 1.524 | Up |
| G037800 | 0.014 | 1.523 | Up |
| AC034102.4 | 0.013 | 1.519 | Up |
| AL133304.3 | 0.046 | 1.517 | Up |
| AL354861.3 | 0.031 | 1.517 | Up |
| CATG00000042562.1 | 0.040 | 1.515 | Up |

| | | | |
|---|---|---|---|
| AC010894.2 | 0.017 | 1.514 | Up |
| AC097467.3 | 0.035 | 1.510 | Up |
| SLC12A9 | 0.047 | 1.507 | Up |
| LINC01848 | 0.050 | 1.507 | Up |
| AL451060.1 | 0.033 | 1.507 | Up |
| G086541 | 0.012 | 1.507 | Up |
| ATXN3 | 0.005 | 1.505 | Up |
| BIG-lncRNA-1653 | 0.015 | 1.505 | Up |
| XLOC_006122 | 0.018 | 1.504 | Up |
| AC007923.1 | 0.040 | 1.503 | Up |
| G026354 | 0.019 | 1.503 | Up |
| SMA4 | 0.007 | 1.503 | Up |
| AC008609.1 | 0.016 | 1.502 | Up |
| G030084 | 0.050 | 1.502 | Up |
| AC005523.2 | 0.037 | 1.501 | Up |
| AC000068.2 | 0.024 | 1.501 | Up |
| AC021422.1 | 0.044 | 1.500 | Up |
| LINC01873 | 0.005 | 1.500 | Up |
| CCDC63 | 0.041 | 1.499 | Up |
| LINC01282 | 0.045 | 1.499 | Up |
| GSE1 | 0.003 | 1.497 | Up |
| XLOC_005778 | 0.048 | 1.497 | Up |
| AC004596.1 | 0.040 | 1.497 | Up |
| AL513008.1 | 0.030 | 1.494 | Up |
| LRRC75A-AS1 | 0.029 | 1.493 | Up |
| G031384 | 0.003 | 1.493 | Up |
| G051925 | 0.049 | 1.492 | Up |
| STK25 | 0.020 | 1.492 | Up |
| G059684 | 0.049 | 1.492 | Up |
| B9D1 | 0.009 | 1.490 | Up |
| MEG3 | 0.005 | 1.490 | Up |
| CATG00000088944.1 | 0.042 | 1.490 | Up |
| AC008735.1 | 0.034 | 1.488 | Up |
| AC114752.2 | 0.043 | 1.487 | Up |
| ALMS1-IT1 | 0.045 | 1.487 | Up |
| LINC01715 | 0.021 | 1.487 | Up |
| G075474 | 0.006 | 1.486 | Up |
| XLOC_012033 | 0.037 | 1.486 | Up |
| LINC01729 | 0.000 | 1.485 | Up |
| FLVCR1-DT | 0.025 | 1.485 | Up |
| CARD8-AS1 | 0.008 | 1.484 | Up |
| G078168 | 0.035 | 1.483 | Up |
| AC087289.4 | 0.034 | 1.481 | Up |
| AC008736.1 | 0.048 | 1.481 | Up |
| AC011446.2 | 0.024 | 1.481 | Up |
| UBOX5-AS1 | 0.010 | 1.480 | Up |
| G029129 | 0.004 | 1.480 | Up |
| PCAT7 | 0.023 | 1.478 | Up |
| ABHD14B | 0.020 | 1.477 | Up |
| AC098487.1 | 0.048 | 1.477 | Up |
| PC | 0.015 | 1.476 | Up |
| CATG00000020477.1 | 0.039 | 1.473 | Up |
| CATG00000002414.1 | 0.021 | 1.473 | Up |
| AC009163.3 | 0.048 | 1.472 | Up |
| G045363 | 0.032 | 1.471 | Up |
| SYS1-DBNDD2 | 0.013 | 1.470 | Up |
| NAV2-AS4 | 0.045 | 1.469 | Up |
| AL031772.1 | 0.015 | 1.468 | Up |
| AL157871.1 | 0.039 | 1.468 | Up |
| TXNDC9 | 0.009 | 1.467 | Up |
| SERTAD4-AS1 | 0.011 | 1.466 | Up |
| RP11-640M9.1 | 0.015 | 1.463 | Up |
| AC007285.1 | 0.037 | 1.463 | Up |
| G069140 | 0.012 | 1.463 | Up |
| AC009656.1 | 0.047 | 1.461 | Up |

| | | | |
|---|---|---|---|
| SURF6 | 0.041 | 1.461 | Up |
| AL627443.1 | 0.000 | 1.459 | Up |
| CATG00000047720.1 | 0.002 | 1.458 | Up |
| G048199 | 0.036 | 1.458 | Up |
| LINC01040 | 0.039 | 1.457 | Up |
| AC016245.1 | 0.047 | 1.457 | Up |
| XLOC_007866 | 0.026 | 1.456 | Up |
| XLOC_013235 | 0.040 | 1.455 | Up |
| CATG00000102375.1 | 0.021 | 1.454 | Up |
| AC110285.1 | 0.041 | 1.453 | Up |
| AC069213.1 | 0.033 | 1.453 | Up |
| AL158825.2 | 0.030 | 1.450 | Up |
| AC011625.1 | 0.016 | 1.450 | Up |
| FAM27E2 | 0.047 | 1.449 | Up |
| TSPAN32 | 0.016 | 1.446 | Up |
| CATG00000056163.1 | 0.046 | 1.445 | Up |
| MGC15885 | 0.021 | 1.445 | Up |
| ADAMTS13 | 0.050 | 1.443 | Up |
| LINC01919 | 0.017 | 1.442 | Up |
| CCDC37-DT | 0.042 | 1.441 | Up |
| AC137056.1 | 0.046 | 1.440 | Up |
| ZNF671 | 0.025 | 1.440 | Up |
| RAI1-AS1 | 0.010 | 1.440 | Up |
| AC008132.13 | 0.012 | 1.439 | Up |
| SNX22 | 0.021 | 1.436 | Up |
| CATG00000077438.1 | 0.011 | 1.436 | Up |
| AL360181.1 | 0.009 | 1.436 | Up |
| RNF212B | 0.011 | 1.435 | Up |
| AC008869.1 | 0.049 | 1.433 | Up |
| MIR503HG | 0.046 | 1.432 | Up |
| HOXA-AS3 | 0.028 | 1.432 | Up |
| LINC02166 | 0.035 | 1.431 | Up |
| LEF1-AS1 | 0.026 | 1.431 | Up |
| A1BG | 0.001 | 1.428 | Up |
| ARHGAP26-AS1 | 0.017 | 1.428 | Up |
| ITGB2-AS1 | 0.003 | 1.428 | Up |
| COL22A1 | 0.005 | 1.427 | Up |
| AC022150.2 | 0.033 | 1.426 | Up |
| TSPEAR-AS1 | 0.009 | 1.426 | Up |
| AC211486.5 | 0.048 | 1.424 | Up |
| CXCL12 | 0.037 | 1.423 | Up |
| CATG00000013703.1 | 0.030 | 1.423 | Up |
| CATG00000031910.1 | 0.014 | 1.422 | Up |
| AC105450.1 | 0.028 | 1.421 | Up |
| G021483 | 0.041 | 1.421 | Up |
| AL391415.1 | 0.033 | 1.420 | Up |
| AC010761.4 | 0.037 | 1.420 | Up |
| AC068397.2 | 0.003 | 1.420 | Up |
| LINC02411 | 0.040 | 1.416 | Up |
| AC012313.1 | 0.015 | 1.415 | Up |
| PLBD1-AS1 | 0.034 | 1.414 | Up |
| AC011389.1 | 0.030 | 1.413 | Up |
| TCL6 | 0.032 | 1.410 | Up |
| ZNF667-AS1 | 0.003 | 1.410 | Up |
| TAF12 | 0.008 | 1.409 | Up |
| AC006037.1 | 0.034 | 1.408 | Up |
| LOC101927278 | 0.036 | 1.408 | Up |
| HOXB-AS3 | 0.005 | 1.402 | Up |
| AP000662.1 | 0.047 | 1.402 | Up |
| NOP53-AS1 | 0.021 | 1.399 | Up |
| RTCA-AS1 | 0.049 | 1.398 | Up |
| YPEL4 | 0.009 | 1.397 | Up |
| SRRM2-AS1 | 0.014 | 1.396 | Up |
| AC074389.2 | 0.023 | 1.396 | Up |
| AC015712.4 | 0.005 | 1.396 | Up |

| | | | |
|---|---|---|---|
| CXCL16 | 0.021 | 1.393 | Up |
| PDE4DIP | 0.025 | 1.389 | Up |
| XLOC_007476 | 0.032 | 1.388 | Up |
| AC036103.1 | 0.024 | 1.385 | Up |
| G016623 | 0.025 | 1.384 | Up |
| AC022113.2 | 0.026 | 1.381 | Up |
| DSCAM-IT1 | 0.038 | 1.378 | Up |
| CAPN10-DT | 0.007 | 1.376 | Up |
| LINC01680 | 0.015 | 1.376 | Up |
| LINC00927 | 0.039 | 1.376 | Up |
| NALT1 | 0.010 | 1.373 | Up |
| AC063980.1 | 0.027 | 1.373 | Up |
| AC004943.1 | 0.006 | 1.370 | Up |
| AL590369.1 | 0.044 | 1.369 | Up |
| XLOC_001449 | 0.037 | 1.369 | Up |
| Z98949.1 | 0.015 | 1.369 | Up |
| AC098587.1 | 0.018 | 1.369 | Up |
| MIR663AHG | 0.035 | 1.367 | Up |
| LRRC23 | 0.013 | 1.366 | Up |
| AC129507.4 | 0.038 | 1.366 | Up |
| MLPH | 0.037 | 1.363 | Up |
| LINC01845 | 0.022 | 1.360 | Up |
| PGM5-AS1 | 0.006 | 1.359 | Up |
| AC020659.2 | 0.039 | 1.358 | Up |
| BIG-lncRNA-1075 | 0.011 | 1.357 | Up |
| AC083949.1 | 0.025 | 1.356 | Up |
| CATG00000100625.1 | 0.004 | 1.355 | Up |
| LINC02247 | 0.043 | 1.353 | Up |
| G021071 | 0.045 | 1.350 | Up |
| AC073957.2 | 0.047 | 1.350 | Up |
| AP000289.1 | 0.044 | 1.349 | Up |
| NAIF1 | 0.038 | 1.348 | Up |
| G007039 | 0.008 | 1.346 | Up |
| AC108449.3 | 0.004 | 1.343 | Up |
| LINC01894 | 0.045 | 1.343 | Up |
| NAMA | 0.009 | 1.340 | Up |
| LRRC74B | 0.009 | 1.331 | Up |
| AL161449.2 | 0.023 | 1.329 | Up |
| AC005253.1 | 0.038 | 1.328 | Up |
| AC011603.2 | 0.012 | 1.326 | Up |
| ZNF273 | 0.026 | 1.325 | Up |
| LINC02088 | 0.020 | 1.324 | Up |
| CATG00000071146.1 | 0.023 | 1.323 | Up |
| AC009095.1 | 0.020 | 1.323 | Up |
| AP000787.1 | 0.024 | 1.322 | Up |
| AL592486.1 | 0.016 | 1.321 | Up |
| XLOC_006297 | 0.010 | 1.317 | Up |
| B3GALT5-AS1 | 0.043 | 1.316 | Up |
| LINC00648 | 0.041 | 1.315 | Up |
| IPO9-AS1 | 0.003 | 1.314 | Up |
| CATG00000097276.1 | 0.003 | 1.311 | Up |
| LINC00462 | 0.004 | 1.311 | Up |
| IL11RA | 0.018 | 1.309 | Up |
| INSYN2B | 0.047 | 1.308 | Up |
| AC022726.1 | 0.018 | 1.307 | Up |
| LINC00092 | 0.031 | 1.303 | Up |
| LINC02021 | 0.013 | 1.302 | Up |
| AC005618.1 | 0.003 | 1.302 | Up |
| AY927475 | 0.026 | 1.301 | Up |
| AP004608.1 | 0.007 | 1.298 | Up |
| LINC01006 | 0.011 | 1.291 | Up |
| LINC02210-CRHR1 | 0.041 | 1.291 | Up |
| PLA2G16 | 0.029 | 1.290 | Up |
| LINC01856 | 0.024 | 1.289 | Up |
| AC012668.3 | 0.043 | 1.287 | Up |

| | | | |
|---|---|---|---|
| Z99716.1 | 0.020 | 1.282 | Up |
| AL590556.3 | 0.043 | 1.280 | Up |
| SNHG1 | 0.022 | 1.278 | Up |
| AL596223.2 | 0.023 | 1.277 | Up |
| SNHG28 | 0.003 | 1.277 | Up |
| AC092574.2 | 0.013 | 1.277 | Up |
| CBR3-AS1 | 0.003 | 1.277 | Up |
| LINC01789 | 0.026 | 1.276 | Up |
| CLDN10-AS1 | 0.034 | 1.275 | Up |
| CATG00000058417.1 | 0.018 | 1.274 | Up |
| SNHG4 | 0.001 | 1.267 | Up |
| LINC02041 | 0.049 | 1.267 | Up |
| AC034231.1 | 0.038 | 1.264 | Up |
| C22orf34 | 0.041 | 1.260 | Up |
| G025902 | 0.045 | 1.259 | Up |
| AC015853.2 | 0.042 | 1.249 | Up |
| CATG00000058602.1 | 0.005 | 1.241 | Up |
| AC091979.1 | 0.046 | 1.239 | Up |
| SNHG8 | 0.041 | 1.237 | Up |
| XLOC_000352 | 0.041 | 1.231 | Up |
| MN1 | 0.047 | 1.229 | Up |
| AATBC | 0.044 | 1.228 | Up |
| AC113189.4 | 0.022 | 1.228 | Up |
| AL355974.3 | 0.043 | 1.227 | Up |
| AL353801.1 | 0.016 | 1.225 | Up |
| LIPA | 0.028 | 1.221 | Up |
| G078391 | 0.030 | 1.219 | Up |
| LOC101928911 | 0.044 | 1.216 | Up |
| AC009005.1 | 0.048 | 1.211 | Up |
| PPFIA4 | 0.043 | 1.211 | Up |
| LINC00587 | 0.040 | 1.208 | Up |
| XLOC_002534 | 0.033 | 1.206 | Up |
| PES1 | 0.034 | 1.204 | Up |
| MAP3K14-AS1 | 0.027 | 1.204 | Up |
| AL139339.1 | 0.024 | 1.202 | Up |
| AC005100.1 | 0.001 | 20.362 | Down |
| G090367 | 0.000 | 15.240 | Down |
| AC025674.2 | 0.002 | 12.870 | Down |
| CATG00000031418.1 | 0.004 | 12.035 | Down |
| AC109322.1 | 0.044 | 11.565 | Down |
| TYW5 | 0.036 | 11.362 | Down |
| DUBR | 0.009 | 11.170 | Down |
| AC122685.1 | 0.021 | 10.319 | Down |
| AP001269.4 | 0.025 | 10.156 | Down |
| AL033504.1 | 0.047 | 9.807 | Down |
| ITPR1-DT | 0.006 | 8.961 | Down |
| LINC01960 | 0.006 | 8.875 | Down |
| G073595 | 0.001 | 8.648 | Down |
| TAT-AS1 | 0.003 | 8.596 | Down |
| AC100812.1 | 0.037 | 8.585 | Down |
| AC017015.2 | 0.011 | 8.555 | Down |
| AC068313.1 | 0.006 | 8.514 | Down |
| LINC01815 | 0.018 | 8.473 | Down |
| AC110792.3 | 0.009 | 8.421 | Down |
| AL590560.1 | 0.026 | 8.367 | Down |
| AC087588.2 | 0.017 | 8.233 | Down |
| AL353148.1 | 0.001 | 8.226 | Down |
| LINC00343 | 0.003 | 8.190 | Down |
| BIG-lncRNA-689 | 0.023 | 7.602 | Down |
| AL356289.2 | 0.007 | 7.554 | Down |
| AC010974.1 | 0.042 | 7.542 | Down |
| XLOC_008445 | 0.004 | 7.419 | Down |
| C15orf45 | 0.005 | 7.398 | Down |
| AL157817.1 | 0.044 | 7.365 | Down |
| PAQR9-AS1 | 0.006 | 7.353 | Down |

| | | | |
|---|---|---|---|
| AC073655.1 | 0.034 | 7.352 | Down |
| AL928596.1 | 0.006 | 7.300 | Down |
| AC073389.3 | 0.003 | 7.286 | Down |
| LINC00854 | 0.025 | 7.230 | Down |
| LINC00458 | 0.025 | 7.189 | Down |
| DLGAP1-AS1 | 0.025 | 6.957 | Down |
| AC092794.2 | 0.025 | 6.949 | Down |
| TRMT13 | 0.006 | 6.910 | Down |
| AL021918.3 | 0.047 | 6.825 | Down |
| AL832168 | 0.023 | 6.800 | Down |
| AP001610.2 | 0.040 | 6.724 | Down |
| AL031432.3 | 0.000 | 6.671 | Down |
| AC099684.1 | 0.016 | 6.668 | Down |
| JMJD1C | 0.000 | 6.652 | Down |
| KC877392.1 | 0.025 | 6.614 | Down |
| KIAA1191 | 0.034 | 6.568 | Down |
| AC087477.4 | 0.004 | 6.552 | Down |
| AC025165.4 | 0.005 | 6.466 | Down |
| G053006 | 0.023 | 6.402 | Down |
| CLEC9A | 0.002 | 6.370 | Down |
| SYNJ2-IT1 | 0.045 | 6.362 | Down |
| C6orf163 | 0.002 | 6.338 | Down |
| AC010975.1 | 0.007 | 6.299 | Down |
| AC073476.3 | 0.024 | 6.291 | Down |
| G054986 | 0.021 | 6.257 | Down |
| G075949 | 0.012 | 6.229 | Down |
| DICER1-AS1 | 0.020 | 6.170 | Down |
| AC100803.2 | 0.012 | 6.109 | Down |
| AC078883.1 | 0.046 | 6.108 | Down |
| AL035541.1 | 0.045 | 6.087 | Down |
| AC103876.1 | 0.016 | 6.059 | Down |
| XLOC_006565 | 0.009 | 6.031 | Down |
| LINC01829 | 0.003 | 5.900 | Down |
| GJB7 | 0.001 | 5.889 | Down |
| NLGN4Y | 0.025 | 5.861 | Down |
| BX284668.5 | 0.016 | 5.855 | Down |
| LINC00492 | 0.008 | 5.809 | Down |
| AC025428.2 | 0.006 | 5.728 | Down |
| FGF14-AS1 | 0.002 | 5.726 | Down |
| G018844 | 0.006 | 5.703 | Down |
| DLC1 | 0.043 | 5.703 | Down |
| LINC01605 | 0.004 | 5.684 | Down |
| LINC00993 | 0.017 | 5.650 | Down |
| AL160272.1 | 0.007 | 5.642 | Down |
| G061260 | 0.028 | 5.636 | Down |
| LINC02274 | 0.001 | 5.619 | Down |
| AC097487.1 | 0.013 | 5.585 | Down |
| AC078955.1 | 0.010 | 5.555 | Down |
| AC100835.2 | 0.035 | 5.554 | Down |
| PURPL | 0.009 | 5.501 | Down |
| AC010181.2 | 0.006 | 5.497 | Down |
| AC090772.3 | 0.001 | 5.451 | Down |
| G024277 | 0.011 | 5.448 | Down |
| AL512413.1 | 0.006 | 5.422 | Down |
| AC243830.2 | 0.017 | 5.340 | Down |
| AC079062.1 | 0.008 | 5.298 | Down |
| LINC00261 | 0.016 | 5.245 | Down |
| AC022306.3 | 0.028 | 5.227 | Down |
| AC124784.1 | 0.041 | 5.179 | Down |
| AC040174.2 | 0.023 | 5.170 | Down |
| PLLP | 0.017 | 5.164 | Down |
| AL358074.1 | 0.032 | 5.161 | Down |
| AC074124.1 | 0.024 | 5.155 | Down |
| AC024145.1 | 0.021 | 5.134 | Down |
| AC009754.1 | 0.002 | 5.126 | Down |

| | | | |
|---|---|---|---|
| AC010834.2 | 0.008 | 5.125 | Down |
| AL034347.1 | 0.046 | 5.094 | Down |
| BX284656.2 | 0.009 | 5.081 | Down |
| INF2 | 0.026 | 5.051 | Down |
| G031475 | 0.006 | 5.022 | Down |
| G052829 | 0.004 | 5.019 | Down |
| VIPR1-AS1 | 0.020 | 5.015 | Down |
| LINC00518 | 0.031 | 5.002 | Down |
| AC093909.2 | 0.024 | 4.966 | Down |
| MED21 | 0.017 | 4.962 | Down |
| ZNF609 | 0.019 | 4.942 | Down |
| CATG00000074437.1 | 0.004 | 4.929 | Down |
| AC023830.3 | 0.021 | 4.911 | Down |
| AC004872.1 | 0.031 | 4.907 | Down |
| AC016383.1 | 0.035 | 4.867 | Down |
| AC107396.1 | 0.035 | 4.859 | Down |
| TSL | 0.011 | 4.831 | Down |
| SOX2-OT | 0.014 | 4.804 | Down |
| RFX3-AS1 | 0.028 | 4.785 | Down |
| SH2D1A | 0.020 | 4.781 | Down |
| AC239803.1 | 0.005 | 4.766 | Down |
| AP001782.1 | 0.013 | 4.761 | Down |
| AC093725.1 | 0.044 | 4.739 | Down |
| AC097478.1 | 0.049 | 4.726 | Down |
| AL138689.1 | 0.012 | 4.726 | Down |
| AC119403.1 | 0.033 | 4.725 | Down |
| AL117339.4 | 0.004 | 4.713 | Down |
| AC010105.1 | 0.050 | 4.665 | Down |
| G010647 | 0.015 | 4.664 | Down |
| TOR1A | 0.030 | 4.662 | Down |
| BRI3 | 0.001 | 4.662 | Down |
| AP001269.1 | 0.030 | 4.637 | Down |
| STARD13 | 0.020 | 4.585 | Down |
| AF117829.1 | 0.019 | 4.583 | Down |
| G033488 | 0.047 | 4.581 | Down |
| LNCAROD | 0.011 | 4.568 | Down |
| LINC00921 | 0.014 | 4.567 | Down |
| G047388 | 0.034 | 4.561 | Down |
| XLOC_011403 | 0.022 | 4.554 | Down |
| XLOC_003852 | 0.041 | 4.532 | Down |
| AC020913.1 | 0.003 | 4.529 | Down |
| AF064860.1 | 0.009 | 4.526 | Down |
| LINC00887 | 0.012 | 4.524 | Down |
| G023288 | 0.009 | 4.510 | Down |
| CYSTM1 | 0.034 | 4.500 | Down |
| AC103879.1 | 0.043 | 4.480 | Down |
| G060775 | 0.007 | 4.479 | Down |
| AL031770.1 | 0.004 | 4.469 | Down |
| G013248 | 0.013 | 4.436 | Down |
| G051937 | 0.002 | 4.402 | Down |
| AC145207.1 | 0.023 | 4.376 | Down |
| G065722 | 0.005 | 4.349 | Down |
| LINC02217 | 0.023 | 4.338 | Down |
| AC090159.1 | 0.022 | 4.325 | Down |
| G000859 | 0.025 | 4.316 | Down |
| AC008429.1 | 0.014 | 4.308 | Down |
| AC073525.1 | 0.024 | 4.299 | Down |
| LINC00658 | 0.007 | 4.282 | Down |
| LINC01510 | 0.026 | 4.274 | Down |
| XLOC_012600 | 0.034 | 4.274 | Down |
| CATG00000098782.1 | 0.050 | 4.274 | Down |
| AL118522.1 | 0.032 | 4.253 | Down |
| G064142 | 0.019 | 4.243 | Down |
| G078311 | 0.036 | 4.235 | Down |
| LINC00903 | 0.005 | 4.210 | Down |

| | | | |
|---|---|---|---|
| AL355516.1 | 0.020 | 4.208 | Down |
| CERS6-AS1 | 0.003 | 4.199 | Down |
| AC010327.3 | 0.031 | 4.194 | Down |
| LINC01727 | 0.027 | 4.184 | Down |
| PACRG-AS2 | 0.033 | 4.183 | Down |
| G075886 | 0.011 | 4.160 | Down |
| AC105411.1 | 0.008 | 4.127 | Down |
| CATG00000089573.1 | 0.031 | 4.126 | Down |
| AC026462.5 | 0.043 | 4.120 | Down |
| AC139713.2 | 0.050 | 4.104 | Down |
| AL353611.1 | 0.000 | 4.095 | Down |
| SEPT2 | 0.012 | 4.093 | Down |
| RABL2A | 0.002 | 4.087 | Down |
| ZSCAN10 | 0.049 | 4.073 | Down |
| AL354956.1 | 0.043 | 4.067 | Down |
| AL357518.1 | 0.008 | 4.037 | Down |
| GHc-362H12.3 | 0.019 | 4.021 | Down |
| CATG00000014027.1 | 0.012 | 4.015 | Down |
| G043364 | 0.033 | 4.013 | Down |
| G060418 | 0.033 | 4.010 | Down |
| LINC02212 | 0.012 | 3.999 | Down |
| AL138885.3 | 0.000 | 3.986 | Down |
| XLOC_001745 | 0.009 | 3.974 | Down |
| LINC01741 | 0.014 | 3.971 | Down |
| AC004448.3 | 0.010 | 3.965 | Down |
| MIR2117HG | 0.010 | 3.961 | Down |
| MIR22HG | 0.018 | 3.957 | Down |
| TCF12 | 0.020 | 3.951 | Down |
| AL392046.1 | 0.025 | 3.944 | Down |
| GATA3-AS1 | 0.005 | 3.919 | Down |
| BIG-lncRNA-945 | 0.004 | 3.917 | Down |
| MACROD2-AS1 | 0.022 | 3.905 | Down |
| AL356108.1 | 0.017 | 3.877 | Down |
| MCM9 | 0.027 | 3.862 | Down |
| AC006058.3 | 0.002 | 3.861 | Down |
| G005560 | 0.007 | 3.861 | Down |
| LAMTOR5-AS1 | 0.027 | 3.845 | Down |
| DHRS4-AS1 | 0.015 | 3.844 | Down |
| AK303572 | 0.011 | 3.833 | Down |
| RARA-AS1 | 0.004 | 3.831 | Down |
| NBPF9 | 0.003 | 3.831 | Down |
| G027402 | 0.010 | 3.831 | Down |
| KLHL3 | 0.009 | 3.812 | Down |
| LINC01065 | 0.028 | 3.808 | Down |
| G075334 | 0.026 | 3.806 | Down |
| G021002 | 0.047 | 3.797 | Down |
| FLJ37035 | 0.027 | 3.791 | Down |
| AC006205.2 | 0.006 | 3.790 | Down |
| CATG00000019978.1 | 0.007 | 3.786 | Down |
| G056731 | 0.028 | 3.763 | Down |
| BIG-lncRNA-959 | 0.024 | 3.754 | Down |
| LINC01607 | 0.005 | 3.748 | Down |
| BX664615.2 | 0.031 | 3.746 | Down |
| AC091729.3 | 0.043 | 3.736 | Down |
| G048880 | 0.040 | 3.734 | Down |
| INTS6-AS1 | 0.005 | 3.732 | Down |
| ZNF346-IT1 | 0.044 | 3.730 | Down |
| WDR11-AS1 | 0.041 | 3.715 | Down |
| CATG00000081626.1 | 0.025 | 3.711 | Down |
| AL353651.3 | 0.024 | 3.710 | Down |
| AC105285.1 | 0.011 | 3.704 | Down |
| AP000346.1 | 0.033 | 3.689 | Down |
| AC018926.2 | 0.029 | 3.680 | Down |
| LINC02557 | 0.004 | 3.678 | Down |
| AC037441.1 | 0.036 | 3.677 | Down |

| | | | |
|---|---|---|---|
| AC068533.3 | 0.008 | 3.668 | Down |
| AC002070.1 | 0.018 | 3.662 | Down |
| AC011467.1 | 0.002 | 3.651 | Down |
| AC104002.3 | 0.033 | 3.650 | Down |
| FMO5 | 0.043 | 3.644 | Down |
| XLOC_009198 | 0.019 | 3.633 | Down |
| AC105001.1 | 0.035 | 3.603 | Down |
| TMEM138 | 0.014 | 3.602 | Down |
| LINC01173 | 0.015 | 3.602 | Down |
| LINC02482 | 0.008 | 3.593 | Down |
| BIG-lncRNA-489 | 0.004 | 3.586 | Down |
| AC010326.3 | 0.008 | 3.574 | Down |
| LINC00599 | 0.016 | 3.567 | Down |
| SPATA41 | 0.018 | 3.567 | Down |
| G011570 | 0.000 | 3.525 | Down |
| AL356124.2 | 0.041 | 3.518 | Down |
| WWC2-AS1 | 0.041 | 3.513 | Down |
| AC105177.1 | 0.017 | 3.502 | Down |
| AL353747.3 | 0.014 | 3.490 | Down |
| XLOC_006121 | 0.007 | 3.479 | Down |
| Z49979 | 0.017 | 3.478 | Down |
| GSE61474_XLOC_060526 | 0.049 | 3.477 | Down |
| AC005363.2 | 0.017 | 3.474 | Down |
| PLCB1-IT1 | 0.005 | 3.461 | Down |
| MIR3681HG | 0.008 | 3.460 | Down |
| AC135782.1 | 0.022 | 3.457 | Down |
| G046072 | 0.026 | 3.439 | Down |
| PRC1-AS1 | 0.047 | 3.435 | Down |
| G002630 | 0.006 | 3.426 | Down |
| CPNE1 | 0.030 | 3.419 | Down |
| LINC00271 | 0.000 | 3.416 | Down |
| MOSPD1 | 0.032 | 3.415 | Down |
| C7orf65 | 0.045 | 3.412 | Down |
| CATG00000020104.1 | 0.003 | 3.406 | Down |
| LOC390705 | 0.048 | 3.406 | Down |
| G070055 | 0.023 | 3.404 | Down |
| ATP6AP1L | 0.045 | 3.403 | Down |
| AC008592.5 | 0.023 | 3.400 | Down |
| AC008088.1 | 0.047 | 3.396 | Down |
| STX18-AS1 | 0.018 | 3.394 | Down |
| XLOC_004632 | 0.036 | 3.389 | Down |
| LINC01833 | 0.025 | 3.380 | Down |
| XLOC_006249 | 0.011 | 3.373 | Down |
| AC136475.9 | 0.028 | 3.371 | Down |
| BCAR4 | 0.015 | 3.368 | Down |
| AC138331.1 | 0.034 | 3.366 | Down |
| AC090515.2 | 0.016 | 3.364 | Down |
| AL355432.1 | 0.035 | 3.358 | Down |
| ZIM2-AS1 | 0.019 | 3.355 | Down |
| CATG00000116243.1 | 0.010 | 3.352 | Down |
| GPC5-AS1 | 0.005 | 3.348 | Down |
| AC036176.3 | 0.043 | 3.343 | Down |
| B4GALT1-AS1 | 0.012 | 3.332 | Down |
| AC011243.1 | 0.007 | 3.322 | Down |
| LINC00327 | 0.039 | 3.322 | Down |
| G052480 | 0.014 | 3.312 | Down |
| G082347 | 0.004 | 3.309 | Down |
| BIG-lncRNA-444 | 0.039 | 3.305 | Down |
| AC078851.1 | 0.003 | 3.289 | Down |
| AC000123.1 | 0.008 | 3.284 | Down |
| AP000561.1 | 0.027 | 3.262 | Down |
| XLOC_007148 | 0.018 | 3.259 | Down |
| CATG00000056759.1 | 0.048 | 3.257 | Down |
| SH3D19 | 0.034 | 3.253 | Down |
| AP001961.1 | 0.001 | 3.233 | Down |

| | | | |
|---|---|---|---|
| AC023983.2 | 0.017 | 3.232 | Down |
| CATG00000067089.1 | 0.047 | 3.215 | Down |
| AC104389.4 | 0.030 | 3.209 | Down |
| AL121917.2 | 0.013 | 3.207 | Down |
| AC021683.5 | 0.045 | 3.207 | Down |
| CATG00000052789.1 | 0.021 | 3.205 | Down |
| AC011477.3 | 0.017 | 3.196 | Down |
| G025399 | 0.039 | 3.196 | Down |
| PARG | 0.002 | 3.196 | Down |
| AL109920.1 | 0.008 | 3.191 | Down |
| CATG00000054624.1 | 0.041 | 3.181 | Down |
| ZRANB2-AS2 | 0.048 | 3.171 | Down |
| AL079303.1 | 0.032 | 3.167 | Down |
| G006890 | 0.034 | 3.162 | Down |
| AC099329.2 | 0.000 | 3.161 | Down |
| LINC01234 | 0.024 | 3.160 | Down |
| AC011247.1 | 0.027 | 3.158 | Down |
| PRRT3-AS1 | 0.017 | 3.157 | Down |
| PRAMENP | 0.044 | 3.155 | Down |
| G041118 | 0.015 | 3.154 | Down |
| G005587 | 0.010 | 3.150 | Down |
| G031275 | 0.032 | 3.145 | Down |
| PPP4R1-AS1 | 0.026 | 3.138 | Down |
| G061386 | 0.017 | 3.136 | Down |
| AC005837.3 | 0.014 | 3.128 | Down |
| FLJ33534 | 0.033 | 3.127 | Down |
| AC009902.2 | 0.003 | 3.127 | Down |
| G001001 | 0.046 | 3.123 | Down |
| XLOC_005549 | 0.011 | 3.122 | Down |
| F10-AS1 | 0.047 | 3.112 | Down |
| CATG00000049022.1 | 0.027 | 3.107 | Down |
| JAKMIP2-AS1 | 0.009 | 3.105 | Down |
| AC109347.1 | 0.001 | 3.105 | Down |
| AC012456.2 | 0.034 | 3.099 | Down |
| G075573 | 0.041 | 3.097 | Down |
| U95743.1 | 0.010 | 3.095 | Down |
| AC010609.1 | 0.021 | 3.091 | Down |
| AC080129.2 | 0.015 | 3.090 | Down |
| DLGAP2 | 0.036 | 3.086 | Down |
| AC012254.3 | 0.026 | 3.081 | Down |
| G055010 | 0.008 | 3.077 | Down |
| CATG00000074316.1 | 0.020 | 3.076 | Down |
| G011374 | 0.039 | 3.066 | Down |
| AC025419.1 | 0.011 | 3.056 | Down |
| AC021088.1 | 0.006 | 3.047 | Down |
| AL157414.2 | 0.011 | 3.031 | Down |
| AC010745.4 | 0.046 | 3.017 | Down |
| AC008743.1 | 0.033 | 3.015 | Down |
| EPN1 | 0.012 | 3.014 | Down |
| NAPA-AS1 | 0.033 | 3.007 | Down |
| G077679 | 0.006 | 3.000 | Down |
| AC108067.1 | 0.034 | 3.000 | Down |
| G008684 | 0.010 | 2.998 | Down |
| AL049749.1 | 0.038 | 2.997 | Down |
| XLOC_012244 | 0.019 | 2.990 | Down |
| G051748 | 0.013 | 2.985 | Down |
| AC006206.2 | 0.012 | 2.978 | Down |
| AP003119.2 | 0.013 | 2.976 | Down |
| CATG00000036157.1 | 0.034 | 2.975 | Down |
| XLOC_006034 | 0.044 | 2.974 | Down |
| AC004057.1 | 0.000 | 2.966 | Down |
| MIR193BHG | 0.017 | 2.963 | Down |
| LINC00408 | 0.003 | 2.961 | Down |
| XLOC_012379 | 0.020 | 2.949 | Down |
| AL035587.1 | 0.016 | 2.942 | Down |

| | | | |
|---|---|---|---|
| AC123768.1 | 0.018 | 2.937 | Down |
| XLOC_007407 | 0.023 | 2.937 | Down |
| ANAPC7 | 0.036 | 2.935 | Down |
| LINC01047 | 0.041 | 2.930 | Down |
| CATG00000075679.1 | 0.009 | 2.916 | Down |
| G063931 | 0.031 | 2.911 | Down |
| AL137782.1 | 0.010 | 2.911 | Down |
| XLOC_008811 | 0.001 | 2.909 | Down |
| SALRNA2 | 0.009 | 2.909 | Down |
| G029518 | 0.048 | 2.904 | Down |
| CDC37L1-DT | 0.012 | 2.901 | Down |
| CATG00000058727.1 | 0.021 | 2.899 | Down |
| CATG00000087063.1 | 0.024 | 2.894 | Down |
| AL136234.1 | 0.049 | 2.894 | Down |
| VAC14-AS1 | 0.042 | 2.890 | Down |
| AC069200.1 | 0.042 | 2.889 | Down |
| AC034268.2 | 0.013 | 2.882 | Down |
| AL122125.1 | 0.019 | 2.881 | Down |
| AL353708.2 | 0.037 | 2.880 | Down |
| G057757 | 0.017 | 2.879 | Down |
| G045626 | 0.026 | 2.873 | Down |
| AL157392.3 | 0.047 | 2.868 | Down |
| ICAM2 | 0.024 | 2.867 | Down |
| FAM167A | 0.007 | 2.864 | Down |
| GSE61474_XLOC_017415 | 0.007 | 2.862 | Down |
| AC245014.3 | 0.041 | 2.860 | Down |
| CATG00000094268.1 | 0.011 | 2.850 | Down |
| G018145 | 0.050 | 2.849 | Down |
| ABCC5-AS1 | 0.025 | 2.847 | Down |
| AC026786.2 | 0.018 | 2.844 | Down |
| G018076 | 0.043 | 2.837 | Down |
| LINC01531 | 0.037 | 2.837 | Down |
| PRH1 | 0.005 | 2.836 | Down |
| LINC00470 | 0.034 | 2.835 | Down |
| LYST | 0.030 | 2.834 | Down |
| XLOC_008430 | 0.013 | 2.831 | Down |
| AC018450.1 | 0.026 | 2.825 | Down |
| XLOC_003234 | 0.040 | 2.819 | Down |
| TCF4 | 0.038 | 2.818 | Down |
| LINC01876 | 0.032 | 2.818 | Down |
| G024234 | 0.003 | 2.813 | Down |
| G079621 | 0.030 | 2.809 | Down |
| NRG1-IT1 | 0.013 | 2.808 | Down |
| G039382 | 0.005 | 2.807 | Down |
| AC130650.2 | 0.004 | 2.803 | Down |
| G082085 | 0.027 | 2.802 | Down |
| G000090 | 0.011 | 2.801 | Down |
| XLOC_004448 | 0.018 | 2.799 | Down |
| AL512643.1 | 0.015 | 2.798 | Down |
| MIR548A1HG | 0.027 | 2.797 | Down |
| AC092287.1 | 0.023 | 2.796 | Down |
| AK023372 | 0.027 | 2.791 | Down |
| G002513 | 0.044 | 2.789 | Down |
| ZNF451-AS1 | 0.022 | 2.785 | Down |
| SCLT1 | 0.040 | 2.784 | Down |
| MRGPRG-AS1 | 0.030 | 2.781 | Down |
| CATG00000007614.1 | 0.019 | 2.767 | Down |
| G062855 | 0.037 | 2.764 | Down |
| XLOC_006471 | 0.007 | 2.763 | Down |
| LINC00642 | 0.018 | 2.761 | Down |
| RERG | 0.026 | 2.760 | Down |
| IL12A-AS1 | 0.044 | 2.753 | Down |
| AC011500.3 | 0.047 | 2.749 | Down |
| AC016727.1 | 0.019 | 2.748 | Down |
| CATG00000089879.1 | 0.031 | 2.743 | Down |

| | | | |
|---|---|---|---|
| AC092634.5 | 0.043 | 2.742 | Down |
| XLOC_001415 | 0.049 | 2.739 | Down |
| LINC01023 | 0.044 | 2.738 | Down |
| C12orf60 | 0.026 | 2.735 | Down |
| G023046 | 0.039 | 2.731 | Down |
| AC107905.1 | 0.024 | 2.725 | Down |
| LINC01900 | 0.014 | 2.720 | Down |
| SPAG5-AS1 | 0.005 | 2.718 | Down |
| INHBA | 0.046 | 2.717 | Down |
| AC096669.1 | 0.035 | 2.715 | Down |
| AC114810.1 | 0.002 | 2.715 | Down |
| AC009502.1 | 0.047 | 2.715 | Down |
| HSD52 | 0.032 | 2.715 | Down |
| G000878 | 0.025 | 2.713 | Down |
| AP000708.1 | 0.027 | 2.712 | Down |
| LINC01844 | 0.006 | 2.702 | Down |
| LINC00893 | 0.045 | 2.702 | Down |
| CATG00000051535.1 | 0.030 | 2.701 | Down |
| CFLAR-AS1 | 0.037 | 2.697 | Down |
| XLOC_008346 | 0.009 | 2.695 | Down |
| DLGAP4-AS1 | 0.006 | 2.695 | Down |
| LOC100506858 | 0.050 | 2.695 | Down |
| AL356019.2 | 0.004 | 2.694 | Down |
| CATG00000080149.1 | 0.008 | 2.693 | Down |
| XLOC_002440 | 0.036 | 2.691 | Down |
| AC004147.3 | 0.017 | 2.689 | Down |
| LINC02574 | 0.003 | 2.685 | Down |
| MACROD2-IT1 | 0.039 | 2.684 | Down |
| LINC01179 | 0.011 | 2.681 | Down |
| G080744 | 0.021 | 2.681 | Down |
| G047816 | 0.005 | 2.680 | Down |
| CATG00000078872.1 | 0.004 | 2.678 | Down |
| AL356218.1 | 0.037 | 2.670 | Down |
| CATG00000047451.1 | 0.006 | 2.667 | Down |
| AC026979.1 | 0.029 | 2.666 | Down |
| CATG00000085471.1 | 0.001 | 2.666 | Down |
| LOC100133091 | 0.003 | 2.660 | Down |
| LINC00910 | 0.018 | 2.660 | Down |
| CATG00000061658.1 | 0.025 | 2.655 | Down |
| CEP85 | 0.005 | 2.655 | Down |
| LINC02448 | 0.021 | 2.654 | Down |
| CATG00000055976.1 | 0.028 | 2.648 | Down |
| CATG00000059733.1 | 0.023 | 2.648 | Down |
| AL683807.1 | 0.040 | 2.636 | Down |
| XLOC_002462 | 0.009 | 2.634 | Down |
| AC093462.1 | 0.047 | 2.631 | Down |
| AC108134.1 | 0.032 | 2.619 | Down |
| G083871 | 0.023 | 2.619 | Down |
| AC027328.1 | 0.004 | 2.619 | Down |
| TINCR | 0.018 | 2.618 | Down |
| AC106782.6 | 0.037 | 2.614 | Down |
| COPS5 | 0.020 | 2.612 | Down |
| G046246 | 0.045 | 2.609 | Down |
| AJ009632.2 | 0.011 | 2.609 | Down |
| AL157871.2 | 0.024 | 2.607 | Down |
| LILRA1 | 0.035 | 2.599 | Down |
| ZNF587 | 0.015 | 2.597 | Down |
| MIR3945HG | 0.013 | 2.594 | Down |
| G057319 | 0.002 | 2.593 | Down |
| ELOA-AS1 | 0.046 | 2.593 | Down |
| AC011632.2 | 0.017 | 2.591 | Down |
| LINC01166 | 0.043 | 2.586 | Down |
| AC008667.4 | 0.042 | 2.586 | Down |
| AC012676.3 | 0.025 | 2.583 | Down |
| PCAT18 | 0.010 | 2.583 | Down |

| | | | |
|---|---|---|---|
| DPP10-AS1 | 0.026 | 2.575 | Down |
| AC131157.1 | 0.046 | 2.574 | Down |
| G037127 | 0.025 | 2.572 | Down |
| AC025171.2 | 0.040 | 2.571 | Down |
| MRPS14 | 0.034 | 2.570 | Down |
| AC006207.1 | 0.002 | 2.567 | Down |
| AL137847.2 | 0.026 | 2.566 | Down |
| MACROD2 | 0.004 | 2.565 | Down |
| SERTAD2 | 0.014 | 2.565 | Down |
| AC018647.2 | 0.017 | 2.564 | Down |
| AC011511.5 | 0.048 | 2.562 | Down |
| LINC01799 | 0.044 | 2.562 | Down |
| TAB3-AS1 | 0.015 | 2.560 | Down |
| G073705 | 0.003 | 2.559 | Down |
| MIR4458HG | 0.018 | 2.552 | Down |
| AL356489.1 | 0.014 | 2.549 | Down |
| RHOQ | 0.002 | 2.547 | Down |
| CATG00000081948.1 | 0.034 | 2.546 | Down |
| AC106875.1 | 0.002 | 2.544 | Down |
| AL109811.3 | 0.021 | 2.542 | Down |
| AC241644.3 | 0.038 | 2.540 | Down |
| CATG00000053131.1 | 0.007 | 2.537 | Down |
| AL163636.1 | 0.006 | 2.537 | Down |
| TTLL7-IT1 | 0.008 | 2.531 | Down |
| AC107884.1 | 0.018 | 2.525 | Down |
| ZDHHC20-IT1 | 0.014 | 2.524 | Down |
| G036887 | 0.013 | 2.515 | Down |
| BIG-lncRNA-1299 | 0.027 | 2.513 | Down |
| CATG00000025244.1 | 0.011 | 2.509 | Down |
| LINC02201 | 0.017 | 2.507 | Down |
| AP001180.1 | 0.007 | 2.505 | Down |
| AP000894.2 | 0.015 | 2.503 | Down |
| G089953 | 0.017 | 2.498 | Down |
| CXXC1 | 0.011 | 2.498 | Down |
| OGA | 0.003 | 2.498 | Down |
| XLOC_013863 | 0.038 | 2.496 | Down |
| G014697 | 0.042 | 2.495 | Down |
| AF127577.3 | 0.004 | 2.491 | Down |
| CATG00000083252.1 | 0.011 | 2.490 | Down |
| XLOC_012965 | 0.027 | 2.488 | Down |
| CASC19 | 0.022 | 2.488 | Down |
| AL132819.1 | 0.047 | 2.486 | Down |
| LINC02202 | 0.008 | 2.485 | Down |
| AL357060.1 | 0.016 | 2.485 | Down |
| HTR2A-AS1 | 0.023 | 2.482 | Down |
| ENTPD3-AS1 | 0.008 | 2.480 | Down |
| G059378 | 0.000 | 2.477 | Down |
| TM4SF19-AS1 | 0.005 | 2.477 | Down |
| AL390760.1 | 0.031 | 2.474 | Down |
| AC018529.2 | 0.025 | 2.470 | Down |
| G082098 | 0.018 | 2.468 | Down |
| LINC01581 | 0.023 | 2.466 | Down |
| AC092326.1 | 0.040 | 2.462 | Down |
| AC012370.2 | 0.046 | 2.462 | Down |
| LINC01762 | 0.039 | 2.459 | Down |
| AP005271.1 | 0.009 | 2.459 | Down |
| CATG00000012794.1 | 0.023 | 2.459 | Down |
| AL355578.1 | 0.023 | 2.458 | Down |
| AC024610.2 | 0.038 | 2.456 | Down |
| G089911 | 0.005 | 2.455 | Down |
| CATG00000111502.1 | 0.029 | 2.455 | Down |
| AL445430.1 | 0.029 | 2.452 | Down |
| XLOC_004188 | 0.014 | 2.450 | Down |
| G051620 | 0.002 | 2.446 | Down |
| XLOC_003778 | 0.023 | 2.445 | Down |

| | | | |
|---|---|---|---|
| G060924 | 0.026 | 2.444 | Down |
| CATG00000063081.1 | 0.020 | 2.443 | Down |
| LINC01609 | 0.003 | 2.443 | Down |
| TMEM50B | 0.013 | 2.440 | Down |
| CASP8 | 0.017 | 2.440 | Down |
| G079019 | 0.042 | 2.439 | Down |
| AC009117.2 | 0.031 | 2.438 | Down |
| LINC02519 | 0.025 | 2.432 | Down |
| AC073834.1 | 0.044 | 2.429 | Down |
| CATG00000055087.1 | 0.006 | 2.428 | Down |
| XLOC_009229 | 0.017 | 2.424 | Down |
| AC003005.2 | 0.031 | 2.423 | Down |
| AP002754.1 | 0.011 | 2.423 | Down |
| AC009950.1 | 0.040 | 2.419 | Down |
| ADAM29 | 0.028 | 2.417 | Down |
| G021171 | 0.004 | 2.417 | Down |
| G014177 | 0.022 | 2.409 | Down |
| DGKE | 0.019 | 2.406 | Down |
| G029222 | 0.047 | 2.405 | Down |
| XLOC_003721 | 0.026 | 2.399 | Down |
| BIG-lncRNA-582 | 0.032 | 2.398 | Down |
| G087922 | 0.008 | 2.395 | Down |
| AL163051.2 | 0.003 | 2.393 | Down |
| G011132 | 0.045 | 2.392 | Down |
| MAPK10 | 0.010 | 2.392 | Down |
| LINC01818 | 0.013 | 2.391 | Down |
| AC092535.4 | 0.003 | 2.390 | Down |
| LINC00407 | 0.042 | 2.389 | Down |
| AC109587.1 | 0.019 | 2.389 | Down |
| RFC5 | 0.030 | 2.387 | Down |
| CATG00000036596.1 | 0.048 | 2.384 | Down |
| AMZ1 | 0.004 | 2.380 | Down |
| LINC02147 | 0.026 | 2.380 | Down |
| G064137 | 0.049 | 2.378 | Down |
| AC009163.6 | 0.028 | 2.377 | Down |
| AL356356.1 | 0.046 | 2.376 | Down |
| XLOC_000903 | 0.038 | 2.374 | Down |
| XLOC_000610 | 0.001 | 2.374 | Down |
| AP003469.1 | 0.001 | 2.366 | Down |
| AC107993.1 | 0.003 | 2.363 | Down |
| AC097724.1 | 0.048 | 2.361 | Down |
| G090197 | 0.017 | 2.360 | Down |
| AL445471.2 | 0.021 | 2.357 | Down |
| AC113414.1 | 0.049 | 2.355 | Down |
| AC010536.2 | 0.007 | 2.350 | Down |
| AC006299.1 | 0.019 | 2.346 | Down |
| CATG00000055118.1 | 0.001 | 2.346 | Down |
| G024626 | 0.005 | 2.342 | Down |
| G063055 | 0.047 | 2.339 | Down |
| CATG00000077150.1 | 0.037 | 2.338 | Down |
| AP005137.2 | 0.044 | 2.336 | Down |
| AL121672.3 | 0.033 | 2.331 | Down |
| AL365204.1 | 0.009 | 2.325 | Down |
| AL023584.2 | 0.044 | 2.321 | Down |
| AL138900.3 | 0.049 | 2.316 | Down |
| XLOC_004514 | 0.040 | 2.316 | Down |
| AC009948.1 | 0.041 | 2.312 | Down |
| AC068473.5 | 0.039 | 2.301 | Down |
| ATP2A1 | 0.019 | 2.297 | Down |
| AC006064.2 | 0.032 | 2.295 | Down |
| AL139121.1 | 0.036 | 2.294 | Down |
| LYPLA2P2 | 0.046 | 2.294 | Down |
| LINC01119 | 0.008 | 2.294 | Down |
| AL355103.1 | 0.010 | 2.292 | Down |
| G018505 | 0.005 | 2.291 | Down |

| | | | |
|---|---|---|---|
| MIR4432HG | 0.047 | 2.291 | Down |
| LINC01185 | 0.027 | 2.288 | Down |
| LINC01931 | 0.025 | 2.284 | Down |
| REEP5 | 0.019 | 2.280 | Down |
| PHACTR2-AS1 | 0.027 | 2.276 | Down |
| AC025887.1 | 0.018 | 2.275 | Down |
| AC091939.1 | 0.023 | 2.274 | Down |
| CATG00000022510.1 | 0.004 | 2.264 | Down |
| FGF14-AS2 | 0.009 | 2.262 | Down |
| LINC01861 | 0.035 | 2.261 | Down |
| LINC02246 | 0.044 | 2.259 | Down |
| LINC00167 | 0.011 | 2.255 | Down |
| AC018462.1 | 0.009 | 2.254 | Down |
| XLOC_008997 | 0.044 | 2.253 | Down |
| AC074117.1 | 0.036 | 2.253 | Down |
| CATG00000055540.1 | 0.047 | 2.253 | Down |
| TYW3 | 0.035 | 2.252 | Down |
| AC020594.1 | 0.044 | 2.249 | Down |
| AC092958.1 | 0.037 | 2.249 | Down |
| AC005165.1 | 0.022 | 2.247 | Down |
| AC125793.1 | 0.032 | 2.245 | Down |
| AC010148.1 | 0.040 | 2.243 | Down |
| ELL3 | 0.048 | 2.241 | Down |
| AC013509.1 | 0.022 | 2.233 | Down |
| TSPEAR-AS2 | 0.045 | 2.232 | Down |
| PAXBP1-AS1 | 0.048 | 2.229 | Down |
| XLOC_002175 | 0.021 | 2.229 | Down |
| LINC00942 | 0.013 | 2.228 | Down |
| XLOC_010360 | 0.003 | 2.227 | Down |
| AC018682.2 | 0.046 | 2.227 | Down |
| G090012 | 0.037 | 2.224 | Down |
| AX746920 | 0.036 | 2.220 | Down |
| RCC1 | 0.023 | 2.218 | Down |
| LINC00583 | 0.004 | 2.210 | Down |
| AC132872.1 | 0.009 | 2.207 | Down |
| AL442638.1 | 0.023 | 2.206 | Down |
| AC013470.2 | 0.011 | 2.204 | Down |
| XLOC_011620 | 0.016 | 2.197 | Down |
| XLOC_007341 | 0.046 | 2.195 | Down |
| LINC00881 | 0.045 | 2.194 | Down |
| METTL1 | 0.007 | 2.191 | Down |
| G061296 | 0.011 | 2.190 | Down |
| AF130417.1 | 0.001 | 2.188 | Down |
| CATG00000049425.1 | 0.004 | 2.184 | Down |
| G061332 | 0.019 | 2.179 | Down |
| AC010261.2 | 0.020 | 2.176 | Down |
| AC008080.1 | 0.011 | 2.175 | Down |
| AC007388.1 | 0.008 | 2.175 | Down |
| LINC02243 | 0.050 | 2.174 | Down |
| MKRN2OS | 0.022 | 2.172 | Down |
| AC003682.1 | 0.030 | 2.170 | Down |
| LINC01385 | 0.049 | 2.170 | Down |
| XLOC_013842 | 0.023 | 2.169 | Down |
| G032076 | 0.012 | 2.168 | Down |
| LINC02504 | 0.025 | 2.167 | Down |
| CATG00000093299.1 | 0.005 | 2.166 | Down |
| CATG00000063838.1 | 0.019 | 2.163 | Down |
| DARS-AS1 | 0.018 | 2.162 | Down |
| AC117465.1 | 0.014 | 2.162 | Down |
| XLOC_008965 | 0.048 | 2.161 | Down |
| AC103409.1 | 0.031 | 2.161 | Down |
| ZNF337-AS1 | 0.029 | 2.161 | Down |
| STARD7-AS1 | 0.009 | 2.160 | Down |
| Z98884.2 | 0.039 | 2.158 | Down |
| PABPC1L | 0.006 | 2.158 | Down |

| | | | |
|---|---|---|---|
| AC007493.1 | 0.006 | 2.158 | Down |
| NAALADL2-AS3 | 0.011 | 2.151 | Down |
| XLOC_007348 | 0.008 | 2.149 | Down |
| G065797 | 0.000 | 2.148 | Down |
| G036785 | 0.023 | 2.144 | Down |
| LINC01093 | 0.009 | 2.140 | Down |
| G027878 | 0.007 | 2.139 | Down |
| G064228 | 0.048 | 2.134 | Down |
| CATG00000001164.1 | 0.047 | 2.130 | Down |
| SEC11C | 0.035 | 2.128 | Down |
| KCNJ2-AS1 | 0.013 | 2.127 | Down |
| LINC00678 | 0.021 | 2.124 | Down |
| FAM27E3 | 0.026 | 2.123 | Down |
| AL133415.1 | 0.035 | 2.121 | Down |
| AL929472.2 | 0.004 | 2.119 | Down |
| AL158071.4 | 0.018 | 2.118 | Down |
| CATG00000078853.1 | 0.022 | 2.117 | Down |
| AF052103 | 0.019 | 2.116 | Down |
| AL158198.1 | 0.008 | 2.114 | Down |
| AC026691.1 | 0.017 | 2.113 | Down |
| AC093326.1 | 0.022 | 2.113 | Down |
| AC007179.2 | 0.042 | 2.112 | Down |
| RFX2 | 0.016 | 2.112 | Down |
| POLR3C | 0.002 | 2.110 | Down |
| CCDC18-AS1 | 0.045 | 2.110 | Down |
| AC023389.1 | 0.026 | 2.109 | Down |
| C8orf34-AS1 | 0.019 | 2.109 | Down |
| LINC01405 | 0.048 | 2.106 | Down |
| AP001893.1 | 0.015 | 2.104 | Down |
| XLOC_008665 | 0.037 | 2.104 | Down |
| AC120114.1 | 0.019 | 2.103 | Down |
| ACOXL-AS1 | 0.043 | 2.102 | Down |
| COG5 | 0.022 | 2.102 | Down |
| RGPD4-AS1 | 0.002 | 2.098 | Down |
| GABRE | 0.014 | 2.097 | Down |
| G060408 | 0.024 | 2.097 | Down |
| AC022601.1 | 0.001 | 2.096 | Down |
| CATG00000096753.1 | 0.001 | 2.095 | Down |
| AC096577.1 | 0.050 | 2.093 | Down |
| AP000472.1 | 0.047 | 2.091 | Down |
| XLOC_006182 | 0.018 | 2.091 | Down |
| FKRP | 0.010 | 2.089 | Down |
| AC095032.1 | 0.012 | 2.087 | Down |
| AC008700.1 | 0.039 | 2.086 | Down |
| CATG00000117623.1 | 0.019 | 2.086 | Down |
| G066688 | 0.046 | 2.084 | Down |
| LINC02020 | 0.029 | 2.082 | Down |
| IMP3 | 0.015 | 2.082 | Down |
| LINC01413 | 0.045 | 2.081 | Down |
| TRAF3IP2-AS1 | 0.049 | 2.081 | Down |
| LINC00705 | 0.039 | 2.076 | Down |
| AL583722.3 | 0.010 | 2.076 | Down |
| HDAC2-AS2 | 0.000 | 2.075 | Down |
| LINC02366 | 0.039 | 2.073 | Down |
| MIR7-3HG | 0.012 | 2.072 | Down |
| AC011410.1 | 0.017 | 2.071 | Down |
| AC007384.1 | 0.046 | 2.070 | Down |
| XLOC_010149 | 0.011 | 2.070 | Down |
| GLYR1 | 0.001 | 2.069 | Down |
| AC105393.1 | 0.027 | 2.069 | Down |
| G035903 | 0.004 | 2.069 | Down |
| G002089 | 0.002 | 2.067 | Down |
| AC114763.1 | 0.007 | 2.066 | Down |
| LINC00354 | 0.000 | 2.063 | Down |
| NCK1-DT | 0.038 | 2.059 | Down |

| | | | |
|---|---|---|---|
| AL121983.2 | 0.016 | 2.056 | Down |
| RP11-395B7.2 | 0.021 | 2.054 | Down |
| AL360267.2 | 0.015 | 2.050 | Down |
| HACE1 | 0.048 | 2.048 | Down |
| AP002748.3 | 0.022 | 2.047 | Down |
| LINC01630 | 0.040 | 2.046 | Down |
| XLOC_010373 | 0.039 | 2.045 | Down |
| TIGD6 | 0.010 | 2.044 | Down |
| AL138762.1 | 0.043 | 2.043 | Down |
| AL355073.2 | 0.025 | 2.043 | Down |
| AC006487.2 | 0.023 | 2.043 | Down |
| AC025035.1 | 0.043 | 2.041 | Down |
| AC073530.1 | 0.049 | 2.038 | Down |
| G067375 | 0.038 | 2.038 | Down |
| AL669831.5 | 0.041 | 2.035 | Down |
| SMIM10L2B-AS1 | 0.041 | 2.031 | Down |
| LINC00837 | 0.018 | 2.031 | Down |
| CATG00000063834.1 | 0.016 | 2.031 | Down |
| MIR4435-2HG | 0.031 | 2.029 | Down |
| AC022509.1 | 0.049 | 2.028 | Down |
| C15orf54 | 0.028 | 2.027 | Down |
| ZNF582-AS1 | 0.033 | 2.024 | Down |
| LINC01485 | 0.020 | 2.024 | Down |
| AC005225.2 | 0.016 | 2.024 | Down |
| XLOC_011138 | 0.015 | 2.016 | Down |
| DPH5 | 0.039 | 2.014 | Down |
| LINC02328 | 0.027 | 2.013 | Down |
| AC104211.3 | 0.009 | 2.010 | Down |
| AC114912.1 | 0.024 | 2.010 | Down |
| LINC02430 | 0.035 | 2.006 | Down |
| LINC01588 | 0.013 | 2.006 | Down |
| G086260 | 0.011 | 2.003 | Down |
| XLOC_010681 | 0.022 | 2.002 | Down |
| BDNF-AS | 0.048 | 1.999 | Down |
| AC113361.1 | 0.002 | 1.999 | Down |
| CATG00000004492.1 | 0.007 | 1.998 | Down |
| AC099566.1 | 0.017 | 1.997 | Down |
| AC005518.1 | 0.004 | 1.997 | Down |
| AC090337.2 | 0.028 | 1.996 | Down |
| AC046195.2 | 0.028 | 1.995 | Down |
| SNHG16 | 0.014 | 1.993 | Down |
| G082398 | 0.003 | 1.992 | Down |
| LINC01538 | 0.020 | 1.987 | Down |
| CATG00000065359.1 | 0.022 | 1.986 | Down |
| Z69666.1 | 0.006 | 1.985 | Down |
| CCDC66 | 0.032 | 1.984 | Down |
| NRSN1 | 0.025 | 1.984 | Down |
| G087600 | 0.020 | 1.984 | Down |
| PEAK1 | 0.046 | 1.977 | Down |
| G047908 | 0.032 | 1.977 | Down |
| C3orf85 | 0.029 | 1.976 | Down |
| AC068768.1 | 0.049 | 1.976 | Down |
| AC005856.1 | 0.018 | 1.976 | Down |
| CATG00000061649.1 | 0.021 | 1.972 | Down |
| LINC02433 | 0.047 | 1.972 | Down |
| CATG00000077933.1 | 0.034 | 1.972 | Down |
| ADGRF1 | 0.039 | 1.970 | Down |
| TMEM161B-AS1 | 0.042 | 1.970 | Down |
| FLVCR2 | 0.006 | 1.968 | Down |
| G074144 | 0.001 | 1.965 | Down |
| KIAA0087 | 0.023 | 1.964 | Down |
| AL356753.1 | 0.018 | 1.963 | Down |
| AC239802.2 | 0.015 | 1.963 | Down |
| G065137 | 0.027 | 1.962 | Down |
| G086106 | 0.035 | 1.961 | Down |

| | | | |
|---|---|---|---|
| XLOC_003820 | 0.040 | 1.957 | Down |
| AP001781.1 | 0.048 | 1.957 | Down |
| AC090844.3 | 0.030 | 1.956 | Down |
| MMP24OS | 0.038 | 1.954 | Down |
| PHTF2 | 0.018 | 1.950 | Down |
| AC022915.2 | 0.002 | 1.949 | Down |
| LINC01959 | 0.016 | 1.947 | Down |
| PKN2-AS1 | 0.037 | 1.947 | Down |
| ZNF852 | 0.023 | 1.946 | Down |
| G061722 | 0.029 | 1.945 | Down |
| AC012174.1 | 0.024 | 1.944 | Down |
| HOXA9 | 0.030 | 1.944 | Down |
| AC138761.3 | 0.021 | 1.943 | Down |
| AC087761.1 | 0.012 | 1.943 | Down |
| PDP2 | 0.011 | 1.938 | Down |
| AC011498.2 | 0.018 | 1.937 | Down |
| SET | 0.046 | 1.937 | Down |
| AC104463.1 | 0.043 | 1.937 | Down |
| CATG00000013574.1 | 0.011 | 1.934 | Down |
| ARHGAP33 | 0.014 | 1.934 | Down |
| USP12-AS1 | 0.012 | 1.926 | Down |
| XLOC_007305 | 0.027 | 1.926 | Down |
| AC005692.3 | 0.040 | 1.924 | Down |
| AL391361.3 | 0.013 | 1.923 | Down |
| LINC02438 | 0.007 | 1.922 | Down |
| PAUPAR | 0.005 | 1.922 | Down |
| ENO1 | 0.016 | 1.921 | Down |
| ADGRV1 | 0.009 | 1.920 | Down |
| BRD3OS | 0.001 | 1.919 | Down |
| AL031598.1 | 0.009 | 1.918 | Down |
| ART3 | 0.048 | 1.917 | Down |
| LINC01511 | 0.028 | 1.914 | Down |
| CATG00000101288.1 | 0.000 | 1.913 | Down |
| ESRG | 0.013 | 1.911 | Down |
| AC116666.1 | 0.037 | 1.911 | Down |
| AC012467.2 | 0.024 | 1.910 | Down |
| CATG00000001057.1 | 0.019 | 1.910 | Down |
| KCNJ6-AS1 | 0.008 | 1.909 | Down |
| PNMA2 | 0.025 | 1.909 | Down |
| CATG00000017025.1 | 0.048 | 1.909 | Down |
| BVES-AS1 | 0.002 | 1.905 | Down |
| BX539320.1 | 0.049 | 1.904 | Down |
| AL034345.2 | 0.013 | 1.902 | Down |
| LINC01606 | 0.017 | 1.901 | Down |
| AL109628.2 | 0.025 | 1.900 | Down |
| CATG00000009958.1 | 0.042 | 1.898 | Down |
| CATG00000030400.1 | 0.018 | 1.898 | Down |
| AC138207.5 | 0.042 | 1.898 | Down |
| XLOC_010545 | 0.030 | 1.898 | Down |
| AC008734.2 | 0.039 | 1.897 | Down |
| G015350 | 0.036 | 1.895 | Down |
| AC115102.1 | 0.025 | 1.895 | Down |
| TESMIN | 0.011 | 1.894 | Down |
| AC116424.1 | 0.016 | 1.894 | Down |
| PVT1 | 0.009 | 1.891 | Down |
| AC010654.1 | 0.037 | 1.891 | Down |
| AL157895.1 | 0.035 | 1.889 | Down |
| AC025580.2 | 0.009 | 1.888 | Down |
| NFATC3 | 0.041 | 1.888 | Down |
| XLOC_000619 | 0.046 | 1.888 | Down |
| BACE1-AS | 0.020 | 1.884 | Down |
| G022744 | 0.023 | 1.880 | Down |
| AP001595.1 | 0.012 | 1.878 | Down |
| AP000924.1 | 0.016 | 1.877 | Down |
| AC027031.1 | 0.037 | 1.877 | Down |

| | | | |
|---|---|---|---|
| XLOC_011515 | 0.024 | 1.876 | Down |
| CATG00000005514.1 | 0.016 | 1.874 | Down |
| LINC01620 | 0.037 | 1.874 | Down |
| AC073284.1 | 0.016 | 1.873 | Down |
| G015366 | 0.009 | 1.873 | Down |
| LINC01114 | 0.024 | 1.870 | Down |
| CENPA | 0.016 | 1.870 | Down |
| TMEM9B-AS1 | 0.025 | 1.869 | Down |
| AP005482.2 | 0.016 | 1.867 | Down |
| G089016 | 0.001 | 1.867 | Down |
| AC093690.1 | 0.002 | 1.856 | Down |
| XLOC_010647 | 0.004 | 1.856 | Down |
| AC092471.1 | 0.025 | 1.854 | Down |
| WWTR1 | 0.043 | 1.853 | Down |
| LINC02245 | 0.010 | 1.852 | Down |
| AC092675.1 | 0.040 | 1.849 | Down |
| G065980 | 0.033 | 1.849 | Down |
| AC099524.1 | 0.016 | 1.844 | Down |
| AC093330.1 | 0.008 | 1.844 | Down |
| AL139158.3 | 0.032 | 1.843 | Down |
| AL133343.2 | 0.043 | 1.841 | Down |
| CR381653.1 | 0.013 | 1.837 | Down |
| AC002470.1 | 0.009 | 1.836 | Down |
| CATG00000013627.1 | 0.001 | 1.835 | Down |
| LINC00547 | 0.022 | 1.833 | Down |
| AL589826.2 | 0.008 | 1.833 | Down |
| AC096711.3 | 0.027 | 1.832 | Down |
| AC108865.2 | 0.030 | 1.830 | Down |
| JPX | 0.021 | 1.828 | Down |
| XLOC_006944 | 0.042 | 1.827 | Down |
| AC024619.2 | 0.014 | 1.826 | Down |
| AC022966.1 | 0.003 | 1.825 | Down |
| AC003102.1 | 0.011 | 1.824 | Down |
| NRXN1 | 0.013 | 1.823 | Down |
| AC010680.4 | 0.004 | 1.822 | Down |
| AKR1C8P | 0.048 | 1.820 | Down |
| MIAT | 0.039 | 1.820 | Down |
| CSF2RA | 0.008 | 1.817 | Down |
| CALU | 0.030 | 1.813 | Down |
| AC002383.1 | 0.040 | 1.812 | Down |
| AC103724.4 | 0.023 | 1.811 | Down |
| LINC00571 | 0.028 | 1.811 | Down |
| AL033384.1 | 0.043 | 1.810 | Down |
| MAGI1 | 0.013 | 1.809 | Down |
| PER4 | 0.017 | 1.808 | Down |
| SPATA3-AS1 | 0.035 | 1.808 | Down |
| XLOC_000569 | 0.023 | 1.808 | Down |
| G026452 | 0.035 | 1.807 | Down |
| LINC02232 | 0.042 | 1.806 | Down |
| AL157786.1 | 0.025 | 1.801 | Down |
| AL513327.1 | 0.020 | 1.801 | Down |
| AL136460.1 | 0.005 | 1.801 | Down |
| G011575 | 0.010 | 1.797 | Down |
| LINC00963 | 0.007 | 1.796 | Down |
| AC093157.1 | 0.031 | 1.795 | Down |
| CHD1L | 0.026 | 1.795 | Down |
| G087414 | 0.000 | 1.795 | Down |
| AP002856.1 | 0.038 | 1.791 | Down |
| SBF2-AS1 | 0.014 | 1.790 | Down |
| G020427 | 0.026 | 1.786 | Down |
| AJ003147.1 | 0.009 | 1.781 | Down |
| SIPA1L1 | 0.044 | 1.781 | Down |
| AL139317.1 | 0.036 | 1.780 | Down |
| SLC14A2-AS1 | 0.004 | 1.778 | Down |
| AC016831.6 | 0.001 | 1.777 | Down |

| | | | |
|---|---|---|---|
| FCRL3 | 0.002 | 1.775 | Down |
| LINC01378 | 0.004 | 1.775 | Down |
| SRD5A3-AS1 | 0.041 | 1.774 | Down |
| LMBR1 | 0.029 | 1.774 | Down |
| ZMIZ1-AS1 | 0.027 | 1.773 | Down |
| G044057 | 0.001 | 1.772 | Down |
| AC020891.1 | 0.002 | 1.772 | Down |
| AC002064.2 | 0.049 | 1.771 | Down |
| CATG00000051486.1 | 0.020 | 1.770 | Down |
| LINC00707 | 0.045 | 1.767 | Down |
| AL035706.1 | 0.030 | 1.767 | Down |
| VLDLR-AS1 | 0.042 | 1.765 | Down |
| CATG00000011583.1 | 0.004 | 1.764 | Down |
| AC012640.2 | 0.016 | 1.763 | Down |
| AC109466.1 | 0.045 | 1.762 | Down |
| LOC105376306 | 0.015 | 1.762 | Down |
| DPH3 | 0.012 | 1.759 | Down |
| AC104012.1 | 0.006 | 1.757 | Down |
| AC016822.1 | 0.013 | 1.755 | Down |
| AL158829.1 | 0.047 | 1.753 | Down |
| G035480 | 0.003 | 1.753 | Down |
| AL132712.2 | 0.015 | 1.752 | Down |
| AC072062.1 | 0.034 | 1.749 | Down |
| G043065 | 0.022 | 1.748 | Down |
| CATG00000030927.1 | 0.030 | 1.746 | Down |
| AL360294.1 | 0.010 | 1.744 | Down |
| SNX19 | 0.001 | 1.744 | Down |
| AC137810.1 | 0.032 | 1.743 | Down |
| AC023442.2 | 0.040 | 1.741 | Down |
| OSTM1 | 0.022 | 1.739 | Down |
| RBMS3-AS2 | 0.022 | 1.733 | Down |
| AC104574.2 | 0.034 | 1.732 | Down |
| MMP25-AS1 | 0.006 | 1.730 | Down |
| XLOC_006094 | 0.020 | 1.729 | Down |
| AP002812.3 | 0.022 | 1.728 | Down |
| AC004466.1 | 0.030 | 1.728 | Down |
| SHQ1 | 0.032 | 1.727 | Down |
| ACTA2-AS1 | 0.006 | 1.726 | Down |
| CACNA1G-AS1 | 0.026 | 1.725 | Down |
| G080931 | 0.008 | 1.724 | Down |
| TTTY1 | 0.018 | 1.723 | Down |
| CINP | 0.014 | 1.723 | Down |
| CATG00000096657.1 | 0.025 | 1.723 | Down |
| DLGAP1-AS3 | 0.022 | 1.722 | Down |
| TCP10L2 | 0.040 | 1.721 | Down |
| RHOH | 0.009 | 1.718 | Down |
| SYNRG | 0.004 | 1.717 | Down |
| AC116667.2 | 0.036 | 1.715 | Down |
| G018581 | 0.015 | 1.715 | Down |
| G072579 | 0.018 | 1.713 | Down |
| BIG-lncRNA-464 | 0.014 | 1.713 | Down |
| LOC101409256 | 0.022 | 1.713 | Down |
| AL358393.1 | 0.042 | 1.712 | Down |
| CATG00000058838.1 | 0.043 | 1.710 | Down |
| AC018866.1 | 0.043 | 1.709 | Down |
| CEP41 | 0.034 | 1.708 | Down |
| AC107464.3 | 0.013 | 1.708 | Down |
| LINC00598 | 0.032 | 1.707 | Down |
| Z95114.2 | 0.041 | 1.703 | Down |
| G045213 | 0.017 | 1.702 | Down |
| BIG-lncRNA-454 | 0.044 | 1.700 | Down |
| AL035696.1 | 0.009 | 1.696 | Down |
| AC069540.2 | 0.008 | 1.695 | Down |
| AC002454.1 | 0.021 | 1.695 | Down |
| CATG00000031078.1 | 0.008 | 1.694 | Down |

| | | | |
|---|---|---|---|
| XLOC_000609 | 0.023 | 1.693 | Down |
| AP000281.2 | 0.024 | 1.693 | Down |
| AC015909.1 | 0.028 | 1.691 | Down |
| CATG00000100994.1 | 0.015 | 1.691 | Down |
| KIRREL3 | 0.023 | 1.690 | Down |
| G073008 | 0.001 | 1.689 | Down |
| CATG00000103390.1 | 0.037 | 1.689 | Down |
| AL355512.1 | 0.033 | 1.689 | Down |
| AC090510.1 | 0.006 | 1.688 | Down |
| FCAMR | 0.011 | 1.687 | Down |
| FAM157C | 0.016 | 1.686 | Down |
| G079867 | 0.047 | 1.685 | Down |
| AC002451.1 | 0.030 | 1.685 | Down |
| TAB2-AS1 | 0.028 | 1.682 | Down |
| KIF25-AS1 | 0.039 | 1.682 | Down |
| LINC00607 | 0.022 | 1.681 | Down |
| HCG20 | 0.041 | 1.680 | Down |
| G072542 | 0.039 | 1.678 | Down |
| CATG00000013944.1 | 0.044 | 1.677 | Down |
| CATG00000040290.1 | 0.024 | 1.675 | Down |
| AC090673.1 | 0.028 | 1.674 | Down |
| XLOC_005521 | 0.025 | 1.674 | Down |
| G011248 | 0.018 | 1.672 | Down |
| AL022068.1 | 0.007 | 1.672 | Down |
| CATG00000049227.1 | 0.045 | 1.671 | Down |
| AC027307.2 | 0.016 | 1.668 | Down |
| LINC02588 | 0.044 | 1.666 | Down |
| AC136618.1 | 0.034 | 1.665 | Down |
| XLOC_007595 | 0.040 | 1.663 | Down |
| SH2B1 | 0.020 | 1.663 | Down |
| OSBPL10-AS1 | 0.049 | 1.663 | Down |
| DOCK6 | 0.044 | 1.663 | Down |
| AL133227.1 | 0.014 | 1.662 | Down |
| AL031656.1 | 0.010 | 1.662 | Down |
| FLJ45513 | 0.037 | 1.661 | Down |
| VEZT | 0.047 | 1.660 | Down |
| AC007220.1 | 0.012 | 1.658 | Down |
| AL138831.1 | 0.009 | 1.658 | Down |
| LINC02319 | 0.035 | 1.656 | Down |
| G016237 | 0.009 | 1.656 | Down |
| XLOC_010677 | 0.003 | 1.654 | Down |
| XLOC_005944 | 0.029 | 1.653 | Down |
| G060478 | 0.004 | 1.652 | Down |
| IBA57-DT | 0.038 | 1.651 | Down |
| LINC02120 | 0.025 | 1.650 | Down |
| AC090531.1 | 0.031 | 1.649 | Down |
| AC093772.1 | 0.014 | 1.649 | Down |
| BDNF | 0.029 | 1.646 | Down |
| RAP2C-AS1 | 0.041 | 1.646 | Down |
| AC017116.1 | 0.006 | 1.643 | Down |
| MGAM | 0.014 | 1.642 | Down |
| CATG00000106063.1 | 0.014 | 1.641 | Down |
| AL732314.6 | 0.038 | 1.641 | Down |
| AC073911.1 | 0.016 | 1.636 | Down |
| ELOVL2-AS1 | 0.050 | 1.635 | Down |
| AC008406.1 | 0.041 | 1.635 | Down |
| MTBP | 0.008 | 1.635 | Down |
| AC012511.1 | 0.038 | 1.634 | Down |
| CATG00000052264.1 | 0.029 | 1.627 | Down |
| AC092127.1 | 0.026 | 1.627 | Down |
| TYMS | 0.016 | 1.626 | Down |
| AC019133.1 | 0.021 | 1.626 | Down |
| CBR1 | 0.010 | 1.626 | Down |
| LINC00222 | 0.048 | 1.625 | Down |
| HUNK-AS1 | 0.038 | 1.624 | Down |

| | | | |
|---|---|---|---|
| RAD51D | 0.000 | 1.624 | Down |
| L34079.3 | 0.034 | 1.623 | Down |
| CKMT2-AS1 | 0.036 | 1.619 | Down |
| G007125 | 0.039 | 1.619 | Down |
| AC034114.2 | 0.019 | 1.618 | Down |
| AC127024.4 | 0.006 | 1.614 | Down |
| TMEM237 | 0.022 | 1.614 | Down |
| AL161725.1 | 0.001 | 1.610 | Down |
| AL139286.2 | 0.014 | 1.610 | Down |
| XLOC_006904 | 0.003 | 1.610 | Down |
| XLOC_007374 | 0.007 | 1.607 | Down |
| NEGR1-IT1 | 0.039 | 1.604 | Down |
| LINC01018 | 0.013 | 1.603 | Down |
| AC090607.4 | 0.007 | 1.603 | Down |
| AL391883.1 | 0.044 | 1.603 | Down |
| INSYN1-AS1 | 0.008 | 1.602 | Down |
| CHN1 | 0.039 | 1.601 | Down |
| LCN8 | 0.044 | 1.599 | Down |
| LINC01725 | 0.040 | 1.592 | Down |
| NALCN-AS1 | 0.004 | 1.591 | Down |
| AC092111.2 | 0.017 | 1.589 | Down |
| LINC00667 | 0.025 | 1.587 | Down |
| AC015969.1 | 0.007 | 1.587 | Down |
| AL136301.1 | 0.027 | 1.585 | Down |
| XLOC_001990 | 0.011 | 1.583 | Down |
| AL160313.1 | 0.012 | 1.583 | Down |
| AC016866.1 | 0.026 | 1.581 | Down |
| AC027682.3 | 0.033 | 1.580 | Down |
| CCDC144NL-AS1 | 0.005 | 1.579 | Down |
| NIFK-AS1 | 0.024 | 1.578 | Down |
| AC106895.2 | 0.019 | 1.575 | Down |
| AL691403.1 | 0.040 | 1.574 | Down |
| AC105233.5 | 0.044 | 1.574 | Down |
| CATG00000075725.1 | 0.014 | 1.573 | Down |
| AL121906.2 | 0.039 | 1.573 | Down |
| PRKCZ-AS1 | 0.050 | 1.571 | Down |
| USP12-AS2 | 0.018 | 1.570 | Down |
| CATG00000072894.1 | 0.049 | 1.569 | Down |
| TBX22 | 0.002 | 1.566 | Down |
| PRIMPOL | 0.042 | 1.566 | Down |
| KLHL22 | 0.045 | 1.566 | Down |
| AC008592.1 | 0.021 | 1.565 | Down |
| LINC00461 | 0.016 | 1.564 | Down |
| SP2-AS1 | 0.013 | 1.562 | Down |
| XLOC_003967 | 0.031 | 1.562 | Down |
| SNHG14 | 0.017 | 1.561 | Down |
| AC090912.1 | 0.020 | 1.561 | Down |
| AC018685.1 | 0.013 | 1.558 | Down |
| CATG00000040000.1 | 0.018 | 1.556 | Down |
| TTLL7 | 0.015 | 1.555 | Down |
| CATG00000114545.1 | 0.011 | 1.555 | Down |
| G080668 | 0.047 | 1.555 | Down |
| SNHG12 | 0.030 | 1.553 | Down |
| DPH6-DT | 0.011 | 1.551 | Down |
| AC245164.1 | 0.030 | 1.550 | Down |
| AC079910.1 | 0.049 | 1.547 | Down |
| AK128153 | 0.024 | 1.547 | Down |
| AL138976.2 | 0.040 | 1.544 | Down |
| G031235 | 0.006 | 1.544 | Down |
| HRAT5 | 0.015 | 1.544 | Down |
| AC073342.2 | 0.028 | 1.543 | Down |
| AL031386.1 | 0.016 | 1.542 | Down |
| CATG00000103478.1 | 0.032 | 1.542 | Down |
| AC108062.1 | 0.005 | 1.541 | Down |
| AL513283.1 | 0.014 | 1.541 | Down |

| | | | |
|---|---|---|---|
| AL359313.1 | 0.022 | 1.539 | Down |
| LINC00923 | 0.044 | 1.538 | Down |
| BIG-lncRNA-81 | 0.001 | 1.537 | Down |
| AC010331.1 | 0.041 | 1.537 | Down |
| LOC101929431 | 0.000 | 1.535 | Down |
| AC087501.3 | 0.036 | 1.532 | Down |
| LINC02537 | 0.012 | 1.530 | Down |
| LINC01874 | 0.004 | 1.529 | Down |
| G077775 | 0.032 | 1.528 | Down |
| G017283 | 0.038 | 1.527 | Down |
| AC100843.1 | 0.029 | 1.526 | Down |
| LINC01560 | 0.022 | 1.525 | Down |
| AP000233.2 | 0.032 | 1.524 | Down |
| APP | 0.037 | 1.524 | Down |
| LINC01633 | 0.019 | 1.523 | Down |
| AL049637.1 | 0.049 | 1.523 | Down |
| LINC02362 | 0.015 | 1.522 | Down |
| XLOC_010714 | 0.046 | 1.521 | Down |
| XLOC_010783 | 0.046 | 1.517 | Down |
| LINC00173 | 0.042 | 1.516 | Down |
| AC004156.1 | 0.048 | 1.516 | Down |
| G008066 | 0.027 | 1.515 | Down |
| CATG00000059739.1 | 0.034 | 1.512 | Down |
| BX005266.2 | 0.036 | 1.511 | Down |
| AC126696.3 | 0.003 | 1.511 | Down |
| XLOC_006384 | 0.029 | 1.509 | Down |
| G063729 | 0.038 | 1.505 | Down |
| AL109954.1 | 0.003 | 1.505 | Down |
| PCAT6 | 0.030 | 1.503 | Down |
| AC138123.2 | 0.041 | 1.502 | Down |
| AC021321.1 | 0.005 | 1.501 | Down |
| AL356599.1 | 0.031 | 1.500 | Down |
| CATG00000051921.1 | 0.029 | 1.499 | Down |
| XLOC_002094 | 0.001 | 1.499 | Down |
| AL160191.1 | 0.036 | 1.498 | Down |
| G017203 | 0.046 | 1.496 | Down |
| AP001372.3 | 0.035 | 1.495 | Down |
| BIG-lncRNA-1074 | 0.009 | 1.495 | Down |
| G028667 | 0.022 | 1.495 | Down |
| SNHG17 | 0.017 | 1.494 | Down |
| LINC00652 | 0.010 | 1.493 | Down |
| RP1L1 | 0.049 | 1.491 | Down |
| XLOC_011701 | 0.031 | 1.490 | Down |
| CATG00000017432.1 | 0.043 | 1.489 | Down |
| AC111194.2 | 0.019 | 1.488 | Down |
| LRRC29 | 0.032 | 1.488 | Down |
| AC005256.1 | 0.006 | 1.488 | Down |
| DEFB124 | 0.034 | 1.486 | Down |
| SPATA9 | 0.046 | 1.484 | Down |
| G011384 | 0.048 | 1.481 | Down |
| G056235 | 0.015 | 1.481 | Down |
| G000411 | 0.049 | 1.480 | Down |
| CATG00000035108.1 | 0.009 | 1.478 | Down |
| AL590432.1 | 0.041 | 1.476 | Down |
| LINC02531 | 0.019 | 1.475 | Down |
| XLOC_002707 | 0.001 | 1.474 | Down |
| AC138024.1 | 0.028 | 1.473 | Down |
| AP000426.1 | 0.042 | 1.472 | Down |
| SLC22A23 | 0.009 | 1.470 | Down |
| FLG-AS1 | 0.038 | 1.469 | Down |
| STEAP1B | 0.021 | 1.469 | Down |
| LINC01482 | 0.036 | 1.467 | Down |
| CATG00000015362.1 | 0.030 | 1.466 | Down |
| AC015818.9 | 0.003 | 1.466 | Down |
| GSE61474_XLOC_003354 | 0.037 | 1.466 | Down |

| | | | |
|---|---|---|---|
| G071435 | 0.023 | 1.465 | Down |
| FKBP9P1 | 0.023 | 1.463 | Down |
| CATG00000042245.1 | 0.032 | 1.462 | Down |
| AC078880.3 | 0.006 | 1.462 | Down |
| LINC02197 | 0.030 | 1.461 | Down |
| XLOC_004495 | 0.002 | 1.459 | Down |
| AL355493.2 | 0.006 | 1.454 | Down |
| LINC02407 | 0.006 | 1.454 | Down |
| SLC2A9 | 0.047 | 1.454 | Down |
| ALPK1 | 0.036 | 1.453 | Down |
| AC008105.1 | 0.038 | 1.452 | Down |
| AC104237.1 | 0.033 | 1.451 | Down |
| CATG00000029348.1 | 0.044 | 1.451 | Down |
| AL645465.1 | 0.037 | 1.448 | Down |
| GIGYF2 | 0.045 | 1.446 | Down |
| SOX9-AS1 | 0.031 | 1.446 | Down |
| CATG00000046856.1 | 0.025 | 1.445 | Down |
| CCDC92 | 0.047 | 1.443 | Down |
| AC025470.2 | 0.032 | 1.438 | Down |
| AC233976.1 | 0.040 | 1.438 | Down |
| AL591885.1 | 0.023 | 1.436 | Down |
| LINC02348 | 0.020 | 1.435 | Down |
| AP001825.1 | 0.030 | 1.434 | Down |
| AL354732.1 | 0.005 | 1.434 | Down |
| ZNF213-AS1 | 0.035 | 1.432 | Down |
| AP002433.2 | 0.030 | 1.431 | Down |
| AL078623.1 | 0.014 | 1.428 | Down |
| AC006213.1 | 0.014 | 1.422 | Down |
| STAC | 0.040 | 1.422 | Down |
| AC004690.2 | 0.013 | 1.420 | Down |
| PRKCE | 0.044 | 1.419 | Down |
| CATG00000054726.1 | 0.012 | 1.418 | Down |
| AL138828.1 | 0.046 | 1.417 | Down |
| AC105206.1 | 0.014 | 1.416 | Down |
| G064452 | 0.008 | 1.413 | Down |
| AL354702.2 | 0.024 | 1.413 | Down |
| AC067930.3 | 0.022 | 1.412 | Down |
| AC125421.1 | 0.039 | 1.411 | Down |
| CATG00000048945.1 | 0.040 | 1.410 | Down |
| AC015660.2 | 0.037 | 1.410 | Down |
| RP11-144G6.12 | 0.014 | 1.409 | Down |
| AC131025.2 | 0.002 | 1.407 | Down |
| AC136285.3 | 0.043 | 1.406 | Down |
| AL391361.2 | 0.012 | 1.404 | Down |
| XLOC_012666 | 0.022 | 1.403 | Down |
| XLOC_009920 | 0.050 | 1.401 | Down |
| LINC01775 | 0.039 | 1.398 | Down |
| LINC00687 | 0.020 | 1.397 | Down |
| XLOC_005578 | 0.004 | 1.396 | Down |
| AC090559.1 | 0.040 | 1.395 | Down |
| AC025154.1 | 0.046 | 1.393 | Down |
| BIG-lncRNA-907 | 0.020 | 1.392 | Down |
| CATG00000106168.1 | 0.042 | 1.388 | Down |
| G012838 | 0.040 | 1.384 | Down |
| C2orf92 | 0.021 | 1.384 | Down |
| AL139287.1 | 0.047 | 1.383 | Down |
| LINC01535 | 0.031 | 1.381 | Down |
| AC009063.2 | 0.030 | 1.380 | Down |
| G036546 | 0.026 | 1.379 | Down |
| G008785 | 0.000 | 1.378 | Down |
| CATG00000067103.1 | 0.028 | 1.374 | Down |
| NAT1 | 0.047 | 1.367 | Down |
| AC113192.5 | 0.001 | 1.366 | Down |
| AC093802.2 | 0.043 | 1.362 | Down |
| AL021396.1 | 0.015 | 1.362 | Down |

| | | | |
|---|---|---|---|
| CSNK1D | 0.007 | 1.362 | Down |
| AK023948 | 0.038 | 1.360 | Down |
| TMEM254-AS1 | 0.049 | 1.360 | Down |
| AL603839.2 | 0.041 | 1.358 | Down |
| AL139241.1 | 0.039 | 1.355 | Down |
| G055549 | 0.003 | 1.353 | Down |
| XLOC_003880 | 0.026 | 1.349 | Down |
| AC004771.5 | 0.041 | 1.348 | Down |
| ADD3-AS1 | 0.009 | 1.347 | Down |
| XLOC_009831 | 0.044 | 1.347 | Down |
| LINC02272 | 0.000 | 1.347 | Down |
| G006749 | 0.017 | 1.343 | Down |
| AC156455.1 | 0.031 | 1.342 | Down |
| INO80-AS1 | 0.030 | 1.341 | Down |
| G056200 | 0.017 | 1.339 | Down |
| CATG00000009721.1 | 0.005 | 1.338 | Down |
| LINC02193 | 0.019 | 1.338 | Down |
| ZFC3H1 | 0.042 | 1.337 | Down |
| AC010401.1 | 0.015 | 1.329 | Down |
| MEIS1-AS2 | 0.001 | 1.329 | Down |
| CACFD1 | 0.000 | 1.328 | Down |
| G009751 | 0.044 | 1.328 | Down |
| AP001189.6 | 0.041 | 1.327 | Down |
| CATG00000106576.1 | 0.028 | 1.327 | Down |
| AC018521.1 | 0.046 | 1.324 | Down |
| SUGT1P1 | 0.001 | 1.324 | Down |
| GRM7-AS3 | 0.002 | 1.322 | Down |
| REXO4 | 0.045 | 1.322 | Down |
| CATG00000018535.1 | 0.004 | 1.322 | Down |
| HNRNPUL2-BSCL2 | 0.017 | 1.320 | Down |
| AL355075.2 | 0.005 | 1.318 | Down |
| G024540 | 0.032 | 1.318 | Down |
| AKR1C2 | 0.003 | 1.318 | Down |
| AL022238.2 | 0.045 | 1.316 | Down |
| AL157413.1 | 0.002 | 1.316 | Down |
| MRPS27 | 0.035 | 1.316 | Down |
| AC020978.5 | 0.049 | 1.307 | Down |
| AC009163.1 | 0.034 | 1.307 | Down |
| XLOC_010146 | 0.014 | 1.306 | Down |
| XLOC_001460 | 0.047 | 1.306 | Down |
| G074709 | 0.027 | 1.301 | Down |
| TMEM120B | 0.001 | 1.301 | Down |
| AC027031.2 | 0.002 | 1.299 | Down |
| MGLL | 0.024 | 1.299 | Down |
| AC078852.1 | 0.002 | 1.295 | Down |
| AL022238.4 | 0.039 | 1.295 | Down |
| G037088 | 0.021 | 1.294 | Down |
| LINC02471 | 0.010 | 1.292 | Down |
| AC009411.1 | 0.045 | 1.291 | Down |
| SNX17 | 0.003 | 1.290 | Down |
| XLOC_006156 | 0.019 | 1.289 | Down |
| AL583808.1 | 0.020 | 1.288 | Down |
| LINC00359 | 0.024 | 1.287 | Down |
| AL160286.2 | 0.019 | 1.286 | Down |
| G054239 | 0.002 | 1.286 | Down |
| AC115676.1 | 0.004 | 1.284 | Down |
| ABCA13 | 0.043 | 1.277 | Down |
| G036192 | 0.042 | 1.273 | Down |
| AC004908.3 | 0.005 | 1.272 | Down |
| LCMT1-AS2 | 0.027 | 1.265 | Down |
| TUSC8 | 0.001 | 1.265 | Down |
| CATG00000076041.1 | 0.008 | 1.264 | Down |
| PRKAG2-AS1 | 0.035 | 1.264 | Down |
| G029713 | 0.025 | 1.264 | Down |
| AL645939.5 | 0.020 | 1.263 | Down |

| | | | |
|---|---|---|---|
| DKFZP434K028 | 0.045 | 1.262 | Down |
| AC116456.1 | 0.026 | 1.261 | Down |
| GPR17 | 0.011 | 1.256 | Down |
| Z83844.2 | 0.037 | 1.255 | Down |
| XLOC_008610 | 0.022 | 1.248 | Down |
| MIR9-3HG | 0.018 | 1.245 | Down |
| AC004466.2 | 0.023 | 1.245 | Down |
| LINC01122 | 0.034 | 1.244 | Down |
| LINC01711 | 0.024 | 1.240 | Down |
| LINC02443 | 0.007 | 1.237 | Down |
| LINC01070 | 0.030 | 1.234 | Down |
| EXTL3-AS1 | 0.037 | 1.231 | Down |
| SPTA1 | 0.013 | 1.229 | Down |
| AC025871.1 | 0.019 | 1.223 | Down |
| CATG00000058740.1 | 0.014 | 1.216 | Down |
| AP001604.1 | 0.010 | 1.214 | Down |
| XLOC_013696 | 0.022 | 1.207 | Down |
| CATG00000069831.1 | 0.021 | 1.202 | Down |
| AC091167.5 | 0.006 | 1.201 | Down |

**Supplementary Table 14. KEGG analysis of the anesthesia/surgery (> 3 hours) deregulated mRNAs in serum from pediatric patients < 4 years old**

| Enriched Pathway | Genes | P value |
|---|---|---|
| *Pathways of Neurodegeneration* | APC2, FRAT2, NDUFA10, NDUFA11, NDUFA7, NDUFB1, TANK, CALML5, EIF2S1, FZD5, GRIN2A, MAP1LC3C, MT-CYB, MT-CYB, MT-CO3, MAP2K2, PSMA2, PSMD14 RYR2, UQCRQ, UQCR11, and UBEG | 0.650 |
| *Neutrophil Extracellular Trap formation* | H2AC25, H2AC6, H2AZ1, H3C11, H4C1, HDAC4, MAP2K2, MPO, PRKCA, H3-5, H3-4 | 1.000 |
| *Necroptosis* | H2AC25, H2AC6, H2AZ1, PGAM5, CHMP3, CHMP4C, CHMP7, FTH1, FTL, STAT3, STAT6 | 0.650 |
| *Ferroptosis* | FTH1, FTL, MAP1LC3C, SLC3A2 and SLC39A8 | 0.780 |

KEGG pathway associated with 1492 mRNA transcripts that met thresholds of (±1.2-fold difference; $p < 0.05$) between the post and pre-surgery sample in the pediatric patients.

**Supplementary Table 15. Dysregulated synaptic mRNAs in serum from anesthetic exposed pediatric patients < 4 years old and their related cellular components and biological process**

| Gene ID | Gene name | GO Term Name | GO Domain |
|---|---|---|---|
| ABHD17C | abhydrolase domain containing 17C, depalmitoylase | regulation of postsynapse organization (GO:0099175) | BP |
| ACTN1 | actinin alpha 1 | structural constituent of postsynapse (GO:0099186) | BP |
| ADORA2A | adenosine A2a receptor | chemical synaptic transmission (GO:0099171) | BP |
| ANO6 | anoctamin 6 | regulation of postsynaptic membrane potential (GO:0060078) | BP |
| ARHGEF15 | Rho guanine nucleotide exchange factor 15 | regulation of postsynapse assembly (GO:0150052) | BP |
| ATG7 | autophagy related 7 | synaptic vesicle cycle (GO:0099504) | BP |
| ATP6V0E2 | ATPase H+ transporting V0 subunit e2 | synaptic vesicle proton loading (GO:0097401) | BP |
| ATP6V1B2 | ATPase H+ transporting V1 subunit B2 | synaptic vesicle proton loading (GO:0097401) | BP |
| BCAN | brevican | synapse maturation (GO:0060074) | BP |
| BSN | bassoon presynaptic cytomatrix protein | zone (GO:0098882), regulation of synaptic vesicle cycle (GO:0098693), synaptic vesicle clustering (GO:0097091) | BP |
| CACNB2 | calcium voltage-gated channel auxiliary subunit beta 2 | voltage-gated calcium channel activity involved in regulation of presynaptic cytosolic calcium levels (GO:0099626) | BP |
| CALB2 | calbindin 2 | calcium ion binding involved in regulation of presynaptic cytosolic calcium levels (GO:0099534) | BP |
| CAP2 | cyclase associated actin cytoskeleton regulatory protein 2 | presynaptic actin cytoskeleton organization (GO:0099140), regulation of modification of postsynaptic actin cytoskeleton (GO:1905274) | BP |
| CASP3 | caspase 3 | regulation of synaptic vesicle cycle (GO:0098693) | BP |
| CASR | calcium sensing receptor | regulation of presynaptic membrane potential (GO:0099505) | BP |
| CBLB | Cbl proto-oncogene B | regulation protein catabolic process at postsynapse (GO:0140252), regulation of postsynaptic neurotransmitter receptor endocytosis (GO:0099149) | BP |
| CBLN2 | cerebellin 2 precursor | trans-synaptic signaling, modulating synaptic transmission (GO:0099550) | BP |

| | | | |
|---|---|---|---|
| CHRM1 | cholinergic receptor muscarinic 1 | membrane potential (GO:0060078), postsynaptic modulation of chemical synaptic transmission (GO:0099170) | BP |
| CNR2 | cannabinoid receptor 2 | trans-synaptic signaling by endocannabinoid, modulating synaptic transmission (GO:0099553) | BP |
| CNTNAP2 | contactin associated protein 2 | regulation of postsynaptic membrane neurotransmitter receptor levels (GO:0099072) | BP |
| CTBP1 | C-terminal binding protein 1 | presynapse to nucleus signaling pathway (GO:0099526), synaptic vesicle endocytosis (GO:0048488), synaptic vesicle clustering (GO:0097091) | BP |
| CTSD | cathepsin D | synaptic vesicle endosomal processing (GO:0099532) | BP |
| DIXDC1 | DIX domain containing 1 | modification of postsynaptic actin cytoskeleton (GO:0098885) | BP |
| DRD1 | dopamine receptor D1 | modification of postsynaptic structure (GO:0099010), presynaptic modulation of chemical synaptic transmission (GO:0099171), postsynaptic modulation of chemical synaptic | BP |
| EEF2K | eukaryotic elongation factor 2 kinase | regulation of translation at postsynapse (GO:0140245) | BP |
| EIF4E | eukaryotic translation initiation factor 4E | regulation of translation at postsynapse, modulating synaptic transmission (GO:0099578) | BP |
| ELFN1 | extracellular leucine rich repeat and fibronectin type III domain containing 1 | synapse adhesion between pre- and post-synapse (GO:0099560) | BP |
| ERBIN | erbb2 interacting protein | regulation of postsynaptic membrane neurotransmitter receptor levels (GO:0099072) | BP |
| FZD5 | frizzled class receptor 5 | presynapse assembly (GO:0099054) | BP |
| GABBR1 | gamma-aminobutyric acid type B receptor subunit 1 | G protein-coupled neurotransmitter receptor activity involved in regulation of presynaptic membrane potential (GO:0150047), G protein-coupled neurotransmitter receptor activity involved in regulation of postsynaptic membrane potential | BP |
| GABRB2 | gamma-aminobutyric acid type A receptor subunit beta2 | ligand-gated ion channel activity involved in regulation of presynaptic membrane potential (GO:0099507), transmitter-gated ion channel activity involved in regulation of postsynaptic membrane potential (GO:1904315) | BP |

| | | | |
|---|---|---|---|
| GAP43 | growth associated protein 43 | regulation of postsynaptic specialization assembly (GO:0099150) | BP |
| GNAI1 | G protein subunit alpha i1 | neurotransmitter receptor localization to postsynaptic specialization membrane (GO:0099645) | BP |
| GRIN2A | glutamate ionotropic receptor NMDA type subunit 2A | ligand-gated ion channel activity involved in regulation of presynaptic membrane potential (GO:0099507) | BP |
| GRIN2A | glutamate ionotropic receptor NMDA type subunit 2A | transmitter-gated ion channel activity involved in regulation of postsynaptic membrane potential (GO:1904315) | BP |
| HNRNPK | heterogeneous nuclear ribonucleoprotein K | modulation of chemical synaptic transmission (GO:0050804), regulation of postsynapse organization (GO:0099175) | BP |
| HNRNPM | heterogeneous nuclear ribonucleoprotein M | regulation of postsynapse organization (GO:0099175) | BP |
| HTR7 | 5-hydroxytryptamine receptor 7 | presynaptic modulation of chemical synaptic transmission (GO:0099171) | BP |
| IGF1 | insulin like growth factor 1 | postsynaptic modulation of chemical synaptic transmission (GO:0099170) | BP |
| KCNA4 | potassium voltage-gated channel subfamily A member 4 | integral component of presynaptic membrane (GO:0099056), voltage-gated ion channel activity involved in regulation of presynaptic membrane potential (GO:0099508) | CC |
| KCNJ8 | potassium inwardly rectifying channel subfamily J member 8 | voltage-gated ion channel activity involved in regulation of presynaptic membrane potential (GO:0099508) | BP |
| KCNJ9 | potassium inwardly rectifying channel subfamily J member 9 | regulation of presynaptic membrane potential (GO:0099505) | BP |
| KCTD8 | potassium channel tetramerization domain containing 8 | G protein-coupled neurotransmitter receptor activity involved in regulation of presynaptic membrane potential (GO:0150047) | BP |
| KIF21B | kinesin family member 21B | retrograde dendritic transport (GO:0098934), regulation of postsynaptic membrane neurotransmitter receptor levels (GO:0099072) | BP |
| KPNA1 | karyopherin subunit alpha 1 | postsynapse to nucleus signaling pathway (GO:0099527) | BP |
| MARCKSL1 | MARCKS like 1 | regulation of presynaptic cytosolic calcium levels (GO:0099509) | BP |

| | | | |
|---|---|---|---|
| NAPB | NSF attachment protein beta | regulation of synaptic vesicle priming (GO:0010807) | BP |
| NEO1 | neogenin 1 | trans-synaptic signaling, modulating synaptic transmission (GO:0099550) | BP |
| NGDN | neuroguidin | postsynapse (GO:0140245), postsynaptic modulation of chemical synaptic transmission (GO:0099170) | BP |
| NGEF | neuronal guanine nucleotide exchange factor | regulation of synapse disassembly (GO:1905806) | BP |
| PLAT | plasminogen activator, tissue type | trans-synaptic signaling by BDNF, modulating synaptic transmission (GO:0099183) | BP |
| PRKCA | protein kinase C alpha | regulation of synaptic vesicle exocytosis (GO:2000300) | BP |
| PRKCI | protein kinase C iota | regulation of postsynaptic membrane neurotransmitter receptor levels (GO:0099072) | BP |
| PRMT8 | protein arginine methyltransferase 8 | regulation of modification of postsynaptic actin cytoskeleton (GO:1905274) | BP |
| PSEN2 | presenilin 2 | protein catabolic process at postsynapse (GO:0140249), regulation of postsynapse organization (GO:0099175) | BP |
| PUM2 | pumilio RNA binding family member 2 | regulation of postsynapse assembly (GO:0150052) | BP |
| RBMX | RNA binding motif protein X-linked | regulation of postsynapse organization (GO:0099175) | BP |
| RDX | radixin | regulation of postsynaptic neurotransmitter receptor diffusion trapping (GO:0150054) | BP |
| RELN | reelin | synaptic vesicle exocytosis (GO:0016079), regulation of synapse maturation (GO:0090128), integral component of postsynaptic membrane (GO:0099055) | BP |
| RHOG | ras homolog family member G | regulation of postsynapse assembly (GO:0150052) | BP |
| RNF216 | ring finger protein 216 | protein catabolic process, modulating synaptic transmission (GO:0099546), regulation of postsynaptic neurotransmitter receptor endocytosis | BP |
| ROBO2 | roundabout guidance receptor 2 | synapse maturation (GO:0060074) | BP |

| | | | |
|---|---|---|---|
| ROR2 | receptor tyrosine kinase like orphan receptor 2 | regulation of synaptic signaling by nitric oxide (GO:0150045), modulation of chemical synaptic transmission (GO:0050804), regulation of postsynapse organization (GO:0099175) | BP |
| RPL17 | ribosomal protein L17 | translation at presynapse (GO:0140236), translation at postsynapse (GO:0140242) | BP |
| RPL22 | ribosomal protein L22 | translation at presynapse (GO:0140236), translation at postsynapse (GO:0140242) | BP |
| RPL26 | ribosomal protein L26 | translation at presynapse (GO:0140236), translation at postsynapse (GO:0140242) | BP |
| RPL29 | ribosomal protein L29 | translation at presynapse (GO:0140236), translation at postsynapse (GO:0140242) | BP |
| RPL37A | ribosomal protein L37a | translation at presynapse (GO:0140236), translation at postsynapse (GO:0140242) | BP |
| RPS24 | ribosomal protein S24 | translation at presynapse (GO:0140236), translation at postsynapse (GO:0140242) | BP |
| RPS28 | ribosomal protein S28 | translation at presynapse (GO:0140236), translation at postsynapse (GO:0140242) | BP |
| SCN2A | sodium voltage-gated channel alpha subunit 2 | voltage-gated ion channel activity involved in regulation of presynaptic membrane potential (GO:0099508) | BP |
| SDCBP | syndecan binding protein | presynapse assembly (GO:0099054) | BP |
| SHANK3 | SH3 and multiple ankyrin repeat domains 3 | structural constituent of postsynaptic density (GO:0098919), regulation of postsynapse organization (GO:0099175), modulation of chemical synaptic transmission | BP |
| SLC6A17 | solute carrier family 6 member 17 | metabolism | BP |
| SLIT2 | slit guidance ligand 2 | synapse maturation (GO:0060074) | BP |
| SORT1 | sortilin 1 | maintenance of synapse structure (GO:0099558) | BP |
| STAT3 | signal transducer and activator of transcription 3 | postsynapse to nucleus signaling pathway (GO:0099527), modulation of chemical synaptic transmission (GO:0050804) | BP |

| | | | |
|---|---|---|---|
| STX4 | syntaxin 4 | exocytic insertion of neurotransmitter receptor to postsynaptic membrane (GO:0098967), regulation of postsynaptic membrane neurotransmitter receptor levels (GO:0099072) | BP |
| SYT17 | synaptotagmin 17 | regulation of postsynaptic neurotransmitter receptor endocytosis (GO:0099149) | BP |
| SYT2 | synaptotagmin 2 | calcium-dependent activation of synaptic vesicle fusion (GO:0099502) | BP |
| TANC2 | tetratricopeptide repeat, ankyrin repeat and coiled-coil containing 2 | structural constituent of postsynaptic density (GO:0098919) | BP |
| UNC13A | unc-13 homolog A | synaptic vesicle exocytosis (GO:0016079) | BP |
| VHL | von Hippel-Lindau tumor suppressor | regulation of postsynapse organization (GO:0099175), regulation protein catabolic process at postsynapse (GO:0140252) | BP |
| VAPB | VAMP associated protein B and C | presynaptic cytoskeleton (GO:0099569) | CC |
| ACP4 | acid phosphatase 4 | integral component of synaptic membrane (GO:0099699) | CC |
| ACTN1 | actinin alpha 1 | synapse (GO:0045202), postsynaptic actin cytoskeleton (GO:0098871) | CC |
| ADD1 | adducin 1 | presynapse (GO:0098793), postsynapse (GO:0098794), synapse (GO:0045202) | CC |
| ADORA2A | adenosine A2a receptor | integral component of presynaptic membrane (GO:0099056), integral component of postsynaptic membrane (GO:0099055) | CC |
| AKAP1 | A-kinase anchoring protein 1 | postsynaptic specialization, intracellular component (GO:0099091) | CC |
| ANO6 | anoctamin 6 | integral component of synaptic membrane (GO:0099699) | CC |
| AP1S1 | adaptor related protein complex 1 subunit sigma 1 | presynapse (GO:0098793) | CC |
| APC2 | APC regulator of WNT signaling pathway 2 | postsynapse (GO:0098794) | CC |
| APLP2 | amyloid beta precursor like protein 2 | spine apparatus (GO:0097444) | CC |
| APOA4 | apolipoprotein A4 | synapse (GO:0045202) | CC |
| ARHGEF15 | Rho guanine nucleotide exchange factor 15 | postsynapse (GO:0098794) | CC |

| | | | |
|---|---|---|---|
| ATP6V0E2 | ATPase H+ transporting V0 subunit e2 | integral component of synaptic vesicle membrane (GO:0030285) | CC |
| ATP6V1B2 | ATPase H+ transporting V1 subunit B2 | extrinsic component of synaptic vesicle membrane (GO:0098850) | CC |
| ATR | ATR serine/threonine kinase | extrinsic component of synaptic vesicle membrane (GO:0098850), postsynapse (GO:0098794) | CC |
| BCAN | brevican | perisynaptic extracellular matrix (GO:0098966), perisynaptic extracellular matrix (GO:0098966) | CC |
| BCL11A | BCL11 transcription factor A | postsynapse (GO:0098794) | CC |
| BSN | bassoon presynaptic cytomatrix protein | presynaptic active zone cytoplasmic component (GO:0098831) | CC |
| CALB2 | calbindin 2 | synapse (GO:0045202) | CC |
| CAP2 | cyclase associated actin cytoskeleton regulatory protein 2 | postsynapse (GO:0098794) | CC |
| CASP3 | caspase 3 | postsynaptic density (GO:0014069) | CC |
| CASR | calcium sensing receptor | integral component of presynaptic membrane (GO:0099056) | CC |
| CBLB | Cbl proto-oncogene B | postsynapse (GO:0098794) | CC |
| CBLN2 | cerebellin 2 precursor | synaptic cleft (GO:0043083) | CC |
| CHRM1 | cholinergic receptor muscarinic 1 | integral component of postsynaptic membrane (GO:0099055), integral component of postsynaptic density membrane (GO:0099061), integral component of presynaptic membrane (GO:0099056) | CC |
| CNR2 | cannabinoid receptor 2 | integral component of postsynaptic membrane (GO:0099055) | CC |
| CNTNAP2 | contactin associated protein 2 | integral component of presynaptic membrane (GO:0099056) | CC |
| CPNE7 | copine 7 | synapse (GO:0045202) | CC |
| CPSF2 | cleavage and polyadenylation specific factor 2 | postsynapse (GO:0098794) | CC |
| CTBP1 | C-terminal binding protein 1 | cytoplasmic component (GO:0098831), extrinsic component of presynaptic endocytic zone membrane (GO:0098894) | CC |
| CTSD | cathepsin D | presynaptic endosome (GO:0098830) | CC |
| DIXDC1 | DIX domain containing 1 | postsynapse (GO:0098794) | CC |
| DRD1 | dopamine receptor D1 | integral component of presynaptic membrane (GO:0099056), integral component of postsynaptic membrane (GO:0099055), integral component of presynaptic membrane (GO:0099056) | CC |

| | | | |
|---|---|---|---|
| EEF2K | eukaryotic elongation factor 2 kinase | postsynaptic density (GO:0014069) | CC |
| EIF2B2 | eukaryotic translation initiation factor 2B subunit beta | postsynaptic ER | CC |
| EIF2S1 | eukaryotic translation initiation factor 2 subunit alpha | synapse (GO:0045202) | CC |
| EIF4E | eukaryotic translation initiation factor 4E | postsynapse (GO:0098794), postsynaptic cytosol (GO:0099524) | CC |
| ELFN1 | extracellular leucine rich repeat and fibronectin type III domain containing 1 | integral component of postsynaptic density membrane (GO:0099061) | CC |
| ERBIN | erbb2 interacting protein | postsynaptic specialization (GO:0099572), postsynapse (GO:0098794) | CC |
| FZD5 | frizzled class receptor 5 | synapse (GO:0045202) | CC |
| GABBR1 | gamma-aminobutyric acid type B receptor subunit 1 | integral component of presynaptic membrane (GO:0099056), integral component of postsynaptic specialization membrane (GO:0099060), integral component of postsynaptic membrane (GO:0099055), integral component of presynaptic active zone membrane (GO:0099059) | CC |
| GABRB2 | gamma-aminobutyric acid type A receptor subunit beta2 | integral component of presynaptic active zone membrane (GO:0099059), integral component of postsynaptic specialization membrane (GO:0099060) | CC |
| GAP43 | growth associated protein 43 | presynapse (GO:0098793) | CC |
| GNAI1 | G protein subunit alpha i1 | postsynapse (GO:0098794) | CC |
| GNAZ | G protein subunit alpha z | synapse (GO:0045202) | CC |
| GRIN2A | glutamate ionotropic receptor NMDA type subunit 2A | integral component of presynaptic membrane (GO:0099056), integral component of postsynaptic density membrane (GO:0099061) | CC |
| GUCY1B1 | guanylate cyclase 1 soluble subunit beta 1 | presynaptic active zone cytoplasmic component (GO:0098831) | CC |
| HNRNPA1 | heterogeneous nuclear ribonucleoprotein A1 | synapse (GO:0045202) | CC |
| HNRNPK | heterogeneous nuclear ribonucleoprotein K | postsynapse (GO:0098794) | CC |
| HNRNPM | heterogeneous nuclear ribonucleoprotein M | [illegible] (GO:0014069), synapse (GO:0045202) | CC |
| HTR7 | 5-hydroxytryptamine receptor 7 | integral component of presynaptic membrane (GO:0099056), integral component of postsynaptic membrane (GO:0099055) | CC |

| | | | |
|---|---|---|---|
| IGF1 | insulin like growth factor 1 | neuronal dense core vesicle lumen (GO:0099013) | CC |
| KCNAB2 | potassium voltage-gated channel subfamily A regulatory beta subunit 2 | integral component of postsynaptic density membrane (GO:0099061) | CC |
| KCNJ8 | potassium inwardly rectifying channel subfamily J member 8 | integral component of presynaptic active zone membrane (GO:0099059) | CC |
| KCNJ9 | potassium inwardly rectifying channel subfamily J member 9 | integral component of presynaptic membrane (GO:0099056) | CC |
| KCTD8 | potassium channel tetramerization domain containing 8 | postsynapse (GO:0098794), presynaptic active zone membrane (GO:0048787), presynaptic membrane (GO:0042734), | CC |
| KIF21B | kinesin family member 21B | postsynaptic endosome (GO:0098845) | CC |
| KPNA1 | karyopherin subunit alpha 1 | postsynaptic density (GO:0014069) | CC |
| MARCKSL1 | MARCKS like 1 | presynaptic membrane (GO:0099026), synaptic vesicle (GO:0008021), presynaptic cytosol (GO:0099523) | CC |
| MPP1 | MAGUK p55 scaffold protein 1 | postsynapse (GO:0098794) | CC |
| NAPB | NSF attachment protein beta | presynapse (GO:0098793), postsynapse (GO:0098794), presynaptic active zone membrane (GO:0048787) | CC |
| NEO1 | neogenin 1 | integral component of postsynaptic density membrane (GO:0099061) | CC |
| NGDN | neuroguidin | postsynapse (GO:0098794) | CC |
| NGEF | neuronal guanine nucleotide exchange factor | postsynapse (GO:0098794) | CC |
| PDYN | prodynorphin | synaptic vesicle (GO:0008021), neuronal dense core vesicle (GO:0098992) | CC |
| PLAT | plasminogen activator, tissue type | postsynapse (GO:0098794), neuronal dense core vesicle (GO:0098992), extrasynaptic space (GO:0099544), postsynapse (GO:0098794) | CC |
| PPT1 | palmitoyl-protein thioesterase 1 | synaptic vesicle membrane (GO:0030672), presynaptic cytosol (GO:0099523) | CC |
| PRKCA | protein kinase C alpha | presynaptic cytosol (GO:0099523) | CC |
| PRMT8 | protein arginine methyltransferase 8 | postsynapse (GO:0098794) | CC |
| PSEN2 | presenilin 2 | integral component of presynaptic membrane (GO:0099056) | CC |

| | | | |
|---|---|---|---|
| PTPRN2 | protein tyrosine phosphatase receptor type N2 | integral component of synaptic vesicle membrane (GO:0030285), presynapse (GO:0098793) | CC |
| PUM2 | pumilio RNA binding family member 2 | postsynapse (GO:0098794) | CC |
| RAB6B | RAB6B, member RAS oncogene family | presynapse (GO:0098793) | CC |
| RBMX | RNA binding motif protein X-linked | postsynaptic density (GO:0014069) | CC |
| RELN | reelin | postsynaptic density (GO:0014069) | CC |
| RHOG | ras homolog family member G | postsynapse (GO:0098794) | CC |
| RNF216 | ring finger protein 216 | postsynaptic endocytic zone (GO:0098843) | CC |
| ROBO2 | roundabout guidance receptor 2 | integral component of postsynaptic density membrane (GO:0099061) | CC |
| ROR2 | receptor tyrosine kinase like orphan receptor 2 | postsynapse (GO:0098794) | CC |
| RPL17 | ribosomal protein L17 | synapse (GO:0045202), postsynaptic ribosome, presynaptic ribosome | CC |
| RPL22 | ribosomal protein L22 | presynaptic ribosome, synapse (GO:0045202) | CC |
| RPL26 | ribosomal protein L26 | presynaptic ribosome, synapse (GO:0045202) | CC |
| RPL29 | ribosomal protein L29 | synapse (GO:0045202), postsynaptic ribosome, presynaptic ribosome | CC |
| RPL30 | ribosomal protein L30 | postsynaptic density (GO:0014069), synapse (GO:0045202), postsynaptic ribosome | CC |
| RPL31 | ribosomal protein L31 | synapse (GO:0045202), postsynaptic ribosome | CC |
| RPL37A | ribosomal protein L37a | synapse (GO:0045202), presynaptic ribosome | CC |
| RPS19 | ribosomal protein S19 | [illegible] postsynaptic density (GO:0014069), postsynaptic ribosome | CC |
| RPS21 | ribosomal protein S21 | synapse (GO:0045202), postsynaptic ribosome | CC |
| RPS24 | ribosomal protein S24 | synapse (GO:0045202), postsynaptic ribosome, presynaptic ribosome | CC |
| RPS25 | ribosomal protein S25 | [illegible] postsynaptic density (GO:0014069) | CC |
| RPS28 | ribosomal protein S28 | synapse (GO:0045202),. presynaptic ribosome | CC |

| | | | |
|---|---|---|---|
| RPS3A | ribosomal protein S3A | synapse (GO:0045202), postsynaptic ribosome | CC |
| RPS4X | ribosomal protein S4 X-linked | synapse (GO:0045202) | CC |
| S1PR3 | sphingosine-1-phosphate receptor 3 | presynapse (GO:0098793) | CC |
| S1PR4 | sphingosine-1-phosphate receptor 4 | presynapse (GO:0098793) | CC |
| SCN2A | sodium voltage-gated channel alpha subunit 2 | integral component of presynaptic membrane (GO:0099056) | CC |
| SDCBP | syndecan binding protein | presynapse (GO:0098793) | CC |
| SEPTIN6 | septin 6 | presynapse (GO:0098793) | CC |
| SHANK3 | SH3 and multiple ankyrin repeat domains 3 | intracellular component (GO:0099092), postsynaptic density (GO:0014069) | CC |
| SLC3A2 | solute carrier family 3 member 2 | synapse (GO:0045202) | CC |
| SLC6A17 | solute carrier family 6 member 17 | integral component of synaptic vesicle membrane (GO:0030285) | CC |
| SLIT2 | slit guidance ligand 2 | perisynaptic extracellular matrix (GO:0098966) | CC |
| SRPK2 | SRSF protein kinase 2 | presynapse (GO:0098793) | CC |
| STAT3 | signal transducer and activator of transcription 3 | postsynaptic density (GO:0014069) | CC |
| STX16 | syntaxin 16 | integral component of synaptic vesicle membrane (GO:0030285) | CC |
| STX2 | syntaxin 2 | synapse (GO:0045202) | CC |
| STX4 | syntaxin 4 | postsynapse (GO:0098794), integral component of postsynaptic membrane (GO:0099055) | CC |
| SYT17 | synaptotagmin 17 | postsynapse (GO:0098794) | CC |
| SYT2 | synaptotagmin 2 | integral component of synaptic vesicle membrane (GO:0030285) | CC |
| TANC2 | tetratricopeptide repeat, ankyrin repeat and coiled-coil containing 2 | postsynaptic density, intracellular component (GO:0099092), postsynaptic density (GO:0014069) | CC |
| UNC13A | unc-13 homolog A | presynaptic active zone (GO:0048786) | CC |
| USP14 | ubiquitin specific peptidase 14 | presynaptic cytosol (GO:0099523) | CC |
| VAPB | VAMP associated protein B and C | integral component of presynaptic membrane (GO:0099056) | CC |
| VHL | von Hippel-Lindau tumor suppressor | postsynaptic density (GO:0014069) | CC |
| VPS11 | VPS11 core subunit of CORVET and HOPS complexes | presynaptic active zone (GO:0048786) | CC |

| | | | |
|---|---|---|---|
| WDR7 | WD repeat domain 7 | synaptic vesicle (GO:0008021) | CC |

BP: biological process, CC: cellular component

**Supplementary Table 16. Dysregulated mitochondrial mRNAs in serum from anesthetics exposed pediatric patients < 4 years old**

| Gene ID | Gene name | Process | P value | Fold change | Regulation |
|---|---|---|---|---|---|
| IDE | insulin degrading enzyme | Mitochondrial Signaling | 0.006 | 7.280 | Down |
| QDPR | reductase | Amino Acid Metabolism | 0.021 | 6.672 | Down |
| IMMP2L | inner mitochondrial membrane peptidase subunit 2 | Import & Sorting | 0.010 | 4.864 | Down |
| TXNRD1 | thioredoxin reductase 1 | ROS Defense | 0.008 | 4.486 | Down |
| NDUFB1 | NADH:ubiquinone oxidoreductase subunit B1 | Oxidative Phosphorylation | 0.049 | 4.028 | Down |
| TSTD1 | thiosulfate sulfurtransferase like domain containing 1 | Oxidative Phosphorylation | 0.031 | 4.008 | Down |
| COQ8A | coenzyme Q8A | Phosphorylation | 0.018 | 3.329 | Down |
| MMAA | metabolism of cobalamin associated A | Metabolism of Vitamins & Co-Factors | 0.018 | 3.276 | Down |
| FIS1 | fission, mitochondrial 1 | Mitochondrial Dynamics | 0.012 | 3.188 | Down |
| ARNT | aryl hydrocarbon receptor nuclear translocator | Transcription (nuclear) | 0.025 | 3.044 | Down |
| UQCRQ | ubiquinol-cytochrome c reductase complex III subunit VII | Oxidative Phosphorylation | 0.019 | 3.000 | Down |
| SUOX | sulfite oxidase | Amino Acid Metabolism | 0.008 | 2.948 | Down |
| NDUFA10 | NADH:ubiquinone oxidoreductase subunit A10 | Oxidative Phosphorylation | 0.007 | 2.916 | Down |
| BIRC8 | 8 | Apoptosis | 0.036 | 2.857 | Down |
| COA5 | cytochrome c oxidase assembly factor 5 | Oxidative Phosphorylation | 0.011 | 2.799 | Down |
| HADH | hydroxyacyl-CoA dehydrogenase | Fatty Acid Degradation & Beta-oxidation | 0.035 | 2.791 | Down |
| MRM3 | mitochondrial rRNA methyltransferase 3 | Translation | 0.015 | 2.737 | Down |
| SIRT4 | sirtuin 4 | Mitochondrial Signaling | 0.030 | 2.617 | Down |
| SLC25A53 | 53 | Mitochondrial Carrier | 0.006 | 2.564 | Down |
| DNAJC30 | DnaJ heat shock protein family (Hsp40) member C30 | Protein Stability & Degradation | 0.020 | 2.401 | Down |
| GLYATL2 | glycine-N-acyltransferase like 2 | Fatty Acid Metabolism | 0.003 | 2.366 | Down |
| MMADHC | metabolism of cobalamin associated D | Metabolism of Vitamins & Co-Factors | 0.030 | 2.276 | Down |
| NDUFA7 | NADH:ubiquinone oxidoreductase subunit A7 | Oxidative Phosphorylation | 0.012 | 2.123 | Down |
| CYP27B1 | cytochrome P450 family 27 subfamily B member 1 | Metabolism of Vitamins & Co-Factors | 0.028 | 2.070 | Down |
| CKMT1B | creatine kinase, mitochondrial 1B | Amino Acid Metabolism | 0.023 | 2.038 | Down |
| ME1 | malic enzyme 1 | Pyruvate Metabolism | 0.009 | 2.024 | Down |
| MRPL30 | L30 | Translation | 0.006 | 1.990 | Down |
| ECHS1 | enoyl-CoA hydratase, short chain 1 | Fatty Acid Biosynthesis & Elongation | 0.014 | 1.952 | Down |
| QTRT1 | queuine tRNA-ribosyltransferase catalytic subunit 1 | Translation | 0.024 | 1.849 | Down |
| MRPL33 | L33 | Translation | 0.007 | 1.813 | Down |
| OXA1L | OXA1L mitochondrial inner membrane protein | Oxidative Phosphorylation | 0.013 | 1.811 | Down |
| MTFP1 | mitochondrial fission process 1 | Mitochondrial Dynamics | 0.003 | 1.807 | Down |
| PTCD3 | pentatricopeptide repeat domain 3 | Translation | 0.011 | 1.785 | Down |
| SLC2A5 | solute carrier family 2 member 5 | Glycolysis | 0.019 | 1.784 | Down |
| NLN | neurolysin | Protein Stability & Degradation | 0.003 | 1.650 | Down |
| ACP6 | acid phosphatase 6, lysophosphatidic | Metabolism of Lipids & Lipoproteins | 0.022 | 1.598 | Down |

| | | | | | |
|---|---|---|---|---|---|
| BCL2L2 | BCL2 like 2 | Apoptosis | 0.013 | 1.571 | Down |
| DHX29 | DExH-box helicase 29 | Transcription | 0.002 | 1.439 | Down |
| ATPAF1 | ATP synthase mitochondrial F1 complex assembly factor 1 | Oxidative Phosphorylation | 0.037 | 1.428 | Down |
| DBT | dihydrolipoamide branched chain transacylase E2 | Amino Acid Metabolism | 0.024 | 1.372 | Down |
| CASP3 | caspase 3 | Apoptosis | 0.041 | 1.363 | Down |
| LACTB | lactamase beta | Metabolism of Lipids & Lipoproteins | 0.038 | 1.358 | Down |
| CISD1 | CDGSH iron sulfur domain 1 | Phosphorylation | 0.022 | 1.256 | Down |
| TKFC | triokinase and FMN cyclase | Fructose Metabolism | 0.005 | 1.223 | Down |
| SLC25A25 | 25 | Mitochondrial Carrier | 0.013 | 1.210 | Down |
| SLC25A32 | 32 | Metabolism | 0.012 | 1.209 | Up |
| HK3 | hexokinase 3 | Glycolysis | 0.042 | 1.212 | Up |
| TMEM143 | transmembrane protein 143 | Unknown | 0.031 | 1.253 | Up |
| NARF | nuclear prelamin A recognition factor | Fe-S Cluster Biosynthesis | 0.046 | 1.262 | Up |
| ACSS3 | acyl-CoA synthetase short chain family member 3 | Metabolism of Lipids & Lipoproteins | 0.002 | 1.277 | Up |
| CAT | catalase | ROS Defense | 0.038 | 1.277 | Up |
| AKAP1 | A-kinase anchoring protein 1 | Mitochondrial Signaling | 0.001 | 1.287 | Up |
| ATP5PD | ATP synthase peripheral stalk subunit d | Oxidative Phosphorylation | 0.013 | 1.360 | Up |
| SLC25A40 | 40 | Mitochondrial Carrier | 0.033 | 1.384 | Up |
| NUBP2 | nucleotide binding protein 2 | Fe-S Cluster Biosynthesis | 0.041 | 1.466 | Up |
| YRDC | yrdC N6-threonylcarbamoyltransferase domain containing | Replication & Transcription | 0.026 | 1.487 | Up |
| TAZ | WW domain containing transcription regulator 1 | Cardiolipin Biosynthesis | 0.028 | 1.519 | Up |
| ATIC | 5-aminoimidazole-4-carboxamide ribonucleotide formyltransferase/IMP cyclohydrolase | Nucleotide Metabolism | 0.014 | 1.522 | Up |
| MRPS11 | S11 | Translation | 0.038 | 1.525 | Up |
| PKM | pyruvate kinase M1/2 | Glycolysis | 0.034 | 1.579 | Up |
| CASP9 | caspase 9 | Apoptosis | 0.020 | 1.641 | Up |
| SLC25A43 | 43 | Mitochondrial Carrier | 0.023 | 1.680 | Up |
| ACSS1 | acyl-CoA synthetase short chain family member 1 | Pyruvate Metabolism | 0.031 | 1.682 | Up |
| PDPR | pyruvate dehydrogenase phosphatase regulatory subunit | Pyruvate Metabolism | 0.043 | 1.694 | Up |
| PRELID3B | PRELI domain containing 3B | Metabolism of Lipids & Lipoproteins | 0.048 | 1.699 | Up |
| ABHD16A | abhydrolase domain containing 16A | Amino Acid Metabolism | 0.013 | 1.721 | Up |
| NDUFB10 | NADH:ubiquinone oxidoreductase subunit B10 | Oxidative Phosphorylation | 0.028 | 1.725 | Up |
| PYCR1 | 1 | Amino Acid Metabolism | 0.033 | 1.774 | Up |
| LDHA | lactate dehydrogenase A | Glycolysis | 0.017 | 1.775 | Up |
| MFF | mitochondrial fission factor | Mitochondrial Dynamics | 0.007 | 2.197 | Up |
| COQ7 | coenzyme Q7, hydroxylase | Ubiquinone Biosynthesis | 0.037 | 2.341 | Up |
| MRPL57 | L57 | Translation | 0.037 | 2.741 | Up |
| PPARA | peroxisome proliferator activated receptor alpha | Transcription (nuclear) | 0.014 | 3.085 | Up |
| MT-CYB | mitochondrially encoded cytochrome b | Oxidative Phosphorylation (mt) | 0.024 | 3.182 | Up |
| SLC25A14 | 14 | Mitochondrial Carrier | 0.023 | 3.276 | Up |
| MRPS16 | S16 | Translation | 0.027 | 3.397 | Up |

| | | | | | |
|---|---|---|---|---|---|
| ALDH6A1 | aldehyde dehydrogenase 6 family member A1 | Amino Acid Metabolism | 0.040 | 3.640 | Up |
| MT-ND3 | mitochondrially encoded NADH dehydrogenase 3 | Oxidative Phosphorylation (mt) | 0.024 | 4.353 | Up |
| TSFM | Ts translation elongation factor, mitochondrial | Translation | 0.013 | 4.770 | Up |
| PNPT1 | polyribonucleotide nucleotidyltransferase 1 | Translation | 0.015 | 5.051 | Up |
| NDUFA11 | NADH:ubiquinone oxidoreductase subunit A11 | Oxidative Phosphorylation | 0.036 | 5.641 | Up |
| TOP3A | DNA topoisomerase III alpha | Transcription | 0.022 | 5.787 | Up |
| PTCD1 | pentatricopeptide repeat domain 1 | Translation | 0.007 | 5.969 | Up |
| PCCB | propionyl-CoA carboxylase subunit beta | Fatty Acid Metabolism | 0.004 | 6.326 | Up |
| PGAM5 | PGAM family member 5, mitochondrial serine/threonine protein phosphatase | Mitophagy | 0.004 | 6.685 | Up |
| RPS19 | ribosomal protein S19 | Translation | 0.012 | 6.868 | Up |
| OPA3 | OPA3 outer mitochondrial membrane lipid metabolism regulator | Metabolism of Lipids & Lipoproteins | 0.005 | 7.428 | Up |
| UQCR11 | ubiquinol-cytochrome c reductase, complex III subunit XI | Oxidative Phosphorylation | 0.000 | 7.901 | Up |
| MT-CO3 | mitochondrially encoded cytochrome c oxidase III | Oxidative Phosphorylation (mt) | 0.017 | 16.032 | Up |

**Supplementary Table 17. Dysregulated Inflammatory mRNAs in serum from anesthetic exposed pediatric patients < 4 years old**

| Gene ID | P value | Fold change | Regulation |
|---|---|---|---|
| FTL | 0.003 | 19.556 | Up |
| PPBP | 0.019 | 16.490 | Up |
| NEO1 | 0.034 | 1.883 | Up |
| GDAP1L1 | 0.000 | 1.848 | Up |
| CTSL | 0.016 | 1.660 | Up |
| PLA2G6 | 0.026 | 1.655 | Up |
| DCUN1D5 | 0.038 | 1.633 | Up |
| SPINK13 | 0.041 | 1.442 | Up |
| UPF1 | 0.038 | 1.442 | Up |
| TPH1 | 0.020 | 1.427 | Up |
| PRMT1 | 0.019 | 1.420 | Up |
| USP14 | 0.044 | 1.411 | Up |
| CYP1A1 | 0.012 | 1.392 | Up |
| CTBP1 | 0.035 | 1.387 | Up |
| NDST1 | 0.017 | 1.367 | Up |
| SORT1 | 0.041 | 1.340 | Up |
| KLF2 (other aliase for CKLF) | 0.030 | 1.338 | Up |
| KNG1 | 0.037 | 1.336 | Up |
| CNDP2 | 0.032 | 1.320 | Up |
| IL23A | 0.026 | 1.318 | Up |
| HSD17B1 | 0.017 | 1.310 | Up |
| ICOSLG | 0.033 | 1.297 | Up |
| AKAP1 | 0.001 | 1.287 | Up |
| GUCY1B1 | 0.020 | 1.283 | Up |
| CAT | 0.038 | 1.277 | Up |
| S1PR3 | 0.045 | 1.249 | Up |
| FGF19 | 0.042 | 1.243 | Up |
| HK3 (other aliase for KLK3) | 0.042 | 1.212 | Up |
| SLC28A3 | 0.001 | 16.609 | Down |
| MCAM | 0.023 | 7.936 | Down |
| CCL8 | 0.006 | 6.435 | Down |
| STAT6 | 0.006 | 6.043 | Down |
| EGR1 | 0.001 | 6.040 | Down |
| IRS2 | 0.006 | 5.936 | Down |
| RPL22 | 0.010 | 5.253 | Down |
| MNDA | 0.002 | 4.906 | Down |
| LGALS8 | 0.006 | 4.782 | Down |
| CLEC1A | 0.004 | 4.578 | Down |
| VEGFC | 0.006 | 4.529 | Down |
| TXNRD1 | 0.008 | 4.486 | Down |
| BMP7 | 0.004 | 4.045 | Down |
| PROKR1 | 0.018 | 3.974 | Down |
| IL13 | 0.031 | 3.924 | Down |
| BRD3 | 0.024 | 3.763 | Down |
| PQBP1 | 0.004 | 3.705 | Down |
| FRZB | 0.005 | 3.474 | Down |
| PRKCQ | 0.020 | 3.104 | Down |
| TSHZ3 | 0.024 | 3.072 | Down |
| MPO | 0.039 | 3.060 | Down |
| HLA-B | 0.009 | 3.037 | Down |
| HSD17B13 | 0.006 | 2.946 | Down |
| FSTL1 | 0.019 | 2.927 | Down |
| LTA4H | 0.026 | 2.814 | Down |
| SLC3A2 | 0.030 | 2.772 | Down |
| JMJD1C | 0.042 | 2.732 | Down |
| HDAC4 | 0.040 | 2.700 | Down |
| DEFA1 | 0.035 | 2.660 | Down |
| TRAF3IP2 | 0.016 | 2.621 | Down |
| SIRT4 | 0.030 | 2.617 | Down |
| ROR1 (Aliase for RORA) | 0.009 | 2.560 | Down |
| FBXO11 | 0.010 | 2.549 | Down |

| | | | |
|---|---|---|---|
| MAP2K2 | 0.015 | 2.498 | Down |
| SLIT2 | 0.044 | 2.454 | Down |
| GPR183 | 0.041 | 2.405 | Down |
| LMNB1 | 0.014 | 2.395 | Down |
| TMED2 | 0.027 | 2.363 | Down |
| TFF1 | 0.009 | 2.263 | Down |
| TFAP2C | 0.039 | 2.249 | Down |
| COCH | 0.022 | 2.222 | Down |
| GC | 0.044 | 2.215 | Down |
| CFI | 0.040 | 2.169 | Down |
| GPR15 | 0.022 | 2.137 | Down |
| FTH1 | 0.036 | 2.126 | Down |
| PGAP1 | 0.029 | 2.111 | Down |
| CYP27B1 | 0.028 | 2.070 | Down |
| SLC39A8 | 0.036 | 2.062 | Down |
| IL23R | 0.002 | 2.043 | Down |
| APOA4 | 0.048 | 1.962 | Down |
| IGF1 | 0.010 | 1.904 | Down |
| PPP1R7 (Aliase for PTK2) | 0.007 | 1.902 | Down |
| DUSP14 | 0.037 | 1.897 | Down |
| TIGIT | 0.015 | 1.764 | Down |
| TSLP | 0.006 | 1.736 | Down |
| TYROBP | 0.031 | 1.680 | Down |
| ACKR1 | 0.022 | 1.639 | Down |
| SCARB1 | 0.019 | 1.604 | Down |
| SHANK3 | 0.042 | 1.603 | Down |
| GAP43 | 0.044 | 1.600 | Down |
| ROR2 (Aliase For RORA) | 0.003 | 1.548 | Down |
| CASR | 0.042 | 1.537 | Down |
| ASH2L | 0.015 | 1.535 | Down |
| GZMM | 0.019 | 1.501 | Down |
| PCSK6 | 0.016 | 1.478 | Down |
| BHLHE41 | 0.041 | 1.473 | Down |
| CRISP3 | 0.046 | 1.470 | Down |
| DDR1 | 0.022 | 1.460 | Down |
| XDH | 0.009 | 1.435 | Down |
| ANG | 0.036 | 1.430 | Down |
| PLAT | 0.023 | 1.427 | Down |
| CCL18 | 0.029 | 1.422 | Down |
| HBA1 | 0.031 | 1.359 | Down |
| CES2 | 0.020 | 1.357 | Down |
| PUM2 | 0.044 | 1.335 | Down |
| ATG7 | 0.017 | 1.321 | Down |
| FCAR | 0.022 | 1.306 | Down |
| PGLYRP1 | 0.006 | 1.303 | Down |
| SERPINF2 | 0.038 | 1.283 | Down |
| ADORA2A | 0.010 | 1.254 | Down |
| DUSP19 | 0.040 | 1.245 | Down |
| CNR2 | 0.024 | 1.206 | Down |

28 up-regulated and 82 down-regulated

**Supplementary Table 18. Brain cell type-specific mRNAs in the serum from pediatric patients < 4 years old with >3-hour anesthesia/surgery**

| Genes | Pvalues | Fold Change | Cell Type |
|---|---|---|---|
| FRMPD2 | 0.028 | 2.296 | astrocytes |
| BHMT2 | 0.007 | 6.419 | astrocytes |
| SLC4A4 | 0.009 | 2.035 | astrocytes |
| BCAN | 0.031 | 2.200 | astrocytes |
| RGR | 0.024 | 2.433 | astrocytes |
| PPARA | 0.014 | 3.085 | astrocytes |
| INTS3 | 0.039 | 2.600 | astrocytes |
| PRKCA | 0.015 | 3.098 | astrocytes |
| BMP7 | 0.004 | 4.045 | astrocytes |
| SUOX | 0.008 | 2.948 | astrocytes |
| PTCHD1 | 0.022 | 2.137 | astrocytes |
| TRAF3IP2 | 0.016 | 2.621 | astrocytes |
| SLC25A53 | 0.006 | 2.564 | astrocytes |
| TFAP2C | 0.039 | 2.249 | astrocytes |
| MACF1 | 0.036 | 3.442 | astrocytes |
| PCDHGA9 | 0.042 | 2.016 | astrocytes |
| IL17D | 0.031 | 5.653 | astrocytes |
| ME1 | 0.009 | 2.024 | astrocytes |
| MUT | 0.012 | 5.212 | astrocytes |
| TPP1 | 0.006 | 3.200 | astrocytes |
| C1ORF35 | 0.009 | 2.801 | endothelial |
| APOL1 | 0.007 | 2.025 | endothelial |
| H2AFZ | 0.003 | 2.103 | endothelial |
| OPA3 | 0.005 | 7.428 | endothelial |
| WNT11 | 0.001 | 2.043 | endothelial |
| RPS21 | 0.002 | 21.240 | endothelial |
| SLC12A7 | 0.018 | 4.898 | endothelial |
| FOXC2 | 0.029 | 7.654 | endothelial |
| CLEC1A | 0.004 | 4.578 | endothelial |
| FBXO25 | 0.023 | 2.567 | endothelial |
| LNX2 | 0.013 | 3.641 | endothelial |
| ANTXR2 | 0.009 | 2.018 | endothelial |
| CFI | 0.040 | 2.169 | endothelial |
| RBMS2 | 0.000 | 5.129 | endothelial |
| FSTL1 | 0.019 | 2.927 | endothelial |
| TRAM2 | 0.024 | 8.822 | endothelial |
| GJC1 | 0.021 | 3.332 | endothelial |
| FRZB | 0.005 | 3.474 | endothelial |
| KRTCAP3 | 0.039 | 2.902 | endothelial |
| SLC39A8 | 0.036 | 2.062 | endothelial |
| CBWD2 | 0.023 | 3.167 | microglia |
| ALOX15B | 0.012 | 7.997 | microglia |
| VHL | 0.011 | 7.374 | microglia |
| CD226 | 0.002 | 2.028 | microglia |
| LIMS1 | 0.021 | 4.321 | microglia |
| ZNF3 | 0.013 | 5.208 | microglia |
| MNDA | 0.002 | 4.906 | microglia |
| CPED1 | 0.011 | 4.065 | microglia |
| GPR183 | 0.041 | 2.405 | microglia |
| ZBTB1 | 0.021 | 2.763 | microglia |
| CEP170 | 0.032 | 2.378 | microglia |
| EIF4E | 0.011 | 2.412 | microglia |
| ATF7IP2 | 0.045 | 2.282 | microglia |
| SOCS6 | 0.049 | 6.171 | microglia |
| SLC25A14 | 0.023 | 3.276 | neurons |
| CALB2 | 0.027 | 2.509 | neurons |
| ANKRD36B | 0.014 | 2.205 | neurons |
| ROBO2 | 0.023 | 2.719 | neurons |
| SAMD12 | 0.049 | 2.419 | neurons |
| XK | 0.038 | 2.058 | neurons |

| | | | |
|---|---|---|---|
| HIST3H2A | 0.031 | 3.674 | neurons |
| GRIN2A | 0.001 | 2.620 | neurons |
| ZNF567 | 0.025 | 2.086 | neurons |
| PTPRN2 | 0.019 | 2.928 | neurons |
| SRRM4 | 0.004 | 7.132 | neurons |
| PRLHR | 0.021 | 5.008 | neurons |
| LRRC39 | 0.001 | 5.291 | neurons |
| FAM153C | 0.017 | 2.047 | neurons |
| THSD7A | 0.005 | 4.860 | neurons |
| PTGFR | 0.038 | 2.511 | neurons |
| COCH | 0.022 | 2.222 | neurons |
| PJA1 | 0.046 | 4.699 | neurons |
| TBC1D30 | 0.042 | 2.047 | neurons |
| AC005477.1 | 0.013 | 3.359 | neurons |
| SLIT2 | 0.044 | 2.454 | neurons |
| CKMT1B | 0.023 | 2.038 | neurons |
| RYR2 | 0.040 | 4.670 | neurons |
| NAPB | 0.039 | 3.752 | neurons |
| PTMA | 0.003 | 11.416 | oligodendrocytes |
| ANKRD28 | 0.048 | 3.704 | oligodendrocytes |
| RNASEH2A | 0.023 | 3.644 | oligodendrocytes |
| PTMA | 0.003 | 11.416 | oligodendrocytes |
| ZNF664 | 0.005 | 3.369 | oligodendrocytes |
| TMEM206 | 0.022 | 2.014 | oligodendrocytes |
| PNPT1 | 0.015 | 5.051 | oligodendrocytes |
| HIST1H2AC | 0.010 | 2.801 | oligodendrocytes |
| POGK | 0.008 | 3.766 | oligodendrocytes |
| STMN4 | 0.018 | 2.459 | oligodendrocytes |
| MARCKSL1 | 0.022 | 5.252 | oligodendrocytes |
| SRPK2 | 0.017 | 2.286 | oligodendrocytes |
| PRKCQ | 0.020 | 3.104 | oligodendrocytes |
| GDE1 | 0.021 | 6.135 | oligodendrocytes |
| FIS1 | 0.012 | 3.188 | oligodendrocytes |
| ANAPC5 | 0.049 | 2.941 | oligodendrocytes |
| KIF13A | 0.022 | 2.350 | oligodendrocytes |
| CENPI | 0.029 | 2.034 | oligodendrocytes |
| CAPN13 | 0.006 | 2.666 | oligodendrocytes |
| SERINC5 | 0.026 | 2.727 | oligodendrocytes |
| SHB | 0.029 | 3.887 | oligodendrocytes |
| GNAI1 | 0.003 | 3.846 | oligodendrocytes |
| GOLGA7 | 0.003 | 4.601 | oligodendrocytes |
| RAPGEF5 | 0.012 | 3.046 | oligodendrocytes |
| RAB28 | 0.020 | 2.944 | oligodendrocytes |
| SLC22A15 | 0.037 | 2.004 | oligodendrocytes |
| AFMID | 0.039 | 2.178 | oligodendrocytes |
| TMEFF2 | 0.018 | 2.561 | oligodendrocytes |
| INSL3 | 0.009 | 2.220 | oligodendrocytes |
| CLIP2 | 0.028 | 2.379 | oligodendrocytes |
| VRK2 | 0.030 | 2.822 | oligodendrocytes |
| ZDHHC9 | 0.011 | 3.214 | oligodendrocytes |
| SET | 0.021 | 2.902 | oligodendrocytes |
| RDX | 0.026 | 5.461 | oligodendrocytes |
| ENOSF1 | 0.022 | 4.912 | oligodendrocytes |
| OTUD6B | 0.000 | 2.590 | oligodendrocytes |
| QDPR | 0.021 | 6.672 | oligodendrocytes |
| TRMT13 | 0.021 | 11.384 | oligodendrocytes |

**Supplementary Table 19. Overlapping mRNAs dysregulated in both the anesthetics ex pediatric patients < 4 years old**

| | Organoids | | | Serum | | |
|---|---|---|---|---|---|---|
| **Gene ID** | **P value** | **Fold change** | **Regulation** | **P-value** | **Fold Chang** | **Regulation** |
| ARHGAP8 | 0.038 | -1.364 | Down | 0.036 | -2.191 | Down |
| BAG1 | 0.025 | -1.238 | Down | 0.045 | -4.343 | Down |
| CKMT1B | 0.006 | -1.230 | Down | 0.023 | -2.038 | Down |
| FUT11 | 0.030 | -1.469 | Down | 0.033 | -1.725 | Down |
| IL17RC | 0.021 | -1.323 | Down | 0.015 | -2.228 | Down |
| LRRC36 | 0.039 | -1.364 | Down | 0.040 | -1.740 | Down |
| MPO | 0.042 | -1.471 | Down | 0.039 | -3.060 | Down |
| PTPRN2 | 0.029 | -1.474 | Down | 0.019 | -2.928 | Down |
| SERINC1 | 0.046 | -1.284 | Down | 0.030 | -2.254 | Down |
| ACTN1 | 0.040 | 1.413 | Up | 0.036 | 1.349 | Up |
| BAG3 | 0.012 | 1.502 | Up | 0.012 | 1.678 | Up |
| DCDC2C | 0.028 | 1.352 | Up | 0.001 | 1.404 | Up |
| FAM120A | 0.035 | 1.409 | Up | 0.037 | 1.831 | Up |
| GAPVD1 | 0.009 | 1.238 | Up | 0.004 | 1.951 | Up |
| KAT2A | 0.025 | 1.295 | Up | 0.013 | 1.474 | Up |
| NUP188 | 0.030 | 1.254 | Up | 0.034 | 2.080 | Up |
| RHBDD3 | 0.012 | 1.232 | Up | 0.043 | 1.667 | Up |
| RTEL1 | 0.001 | 1.231 | Up | 0.026 | 1.761 | Up |
| RWDD4 | 0.045 | 1.210 | Up | 0.017 | 1.235 | Up |
| SCNN1D | 0.026 | 1.403 | Up | 0.012 | 4.103 | Up |
| SHC2 | 0.029 | 1.226 | Up | 0.015 | 1.466 | Up |

9 down-regulated and 12 up-regulated

**posed organoid and serum from**

| [illegible] down between serum and orgnaoids |
|---|
| Same |
| Same |
| Same |
| Same |
| Same |
| Same |
| Same |
| Same |
| Same |
| Same |
| Same |
| Same |
| Same |
| Same |
| Same |
| Same |
| Same |
| Same |
| Same |
| Same |
| Same |

**Supplementary Table 20. Overlapping synaptic mRNAs dysregulated in both the anesthetics exposed organoid and serum from pediatric patients < 4 years old**

| | **Organoids** | | | **Serum** | | | **Comparison of trend of up or down between serum and orgnaoids** |
|---|---|---|---|---|---|---|---|
| **Gene ID** | P value | Fold change | Regulation | P value | Fold change | Regulation | |
| PTPRN2 | 0.029 | 1.474 | Down | 0.019 | 2.928 | Down | Same |
| ACTN1 | 0.040 | 1.413 | Up | 0.036 | 1.349 | Up | Same |

\

**Supplementary Table 21. Overlapping mitochondrial mRNAs dysregulated in both the anesthetics exposed organoid and serum from pediatric patients < 4 years old**

| | Organoids | | | Serum | | | Comparison of trend of up or down between serum and orgnaoids |
|---|---|---|---|---|---|---|---|
| **Gene ID** | **P value** | **Fold change** | **Regulation** | **P value** | **Fold change** | **Regulation** | |
| CKMT1B | 0.006 | -1.230 | Down | 0.023 | -2.038 | Down | Same |

**Supplementary Table 22. Overlapping lncRNAs dysregulated in both the anesthetics exposed organoid and serum from pediatric patients < 4 years old**

| Gene ID | Organoids | | | Serum | | | Comparison of trend of up or down between serum and orgnaoids |
|---|---|---|---|---|---|---|---|
| | P value | Fold change | Regulation | P value | Fold change | Regulation | |
| CCDC144NL-AS | 0.023 | 1.913 | Down | 0.005 | 1.579 | Down | Same |
| DHRS4-AS1 | 0.005 | 1.450 | Down | 0.015 | 3.844 | Down | Same |
| G061722 | 0.002 | 1.202 | Down | 0.029 | 1.945 | Down | Same |
| HRAT5 | 0.017 | 1.737 | Down | 0.015 | 1.544 | Down | Same |
| LINC00408 | 0.017 | 1.662 | Down | 0.003 | 2.961 | Down | Same |
| LINC00652 | 0.021 | 1.441 | Down | 0.010 | 1.493 | Down | Same |
| LINC01023 | 0.029 | 1.507 | Down | 0.044 | 2.738 | Down | Same |
| XLOC_004632 | 0.023 | 1.294 | Down | 0.036 | 3.389 | Down | Same |
| G038394 | 0.048 | 1.292 | Up | 0.023 | 1.977 | Up | Same |
| LACTB2-AS1 | 0.010 | 1.687 | Up | 0.008 | 2.103 | Up | Same |
| LINC00685 | 0.031 | 1.308 | Up | 0.008 | 6.037 | Up | Same |
| LIPE-AS1 | 0.025 | 1.364 | Up | 0.017 | 1.744 | Up | Same |
